# Genetic variants for head size share genes and pathways with cancer

**DOI:** 10.1101/2020.07.15.191114

**Authors:** Maria J. Knol, Raymond A. Poot, Tavia E. Evans, Claudia L. Satizabal, Aniket Mishra, Sandra van der Auwera, Marie-Gabrielle Duperron, Xueqiu Jian, Isabel C. Hostettler, Dianne H.K. van Dam-Nolen, Sander Lamballais, Mikolaj A. Pawlak, Cora E. Lewis, Amaia Carrion-Castillo, Theo G.M. van Erp, Céline S. Reinbold, Jean Shin, Markus Scholz, Asta K. Håberg, Anders Kämpe, Gloria H.Y. Li, Reut Avinun, Joshua R. Atkins, Fang-Chi Hsu, Alyssa R. Amod, Max Lam, Ami Tsuchida, Mariël W.A. Teunissen, Alexa S. Beiser, Frauke Beyer, Joshua C. Bis, Daniel Bos, R. Nick Bryan, Robin Bülow, Svenja Caspers, Gwenaëlle Catheline, Charlotte A.M. Cecil, Shareefa Dalvie, Jean-François Dartigues, Charles DeCarli, Maria Enlund-Cerullo, Judith M. Ford, Barbara Franke, Barry I. Freedman, Nele Friedrich, Melissa J. Green, Simon Haworth, Catherine Helmer, Per Hoffmann, Georg Homuth, M. Kamran Ikram, Clifford R. Jack, Neda Jahanshad, Christiane Jockwitz, Yoichiro Kamatani, Annchen R. Knodt, Shuo Li, Keane Lim, W. T. Longstreth, Fabio Macciardi, The Cohorts for Heart and Aging Research in Genomic Epidemiology (CHARGE) Consortium, The Enhancing NeuroImaging Genetics through Meta-Analysis (ENIGMA) Consortium, Outi Mäkitie, Bernard Mazoyer, Sarah E. Medland, Susumu Miyamoto, Susanne Moebus, Thomas H. Mosley, Ryan Muetzel, Thomas W. Mühleisen, Manabu Nagata, Soichiro Nakahara, Nicholette D. Palmer, Zdenka Pausova, Adrian Preda, Yann Quidé, William R. Reay, Gennady V. Roshchupkin, Reinhold Schmidt, Pamela J. Schreiner, Kazuya Setoh, Chin Yang Shapland, Stephen Sidney, Beate St Pourcain, Jason L. Stein, Yasuharu Tabara, Alexander Teumer, Anne Uhlmann, Aad van der Lugt, Meike W. Vernooij, David J. Werring, B. Gwen Windham, A. Veronica Witte, Katharina Wittfeld, Qiong Yang, Kazumichi Yoshida, Han G. Brunner, Quentin Le Grand, Kang Sim, Dan J. Stein, Donald W. Bowden, Murray J. Cairns, Ahmad R. Hariri, Ching-Lung Cheung, Sture Andersson, Arno Villringer, Tomas Paus, Sven Cichon, Vince D. Calhoun, Fabrice Crivello, Lenore J. Launer, Tonya White, Peter J. Koudstaal, Henry Houlden, Myriam Fornage, Fumihiko Matsuda, Hans J. Grabe, M. Arfan Ikram, Stéphanie Debette, Paul M. Thompson, Sudha Seshadri, Hieab H.H. Adams

**Affiliations:** Department of Epidemiology, Erasmus MC University Medical Center, Rotterdam, the Netherlands; Department of Cell Biology, Erasmus MC University Medical Center, Rotterdam, the Netherlands; Department of Clinical Genetics, Erasmus MC University Medical Center, Rotterdam, the Netherlands; Department of Radiology and Nuclear Medicine, Erasmus MC University Medical Center, Rotterdam, the Netherlands; Glenn Biggs Institute for Alzheimer’s & Neurodegenerative Diseases, UT Health San Antonio, San Antonio, TX, USA; The Framingham Heart Study, Framingham, MA, USA; Department of Neurology, Boston University School of Medicine, Boston, MA, USA; University of Bordeaux, Inserm, Bordeaux Population Health Research Center, team VINTAGE, UMR 1219, Bordeaux, France; Department of Psychiatry and Psychotherapy, University Medicine Greifswald, Greifswald, Germany; German Centre of Neurodegenerative Diseases (DZNE), Site Rostock/Greifswald, Greifswald, Germany; Brown Foundation Institute of Molecular Medicine, McGovern Medical School, University of Texas Health Science Center at Houston, Houston, TX, USA; Stroke Research Centre, University College London, Institute of Neurology, London, UK; Department of Neurosurgery, Klinikum rechts der Isar, University of Munich, Munich, Germany; Department of Neurology and Cerebrovascular Disorders, Poznań University of Medical Sciences, Poznań, Poland; Department of Epidemiology, School of Public Health, University of Alabama at Birmingham Shool of Medicine, Birmingham, AL, USA; Language and Genetics Department, Max Planck Institute for Psycholinguistics, Nijmegen, the Netherlands; Clinical Translational Neuroscience Laboratory, Department of Psychiatry and Human Behavior, University of California Irvine, Irvine, CA, United States; Center for the Neurobiology of Learning and Memory, University of California Irvine, Irvine, CA, USA; Department of Biomedicine, University of Basel, Basel, Switzerland; Institute of Medical Genetics and Pathology, University Hospital Basel, Basel, Switzerland; Department of Psychology, University of Oslo, Oslo, Norway; The Hospital for Sick Children, University of Toronto, Toronto, Canada; Department of Nutritional Sciences, University of Toronto, Toronto, Canada; Institute for Medical Informatics, Statistics and Epidemiology, University of Leipzig, Leipzig, Germany; LIFE Research Center for Civilization Disease, Leipzig, Germany; Department of Neuromedicine and Movement Science, Norwegian University of Science and Technology (NTNU), Trondheim, Norway; Department of Radiology and Nuclear Medicine, St. Olavs University Hospital, Trondheim, Norway; Department of Molecular Medicine and Surgery, Karolinska Institutet, Stockholm, Sweden; Department of Clinical Genetics, Karolinska University Hospital, Stockholm, Sweden; Department of Pharmacology and Pharmacy, Li Ka Shing Faculty of Medicine, The University of Hong Kong, Hong Kong; Laboratory of NeuroGenetics, Department of Psychology & Neuroscience, Duke University, Durham, NC, USA; School of Biomedical Sciences and Pharmacy, The University of Newcastle, Callaghan, NSW, Australia; Centre for Brain and Mental Health Research, Hunter Medical Research Institute, Newcastle, NSW, Australia; Department of Biostatistics and Data Science, Wake Forest School of Medicine, Winston-Salem, NC, USA; Department of Psychiatry and Mental Health, University of Cape Town, Cape Town, South Africa; Research Division, Institute of Mental Health, Singapore; Groupe d’Imagerie Neurofonctionnelle, Institut des Maladies Neurodégénératives, UMR5293, Université de Bordeaux, Bordeaux, France; Groupe d’Imagerie Neurofonctionnelle, Institut des Maladies Neurodégénératives, UMR5293, CNRS, Bordeaux, France; Groupe d’Imagerie Neurofonctionnelle, Institut des Maladies Neurodégénératives, UMR5293, CEA, Bordeaux, France; Department of Neurology, Maastricht University Medical Center+, Maastricht, The Netherlands; Department of Human Genetics, Radboud University Medical Center, Nijmegen, The Netherlands; Department of Biostatistics, Boston University School of Public Health, Boston, MA, USA; Department of Neurology, Max Planck Institute for Cognitive and Brain Sciences, Leipzig, Germany; Collaborative Research Center 1052 Obesity Mechanisms, Faculty of Medicine, University of Leipzig, Leipzig, Germany; Cardiovascular Health Research Unit, Department of Medicine, University of Washington, Seattle, WA, USA; The University of Texas at Austin Dell Medical School, Austin, TX, USA; Institute of Diagnostic Radiology and Neuroradiology, University Medicine Greifswald, Greifswald, Germany; Institute of Neuroscience and Medicine (INM-1), Research Centre Jülich, Jülich, Germany; JARA-BRAIN, Jülich-Aachen Research Alliance, Jülich, Germany; Institute for Anatomy I, Medical Faculty, Heinrich Heine University Du□sseldorf, Du□sseldorf, Germany; University of Bordeaux, CNRS, INCIA, UMR 5287, team NeuroImagerie et Cognition Humaine, Bordeaux, France; EPHE-PSL University, Bordeaux, France; Department of Child and Adolescent Psychiatry, Erasmus MC University Medical Center, Rotterdam, the Netherlands; University of Bordeaux, Inserm, Bordeaux Population Health Research Center, team SEPIA, UMR 1219, Bordeaux, France; Department of Neurology and Center for Neuroscience, University of California at Davis, Sacramento, CA, USA; Children’s Hospital, University of Helsinki and Helsinki University Hospital, Helsinki, Finland; Folkhälsan Research Center, Helsinki, Finland; San Francisco Veterans Administration Medical Center, San Francisco, CA, USA; University of California San Francisco, San Francisco, CA, USA; Department of Psychiatry, Radboud University Medical Center, Nijmegen, The Netherlands; Donders Institute for Brain, Cognition and Behaviour, Radboud University, Nijmegen, The Netherlands; Department of Internal Medicine, Section on Nephrology, Wake Forest School of Medicine, Winston-Salem, NC, USA; Institute of Clinical Chemistry and Laboratory Medicine, University Medicine Greifswald, Greifswald, Germany; School of Psychiatry, the University of New South Wales, Sydney, NSW, Australia; Neuroscience Research Australia, Sydney, NSW, Australia; MRC Integrative Epidemiology Unit, University of Bristol, Bristol, UK; Population Health Sciences, University of Bristol, Bristol, UK; Bristol Dental School, University of Bristol, Bristol, UK; University of Bordeaux, Inserm, Bordeaux Population Health Research Center, team LEHA, UMR 1219, Bordeaux, France; Institute of Human Genetics, University of Bonn Medical School, Bonn, Germany; Interfaculty Institute for Genetics and Functional Genomics, University Medicine Greifswald, Greifswald, Germany; Department of Neurology, Erasmus MC University Medical Center, Rotterdam, the Netherlands; Department of Radiology, Mayo Clinic, Rochester, MN, USA; Imaging Genetics Center, Mark & Mary Stevens Neuroimaging & Informatics Institute, Keck USC School of Medicine, CA, USA; Department of Psychiatry, Psychotherapy and Psychosomatics, RWTH Aachen University, Medical Faculty, Aachen, Germany; Center for Genomic Medicine, Kyoto University Graduate School of Medicine, Kyoto, Japan; Department of Neurology, University of Washington, Seattle, WA, USA; Department of Epidemiology, University of Washington, Seattle, WA, USA; Laboratory of Molecular Psychiatry, Department of Psychiatry and Human Behavior, School of Medicine, University of California, Irvine, CA, USA; Groupe d’Imagerie Neurofonctionnelle, Institut des Maladies Neurodégénératives, Centre National de la Recherche Scientifique, Commissariat à l’Energie Atomique, et Université de Bordeaux, Bordeaux, France; Centre Hospitalo-Universitaire de Bordeaux, Bordeaux, France; Psychiatric Genetics, QIMR Berghofer Medical Research Institute, Brisbane, QLD, Australia; School of Psychology, University of Queensland, Brisbane, QLD, Australia; Faculty of Medicine, University of Queensland, Brisbane, QLD, Australia; Department of Neurosurgery, Kyoto University Graduate School of Medicine, Kyoto, Japan; Institute for Urban Public Health, University of Duisburg-Essen, Essen, Germany; Department of Medicine, Division of Geriatrics, University of Mississippi Medical Center, Jackson, MS, USA; Memory Impairment and Neurodegenerative Dementia (MIND) Center, Jackson, MS, USA; C. and O. Vogt Institute for Brain Research, Medical Faculty, Heinrich Heine University Düsseldorf, Düsseldorf, Germany; Unit 2, Candidate Discovery Science Labs, Drug Discovery Research, Astellas Pharma Inc, 21, Miyukigaoka, Tsukuba, Ibaraki 305-8585, Japan; Department of Biochemistry, Wake Forest School of Medicine, Winston-Salem, NC, USA; Department of Psychiatry, University of California Irvine, Irvine, CA, USA; Clinical Division of Neurogeriatrics, Department of Neurology, Medical University of Graz, Graz, Austria; University of Minnesota School of Public Health, Minneapolis, MN, USA; Kaiser Permanente Division of Research, Oakland, CA, USA; Department of Genetics & UNC Neuroscience Center, University of North Carolina at Chapel Hill, Chapel Hill, NC, USA; Institute for Community Medicine, University Medicine Greifswald, Greifswald, Germany; Department of Human Genetics, Donders Institute for Brain, Cognition, and Behaviour, Radboud University Medical Center, Nijmegen, the Netherlands; Department of Clinical Genetics MUMC+, GROW School of Oncology and developmental biology, and MHeNs School of Mental Health and Neuroscience, Maastricht University, The Netherlands; Bordeaux Population Health, University of Bordeaux INSERM U1219, Bordeaux, France; West Region, Institute of Mental Health, Singapore; Yong Loo Lin School of Medicine, National University of Singapore, Singapore; Lee Kong Chian School of Medicine, Nanyang Technological University, Singapore; MRC Unit on Risk and Resilience, University of Cape Town, Cape Town, South Africa; Centre for Genomic Sciences, Li Ka Shing Faculty of Medicine, The University of Hong Kong, Hong Kong; Department of Medicine, Li Ka Shing Faculty of Medicine, The University of Hong Kong, Hong Kong; Day Clinic for Cognitive Neurology, University Hospital Leipzig, Leipzig, Germany; Bloorview Research Institute, Holland Bloorview Kids Rehabilitation Hospital, Toronto, Canada; Departments of Psychology and Psychiatry, University of Toronto, Toronto, Canada; Tri-institutional Center for Translational Research in Neuroimaging and Data Science (TReNDS), Atlanta, GA, USA; Laboratory of Epidemiology, Demography, and Biometry, Intramural Research Program, National Institute of Aging, The National Institutes of Health, Bethesda, MD, USA; Human Genetics Center, School of Public Health, University of Texas Health Science Center at Houston, Houston, TX, USA; Department of Neurology, Bordeaux University Hospital, Bordeaux, France; Univ. Lille, Inserm, CHU Lille, Institut Pasteur de Lille, U1167 - RID-AGE - Risk Factors and Molecular Determinants of Aging-Related Diseases and Labex Distalz, Lille, France; Department of Biomedical Engineering, Illinois Institute of Technology, Chicago, IL, USA; Rush Alzheimer’s Disease Center, Rush University Medical Center, Chicago, IL, USA; Brain Research Imaging Centre, University of Edinburgh, Edinburgh, UK; Department of Computer Science, Lagos State University, Lagos, Nigeria; Scottish Imaging Network, A Platform for Scientific Excellence (SINAPSE) Collaboration, Department of Neuroimaging Sciences, University of Edinburgh, Edinburgh, UK; Centre for Clinical Brain Sciences, University of Edinburgh, Edinburgh, UK; Edinburgh Imaging, University of Edinburgh, Edinburgh, UK; Centre for Brain Research, Indian Institute of Science, Bangalore, India; Memory Aging & Cognition Centre, Departments of Pharmacology and Psychological Medicine, Yong Loo Lin School of Medicine, National University of Singapore, Singapore; Duke-NUS Medical School, Singapore; Singapore Eye Research Institute, Singapore; Center for Translational and Computational Neuroimmunology, Department of Neurology and the Taub Institute for Research on Alzheimer’s disease and the Aging brain, Columbia University Irving Medical Center, New York NY, USA; Lothian Birth Cohorts, Department of Psychology, University of Edinburgh, Edinburgh, UK; Department of Neurological Sciences, Rush University Medical Center, Chicago, IL, USA; Department of Psychiatry and Behavorial Sciences, Rush University Medical Center, Chicago, IL, USA; Department of Neurology, Johns Hopkins University School of Medicine, Baltimore, MD, USA; Department of Epidemiology, Johns Hopkins University School of Medicine, Baltimore, MD, USA; Icelandic Heart Association, Kopavogur, Iceland; Faculty of Medicine, University of Iceland, Reykjavik, Iceland; Saw Swee Hock School of Public Health, National University of Singapore, Singapore; Department of Pharmacology, National University of Singapore, Singapore; Memory Aging and Cognition Center, National University Health System, Singapore; Institute for Medical Informatics, Statistics and Documentation, Medical University of Graz, Graz, Austria; Department of Cardiology, Leiden University Medical Center, Leiden, the Netherlands; Netherlands Heart Institute, Utrecht, the Netherlands; Netherlands Consortium of Healthy Ageing; Durrer Center for Cardiogenetic Research, Amsterdam, the Netherlands; Centre for Cognitive Ageing and Cognitive Epidemiology, Psychology, University of Edinburgh, Edinburgh, UK; Department of Biology, Maynooth University, Maynooth, Co. Kildare, Ireland; Department of Neurology and Psychiatry, University of Pittsburgh School of Medicine, Pittsburgh, PA, USA; Imaging of Dementia and Aging (IDeA) Laboratory, University of California at Davis, Davis, CA, USA; Faculty of Applied Sciences, Delft University of Technology, Delft, the Netherlands; GeneSTAR Research Program, Johns Hopkins University School of Medicine, Baltimore, MD, USA; Institute for Translational Genomics and Population Sciences, the Lundquist Institute for Biomedical Innovation at Harbor-UCLA Medical Center, Torrance, CA, USA; University of Miami Miller School of Medicine, Miami, FL, USA; Research Unit Genetic Epidemiology, Gottfried Schatz Research Center, Division of Molecular Biology and Biochemistry, Medical University Graz, Graz, Austria; Department of Social and Behavioural Sciences, Harvard T.H. Chan School of Public Health, Boston, MA, USA; Department of Internal Medicine, Section of Gerontology and Geriatrics, Leiden University Medical Center, Leiden, the Netherlands; Department of Radiology, Leiden University Medical Center, Leiden, the Netherlands; UK Dementia Research Centre, University of Edinburgh, Edinburgh, UK; NORMENT (Norwegian Centre for Mental Disorders Research), Institute of Clinical Medicine, University of Oslo, Oslo, Norway; Department of Psychiatric Research, Diakonhjemmet Hospital, Oslo, Norway; Centre for Psychiatry Research, Department of Clinical Neuroscience, Karolinska Institutet & Stockholm Health Care Services, Stockholm County Council, Stockholm, Sweden; Neurology Department, Yale University School of Medicine, New Haven, CT, USA; Department of Genetics, University of Pennsylvania Perelman School of Medicine, Philadelphia, PA, USA; Department of Biomedical and Health Informatics, Children’s Hospital of Philadelphia, Philadelphia, PA, USA; Academic Unit for Psychiatry of Old Age, University of Melbourne, St George’s Hospital, Kew, Australia; National Ageing Research Institute, Parkville, Victoria, Australia; CoE NORMENT, Division of Mental Health and Addiction, Oslo University Hospital, Oslo, Norway; Institute of Clinical Medicine, University of Oslo, Oslo, Norway; Mathematics & Statistics, Murdoch University, Perth, WA, Australia; Translational Medicine, Hospital for Sick Children, University of Toronto, Toronto, Canada; Department of Human Genetics, Brownsville, TX, USA; South Texas Diabetes and Obesity Institute, University of Texas Rio Grande Valley School of Medicine, Brownsville, TX, USA; Department of Psychiatry, UMC Utrecht, Utrecht, the Netherlands; VU university Amsterdam, Department of Biological Psychology & Netherlands Twin Register, Amsterdam, Netherlands; Taub Institute for Research on Alzheimer’s Disease and the Aging Brain, Department of Neurology, College of Physicians and Surgeons, Columbia University, New York, NY, USA; CHeBA (Centre for Healthy Brain Ageing), School of Psychiatry, UNSW Sydney, Sydney, Australia; Dementia Centre for Research Collaboration, University of New South Wales, Sydney, Australia; Department of Psychology and Center for Brain Science, Harvard University, Boston, MA, USA; Department of Radiology, Massachusetts General Hospital, Boston, MA, USA; Department of Psychiatry, Massachusetts General Hospital, Boston, MA, USA; Department of Cognitive Neuroscience, Donders Institute for Brain, Cognition and Behavior, Radboudumc, Nijmegen, the Netherlands; Centre for Neuroimaging & Cognitive Genomics (NICOG), Clinical Neuroimaging Laboratory, NCBES Galway Neuroscience Centre, College of Medicine Nursing and Health Sciences, National University of Ireland Galway, Galway, Ireland; Department of Psychiatry, School of Clinical Sciences, Monash University, Melbourne, Victoria, Australia; University of Queensland, Brisbane, QLD, Australia; University of Sydney, Sydney, Australia; Computational Brain Anatomy (CoBrA) Laboratory, Cerebral Imaging Centre, Douglas Research Centre, Montreal, QC, Canada; Departments of Psychiatry and Biological and Biomedical Engineering, McGill University, Montreal, QC, Canada; Lieber Institute for Brain Development, Baltimore, MD, USA; Department of Psychiatry, Trinity Translational Medicine Institute, Trinity College, Dublin, Ireland; Hospital Universitario Virgen del Rocío, Department of Psychiatry, IBiS, University of Sevilla, Sevilla, Spain; CIBERSAM, Centro Investigación Biomédica en Red Salud Mental, Sevilla, Spain; Avera Institute for Human Genetics, Sioux Falls, SD, USA; Department of Biological Psychology, Vrije Universiteit Amsterdam, Amsterdam, the Netherlands; Amsterdam Public Health Research Institute, Amsterdam UMC, Amsterdam, the Netherlands; Faculty of Health and Institute of Health and Biomedical Innovation, Queensland University of Technology, Brisbane, Australia; MRC-SGDP Centre, Institute of Psychiatry, King’s College London, London, UK; Cell Biology and Gene Expression Section, Laboratory of Neurogenetics, National Institute on Aging, National Institutes of Health, Bethesda, Maryland, USA; Department of Medical Genetics, Oslo University Hospital, Oslo, Norway; NORMENT, Department of Clinical Science, University of Bergen, Bergen, Norway; Janssen Research & Development, LLC, of Johnson & Johnson, San Diego, CA, USA; Division of Psychological and Social Medicine and Developmental Neurosciences, Faculty of Medicine, TU Dresden, Germany; Department of Psychiatry and Psychotherapy, Charité Universitätsmedizin Berlin, corporate member of Freie Universität Berlin, Humboldt-Universität zu Berlin, and Berlin Institute of Health, Berlin, Germany; Bjørknes College, Oslo, Norway; Indiana University School of Medicine, Department of Medical and Molecular Genetics, Indianapolis, IN, USA; Psychiatric and Neurodevelopmental Genetics Unit, Center for Genomic Medicine, Massachusetts General Hospital, Boston, MA, USA; Dr Einar Martens Research Group for Biological Psychiatry, Center for Medical Genetics and Molecular Medicine, Haukeland University Hospital, Bergen, Norway; Department of Psychiatry, Boston Children’s Hospital and Harvard Medical School, Boston, MA, USA; Brigham and Women’s Hospital, Boston, MA, USA; The Broad Institute, Boston, MA, USA; Harvard Medical School, Boston, MA, USA; Karakter, Child and Adolescent Psychiatry, University Center, Nijmegen, the Netherlands; King’s College London, Social, Genetic and Developmental Psychiatry, Institute of Psychiatry, Psychology and Neuroscience, London, UK; Department of Psychiatry, Psychosomatic Medicine and Psychotherapy, University Hospital, Goethe University, Frankfurt, Germany; Queensland Brain Institute, University of Queensland, Brisbane, Australia; University of Groningen, University Medical Center Groningen, Interdisciplinary Center Psychopathology and Emotion regulation, Groningen, the Netherlands; Department of Pathology of Mental Diseases, National Institute of Mental Health, National Center of Neurology and Psychiatry, Kodaira, Tokyo, Japan; School of Medicine & Public Health, University of Newcastle, Callaghan, NSW, Australia; Health Behaviour Research Group, University of Newcastle, Callaghan, NSW, Australia; Genentech, Inc., South San Francisco, CA, USA; Department of Psychiatry, University of Iowa Carver College of Medicine, Iowa City, IA, USA; University of Groningen, University Medical Center Groningen, Department of Child and Adolescent Psychiatry, Groningen, Netherlands; Department of Psychology, Yale University, New Haven, Connecticut, USA; Department of Psychiatry, Yale University, New Haven, Connecticut, USA; Brain Center Rudolf Magnus, Department of Psychiatry, UMC Utrecht, Utrecht, the Netherlands; University of Western Australia, Perth, Australia; Wellcome centre for Integrative Neuroimaging (WIN), Nuffield Department of Clinical Neurosciences, University of Oxford, John Radcliffe Hospital, Oxford, UK; Australian Institute of Machine Learning (AIML), Department of Computer Science, University of Adelaide, SA, Australia; South Australian Health and Medical Research Institute (SAHMRI), North Terrace, Adelaide, SA, Australia; Institute of Science and Technology for Brain-Inspired Intelligence, Fudan University, Shanghai, China; Key Laboratory of Computational Neuroscience and Brain-Inspired Intelligence (Fudan University), Ministry of Education, China; Centre for Population Neuroscience and Precision Medicine, MRC SGDP Centre, IoPPN, King’s College London, London, UK; Institute of Medical Informatics, Biometry and Epidemiology, University of Duisburg-Essen, Essen, Germany; NORMENT – TOP study, Institute of Clinical Medicine, Psychiatry section, University of Oslo, Oslo, Norway; Electrical and Computer Engineering, State University of New York, Oswego, NY, USA; Center for Neuroimaging, Department of Radiology and Imaging Sciences, Indiana University School of Medicine, Indianapolis, IN, USA; Department of Psychiatry, University of California San Diego, CA, USA; Maryland Psychiatric Research Center, Department of Psychiatry, University of Maryland School of Medicine, Baltimore, MD, USA; Brain and Mind Centre, The University of Sydney, Camperdown, NSW, Australia; School of Medical Sciences, University of New South Wales, Sydney, Australia; Division of Psychiatry, University of Edinburgh, Royal Edinburgh Hospital, Edinburgh, Scotland, UK; Hunter New England Health, New Lambton, NSW, Australia; University of Newcastle, Callaghan, NSW, Australia; QIMR Berghofer Medical Research Institute, Brisbane, Queensland, Australia; INSERM U A10 “Trajectoires développementales & psychiatrie”, Université Paris-Saclay, Ecole Normale Supérieure Paris-Saclay, CNRS, Centre Borelli, Gif-sur-Yvette, France; UCL Institute of Neurology, London, United Kingdom and Epilepsy Society, UK; Department of Psychiatry and Behavioral Sciences and Weill Institute for Neurosciences, University of California, San Francisco, CA, USA; Veterans Affairs San Francisco Healthcare System, San Francisco, CA, USA; Department of Radiology, Johns Hopkins University School of Medicine, Baltimore, MD, USA; Genetic Basis of Mood & Anxiety Section, Human Genetics Branch, NIMH Intramural Research Program, NIH, Bethesda, MD, USA; School of Clinical Sciences, Institute of Health and Biomedical Innovation, Queensland University of Technology, Brisbane, Australia; Menzies Institute for Medical Research, University of Tasmania, Hobart, Tasmania, Australia; Centre for Law and Genetics, Faculty of Law, University of Tasmania, Hobart, Tasmania, Australia; Section of Gerontology and Geriatrics, Department of Medicine, University of Perugia, Perugia, Italy; NORMENT Centre, Oslo University Hospital, Oslo, Norway; Clinical Department of Psychiatry and Psychotherapy, Central Institute of Mental Health, Medical Faculty Mannheim, Heidelberg University, Mannheim, Germany; School of Psychology, University of Newcastle, Callaghan, NSW, Australia; Department of Psychiatry, Amsterdam Public Health and Amsterdam Neuroscience, Amsterdam UMC/Vrije Universiteit & GGZinGeest, Amsterdam, the Netherlands; Queensland Centre for Mental Health Research, The University of Queensland, Brisbane, QLD, Australia; Oxford Big Data Institute, Li Ka Shing Centre for Health Information and Discovery, Nuffield Department of Population Health, University of Oxford, Oxford, UK; Departments of Psychiatry and Pediatrics, UT Health San Antonio, San Antonio, TX, USA; Emma Children’s Hospital, Amsterdam UMC, University of Amsterdam, Emma Neuroscience Group, department of Pediatrics, Amsterdam Reproduction & Development, Amsterdam, the Netherlands; Vrije Universiteit, Clinical Neuropsychology section, Amsterdam, the Netherlands; UCLA Center for Neurobehavioral Genetics, Los Angeles, California, USA; Department of Neurology, Hopital Erasme, Université Libre de Bruxelles, Brussels, Belgium; ULB Laboratory of Experimental Neurology, Brussels, Belgium; Melbourne Neuropsychiatry Centre, Department of Psychiatry, University of Melbourne, Melbourne, Victoria, Australia; NorthWestern Mental Health, Sunshine Hospital, Sunshine, Victoria, Australia; Florey Institute for Neuroscience and Mental Health, Parkville, Victoria, Australia; Department of Psychiatry, Amsterdam UMC, Vrije Universiteit, Amsterdam, the Netherlands; Departments of Radiology and Clinical Neurosciences, University of Calgary, Calgary, Alberta, Canada; Priority Research Centres for Brain and Mental Health Research and Stroke and Brain Injury, University of Newcastle, Newcastle, Australia; Hunter Medical Research Institute, Newcastle, NSW, Australia; Department of Genetics & Computational Biology, QIMR Berghofer Medical Research Institute, Brisbane, QLD, Australia; Department of Developmental Disability Neuropsychiatry, School of Psychiatry, University of New South Wales, Sydney, Australia; Department of Genetic Epidemiology in Psychiatry, Central Institute of Mental Health, Medical Faculty Mannheim, University of Heidelberg, Heidelberg, Mannheim, Germany; Department of Radiology and Imaging Sciences, Indiana University School of Medicine, Indianapolis, IN, USA; Program for Translational Research on Adversity and Neurodevelopment (P-TRAN), Edna Bennett Pierce Prevention Research Center, The Pennsylvania State University, Pennsylvania, USA; Neuropsychiatric Institute, The Prince of Wales Hospital, Sydney, Australia; Max Planck Institute of Psychiatry, Munich, Germany; Centre for Population Neuroscience and Stratified Medicine (PONS), Institute of Psychiatry, Psychology and Neuroscience, SGDP-Centre, King’s College London, London, UK; Institute for Science and Technology of Brain-Inspired Intelligence (ISTBI), Fudan University, Shanghai, China; NSW Health Pathology, Newcastle, NSW, Australia; Department of Biostatistics, Epidemiology and Informatics, Institute for Biomedical Informatics, Perelman School of Medicine, University of Pennsylvania, Philadelphia, PA, USA; UCL Queen Square Institute of Neurology, London, UK; Chalfont Centre for Epilepsy, Bucks, UK; Institute of Clinical Medicine, Neurology, University of Eastern Finland, Kuopio, Finland; Peninsula Clinical School, Central Clinical School, Monash University, Australia; Laboratory of Neuro imaging, USC Stevens Neuroimaging and Informatics Institute, University of Southern California, Los Angeles, CA, USA; Neuroimaging Unit, Technological Facilities, Valdecilla Biomedical Research Institute IDIVAL, Spain; Centro Investigación Biomédica en Red de Salud Mental (CIBERSAM), Spain; Psychology Department & Neuroscience Institute, Georgia State University, Atlanta, GA, USA; NORMENT, KG Jebsen Centre for Psychosis Research, Division of Mental Health and Addiction, Oslo University Hospital & Institute of Clinical Medicine, University of Oslo, Oslo, Norway; School of Mental Health and Neuroscience, Faculty of Health, Medicine and Life Sciences, Maastricht University, Maastricht, the Netherlands; Department of Psychiatry, Leiden University Medical Center, Leiden, the Netherlands; Leiden Institute for Brain and Cognition, Leiden University Medical Center, Leiden, the Netherlands; Amsterdam Neuroscience, Amsterdam, the Netherlands; Departments of Psychiatry, Neurology, Neuroscience and Genetic Medicine, Johns Hopkins University, Baltimore, MD, USA; KG Jebsen Centre for Neurodevelopmental Disorders, University of Oslo, Oslo, Norway; Division of Clinical Geriatrics, Department of Neurobiology, Care Sciences and Society, Karolinska Institutet, Stockholm, Sweden; National Institute of Mental Health, National Institutes of Health, Bethesda, MD, USA; Human Genetics Branch, National Institute of Mental Health (NIMH), National Institute of Health (NIH), Bethesda, MD, USA; Centre for Advanced Imaging, University of Queensland, Brisbane, Queensland, Australia

## Abstract

The size of the human head is determined by growth in the first years of life, while the rest of the body typically grows until early adulthood^1^. Such complex developmental processes are regulated by various genes and growth pathways^2^. Rare genetic syndromes have revealed genes that affect head size^3^, but the genetic drivers of variation in head size within the general population remain largely unknown. To elucidate biological pathways underlying the growth of the human head, we performed the largest genome-wide association study on human head size to date (N = 79,107). We identified 67 genetic loci, 50 of which are novel, and found that these loci are preferentially associated with head size and mostly independent from height. In subsequent neuroimaging analyses, the majority of genetic variants demonstrated widespread effects on the brain, whereas the effects of 17 variants could be localized to one or two specific brain regions. Through hypothesis-free approaches, we find a strong overlap of head size variants with both cancer pathways and cancer genes. Gene set analyses showed enrichment for different types of cancer and the p53, Wnt and ErbB signalling pathway. Genes overlapping or close to lead variants – such as *TP53*, *PTEN* and *APC* – were enriched for genes involved in macrocephaly syndromes (up to 37-fold) and high-fidelity cancer genes (up to 9-fold), whereas this enrichment was not seen for human height variants. This indicates that genes regulating early brain and cranial growth are associated with a propensity to neoplasia later in life, irrespective of height. Our results warrant further investigations of the link between head size and cancer, as well as its clinical implications in the general population.

## Main

To gain more insight into the genetic underpinnings of the human head size, we performed a metaanalysis of genome-wide association studies (GWAS) by including samples measuring head size using intracranial volume from magnetic resonance imaging or computed tomography, and tape measured head circumference (**Table S1-S4; Online Methods**). Compared to previous efforts^4,5^, we nearly doubled the sample size (N = 79,107), of which the majority were of European ancestry (N = 75,309). We identified 90 independent genetic variants in 67 loci associated with human head size in the European sample (**Figure 1A; Table S5-S7**), of which 50 loci were novel. Most variants (N = 48) showed consistent directions of association between the European, African (N = 1,356), and Asian (N = 1,335) ancestry samples (**Figure 1B**), while nominally significant heterogeneity was observed for five variants (**Table S6**), suggesting population-specific genetic effects on head size in these loci.

**Figure 1.**
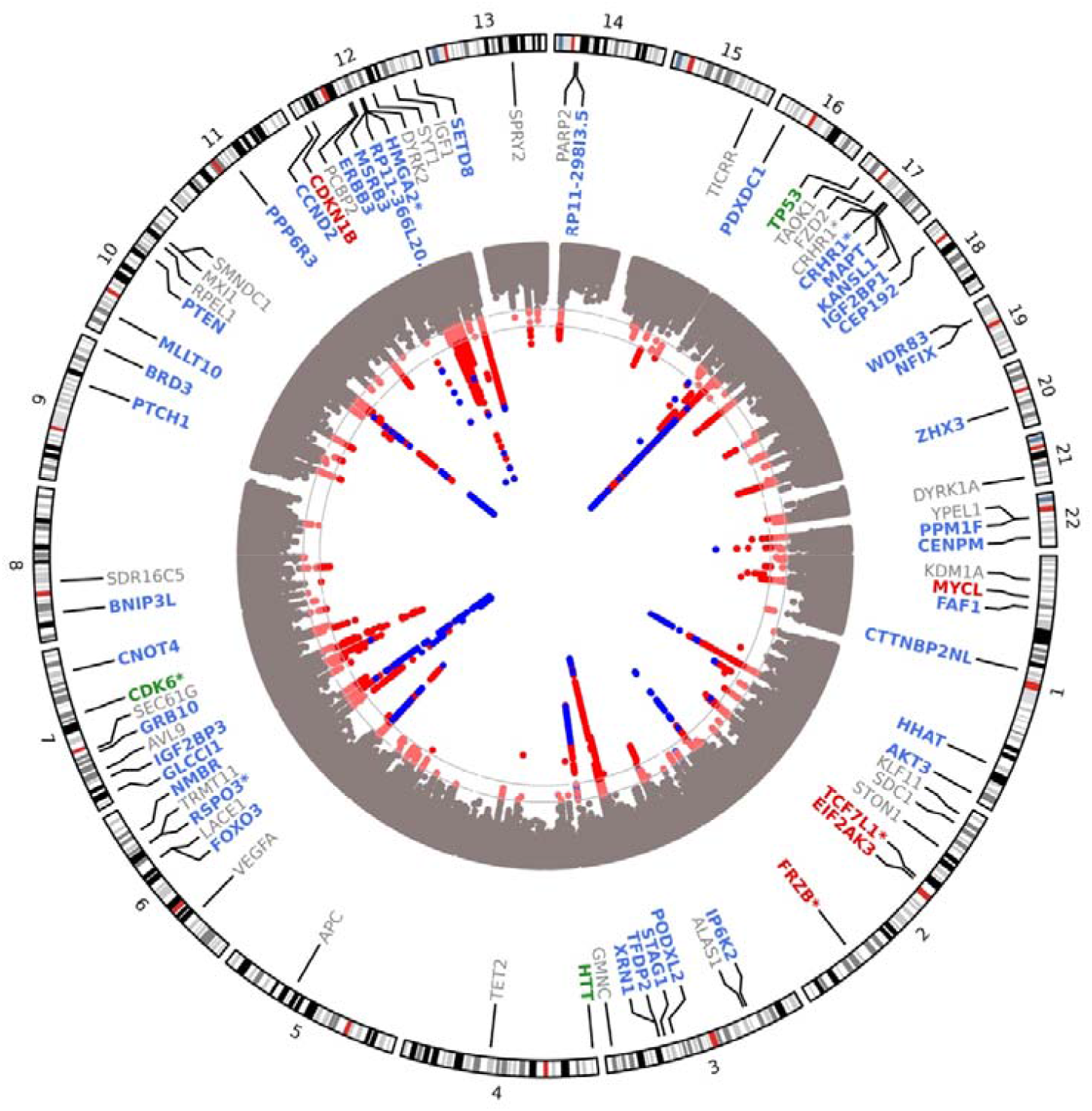
Genome-wide association studies on human head size. **Figure 1A.** Circos Manhattan plot of the European ancestry GWAS on head size, with the grey horizontal lines corresponding to a genome-wide significant (*P* < 5 x 10^−8^) or sub-significant (*P* < 1 x 10^−6^) *P* value threshold. Known genetic variants are depicted in blue, whereas novel variants are depicted in red. For each lead genetic variant, the nearest gene is shown with their corresponding location on the genome. The colour of each gene corresponds to its position to the lead variant: exonic (red), 3’-UTR (green), intronic (blue), intergenic including up- and downstream, exonic and intronic non-coding RNA (grey). Genes that are the nearest gene for more than one locus are denoted with an asterisk (*).

**Figure 1B.**
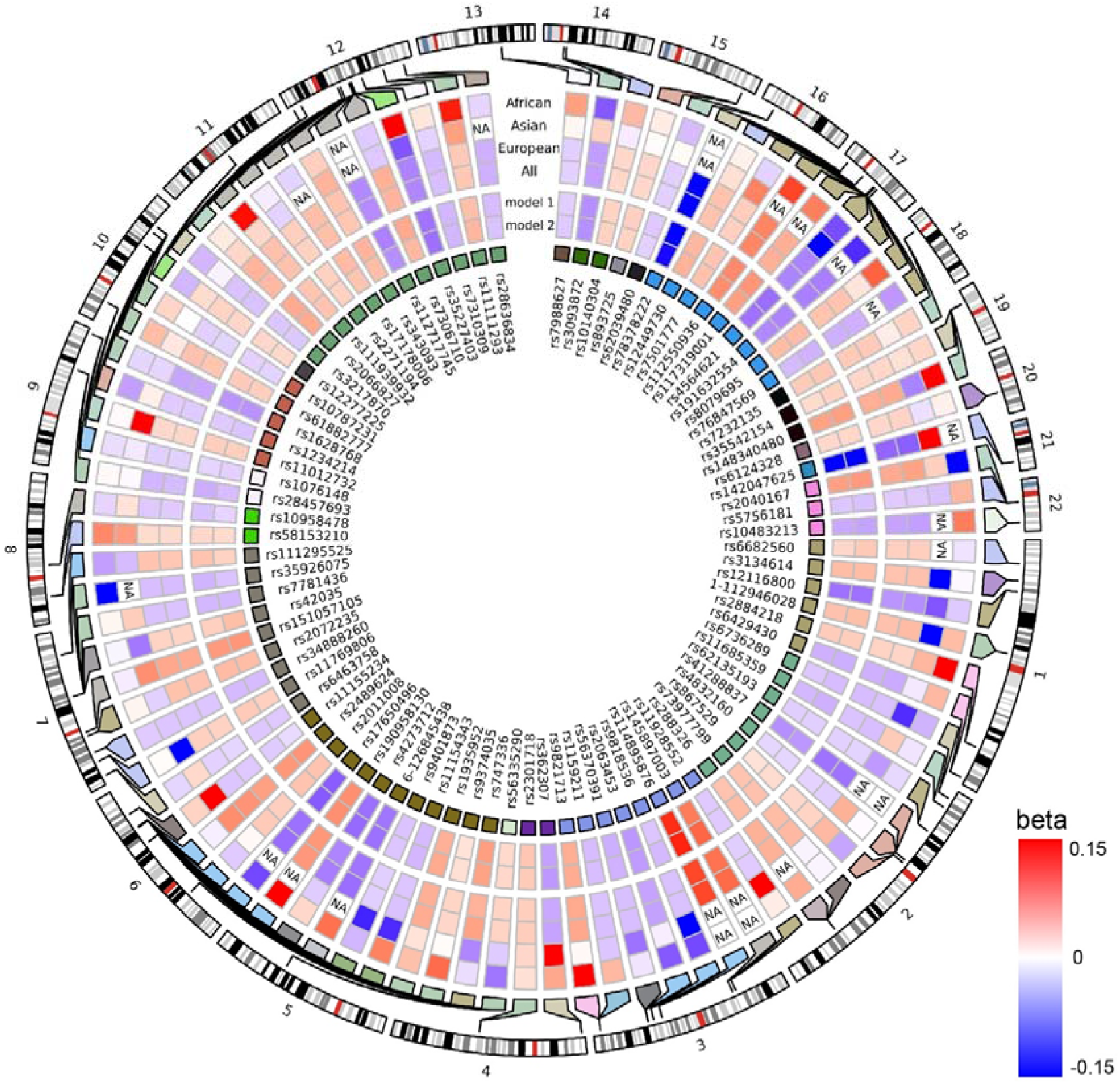
Circos heatmap showing the betas of the 90 identified lead genetic variants in African, Asian and European ancestry sample meta-analysis, as well as the transancestral meta-analysis. In addition, the differences between the height-unadjusted (model 1) and height-adjusted (model 2) meta-analysis is shown. Positive associations are depicted in red, negative associations are depicted in blue.

**Figure 1C.**
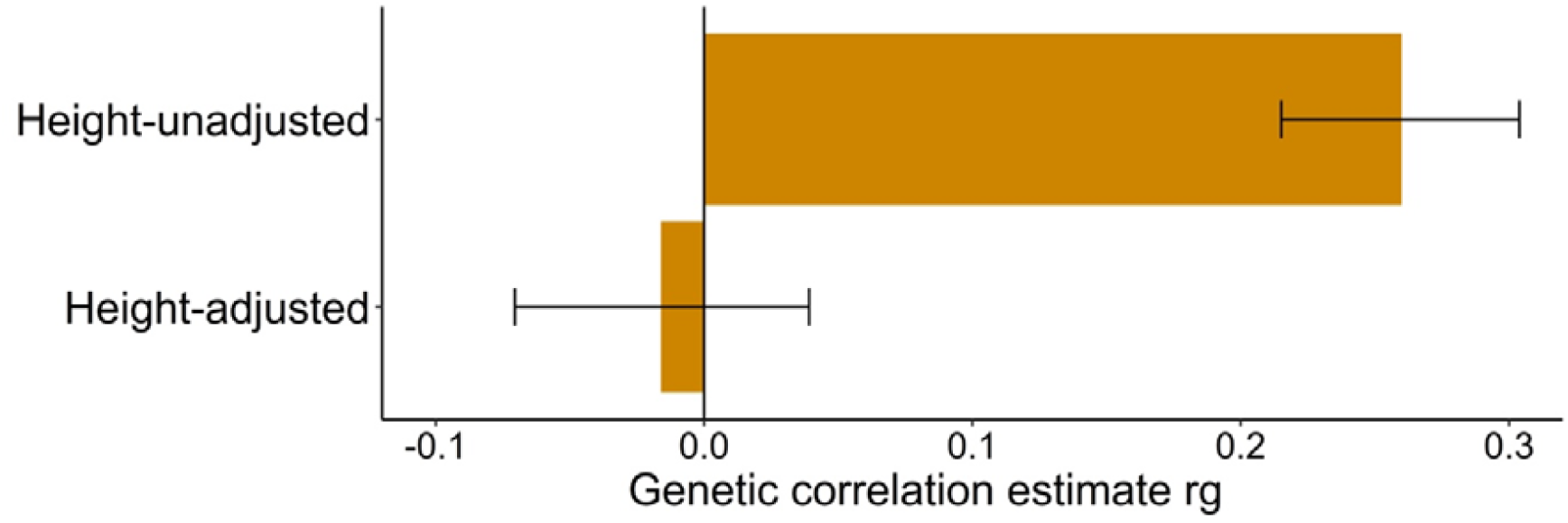
Barplot of the genetic correlation coefficient (ρ_genetic_) of the height-unadjusted and height-adjusted head size genome-wide association study with the height genome-wide association study, with their accompanying 95% confidence intervals.

### Head-specific growth versus general growth

Head growth coincides with growth of the entire body, prompting us to investigate whether variants affecting head size are specific for growth of the human brain and cranium or whether this is driven at least partly by an effect on human body height. We therefore performed an additional height-adjusted head size GWAS in European studies for which height measures were also available (N = 50,424). The genetic correlation between head size and height (ρ_genetic_ = 0.26, *P* = 2.1 x 10^−30^) disappeared in this second model (p_genetic_= −0.02, *P* = 0.58) (**Figure 1C**), confirming the removal of height-associated effects. Importantly, there was no significant attenuation for any of the lead variants’ effect sizes for their association with head size (**Table S6**). We further explored the effect of these variants on the size of other body parts using area measures obtained from bone density scans (N = 3,313). As expected, a polygenic score of the lead variants was associated with the skull area, even after adjusting for height (*P* = 2.1 x 10^−12^). One lead genetic variant (rs12277225) was significantly associated with the L1-L4 spine area (*P* = 1.3 x 10^−5^), but the other lead variants did not affect bone area measures of arm, leg, and spine (**Table S8**). Altogether, this indicates that the effect of the identified variants on head size is predominantly cranium-specific.

### Regional brain volumetric effects

Height is an overall measure reflective of growth in various body parts. Accordingly, head size itself may also reflect growth of specific brain regions. Indeed, 15 lead genetic variants or variants in LD (r^2^>0.6) from 12 genetic loci were previously reported to affect volumes of subregions of the brain (**Figure 2A; Table S9**). We further screened all loci previously associated with these regional brain volumes, and found 16 of those 132 loci to be significantly related with head size in our data set after multiple testing correction (**Table S10**). To determine if the current findings can be localized to specific brain regions, we systematically investigated the 90 independent head size variants in relation to more fine-grained measures of brain morphometry – corrected for head size – in 22,145 individuals (**Figure 2B; Table S11**). Twenty-nine variants were associated with multiple cortical, subcortical, and global brain regions, and for the other 51 variants there was no apparent predilection to influence particular brain regions. However, seventeen variants were preferentially associated with one or two specific cortical or subcortical regions. For example, rs111939932 was associated with nucleus accumbens volume. This intronic variant in *PCBP2* is an eQTL for different genes in multiple tissues, including *ATP5G2* in the nucleus accumbens and basal ganglia of the brain. Further analysis additionally revealed localized effects of this variant on the shape of this structure (**Figure 2C; Table S12**). In the largest GWAS on nucleus accumbens volume to date^6^, this variant was nominally significant (*P* = 0.02), underlining the improved power of the current study to identify novel loci for brain morphometry. Overall, these results suggest that most head size variants are important for generalized brain or cranial growth, while a minority influences regional brain growth.

**Figure 2.**
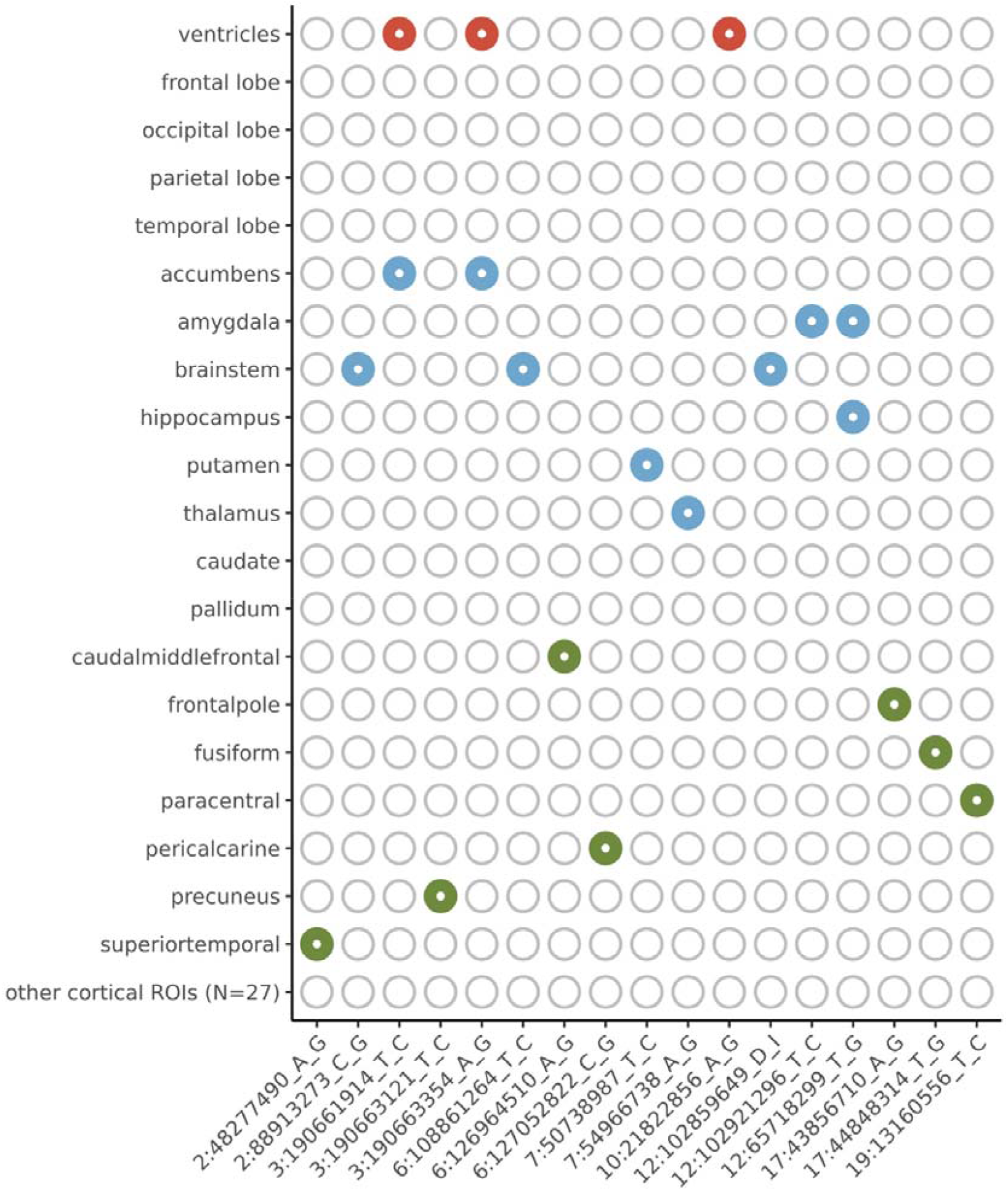
Genetic loci for head size and effects on regional brain volumes. **Figure 2A.** Heatmap showing the genetic loci identified for human head size that overlap with previously identified genetic loci for global brain volumes (depicted in red), subcortical brain volumes (depicted in blue) and cortical regional of interest volumes (depicted in green).

**Figure 2B.**
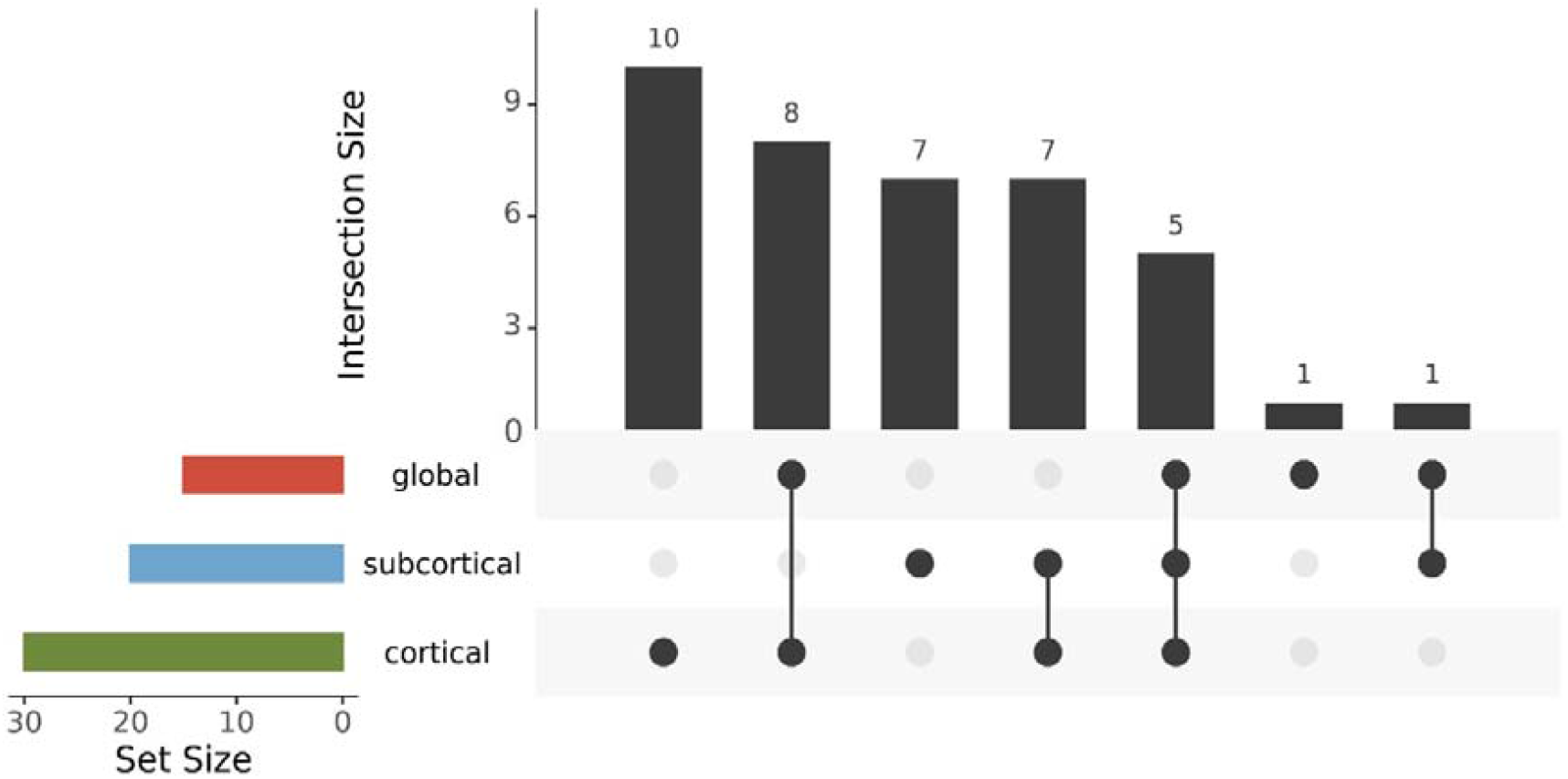
UpSet plot of the different combinations of associations of the identified genetic variants for human head size and regional brain volumes. The intersection size corresponds to the frequency of the combination depicted below the bar. The set size corresponds to the frequency of associations with one of the structures belonging to the brain volume category (i.e., global, subcortical or cortical). Global volumes include the volumes of four brain lobes and the lateral ventricle volumes (depicted in red), subcortical volumes include the volumes of eight subcortical structures (depicted in blue), and the cortical volumes include the volumes of 34 cortical regions of interest (depicted in green).

**Figure 2C.**
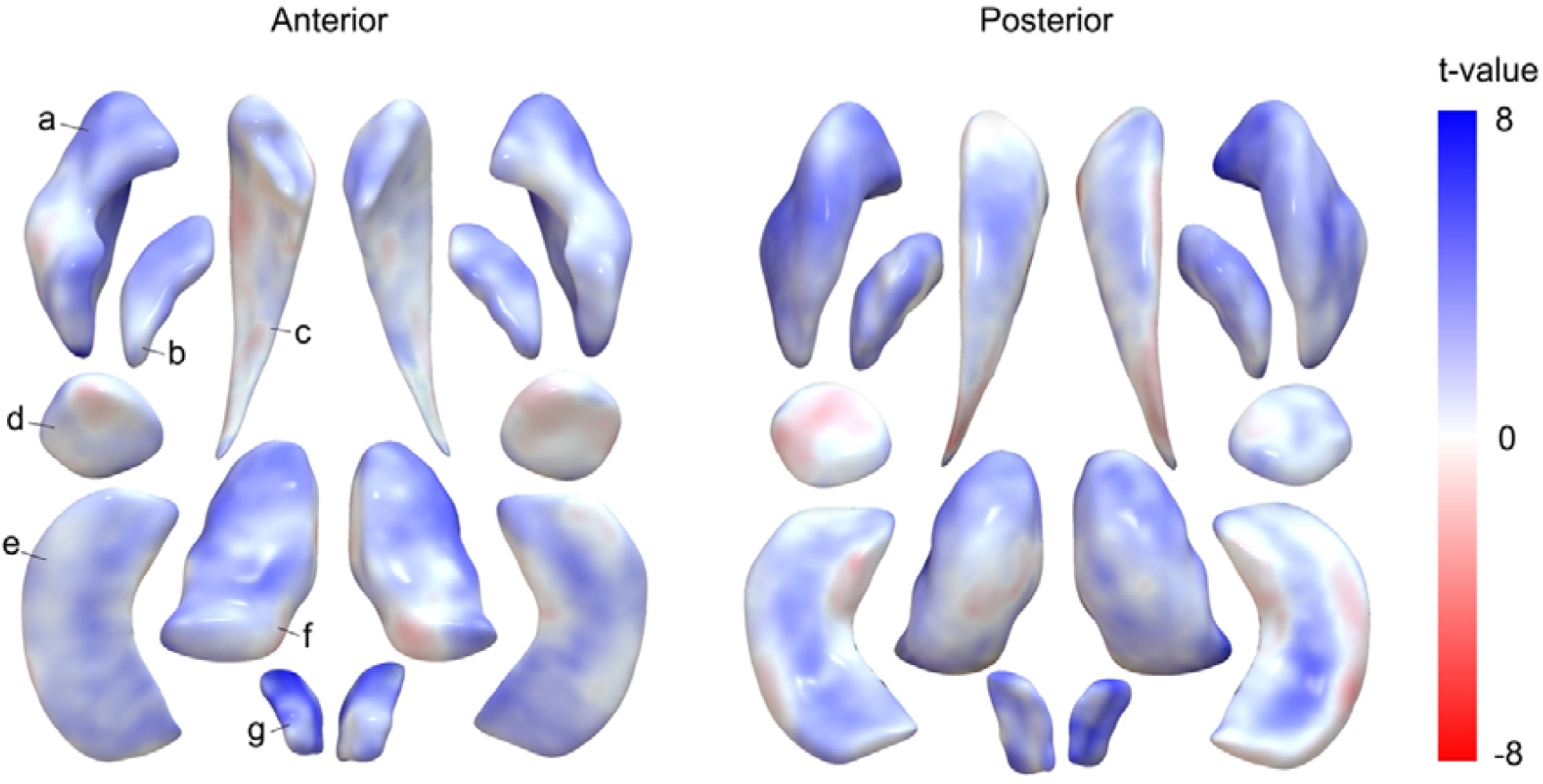
Plot showing the results of the subcortical shape analysis of rs111939932 using log Jacobian determinants. Colours correspond to t-values, with positive associations depicted in blue, and negative associations depicted in red. The letters point to the different subcortical structures: a – putamen; b – pallidum; c – caudate; d – amygdala; e – hippocampus; f – thalamus; g – accumbens.

### Pathway analysis

To obtain novel insights into the biological mechanisms underlying variation in human head size, we performed a hypothesis-free gene set enrichment analysis of all KEGG^7^ gene sets and found 14 to be significantly enriched (**Figure 3A; Table S13**). Nine of those gene sets represent different cancer types that substantially overlap between each other and share underlying biological pathways (**Figure 3B**). The remaining gene sets represent the p53, Wnt and ErbB signalling pathways, which are all involved in tumorigenesis including in the above cancer types^8^. Remarkably, the lead variants were often intragenic for the overlapping 7 genes in the p53 pathway, 8 genes in the Wnt pathway and 6 genes in the ErbB-EGFR pathway (**Figure 3C**), suggesting that modulation of these pathways plays an important role in head size variation.

**Figure 3.**
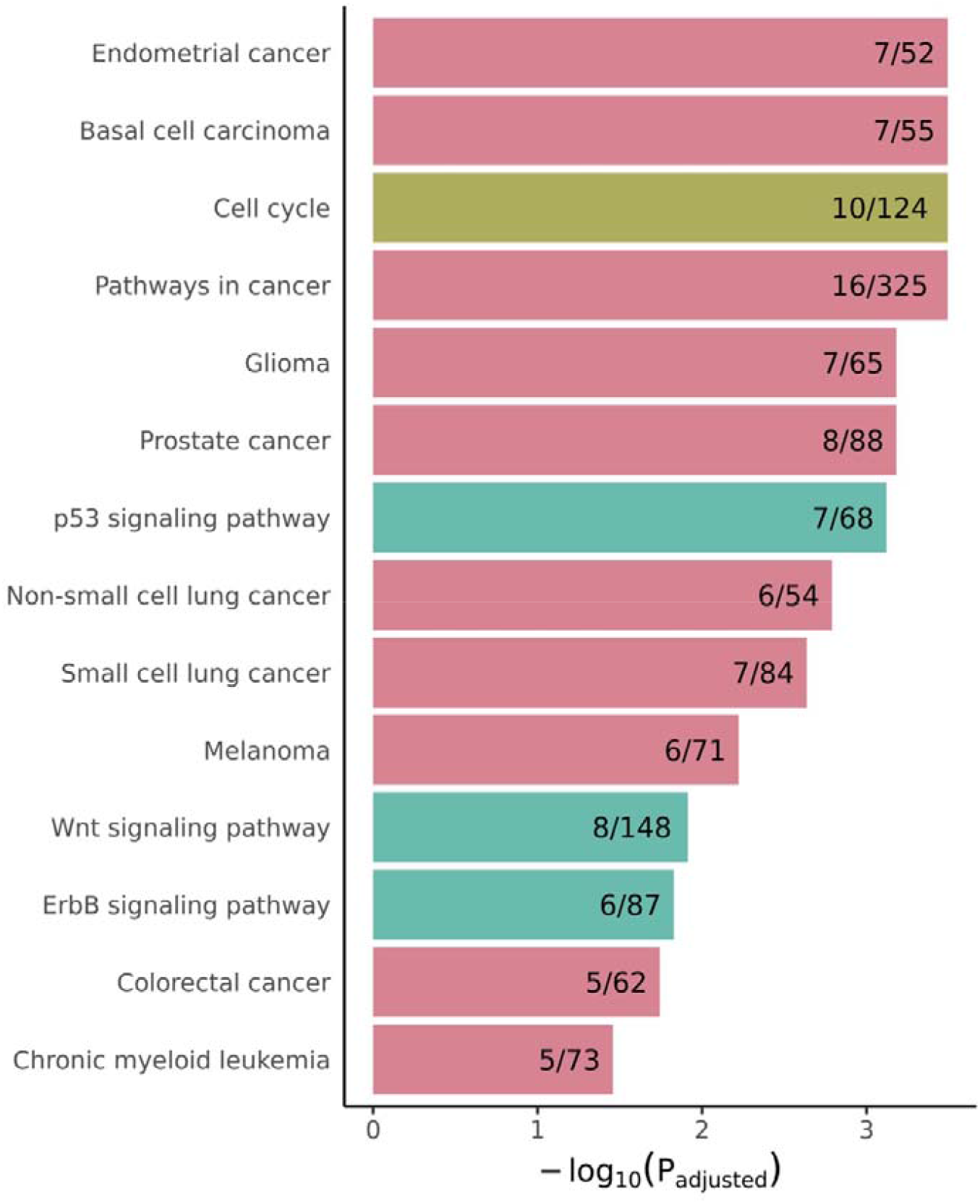
Gene sets enriched in human head size loci. **Figure 3A.** Barplots presenting the significantly enriched KEGG gene sets. On the x-axis the –log_10_ of the adjusted p-value is presented, and the proportion of genes in the gene set that overlap with the genes nearby the genetic loci are shown inside the bars. Colours correspond to different categories of gene sets: cancer gene sets are depicted in pink, cell growth and death gene sets in yellow-green, and signal transduction gene sets in turquoise.

**Figure 3B.**
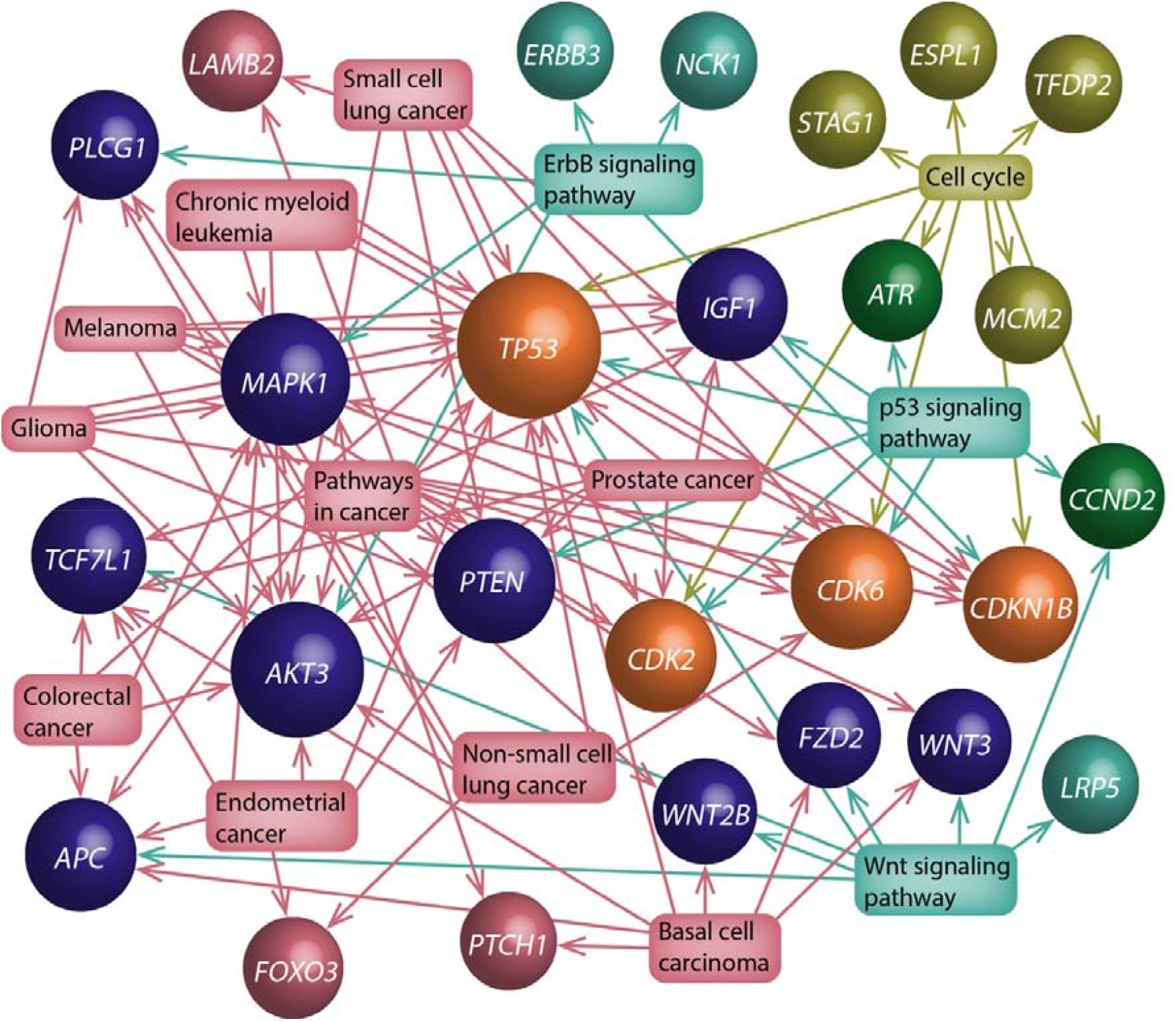
Network graph showing the enriched KEGG gene sets and their included genes near genetic lead variants. Gene sets are shown in squares, with arrows connecting them to the overlapping genes presented as spheres. The colours of the spheres correspond to the gene set category the gene is linked to: only cancer gene sets (pink), only cell growth and death gene sets (yellow-green), only signal transduction gene sets (turquoise), cancer gene sets and cell growth and death gene sets (dark blue), cell growth and death gene sets and signal transduction gene sets (green), or all three gene set categories (orange). The size of a sphere corresponds to the amount of gene sets linked to that gene.

**Figure 3C.**
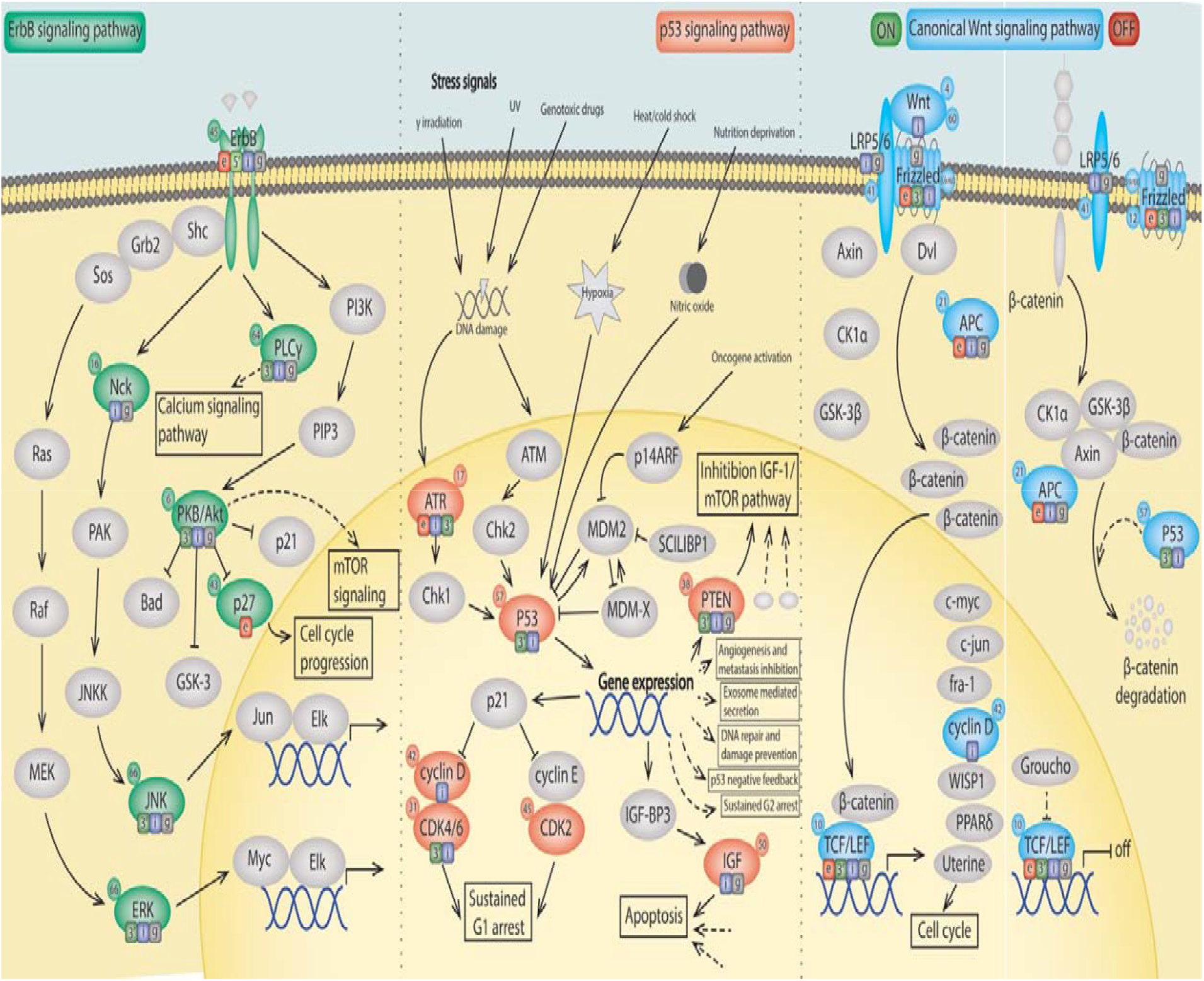
Schematic overview of the significantly enriched signalling pathways with proteins encoded by genes near (< 10 kb) identified genetic loci. Proteins encoded by these genes are coloured (green – ErbB signalling pathway, red – p53 signalling pathway; blue – Wnt signalling pathway), whereas the other proteins are depicted in grey. The circle next to each protein name provides the locus number to which the encoding gene belongs. Locations of lead genetic variants and variants in linkage disequilibrium (r^2^ > 0.6) are shown in the squares within each protein: exonic (e; red), 3’-UTR (3’; green), 5’-UTR (5; light green), intronic (i; blue), intergenic including up- and downstream, exonic and intronic non-coding RNA (g; grey). For Frizzled, not only *FZD2* but also *FRZB* is taken into consideration.

#### P53 signalling pathway

The signalling pathway showing the strongest enrichment was the p53 signalling pathway (*P*_adjusted_ = 7.6 x 10^−4^) (**Figure 3C**). The tumour suppressor protein p53, encoded by *TP53,* is activated by different stress signals to regulate the cell cycle and apoptosis. Our lead signal in this locus was the *TP53* 3’-UTR variant rs78378222 with predicted deleterious effects (CADD = 15.93), which was identified previously^5^. Three other genes in this pathway *(ATR, CDK6* and *PTEN)* also contained 3’-UTR or exonic variants in LD (r^2^ > 0.6) with the identified lead variants. As we identified genes involved in cell cycle arrest and cellular senescence *(CDK6, CDK2* and *CCND2),* apoptosis *(IGF1)* and inhibition of the IGF-1/mTOR pathway *(PTEN),* our results suggest a comprehensive involvement of the p53 signalling pathway in cranial growth. This finding is in line with evidence that p53 signalling regulates both normal and malignant neural stem cell populations^9–11^.

#### Wnt signalling pathway

The Wnt signalling pathway has extensive links to carcinogenesis, but also plays pivotal roles in the developing and adult central nervous system^12,13^, as well as in bone development including cranial growth^14^. Of the eight overlapping genes, three contained exonic or 3’-UTR variants in LD (r^2^>0.6) with identified lead variants *(APC, TP53* and *TCF7L1).* The Wnt signalling pathway gene *FRZB,* not annotated in KEGG, also contained exonic and 3’-UTR variants. In total, 1,948 genetic variants in LD with the identified lead variants (r^2^>0.6), among which 35 exonic variants, are eQTLs for *WNT3* in 27 different tissues including the cerebellar hemispheres. In addition, various exonic, 3’-UTR and 5’-UTR variants in LD with the lead variants are eQTLs for *TCF7L1* in brain tissues. Altogether, these observations suggest that this pathway is critical for brain and cranial growth in humans.

#### ErbB signalling pathway

The third enriched signalling pathway was the ErbB pathway (*P*_ad_j_uste_d = 0.014), also known as the EGFR signalling pathway, with six overlapping genes. Overlapping genes near head size variants are involved in the downstream calcium signalling (*PLCG1*), MAPK signalling *(NCK1* and *MAPK1)* and PI3K-AKT signalling *(ERBB3, AKT3* and *CDKN1B)* pathways. In addition, five genetic variants are eQTLs for *EGFR* in the cerebellum. Interestingly, both *AKT3* and *CDKN1B* have been linked to clinical head size syndromes and cancer risk^15–18^ and contain, respectively, 3’-UTR variants and an exonic variant that reached genome-wide significance in the current study. This ErbB signalling is also increasingly recognized for its involvement in neurodevelopment^19–21^, making it a plausible pathway involved in head size variations.

#### P53, Wnt and ErbB signalling pathway in general growth

Since these signalling pathways have universal roles in cell growth, and thus are not specific for head size, we determined the enrichment for these pathways in the height GWAS. We found that from these three signalling pathways, only the Wnt signalling pathway was significantly enriched in the height GWAS (*P*_adjusted_ = 3.8 x 10^−2^), suggesting that the p53 and ErbB signalling pathways are more specifically involved in processes for head growth rather than generalized body growth.

### Enrichment analyses

Because pathway analyses aggregate all genes in the vicinity of the lead variant, it becomes difficult to discern actual target genes. Given that target genes of GWAS variants are often close to the lead variant^22^, we determined the enrichment of different categories of genes located nearby head size variants stratified by their distance (**Table S14**).

#### OMIM macro- and microcephaly genes

First, we investigated genes mutated in OMIM syndromes associated with abnormal head size, i.e. macrocephaly or microcephaly (**Table S15-16**). We found increasing enrichment for macrocephaly genes with decreasing distance to the lead variants, culminating in a 37-fold enrichment of macrocephaly genes in genes containing an intragenic lead variant. In contrast, microcephaly genes did not enrich upon shorter distance from lead variants (**Figure 4A**). The striking enrichment of macrocephaly genes did not change in the height-adjusted GWAS (**Table S17**). Furthermore, there was only a modest enrichment for macrocephaly genes in the height GWAS, even for the top 67 loci (i.e., the same number of loci as our GWAS; **Table S17**). Macrocephaly genes with intragenic lead variants include *AKT3* (Megalencephaly-polymicrogyria-polydactyly-hydrocephalus syndrome 2), *PTCH1* (Basal cell nevus syndrome), *PTEN* (Cowden syndrome 1), *CCND2* (Megalencephaly-polymicrogyria-polydactyly-hydrocephalus syndrome 3) and *NFIX* (Sotos syndrome 2). We conclude that common genetic variation in genes associated with macrocephaly syndromes, but not microcephaly syndromes, contributes to variation in head size in the general population. Reciprocal to this, genes identified through our GWAS of head size may therefore also identify currently unknown causal genes for macrocephaly. Accordingly, we observed a patient in a previously described intellectual disability cohort^23^ who presented with macrocephaly and had a mutation in *TICRR*, a gene for which a lead variant and variants in LD were eQTLs in twelve different tissues. This gene is involved in the initiation of DNA replication and interacts with *CDK2*^24^, one of the genes nearby another lead variant. Thus, *TICRR* is an interesting candidate for further study in currently undiagnosed macrocephaly syndromes.

**Figure 4.**
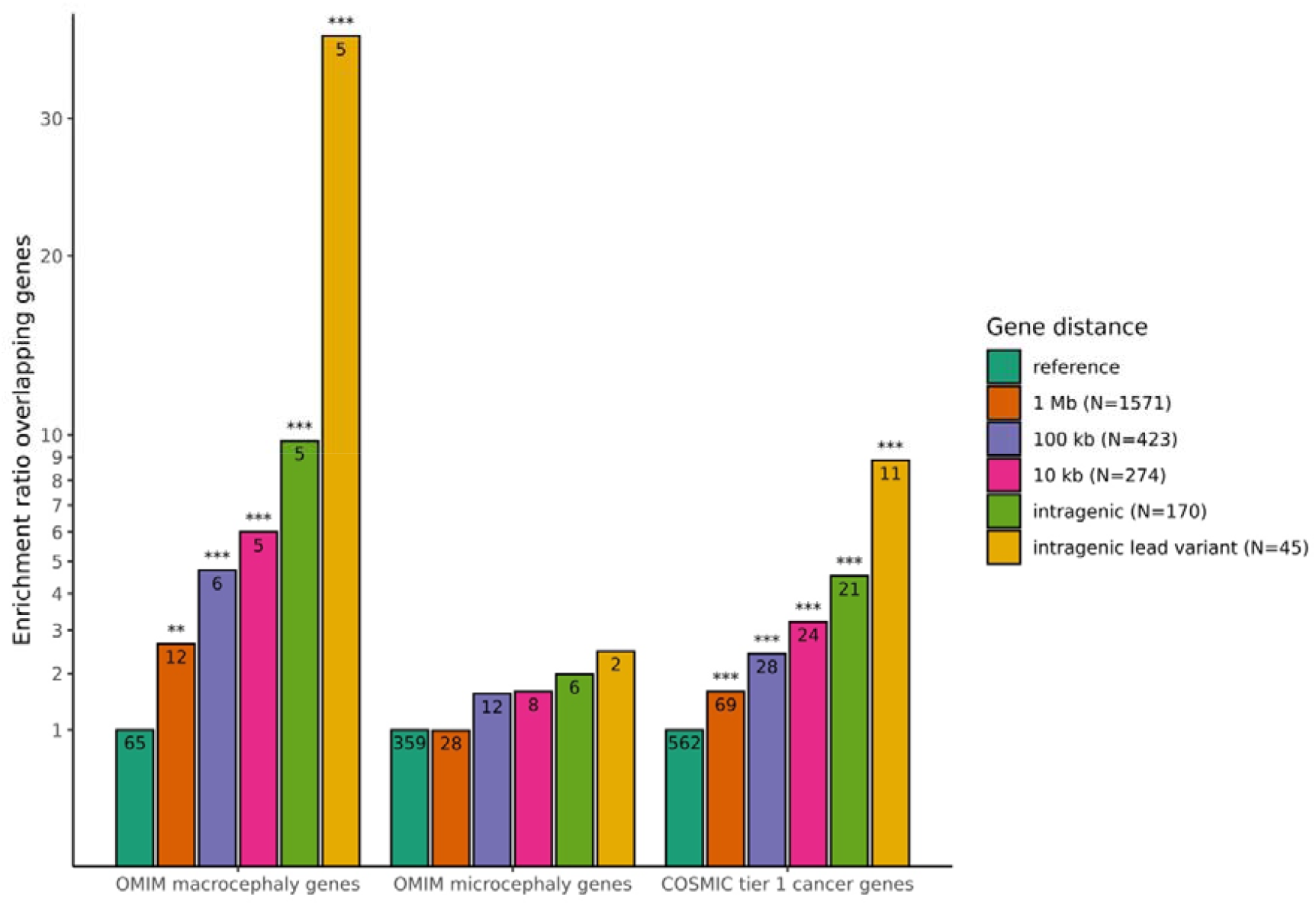
Gene enrichment stratified by distance from lead variants. **Figure 4A.** Enrichment of genes nearby the identified genetic loci for OMIM macrocephaly genes, OMIM microcephaly genes and COSMIC tier 1 genes. Depicted are enrichment of genes within 1 Mb (orange), 100 kb (purple) or 10 kb (pink) of the identified genetic loci, genes within 10 kb of the identified genetic loci with intragenic genetic variants (light green), and genes with intragenic genetic lead variants (yellow), in comparison with genes in the reference genome (dark green). Significant results are denoted by asterisks: \**P* < 0.05; \*\**P* < 0.0125 (0.05 / 4); \*\*\**P* < 0.0025 (0.05 / 4 / 5).

#### Autosomal dominance score

We did not observe a significant enrichment for microcephaly genes (**Figure 4A**). This lack of enrichment is likely due to differences between the microcephaly and macrocephaly gene sets. Notably, macrocephaly typically results from mutations with an autosomal dominant inheritance pattern (64.6%, **Table S15**), whereas microcephaly predominantly involves mutations with an autosomal recessive inheritance pattern (72.3%, **Table S16**). We observed a profound increase for genes with a predicted dominant inheritance pattern closer to our lead variants (**Figure 4B**). However, neither dominant nor recessive microcephaly genes were enriched (**Table S17**) and the predominant recessive inheritance patterns of microcephaly genes could not explain their lack of enrichment. An alternative explanation is that microcephaly syndromes are more clinically heterogeneous and the underlying mechanisms are less specific to brain and cranial growth.

**Figure 4B.**
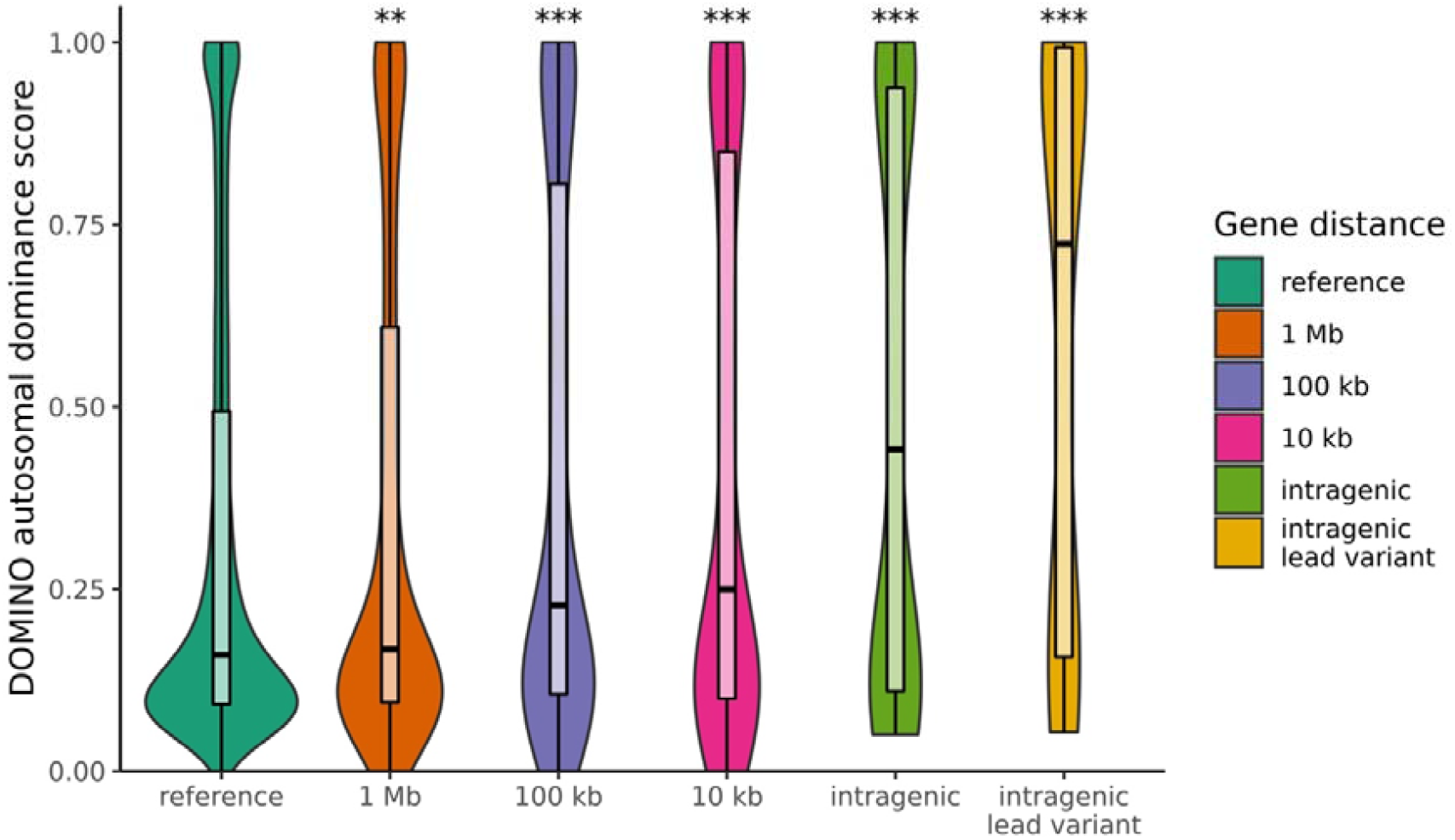
Violin plots and boxplots showing the DOMINO autosomal dominance scores of genes within 1 Mb (orange), 100 kb (purple) or 10 kb (pink) of the identified genetic loci, genes within 10 kb of the identified genetic loci with intragenic genetic variants (light green), and genes with intragenic genetic lead variants (yellow), in comparison with genes in the reference genome (dark green). Significant results are denoted by asterisks: \**P* < 0.05; \*\**P* < 0.01; \*\*\**P* < 0.001.

#### COSMIC tier 1 cancer genes

As our KEGG analysis showed a strong enrichment for cancer pathways (**Figure 3A**), we determined whether cancer genes are also enriched among genes closer to the lead variants (**Figure 4A**). Indeed, there was a 9-fold enrichment for high-fidelity cancer genes (first tier COSMIC^25^) among genes with an intragenic lead variant, which persisted after adjusting for height (**Table S17**). There was only a modest enrichment of cancer genes close to variants from the height GWAS, providing additional evidence that cancer-related genes are specifically important for head size.

#### Gain of function and loss of function

We found that macrocephaly-associated genes were more enriched for high-fidelity cancer genes than microcephaly-associated genes (enrichment ratio 12.9 vs. 3.2, **Table S17**). We therefore investigated whether the same mutation type, i.e. gain of function or loss of function, causes both macrocephaly syndromes as a germ line mutation but also associate with cancer as somatic mutations. We found that this was the case for the vast majority of macrocephaly-associated genes with a defined role in cancer (37 of 41 genes, **Table S15**), i.e. the same type of mutation associates with both macrocephaly and cancer. Moreover, germ line mutations in 14 of these 37 genes, including our GWAS genes *PTEN, PTCH1* and *SUFU,* are associated with a syndrome or condition with a suggested cancer-predisposition (**Table S15**). Our GWAS data and these observations therefore suggest that subtle upregulation of oncogenes and oncogenic pathways or down-regulation of tumor suppressor genes and pathways may increase head size in the general population.

### Implications of the head size and cancer link

The link between cancer and head size is intriguing, with some of the high-fidelity cancer genes being known macrocephaly genes (**Figure 4C**). Germline mutations in two genes are known to be related to clinical syndromes causing both abnormal head sizes and an increased cancer risk, namely the genes *PTEN* (Cowden syndrome) and *PTCH1* (Gorlin syndrome). For both syndromes, patients are routinely screened for macrocephaly as part of the diagnostic criteria, but this relationship is not yet known for other syndromes such as Li-Fraumeni syndrome *(TP53)* or familial adenomatous polyposis syndrome (*APC*), both of which are near lead variants. Our GWAS, however, was performed in the general population, prompting the interesting question whether the link between head size and cancer extends beyond rare genetic syndromes.

**Figure 4C.**
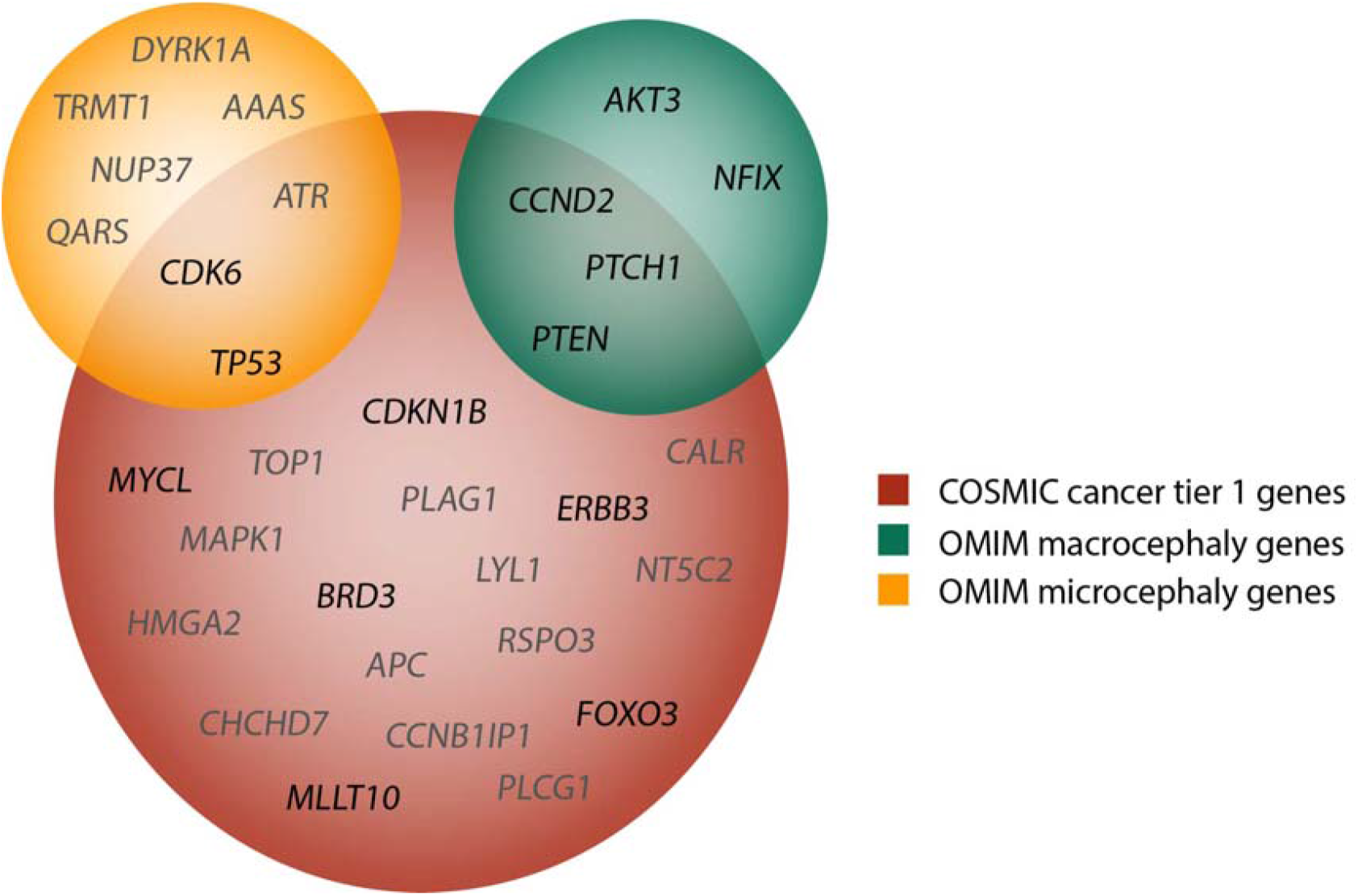
Venn diagram showing the nearby (< 10 kb) genes that overlap with OMIM microcephaly genes (depicted in yellow), OMIM macrocephaly genes (depicted in green) and COSMIC cancer tier 1 genes (depicted in red), and their in-between overlap. Genes with intragenic genetic lead variants are depicted in black, and genes without intragenic genetic lead variants in grey.

Meta-analyses of prospective observational studies found associations between height and increased risk of various forms of cancer^26^, and the few studies on body length and head circumference at birth have shown similar results^27–29^. Our results also indicate that particularly genes associated with early growth rather than later adolescent growth may be associated to neoplasia, since cranial growth is completed around the 6^th^ to 7^th^ year of age whereas height is primarily determined by peripubertal growth. In combination with our findings, the relationship between head size and cancer risk warrants further study, as well as an exploration of its clinical implications.

## Online methods

### Study population

Most studies participate in the Cohorts for Heart and Aging Research in Genomic Epidemiology (CHARGE)^30^ or the Enhancing NeuroImaging Genetics through Meta-Analysis (ENIGMA)^31^ consortium. We also included the results of the most recent head circumference GWAS^5^. A complete overview of the population characteristics is presented in **Table S1**. Each contributing study was approved by their institutional review boards or local ethical committees. Written informed consent was obtained from all study participants.

### Genotyping

Genotyping of individuals was performed on commercially available arrays, and imputed to 1000 Genomes (1KG) or Haplotype Reference Consortium (HRC) imputation panels (**Table S2**). Quality control was performed using the EasyQC software^32^. In each study, genetic variants with an imputation quality r^2^ below 0.3 and a minor allele frequency (MAF) below 0.001 were excluded. Additionally, variants were filtered on study level requiring (*r*^2^ *x MAF xN*)>5.

### Phenotyping

Different methods were used to measure human head size across studies. Briefly, either head circumference was measured, or intracranial volume was measured on computed tomography (CT) or magnetic resonance imaging (MRI) scans. In total, human head size was measured using intracranial volume measured on CT or MRI scans in respectively 1,283 and 57,186 individuals, and using head circumference in 20,524 individuals (**Table S3**). These measures have previously shown to be phenotypically and genetically correlated^4,5,33^, allowing us to perform a combined meta-analyse of different measures of head size.

### Genome-wide association studies

GWAS were performed for each study adjusted for age, age^2^ (if significant), sex, eigenstrat PC1-4 (if significant), study-specific adjustments and case-control status (if applicable). In a second model, additional adjustment for height was made. The METAL software^34^ was used to perform a sample size weighted Z-score meta-analysis. After meta-analysis, genetic variants available in less than 5,000 individuals were excluded. Comparable betas were derived using the formula 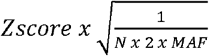 as was done previously^35^. Genomic inflation and polygenic heterogeneity were assessed using the LD score regression software^36^ by comparing the genomic control inflation factor and the LD score regression intercept (**Table S4**).

### Functional annotations

Regional association plots were made with the LocusZoom software^37^. The Functional Mapping and Annotation of Genome-Wide Association Studies (FUMA GWAS) platform^38^ was used to derive the independent genomic loci and genetic lead variants, and to functionally annotate the identified genetic variants. Additionally, enrichment for KEGG^7^ biological pathways was assessed for genes located nearby the identified genetic loci using the default options in FUMA, using hypergeometric tests. Genotype-Tissue Expression (GTEx) v7 was used to identify expression quantitative trait loci (eQTL) for the lead genetic variants and variants in LD (r^2^ > 0.6).

### Effects on anthropomorphic measures and regional brain volumes

The LD score regression software^36,39^ was used to assess genetic correlations with adult height^40^, for both the height-unadjusted and height-adjusted model.

Dual-energy X-ray absorptiometry (DXA) measurements of the UK Biobank imaging subsample (N = 3,313) were used to examine the effect of the identified lead variants on anthropometric measures across the body, i.e. bone area of the arms, legs, pelvis, ribs, spine, trunk and vertebrae L1-L4. In these analyses values more than three standard deviations from the mean were considered outliers and removed from the analyses. We adjusted for age, age^2^, sex and principal components (model 1), and additionally for height (model 2) to correct for an overall growth effect.

To investigate the effects of the identified variants for head size on growth in specific brain regions, we investigated the overlap between the identified loci for head size and previous genome-wide association studies (GWAS) on brain volumes^6,41–44^. We also analysed the associations between the identified lead genetic variants and volumes of four brain lobes, the lateral ventricles, eight subcortical structures and 34 cortical regions of interest in the UK Biobank (N = 22,145). Volumes were derived using the FreeSurfer 6.0 software. Values more than 3.5 standard deviations away from the mean were considered outliers and removed from the analysis. In the first model, we adjusted for age, age^2^, sex and principal components, and in the second model additionally for intracranial volume.

Additionally, we took the lead variants specifically associated with one or two subcortical volumes, and investigated their effects on the shape of seven subcortical structures, i.e. amygdala, caudate nucleus, hippocampus, nucleus accumbens, pallidum, putamen and thalamus. The radial distances and log Jacobian determinants were derived using the ENIGMA-Shape package (http://enigma.usc.edu/ongoing/enigma-shape-analysis/). Volumetric outliers more than 3.5 standard deviations from the mean were removed from the analysis.

We performed 10,000 permutations to define the number of independent DXA, brain volumetric and subcortical shape outcomes. We used this number to define our multiple testing adjusted p-value thresholds for significance, i.e. 0.05 / (number of independent outcomes x number of lead genetic variants).

### Enrichment analyses

We performed enrichment analyses of different gene sets: genes within 1 Mb, 100 kb or 10 kb of the identified genetic loci, genes within 10 kb of the identified genetic loci with intragenic genetic variants, and genes within 10 kb of the identified genetic loci with intragenic genetic lead variants. As a reference, we used the rest of the protein-coding genome.

First, the Online Mendelian Inheritance in Man (OMIM) database^45^ was used to retrieve information on genes related to heritable phenotypes affecting head size. Second, the Catalogue of Somatic Mutations in Cancer (COSMIC) database^25^ was used to extract Tier 1 cancer genes. Taking the rest of the genome as our reference gene set, we calculated the enrichment of these macrocephaly, microcephaly and cancer genes in the abovementioned gene sets.

Lastly, DOMINO^46^, a previously developed machine learning tool, was used to assess if the genes in the different gene sets were more often predicted to harbour dominant changes in comparison with genes in the rest of the genome.

Mean autosomal dominance scores were compared with the reference genome using a Mann-Whitney test. Differences in the proportions for the OMIM macro- and microcephaly genes, intellectual disability genes and COSMIC genes were calculated using a Pearson’s χ^2^ test.

We performed these analyses for the head size height-unadjusted GWAS results, but also the GWAS in the subset of studies for which height was available, the height-adjusted GWAS and the height GWAS^40^. For comparison, we also selected the top 67 loci for the height GWAS, so the results were not driven by a difference in the number of associated loci.

## Supporting information

Supplementary Figure S1. Forest plots

Supplementary Figure S2. Locuszoom plots

## Supplementary Materials

### Supplementary Tables

See separate Excel file.

**Table S1.** List of all contributing studies.

**Table S2.** Population characteristics of new or updated contributing studies.

**Table S3.** Information on genotyping and quality control.

**Table S4.** Phenotyping information.

**Table S5.** Lambda genomic control, LD score regression intercept and ratio for different models.

**Table S6.** Lead genetic variants and their effects on human head size in samples of different ethnicities, with and without adjustment for height.

**Table S7.** Genome-wide significant genetic variants and variants in linkage disequilibrium (r^2^>0.6), including functional annotations.

**Table S8.** Effects of lead genetic variants on bone size area measured using dual-energy X-ray absorptiometry (DXA).

**Table S9.** Overlap between identified loci and previously identified loci in genome-wide association studies of brain volumes.

**Table S10.** The effects of previously identified genetic variants for regional brain volumes in the current genome-wide association study.

**Table S11.** Association of identified lead genetic variants with regional brain volumes.

**Table S12.** Results of the subcortical shape analyses of seven lead genetic variants specifically associated with one or two subcortical structures.

**Table S13.** Kyoto Encyclopaedia of Genes and Genomes (KEGG) pathway analysis.

**Table S14.** Genes in or nearby identified genetic loci (< 1 Mb).

**Table S15.** List of genes linked to macrocephaly in the Online Mendelian Inheritance of Man (OMIM) database.

**Table S16.** List of genes linked to microcephaly in the Online Mendelian Inheritance of Man (OMIM) database.

**Table S17.** Enrichment of micro- and macrocephaly OMIM genes, COSMIC tier 1 cancer genes, intellectual disability trios and autosomal dominance DOMINO score.

### Supplementary Figures

**Figure S1.** Forest plots presenting the study-specific associations of the identified lead genetic variants with human head size.

**Figure S2. Regional plots of the identified genetic loci for human head size (±100 kb).Acknowledgements**

## Acknowledgements

are provided for the studies who contributed new samples in addition to samples in previous efforts^4,5^.

## The 1000BRAINS study

The 1000BRAINS study was funded by the Institute of Neuroscience and Medicine, Research Center Juelich, Germany. We thank the Heinz Nixdorf Foundation (Germany) for the generous support of the Heinz Nixdorf Recall Study on which 1000BRAINS is based. We also thank the scientists and the study staff of the Heinz Nixdorf Recall Study and 1000BRAINS. Funding was also granted by the Initiative and Networking Fund of the Helmholtz Association (Svenja Caspers) and the European Union’s Horizon 2020 Research and Innovation Program under Grant Agreement 785907 (Human Brain Project SGA2; Katrin Amunts, Svenja Caspers, Sven Cichon) and 945539 (Human Brain Project SGA3; Katrin Amunts, Svenja Caspers, Sven Cichon).

## The Three-City Study (Bordeaux and Dijon)

The Three-City Study is conducted under a partnership agreement between the Institut National de la Santé et de la Recherche Médicale (INSERM), the Institut de Santé Publique et Développement of the Victor Segalen Bordeaux 2 University and Sanofi-Aventis. The Fondation pour la Recherche Médicale funded the preparation and initiation of the study. The 3C Study is also supported by the Caisse Nationale Maladie des Travailleurs Salariés, Direction Générale de la Santé, Mutuelle Générale de l’Education Nationale, Institut de la Longévité, Regional Governments of Aquitaine and Bourgogne, Fondation de France, Ministry of Research-INSERM Programme “Cohortes et collections de données biologiques”, French National Research Agency COGINUT ANR-06-PNRA-005, the Fondation Plan Alzheimer (FCS 2009–2012), and the Caisse Nationale pour la Solidarité et l’Autonomie (CNSA). This project has received funding from the European Union’s Horizon 2020 research and innovation programme under grant agreement No 643417, No 640643, the French National Research Agency (ANR), and the University of Bordeaux Initiative of Excellence (IdEX). Part of the computations were performed at the Bordeaux Bioinformatics Center (CBiB), University of Bordeaux and at the CREDIM (Centre de Ressource et Développement en Informatique Médicale) at University of Bordeaux, on a server infrastructure supported by the Fondation Claude Pompidou. The project is supported through the following funding organisations under the aegis of JPND-www.jpnd.eu (BRIDGET project): Australia, National Health and Medical Research Council, Austria, Federal Ministry of Science, Research and Economy; Canada, Canadian Institutes of Health Research; France, French National Research Agency; Germany, Federal Ministry of Education and Research; Netherlands, The Netherlands Organisation for Health Research and Development; United Kingdom, Medical Research Council.

## The Atherosclerosis Risk in Communities (ARIC) study

The Atherosclerosis Risk in Communities study has been funded in whole or in part with Federal funds from the National Heart, Lung, and Blood Institute, National Institutes of Health, Department of Health and Human Services (contract numbers HHSN268201700001I, HHSN268201700002I, HHSN268201700003I, HHSN268201700004I and HHSN268201700005I), R01HL087641, R01HL086694; National Human Genome Research Institute contract U01HG004402; and National Institutes of Health contract HHSN268200625226C. The authors thank the staff and participants of the ARIC study for their important contributions. Infrastructure was partly supported by Grant Number UL1RR025005, a component of the National Institutes of Health and NIH Roadmap for Medical Research. This project was supported in part by National Institute of Neurological Disorders and Stroke grant NS087541.

## The Australian Schizophrenia Research Bank (ASRB)

Data and samples were collected by the Australian Schizophrenia Research Bank (ASRB), supported by the Australian NHMRC, the Pratt Foundation, Ramsay Health Care, and the Viertel Charitable Foundation. The ASRB were also supported by the Schizophrenia Research Institute (Australia), utilizing infrastructure funding from NSW Health and the Macquarie Group Foundation. DNA analysis was supported by the Neurobehavioral Genetics Unit, utilising funding from NSW Health. MC was supported by an NHMRC Senior Research Fellowship (1121474) and MC and MG were supported by NHMRC project grants (1147644 and 1051672).

## The Coronary Artery Risk Development in Young Adults (CARDIA) study

The Coronary Artery Risk Development in Young Adults Study (CARDIA) is conducted and supported by the National Heart, Lung, and Blood Institute (NHLBI) in collaboration with the University of Alabama at Birmingham (HHSN268201800005I & HHSN268201800007I), Northwestern University (HHSN268201800003I), University of Minnesota (HHSN268201800006I), and Kaiser Foundation Research Institute (HHSN268201800004I). CARDIA was also partially supported by the Intramural Research Program of the National Institute on Aging (NIA) and an intra-agency agreement between NIA and NHLBI (AG0005). Genotyping was funded as part of the NHLBI Candidate-gene Association Resource (N01-HC-65226) and the NHGRI Gene Environment Association Studies (GENEVA) (U01-HG004729, U01-HG04424, and U01-HG004446). This manuscript has been reviewed and approved by CARDIA for scientific content.

## CROMIS-2 ICH

The CROMIS-2 study is funded by the Stroke Association and British Heart Foundation. Funding for genotyping was provided by the UCLH/UCL National Institute for Health Research (NIHR) Biomedical Research Centre.

## Diabetes Heart Study (DHS)

This study was supported in part by the National Institutes of Health through R01 HL67348, R01 HL092301, R01 NS058700, R01 NS075107, R01 AG058921 and the General Clinical Research Center at Wake Forest School of Medicine (M01 RR07122, F32 HL085989). The authors thank the investigators, staff, and participants of the DHS for their valuable contributions.

## The Duke Neurogenetics Study (DNS)

The Duke Neurogenetics Study was supported by Duke University as well as US-National Institutes of Health grants R01DA033369 and R01DA031579. RA, ARK, and ARH received further support from US-National Institutes of Health grant R01AG049789.

## Epidemiological Prevention Study Zoetermeer (EPOZ)

We are grateful to all the study participants. We would like to thank Dr. Ir. Natalie Terzikhan for imputing the genetic data.

## Erasmus Stroke Study (ESS)

The ESS was supported by the Stroke Research Foundation and Erasmus MC MRACE grants.

## The Function Biomedical Informatics Research Network (FBIRN)

This work was supported by the National Center for Research Resources at the National Institutes of Health [grant numbers: NIH 1 U24 RR021992 (Function Biomedical Informatics Research Network), NIH 1 U24 RR025736-01 (Biomedical Informatics Research Network Coordinating Center], the National Center for Research Resources and the National Center for Advancing Translational Sciences, National Institutes of Health, through Grant UL1 TR000153, and the National Institutes of Health through 5R01MH094524 and P20GM103472.

## Framingham Heart Study (FHS)

This work was supported by the National Heart, Lung and Blood Institute’s Framingham Heart Study (Contracts N01-HC-25195, HHSN268201500001I, and 75N92019D00031) and its contract with Affymetrix, Inc. for genotyping services (Contract No. N02-HL-6-4278). A portion of this research utilized the Linux Cluster for Genetic Analysis (LinGA-II) funded by the Robert Dawson Evans Endowment of the Department of Medicine at Boston University School of Medicine and Boston Medical Center. This study was also supported by grants from the National Institute of Aging (R01s AG033040, AG033193, AG054076, AG049607, AG008122, AG016495, U01-AG049505, AG052409, AG058589 and RF1AG059421) and the National Institute of Neurological Disorders and Stroke (R01-NS017950). We would like to thank the dedication of the Framingham Study participants, as well as the Framingham Study team, especially investigators and staff from the Neurology group, for their contributions to data collection. Dr. DeCarli is supported by the Alzheimer’s Disease Center (P30 AG 010129). The views expressed in this manuscript are those of the authors and do not necessarily represent the views of the National Heart, Lung, and Blood Institute; the National Institutes of Health; or the U.S. Department of Health and Human Services.

## Generation R

The work was supported by ZonMw (TOP project 91211021 [TW]), the Sophia Foundation (grant S18-20 [RLM]) and the European Union’s Horizon 2020 research and innovation programme (No 733206 LifeCycle Project [SL], No 848158 EarlyCause Project [CAMC] and Marie Skłodowska-Curie grant agreement No 707404 [CAMC]). The Generation R Study is conducted by the Erasmus Medical Center in close collaboration with the School of Law and Faculty of Social Sciences of the Erasmus University Rotterdam, the Municipal Health Service Rotterdam area, Rotterdam, the Rotterdam Homecare Foundation, Rotterdam and the Stichting Trombosedienst & Artsenlaboratorium Rijnmond (STAR-MDC), Rotterdam. We gratefully acknowledge the contribution of children and parents, general practitioners, hospitals, midwives and pharmacies in Rotterdam. The general design of Generation R Study is made possible by financial support from the Erasmus Medical Center, Rotterdam, the Erasmus University Rotterdam, the Netherlands Organization for Health Research and Development (ZonMw), the Netherlands Organisation for Scientific Research (NWO), the Ministry of Health, Welfare and Sport and the Ministry of Youth and Families.

## The Trøndelag Health Study (HUNT)

The HUNT Study is a collaboration between HUNT Research Centre (Faculty of Medicine and Health Sciences, NTNU – Norwegian University of Science and Technology), Nord-Trøndelag County Council, Central Norway Health Authority, and the Norwegian Institute of Public Health. HUNT MRI was funded by the Liaison Committee between the Central Norway Regional Health Authority and the Norwegian University of Science and Technology, and the Norwegian National Advisory Unit for functional MRI.

## The Institute of Mental Health (IMH) study

This work was supported by research grants from the National Healthcare Group, Singapore (SIG/05004; SIG/05028), and the Singapore Bioimaging Consortium (RP C-009/2006) research grants awarded to KS.

## LIFE-Adult

LIFE-Adult is supported by LIFE – Leipzig Research Centre for Civilization Diseases, an organisational unit affiliated to the Medical Faculty of the University of Leipzig. LIFE is funded by means of the European Union, European Regional Development Fund (ERDF) and by funds of the Free State of Saxony within the framework of the excellence initiative (project numbers 713-241202, 713-241202, 14505/2470, 14575/2470). We thank all participants and Kerstin Wirkner, Ulrike Scharrer, Katrin Arelin, Frauke Beyer and everyone involved in MRI data acquisition and analysis.

## The Poznan MS study

Dr Pawlak reported receiving grants from Polish National Science Centre 2011/01/D/NZ4/05801.

## The Rotterdam Study (RS)

The Rotterdam Study is funded by Erasmus Medical Center and Erasmus University, Rotterdam, Netherlands Organization for the Health Research and Development (ZonMw), the Research Institute for Diseases in the Elderly (RIDE), the Ministry of Education, Culture and Science, the Ministry for Health, Welfare and Sports, the European Commission (DG XII), and the Municipality of Rotterdam. The authors are grateful to the study participants, the staff from the Rotterdam Study and the participating general practitioners and pharmacists. The generation and management of GWAS genotype data for the Rotterdam Study (RS I, RS II, RS III) were executed by the Human Genotyping Facility of the Genetic Laboratory of the Department of Internal Medicine, Erasmus MC, Rotterdam, The Netherlands. The GWAS datasets are supported by the Netherlands Organisation of Scientific Research NWO Investments (nr. 175.010.2005.011, 911-03-012), the Genetic Laboratory of the Department of Internal Medicine, Erasmus MC, the Research Institute for Diseases in the Elderly (014-93-015; RIDE2), the Netherlands Genomics Initiative (NGI)/Netherlands Organisation for Scientific Research (NWO) Netherlands Consortium for Healthy Aging (NCHA), project nr. 050-060-810. We thank Pascal Arp, Mila Jhamai, Marijn Verkerk, Lizbeth Herrera and Marjolein Peters, and Carolina Medina-Gomez, for their help in creating the GWAS database, and Karol Estrada, Yurii Aulchenko, and Carolina Medina-Gomez, for the creation and analysis of imputed data. HHHA is supported by ZonMW grant number 916.19.151.

## Study of Health in Pomerania - TREND (SHIP-TREND)

SHIP is part of the Community Medicine Research net of the University of Greifswald, Germany, which is funded by the Federal Ministry of Education and Research (grants no. 01ZZ9603, 01ZZ0103, and 01ZZ0403), the Ministry of Cultural Affairs and the Social Ministry of the Federal State of Mecklenburg-West Pomerania. MRI scans and genome-wide SNP typing in SHIP-TREND have been supported by a joint grant from Siemens Healthineers, Erlangen, Germany and the Federal State of Mecklenburg-West Pomerania. The University of Greifswald is a member of the Caché Campus program of the InterSystems GmbH.

## The Saguenay Youth Study (SYS)

The Canadian Institutes of Health Research and the Heart and Stroke Foundation of Canada fund the SYS. Computations were performed on the GPC supercomputer at the SciNet HPC Consortium. SciNet is funded by: the Canada Foundation for Innovation under the auspices of Compute Canada; the Government of Ontario; Ontario Research Fund - Research Excellence; and the University of Toronto.

## UK Biobank

This research has been conducted using the UK Biobank Resource under Application Number 23509.

## Vitamin D Intervention in Infants (VIDI) study

The VIDI study is supported by The Finnish Medical Foundation, the Academy of Finland, the Sigrid Jusélius Foundation, the Swedish Research Council, the Novo Nordisk Foundation, Finska Läkaresällskapet and Folkhälsan Research Foundation. We want to thank Dr. Helena Hauta-alus, Dr. Elisa Holmlund-Suila, Dr. Saara Valkama and Dr. Jenni Rosendahl for their contribution in acquiring the data.

## The Cohorts for Heart and Aging Research in Genomic Epidemiology (CHARGE) Consortium

Philippe Amouyel^114^, Konstantinos Arfanakis^115,116^, Benjamin S. Aribisala^117,118,119^, Mark E. Bastin^120,121^, Ganesh Chauhan^8,122^, Christopher Chen^123^, Ching-Yu Cheng^124,125^, Philip L. de Jager^126^, Ian J. Deary^127^, Debra A. Fleischman^116,128,129^, Rebecca F. Gottesman^130,131^, Vilmundur Gudnason^132,133^, Saima Hilal^134,135,136^, Edith Hofer^93,137^, Deborah Janowitz^9^, J. Wouter Jukema^138,139,140,141^, David C.M. Liewald^142^, Lorna M. Lopez^143^, Oscar Lopez^144^, Michelle Luciano^142^, Oliver Martinez^145^, Wiro J. Niessen^4,146^, Paul Nyquist^130,147^, Jerome I. Rotter^148^, Tatjana Rundek^149^, Ralph L. Sacco^149^, Helena Schmidt^150^, Henning Tiemeier^53,151^, Stella Trompet^152^, Jeroen van der Grond^153^, Henry Völzke^97^, Joanna M. Wardlaw^120,121,154^, Lisa Yanek^147^, Jingyun Yang^116,128^.

## The Enhancing NeuroImaging Genetics through Meta-Analysis (ENIGMA) Consortium

Ingrid Agartz^155,156,157^, Saud Alhusaini^158^, Laura Almasy^159,160^, David Ames^161,162^, Katrin Amunts^48,49,89^, Ole A. Andreassen^163,164^, Nicola Armstrong^165^, Manon Bernard^166^, John Blangero^167,168^, Laura M.E. Blanken^53^, Marco P. Boks^169^, Dorret I. Boomsma^170^, Adam M. Brickman^171^, Henry Brodaty^172,173^, Randy L. Buckner^174,175,176^, Jan K. Buitelaar^177^, Dara M. Cannon^178^, Vaughan J. Carr^64,65,179^, Stanley V. Catts^180,181^, M. Mallar Chakravarty^182,183^, Qiang Chen^184^, Christopher R.K. Ching^74^, Aiden Corvin^185^, Benedicto Crespo-Facorro^186,187^, Joanne E. Curran^167,168^, Gareth E. Davies^188^, Eco J.C. de Geus^189,190^, Greig I. de Zubicaray^191^, Anouk den Braber^170^, Sylvane Desrivières^192^, Allissa Dillman^193^, Srdjan Djurovic^194,195^, Wayne C. Drevets^196^, Ravi Duggirala^167,168^, Stefan Ehrlich^197^, Susanne Erk^198^, Thomas Espeseth^21,199^, Iryna O. Fedko^170^, Guillen Fernández^61^, Simon E. Fisher^16,61^, Tatiana M. Foroud^200^, Tian Ge^201^, Sudheer Giddaluru^195,202^, David C. Glahn^203^, Aaron L. Goldman^184^, Robert C. Green^204,205,206^, Corina U. Greven^177,207,208^, Oliver Grimm^209^, Narelle K. Hansell^210^, Catharina A. Hartman^211^, Ryota Hashimoto^212^, Andreas Heinz^198^, Frans Henskens^213,214^, Derrek P. Hibar^215^, Beng-Choon Ho^216^, Pieter J. Hoekstra^217^, Avram J. Holmes^176,218,219^, Martine Hoogman^61^, Jouke-Jan Hottenga^170^, Hilleke E. Hulshoff Pol^220^, Assen Jablensky^221^, Mark Jenkinson^222,223,224^, Tianye Jia^225,226,227^, Karl-Heinz Jöckel^228^, Erik G. Jönsson^157,229^, Sungeun Kim^230,231^, Marieke Klein^98,232^, Peter Kochunov^233^, John B. Kwok^234,235^, Stephen M. Lawrie^236^, Stephanie Le Hellard^195,202^, Hervé Lemaître^38^, Carmel Loughland^237,238^, Andre F. Marquand^61^, Nicholas G. Martin^239^, Jean-Luc Martinot^240^, Mar Matarin^241^, Daniel H. Mathalon^242,243^, Karen A. Mather^65,172^, Venkata S. Mattay^130,184,244^, Colm McDonald^178^, Francis J. McMahon^245^, Katie L. McMahon^246^, Rebekah E, McWhirter^247,248^, Patrizia Mecocci^249^, Ingrid Melle^155,250^, Andreas Meyer-Lindenberg^251^, Patricia T. Michie^252^, Yuri Milaneschi^253^, Derek W. Morris^178^, Bryan Mowry^210,254^, Kwangsik Nho^231^, Thomas E. Nichols^255^, Markus N. Nöthen^70^, Rene L. Olvera^256^, Jaap Oosterlaan^257,258^, Roel A. Ophoff^53,259^, Massimo Pandolfo^260,261^, Christos Pantelis^262,263,264^, Irene Pappa^53^, Brenda Penninx^265^, G. Bruce Pike^266^, Paul E. Rasser^238,267,268^, Miguel E. Renteria^269^, Simone Reppermund^172,270^, Marcella Rietschel^271^, Shannon L. Risacher^272^, Nina Romanczuk-Seiferth^198^, Emma Jane Rose^273^, Perminder S. Sachdev^172,274^, Philipp G. Sämann^275^, Andrew J. Saykin^200,272^, Ulrich Schall^238,268^, Peter R. Schofield^65,235^, Sara Schramm^228^, Gunter Schumann^198,276,277^, Rodney Scott^238,268,278^, Li Shen^279^, Sanjay M. Sisodiya^280,281^, Hilkka Soininen^282^, Emma Sprooten^177^, Velandai Srikanth^283^, Vidar M. Steen^195^, Lachlan T. Strike^210^, Anbupalam Thalamuthu^172^, Arthur W. Toga^284^, Paul Tooney^32,267,268^, Diana Tordesillas-Gutiérrez^285,286^, Jessica A. Turner^287^, Maria del C. Valdés Hernández^119,120,142^, Dennis van der Meer^288,289^, Nic J.A. Van der Wee^290,291^, Neeltje E.M. Van Haren^53,169^, Dennis van ‘t Ent^170^, Dick J. Veltman^265,292^, Henrik Walter^198^, Daniel R. Weinberger^184,293^, Michael W. Weiner^59^, Wei Wen^172,274^, Lars T. Westlye^21,163,294^, Eric Westman^295^, Anderson M. Winkler^219,296^, Girma Woldehawariat^297^, Margaret J. Wright^210,298^, Jingqin Wu^32^.

