## Supplementary Figure S1. Forest plots for "Genetic variants for head size share genes and pathways with cancer"

**rs6682560 (A); chr1:23357282; model 1 (I2=24.8, HetP=0.06816)**

| Study | N_total | EAF | Beta | P |
|---|---|---|---|---|
| **Head circumference** | | | | |
| Haworth | 18881 | 0.8326 | 0.0247 | 7.303e-02 |
| VIDI | 754 | 0.8127 | 0.0739 | 2.631e-01 |
| **CT-based** | | | | |
| ESS | 641 | 0.8367 | 0.1262 | 9.463e-02 |
| UCL | 642 | 0.8268 | 0.122 | 9.805e-02 |
| **MR-based** | | | | |
| 1000Brains | 751 | 0.8541 | 0.0659 | 3.672e-01 |
| 3C-Bordeaux | 564 | 0.8081 | 0.0736 | 3.303e-01 |
| 3c-Dijon | 1499 | 0.8174 | 1e-04 | 9.976e-01 |
| AGES | 2226 | 0.8385 | -0.012 | 7.678e-01 |
| ARIC.EA | 1390 | 0.825 | 0.1138 | 2.260e-02 |
| ASPS-Family | 338 | 0.8271 | 0.0151 | 8.820e-01 |
| ASRB | 233 | 0.8102 | -0.1623 | 1.709e-01 |
| CARDIA.EA | 349 | 0.8189 | -0.0552 | 5.739e-01 |
| CHAP | 261 | 0.8314 | 0.0534 | 6.483e-01 |
| CHS | 544 | 0.8418 | 0.1526 | 6.617e-02 |
| DHS | 460 | 0.823 | -0.0389 | 6.529e-01 |
| ENIGMA | 11373 | 0.8304 | 0.0352 | 4.619e-02 |
| EPOZ | 251 | 0.8364 | -0.2875 | 1.718e-02 |
| FHS | 4710 | 0.8176 | 0.0392 | 1.418e-01 |
| GenR | 265 | 0.8349 | 0.1578 | 1.775e-01 |
| HUNT | 851 | 0.8328 | -0.0165 | 7.998e-01 |
| LBC1936 | 577 | 0.8184 | 0.0355 | 6.422e-01 |
| LIFE | 1825 | 0.8377 | -0.0575 | 2.003e-01 |
| LLS | 355 | 0.8208 | 0.1467 | 1.339e-01 |
| MethCT. | 114 | 0.8812 | -0.0872 | 6.704e-01 |
| NOMAS | 141 | 0.8535 | -0.1083 | 5.212e-01 |
| PoznanMS | 204 | 0.8404 | 0.1556 | 2.499e-01 |
| PROSPER | 315 | 0.8539 | 0.0725 | 5.199e-01 |
| RS1 | 1103 | 0.8301 | 0.0846 | 1.356e-01 |
| RS2 | 1159 | 0.8357 | 0.0362 | 5.189e-01 |
| RS3 | 2582 | 0.8291 | 0.0609 | 9.985e-02 |
| RUSH1 | 184 | 0.8316 | 0.202 | 1.489e-01 |
| RUSH2 | 106 | 0.8584 | 0.6576 | 1.191e-03 |
| SHIP | 966 | 0.8474 | -0.0013 | 9.837e-01 |
| SHIP-TREND | 858 | 0.8473 | 0.0628 | 3.500e-01 |
| SYS.adolescents | 980 | 0.8286 | -0.0174 | 7.716e-01 |
| SYS.adults | 594 | 0.824 | -0.0453 | 5.527e-01 |
| TREND | 1084 | 0.8228 | 0.0848 | 1.319e-01 |
| UKB | 14571 | 0.8222 | 0.0634 | 3.553e-05 |
| WHICAP | 83 | 0.8333 | -0.2341 | 2.645e-01 |
| **Non-European** | | | | |
| ARIC.AA | 675 | 0.8715 | 0.0493 | 5.446e-01 |
| CARDIA.AA | 132 | 0.889 | -0.1848 | 3.457e-01 |
| CHAP.black | 321 | 0.8746 | -0.2105 | 7.837e-02 |
| FBIRN | 275 | 0.8891 | 0.0607 | 6.552e-01 |
| NOMAS.black | 168 | 0.8602 | 0.0727 | 6.444e-01 |
| NOMAS.hispanic | 718 | 0.8483 | -0.0395 | 5.912e-01 |
| WHICAP.black | 60 | 0.9143 | 0.4057 | 2.189e-01 |
| **Summary European MA** | **74670** | **0.8288** | **0.039** | **1.851e-08** |
| **Summary African MA** | **1356** | **0.8744** | **-0.017** | **0.7626** |
| **Summary Transancestral MA** | **77133** | **0.8301** | **0.037** | **4.944e-08** |

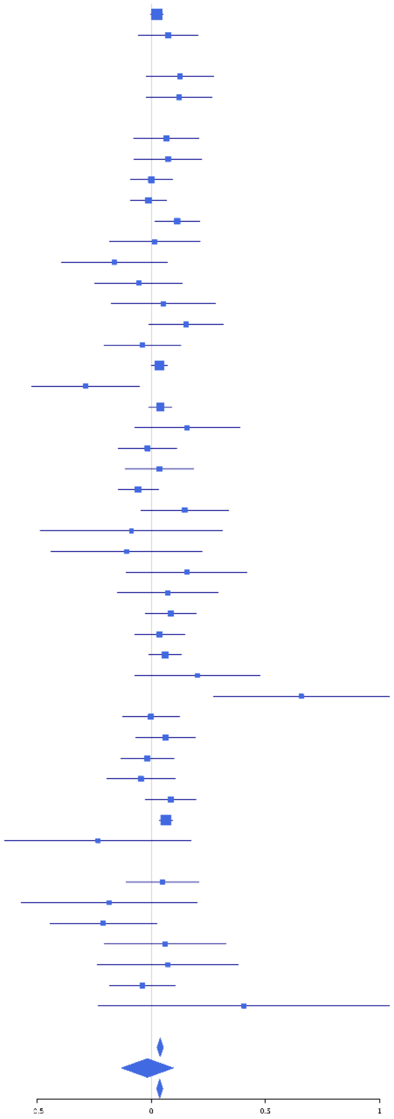

**rs3134614 (C); chr1:40363054; model 1 (I2=5.3, HetP=0.3714)**

| Study | N_total | EAF | Beta | P |
|---|---|---|---|---|
| **Head circumference** | | | | |
| Haworth | 18881 | 0.8792 | 0.0508 | 1.251e-03 |
| VIDI | 728 | 0.8902 | 0.114 | 1.741e-01 |
| **CT-based** | | | | |
| ESS | 641 | 0.8821 | 0.0716 | 4.083e-01 |
| UCL | 642 | 0.8777 | -0.0195 | 8.189e-01 |
| **MR-based** | | | | |
| 1000Brains | 751 | 0.8793 | 0.019 | 8.106e-01 |
| 3C-Bordeaux | 564 | 0.8766 | -0.0391 | 6.661e-01 |
| 3c-Dijon | 1499 | 0.8793 | 0.142 | 1.132e-02 |
| AGES | 2226 | 0.88 | 0.0702 | 1.280e-01 |
| ARIC.EA | 1390 | 0.8666 | -0.0028 | 9.603e-01 |
| ASPS-Family | 338 | 0.8683 | 0.1501 | 1.867e-01 |
| ASRB | 227 | 0.8154 | -0.0105 | 9.308e-01 |
| CARDIA.EA | 349 | 0.8833 | 0.0646 | 5.835e-01 |
| CHAP | 261 | 0.8541 | 0.0281 | 8.203e-01 |
| CHS | 544 | 0.8865 | 0.2021 | 3.456e-02 |
| DHS | 460 | 0.8945 | 0.2292 | 3.270e-02 |
| ENIGMA | 11373 | 0.8745 | 0.0076 | 7.056e-01 |
| EPOZ | 251 | 0.8772 | -0.1564 | 2.502e-01 |
| ERF | 122 | 0.9155 | -0.5388 | 1.926e-02 |
| FHS | 4710 | 0.8694 | 0.0221 | 4.696e-01 |
| GenR | 265 | 0.8855 | -0.0488 | 7.205e-01 |
| HUNT | 851 | 0.8584 | 0.055 | 4.290e-01 |
| LBC1936 | 577 | 0.8802 | 0.0146 | 8.722e-01 |
| LIFE | 1825 | 0.8703 | 0.0289 | 5.577e-01 |
| LLS | 355 | 0.8795 | 0.0191 | 8.679e-01 |
| NOMAS | 138 | 0.8572 | 0.3769 | 3.014e-02 |
| PoznanMS | 204 | 0.8907 | 0.2925 | 6.536e-02 |
| PROSPER | 315 | 0.8908 | 0.0461 | 7.184e-01 |
| RS1 | 1103 | 0.8683 | 0.1139 | 7.040e-02 |
| RS2 | 1159 | 0.8641 | 0.0732 | 2.272e-01 |
| RS3 | 2582 | 0.8734 | 0.1484 | 3.905e-04 |
| RUSH1 | 184 | 0.8641 | -0.1174 | 4.411e-01 |
| RUSH2 | 106 | 0.9123 | -0.0524 | 8.294e-01 |
| SHIP | 966 | 0.8908 | 1e-04 | 9.982e-01 |
| SHIP-TREND | 858 | 0.8851 | 0.045 | 5.517e-01 |
| SYS.adolescents | 980 | 0.9201 | 0.0496 | 5.513e-01 |
| SYS.adults | 594 | 0.9288 | 0.0174 | 8.780e-01 |
| TREND | 1084 | 0.8714 | 0.0982 | 1.263e-01 |
| UKB | 14571 | 0.874 | 0.0556 | 1.630e-03 |
| WHICAP | 83 | 0.8339 | -0.1072 | 6.089e-01 |
| **Non-European** | | | | |
| ARIC.AA | 675 | 0.9775 | -0.0648 | 7.240e-01 |
| CHAP.black | 321 | 0.9752 | 0.1716 | 4.998e-01 |
| FBIRN | 275 | 0.901 | 0.0294 | 8.371e-01 |
| NOMAS.black | 164 | 0.9554 | -0.1618 | 5.461e-01 |
| NOMAS.hispanic | 701 | 0.9118 | 0.1196 | 2.044e-01 |
| SIMES | 201 | 0.9922 | -0.4513 | 3.041e-01 |
| WHICAP.black | 60 | 0.0417 | 0.1744 | 7.042e-01 |
| **Summary European MA** | **74757** | **0.8767** | **0.048** | **8.123e-10** |
| **Summary African MA** | **1220** | **0.9279** | **-0.004** | **0.9572** |
| **Summary Asian MA** | **201** | **0.9922** | **-0.583** | **0.3041** |
| **Summary Transancestral MA** | **77154** | **0.8782** | **0.048** | **9.335e-10** |

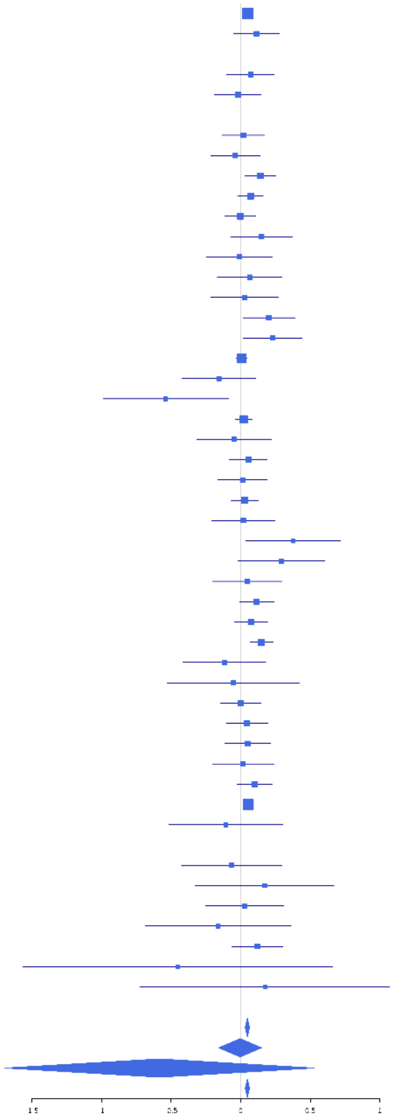

**rs12116800 (A); chr1:51027554; model 1 (I2=17.3, HetP=0.1566)**

| Study | N_total | EAF | Beta | P |
|---|---|---|---|---|
| **Head circumference** | | | | |
| Haworth | 18881 | 0.0514 | -0.0195 | 4.029e-01 |
| **CT-based** | | | | |
| ESS | 641 | 0.0381 | -0.157 | 2.817e-01 |
| UCL | 642 | 0.0486 | 0.0385 | 7.661e-01 |
| **MR-based** | | | | |
| 1000Brains | 751 | 0.0807 | 0.158 | 9.570e-02 |
| 3C-Bordeaux | 564 | 0.0532 | -0.2587 | 5.160e-02 |
| 3c-Dijon | 1499 | 0.0662 | -0.0353 | 6.305e-01 |
| AGES | 2226 | 0.0671 | -0.1578 | 8.434e-03 |
| ARIC.EA | 1390 | 0.04 | -0.089 | 3.581e-01 |
| ASPS-Family | 338 | 0.0705 | -0.3581 | 1.712e-02 |
| CARDIA.EA | 349 | 0.0691 | -0.0248 | 8.684e-01 |
| CHAP | 261 | 0.0744 | -0.2062 | 2.175e-01 |
| CHS | 544 | 0.0709 | -0.0022 | 9.850e-01 |
| DHS | 460 | 0.0693 | -0.0907 | 4.843e-01 |
| ENIGMA | 11373 | 0.0614 | -0.1012 | 2.482e-04 |
| EPOZ | 251 | 0.0358 | 0.1134 | 6.370e-01 |
| ERF | 122 | 0.0754 | -0.0393 | 8.712e-01 |
| FHS | 4710 | 0.055 | -0.0803 | 7.560e-02 |
| GenR | 265 | 0.0412 | 0.301 | 1.684e-01 |
| HUNT | 851 | 0.0664 | -0.1288 | 1.859e-01 |
| LBC1936 | 577 | 0.0658 | -0.2825 | 1.734e-02 |
| LIFE | 1825 | 0.0699 | -0.1019 | 1.167e-01 |
| LLS | 355 | 0.0808 | 0.2 | 1.464e-01 |
| MethCT. | 114 | 0.2002 | -0.0225 | 8.919e-01 |
| NOMAS | 141 | 0.0854 | -0.0098 | 9.633e-01 |
| PoznanMS | 204 | 0.0469 | -0.3474 | 1.381e-01 |
| PROSPER | 315 | 0.0598 | -0.1467 | 3.827e-01 |
| RS1 | 1103 | 0.0354 | -0.1152 | 3.172e-01 |
| RS2 | 1159 | 0.0349 | -0.0094 | 9.342e-01 |
| RS3 | 2582 | 0.037 | -0.0202 | 7.842e-01 |
| RUSH1 | 184 | 0.0859 | 0.1237 | 5.072e-01 |
| RUSH2 | 106 | 0.0472 | 0.001 | 9.978e-01 |
| SHIP | 966 | 0.0712 | -0.0129 | 8.843e-01 |
| SHIP-TREND | 858 | 0.0804 | -0.0357 | 6.878e-01 |
| SYS.adolescents | 980 | 0.059 | -0.1879 | 4.996e-02 |
| SYS.adults | 594 | 0.0582 | -0.076 | 5.402e-01 |
| TREND | 1084 | 0.0413 | 0.1251 | 2.461e-01 |
| UKB | 14571 | 0.0389 | -0.1414 | 3.021e-06 |
| WHICAP | 83 | 0.078 | -0.1627 | 5.756e-01 |
| **Non-European** | | | | |
| ARIC.AA | 675 | 0.29 | -0.019 | 7.510e-01 |
| CARDIA.AA | 132 | 0.265 | 0.1413 | 3.110e-01 |
| CHAP.black | 321 | 0.3074 | -0.1249 | 1.453e-01 |
| FBIRN | 275 | 0.0904 | 0.0324 | 8.278e-01 |
| NOMAS.black | 168 | 0.2568 | -0.0658 | 5.992e-01 |
| NOMAS.hispanic | 718 | 0.1306 | 0.0395 | 6.137e-01 |
| SCES | 209 | 0.0728 | 0.121 | 5.199e-01 |
| SIMES | 201 | 0.1161 | -0.3107 | 4.761e-02 |
| WHICAP.black | 60 | 0.2489 | 0.0017 | 9.933e-01 |
| **Summary European MA** | **73805** | **0.0526** | **-0.074** | **2.68e-10** |
| **Summary African MA** | **1356** | **0.2858** | **-0.034** | **0.424** |
| **Summary Asian MA** | **410** | **0.094** | **-0.111** | **0.3537** |
| **Summary Transancestral MA** | **76678** | **0.058** | **-0.069** | **2.714e-10** |

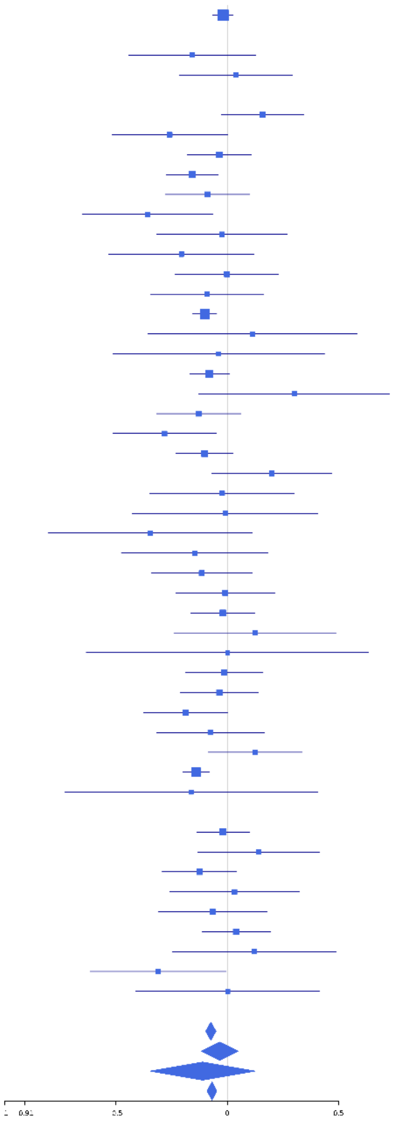

**1-112946028 (CTGTT); chr1:112946028; model 1 (I2=15.7, HetP=0.2406)**

| Study | N_total | EAF | Beta | P |
|---|---|---|---|---|
| **Head circumference** | | | | |
| **CT-based** | | | | |
| **MR-based** | | | | |
| 3c-Dijon | 1499 | 0.344 | 0.0436 | 2.572e-01 |
| AGES | 2226 | 0.3383 | 0.0029 | 9.261e-01 |
| ASPS-Family | 338 | 0.3882 | 0.0405 | 6.076e-01 |
| ASRB | 233 | 0.3297 | 0.0442 | 6.540e-01 |
| CHS | 544 | 0.3481 | 0.04 | 5.293e-01 |
| ENIGMA | 7204 | 0.3603 | 0.0656 | 1.549e-04 |
| ERF | 122 | 0.3103 | 0.0143 | 9.177e-01 |
| FHS | 4710 | 0.351 | 0.0896 | 3.322e-05 |
| HUNT | 851 | 0.3571 | 0.0503 | 3.202e-01 |
| LBC1936 | 577 | 0.3377 | 0.0936 | 1.328e-01 |
| LIFE | 1825 | 0.3838 | 0.0332 | 3.301e-01 |
| LLS | 355 | 0.3587 | 0.1051 | 1.793e-01 |
| MethCT. | 114 | 0.5454 | 0.1256 | 3.451e-01 |
| NOMAS | 141 | 0.4036 | -0.1151 | 3.446e-01 |
| PROSPER | 315 | 0.3566 | -0.0061 | 9.419e-01 |
| SHIP | 966 | 0.3833 | 0.0265 | 5.714e-01 |
| SHIP-TREND | 858 | 0.3805 | 0.017 | 7.321e-01 |
| WHICAP | 83 | 0.4074 | -0.1358 | 3.925e-01 |
| **Non-European** | | | | |
| ARIC.AA | 675 | 0.7219 | 0.0496 | 4.145e-01 |
| CARDIA.AA | 132 | 0.734 | 0.0018 | 9.899e-01 |
| FBIRN | 275 | 0.471 | -0.1305 | 1.278e-01 |
| NOMAS.black | 168 | 0.7094 | 0.1006 | 4.037e-01 |
| NOMAS.hispanic | 718 | 0.4859 | 0.0967 | 6.752e-02 |
| SCES | 209 | 0.5845 | -0.234 | 1.837e-02 |
| SIMES | 201 | 0.4731 | -0.1098 | 2.724e-01 |
| **Summary European MA** | **22847** | **0.358** | **0.053** | **4.71e-08** |
| **Summary African MA** | **975** | **0.7214** | **0.052** | **0.3029** |
| **Summary Asian MA** | **410** | **0.5299** | **-0.172** | **0.0142** |
| **Summary Transancestral MA** | **25339** | **0.3805** | **0.048** | **1.23e-07** |

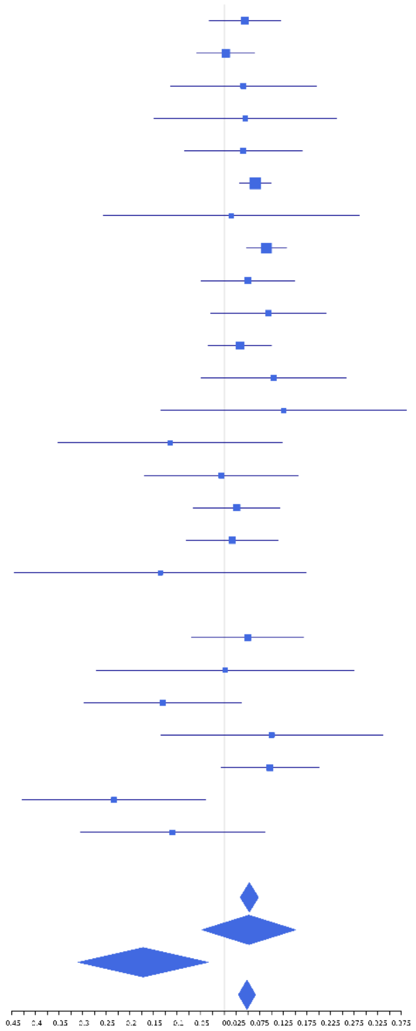

**rs2884218 (T); chr1:210557309; model 1 (I2=27.6, HetP=0.03832)**

| Study | N_total | EAF | Beta | P |
|---|---|---|---|---|
| **Head circumference** | | | | |
| Haworth | 18881 | 0.6994 | 0.0138 | 2.194e-01 |
| HKOS | 888 | 0.7029 | 0.0442 | 3.947e-01 |
| VIDI | 739 | 0.7148 | 0.0648 | 2.613e-01 |
| **CT-based** | | | | |
| ESS | 641 | 0.6943 | 0.0834 | 1.687e-01 |
| UCL | 642 | 0.7273 | -0.0179 | 7.757e-01 |
| **MR-based** | | | | |
| 1000Brains | 751 | 0.734 | 0.1285 | 2.813e-02 |
| 3C-Bordeaux | 564 | 0.7089 | 0.1751 | 7.751e-03 |
| 3c-Dijon | 1499 | 0.7075 | 0.0778 | 5.264e-02 |
| AGES | 2226 | 0.6863 | 0.0414 | 1.997e-01 |
| ARIC.EA | 1390 | 0.6939 | -0.017 | 6.801e-01 |
| ASPS-Family | 338 | 0.7428 | 0.0173 | 8.438e-01 |
| ASRB | 233 | 0.6866 | 0.011 | 9.128e-01 |
| CARDIA.EA | 349 | 0.7101 | 0.147 | 7.810e-02 |
| CHAP | 261 | 0.6993 | -0.0193 | 8.403e-01 |
| CHS | 544 | 0.7217 | 0.0864 | 2.015e-01 |
| DNGS | 513 | 0.7143 | 0.1205 | 8.169e-02 |
| ENIGMA | 11373 | 0.7058 | 0.0449 | 2.032e-03 |
| EPOZ | 251 | 0.7261 | -0.0829 | 4.076e-01 |
| ERF | 122 | 0.6102 | -0.0637 | 6.279e-01 |
| FHS | 4710 | 0.7112 | 0.0046 | 8.398e-01 |
| GenR | 265 | 0.7248 | 0.081 | 4.049e-01 |
| HUNT | 851 | 0.6927 | 0.076 | 1.479e-01 |
| LBC1936 | 577 | 0.6806 | -0.096 | 1.283e-01 |
| LIFE | 1825 | 0.7283 | 0.0334 | 3.699e-01 |
| LLS | 355 | 0.6985 | 0.0261 | 7.494e-01 |
| MethCT. | 114 | 0.7747 | 0.3067 | 5.298e-02 |
| NOMAS | 141 | 0.7567 | -0.0905 | 5.154e-01 |
| PoznanMS | 204 | 0.7341 | -0.0123 | 9.127e-01 |
| PROSPER | 315 | 0.6574 | -6e-04 | 9.941e-01 |
| RS1 | 1103 | 0.6882 | 0.0758 | 9.904e-02 |
| RS2 | 1159 | 0.6885 | 0.0438 | 3.286e-01 |
| RS3 | 2582 | 0.6972 | 0.0632 | 3.680e-02 |
| RUSH1 | 184 | 0.6901 | -0.2974 | 9.089e-03 |
| RUSH2 | 106 | 0.6904 | -0.1517 | 3.099e-01 |
| SHIP | 966 | 0.7126 | -0.0441 | 3.806e-01 |
| SHIP-TREND | 858 | 0.7022 | 0.0561 | 2.883e-01 |
| SYS.adolescents | 980 | 0.748 | 0.0276 | 5.955e-01 |
| SYS.adults | 594 | 0.7521 | 0.0579 | 3.886e-01 |
| TREND | 1084 | 0.7249 | 0.043 | 3.709e-01 |
| UKB | 14571 | 0.6937 | 0.0439 | 5.471e-04 |
| WHICAP | 83 | 0.7226 | -0.1642 | 3.468e-01 |
| **Non-European** | | | | |
| ARIC.AA | 675 | 0.8421 | 0.1243 | 9.578e-02 |
| CARDIA.AA | 132 | 0.831 | 0.3048 | 6.343e-02 |
| CHAP.black | 321 | 0.822 | 0.1587 | 1.250e-01 |
| FBIRN | 275 | 0.715 | 0.0094 | 9.207e-01 |
| IMH | 37 | 0.6228 | -0.2111 | 3.790e-01 |
| NOMAS.black | 168 | 0.848 | 0.4328 | 4.965e-03 |
| NOMAS.hispanic | 718 | 0.739 | -0.0525 | 3.823e-01 |
| SCES | 209 | 0.71 | 0.2027 | 6.017e-02 |
| SIMES | 201 | 0.7294 | 0.0162 | 8.859e-01 |
| WHICAP.black | 60 | 0.8026 | -0.3571 | 1.254e-01 |
| **Summary European MA** | **74830** | **0.7027** | **0.033** | **7.463e-09** |
| **Summary African MA** | **1356** | **0.8352** | **0.164** | **0.001539** |
| **Summary Asian MA** | **1335** | **0.7058** | **0.057** | **0.178** |
| **Summary Transancestral MA** | **78628** | **0.7055** | **0.034** | **4.774e-10** |

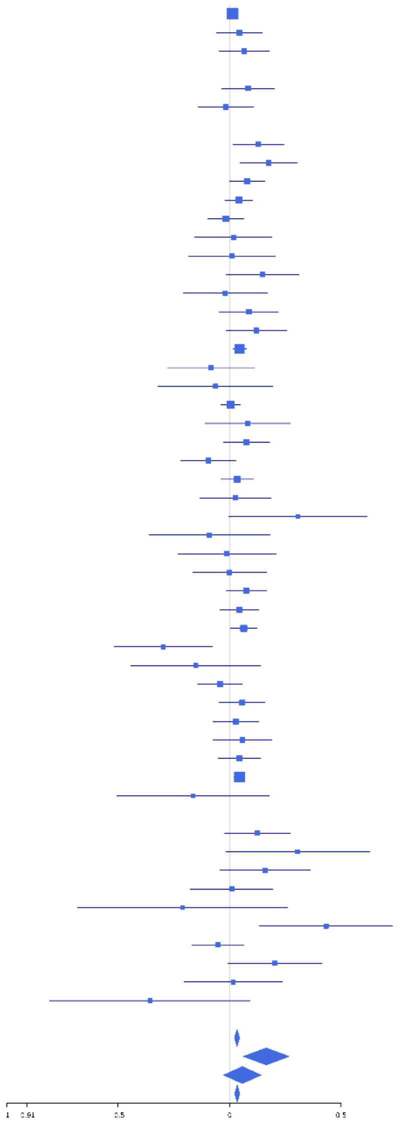

**rs6429430 (T); chr1:243834477; model 1 (I2=0, HetP=0.8582)**

| Study | N_total | EAF | Beta | P |
|---|---|---|---|---|
| **Head circumference** | | | | |
| Haworth | 18881 | 0.8218 | -0.0566 | 2.658e-05 |
| HKOS | 888 | 0.6959 | -7e-04 | 9.897e-01 |
| VIDI | 745 | 0.7901 | -0.1307 | 4.020e-02 |
| **CT-based** | | | | |
| ESS | 641 | 0.8031 | 0.067 | 3.403e-01 |
| UCL | 642 | 0.8158 | -0.1378 | 5.556e-02 |
| **MR-based** | | | | |
| 1000Brains | 751 | 0.833 | -0.0391 | 5.720e-01 |
| 3C-Bordeaux | 564 | 0.8092 | -0.0846 | 2.644e-01 |
| 3c-Dijon | 1499 | 0.8089 | -0.025 | 5.907e-01 |
| AGES | 2226 | 0.7729 | 0.0063 | 8.600e-01 |
| ARIC.EA | 1390 | 0.8162 | -0.0919 | 6.066e-02 |
| ASPS-Family | 338 | 0.8076 | -0.1464 | 1.337e-01 |
| ASRB | 233 | 0.7904 | -0.0552 | 6.283e-01 |
| CARDIA.EA | 349 | 0.8251 | 0.0178 | 8.582e-01 |
| CHAP | 261 | 0.8487 | 0.1252 | 3.062e-01 |
| CHS | 544 | 0.8401 | -0.0485 | 5.580e-01 |
| DNGS | 514 | 0.7572 | -0.0049 | 9.460e-01 |
| ENIGMA | 11373 | 0.8156 | -0.0501 | 3.355e-03 |
| EPOZ | 251 | 0.7981 | -0.1417 | 2.027e-01 |
| ERF | 122 | 0.8265 | -0.2546 | 1.320e-01 |
| FHS | 4710 | 0.811 | -0.0579 | 2.774e-02 |
| GenR | 265 | 0.7833 | 0.0523 | 6.196e-01 |
| HUNT | 851 | 0.8162 | -0.0541 | 3.868e-01 |
| LBC1936 | 577 | 0.8243 | -0.1474 | 5.680e-02 |
| LIFE | 1825 | 0.8339 | -0.0296 | 5.052e-01 |
| LLS | 355 | 0.813 | 0.0287 | 7.654e-01 |
| MethCT. | 114 | 0.7129 | 0.095 | 5.165e-01 |
| NOMAS | 141 | 0.8889 | -0.2433 | 2.015e-01 |
| PoznanMS | 204 | 0.8465 | -0.0621 | 6.514e-01 |
| PROSPER | 315 | 0.8049 | -0.0151 | 8.810e-01 |
| RS1 | 1103 | 0.8036 | 0.002 | 9.700e-01 |
| RS2 | 1159 | 0.8095 | -0.0639 | 2.271e-01 |
| RS3 | 2582 | 0.8052 | -0.0555 | 1.144e-01 |
| RUSH1 | 184 | 0.8402 | -0.1499 | 2.932e-01 |
| RUSH2 | 106 | 0.7877 | -0.1446 | 3.913e-01 |
| SHIP | 966 | 0.8312 | -0.0647 | 2.871e-01 |
| SHIP-TREND | 858 | 0.8491 | -0.0606 | 3.695e-01 |
| SYS.adolescents | 980 | 0.8146 | -0.067 | 2.489e-01 |
| SYS.adults | 594 | 0.8247 | -0.08 | 2.945e-01 |
| TREND | 1084 | 0.8354 | -0.1372 | 1.800e-02 |
| UKB | 14571 | 0.8139 | -0.059 | 8.796e-05 |
| WHICAP | 83 | 0.8513 | -0.1817 | 4.074e-01 |
| **Non-European** | | | | |
| ARIC.AA | 675 | 0.6622 | 0.0518 | 3.675e-01 |
| CARDIA.AA | 132 | 0.669 | -0.1014 | 4.384e-01 |
| CHAP.black | 321 | 0.6909 | 0.1552 | 7.018e-02 |
| FBIRN | 275 | 0.7679 | 0.0992 | 3.272e-01 |
| NOMAS.black | 168 | 0.6957 | -0.0509 | 6.683e-01 |
| NOMAS.hispanic | 718 | 0.7584 | -0.0594 | 3.357e-01 |
| SCES | 209 | 0.6791 | -0.0796 | 4.470e-01 |
| SIMES | 201 | 0.6448 | -0.0095 | 9.259e-01 |
| WHICAP.black | 60 | 0.6427 | 0.2025 | 2.925e-01 |
| **Summary European MA** | **74837** | **0.8156** | **-0.055** | **2.206e-16** |
| **Summary African MA** | **1356** | **0.6729** | **0.055** | **0.1784** |
| **Summary Asian MA** | **1298** | **0.6853** | **-0.015** | **0.7245** |
| **Summary Transancestral MA** | **78598** | **0.8102** | **-0.051** | **3.037e-15** |

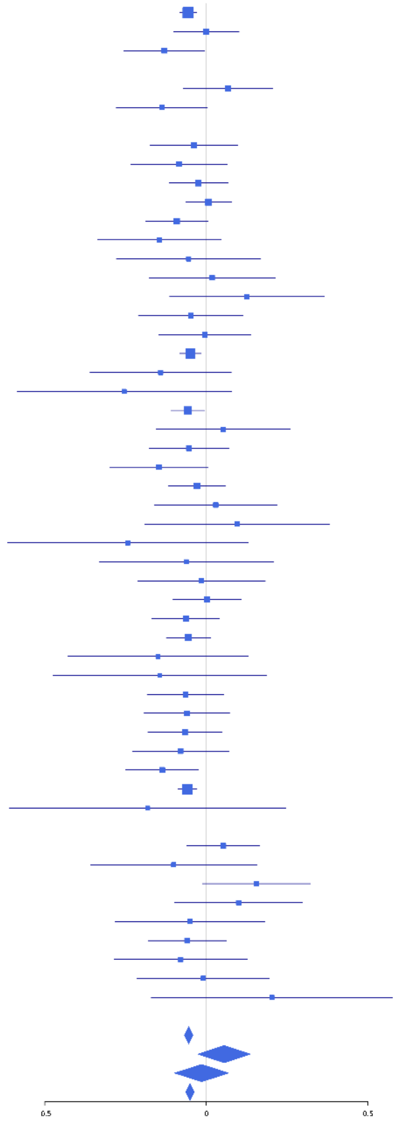

**rs6736289 (C); chr2:10195540; model 1 (I2=2.6, HetP=0.4221)**

| Study | N_total | EAF | Beta | P |
|---|---|---|---|---|
| **Head circumference** | | | | |
| Haworth | 18881 | 0.7922 | -0.0373 | 3.174e-03 |
| HKOS | 888 | 0.8778 | -0.1306 | 7.148e-02 |
| VIDI | 723 | 0.9076 | -0.0311 | 7.322e-01 |
| **CT-based** | | | | |
| ESS | 641 | 0.7869 | 0.0106 | 8.771e-01 |
| UCL | 642 | 0.8002 | -0.0442 | 5.266e-01 |
| **MR-based** | | | | |
| 1000Brains | 751 | 0.8186 | -0.0878 | 1.902e-01 |
| 3C-Bordeaux | 564 | 0.8035 | -0.2291 | 2.334e-03 |
| 3c-Dijon | 1499 | 0.8012 | -0.1477 | 1.246e-03 |
| AGES | 2226 | 0.7897 | -0.1151 | 1.747e-03 |
| ARIC.EA | 1390 | 0.8071 | -0.0943 | 4.977e-02 |
| ASPS-Family | 338 | 0.2411 | -0.0151 | 8.666e-01 |
| ASRB | 233 | 0.79 | -0.0366 | 7.481e-01 |
| CARDIA.EA | 349 | 0.8043 | 0.0927 | 3.311e-01 |
| CHAP | 261 | 0.78 | -0.1038 | 3.269e-01 |
| CHS | 544 | 0.8224 | -0.1023 | 1.974e-01 |
| DHS | 460 | 0.7547 | -0.1771 | 2.084e-02 |
| ENIGMA | 11373 | 0.8038 | -0.0284 | 8.944e-02 |
| EPOZ | 251 | 0.7739 | -0.0377 | 7.243e-01 |
| FHS | 4710 | 0.7976 | 0.0066 | 7.976e-01 |
| GenR | 265 | 0.7884 | -0.1189 | 2.635e-01 |
| HUNT | 851 | 0.798 | -0.0043 | 9.426e-01 |
| LBC1936 | 577 | 0.8028 | -0.0468 | 5.275e-01 |
| LIFE | 1825 | 0.8326 | -0.0843 | 5.752e-02 |
| LLS | 355 | 0.7543 | -0.0642 | 4.609e-01 |
| MethCT. | 114 | 0.7846 | -0.2561 | 1.118e-01 |
| NOMAS | 141 | 0.8588 | 0.0675 | 6.936e-01 |
| PoznanMS | 204 | 0.8603 | 0.058 | 6.847e-01 |
| PROSPER | 315 | 0.7841 | 0.0038 | 9.691e-01 |
| RS1 | 1103 | 0.7918 | -0.0354 | 4.990e-01 |
| RS2 | 1159 | 0.7853 | 0.0576 | 2.551e-01 |
| RS3 | 2582 | 0.7921 | -0.1094 | 1.427e-03 |
| RUSH1 | 184 | 0.8197 | -0.0677 | 6.181e-01 |
| RUSH2 | 106 | 0.8112 | -0.143 | 4.170e-01 |
| SHIP | 966 | 0.8069 | -0.0233 | 6.858e-01 |
| SHIP-TREND | 858 | 0.8293 | -0.0537 | 4.026e-01 |
| SYS.adolescents | 980 | 0.7763 | -0.0026 | 9.618e-01 |
| SYS.adults | 594 | 0.778 | -0.0352 | 6.144e-01 |
| TREND | 1084 | 0.8119 | 0.0169 | 7.587e-01 |
| UKB | 14571 | 0.7919 | -0.0299 | 3.819e-02 |
| WHICAP | 83 | 0.7718 | -0.209 | 2.621e-01 |
| **Non-European** | | | | |
| ARIC.AA | 675 | 0.7421 | -0.0016 | 9.793e-01 |
| CARDIA.AA | 132 | 0.756 | 0.0116 | 9.354e-01 |
| CHAP.black | 321 | 0.7509 | -0.1471 | 1.079e-01 |
| FBIRN | 275 | 0.8123 | -0.0638 | 5.595e-01 |
| NOMAS.black | 168 | 0.7666 | -0.1232 | 3.411e-01 |
| NOMAS.hispanic | 718 | 0.8214 | -0.049 | 4.775e-01 |
| SCES | 209 | 0.849 | -0.191 | 1.622e-01 |
| SIMES | 201 | 0.8472 | -0.0944 | 4.959e-01 |
| WHICAP.black | 60 | 0.7457 | -2e-04 | 9.988e-01 |
| **Summary European MA** | **74639** | **0.7953** | **-0.04** | **3.318e-10** |
| **Summary African MA** | **1356** | **0.7487** | **-0.049** | **0.2667** |
| **Summary Asian MA** | **1298** | **0.8684** | **-0.135** | **0.02036** |
| **Summary Transancestral MA** | **78400** | **0.796** | **-0.042** | **1.603e-11** |

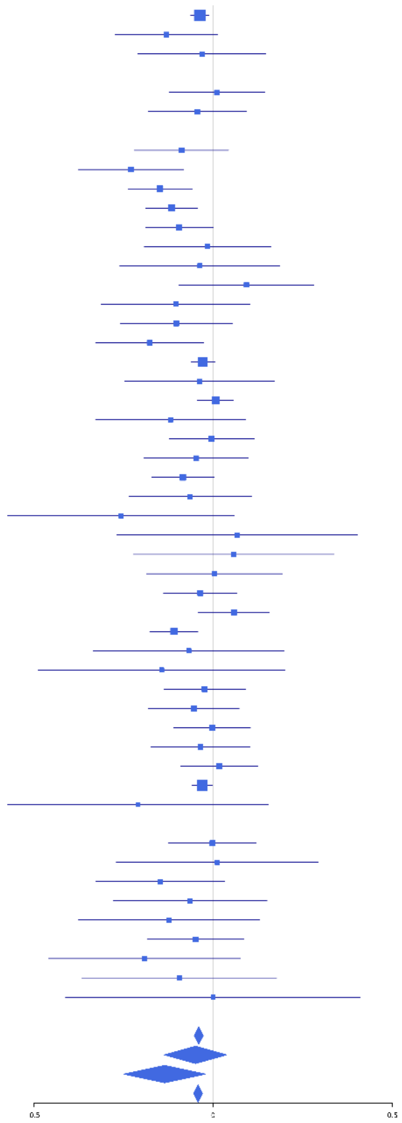

**rs11685359 (T); chr2:20349433; model 1 (I2=0, HetP=0.5754)**

| Study | N_total | EAF | Beta | P |
|---|---|---|---|---|
| **Head circumference** | | | | |
| Haworth | 18881 | 0.156 | -0.0153 | 2.809e-01 |
| HKOS | 888 | 0.2427 | 0.043 | 4.376e-01 |
| VIDI | 740 | 0.2352 | 0.0968 | 1.147e-01 |
| **CT-based** | | | | |
| ESS | 641 | 0.1757 | -0.1092 | 1.368e-01 |
| UCL | 642 | 0.1575 | -0.0907 | 2.363e-01 |
| **MR-based** | | | | |
| 1000Brains | 751 | 0.1493 | 0.0815 | 2.609e-01 |
| 3C-Bordeaux | 564 | 0.1134 | -0.0806 | 3.911e-01 |
| 3c-Dijon | 1499 | 0.1195 | -0.0272 | 6.288e-01 |
| AGES | 2226 | 0.1698 | -0.0424 | 2.882e-01 |
| ARIC.EA | 1390 | 0.159 | -0.0898 | 8.325e-02 |
| ASPS-Family | 338 | 0.1303 | -0.1626 | 1.547e-01 |
| ASRB | 232 | 0.1581 | -0.0793 | 5.338e-01 |
| CARDIA.EA | 349 | 0.1585 | -0.1492 | 1.498e-01 |
| CHAP | 261 | 0.1453 | 0.1097 | 3.779e-01 |
| CHS | 544 | 0.1287 | -0.2018 | 2.583e-02 |
| DHS | 460 | 0.1381 | -0.0941 | 3.246e-01 |
| DNGS | 495 | 0.1654 | -0.1238 | 1.485e-01 |
| ENIGMA | 11373 | 0.1597 | -0.0648 | 3.431e-04 |
| EPOZ | 251 | 0.1757 | -0.0185 | 8.747e-01 |
| ERF | 122 | 0.1186 | -0.3359 | 8.984e-02 |
| FHS | 4710 | 0.1394 | -0.0417 | 1.611e-01 |
| GenR | 265 | 0.1425 | -0.0936 | 4.513e-01 |
| HUNT | 851 | 0.1758 | -0.0508 | 4.257e-01 |
| LBC1936 | 577 | 0.1427 | -0.0278 | 7.413e-01 |
| LIFE | 1825 | 0.165 | -0.1031 | 2.096e-02 |
| LLS | 355 | 0.1638 | 0.0426 | 6.742e-01 |
| MethCT. | 114 | 0.1592 | -0.0691 | 7.022e-01 |
| NOMAS | 141 | 0.1036 | -0.1376 | 4.825e-01 |
| PoznanMS | 204 | 0.1359 | 0.1501 | 2.986e-01 |
| PROSPER | 315 | 0.1504 | 0.0115 | 9.182e-01 |
| RS1 | 1103 | 0.1604 | -0.0513 | 3.767e-01 |
| RS2 | 1159 | 0.1618 | -0.0149 | 7.908e-01 |
| RS3 | 2582 | 0.1622 | -0.0852 | 2.392e-02 |
| RUSH1 | 184 | 0.1522 | 0.0327 | 8.222e-01 |
| RUSH2 | 106 | 0.1722 | 0.1015 | 5.779e-01 |
| SHIP | 966 | 0.1765 | 0.0385 | 5.187e-01 |
| SHIP-TREND | 858 | 0.1643 | -0.0339 | 6.023e-01 |
| SYS.adolescents | 980 | 0.1328 | -0.0409 | 5.390e-01 |
| SYS.adults | 594 | 0.1443 | -0.062 | 4.529e-01 |
| TREND | 1084 | 0.165 | -0.0441 | 4.461e-01 |
| UKB | 14571 | 0.1552 | -0.0448 | 5.581e-03 |
| WHICAP | 83 | 0.1046 | 0.2166 | 3.958e-01 |
| **Non-European** | | | | |
| ARIC.AA | 675 | 0.0294 | -0.1821 | 2.586e-01 |
| CHAP.black | 321 | 0.0434 | 0.0194 | 9.205e-01 |
| FBIRN | 275 | 0.1365 | 0.1122 | 3.676e-01 |
| NOMAS.black | 168 | 0.0268 | 0.4804 | 1.569e-01 |
| NOMAS.hispanic | 716 | 0.1029 | 0.0298 | 7.319e-01 |
| SCES | 209 | 0.258 | 0.1422 | 2.033e-01 |
| SIMES | 201 | 0.2874 | -0.0708 | 5.208e-01 |
| WHICAP.black | 60 | 0.0419 | 0.2351 | 6.078e-01 |
| **Summary European MA** | **75272** | **0.1559** | **-0.042** | **5.082e-09** |
| **Summary African MA** | **1224** | **0.0333** | **-0.017** | **0.881** |
| **Summary Asian MA** | **1298** | **0.2521** | **0.041** | **0.3682** |
| **Summary Transancestral MA** | **78899** | **0.155** | **-0.039** | **3.016e-08** |

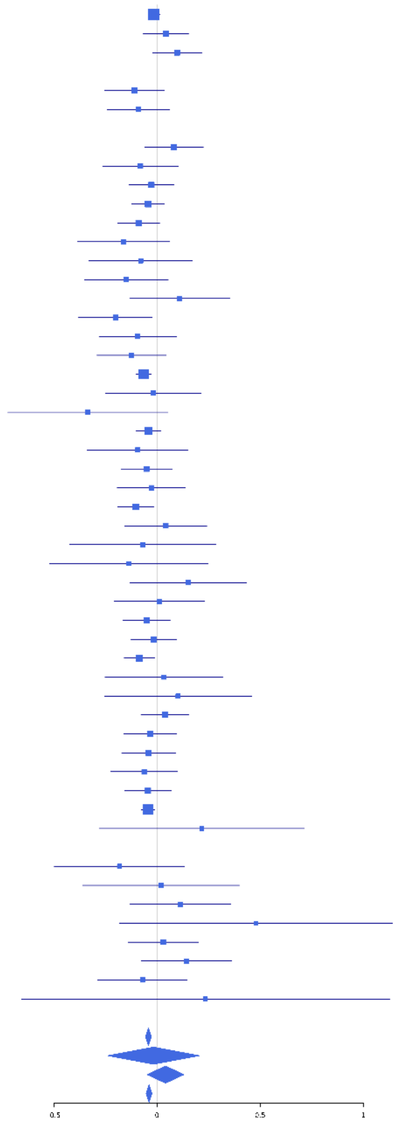

**rs62135193 (T); chr2:48755477; model 1 (I2=4.9, HetP=0.3747)**

| Study | N_total | EAF | Beta | P |
|---|---|---|---|---|
| **Head circumference** | | | | |
| Haworth | 18881 | 0.5431 | -0.0285 | 5.966e-03 |
| HKOS | 888 | 0.1486 | -0.0916 | 1.697e-01 |
| VIDI | 676 | 0.4155 | 0.0095 | 8.638e-01 |
| **CT-based** | | | | |
| ESS | 641 | 0.5436 | -0.052 | 3.538e-01 |
| UCL | 642 | 0.5556 | 0.039 | 4.870e-01 |
| **MR-based** | | | | |
| 1000Brains | 751 | 0.5503 | -0.1324 | 1.087e-02 |
| 3C-Bordeaux | 564 | 0.5842 | -0.0281 | 6.420e-01 |
| 3c-Dijon | 1499 | 0.575 | -0.0798 | 3.067e-02 |
| AGES | 2226 | 0.4798 | -0.0322 | 2.824e-01 |
| ARIC.EA | 1390 | 0.5441 | -0.0915 | 1.621e-02 |
| ASPS-Family | 338 | 0.5308 | -0.0902 | 2.421e-01 |
| ASRB | 233 | 0.5532 | -0.0215 | 8.172e-01 |
| CARDIA.EA | 349 | 0.5484 | -0.0189 | 8.034e-01 |
| CHAP | 261 | 0.5897 | -0.2402 | 7.410e-03 |
| CHS | 544 | 0.5572 | 0.0108 | 8.596e-01 |
| ENIGMA | 11373 | 0.5415 | -0.0148 | 2.674e-01 |
| EPOZ | 251 | 0.5151 | 0.0037 | 9.671e-01 |
| ERF | 122 | 0.6297 | 0.0166 | 9.002e-01 |
| FHS | 4710 | 0.5485 | -0.0292 | 1.588e-01 |
| GenR | 265 | 0.5385 | 0.0536 | 5.386e-01 |
| HUNT | 851 | 0.498 | -0.0087 | 8.571e-01 |
| LBC1936 | 577 | 0.5621 | 0.063 | 2.886e-01 |
| LIFE | 1825 | 0.5281 | -0.033 | 3.196e-01 |
| LLS | 355 | 0.5459 | -0.0058 | 9.388e-01 |
| MethCT. | 114 | 0.4888 | 0.034 | 7.972e-01 |
| NOMAS | 141 | 0.5961 | 0.0078 | 9.489e-01 |
| PoznanMS | 204 | 0.487 | -0.1231 | 2.139e-01 |
| PROSPER | 315 | 0.5462 | -0.1308 | 1.021e-01 |
| RS1 | 1103 | 0.5416 | -0.1065 | 1.271e-02 |
| RS2 | 1159 | 0.5267 | -0.0569 | 1.717e-01 |
| RS3 | 2582 | 0.5406 | -0.067 | 1.651e-02 |
| RUSH1 | 184 | 0.5964 | -0.0885 | 4.060e-01 |
| RUSH2 | 106 | 0.5487 | -0.065 | 6.389e-01 |
| SHIP | 966 | 0.5333 | 0.0317 | 4.871e-01 |
| SHIP-TREND | 858 | 0.528 | 0.097 | 4.529e-02 |
| SYS.adolescents | 980 | 0.5391 | -0.0648 | 1.531e-01 |
| SYS.adults | 594 | 0.5539 | -0.0028 | 9.619e-01 |
| TREND | 1084 | 0.5373 | -0.1122 | 9.279e-03 |
| UKB | 14571 | 0.5384 | -0.0169 | 1.510e-01 |
| WHICAP | 83 | 0.5676 | -0.0675 | 6.675e-01 |
| **Non-European** | | | | |
| ARIC.AA | 675 | 0.4773 | -0.0477 | 3.818e-01 |
| CARDIA.AA | 132 | 0.479 | -0.0357 | 7.720e-01 |
| CHAP.black | 321 | 0.4871 | -0.013 | 8.687e-01 |
| FBIRN | 275 | 0.5017 | -0.1212 | 1.566e-01 |
| IMH | 37 | 0.1491 | 0.4337 | 1.838e-01 |
| NOMAS.black | 168 | 0.4854 | 0.0493 | 6.520e-01 |
| NOMAS.hispanic | 718 | 0.5729 | -0.0966 | 7.067e-02 |
| SCES | 209 | 0.1611 | -0.0969 | 4.668e-01 |
| SIMES | 201 | 0.1726 | -0.0111 | 9.088e-01 |
| WHICAP.black | 60 | 0.4936 | -0.0256 | 8.892e-01 |
| **Summary European MA** | **74254** | **0.5394** | **-0.029** | **2.363e-08** |
| **Summary African MA** | **1356** | **0.4815** | **-0.025** | **0.5103** |
| **Summary Asian MA** | **1335** | **0.1542** | **-0.066** | **0.2184** |
| **Summary Transancestral MA** | **78052** | **0.5319** | **-0.03** | **2.834e-09** |

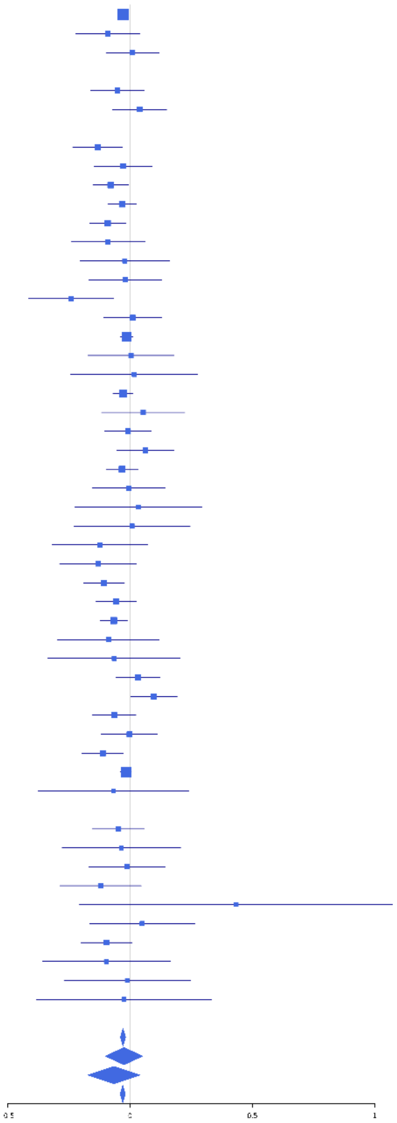

**rs41288837 (T); chr2:85531116; model 1 (I2=18.6, HetP=0.1725)**

| Study | N_total | EAF | Beta | P |
|---|---|---|---|---|
| **Head circumference** | | | | |
| Haworth | 17737 | 0.029 | -0.1519 | 1.588e-06 |
| VIDI | 755 | 0.0295 | 0.2933 | 5.412e-02 |
| **CT-based** | | | | |
| ESS | 641 | 0.031 | -0.0524 | 7.454e-01 |
| UCL | 642 | 0.0251 | -0.0419 | 8.145e-01 |
| **MR-based** | | | | |
| 1000Brains | 751 | 0.0332 | -0.0944 | 5.125e-01 |
| 3C-Bordeaux | 564 | 0.0153 | -0.1294 | 5.936e-01 |
| 3c-Dijon | 1499 | 0.0164 | -0.0095 | 9.475e-01 |
| AGES | 2226 | 0.0329 | -0.1683 | 4.510e-02 |
| ARIC.EA | 1390 | 0.0244 | 0.0285 | 8.166e-01 |
| ASPS-Family | 338 | 0.021 | -0.3193 | 2.344e-01 |
| ASRB | 233 | 0.029 | 0.016 | 9.534e-01 |
| CHS | 544 | 0.0218 | -0.5869 | 4.747e-03 |
| ENIGMA | 11265 | 0.0287 | -0.1088 | 6.412e-03 |
| EPOZ | 251 | 0.0285 | 0.3336 | 2.138e-01 |
| ERF | 122 | 0.0221 | 0.0409 | 9.252e-01 |
| FHS | 4710 | 0.0273 | -0.0865 | 1.707e-01 |
| GenR | 265 | 0.022 | 0.1598 | 5.899e-01 |
| HUNT | 851 | 0.0271 | -0.0724 | 6.277e-01 |
| LBC1936 | 577 | 0.0314 | -0.0447 | 7.911e-01 |
| LIFE | 1825 | 0.0294 | 0.007 | 9.431e-01 |
| LLS | 355 | 0.0213 | -0.3583 | 1.680e-01 |
| NOMAS | 141 | 0.0223 | -1.072 | 8.798e-03 |
| PoznanMS | 204 | 0.0291 | -0.2742 | 3.517e-01 |
| PROSPER | 315 | 0.0351 | -0.124 | 5.665e-01 |
| RS1 | 1103 | 0.0326 | -0.0541 | 6.519e-01 |
| RS2 | 1159 | 0.0322 | -0.1095 | 3.524e-01 |
| RS3 | 2582 | 0.031 | -0.0856 | 2.863e-01 |
| SHIP | 966 | 0.0346 | 0.1242 | 3.186e-01 |
| SHIP-TREND | 858 | 0.0274 | -0.1182 | 4.248e-01 |
| SYS.adolescents | 980 | 0.0457 | -0.0155 | 8.866e-01 |
| SYS.adults | 594 | 0.0379 | 0.2751 | 7.029e-02 |
| TREND | 1084 | 0.0335 | 0.0167 | 8.888e-01 |
| UKB | 14571 | 0.0252 | -0.0711 | 5.687e-02 |
| WHICAP | 83 | 0.0129 | -0.099 | 8.860e-01 |
| **Non-European** | | | | |
| **Summary European MA** | **72181** | **0.0282** | **-0.091** | **1.029e-08** |
| **Summary Transancestral MA** | **72181** | **0.0282** | **-0.091** | **1.029e-08** |

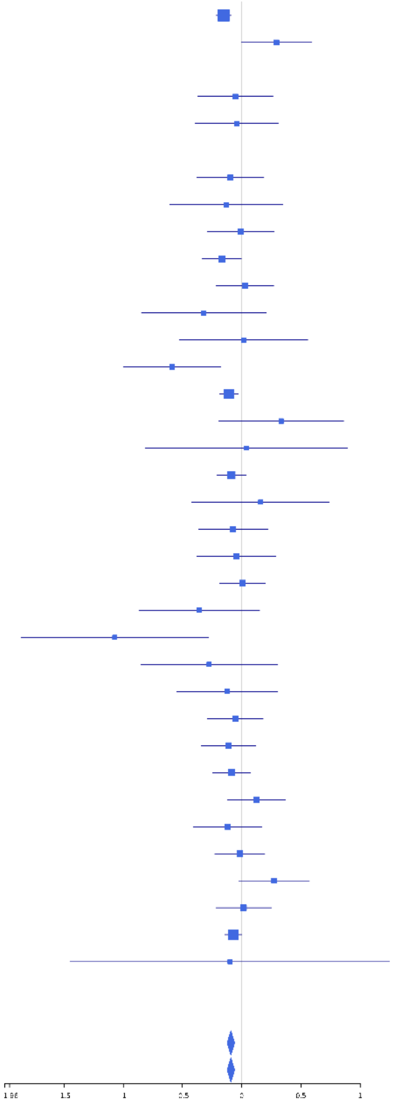

#### rs4832160 (A); chr2:85539170; model 1 (I2=18.1, HetP=0.1377)

| Study | N_total | EAF | Beta | P |
|---|---|---|---|---|
| **Head circumference** | | | | |
| Haworth | 18881 | 0.4896 | -0.0428 | 3.141e-05 |
| HKOS | 888 | 0.0991 | 0.0041 | 9.587e-01 |
| VIDI | 733 | 0.4319 | 0.045 | 3.936e-01 |
| **CT-based** | | | | |
| ESS | 641 | 0.4521 | -0.0593 | 2.907e-01 |
| UCL | 642 | 0.5016 | -0.1524 | 6.324e-03 |
| **MR-based** | | | | |
| 1000Brains | 751 | 0.4851 | 0.0351 | 4.969e-01 |
| 3C-Bordeaux | 564 | 0.4787 | -0.1251 | 3.628e-02 |
| 3c-Dijon | 1499 | 0.4906 | 0.0467 | 2.013e-01 |
| AGES | 2226 | 0.5125 | -0.0376 | 2.096e-01 |
| ARIC.EA | 1390 | 0.4799 | -0.0377 | 3.200e-01 |
| ASPS-Family | 338 | 0.4944 | -0.1799 | 1.940e-02 |
| ASRB | 233 | 0.5208 | -0.0658 | 4.785e-01 |
| CARDIA.EA | 349 | 0.4763 | 0.0549 | 4.690e-01 |
| CHAP | 261 | 0.4423 | -0.147 | 9.645e-02 |
| CHS | 544 | 0.4835 | 0.0228 | 7.071e-01 |
| DNGS | 510 | 0.367 | 0.0157 | 8.087e-01 |
| ENIGMA | 11373 | 0.4951 | -0.017 | 2.009e-01 |
| EPOZ | 251 | 0.5049 | -0.1119 | 2.101e-01 |
| ERF | 122 | 0.4939 | 0.0622 | 6.270e-01 |
| FHS | 4710 | 0.4808 | -0.0329 | 1.112e-01 |
| GenR | 265 | 0.456 | -0.1213 | 1.643e-01 |
| HUNT | 851 | 0.5127 | 0.0186 | 7.014e-01 |
| LBC1936 | 577 | 0.5174 | -0.0129 | 8.265e-01 |
| LIFE | 1825 | 0.4878 | -0.064 | 5.323e-02 |
| LLS | 355 | 0.5058 | 0.0019 | 9.804e-01 |
| MethCT. | 114 | 0.4638 | 0.0704 | 5.961e-01 |
| NOMAS | 141 | 0.4474 | 0.0275 | 8.187e-01 |
| PoznanMS | 204 | 0.4974 | 0.0572 | 5.632e-01 |
| PROSPER | 315 | 0.4828 | 0.0324 | 6.845e-01 |
| RS1 | 1103 | 0.507 | -0.0188 | 6.591e-01 |
| RS2 | 1159 | 0.502 | 0.0169 | 6.849e-01 |
| RS3 | 2582 | 0.5 | -0.0391 | 1.602e-01 |
| RUSH1 | 184 | 0.4927 | 0.164 | 1.175e-01 |
| RUSH2 | 106 | 0.5014 | -0.2944 | 3.455e-02 |
| SHIP | 966 | 0.5138 | -0.0417 | 3.600e-01 |
| SHIP-TREND | 858 | 0.5105 | 0.0611 | 2.062e-01 |
| SYS.adolescents | 980 | 0.4765 | -0.1031 | 2.263e-02 |
| SYS.adults | 594 | 0.4653 | -0.0397 | 4.948e-01 |
| TREND | 1084 | 0.5171 | 0.036 | 4.025e-01 |
| UKB | 14571 | 0.4856 | -0.0373 | 1.470e-03 |
| WHICAP | 83 | 0.4478 | 0.0869 | 5.791e-01 |
| **Non-European** | | | | |
| ARIC.AA | 675 | 0.4665 | 0.0153 | 7.795e-01 |
| CARDIA.AA | 132 | 0.445 | -0.0248 | 8.412e-01 |
| CHAP.black | 321 | 0.469 | -0.0276 | 7.274e-01 |
| FBIRN | 275 | 0.3669 | 0.0181 | 8.377e-01 |
| NOMAS.black | 168 | 0.4564 | 0.0206 | 8.510e-01 |
| NOMAS.hispanic | 718 | 0.427 | -0.0472 | 3.770e-01 |
| SCES | 209 | 0.1099 | 0.013 | 9.335e-01 |
| SIMES | 201 | 0.1303 | 0.3267 | 2.647e-02 |
| WHICAP.black | 60 | 0.4281 | -0.0911 | 6.233e-01 |
| **Summary European MA** | **74821** | **0.4895** | **-0.031** | **3.175e-09** |
| **Summary African MA** | **1356** | **0.462** | **-0.003** | **0.9427** |
| **Summary Asian MA** | **1298** | **0.1057** | **0.061** | **0.3423** |
| **Summary Transancestral MA** | **78582** | **0.4816** | **-0.029** | **1.076e-08** |

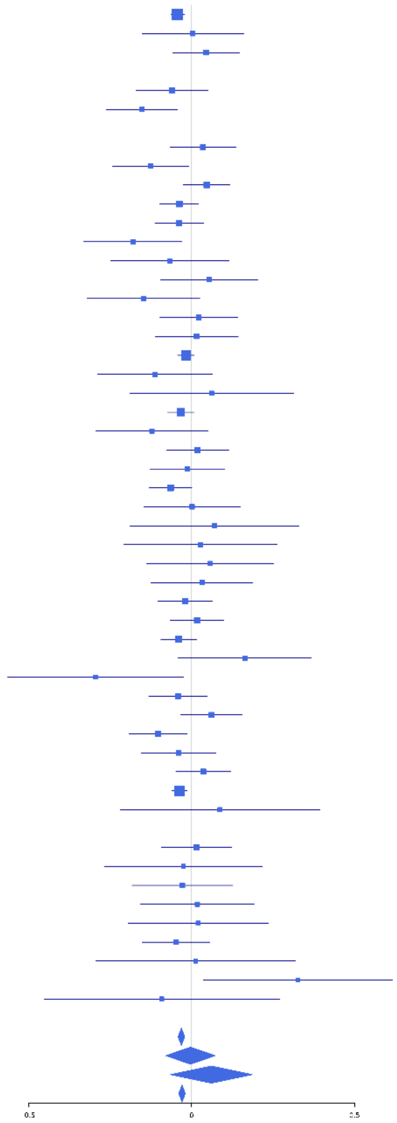

**rs867529 (C); chr2:88913273; model 1 (I2=9.9, HetP=0.2761)**

| Study | N_total | EAF | Beta | P |
|---|---|---|---|---|
| **Head circumference** | | | | |
| Haworth | 18881 | 0.2813 | 0.0376 | 9.966e-04 |
| HKOS | 888 | 0.5107 | 0.0157 | 7.406e-01 |
| VIDI | 755 | 0.3352 | -4e-04 | 9.945e-01 |
| **CT-based** | | | | |
| ESS | 641 | 0.2936 | -0.1329 | 3.021e-02 |
| UCL | 642 | 0.3009 | -0.0429 | 4.809e-01 |
| **MR-based** | | | | |
| 1000Brains | 751 | 0.282 | 0.021 | 7.143e-01 |
| 3C-Bordeaux | 564 | 0.2769 | 0.0828 | 2.141e-01 |
| 3c-Dijon | 1499 | 0.2758 | 0.0859 | 3.557e-02 |
| AGES | 2226 | 0.2573 | 0.0545 | 1.120e-01 |
| ARIC.EA | 1390 | 0.2853 | 0.0394 | 3.485e-01 |
| ASPS-Family | 338 | 0.2411 | 0.0708 | 4.313e-01 |
| ASRB | 233 | 0.2635 | 0.1332 | 2.064e-01 |
| CARDIA.EA | 349 | 0.2809 | 0.1481 | 7.860e-02 |
| CHAP | 261 | 0.2951 | 0.0235 | 8.069e-01 |
| CHS | 544 | 0.2583 | 0.008 | 9.075e-01 |
| ENIGMA | 11373 | 0.2834 | 0.0602 | 4.271e-05 |
| EPOZ | 251 | 0.3223 | -0.0688 | 4.714e-01 |
| ERF | 122 | 0.3407 | -0.0894 | 5.081e-01 |
| FHS | 4710 | 0.267 | -0.0221 | 3.439e-01 |
| GenR | 265 | 0.2742 | 0.1471 | 1.308e-01 |
| HUNT | 851 | 0.3015 | 0.0394 | 4.558e-01 |
| LBC1936 | 577 | 0.2702 | -0.0658 | 3.207e-01 |
| LIFE | 1825 | 0.2566 | 0.028 | 4.591e-01 |
| LLS | 355 | 0.3226 | 0.0257 | 7.489e-01 |
| MethCT. | 114 | 0.1916 | -0.0182 | 9.143e-01 |
| NOMAS | 141 | 0.2619 | -0.0169 | 9.007e-01 |
| PoznanMS | 204 | 0.2594 | 0.1261 | 2.643e-01 |
| PROSPER | 315 | 0.3154 | -0.138 | 1.074e-01 |
| RS1 | 1103 | 0.2961 | 0.0127 | 7.856e-01 |
| RS2 | 1159 | 0.3066 | 0.0288 | 5.219e-01 |
| RS3 | 2582 | 0.2858 | 0.048 | 1.193e-01 |
| RUSH1 | 184 | 0.3224 | -0.0222 | 8.422e-01 |
| RUSH2 | 106 | 0.3113 | -0.0179 | 9.042e-01 |
| SHIP | 966 | 0.262 | 0.0898 | 8.308e-02 |
| SHIP-TREND | 858 | 0.3048 | 0.0621 | 2.364e-01 |
| SYS.adolescents | 980 | 0.234 | -0.0065 | 9.035e-01 |
| SYS.adults | 594 | 0.2521 | 0.1204 | 7.150e-02 |
| TREND | 1084 | 0.2727 | 0.1115 | 2.092e-02 |
| UKB | 14571 | 0.2811 | 0.0199 | 1.273e-01 |
| WHICAP | 83 | 0.179 | 0.1282 | 5.286e-01 |
| **Non-European** | | | | |
| ARIC.AA | 675 | 0.0688 | -0.1182 | 2.718e-01 |
| CARDIA.AA | 132 | 0.062 | -0.1524 | 5.507e-01 |
| CHAP.black | 321 | 0.056 | 0.2182 | 2.045e-01 |
| FBIRN | 275 | 0.2969 | -0.1046 | 2.635e-01 |
| IMH | 37 | 0.4825 | 0.0233 | 9.199e-01 |
| NOMAS.black | 168 | 0.1181 | -0.1707 | 3.141e-01 |
| NOMAS.hispanic | 718 | 0.2147 | 0.0245 | 7.035e-01 |
| SCES | 209 | 0.4805 | 0.1874 | 5.564e-02 |
| SIMES | 201 | 0.4948 | 0.1011 | 3.115e-01 |
| WHICAP.black | 60 | 0.0417 | -0.168 | 7.146e-01 |
| **Summary European MA** | **74333** | **0.2802** | **0.034** | **6.415e-09** |
| **Summary African MA** | **1356** | **0.07** | **-0.058** | **0.4381** |
| **Summary Asian MA** | **1335** | **0.5028** | **0.056** | **0.1508** |
| **Summary Transancestral MA** | **78131** | **0.2797** | **0.032** | **1.101e-08** |

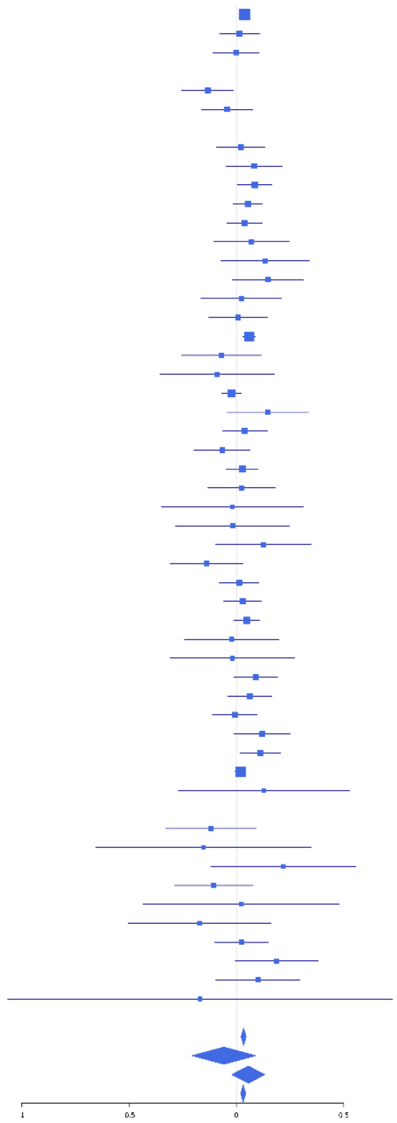

**rs73977799 (A); chr2:183699768; model 1 (I2=0, HetP=0.7725)**

| Study | N_total | EAF | Beta | P |
|---|---|---|---|---|
| **Head circumference** | | | | |
| Haworth | 18881 | 0.0938 | 0.0497 | 4.837e-03 |
| HKOS | 888 | 0.0315 | -0.0073 | 9.570e-01 |
| VIDI | 755 | 0.0306 | -0.1674 | 2.628e-01 |
| **CT-based** | | | | |
| ESS | 641 | 0.0935 | 0.0903 | 3.469e-01 |
| UCL | 642 | 0.0776 | 0.0717 | 4.919e-01 |
| **MR-based** | | | | |
| 1000Brains | 751 | 0.083 | -0.041 | 6.612e-01 |
| 3C-Bordeaux | 564 | 0.1038 | -0.0565 | 5.627e-01 |
| 3c-Dijon | 1499 | 0.0957 | 0.0719 | 2.470e-01 |
| ARIC.EA | 1390 | 0.0806 | -0.0339 | 6.260e-01 |
| ASPS-Family | 338 | 0.1163 | 0.0703 | 5.579e-01 |
| ASRB | 233 | 0.0578 | 0.1246 | 5.309e-01 |
| CARDIA.EA | 349 | 0.0845 | 0.0015 | 9.909e-01 |
| CHAP | 261 | 0.0729 | 0.0285 | 8.656e-01 |
| ENIGMA | 11373 | 0.0824 | 0.0394 | 1.018e-01 |
| EPOZ | 251 | 0.1008 | 0.156 | 2.929e-01 |
| FHS | 4710 | 0.0841 | 0.0654 | 7.821e-02 |
| GenR | 265 | 0.1 | 0.2099 | 1.470e-01 |
| HUNT | 851 | 0.0925 | 0.1086 | 1.941e-01 |
| LBC1936 | 577 | 0.0693 | -0.2079 | 7.286e-02 |
| LIFE | 1825 | 0.0825 | 0.0989 | 1.002e-01 |
| LLS | 355 | 0.0763 | 0.1989 | 1.594e-01 |
| NOMAS | 141 | 0.0873 | -0.1871 | 3.767e-01 |
| PoznanMS | 204 | 0.0805 | 0.16 | 3.793e-01 |
| PROSPER | 315 | 0.0824 | -0.0322 | 8.246e-01 |
| RS1 | 1103 | 0.0934 | 0.1183 | 1.059e-01 |
| RS2 | 1159 | 0.0998 | 0.1435 | 3.840e-02 |
| RS3 | 2582 | 0.1005 | 0.0107 | 8.169e-01 |
| RUSH1 | 184 | 0.0728 | -0.0357 | 8.586e-01 |
| RUSH2 | 106 | 0.0834 | 0.357 | 1.537e-01 |
| SHIP | 966 | 0.0817 | 0.0843 | 3.103e-01 |
| SHIP-TREND | 858 | 0.0775 | 0.049 | 5.876e-01 |
| SYS.adolescents | 980 | 0.0687 | 0.1617 | 7.036e-02 |
| SYS.adults | 594 | 0.0645 | 0.0195 | 8.691e-01 |
| TREND | 1084 | 0.089 | -0.0207 | 7.845e-01 |
| UKB | 14571 | 0.0819 | 0.0822 | 1.207e-04 |
| WHICAP | 83 | 0.0705 | -0.5236 | 8.823e-02 |
| **Non-European** | | | | |
| ARIC.AA | 675 | 0.1641 | -0.0271 | 7.125e-01 |
| CARDIA.AA | 132 | 0.163 | 0.123 | 4.607e-01 |
| CHAP.black | 321 | 0.1305 | 0.0532 | 6.504e-01 |
| FBIRN | 275 | 0.0819 | 0.1082 | 4.868e-01 |
| NOMAS.black | 168 | 0.1854 | -0.1334 | 3.437e-01 |
| NOMAS.hispanic | 718 | 0.1286 | 0.0607 | 4.415e-01 |
| WHICAP.black | 60 | 0.0901 | 0.0223 | 9.443e-01 |
| **Summary European MA** | **71441** | **0.0863** | **0.054** | **7.797e-09** |
| **Summary African MA** | **1356** | **0.1554** | **-0.007** | **0.8982** |
| **Summary Asian MA** | **888** | **0.0315** | **-0.007** | **0.957** |
| **Summary Transancestral MA** | **74678** | **0.0873** | **0.053** | **9.42e-09** |

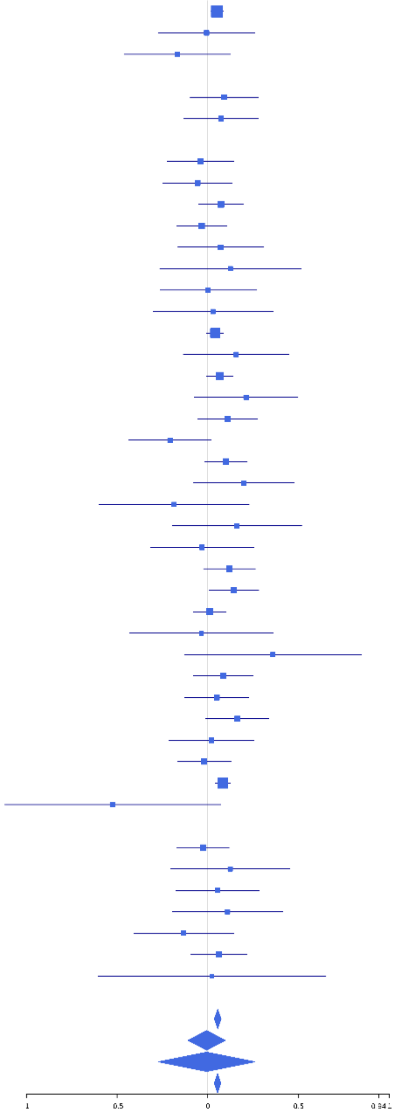

**rs288326 (A); chr2:183703336; model 1 (I2=0, HetP=0.9901)**

| Study | N_total | EAF | Beta | P |
|---|---|---|---|---|
| **Head circumference** | | | | |
| Haworth | 18881 | 0.1241 | 0.085 | 4.620e-08 |
| VIDI | 755 | 0.0629 | -0.0208 | 8.444e-01 |
| **CT-based** | | | | |
| UCL | 642 | 0.1066 | 0.1621 | 7.321e-02 |
| **MR-based** | | | | |
| 1000Brains | 751 | 0.1207 | -0.0024 | 9.761e-01 |
| 3C-Bordeaux | 564 | 0.1305 | 0.0721 | 4.150e-01 |
| 3c-Dijon | 1499 | 0.1287 | 0.0484 | 3.747e-01 |
| AGES | 2226 | 0.0799 | 0.0569 | 3.033e-01 |
| ARIC.EA | 1390 | 0.1148 | 0.0359 | 5.458e-01 |
| ASPS-Family | 338 | 0.152 | -0.0464 | 6.648e-01 |
| ASRB | 233 | 0.1499 | -0.0658 | 6.124e-01 |
| CARDIA.EA | 349 | 0.1231 | 0.1242 | 2.812e-01 |
| CHAP | 261 | 0.111 | 0.096 | 4.913e-01 |
| CHS | 544 | 0.1009 | 0.0062 | 9.509e-01 |
| DNGS | 516 | 0.0775 | 0.0645 | 5.797e-01 |
| ENIGMA | 11373 | 0.1251 | 0.0623 | 1.888e-03 |
| EPOZ | 251 | 0.1385 | 0.1522 | 2.389e-01 |
| ERF | 122 | 0.0526 | -0.0915 | 7.496e-01 |
| FHS | 4710 | 0.1152 | 0.0683 | 3.425e-02 |
| GenR | 265 | 0.1152 | 0.1863 | 1.710e-01 |
| HUNT | 851 | 0.1119 | 0.0334 | 6.643e-01 |
| LBC1936 | 577 | 0.12 | 0.0855 | 3.452e-01 |
| LIFE | 1825 | 0.1162 | 0.065 | 2.081e-01 |
| LLS | 355 | 0.173 | 0.0127 | 8.980e-01 |
| NOMAS | 141 | 0.125 | -0.0079 | 9.648e-01 |
| PoznanMS | 204 | 0.1057 | -0.0855 | 5.954e-01 |
| PROSPER | 315 | 0.1263 | 0.1962 | 1.018e-01 |
| RS1 | 1103 | 0.1314 | 0.1163 | 6.497e-02 |
| RS2 | 1159 | 0.1415 | 0.0205 | 7.305e-01 |
| RS3 | 2582 | 0.1365 | 0.0825 | 4.171e-02 |
| RUSH1 | 184 | 0.1143 | -0.0354 | 8.295e-01 |
| RUSH2 | 106 | 0.0849 | 0.3615 | 1.457e-01 |
| SHIP | 966 | 0.1258 | 0.0495 | 4.702e-01 |
| SHIP-TREND | 858 | 0.1142 | 0.1163 | 1.257e-01 |
| SYS.adolescents | 980 | 0.0965 | 0.1008 | 1.874e-01 |
| SYS.adults | 594 | 0.1083 | -0.0433 | 6.423e-01 |
| TREND | 1084 | 0.118 | 0.0379 | 5.685e-01 |
| UKB | 14571 | 0.1219 | 0.0303 | 9.063e-02 |
| WHICAP | 83 | 0.128 | 0.0423 | 8.564e-01 |
| **Non-European** | | | | |
| ARIC.AA | 675 | 0.0225 | -0.0885 | 6.295e-01 |
| CHAP.black | 321 | 0.0343 | 0.0971 | 6.541e-01 |
| FBIRN | 275 | 0.0904 | 0.1042 | 4.838e-01 |
| NOMAS.black | 168 | 0.0476 | 0.2593 | 3.129e-01 |
| NOMAS.hispanic | 718 | 0.0749 | 0.0279 | 7.811e-01 |
| WHICAP.black | 60 | 0.0169 | 0.9419 | 1.894e-01 |
| **Summary European MA** | **74208** | **0.1206** | **0.061** | **2.152e-14** |
| **Summary African MA** | **1224** | **0.0288** | **0.065** | **0.5922** |
| **Summary Transancestral MA** | **76425** | **0.1185** | **0.061** | **1.771e-14** |

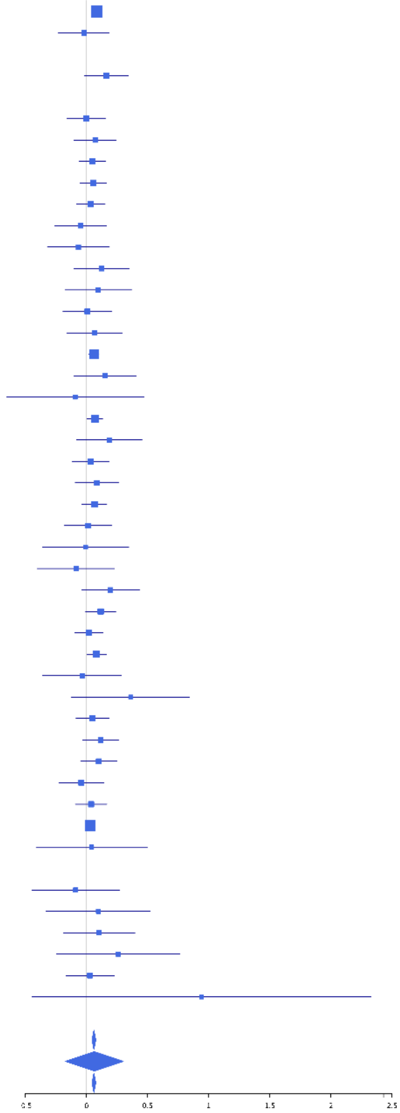

**rs11928552 (T); chr3:48739883; model 1 (I2=17.1, HetP=0.1688)**

| Study | N_total | EAF | Beta | P |
|---|---|---|---|---|
| **Head circumference** | | | | |
| Haworth | 12731 | 0.3693 | -0.0344 | 7.937e-03 |
| HKOS | 888 | 0.0315 | 0.2002 | 1.404e-01 |
| VIDI | 742 | 0.3316 | -0.138 | 1.255e-02 |
| **CT-based** | | | | |
| ESS | 641 | 0.346 | -0.0214 | 7.151e-01 |
| UCL | 642 | 0.3629 | -0.1164 | 4.481e-02 |
| **MR-based** | | | | |
| 1000Brains | 751 | 0.3947 | 0.0023 | 9.646e-01 |
| 3C-Bordeaux | 564 | 0.374 | -0.0524 | 3.946e-01 |
| 3c-Dijon | 1499 | 0.4382 | -0.0286 | 4.376e-01 |
| AGES | 2226 | 0.3841 | -0.0149 | 6.277e-01 |
| ARIC.EA | 1390 | 0.3685 | 0.0512 | 1.930e-01 |
| ASPS-Family | 338 | 0.4872 | -0.2602 | 7.213e-04 |
| ASRB | 233 | 0.3882 | -0.0087 | 9.272e-01 |
| CHS | 544 | 0.4122 | -0.0215 | 7.273e-01 |
| DNGS | 502 | 0.2552 | -0.0286 | 6.926e-01 |
| ENIGMA | 11373 | 0.4163 | -0.0209 | 1.205e-01 |
| EPOZ | 251 | 0.4146 | 0.0953 | 2.927e-01 |
| ERF | 122 | 0.4558 | -0.0526 | 6.823e-01 |
| FHS | 4710 | 0.4129 | -0.0125 | 5.505e-01 |
| GenR | 265 | 0.3793 | -0.0423 | 6.367e-01 |
| HUNT | 851 | 0.4362 | -0.0652 | 1.823e-01 |
| LBC1936 | 577 | 0.4119 | -0.0276 | 6.444e-01 |
| LIFE | 1825 | 0.4407 | -0.0774 | 2.034e-02 |
| LLS | 355 | 0.4574 | 0.0588 | 4.356e-01 |
| MethCT. | 114 | 0.4409 | 0.103 | 4.404e-01 |
| NOMAS | 141 | 0.3587 | 0.0829 | 5.055e-01 |
| PoznanMS | 204 | 0.3578 | 0.0816 | 4.298e-01 |
| PROSPER | 315 | 0.4352 | -0.1933 | 1.619e-02 |
| RS1 | 1103 | 0.3948 | 0.0215 | 6.215e-01 |
| RS2 | 1159 | 0.3855 | -0.0719 | 9.170e-02 |
| RS3 | 2582 | 0.3928 | -0.063 | 2.706e-02 |
| SHIP | 966 | 0.4411 | -0.0166 | 7.173e-01 |
| SHIP-TREND | 858 | 0.4078 | -0.0624 | 2.045e-01 |
| SYS.adolescents | 980 | 0.3888 | -0.0939 | 4.269e-02 |
| SYS.adults | 594 | 0.3854 | -0.0854 | 1.521e-01 |
| TREND | 1084 | 0.3983 | -0.0085 | 8.474e-01 |
| UKB | 14571 | 0.3538 | -0.0394 | 1.308e-03 |
| WHICAP | 83 | 0.3243 | 0.0403 | 8.086e-01 |
| **Non-European** | | | | |
| ARIC.AA | 675 | 0.4552 | -0.0068 | 9.006e-01 |
| CARDIA.AA | 132 | 0.429 | -0.0066 | 9.575e-01 |
| FBIRN | 275 | 0.314 | 0.0486 | 5.971e-01 |
| NOMAS.black | 168 | 0.4308 | 0.1348 | 2.229e-01 |
| NOMAS.hispanic | 718 | 0.3826 | -0.0833 | 1.253e-01 |
| WHICAP.black | 60 | 0.3833 | -0.0362 | 8.477e-01 |
| **Summary European MA** | **67772** | **0.3868** | **-0.034** | **1.173e-09** |
| **Summary African MA** | **1035** | **0.4437** | **0.014** | **0.7453** |
| **Summary Asian MA** | **888** | **0.0315** | **0.2** | **0.1404** |
| **Summary Transancestral MA** | **70802** | **0.3829** | **-0.032** | **5.272e-09** |

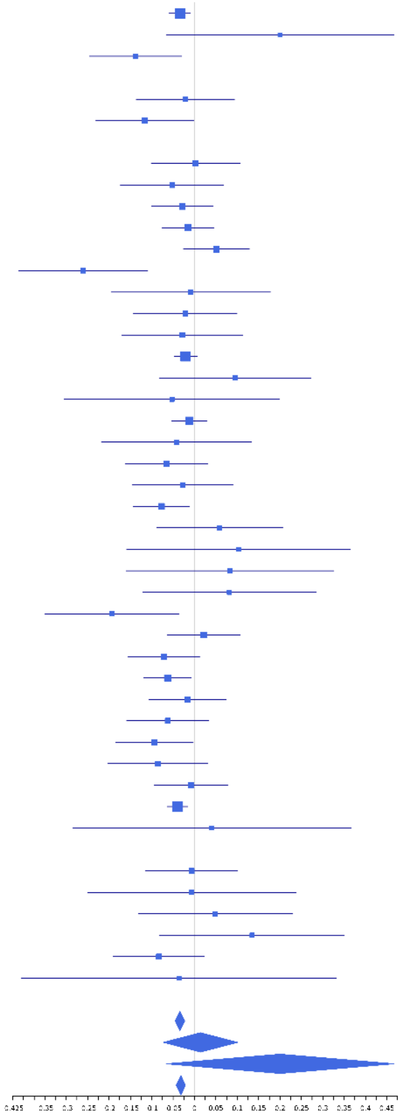

**rs145897003 (A); chr3:52210649; model 1 (I2=41.4, HetP=0.01856)**

| Study | N_total | EAF | Beta | P |
|---|---|---|---|---|
| **Head circumference** | | | | |
| Haworth | 18584 | 0.0236 | 0.1092 | 1.408e-03 |
| VIDI | 738 | 0.012 | -0.056 | 8.149e-01 |
| **CT-based** | | | | |
| ESS | 641 | 0.0185 | 0.5247 | 1.136e-02 |
| UCL | 642 | 0.0133 | -0.0825 | 7.349e-01 |
| **MR-based** | | | | |
| 1000Brains | 751 | 0.0212 | 5e-04 | 9.979e-01 |
| 3C-Bordeaux | 564 | 0.0134 | 0.2752 | 2.883e-01 |
| 3c-Dijon | 1499 | 0.0166 | 0.3548 | 1.292e-02 |
| ARIC.EA | 1390 | 0.0206 | 0.3727 | 5.205e-03 |
| DNGS | 509 | 0.0123 | 0.4699 | 9.881e-02 |
| ENIGMA | 4691 | 0.0275 | 0.1377 | 2.916e-02 |
| HUNT | 851 | 0.0321 | -0.1002 | 4.658e-01 |
| LBC1936 | 577 | 0.022 | -0.4915 | 1.432e-02 |
| LIFE | 1825 | 0.0159 | -0.0145 | 9.124e-01 |
| NOMAS | 141 | 0.0172 | 0.1617 | 7.243e-01 |
| PROSPER | 315 | 0.0222 | -0.1251 | 6.430e-01 |
| RS1 | 1103 | 0.02 | 0.2162 | 1.550e-01 |
| RS2 | 1159 | 0.0217 | 0.1038 | 4.667e-01 |
| RS3 | 2582 | 0.0198 | -0.0133 | 8.940e-01 |
| SHIP | 966 | 0.0256 | 0.3216 | 2.578e-02 |
| SHIP-TREND | 858 | 0.0211 | 0.2191 | 1.929e-01 |
| TREND | 1084 | 0.0254 | 0.0485 | 7.230e-01 |
| UKB | 14571 | 0.0225 | 0.1008 | 1.068e-02 |
| WHICAP | 83 | 0.0171 | 0.1976 | 7.426e-01 |
| **Non-European** | | | | |
| NOMAS.hispanic | 718 | 0.0106 | -0.5411 | 3.605e-02 |
| **Summary European MA** | **56124** | **0.0224** | **0.111** | **3.755e-08** |
| **Summary Transancestral MA** | **56842** | **0.0222** | **0.105** | **1.681e-07** |

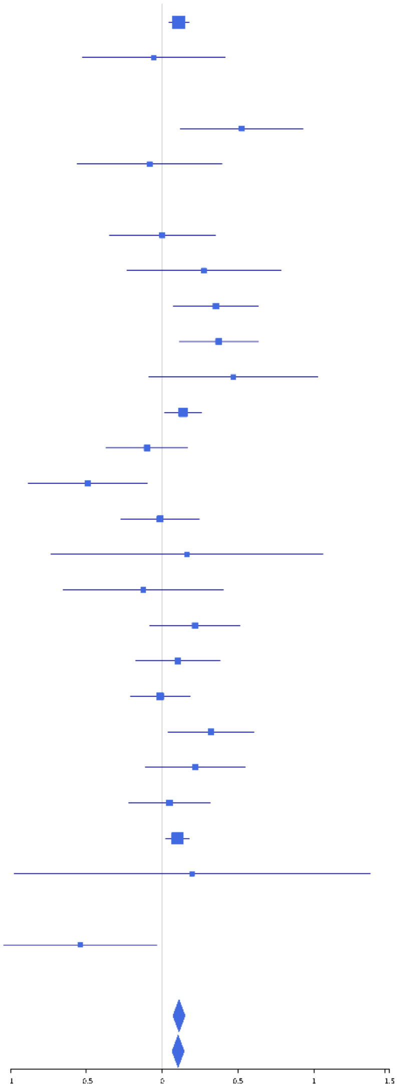

**rs114895876 (A); chr3:127352970; model 1 (I2=21.5, HetP=0.1663)**

| Study | N_total | EAF | Beta | P |
|---|---|---|---|---|
| **Head circumference** | | | | |
| Haworth | 16682 | 0.987 | 0.1042 | 3.094e-02 |
| VIDI | 749 | 0.9591 | 0.217 | 9.655e-02 |
| **CT-based** | | | | |
| UCL | 642 | 0.9898 | 0.065 | 8.150e-01 |
| **MR-based** | | | | |
| 1000Brains | 751 | 0.9828 | 0.2672 | 1.791e-01 |
| 3c-Dijon | 1499 | 0.9899 | 0.1495 | 4.140e-01 |
| AGES | 2226 | 0.9916 | -0.0034 | 9.830e-01 |
| ARIC.EA | 1390 | 0.9869 | 0.1823 | 2.741e-01 |
| ENIGMA | 10372 | 0.9864 | 0.1292 | 3.114e-02 |
| FHS | 4710 | 0.9871 | 0.0871 | 3.395e-01 |
| HUNT | 851 | 0.9784 | 0.3935 | 1.829e-02 |
| LBC1936 | 577 | 0.9924 | 0.464 | 1.710e-01 |
| LIFE | 1825 | 0.9867 | 0.1683 | 2.445e-01 |
| LLS | 355 | 0.9869 | -0.0912 | 7.828e-01 |
| NOMAS | 141 | 0.9888 | -0.1788 | 7.522e-01 |
| PoznanMS | 204 | 0.9712 | 0.1388 | 6.390e-01 |
| PROSPER | 315 | 0.991 | -0.9161 | 2.970e-02 |
| RS1 | 1103 | 0.9854 | 0.0965 | 5.870e-01 |
| RS2 | 1159 | 0.9852 | 0.0014 | 9.937e-01 |
| RS3 | 2582 | 0.9861 | 0.085 | 4.739e-01 |
| SHIP | 966 | 0.9772 | 0.092 | 5.460e-01 |
| SHIP-TREND | 858 | 0.979 | 0.2301 | 1.722e-01 |
| SYS.adolescents | 980 | 0.9897 | -0.3295 | 1.403e-01 |
| TREND | 1084 | 0.9763 | 0.1694 | 2.304e-01 |
| UKB | 14571 | 0.9868 | 0.2212 | 1.661e-05 |
| WHICAP | 83 | 0.9863 | -1.6459 | 1.595e-02 |
| **Non-European** | | | | |
| **Summary European MA** | **66675** | **0.9862** | **0.131** | **2.527e-08** |
| **Summary Transancestral MA** | **66675** | **0.9862** | **0.131** | **2.527e-08** |

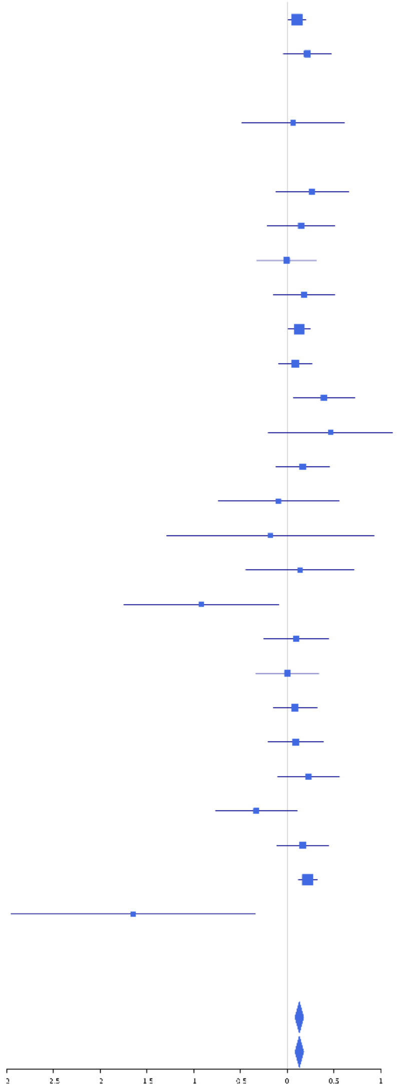

#### rs9818536 (T); chr3:136115295; model 1 (I2=19.5, HetP=0.1209)

| Study | N_total | EAF | Beta | P |
|---|---|---|---|---|
| **Head circumference** | | | | |
| Haworth | 18881 | 0.2494 | -0.0279 | 1.954e-02 |
| VIDI | 755 | 0.2316 | -0.0224 | 7.132e-01 |
| **CT-based** | | | | |
| ESS | 641 | 0.2398 | -0.1618 | 1.335e-02 |
| UCL | 642 | 0.2281 | -0.0128 | 8.476e-01 |
| **MR-based** | | | | |
| 1000Brains | 751 | 0.2494 | -0.0489 | 4.125e-01 |
| 3C-Bordeaux | 564 | 0.2231 | -0.1806 | 1.185e-02 |
| 3c-Dijon | 1499 | 0.221 | -0.0604 | 1.696e-01 |
| AGES | 2226 | 0.2699 | -0.0675 | 4.567e-02 |
| ARIC.EA | 1390 | 0.2416 | -0.0194 | 6.616e-01 |
| ASPS-Family | 338 | 0.2595 | -0.0395 | 6.524e-01 |
| ASRB | 233 | 0.2322 | 0.0482 | 6.612e-01 |
| CARDIA.EA | 349 | 0.239 | -0.0929 | 2.952e-01 |
| CHAP | 261 | 0.2406 | 0.0631 | 5.384e-01 |
| CHS | 544 | 0.2394 | -0.0467 | 5.105e-01 |
| DHS | 460 | 0.2208 | 0.1231 | 1.215e-01 |
| DNGS | 516 | 0.1559 | 0.0215 | 8.019e-01 |
| ENIGMA | 11373 | 0.2357 | -0.0187 | 2.303e-01 |
| EPOZ | 251 | 0.2476 | -0.0141 | 8.918e-01 |
| ERF | 122 | 0.246 | -0.2221 | 1.351e-01 |
| FHS | 4710 | 0.2497 | -0.0594 | 1.258e-02 |
| GenR | 265 | 0.208 | 0.0456 | 6.702e-01 |
| HUNT | 851 | 0.2483 | 0.0376 | 5.019e-01 |
| LBC1936 | 577 | 0.2465 | -0.0418 | 5.405e-01 |
| LIFE | 1825 | 0.2486 | -0.1066 | 5.431e-03 |
| LLS | 355 | 0.2331 | -0.0581 | 5.128e-01 |
| MethCT. | 114 | 0.0919 | 0.1935 | 3.985e-01 |
| NOMAS | 141 | 0.2574 | -0.5302 | 1.549e-04 |
| PoznanMS | 204 | 0.2512 | -0.0774 | 4.980e-01 |
| PROSPER | 315 | 0.2342 | -0.0212 | 8.219e-01 |
| RS1 | 1103 | 0.2471 | -0.036 | 4.651e-01 |
| RS2 | 1159 | 0.242 | -0.0491 | 3.110e-01 |
| RS3 | 2582 | 0.2431 | -0.0213 | 5.117e-01 |
| RUSH1 | 184 | 0.2288 | 0.0688 | 5.806e-01 |
| RUSH2 | 106 | 0.2398 | 0.0901 | 5.766e-01 |
| SHIP | 966 | 0.2315 | -0.1263 | 1.940e-02 |
| SHIP-TREND | 858 | 0.256 | 0.011 | 8.429e-01 |
| SYS.adolescents | 980 | 0.2204 | -0.0848 | 1.198e-01 |
| SYS.adults | 594 | 0.2114 | -0.1627 | 2.207e-02 |
| TREND | 1084 | 0.2361 | 0.0182 | 7.198e-01 |
| UKB | 14571 | 0.2361 | -0.0525 | 1.397e-04 |
| WHICAP | 83 | 0.2785 | -0.1939 | 2.663e-01 |
| **Non-European** | | | | |
| ARIC.AA | 675 | 0.0384 | -0.0966 | 4.953e-01 |
| CARDIA.AA | 132 | 0.046 | -0.3053 | 2.988e-01 |
| CHAP.black | 321 | 0.0639 | 0.0768 | 6.347e-01 |
| FBIRN | 275 | 0.1416 | 0.097 | 4.288e-01 |
| NOMAS.black | 168 | 0.0428 | -0.6523 | 1.661e-02 |
| NOMAS.hispanic | 718 | 0.1318 | -0.0164 | 8.339e-01 |
| SIMES | 201 | 0.0117 | -0.1934 | 6.765e-01 |
| WHICAP.black | 60 | 0.0374 | 0.3127 | 5.182e-01 |
| **Summary European MA** | **75309** | **0.2413** | **-0.04** | **2.22e-11** |
| **Summary African MA** | **1356** | **0.0457** | **-0.118** | **0.2001** |
| **Summary Asian MA** | **201** | **0.0117** | **-0.193** | **0.6765** |
| **Summary Transancestral MA** | **77973** | **0.2357** | **-0.04** | **1.994e-11** |

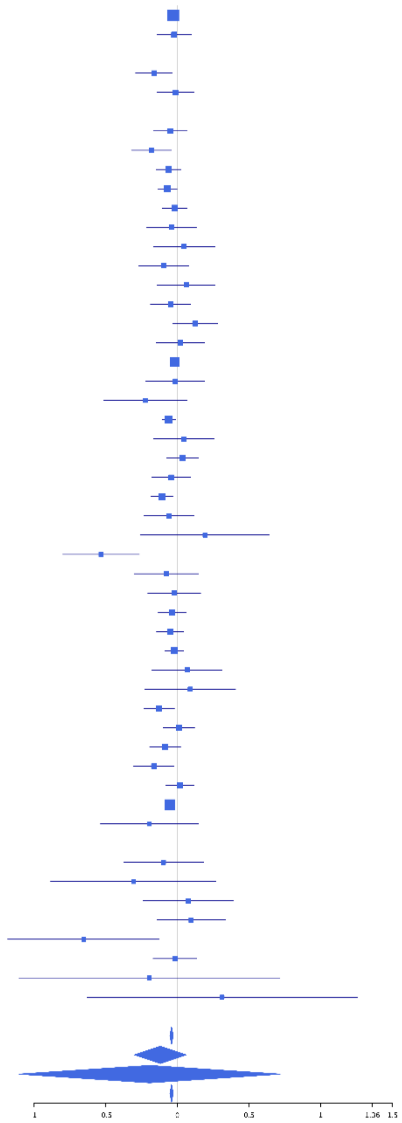

**rs2063453 (T); chr3:141715543; model 1 (I2=0, HetP=0.7585)**

| Study | N_total | EAF | Beta | P |
|---|---|---|---|---|
| **Head circumference** | | | | |
| Haworth | 18881 | 0.8401 | -0.0483 | 5.694e-04 |
| HKOS | 888 | 0.8373 | -0.0663 | 3.020e-01 |
| VIDI | 750 | 0.8287 | -0.0737 | 2.824e-01 |
| **CT-based** | | | | |
| ESS | 641 | 0.8438 | -0.0687 | 3.721e-01 |
| UCL | 642 | 0.8284 | -0.009 | 9.025e-01 |
| **MR-based** | | | | |
| 1000Brains | 751 | 0.845 | -0.1621 | 2.332e-02 |
| 3C-Bordeaux | 564 | 0.845 | 0.001 | 9.907e-01 |
| 3c-Dijon | 1499 | 0.843 | -0.0346 | 4.911e-01 |
| AGES | 2226 | 0.8225 | -0.0069 | 8.594e-01 |
| ARIC.EA | 1390 | 0.8446 | -0.0183 | 7.268e-01 |
| ASPS-Family | 338 | 0.8476 | -0.0443 | 6.790e-01 |
| ASRB | 233 | 0.8396 | -0.0635 | 6.152e-01 |
| CARDIA.EA | 349 | 0.8419 | 0.0252 | 8.076e-01 |
| CHAP | 261 | 0.8534 | 0.0738 | 5.514e-01 |
| CHS | 544 | 0.8564 | -0.0807 | 3.501e-01 |
| DHS | 460 | 0.8412 | -0.1704 | 5.889e-02 |
| DNGS | 516 | 0.8585 | -0.0659 | 4.610e-01 |
| ENIGMA | 11373 | 0.8472 | -0.0527 | 4.261e-03 |
| EPOZ | 251 | 0.8444 | 0.0344 | 7.806e-01 |
| ERF | 122 | 0.8561 | -0.1167 | 5.224e-01 |
| FHS | 4710 | 0.8455 | -0.0949 | 8.734e-04 |
| GenR | 265 | 0.8631 | 0.0631 | 6.178e-01 |
| HUNT | 851 | 0.8643 | -0.2254 | 1.450e-03 |
| LBC1936 | 577 | 0.8305 | -0.0489 | 5.334e-01 |
| LIFE | 1825 | 0.8396 | -0.0851 | 5.926e-02 |
| LLS | 355 | 0.8618 | -0.2169 | 4.616e-02 |
| MethCT. | 114 | 0.8738 | -0.0447 | 8.226e-01 |
| NOMAS | 141 | 0.8929 | -0.1718 | 3.739e-01 |
| PoznanMS | 204 | 0.8358 | -0.1044 | 4.346e-01 |
| PROSPER | 315 | 0.8571 | -0.0964 | 3.971e-01 |
| RS1 | 1103 | 0.8422 | 0.0012 | 9.838e-01 |
| RS2 | 1159 | 0.8438 | -0.08 | 1.620e-01 |
| RS3 | 2582 | 0.8465 | -0.0655 | 8.944e-02 |
| RUSH1 | 184 | 0.837 | -0.1565 | 2.688e-01 |
| RUSH2 | 106 | 0.8444 | -0.1815 | 3.406e-01 |
| SHIP | 966 | 0.8452 | 0.0294 | 6.404e-01 |
| SHIP-TREND | 858 | 0.8397 | -0.1292 | 4.985e-02 |
| SYS.adolescents | 980 | 0.8522 | -0.0321 | 6.146e-01 |
| SYS.adults | 594 | 0.8408 | -0.1025 | 1.964e-01 |
| TREND | 1084 | 0.8467 | -0.0979 | 1.009e-01 |
| UKB | 14571 | 0.8475 | -0.0699 | 1.767e-05 |
| WHICAP | 83 | 0.9138 | -0.5263 | 6.073e-02 |
| **Non-European** | | | | |
| ARIC.AA | 675 | 0.9662 | -0.2898 | 5.440e-02 |
| CHAP.black | 321 | 0.9687 | 0.0673 | 7.667e-01 |
| FBIRN | 275 | 0.8788 | 0.1691 | 1.967e-01 |
| NOMAS.black | 166 | 0.9613 | 0.1528 | 5.923e-01 |
| NOMAS.hispanic | 715 | 0.8936 | -0.0491 | 5.672e-01 |
| SCES | 209 | 0.8531 | -0.0109 | 9.373e-01 |
| SIMES | 201 | 0.8779 | 0.2678 | 7.997e-02 |
| WHICAP.black | 60 | 0.9751 | 0.3246 | 5.822e-01 |
| **Summary European MA** | **75304** | **0.8441** | **-0.061** | **1.214e-17** |
| **Summary African MA** | **1222** | **0.9666** | **-0.108** | **0.3379** |
| **Summary Asian MA** | **1298** | **0.8461** | **-0.011** | **0.8444** |
| **Summary Transancestral MA** | **78928** | **0.8466** | **-0.059** | **2.173e-17** |

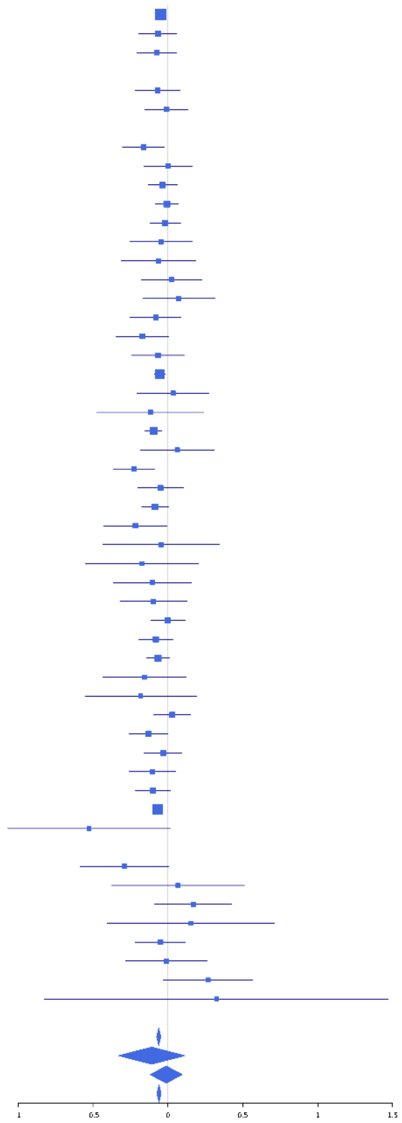

**rs56370391 (A); chr3:142084647; model 1 (I2=0, HetP=0.617)**

| Study | N_total | EAF | Beta | P |
|---|---|---|---|---|
| **Head circumference** | | | | |
| Haworth | 18881 | 0.8664 | -0.02 | 1.847e-01 |
| HKOS | 888 | 0.9302 | -0.0648 | 4.863e-01 |
| VIDI | 745 | 0.8584 | -0.0154 | 8.364e-01 |
| **CT-based** | | | | |
| ESS | 641 | 0.8686 | -0.0685 | 4.071e-01 |
| UCL | 642 | 0.8683 | -0.1108 | 1.797e-01 |
| **MR-based** | | | | |
| 1000Brains | 751 | 0.8638 | -0.219 | 3.706e-03 |
| 3C-Bordeaux | 564 | 0.871 | -0.018 | 8.391e-01 |
| 3c-Dijon | 1499 | 0.8692 | -0.0701 | 1.958e-01 |
| AGES | 2226 | 0.8788 | 1e-04 | 9.985e-01 |
| ARIC.EA | 1390 | 0.8619 | 0.0213 | 6.988e-01 |
| ASPS-Family | 338 | 0.6072 | -0.015 | 8.489e-01 |
| ASRB | 233 | 0.8848 | 0.1196 | 4.109e-01 |
| CARDIA.EA | 349 | 0.8521 | -0.0992 | 3.521e-01 |
| CHAP | 261 | 0.8501 | -0.1187 | 3.338e-01 |
| CHS | 544 | 0.8641 | 0.0073 | 9.338e-01 |
| DHS | 460 | 0.8634 | -0.0217 | 8.209e-01 |
| DNGS | 516 | 0.8741 | -0.2356 | 1.234e-02 |
| ENIGMA | 11373 | 0.86 | -0.0267 | 1.625e-01 |
| EPOZ | 251 | 0.8899 | 0.1093 | 4.428e-01 |
| ERF | 122 | 0.848 | 0.0471 | 7.919e-01 |
| FHS | 4710 | 0.8649 | -0.0663 | 2.784e-02 |
| GenR | 265 | 0.8697 | -0.0991 | 4.426e-01 |
| HUNT | 851 | 0.8616 | -0.0776 | 2.686e-01 |
| LBC1936 | 577 | 0.8622 | -0.0822 | 3.361e-01 |
| LIFE | 1825 | 0.8583 | -0.0553 | 2.441e-01 |
| LLS | 355 | 0.886 | -0.1016 | 3.900e-01 |
| MethCT. | 114 | 0.8502 | 0.2067 | 2.652e-01 |
| NOMAS | 141 | 0.899 | 0.2079 | 2.947e-01 |
| PoznanMS | 204 | 0.823 | 0.0472 | 7.161e-01 |
| PROSPER | 315 | 0.8747 | -0.1729 | 1.508e-01 |
| RS1 | 1103 | 0.8644 | 0.0348 | 5.759e-01 |
| RS2 | 1159 | 0.8643 | -0.0687 | 2.578e-01 |
| RS3 | 2582 | 0.8602 | -0.0051 | 8.997e-01 |
| RUSH1 | 184 | 0.8449 | -0.0671 | 6.419e-01 |
| RUSH2 | 106 | 0.8774 | 0.0094 | 9.638e-01 |
| SHIP | 966 | 0.8546 | -0.1966 | 2.383e-03 |
| SHIP-TREND | 858 | 0.8654 | -0.0478 | 4.991e-01 |
| SYS.adolescents | 980 | 0.7922 | -0.0046 | 9.342e-01 |
| SYS.adults | 594 | 0.8087 | -0.0887 | 2.288e-01 |
| TREND | 1084 | 0.8573 | 0.0276 | 6.532e-01 |
| UKB | 14571 | 0.862 | -0.0717 | 2.453e-05 |
| WHICAP | 83 | 0.9074 | 0.1457 | 5.878e-01 |
| **Non-European** | | | | |
| ARIC.AA | 675 | 0.8606 | -0.0077 | 9.222e-01 |
| CARDIA.AA | 132 | 0.849 | 0.143 | 4.056e-01 |
| CHAP.black | 321 | 0.8424 | -0.1032 | 3.411e-01 |
| FBIRN | 275 | 0.8771 | 0.0895 | 4.915e-01 |
| NOMAS.black | 168 | 0.8255 | 0.0931 | 5.178e-01 |
| NOMAS.hispanic | 718 | 0.8847 | -0.0244 | 7.682e-01 |
| SCES | 209 | 0.9136 | -0.0394 | 8.213e-01 |
| SIMES | 201 | 0.953 | -0.3097 | 1.887e-01 |
| WHICAP.black | 60 | 0.8666 | -0.3514 | 1.963e-01 |
| **Summary European MA** | **75299** | **0.8614** | **-0.042** | **1.96e-08** |
| **Summary African MA** | **1356** | **0.8511** | **-0.017** | **0.7514** |
| **Summary Asian MA** | **1298** | **0.9311** | **-0.092** | **0.2365** |
| **Summary Transancestral MA** | **79060** | **0.8627** | **-0.041** | **1.925e-08** |

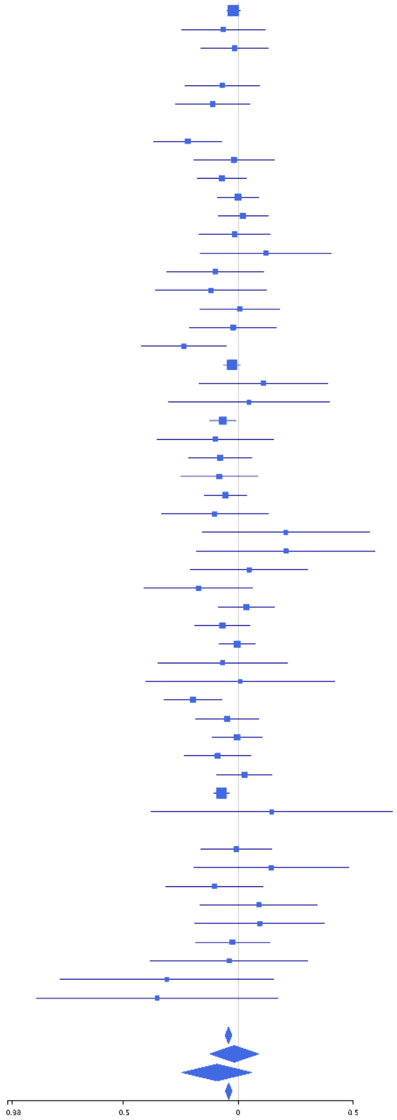

**rs1159211 (T); chr3:190666643; model 1 (I2=0, HetP=0.6776)**

| Study | N_total | EAF | Beta | P |
|---|---|---|---|---|
| **Head circumference** | | | | |
| Haworth | 18881 | 0.3561 | -0.0268 | 1.232e-02 |
| HKOS | 888 | 0.3198 | -0.0469 | 3.571e-01 |
| VIDI | 755 | 0.4092 | -0.0674 | 1.981e-01 |
| **CT-based** | | | | |
| UCL | 642 | 0.3966 | 0.0402 | 4.805e-01 |
| **MR-based** | | | | |
| 1000Brains | 751 | 0.3747 | 0.0019 | 9.713e-01 |
| 3C-Bordeaux | 564 | 0.372 | -0.1341 | 2.993e-02 |
| 3c-Dijon | 1499 | 0.3791 | -0.0707 | 6.033e-02 |
| AGES | 2226 | 0.379 | -0.0806 | 9.103e-03 |
| ARIC.EA | 1390 | 0.3715 | -0.0828 | 3.483e-02 |
| ASPS-Family | 338 | 0.3356 | -0.037 | 6.496e-01 |
| ASRB | 233 | 0.379 | -0.1596 | 9.623e-02 |
| CARDIA.EA | 349 | 0.3613 | -0.1499 | 5.719e-02 |
| CHAP | 261 | 0.3888 | -0.0343 | 7.025e-01 |
| CHS | 544 | 0.382 | -0.1763 | 4.717e-03 |
| DNGS | 515 | 0.3299 | -0.0754 | 2.556e-01 |
| ENIGMA | 10526 | 0.3731 | -0.0608 | 2.007e-05 |
| EPOZ | 251 | 0.3718 | -0.0198 | 8.306e-01 |
| ERF | 122 | 0.5625 | 0.0213 | 8.693e-01 |
| FHS | 4710 | 0.3598 | -0.0141 | 5.117e-01 |
| GenR | 265 | 0.3582 | -0.1306 | 1.492e-01 |
| HUNT | 851 | 0.3317 | -0.0894 | 8.261e-02 |
| LBC1936 | 577 | 0.365 | 0.0526 | 3.892e-01 |
| LIFE | 1825 | 0.3586 | -0.049 | 1.559e-01 |
| LLS | 355 | 0.408 | 0.0017 | 9.828e-01 |
| MethCT. | 114 | 0.3514 | -0.1472 | 2.889e-01 |
| NOMAS | 141 | 0.3143 | 0.0503 | 6.953e-01 |
| PoznanMS | 204 | 0.3531 | -0.0058 | 9.553e-01 |
| PROSPER | 315 | 0.3756 | -0.0649 | 4.300e-01 |
| RS1 | 1103 | 0.3852 | -0.0254 | 5.609e-01 |
| RS2 | 1159 | 0.3764 | -0.0442 | 3.027e-01 |
| RS3 | 2582 | 0.3827 | -0.0439 | 1.249e-01 |
| RUSH1 | 184 | 0.3795 | 0.0141 | 8.956e-01 |
| RUSH2 | 106 | 0.3963 | 0.0579 | 6.810e-01 |
| SHIP | 966 | 0.3554 | 0.0243 | 6.095e-01 |
| SHIP-TREND | 858 | 0.3529 | -0.0711 | 1.598e-01 |
| SYS.adolescents | 980 | 0.3486 | -0.0356 | 4.524e-01 |
| SYS.adults | 594 | 0.3587 | -0.0191 | 7.524e-01 |
| TREND | 1084 | 0.3705 | -0.0016 | 9.709e-01 |
| UKB | 14571 | 0.3829 | -0.0403 | 8.200e-04 |
| WHICAP | 83 | 0.379 | -0.0214 | 8.938e-01 |
| **Non-European** | | | | |
| ARIC.AA | 675 | 0.3455 | 0.0104 | 8.566e-01 |
| CARDIA.AA | 132 | 0.318 | 0.0204 | 8.775e-01 |
| CHAP.black | 321 | 0.2907 | 0.1356 | 1.198e-01 |
| FBIRN | 275 | 0.3003 | -0.0512 | 5.827e-01 |
| NOMAS.black | 168 | 0.2524 | 0.1694 | 1.793e-01 |
| NOMAS.hispanic | 718 | 0.3304 | 0.0387 | 4.907e-01 |
| SCES | 209 | 0.3588 | 0.0171 | 8.663e-01 |
| SIMES | 201 | 0.3675 | -0.0496 | 6.219e-01 |
| WHICAP.black | 60 | 0.3565 | -0.0663 | 7.290e-01 |
| **Summary European MA** | **73360** | **0.3693** | **-0.041** | **3.699e-14** |
| **Summary African MA** | **1356** | **0.3188** | **0.055** | **0.1828** |
| **Summary Asian MA** | **1298** | **0.3335** | **-0.037** | **0.3745** |
| **Summary Transancestral MA** | **77121** | **0.3672** | **-0.039** | **2.303e-13** |

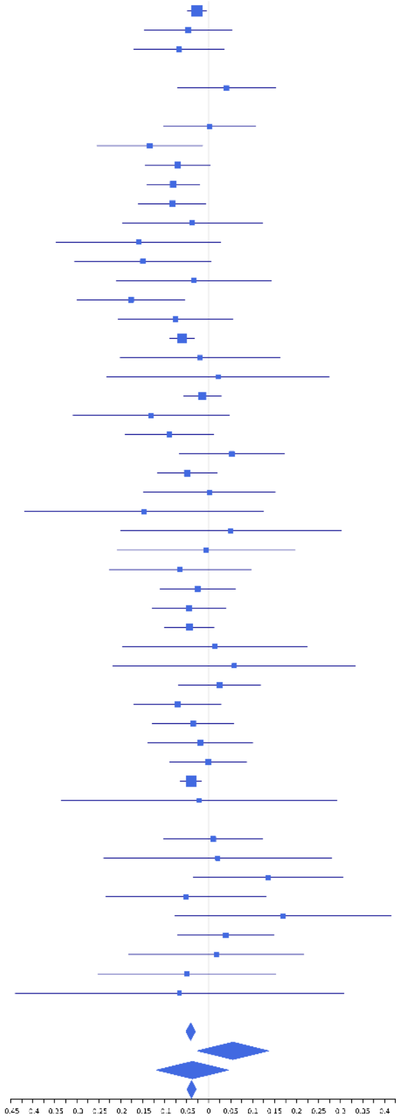

**rs9821713 (C); chr3:190678438; model 1 (I2=0, HetP=0.5005)**

| Study | N_total | EAF | Beta | P |
|---|---|---|---|---|
| **Head circumference** | | | | |
| Haworth | 18881 | 0.0913 | 0.0641 | 3.407e-04 |
| HKOS | 888 | 0.018 | -0.0228 | 8.979e-01 |
| VIDI | 724 | 0.1118 | -0.1178 | 1.580e-01 |
| **CT-based** | | | | |
| ESS | 641 | 0.0842 | -0.0108 | 9.149e-01 |
| UCL | 642 | 0.094 | 0.1305 | 1.723e-01 |
| **MR-based** | | | | |
| 1000Brains | 751 | 0.0818 | -0.072 | 4.447e-01 |
| 3C-Bordeaux | 564 | 0.0732 | 0.1455 | 2.034e-01 |
| 3c-Dijon | 1499 | 0.0762 | 0.154 | 2.528e-02 |
| AGES | 2226 | 0.0999 | 0.063 | 2.074e-01 |
| ARIC.EA | 1390 | 0.0886 | 0.0776 | 2.449e-01 |
| ASPS-Family | 338 | 0.0576 | 0.2527 | 1.261e-01 |
| ASRB | 233 | 0.0919 | -0.1969 | 2.208e-01 |
| CARDIA.EA | 349 | 0.0923 | 0.2098 | 1.086e-01 |
| CHAP | 261 | 0.0824 | 0.1113 | 4.849e-01 |
| ENIGMA | 11373 | 0.0846 | 0.1103 | 3.660e-06 |
| EPOZ | 251 | 0.097 | -0.0917 | 5.431e-01 |
| ERF | 122 | 0.0987 | -0.1934 | 3.676e-01 |
| FHS | 4710 | 0.0795 | 0.0125 | 7.439e-01 |
| GenR | 265 | 0.0689 | 0.2062 | 2.293e-01 |
| HUNT | 851 | 0.1217 | 0.1614 | 2.945e-02 |
| LBC1936 | 577 | 0.0921 | -2e-04 | 9.983e-01 |
| LIFE | 1825 | 0.08 | 0.0694 | 2.556e-01 |
| LLS | 355 | 0.0803 | 0.1407 | 3.084e-01 |
| NOMAS | 141 | 0.0476 | -0.3504 | 2.123e-01 |
| PoznanMS | 204 | 0.0804 | 0.1932 | 2.888e-01 |
| PROSPER | 315 | 0.105 | 0.0706 | 5.874e-01 |
| RS1 | 1103 | 0.0951 | 0.0902 | 2.137e-01 |
| RS2 | 1159 | 0.1004 | 0.0608 | 3.796e-01 |
| RS3 | 2582 | 0.0919 | 0.1257 | 9.015e-03 |
| RUSH1 | 184 | 0.0894 | 0.0901 | 6.228e-01 |
| RUSH2 | 106 | 0.104 | 0.1984 | 3.797e-01 |
| SHIP | 966 | 0.0904 | 0.102 | 1.986e-01 |
| SHIP-TREND | 858 | 0.0908 | 0.1177 | 1.615e-01 |
| SYS.adolescents | 980 | 0.0683 | 0.0877 | 3.274e-01 |
| SYS.adults | 594 | 0.0691 | -0.0024 | 9.829e-01 |
| TREND | 1084 | 0.0874 | 0.0035 | 9.634e-01 |
| UKB | 14571 | 0.0958 | 0.0445 | 2.543e-02 |
| WHICAP | 83 | 0.0317 | 1.0995 | 1.523e-02 |
| **Non-European** | | | | |
| ARIC.AA | 675 | 0.0158 | 0.1342 | 5.382e-01 |
| CHAP.black | 321 | 0.0368 | 0.2369 | 2.594e-01 |
| NOMAS.black | 168 | 0.021 | 0.1674 | 6.606e-01 |
| NOMAS.hispanic | 718 | 0.0437 | -0.1053 | 4.150e-01 |
| SIMES | 201 | 0.0326 | 0.0927 | 7.410e-01 |
| WHICAP.black | 60 | 0.0333 | 0.3556 | 4.875e-01 |
| **Summary European MA** | **73758** | **0.0896** | **0.068** | **1.114e-13** |
| **Summary African MA** | **1224** | **0.0229** | **0.183** | **0.1767** |
| **Summary Asian MA** | **1089** | **0.0207** | **0.004** | **0.9792** |
| **Summary Transancestral MA** | **76789** | **0.0871** | **0.067** | **1.662e-13** |

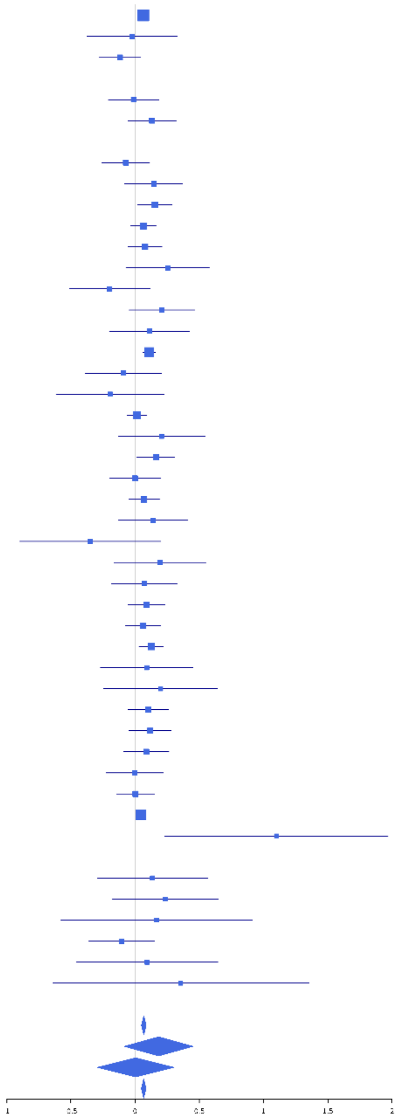

**rs362307 (T); chr4:3241845; model 1 (I2=20.2, HetP=0.1341)**

| Study | N_total | EAF | Beta | P |
|---|---|---|---|---|
| **Head circumference** | | | | |
| Haworth | 18881 | 0.0794 | -0.0443 | 1.969e-02 |
| HKOS | 888 | 0.0169 | 0.1766 | 3.374e-01 |
| VIDI | 749 | 0.0607 | -0.2444 | 2.426e-02 |
| **CT-based** | | | | |
| ESS | 641 | 0.073 | -0.2331 | 2.995e-02 |
| UCL | 642 | 0.0682 | -0.0021 | 9.851e-01 |
| **MR-based** | | | | |
| 1000Brains | 751 | 0.0652 | -0.1444 | 1.673e-01 |
| 3C-Bordeaux | 564 | 0.0612 | 0.0237 | 8.483e-01 |
| 3c-Dijon | 1499 | 0.0679 | -0.1234 | 8.935e-02 |
| AGES | 2226 | 0.0972 | -0.0729 | 1.498e-01 |
| ARIC.EA | 1390 | 0.078 | -0.0292 | 6.794e-01 |
| ASPS-Family | 338 | 0.0632 | -0.1205 | 4.459e-01 |
| CARDIA.EA | 349 | 0.0939 | 0.0112 | 9.317e-01 |
| CHS | 544 | 0.0694 | 0.124 | 2.985e-01 |
| ENIGMA | 11373 | 0.0829 | -0.0333 | 1.656e-01 |
| EPOZ | 251 | 0.0594 | 0.2829 | 1.338e-01 |
| ERF | 122 | 0.1011 | -0.2572 | 2.257e-01 |
| FHS | 4710 | 0.0771 | -0.0194 | 6.151e-01 |
| GenR | 265 | 0.0556 | -0.1122 | 5.541e-01 |
| HUNT | 851 | 0.0886 | -0.2564 | 2.651e-03 |
| LBC1936 | 577 | 0.0869 | -0.0392 | 7.073e-01 |
| LIFE | 1825 | 0.0897 | 0.029 | 6.166e-01 |
| LLS | 355 | 0.0652 | -0.2343 | 1.230e-01 |
| NOMAS | 141 | 0.0752 | -0.1951 | 3.889e-01 |
| PoznanMS | 204 | 0.09 | -0.1161 | 5.021e-01 |
| PROSPER | 315 | 0.0749 | 0.2426 | 1.090e-01 |
| RS1 | 1103 | 0.0679 | -0.1915 | 2.364e-02 |
| RS2 | 1159 | 0.0657 | -0.058 | 4.892e-01 |
| RS3 | 2582 | 0.072 | -0.0896 | 9.601e-02 |
| SHIP | 966 | 0.0861 | 0.0293 | 7.178e-01 |
| SHIP-TREND | 858 | 0.0897 | -0.0334 | 6.920e-01 |
| SYS.adolescents | 980 | 0.0957 | -0.1418 | 6.476e-02 |
| SYS.adults | 594 | 0.0975 | -0.0995 | 3.090e-01 |
| TREND | 1084 | 0.0863 | -0.1632 | 3.308e-02 |
| UKB | 14571 | 0.0729 | -0.0812 | 3.135e-04 |
| WHICAP | 83 | 0.0278 | -0.5066 | 2.867e-01 |
| **Non-European** | | | | |
| ARIC.AA | 675 | 0.0232 | 0.194 | 2.831e-01 |
| FBIRN | 275 | 0.0597 | -0.1265 | 4.827e-01 |
| NOMAS.black | 168 | 0.0214 | -0.4637 | 2.205e-01 |
| NOMAS.hispanic | 718 | 0.0613 | 0.0932 | 3.971e-01 |
| SIMES | 201 | 0.0293 | 0.1417 | 5.821e-01 |
| **Summary European MA** | **73543** | **0.0783** | **-0.06** | **7.954e-10** |
| **Summary African MA** | **843** | **0.0228** | **0.068** | **0.6792** |
| **Summary Asian MA** | **1089** | **0.0192** | **0.172** | **0.2701** |
| **Summary Transancestral MA** | **76468** | **0.0766** | **-0.056** | **6.167e-09** |

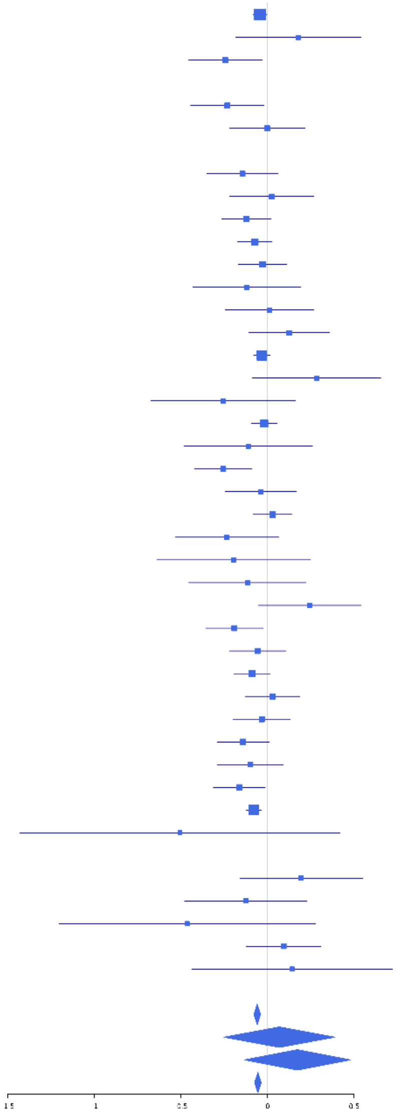

**rs2301718 (A); chr4:106009763; model 1 (I2=21.7, HetP=0.0877)**

| Study | N_total | EAF | Beta | P |
|---|---|---|---|---|
| **Head circumference** | | | | |
| Haworth | 18881 | 0.2179 | 0.0396 | 1.527e-03 |
| HKOS | 888 | 0.7376 | 0.0209 | 6.988e-01 |
| VIDI | 755 | 0.3775 | 0.0696 | 1.901e-01 |
| **CT-based** | | | | |
| ESS | 641 | 0.2435 | -0.0846 | 1.936e-01 |
| UCL | 642 | 0.2179 | 0.0312 | 6.444e-01 |
| **MR-based** | | | | |
| 1000Brains | 751 | 0.2546 | 0.0619 | 2.963e-01 |
| 3C-Bordeaux | 564 | 0.202 | 0.0441 | 5.525e-01 |
| 3c-Dijon | 1499 | 0.2049 | 0.0336 | 4.581e-01 |
| AGES | 2226 | 0.212 | 0.0051 | 8.897e-01 |
| ARIC.EA | 1390 | 0.2268 | 0.0339 | 4.548e-01 |
| ASPS-Family | 338 | 0.1603 | 0.008 | 9.392e-01 |
| ASRB | 233 | 0.2236 | -0.0666 | 5.498e-01 |
| CARDIA.EA | 349 | 0.2295 | 0.0947 | 2.927e-01 |
| CHAP | 261 | 0.2088 | 0.1221 | 2.581e-01 |
| CHS | 544 | 0.2013 | 0.0874 | 2.476e-01 |
| DHS | 460 | 0.2161 | 0.0261 | 7.442e-01 |
| DNGS | 491 | 0.3429 | -0.0294 | 6.625e-01 |
| ENIGMA | 11373 | 0.2291 | 0.0236 | 1.355e-01 |
| EPOZ | 251 | 0.2156 | -0.0885 | 4.154e-01 |
| ERF | 122 | 0.1728 | 0.0962 | 5.703e-01 |
| FHS | 4710 | 0.2279 | 0.0538 | 2.852e-02 |
| GenR | 265 | 0.2316 | -0.0466 | 6.502e-01 |
| HUNT | 851 | 0.2278 | 0.0913 | 1.143e-01 |
| LBC1936 | 577 | 0.223 | 0.1523 | 3.123e-02 |
| LIFE | 1825 | 0.2715 | -0.0204 | 5.844e-01 |
| LLS | 355 | 0.2366 | -0.1566 | 7.624e-02 |
| MethCT. | 114 | 0.2139 | 0.1344 | 4.054e-01 |
| NOMAS | 141 | 0.2607 | 0.438 | 1.558e-03 |
| PoznanMS | 204 | 0.2768 | 0.06 | 5.881e-01 |
| PROSPER | 315 | 0.254 | -0.0214 | 8.153e-01 |
| RS1 | 1103 | 0.2297 | -0.1132 | 2.526e-02 |
| RS2 | 1159 | 0.2254 | 0.1039 | 3.659e-02 |
| RS3 | 2582 | 0.2285 | 0.0567 | 8.690e-02 |
| RUSH1 | 184 | 0.231 | 0.0329 | 7.903e-01 |
| RUSH2 | 106 | 0.2028 | 0.1208 | 4.813e-01 |
| SHIP | 966 | 0.2723 | 0.0295 | 5.639e-01 |
| SHIP-TREND | 858 | 0.2721 | 0.037 | 4.953e-01 |
| SYS.adolescents | 980 | 0.2439 | 0.0958 | 6.848e-02 |
| SYS.adults | 594 | 0.2475 | 0.157 | 1.955e-02 |
| TREND | 1084 | 0.2634 | 0.1136 | 2.003e-02 |
| UKB | 14571 | 0.2092 | 0.0503 | 4.742e-04 |
| WHICAP | 83 | 0.1914 | 0.0122 | 9.503e-01 |
| **Non-European** | | | | |
| ARIC.AA | 675 | 0.0404 | 0.0122 | 9.302e-01 |
| CARDIA.AA | 132 | 0.048 | -0.0821 | 7.756e-01 |
| CHAP.black | 321 | 0.0592 | 0.2818 | 9.292e-02 |
| FBIRN | 275 | 0.2628 | -0.0365 | 7.066e-01 |
| IMH | 37 | 0.6579 | -0.356 | 1.462e-01 |
| NOMAS.black | 168 | 0.0387 | -0.0368 | 8.970e-01 |
| NOMAS.hispanic | 717 | 0.1772 | 0.131 | 5.864e-02 |
| SCES | 209 | 0.7201 | 0.016 | 8.832e-01 |
| SIMES | 201 | 0.5672 | 0.1647 | 1.029e-01 |
| WHICAP.black | 60 | 0.0917 | -0.7398 | 2.312e-02 |
| **Summary European MA** | **75284** | **0.2259** | **0.039** | **2.627e-10** |
| **Summary African MA** | **1356** | **0.0476** | **0.024** | **0.7895** |
| **Summary Asian MA** | **1335** | **0.707** | **0.033** | **0.4445** |
| **Summary Transancestral MA** | **79081** | **0.2306** | **0.039** | **8.611e-11** |

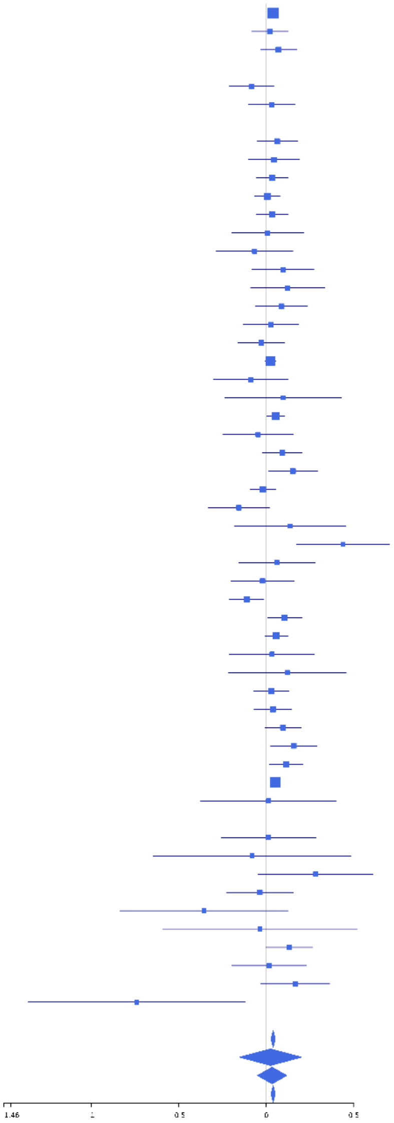

**rs56335290 (A); chr5:112036634; model 1 (I2=0, HetP=0.4918)**

| Study | N_total | EAF | Beta | P |
|---|---|---|---|---|
| **Head circumference** | | | | |
| Haworth | 18881 | 0.2314 | 0.0387 | 1.463e-03 |
| HKOS | 888 | 0.165 | -0.0366 | 5.667e-01 |
| VIDI | 726 | 0.2969 | 0.0229 | 6.909e-01 |
| **CT-based** | | | | |
| ESS | 641 | 0.2189 | -0.048 | 4.778e-01 |
| UCL | 642 | 0.2226 | 0.0332 | 6.203e-01 |
| **MR-based** | | | | |
| 1000Brains | 751 | 0.2477 | 0.0144 | 8.099e-01 |
| 3C-Bordeaux | 564 | 0.2098 | 0.0396 | 5.880e-01 |
| 3c-Dijon | 1499 | 0.2124 | 0.0053 | 9.061e-01 |
| AGES | 2226 | 0.2056 | 0.0576 | 1.205e-01 |
| ARIC.EA | 1390 | 0.2211 | 0.0542 | 2.351e-01 |
| ASPS-Family | 338 | 0.187 | 0.1866 | 5.844e-02 |
| ASRB | 233 | 0.2348 | -0.0425 | 6.976e-01 |
| CARDIA.EA | 349 | 0.2251 | 0.1754 | 5.303e-02 |
| CHAP | 261 | 0.2154 | -0.0859 | 4.202e-01 |
| CHS | 544 | 0.2142 | 0.0566 | 4.434e-01 |
| ENIGMA | 11373 | 0.2237 | 0.0323 | 4.204e-02 |
| EPOZ | 251 | 0.2114 | -0.1929 | 7.753e-02 |
| ERF | 122 | 0.2075 | 0.0751 | 6.342e-01 |
| FHS | 4710 | 0.2128 | 0.0788 | 1.742e-03 |
| GenR | 265 | 0.1932 | 0.198 | 7.192e-02 |
| HUNT | 851 | 0.2206 | 0.046 | 4.311e-01 |
| LBC1936 | 577 | 0.2193 | 0.0388 | 5.847e-01 |
| LIFE | 1825 | 0.2446 | -0.0232 | 5.473e-01 |
| LLS | 355 | 0.2374 | 0.1903 | 3.092e-02 |
| MethCT. | 114 | 0.1436 | 0.1922 | 3.086e-01 |
| NOMAS | 141 | 0.2136 | 0.009 | 9.510e-01 |
| PoznanMS | 204 | 0.2608 | -0.0581 | 6.067e-01 |
| PROSPER | 315 | 0.2423 | -0.0584 | 5.301e-01 |
| RS1 | 1103 | 0.2061 | 0.0497 | 3.448e-01 |
| RS2 | 1159 | 0.2143 | 0.0792 | 1.177e-01 |
| RS3 | 2582 | 0.2104 | 0.0717 | 3.562e-02 |
| RUSH2 | 106 | 0.2161 | 0.2915 | 8.368e-02 |
| SHIP | 966 | 0.2485 | 0.0307 | 5.598e-01 |
| SHIP-TREND | 858 | 0.2286 | 0.0048 | 9.330e-01 |
| SYS.adolescents | 980 | 0.1919 | 0.0353 | 5.382e-01 |
| SYS.adults | 594 | 0.2061 | 0.0078 | 9.131e-01 |
| TREND | 1084 | 0.2511 | -0.0554 | 2.632e-01 |
| UKB | 14571 | 0.2243 | 0.0209 | 1.372e-01 |
| WHICAP | 83 | 0.2155 | 0.0225 | 9.058e-01 |
| **Non-European** | | | | |
| ARIC.AA | 675 | 0.0555 | -0.1721 | 1.477e-01 |
| CHAP.black | 321 | 0.0835 | 0.0544 | 7.037e-01 |
| FBIRN | 275 | 0.1843 | 0.0015 | 9.890e-01 |
| NOMAS.black | 168 | 0.0652 | 0.0197 | 9.289e-01 |
| NOMAS.hispanic | 718 | 0.1364 | 0.1526 | 4.753e-02 |
| SCES | 209 | 0.2111 | -0.0988 | 4.097e-01 |
| SIMES | 201 | 0.1871 | 0.051 | 6.885e-01 |
| WHICAP.black | 60 | 0.0328 | -0.5238 | 3.112e-01 |
| **Summary European MA** | **74120** | **0.2247** | **0.035** | **2.318e-08** |
| **Summary African MA** | **1224** | **0.0631** | **-0.089** | **0.2839** |
| **Summary Asian MA** | **1298** | **0.1758** | **-0.033** | **0.5176** |
| **Summary Transancestral MA** | **77749** | **0.2202** | **0.033** | **4.585e-08** |

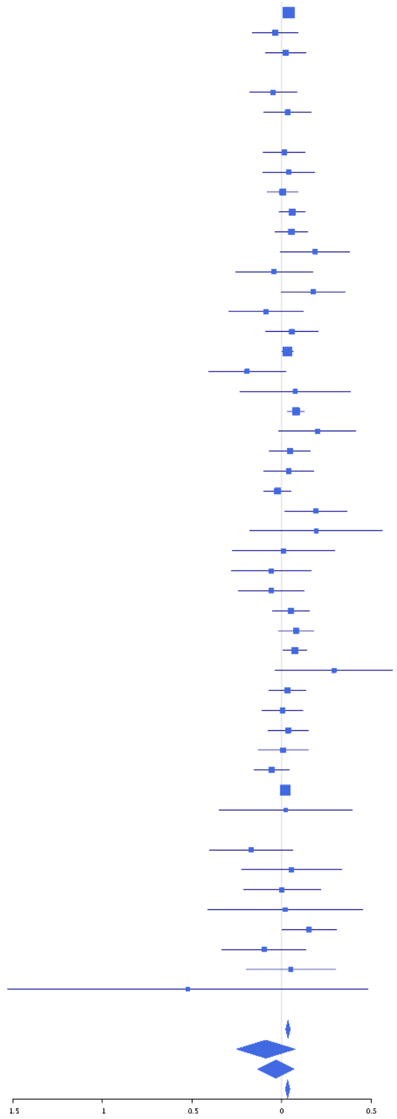

**rs747336 (C); chr6:43832966; model 1 (I2=0, HetP=0.8395)**

| Study | N_total | EAF | Beta | P |
|---|---|---|---|---|
| **Head circumference** | | | | |
| Haworth | 18881 | 0.0442 | 0.0795 | 1.510e-03 |
| HKOS | 888 | 0.0124 | 0.159 | 4.586e-01 |
| VIDI | 748 | 0.0542 | 0.102 | 3.720e-01 |
| **CT-based** | | | | |
| ESS | 641 | 0.0598 | -0.0826 | 4.835e-01 |
| UCL | 642 | 0.0408 | -0.0728 | 6.061e-01 |
| **MR-based** | | | | |
| 1000Brains | 751 | 0.0423 | 0.0734 | 5.672e-01 |
| 3C-Bordeaux | 564 | 0.0391 | 0.0865 | 5.735e-01 |
| 3c-Dijon | 1499 | 0.0443 | 0.1996 | 2.455e-02 |
| AGES | 2226 | 0.0463 | 0.0795 | 2.650e-01 |
| ARIC.EA | 1390 | 0.0464 | 0.1103 | 2.213e-01 |
| ASRB | 233 | 0.0803 | -0.0946 | 5.798e-01 |
| CHAP | 261 | 0.0606 | 0.2058 | 2.631e-01 |
| CHS | 544 | 0.0615 | 0.156 | 2.160e-01 |
| ENIGMA | 8610 | 0.0442 | 0.0558 | 1.327e-01 |
| EPOZ | 251 | 0.0408 | 0.1759 | 4.361e-01 |
| FHS | 4710 | 0.0314 | -0.0077 | 8.960e-01 |
| GenR | 265 | 0.0916 | 0.095 | 5.282e-01 |
| HUNT | 851 | 0.0484 | 0.0907 | 4.221e-01 |
| LBC1936 | 577 | 0.0494 | 0.0951 | 4.842e-01 |
| LIFE | 1825 | 0.0438 | 0.1623 | 4.486e-02 |
| LLS | 355 | 0.0429 | 0.2561 | 1.670e-01 |
| MethCT. | 114 | 0.2034 | -0.0252 | 8.783e-01 |
| PoznanMS | 204 | 0.0568 | -0.025 | 9.069e-01 |
| PROSPER | 315 | 0.041 | 0.0171 | 9.326e-01 |
| RS1 | 1103 | 0.0465 | 0.1587 | 1.167e-01 |
| RS2 | 1159 | 0.0545 | 0.0335 | 7.147e-01 |
| RS3 | 2582 | 0.0511 | 0.1635 | 9.703e-03 |
| RUSH2 | 106 | 0.0377 | -0.0353 | 9.223e-01 |
| SHIP-TREND | 858 | 0.0355 | 0.0297 | 8.195e-01 |
| SYS.adolescents | 980 | 0.0526 | -0.0158 | 8.763e-01 |
| SYS.adults | 594 | 0.0502 | -0.002 | 9.882e-01 |
| TREND | 1084 | 0.045 | 0.0125 | 9.034e-01 |
| UKB | 14571 | 0.0477 | 0.0681 | 1.323e-02 |
| WHICAP | 83 | 0.0384 | -0.553 | 1.750e-01 |
| **Non-European** | | | | |
| ARIC.AA | 675 | 0.3636 | -0.0048 | 9.329e-01 |
| CARDIA.AA | 132 | 0.35 | -0.1241 | 3.361e-01 |
| CHAP.black | 321 | 0.3567 | 0.0054 | 9.474e-01 |
| FBIRN | 275 | 0.1007 | -0.1027 | 4.690e-01 |
| NOMAS.black | 168 | 0.3625 | -0.0918 | 4.198e-01 |
| NOMAS.hispanic | 718 | 0.1588 | -0.0627 | 3.850e-01 |
| SCES | 209 | 0.0205 | -0.6524 | 5.906e-02 |
| SIMES | 201 | 0.0899 | -0.2154 | 2.160e-01 |
| WHICAP.black | 60 | 0.371 | 0.0688 | 7.172e-01 |
| **Summary European MA** | **69463** | **0.0454** | **0.072** | **2.217e-08** |
| **Summary African MA** | **1356** | **0.3608** | **-0.021** | **0.5925** |
| **Summary Asian MA** | **1298** | **0.0257** | **-0.078** | **0.5278** |
| **Summary Transancestral MA** | **73224** | **0.0525** | **0.06** | **2.532e-07** |

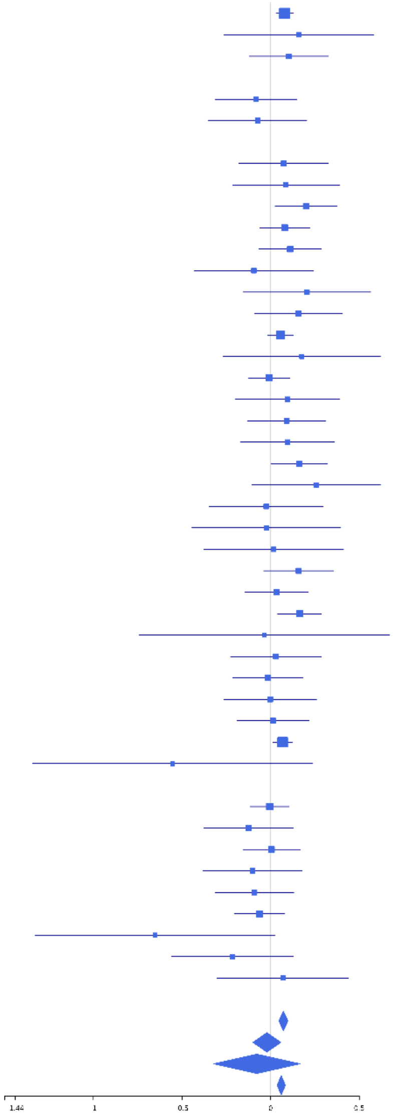

**rs9374035 (A); chr6:108859593; model 1 (I2=20.7, HetP=0.1077)**

| Study | N_total | EAF | Beta | P |
|---|---|---|---|---|
| **Head circumference** | | | | |
| Haworth | 18881 | 0.6536 | 0.0407 | 1.766e-04 |
| HKOS | 888 | 0.6639 | 0.0089 | 8.593e-01 |
| VIDI | 605 | 0.6692 | 0.072 | 2.390e-01 |
| **CT-based** | | | | |
| ESS | 641 | 0.6708 | -0.0159 | 7.886e-01 |
| UCL | 642 | 0.6693 | 0.0958 | 1.063e-01 |
| **MR-based** | | | | |
| 1000Brains | 751 | 0.6682 | -0.0111 | 8.396e-01 |
| 3C-Bordeaux | 564 | 0.6502 | -0.0457 | 4.645e-01 |
| 3c-Dijon | 1499 | 0.6599 | 0.0024 | 9.495e-01 |
| AGES | 2226 | 0.7025 | 5e-04 | 9.869e-01 |
| ARIC.EA | 1390 | 0.6647 | -0.0058 | 8.848e-01 |
| ASPS-Family | 338 | 0.6465 | -0.018 | 8.230e-01 |
| ASRB | 233 | 0.6535 | 0.2516 | 1.037e-02 |
| CARDIA.EA | 349 | 0.6994 | 0.166 | 4.436e-02 |
| CHAP | 261 | 0.6533 | 0.1409 | 1.268e-01 |
| CHS | 544 | 0.6531 | -0.0535 | 4.012e-01 |
| DNGS | 516 | 0.6735 | 1e-04 | 9.992e-01 |
| ENIGMA | 10875 | 0.681 | 0.0587 | 5.386e-05 |
| EPOZ | 251 | 0.6659 | 0.1418 | 1.339e-01 |
| ERF | 122 | 0.7082 | 0.0991 | 4.813e-01 |
| FHS | 4710 | 0.6676 | 0.0472 | 3.094e-02 |
| GenR | 265 | 0.6722 | 0.1057 | 2.536e-01 |
| HUNT | 851 | 0.6596 | -0.0116 | 8.202e-01 |
| LBC1936 | 577 | 0.666 | 0.086 | 1.681e-01 |
| LIFE | 1825 | 0.6532 | 0.0108 | 7.569e-01 |
| LLS | 355 | 0.6775 | 0.0332 | 6.788e-01 |
| MethCT. | 114 | 0.6685 | -0.1068 | 4.479e-01 |
| PoznanMS | 204 | 0.6346 | 0.0694 | 5.000e-01 |
| PROSPER | 315 | 0.6571 | -0.1016 | 2.258e-01 |
| RS1 | 1103 | 0.6701 | -0.0291 | 5.206e-01 |
| RS2 | 1159 | 0.6694 | 0.0527 | 2.326e-01 |
| RS3 | 2582 | 0.6667 | 0.0913 | 1.989e-03 |
| RUSH2 | 106 | 0.6179 | -0.0852 | 5.476e-01 |
| SHIP | 966 | 0.6564 | -0.0776 | 1.057e-01 |
| SHIP-TREND | 858 | 0.6457 | 0.0373 | 4.599e-01 |
| SYS.adolescents | 980 | 0.7005 | 0.0911 | 6.478e-02 |
| SYS.adults | 594 | 0.6835 | 0.0098 | 8.753e-01 |
| TREND | 1084 | 0.646 | 0.0536 | 2.326e-01 |
| UKB | 14571 | 0.6593 | 0.0048 | 6.977e-01 |
| WHICAP | 83 | 0.7284 | 0.2612 | 1.386e-01 |
| **Non-European** | | | | |
| ARIC.AA | 675 | 0.5849 | 0.1566 | 4.569e-03 |
| CARDIA.AA | 132 | 0.583 | 0.2216 | 7.589e-02 |
| CHAP.black | 321 | 0.6495 | 0.0105 | 8.993e-01 |
| IMH | 37 | 0.7105 | -0.0879 | 7.317e-01 |
| NOMAS.black | 168 | 0.6117 | 0.0684 | 5.421e-01 |
| NOMAS.hispanic | 718 | 0.6858 | 0.0629 | 2.689e-01 |
| SCES | 209 | 0.6675 | -0.0118 | 9.094e-01 |
| SIMES | 200 | 0.655 | 0.0114 | 9.143e-01 |
| WHICAP.black | 60 | 0.575 | -0.0464 | 8.031e-01 |
| **Summary European MA** | **73876** | **0.6644** | **0.032** | **5.046e-09** |
| **Summary African MA** | **1356** | **0.6029** | **0.109** | **0.005463** |
| **Summary Asian MA** | **1334** | **0.6644** | **0.003** | **0.9329** |
| **Summary Transancestral MA** | **77398** | **0.6635** | **0.033** | **6.948e-10** |

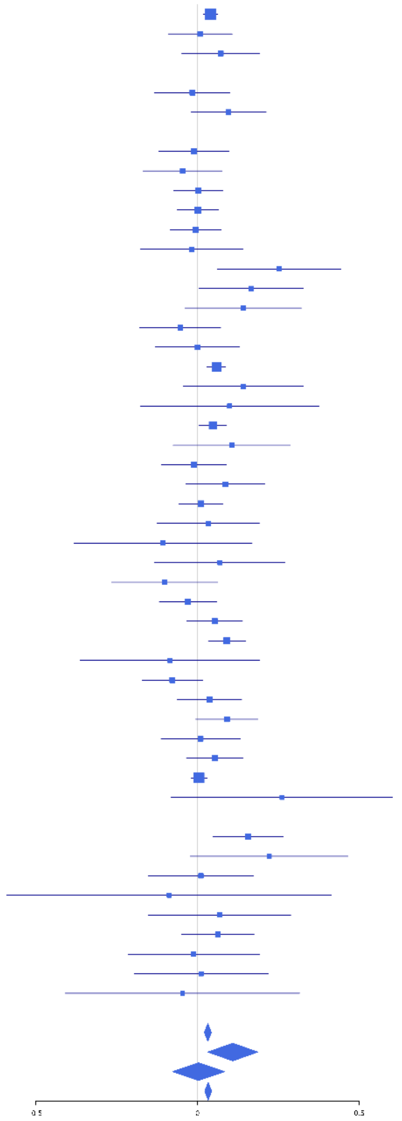

**rs1935952 (C); chr6:108998905; model 1 (I2=35.6, HetP=0.008431)**

| Study | N_total | EAF | Beta | P |
|---|---|---|---|---|
| **Head circumference** | | | | |
| Haworth | 18881 | 0.6915 | 0.0571 | 2.942e-07 |
| HKOS | 888 | 0.6712 | 0.0169 | 7.376e-01 |
| VIDI | 750 | 0.6473 | 0.1043 | 5.380e-02 |
| **CT-based** | | | | |
| ESS | 641 | 0.6756 | 0.2604 | 1.275e-05 |
| UCL | 642 | 0.6936 | 0.0047 | 9.382e-01 |
| **MR-based** | | | | |
| 1000Brains | 751 | 0.7065 | 0.0715 | 2.076e-01 |
| 3C-Bordeaux | 564 | 0.6863 | -0.1367 | 3.360e-02 |
| 3c-Dijon | 1499 | 0.6835 | -0.0202 | 6.068e-01 |
| AGES | 2226 | 0.7343 | -0.0074 | 8.275e-01 |
| ARIC.EA | 1390 | 0.7082 | 0.1 | 1.658e-02 |
| ASPS-Family | 338 | 0.6797 | 0.0911 | 2.691e-01 |
| ASRB | 233 | 0.6894 | -0.0057 | 9.542e-01 |
| CARDIA.EA | 349 | 0.7061 | 0.1106 | 1.833e-01 |
| CHAP | 261 | 0.6872 | 0.1157 | 2.211e-01 |
| CHS | 544 | 0.7292 | 0.0618 | 3.650e-01 |
| ENIGMA | 11373 | 0.7059 | 0.0706 | 1.230e-06 |
| EPOZ | 251 | 0.7436 | 0.0621 | 5.432e-01 |
| ERF | 122 | 0.8151 | 0.1796 | 2.761e-01 |
| FHS | 4710 | 0.6905 | 0.1013 | 5.469e-06 |
| GenR | 265 | 0.6255 | 0.0834 | 3.527e-01 |
| HUNT | 851 | 0.7248 | 0.1002 | 6.476e-02 |
| LBC1936 | 577 | 0.7282 | -0.0836 | 2.061e-01 |
| LIFE | 1825 | 0.694 | 0.0654 | 6.868e-02 |
| LLS | 355 | 0.7348 | 0.0543 | 5.229e-01 |
| MethCT. | 114 | 0.4165 | 0.0841 | 5.311e-01 |
| NOMAS | 141 | 0.6888 | 0.4417 | 7.909e-04 |
| PoznanMS | 204 | 0.671 | 0.1061 | 3.139e-01 |
| PROSPER | 315 | 0.7285 | 0.0347 | 6.985e-01 |
| RS1 | 1103 | 0.7207 | 0.0314 | 5.087e-01 |
| RS2 | 1159 | 0.7202 | 0.1117 | 1.576e-02 |
| RS3 | 2582 | 0.7185 | 0.0779 | 1.177e-02 |
| RUSH1 | 184 | 0.7011 | -0.0966 | 3.976e-01 |
| RUSH2 | 106 | 0.6925 | -0.1898 | 2.054e-01 |
| SHIP | 966 | 0.6957 | 0.0707 | 1.532e-01 |
| SHIP-TREND | 858 | 0.6977 | 0.1479 | 5.004e-03 |
| SYS.adolescents | 980 | 0.6724 | 0.0196 | 6.838e-01 |
| SYS.adults | 594 | 0.6725 | 0.0192 | 7.555e-01 |
| TREND | 1084 | 0.7094 | 0.0732 | 1.221e-01 |
| UKB | 14571 | 0.7101 | 0.068 | 1.396e-07 |
| WHICAP | 83 | 0.6914 | 0.2732 | 1.081e-01 |
| **Non-European** | | | | |
| ARIC.AA | 675 | 0.1576 | 0.0765 | 3.058e-01 |
| CARDIA.AA | 132 | 0.168 | -0.057 | 7.296e-01 |
| CHAP.black | 321 | 0.1854 | -0.0315 | 7.565e-01 |
| FBIRN | 275 | 0.6126 | -0.1025 | 2.424e-01 |
| NOMAS.black | 168 | 0.2195 | 0.1383 | 2.956e-01 |
| NOMAS.hispanic | 718 | 0.5069 | 0.0653 | 2.160e-01 |
| SCES | 209 | 0.6835 | 0.1928 | 6.675e-02 |
| SIMES | 201 | 0.6406 | -0.1269 | 2.230e-01 |
| WHICAP.black | 60 | 0.2417 | 0.149 | 4.878e-01 |
| **Summary European MA** | **74328** | **0.7015** | **0.064** | **7.397e-30** |
| **Summary African MA** | **1356** | **0.1766** | **0.049** | **0.3282** |
| **Summary Asian MA** | **1298** | **0.6684** | **0.022** | **0.5938** |
| **Summary Transancestral MA** | **78089** | **0.6893** | **0.062** | **7.914e-30** |

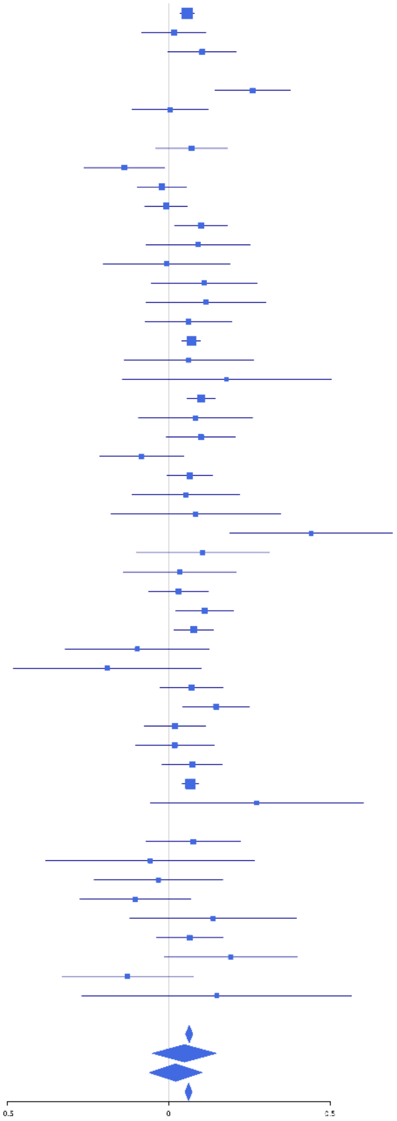

#### rs11154343 (T); chr6:126412953; model 1 (I2=0, HetP=0.9004)

| Study | N_total | EAF | Beta | P |
|---|---|---|---|---|
| **Head circumference** | | | | |
| Haworth | 18881 | 0.3051 | -0.0281 | 1.170e-02 |
| HKOS | 888 | 0.0158 | -0.0831 | 6.626e-01 |
| VIDI | 673 | 0.2372 | -0.0533 | 4.056e-01 |
| **CT-based** | | | | |
| ESS | 641 | 0.2737 | -0.0625 | 3.188e-01 |
| UCL | 642 | 0.3237 | 0.0196 | 7.428e-01 |
| **MR-based** | | | | |
| 1000Brains | 751 | 0.3181 | 0.034 | 5.399e-01 |
| 3C-Bordeaux | 564 | 0.3177 | -0.0059 | 9.256e-01 |
| 3c-Dijon | 1499 | 0.3266 | -0.0451 | 2.465e-01 |
| AGES | 2226 | 0.3456 | -0.0481 | 1.267e-01 |
| ARIC.EA | 1390 | 0.3143 | -0.0619 | 1.299e-01 |
| ASPS-Family | 338 | 0.2766 | -0.0383 | 6.553e-01 |
| ASRB | 233 | 0.3466 | -0.1161 | 2.342e-01 |
| CARDIA.EA | 349 | 0.3474 | -0.0249 | 7.546e-01 |
| CHAP | 261 | 0.3477 | 0.0674 | 4.641e-01 |
| CHS | 544 | 0.3383 | 4e-04 | 9.945e-01 |
| DNGS | 516 | 0.187 | -0.0283 | 7.227e-01 |
| ENIGMA | 11373 | 0.309 | -0.052 | 2.872e-04 |
| EPOZ | 251 | 0.2922 | 0.1126 | 2.515e-01 |
| ERF | 122 | 0.2264 | 0.1704 | 2.652e-01 |
| FHS | 4710 | 0.346 | -0.0346 | 1.106e-01 |
| GenR | 265 | 0.2468 | -0.0841 | 4.034e-01 |
| HUNT | 851 | 0.3403 | -0.0978 | 5.594e-02 |
| LBC1936 | 577 | 0.3242 | -0.0138 | 8.264e-01 |
| LIFE | 1825 | 0.3208 | -0.0615 | 8.333e-02 |
| LLS | 355 | 0.2602 | -0.0514 | 5.476e-01 |
| MethCT. | 114 | 0.1308 | -0.0479 | 8.070e-01 |
| NOMAS | 140 | 0.3393 | -0.1332 | 2.931e-01 |
| PoznanMS | 204 | 0.3385 | 0.0437 | 6.761e-01 |
| PROSPER | 315 | 0.2968 | -0.0849 | 3.307e-01 |
| RS1 | 1103 | 0.3026 | -0.0682 | 1.410e-01 |
| RS2 | 1159 | 0.3128 | -0.0898 | 4.497e-02 |
| RS3 | 2582 | 0.2968 | -0.071 | 1.971e-02 |
| RUSH1 | 184 | 0.2799 | 0.1088 | 3.501e-01 |
| RUSH2 | 106 | 0.3444 | 0.0019 | 9.898e-01 |
| SHIP | 966 | 0.3069 | -0.0485 | 3.261e-01 |
| SHIP-TREND | 858 | 0.313 | 0.0106 | 8.387e-01 |
| SYS.adolescents | 980 | 0.3678 | -0.0551 | 2.394e-01 |
| SYS.adults | 594 | 0.3788 | -0.0279 | 6.408e-01 |
| TREND | 1084 | 0.2992 | -0.0837 | 7.466e-02 |
| UKB | 14571 | 0.3181 | -0.0369 | 3.377e-03 |
| WHICAP | 83 | 0.2987 | -0.1385 | 4.165e-01 |
| **Non-European** | | | | |
| ARIC.AA | 675 | 0.065 | 0.1595 | 1.485e-01 |
| CARDIA.AA | 132 | 0.059 | -0.0797 | 7.600e-01 |
| CHAP.black | 321 | 0.0545 | -0.1107 | 5.249e-01 |
| FBIRN | 275 | 0.2065 | -0.0315 | 7.656e-01 |
| NOMAS.black | 163 | 0.0774 | 0.0694 | 7.383e-01 |
| NOMAS.hispanic | 715 | 0.1807 | 0.0369 | 5.916e-01 |
| SIMES | 201 | 0.0398 | -0.2954 | 2.478e-01 |
| WHICAP.black | 60 | 0.1083 | 0.7059 | 1.979e-02 |
| **Summary European MA** | **74766** | **0.3136** | **-0.039** | **1.402e-12** |
| **Summary African MA** | **1351** | **0.0653** | **0.095** | **0.2213** |
| **Summary Asian MA** | **1089** | **0.0202** | **-0.136** | **0.3731** |
| **Summary Transancestral MA** | **78310** | **0.3034** | **-0.038** | **7.822e-12** |

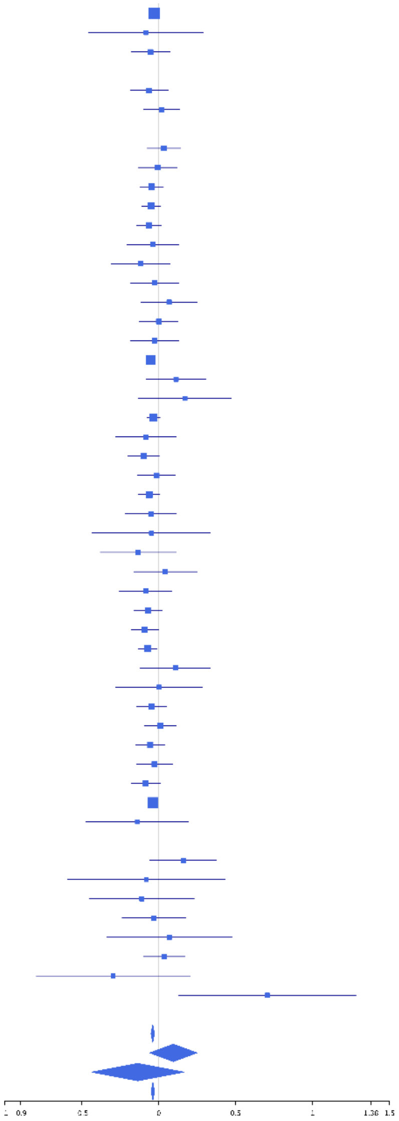

**rs9401873 (T); chr6:126623120; model 1 (I2=11.2, HetP=0.254)**

| Study | N_total | EAF | Beta | P |
|---|---|---|---|---|
| **Head circumference** | | | | |
| Haworth | 18881 | 0.6341 | -0.012 | 2.610e-01 |
| HKOS | 888 | 0.0175 | -0.1471 | 4.168e-01 |
| VIDI | 755 | 0.5913 | -0.0731 | 1.629e-01 |
| **CT-based** | | | | |
| ESS | 641 | 0.6271 | 0.018 | 7.549e-01 |
| UCL | 642 | 0.6732 | 0.0981 | 9.934e-02 |
| **MR-based** | | | | |
| 1000Brains | 751 | 0.6608 | 0.0463 | 3.955e-01 |
| 3C-Bordeaux | 564 | 0.6568 | -0.0485 | 4.398e-01 |
| 3c-Dijon | 1499 | 0.6429 | -6e-04 | 9.882e-01 |
| AGES | 2226 | 0.6436 | -0.0047 | 8.802e-01 |
| ARIC.EA | 1390 | 0.6517 | 0.0221 | 5.780e-01 |
| ASPS-Family | 338 | 0.6386 | 0.0281 | 7.256e-01 |
| ASRB | 233 | 0.6598 | 0.0563 | 5.651e-01 |
| CARDIA.EA | 349 | 0.6546 | -0.0182 | 8.187e-01 |
| CHAP | 261 | 0.6798 | -0.1426 | 1.296e-01 |
| CHS | 544 | 0.6552 | 0.1428 | 2.515e-02 |
| ENIGMA | 11373 | 0.6454 | -0.0463 | 8.286e-04 |
| EPOZ | 251 | 0.6746 | 0.0374 | 6.944e-01 |
| ERF | 122 | 0.6233 | -0.1344 | 3.094e-01 |
| FHS | 4710 | 0.6502 | -0.0873 | 5.313e-05 |
| GenR | 265 | 0.5904 | 0.0188 | 8.310e-01 |
| HUNT | 851 | 0.6385 | -0.0564 | 2.641e-01 |
| LBC1936 | 577 | 0.6239 | -0.058 | 3.395e-01 |
| LIFE | 1825 | 0.657 | -0.0497 | 1.546e-01 |
| LLS | 355 | 0.6708 | 0.0743 | 3.521e-01 |
| MethCT. | 114 | 0.498 | 0.004 | 9.759e-01 |
| NOMAS | 141 | 0.6036 | -0.0555 | 6.492e-01 |
| PoznanMS | 204 | 0.6876 | 0.0157 | 8.834e-01 |
| PROSPER | 315 | 0.6675 | -0.109 | 1.975e-01 |
| RS1 | 1103 | 0.6648 | -0.0654 | 1.473e-01 |
| RS2 | 1159 | 0.6645 | 0.0163 | 7.114e-01 |
| RS3 | 2582 | 0.6557 | -0.02 | 4.948e-01 |
| RUSH1 | 184 | 0.6766 | -0.1328 | 2.350e-01 |
| RUSH2 | 106 | 0.638 | -0.1515 | 2.916e-01 |
| SHIP | 966 | 0.6703 | -0.129 | 7.818e-03 |
| SHIP-TREND | 858 | 0.6619 | -0.0243 | 6.335e-01 |
| SYS.adolescents | 980 | 0.6793 | 0.0193 | 6.905e-01 |
| SYS.adults | 594 | 0.6889 | 0.0142 | 8.210e-01 |
| TREND | 1084 | 0.6734 | -0.032 | 4.856e-01 |
| UKB | 14571 | 0.6558 | -0.0509 | 3.684e-05 |
| WHICAP | 83 | 0.6274 | -0.0817 | 6.125e-01 |
| **Non-European** | | | | |
| ARIC.AA | 675 | 0.569 | -0.0496 | 3.668e-01 |
| CARDIA.AA | 132 | 0.576 | -0.1716 | 1.681e-01 |
| CHAP.black | 321 | 0.5748 | -0.0038 | 9.629e-01 |
| FBIRN | 275 | 0.5034 | 0.0324 | 7.044e-01 |
| NOMAS.black | 167 | 0.5684 | -0.0542 | 6.244e-01 |
| NOMAS.hispanic | 718 | 0.528 | -0.0507 | 3.374e-01 |
| SCES | 209 | 0.03 | -0.0275 | 9.234e-01 |
| SIMES | 201 | 0.1527 | -0.2435 | 8.028e-02 |
| WHICAP.black | 60 | 0.6229 | 0.036 | 8.494e-01 |
| **Summary European MA** | **74333** | **0.6478** | **-0.031** | **1.354e-08** |
| **Summary African MA** | **1355** | **0.5734** | **-0.047** | **0.2218** |
| **Summary Asian MA** | **1298** | **0.0404** | **-0.139** | **0.1619** |
| **Summary Transancestral MA** | **78093** | **0.6346** | **-0.031** | **2.678e-09** |

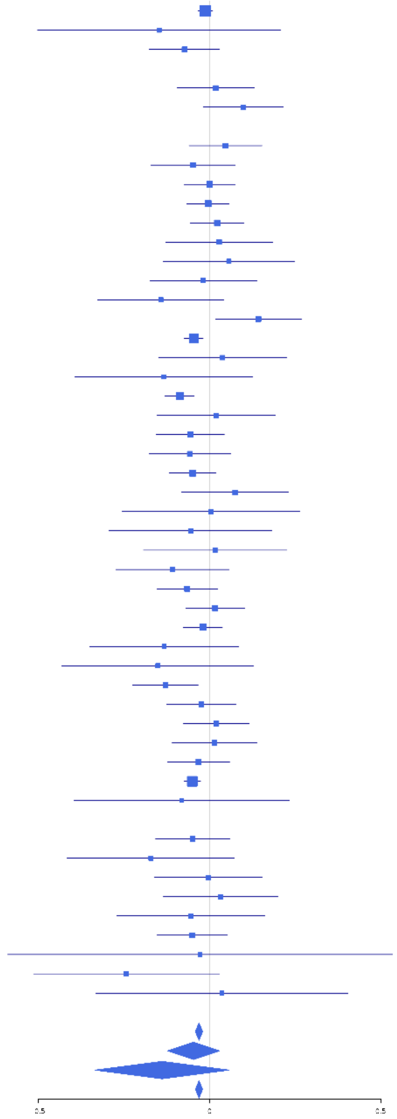

**6-126845438 (A); chr6:126845438; model 1 (I2=29.7, HetP=0.1265)**

| Study | N_total | EAF | Beta | P |
|---|---|---|---|---|
| **Head circumference** | | | | |
| **CT-based** | | | | |
| **MR-based** | | | | |
| 3c-Dijon | 1499 | 0.0846 | -0.1077 | 1.008e-01 |
| AGES | 2226 | 0.0843 | -0.0479 | 3.752e-01 |
| CHS | 544 | 0.0829 | -0.3178 | 3.857e-03 |
| ENIGMA | 7204 | 0.0847 | -0.134 | 7.482e-06 |
| FHS | 4710 | 0.0868 | -0.109 | 2.889e-03 |
| HUNT | 851 | 0.0802 | -0.1182 | 1.856e-01 |
| LBC1936 | 577 | 0.1046 | 0.0306 | 7.502e-01 |
| LIFE | 1825 | 0.0909 | -0.0287 | 6.177e-01 |
| LLS | 355 | 0.0808 | 0.0325 | 8.131e-01 |
| NOMAS | 141 | 0.0602 | -0.3019 | 2.298e-01 |
| PROSPER | 315 | 0.0791 | -0.2349 | 1.117e-01 |
| SHIP | 966 | 0.0985 | -0.0509 | 5.051e-01 |
| SHIP-TREND | 858 | 0.0854 | 0.1134 | 1.897e-01 |
| WHICAP | 83 | 0.0618 | 0.1679 | 6.042e-01 |
| **Non-European** | | | | |
| ARIC.AA | 675 | 0.0159 | 0.1055 | 6.279e-01 |
| FBIRN | 275 | 0.0631 | 0.044 | 8.023e-01 |
| **Summary European MA** | **22154** | **0.0861** | **-0.094** | **2.939e-08** |
| **Summary African MA** | **675** | **0.0159** | **0.106** | **0.6279** |
| **Summary Transancestral MA** | **23104** | **0.0838** | **-0.089** | **1.039e-07** |

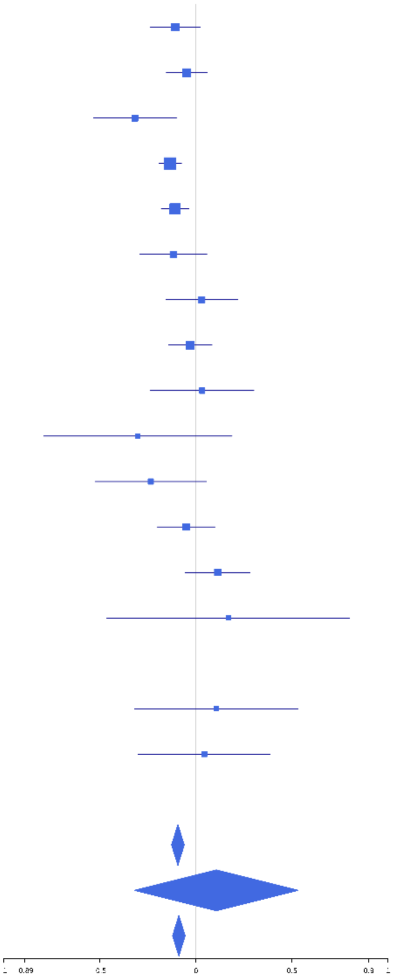

**rs4273712 (A); chr6:126964510; model 1 (I2=26.4, HetP=0.04646)**

| Study | N_total | EAF | Beta | P |
|---|---|---|---|---|
| **Head circumference** | | | | |
| Haworth | 18881 | 0.7172 | -0.0557 | 1.035e-06 |
| HKOS | 888 | 0.5298 | -0.0181 | 7.038e-01 |
| VIDI | 755 | 0.7227 | -0.1085 | 5.962e-02 |
| **CT-based** | | | | |
| ESS | 641 | 0.7303 | -0.1493 | 1.766e-02 |
| UCL | 642 | 0.7594 | -0.0954 | 1.440e-01 |
| **MR-based** | | | | |
| 1000Brains | 751 | 0.7397 | -0.0675 | 2.514e-01 |
| 3C-Bordeaux | 564 | 0.7194 | -0.0691 | 2.979e-01 |
| 3c-Dijon | 1499 | 0.7216 | -0.0984 | 1.575e-02 |
| AGES | 2226 | 0.7259 | -0.1207 | 3.274e-04 |
| ARIC.EA | 1390 | 0.7288 | -0.1836 | 1.680e-05 |
| ASPS-Family | 338 | 0.7685 | 2e-04 | 9.988e-01 |
| ASRB | 233 | 0.708 | 0.0908 | 3.741e-01 |
| CARDIA.EA | 349 | 0.7179 | 0.0617 | 4.631e-01 |
| CHAP | 261 | 0.7567 | -0.0659 | 5.190e-01 |
| CHS | 544 | 0.7142 | -0.1253 | 6.187e-02 |
| DNGS | 516 | 0.6689 | -0.0531 | 4.223e-01 |
| ENIGMA | 11373 | 0.7315 | -0.067 | 7.652e-06 |
| EPOZ | 251 | 0.7346 | -0.1442 | 1.535e-01 |
| ERF | 122 | 0.7458 | -0.1782 | 2.255e-01 |
| FHS | 4710 | 0.7295 | -0.116 | 5.739e-07 |
| GenR | 265 | 0.7183 | -0.0406 | 6.742e-01 |
| HUNT | 851 | 0.7228 | -0.0824 | 1.281e-01 |
| LBC1936 | 577 | 0.7501 | -0.0224 | 7.411e-01 |
| LIFE | 1825 | 0.7389 | -0.0867 | 2.148e-02 |
| LLS | 355 | 0.7 | -0.0703 | 3.910e-01 |
| MethCT. | 114 | 0.7922 | 0.3271 | 4.507e-02 |
| NOMAS | 141 | 0.7966 | -0.2484 | 9.555e-02 |
| PoznanMS | 204 | 0.7751 | -0.0977 | 4.098e-01 |
| PROSPER | 315 | 0.7556 | -0.0168 | 8.563e-01 |
| RS1 | 1103 | 0.7313 | -0.0924 | 5.434e-02 |
| RS2 | 1159 | 0.7262 | -0.0976 | 3.608e-02 |
| RS3 | 2582 | 0.7228 | -0.0976 | 1.690e-03 |
| RUSH1 | 184 | 0.7444 | -0.041 | 7.321e-01 |
| RUSH2 | 106 | 0.7594 | -0.2619 | 1.063e-01 |
| SHIP | 966 | 0.7373 | -0.1411 | 6.466e-03 |
| SHIP-TREND | 858 | 0.7488 | -0.0758 | 1.736e-01 |
| SYS.adolescents | 980 | 0.7658 | 0.0223 | 6.752e-01 |
| SYS.adults | 594 | 0.7694 | 0.0276 | 6.887e-01 |
| TREND | 1084 | 0.7399 | -0.1088 | 2.644e-02 |
| UKB | 14571 | 0.7352 | -0.1127 | 2.047e-17 |
| WHICAP | 83 | 0.7284 | 0.0447 | 7.990e-01 |
| **Non-European** | | | | |
| ARIC.AA | 675 | 0.9193 | -0.074 | 4.587e-01 |
| CARDIA.AA | 132 | 0.915 | -0.29 | 1.887e-01 |
| CHAP.black | 321 | 0.9019 | 0.2181 | 1.012e-01 |
| FBIRN | 275 | 0.698 | 0.1015 | 2.751e-01 |
| IMH | 37 | 0.5702 | -0.0077 | 9.735e-01 |
| NOMAS.black | 168 | 0.9055 | 0.2576 | 1.691e-01 |
| NOMAS.hispanic | 718 | 0.7596 | -0.0622 | 3.138e-01 |
| SCES | 209 | 0.5574 | -0.0101 | 9.179e-01 |
| SIMES | 200 | 0.5625 | -0.1247 | 2.170e-01 |
| WHICAP.black | 60 | 0.8917 | 0.1454 | 6.224e-01 |
| **Summary European MA** | **74849** | **0.729** | **-0.083** | **1.268e-46** |
| **Summary African MA** | **1356** | **0.9118** | **0.031** | **0.6511** |
| **Summary Asian MA** | **1334** | **0.5402** | **-0.032** | **0.404** |
| **Summary Transancestral MA** | **78646** | **0.7292** | **-0.079** | **1.739e-44** |

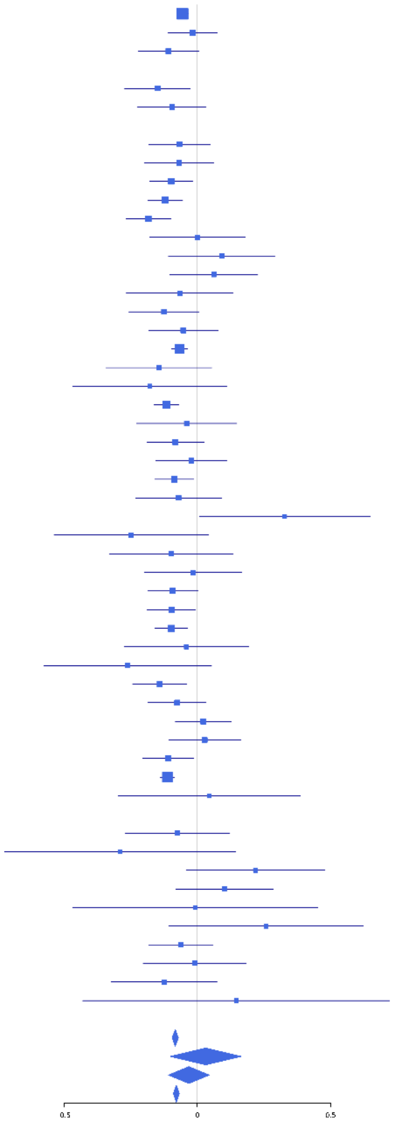

**rs190958130 (C); chr6:127052822; model 1 (I2=0, HetP=0.9153)**

| Study | N_total | EAF | Beta | P |
|---|---|---|---|---|
| **Head circumference** | | | | |
| Haworth | 18881 | 0.9596 | 0.0525 | 4.462e-02 |
| VIDI | 755 | 0.9833 | 0.1534 | 4.450e-01 |
| **CT-based** | | | | |
| ESS | 641 | 0.9636 | -0.0018 | 9.902e-01 |
| UCL | 642 | 0.9514 | 0.1498 | 2.485e-01 |
| **MR-based** | | | | |
| 1000Brains | 751 | 0.968 | -0.0183 | 9.006e-01 |
| 3C-Bordeaux | 564 | 0.9654 | 0.3685 | 2.406e-02 |
| 3c-Dijon | 1499 | 0.9769 | 0.1561 | 1.988e-01 |
| AGES | 2226 | 0.9757 | 0.1668 | 8.678e-02 |
| ARIC.EA | 1390 | 0.9609 | 0.0606 | 5.359e-01 |
| ASPS-Family | 338 | 0.9672 | 0.068 | 7.525e-01 |
| CHS | 544 | 0.9735 | -0.0262 | 8.896e-01 |
| ENIGMA | 9784 | 0.9664 | 0.0826 | 3.732e-02 |
| EPOZ | 251 | 0.9547 | -0.1515 | 4.804e-01 |
| ERF | 122 | 0.9606 | -0.2185 | 5.068e-01 |
| FHS | 4710 | 0.9676 | 0.1083 | 6.261e-02 |
| GenR | 265 | 0.9661 | 0.1548 | 5.187e-01 |
| HUNT | 851 | 0.9521 | 0.1019 | 3.692e-01 |
| LBC1936 | 577 | 0.9799 | 0.0992 | 6.363e-01 |
| LIFE | 1825 | 0.9633 | 0.0378 | 6.681e-01 |
| LLS | 355 | 0.9714 | -0.0126 | 9.550e-01 |
| NOMAS | 141 | 0.9804 | 0.3312 | 4.420e-01 |
| PROSPER | 315 | 0.961 | 0.1938 | 3.464e-01 |
| RS1 | 1103 | 0.9614 | 0.1177 | 2.864e-01 |
| RS2 | 1159 | 0.9598 | 0.2527 | 1.681e-02 |
| RS3 | 2582 | 0.9651 | 0.1491 | 4.929e-02 |
| SHIP | 966 | 0.9664 | 0.0259 | 8.379e-01 |
| SHIP-TREND | 858 | 0.9657 | -0.1244 | 3.483e-01 |
| SYS.adolescents | 980 | 0.9716 | 0.3063 | 2.434e-02 |
| SYS.adults | 594 | 0.971 | -0.1348 | 4.360e-01 |
| TREND | 1084 | 0.9649 | -0.0217 | 8.529e-01 |
| UKB | 14571 | 0.9592 | 0.0817 | 5.762e-03 |
| WHICAP | 83 | 0.9788 | -0.1563 | 7.724e-01 |
| **Non-European** | | | | |
| WHICAP.black | 60 | 0.9853 | 0.7115 | 3.524e-01 |
| **Summary European MA** | **71407** | **0.9634** | **0.08** | **1.59e-08** |
| **Summary African MA** | **60** | **0.9853** | **0.705** | **0.3524** |
| **Summary Transancestral MA** | **71467** | **0.9634** | **0.08** | **1.378e-08** |

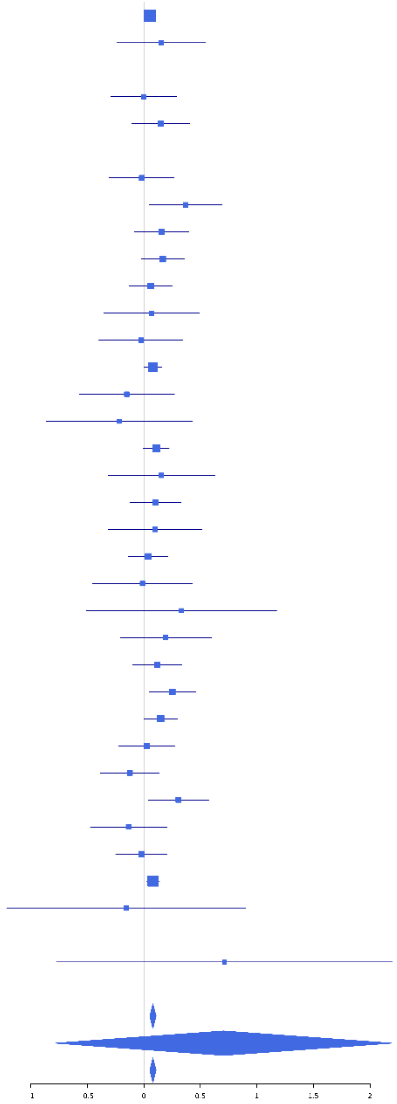

**rs17650496 (A); chr6:127270764; model 1 (I2=0, HetP=0.976)**

| Study | N_total | EAF | Beta | P |
|---|---|---|---|---|
| **Head circumference** | | | | |
| Haworth | 18881 | 0.9317 | -0.0648 | 1.528e-03 |
| VIDI | 747 | 0.9634 | 0.1114 | 4.196e-01 |
| **CT-based** | | | | |
| ESS | 641 | 0.9362 | -0.0957 | 4.027e-01 |
| UCL | 642 | 0.9538 | -0.2585 | 5.183e-02 |
| **MR-based** | | | | |
| 1000Brains | 751 | 0.9611 | -0.0874 | 5.128e-01 |
| 3C-Bordeaux | 564 | 0.9493 | -0.293 | 3.131e-02 |
| AGES | 2226 | 0.9627 | -0.1 | 2.058e-01 |
| ARIC.EA | 1390 | 0.9466 | -0.1352 | 1.092e-01 |
| ASPS-Family | 338 | 0.9641 | -0.1798 | 3.847e-01 |
| ENIGMA | 10797 | 0.941 | -0.1048 | 2.845e-04 |
| EPOZ | 251 | 0.9368 | -0.02 | 9.136e-01 |
| FHS | 4710 | 0.9407 | -0.1073 | 1.389e-02 |
| GenR | 265 | 0.9448 | -0.3131 | 9.982e-02 |
| HUNT | 851 | 0.9554 | -0.1538 | 1.902e-01 |
| LIFE | 1825 | 0.9618 | -0.1035 | 2.311e-01 |
| LLS | 355 | 0.9576 | -0.1881 | 3.126e-01 |
| NOMAS | 141 | 0.8985 | -0.0487 | 8.050e-01 |
| PoznanMS | 204 | 0.9379 | -0.1758 | 3.913e-01 |
| PROSPER | 315 | 0.9674 | -0.327 | 1.451e-01 |
| RS1 | 1103 | 0.9487 | -0.0722 | 4.542e-01 |
| RS2 | 1159 | 0.9461 | -0.156 | 8.989e-02 |
| RS3 | 2582 | 0.9491 | -0.0879 | 1.652e-01 |
| SHIP | 966 | 0.9468 | -0.0848 | 4.034e-01 |
| SHIP-TREND | 858 | 0.9448 | -0.144 | 1.736e-01 |
| SYS.adolescents | 980 | 0.9642 | -0.109 | 3.695e-01 |
| SYS.adults | 594 | 0.9601 | 0.0496 | 7.373e-01 |
| TREND | 1084 | 0.9456 | -0.0157 | 8.682e-01 |
| UKB | 14571 | 0.9458 | -0.1097 | 2.269e-05 |
| WHICAP | 83 | 0.9353 | -0.0334 | 9.160e-01 |
| **Non-European** | | | | |
| ARIC.AA | 675 | 0.9772 | -0.1714 | 3.473e-01 |
| FBIRN | 275 | 0.9369 | 0.1732 | 3.242e-01 |
| NOMAS.hispanic | 718 | 0.8981 | 0.0266 | 7.608e-01 |
| WHICAP.black | 60 | 0.9754 | 0.4491 | 4.492e-01 |
| **Summary European MA** | **69874** | **0.943** | **-0.095** | **2.304e-16** |
| **Summary African MA** | **735** | **0.9771** | **-0.119** | **0.4937** |
| **Summary Transancestral MA** | **71602** | **0.9429** | **-0.092** | **6.307e-16** |

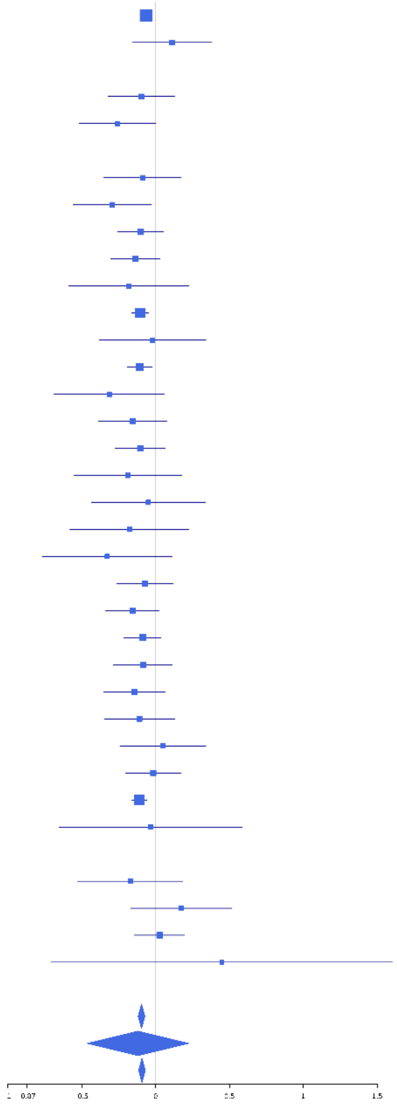

**rs2011008 (C); chr6:127425944; model 1 (I2=2.5, HetP=0.4229)**

| Study | N_total | EAF | Beta | P |
|---|---|---|---|---|
| **Head circumference** | | | | |
| Haworth | 18881 | 0.3102 | 0.0266 | 1.640e-02 |
| HKOS | 888 | 0.5884 | 0.0155 | 7.479e-01 |
| VIDI | 749 | 0.259 | 0.0381 | 5.187e-01 |
| **CT-based** | | | | |
| ESS | 641 | 0.3194 | 0.0503 | 4.010e-01 |
| UCL | 642 | 0.2939 | -0.0023 | 9.705e-01 |
| **MR-based** | | | | |
| 1000Brains | 751 | 0.2809 | -0.1237 | 3.149e-02 |
| 3C-Bordeaux | 564 | 0.292 | 0.0891 | 1.742e-01 |
| 3c-Dijon | 1499 | 0.2847 | 0.1187 | 3.341e-03 |
| AGES | 2226 | 0.3581 | 0.0599 | 5.533e-02 |
| ARIC.EA | 1390 | 0.2946 | 0.0707 | 8.933e-02 |
| ASPS-Family | 338 | 0.2264 | -0.0031 | 9.729e-01 |
| ASRB | 233 | 0.2971 | 0.1291 | 2.040e-01 |
| CARDIA.EA | 349 | 0.3047 | 0.0149 | 8.567e-01 |
| CHAP | 261 | 0.3092 | 0.1867 | 4.979e-02 |
| CHS | 544 | 0.2956 | 0.0565 | 3.951e-01 |
| ENIGMA | 11373 | 0.2934 | 0.0415 | 4.355e-03 |
| EPOZ | 251 | 0.332 | 0.1086 | 2.518e-01 |
| ERF | 122 | 0.2966 | 0.1315 | 3.485e-01 |
| FHS | 4710 | 0.2919 | 0.0233 | 3.051e-01 |
| GenR | 265 | 0.3465 | 0.0067 | 9.420e-01 |
| HUNT | 851 | 0.2773 | 0.0697 | 1.981e-01 |
| LBC1936 | 577 | 0.299 | 0.0213 | 7.403e-01 |
| LIFE | 1825 | 0.2585 | 0.0239 | 5.283e-01 |
| LLS | 355 | 0.3042 | 0.1437 | 7.806e-02 |
| MethCT. | 114 | 0.4697 | -0.1282 | 3.343e-01 |
| NOMAS | 141 | 0.3041 | 0.026 | 8.412e-01 |
| PoznanMS | 204 | 0.2204 | 0.1919 | 1.081e-01 |
| PROSPER | 315 | 0.3167 | 0.0687 | 4.226e-01 |
| RS1 | 1103 | 0.3122 | 0.1195 | 9.300e-03 |
| RS2 | 1159 | 0.3047 | 0.0912 | 4.318e-02 |
| RS3 | 2582 | 0.3017 | 0.0825 | 6.531e-03 |
| RUSH1 | 184 | 0.2761 | 0.118 | 3.129e-01 |
| RUSH2 | 106 | 0.2743 | 0.2294 | 1.395e-01 |
| SHIP | 966 | 0.2757 | 0.0549 | 2.808e-01 |
| SHIP-TREND | 858 | 0.2597 | 0.0176 | 7.494e-01 |
| SYS.adolescents | 980 | 0.3152 | 0.0758 | 1.190e-01 |
| SYS.adults | 594 | 0.3209 | -0.0367 | 5.543e-01 |
| TREND | 1084 | 0.2619 | 0.0013 | 9.792e-01 |
| UKB | 14571 | 0.2958 | 0.0373 | 3.664e-03 |
| WHICAP | 83 | 0.3333 | 0.0734 | 6.568e-01 |
| **Non-European** | | | | |
| ARIC.AA | 675 | 0.4511 | -0.0871 | 1.113e-01 |
| CARDIA.AA | 132 | 0.455 | 0.0278 | 8.222e-01 |
| CHAP.black | 321 | 0.4775 | 0.0948 | 2.309e-01 |
| FBIRN | 275 | 0.3464 | -0.1295 | 1.496e-01 |
| NOMAS.black | 168 | 0.4474 | -0.0757 | 4.912e-01 |
| NOMAS.hispanic | 718 | 0.4222 | 0.0424 | 4.272e-01 |
| SCES | 209 | 0.5771 | -0.0096 | 9.226e-01 |
| SIMES | 201 | 0.4923 | -0.027 | 7.862e-01 |
| WHICAP.black | 60 | 0.4583 | -0.1451 | 4.316e-01 |
| **Summary European MA** | **74327** | **0.2989** | **0.041** | **3.164e-13** |
| **Summary African MA** | **1356** | **0.4576** | **-0.034** | **0.3799** |
| **Summary Asian MA** | **1298** | **0.5717** | **0.005** | **0.9044** |
| **Summary Transancestral MA** | **78088** | **0.3077** | **0.038** | **3.33e-12** |

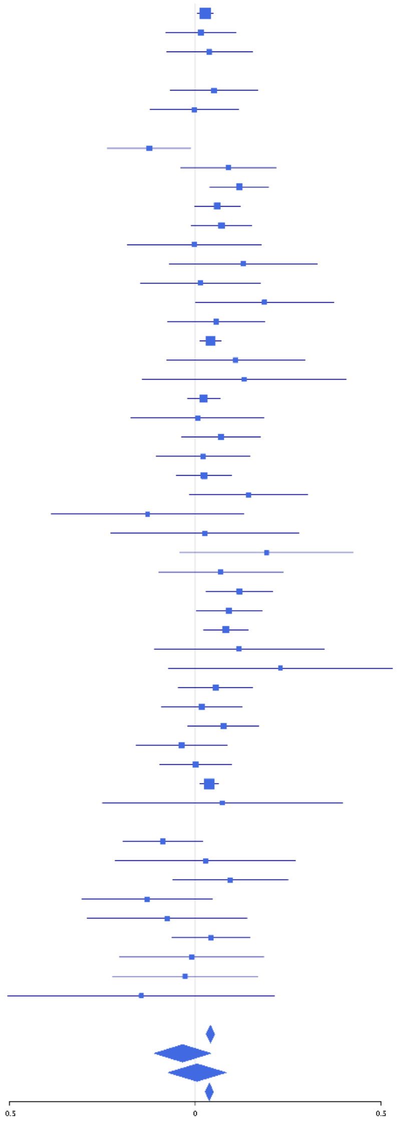

**rs2489624 (A); chr6:127456394; model 1 (I2=18.8, HetP=0.1311)**

| Study | N_total | EAF | Beta | P |
|---|---|---|---|---|
| **Head circumference** | | | | |
| Haworth | 18881 | 0.0333 | 0.0807 | 4.867e-03 |
| HKOS | 888 | 0.0918 | 0.1259 | 1.255e-01 |
| VIDI | 755 | 0.0468 | -0.0869 | 4.763e-01 |
| **CT-based** | | | | |
| ESS | 641 | 0.0333 | 0.1424 | 3.602e-01 |
| UCL | 642 | 0.0321 | 0.4075 | 1.004e-02 |
| **MR-based** | | | | |
| 1000Brains | 751 | 0.0275 | 0.0553 | 7.264e-01 |
| 3C-Bordeaux | 564 | 0.0243 | 0.173 | 3.715e-01 |
| 3c-Dijon | 1499 | 0.0252 | 0.1821 | 1.184e-01 |
| AGES | 2226 | 0.0454 | 0.2512 | 4.868e-04 |
| ARIC.EA | 1390 | 0.0289 | -0.0524 | 6.434e-01 |
| ASPS-Family | 338 | 0.0353 | 0.2285 | 2.730e-01 |
| ASRB | 233 | 0.0337 | 0.021 | 9.343e-01 |
| CARDIA.EA | 349 | 0.0297 | 0.1182 | 5.961e-01 |
| CHAP | 261 | 0.0233 | 0.6638 | 2.295e-02 |
| CHS | 544 | 0.0362 | -0.029 | 8.577e-01 |
| DNGS | 515 | 0.0735 | 0.0671 | 5.745e-01 |
| ENIGMA | 11373 | 0.0307 | 0.054 | 1.603e-01 |
| EPOZ | 251 | 0.0284 | 0.1034 | 7.003e-01 |
| FHS | 4710 | 0.0338 | -0.0113 | 8.427e-01 |
| GenR | 265 | 0.0647 | 0.0788 | 6.556e-01 |
| HUNT | 851 | 0.046 | -0.0524 | 6.505e-01 |
| LBC1936 | 577 | 0.0189 | 0.1507 | 4.855e-01 |
| LIFE | 1825 | 0.0304 | 0.0875 | 3.638e-01 |
| LLS | 355 | 0.0344 | -0.3297 | 1.094e-01 |
| MethCT. | 114 | 0.2143 | 0.2129 | 1.870e-01 |
| NOMAS | 141 | 0.0286 | 0.0539 | 8.802e-01 |
| PoznanMS | 204 | 0.042 | 0.3901 | 1.140e-01 |
| PROSPER | 315 | 0.0323 | 0.0385 | 8.640e-01 |
| RS1 | 1103 | 0.0221 | 0.3409 | 1.867e-02 |
| RS2 | 1159 | 0.0267 | -0.1849 | 1.513e-01 |
| RS3 | 2582 | 0.0269 | 0.1105 | 1.991e-01 |
| RUSH1 | 184 | 0.0355 | -0.0606 | 8.301e-01 |
| RUSH2 | 106 | 0.019 | 0.6087 | 2.293e-01 |
| SHIP | 966 | 0.0427 | 0.045 | 6.895e-01 |
| SHIP-TREND | 858 | 0.0331 | 0.1653 | 2.213e-01 |
| SYS.adolescents | 980 | 0.027 | 0.0176 | 8.998e-01 |
| SYS.adults | 594 | 0.0289 | 0.1447 | 4.039e-01 |
| TREND | 1084 | 0.0293 | -0.034 | 7.896e-01 |
| UKB | 14571 | 0.0264 | 0.1205 | 9.810e-04 |
| WHICAP | 83 | 0.0639 | 0.2091 | 5.117e-01 |
| **Non-European** | | | | |
| ARIC.AA | 675 | 0.2717 | 0.0628 | 3.047e-01 |
| CARDIA.AA | 132 | 0.266 | 0.1716 | 2.179e-01 |
| CHAP.black | 321 | 0.2649 | -0.0919 | 3.048e-01 |
| FBIRN | 275 | 0.0717 | 0.2435 | 1.419e-01 |
| NOMAS.black | 168 | 0.2329 | -0.182 | 1.603e-01 |
| NOMAS.hispanic | 718 | 0.1741 | -0.1275 | 6.743e-02 |
| SCES | 209 | 0.1038 | 0.0308 | 8.480e-01 |
| SIMES | 201 | 0.0936 | -0.0247 | 8.850e-01 |
| WHICAP.black | 60 | 0.1721 | -0.4044 | 1.005e-01 |
| **Summary European MA** | **74726** | **0.0315** | **0.083** | **2.112e-08** |
| **Summary African MA** | **1356** | **0.2603** | **-0.01** | **0.8178** |
| **Summary Asian MA** | **1298** | **0.094** | **0.087** | **0.198** |
| **Summary Transancestral MA** | **78487** | **0.0382** | **0.073** | **2.633e-08** |

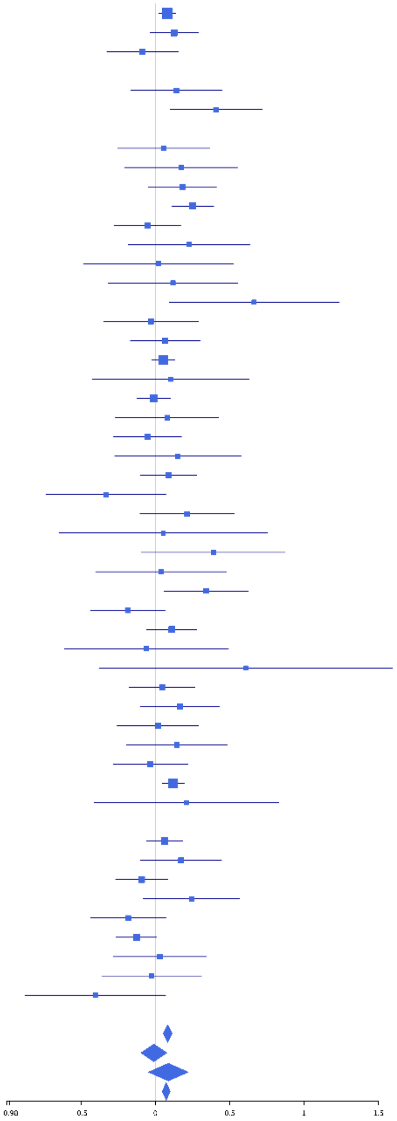

**rs11155234 (T); chr6:142408707; model 1 (I2=22.7, HetP=0.08519)**

| Study | N_total | EAF | Beta | P |
|---|---|---|---|---|
| **Head circumference** | | | | |
| Haworth | 18881 | 0.2177 | -0.0497 | 6.667e-05 |
| VIDI | 680 | 0.1089 | -0.1432 | 1.005e-01 |
| **CT-based** | | | | |
| ESS | 641 | 0.1905 | -0.1773 | 1.267e-02 |
| UCL | 642 | 0.2273 | -0.0217 | 7.441e-01 |
| **MR-based** | | | | |
| 1000Brains | 751 | 0.1979 | 0.0391 | 5.466e-01 |
| 3C-Bordeaux | 564 | 0.2398 | -0.1548 | 2.684e-02 |
| 3c-Dijon | 1499 | 0.239 | 0.0367 | 3.911e-01 |
| AGES | 2226 | 0.2112 | 0.0117 | 7.496e-01 |
| ARIC.EA | 1390 | 0.2028 | -0.0467 | 3.224e-01 |
| ASPS-Family | 338 | 0.2322 | -0.0637 | 4.847e-01 |
| ASRB | 233 | 0.2144 | 0.1045 | 3.557e-01 |
| CARDIA.EA | 349 | 0.2044 | -0.2069 | 2.749e-02 |
| CHAP | 261 | 0.2356 | -0.0437 | 6.717e-01 |
| CHS | 544 | 0.2133 | -0.0064 | 9.306e-01 |
| DNGS | 503 | 0.1373 | -0.1012 | 2.697e-01 |
| ENIGMA | 11373 | 0.2089 | -0.0218 | 1.808e-01 |
| EPOZ | 251 | 0.1924 | 0.1272 | 2.616e-01 |
| ERF | 122 | 0.1314 | -0.2257 | 2.338e-01 |
| FHS | 4710 | 0.2308 | -0.0324 | 1.850e-01 |
| GenR | 265 | 0.1691 | -0.0015 | 9.897e-01 |
| HUNT | 851 | 0.1931 | 0.0017 | 9.781e-01 |
| LBC1936 | 577 | 0.2121 | -0.134 | 6.270e-02 |
| LIFE | 1825 | 0.2138 | -0.0254 | 5.303e-01 |
| LLS | 355 | 0.2312 | 0.0652 | 4.640e-01 |
| MethCT. | 114 | 0.0842 | -0.342 | 1.515e-01 |
| NOMAS | 141 | 0.2479 | 0.0633 | 6.469e-01 |
| PoznanMS | 204 | 0.2201 | -0.0096 | 9.363e-01 |
| PROSPER | 315 | 0.1858 | 0.317 | 1.965e-03 |
| RS1 | 1103 | 0.2013 | -0.0518 | 3.292e-01 |
| RS2 | 1159 | 0.201 | -0.0282 | 5.867e-01 |
| RS3 | 2582 | 0.1974 | -0.0398 | 2.548e-01 |
| RUSH1 | 184 | 0.2141 | -0.1539 | 2.273e-01 |
| RUSH2 | 106 | 0.1773 | -0.0448 | 8.038e-01 |
| SHIP | 966 | 0.2036 | 0.0506 | 3.707e-01 |
| SHIP-TREND | 858 | 0.2224 | -0.127 | 2.892e-02 |
| SYS.adolescents | 980 | 0.2841 | -0.0444 | 3.752e-01 |
| SYS.adults | 594 | 0.268 | -0.0287 | 6.611e-01 |
| TREND | 1084 | 0.1983 | -0.0714 | 1.854e-01 |
| UKB | 14571 | 0.2009 | -0.0429 | 3.375e-03 |
| WHICAP | 83 | 0.2142 | 0.1122 | 5.552e-01 |
| **Non-European** | | | | |
| ARIC.AA | 675 | 0.0455 | 0.0494 | 7.055e-01 |
| CARDIA.AA | 132 | 0.049 | 0.3601 | 2.067e-01 |
| CHAP.black | 321 | 0.0498 | 0.1609 | 3.757e-01 |
| FBIRN | 275 | 0.1399 | 0.1388 | 2.599e-01 |
| NOMAS.black | 168 | 0.06 | -0.4089 | 7.695e-02 |
| NOMAS.hispanic | 718 | 0.1501 | -0.0106 | 8.859e-01 |
| SIMES | 201 | 0.0342 | 0.3196 | 2.373e-01 |
| WHICAP.black | 60 | 0.0496 | -0.1577 | 7.095e-01 |
| **Summary European MA** | **74761** | **0.2109** | **-0.037** | **7.248e-09** |
| **Summary African MA** | **1356** | **0.0488** | **0.035** | **0.696** |
| **Summary Asian MA** | **201** | **0.0342** | **0.324** | **0.2373** |
| **Summary Transancestral MA** | **77425** | **0.2066** | **-0.035** | **2.483e-08** |

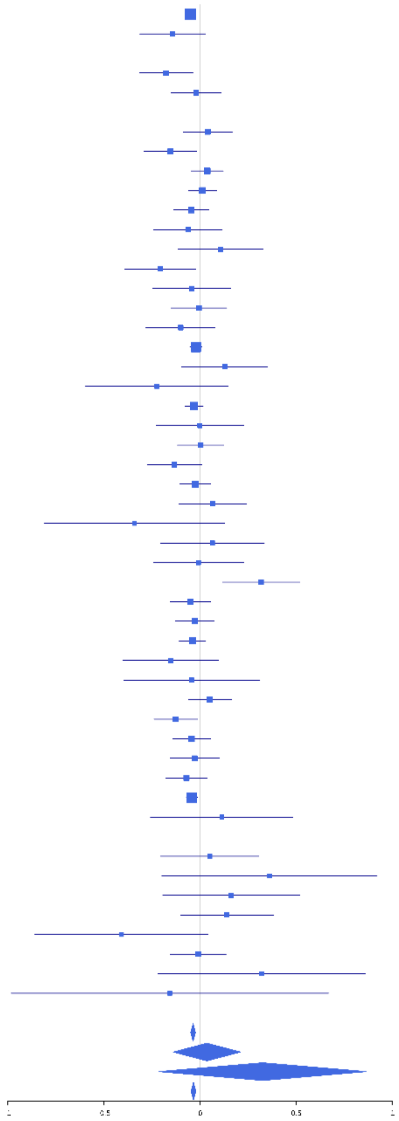

**rs6463758 (A); chr7:8117636; model 1 (I2=9.4, HetP=0.288)**

| Study | N_total | EAF | Beta | P |
|---|---|---|---|---|
| **Head circumference** | | | | |
| Haworth | 18881 | 0.4382 | -0.0157 | 1.312e-01 |
| HKOS | 888 | 0.4696 | 0.084 | 7.730e-02 |
| VIDI | 750 | 0.5269 | -0.0708 | 1.711e-01 |
| **CT-based** | | | | |
| ESS | 641 | 0.4485 | -0.0767 | 1.719e-01 |
| UCL | 642 | 0.4663 | -0.0261 | 6.414e-01 |
| **MR-based** | | | | |
| 1000Brains | 751 | 0.4382 | -0.1293 | 1.312e-02 |
| 3C-Bordeaux | 564 | 0.4205 | -0.0072 | 9.045e-01 |
| 3c-Dijon | 1499 | 0.4516 | 0.009 | 8.074e-01 |
| AGES | 2226 | 0.4636 | -0.0254 | 3.976e-01 |
| ARIC.EA | 1390 | 0.4499 | 0.0368 | 3.343e-01 |
| ASPS-Family | 338 | 0.4606 | 0.0036 | 9.627e-01 |
| CARDIA.EA | 349 | 0.4351 | -0.1344 | 7.844e-02 |
| CHAP | 261 | 0.4114 | -0.1037 | 2.448e-01 |
| CHS | 544 | 0.4641 | -0.0163 | 7.885e-01 |
| DNGS | 512 | 0.4088 | 0.0213 | 7.380e-01 |
| ENIGMA | 11373 | 0.4781 | -0.0361 | 6.553e-03 |
| EPOZ | 251 | 0.4539 | 0.0033 | 9.701e-01 |
| ERF | 122 | 0.4891 | 0.0142 | 9.119e-01 |
| FHS | 4710 | 0.4592 | -0.0832 | 5.752e-05 |
| GenR | 265 | 0.4163 | 0.055 | 5.325e-01 |
| HUNT | 851 | 0.5215 | -0.0617 | 2.038e-01 |
| LBC1936 | 577 | 0.5037 | -0.0465 | 4.299e-01 |
| LIFE | 1825 | 0.4866 | 0.008 | 8.088e-01 |
| LLS | 355 | 0.5032 | -0.0573 | 4.455e-01 |
| MethCT. | 114 | 0.4054 | -0.1422 | 2.921e-01 |
| NOMAS | 141 | 0.4266 | 0.2103 | 8.299e-02 |
| PoznanMS | 204 | 0.4811 | -0.0744 | 4.527e-01 |
| PROSPER | 315 | 0.4722 | -0.0931 | 2.430e-01 |
| RS1 | 1103 | 0.4589 | -0.0454 | 2.884e-01 |
| RS2 | 1159 | 0.4629 | -0.0221 | 5.959e-01 |
| RS3 | 2582 | 0.4652 | -0.0592 | 3.382e-02 |
| RUSH1 | 184 | 0.4511 | -0.0064 | 9.513e-01 |
| RUSH2 | 106 | 0.425 | 0.0796 | 5.680e-01 |
| SHIP | 966 | 0.4877 | -0.0493 | 2.785e-01 |
| SHIP-TREND | 858 | 0.5045 | -0.0379 | 4.326e-01 |
| SYS.adolescents | 980 | 0.4594 | -0.0372 | 4.124e-01 |
| SYS.adults | 594 | 0.4568 | -3e-04 | 9.950e-01 |
| TREND | 1084 | 0.4669 | 0.0157 | 7.159e-01 |
| UKB | 14571 | 0.443 | -0.0277 | 1.883e-02 |
| WHICAP | 83 | 0.4105 | 0.1327 | 4.030e-01 |
| **Non-European** | | | | |
| ARIC.AA | 675 | 0.3056 | 0.0181 | 7.593e-01 |
| CARDIA.AA | 132 | 0.303 | -0.218 | 1.034e-01 |
| CHAP.black | 321 | 0.2491 | 0.0166 | 8.561e-01 |
| FBIRN | 275 | 0.401 | 0.1728 | 4.806e-02 |
| NOMAS.black | 168 | 0.344 | -0.1113 | 3.341e-01 |
| NOMAS.hispanic | 718 | 0.4173 | 0.0518 | 3.334e-01 |
| SCES | 209 | 0.563 | -0.1525 | 1.219e-01 |
| SIMES | 201 | 0.5634 | -0.0196 | 8.374e-01 |
| WHICAP.black | 60 | 0.2937 | -0.0928 | 6.452e-01 |
| **Summary European MA** | **74607** | **0.456** | **-0.029** | **1.971e-08** |
| **Summary African MA** | **1356** | **0.2962** | **-0.027** | **0.5218** |
| **Summary Asian MA** | **1298** | **0.4992** | **0.03** | **0.4474** |
| **Summary Transancestral MA** | **78368** | **0.4533** | **-0.027** | **1.19e-07** |

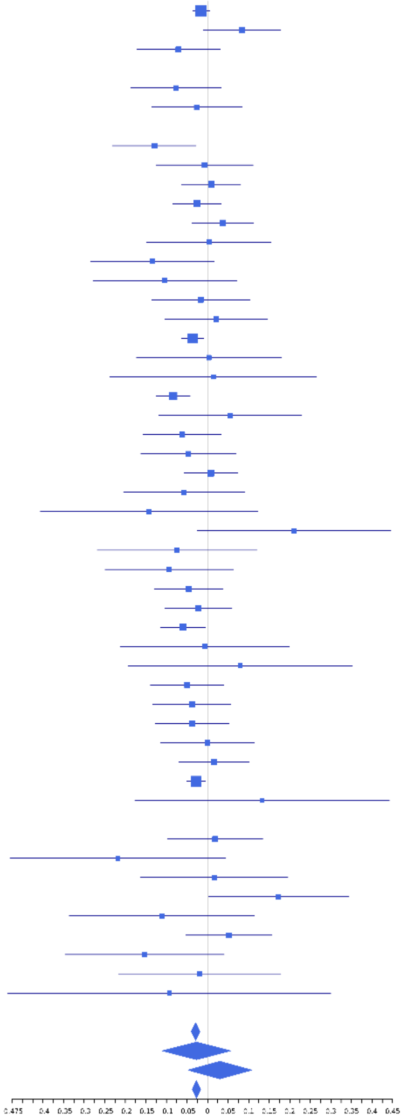

**rs11769806 (A); chr7:23480132; model 1 (I2=31.7, HetP=0.02061)**

| Study | N_total | EAF | Beta | P |
|---|---|---|---|---|
| **Head circumference** | | | | |
| Haworth | 18881 | 0.7574 | 0.0376 | 1.740e-03 |
| VIDI | 754 | 0.8188 | 0.0844 | 2.070e-01 |
| **CT-based** | | | | |
| ESS | 641 | 0.7642 | 0.0659 | 3.163e-01 |
| UCL | 642 | 0.7406 | 0.0707 | 2.669e-01 |
| **MR-based** | | | | |
| 1000Brains | 751 | 0.7809 | -0.0038 | 9.514e-01 |
| 3C-Bordeaux | 564 | 0.7562 | 0.0779 | 2.617e-01 |
| 3c-Dijon | 1499 | 0.7627 | 0.0228 | 5.949e-01 |
| AGES | 2226 | 0.7753 | 0.0791 | 2.750e-02 |
| ARIC.EA | 1390 | 0.7587 | 0.062 | 1.617e-01 |
| ASPS-Family | 338 | 0.8349 | -0.1768 | 8.791e-02 |
| ASRB | 233 | 0.769 | 0.0247 | 8.218e-01 |
| CARDIA.EA | 349 | 0.7584 | 0.0582 | 5.106e-01 |
| CHAP | 261 | 0.745 | -0.1179 | 2.414e-01 |
| CHS | 544 | 0.7395 | -0.1231 | 7.474e-02 |
| DHS | 460 | 0.772 | -0.1599 | 4.190e-02 |
| ENIGMA | 11373 | 0.7684 | 0.0373 | 1.750e-02 |
| EPOZ | 251 | 0.7822 | -0.0995 | 3.575e-01 |
| ERF | 122 | 0.7015 | 0.0827 | 5.547e-01 |
| FHS | 4710 | 0.761 | 0.0259 | 2.846e-01 |
| GenR | 265 | 0.7714 | -0.0205 | 8.427e-01 |
| HUNT | 851 | 0.7719 | 0.0555 | 3.373e-01 |
| LBC1936 | 577 | 0.7554 | -0.0713 | 2.978e-01 |
| LIFE | 1825 | 0.7963 | 0.0846 | 3.975e-02 |
| LLS | 355 | 0.7492 | 0.1408 | 1.039e-01 |
| MethCT. | 114 | 0.8536 | 0.0864 | 6.450e-01 |
| NOMAS | 141 | 0.7378 | 0.0673 | 6.203e-01 |
| PoznanMS | 204 | 0.8122 | -0.2515 | 4.728e-02 |
| PROSPER | 315 | 0.7655 | 0.1365 | 1.464e-01 |
| RS1 | 1103 | 0.7648 | 0.0989 | 4.887e-02 |
| RS2 | 1159 | 0.7565 | 0.0286 | 5.535e-01 |
| RS3 | 2582 | 0.7626 | 0.0514 | 1.160e-01 |
| RUSH1 | 184 | 0.77 | -0.0185 | 8.819e-01 |
| RUSH2 | 106 | 0.7819 | 0.1187 | 4.771e-01 |
| SHIP | 966 | 0.7856 | -0.0175 | 7.529e-01 |
| SHIP-TREND | 858 | 0.761 | -0.0591 | 2.966e-01 |
| SYS.adolescents | 980 | 0.7076 | 0.1736 | 4.732e-04 |
| SYS.adults | 594 | 0.6943 | 0.0875 | 1.644e-01 |
| TREND | 1084 | 0.7756 | -0.0586 | 2.554e-01 |
| UKB | 14571 | 0.7559 | 0.0466 | 6.408e-04 |
| WHICAP | 83 | 0.7317 | 0.0368 | 8.339e-01 |
| **Non-European** | | | | |
| ARIC.AA | 675 | 0.8755 | 0.0191 | 8.165e-01 |
| CARDIA.AA | 132 | 0.861 | -0.3357 | 5.915e-02 |
| CHAP.black | 321 | 0.846 | 0.0571 | 6.020e-01 |
| FBIRN | 275 | 0.7952 | -0.2206 | 3.772e-02 |
| NOMAS.black | 168 | 0.8842 | 0.1217 | 4.764e-01 |
| NOMAS.hispanic | 718 | 0.7899 | -0.055 | 3.962e-01 |
| SIMES | 201 | 0.9407 | -0.1638 | 4.363e-01 |
| WHICAP.black | 60 | 0.8496 | -0.0386 | 8.808e-01 |
| **Summary European MA** | **74792** | **0.7618** | **0.038** | **6.282e-10** |
| **Summary African MA** | **1356** | **0.867** | **0.003** | **0.9619** |
| **Summary Asian MA** | **201** | **0.9407** | **-0.164** | **0.4363** |
| **Summary Transancestral MA** | **77456** | **0.7646** | **0.035** | **4.776e-09** |

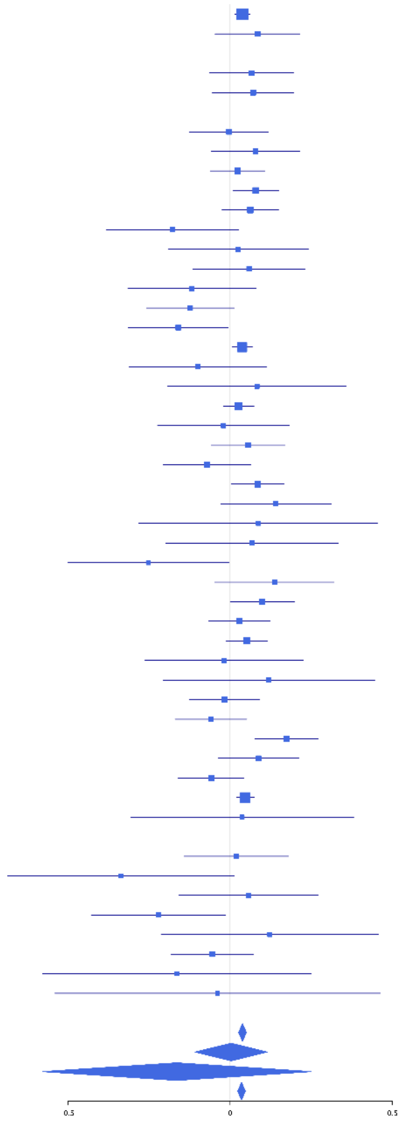

**rs34888260 (A); chr7:32876315; model 1 (I2=0, HetP=0.8554)**

| Study | N_total | EAF | Beta | P |
|---|---|---|---|---|
| **Head circumference** | | | | |
| Haworth | 18881 | 0.3371 | -0.0196 | 7.136e-02 |
| HKOS | 888 | 0.4032 | -0.0499 | 3.021e-01 |
| VIDI | 749 | 0.3528 | -0.0703 | 1.937e-01 |
| **CT-based** | | | | |
| ESS | 641 | 0.3078 | -0.0879 | 1.461e-01 |
| UCL | 642 | 0.3111 | 0.0194 | 7.475e-01 |
| **MR-based** | | | | |
| 1000Brains | 751 | 0.3404 | -0.0255 | 6.396e-01 |
| 3C-Bordeaux | 564 | 0.3297 | -0.0858 | 1.760e-01 |
| 3c-Dijon | 1499 | 0.3319 | -0.1214 | 1.744e-03 |
| AGES | 2226 | 0.3426 | -0.0562 | 7.484e-02 |
| ARIC.EA | 1390 | 0.3399 | -0.0211 | 5.979e-01 |
| ASPS-Family | 338 | 0.3718 | -0.0388 | 6.254e-01 |
| ASRB | 233 | 0.3259 | 0.062 | 5.312e-01 |
| CARDIA.EA | 349 | 0.3566 | -0.0386 | 6.251e-01 |
| CHAP | 261 | 0.3338 | -0.0625 | 5.015e-01 |
| CHS | 544 | 0.3467 | -0.1171 | 6.598e-02 |
| ENIGMA | 11373 | 0.334 | -0.0346 | 1.399e-02 |
| EPOZ | 251 | 0.321 | -0.0433 | 6.506e-01 |
| ERF | 122 | 0.3536 | -0.1643 | 2.199e-01 |
| FHS | 4710 | 0.3405 | -0.0726 | 8.413e-04 |
| GenR | 265 | 0.2969 | 0.001 | 9.917e-01 |
| HUNT | 851 | 0.3098 | 0.0442 | 3.995e-01 |
| LBC1936 | 577 | 0.3072 | -0.1464 | 2.180e-02 |
| LIFE | 1825 | 0.3439 | 0.0116 | 7.399e-01 |
| LLS | 355 | 0.3348 | 0.0333 | 6.753e-01 |
| MethCT. | 114 | 0.2015 | 0.0048 | 9.769e-01 |
| NOMAS | 141 | 0.3802 | -0.2902 | 1.938e-02 |
| PoznanMS | 204 | 0.3494 | 0.0325 | 7.542e-01 |
| PROSPER | 315 | 0.3463 | -0.0541 | 5.185e-01 |
| RS1 | 1103 | 0.3406 | -0.0829 | 6.506e-02 |
| RS2 | 1159 | 0.3324 | -0.0217 | 6.225e-01 |
| RS3 | 2582 | 0.3292 | -0.0518 | 8.038e-02 |
| RUSH1 | 184 | 0.3926 | -0.0733 | 4.931e-01 |
| RUSH2 | 106 | 0.2946 | 0.086 | 5.696e-01 |
| SHIP | 966 | 0.3522 | -0.0043 | 9.278e-01 |
| SHIP-TREND | 858 | 0.3465 | -0.0607 | 2.316e-01 |
| SYS.adolescents | 980 | 0.2801 | -0.046 | 3.603e-01 |
| SYS.adults | 594 | 0.2808 | -0.0375 | 5.610e-01 |
| TREND | 1084 | 0.3505 | -0.0509 | 2.589e-01 |
| UKB | 14571 | 0.3233 | -0.0349 | 5.338e-03 |
| WHICAP | 83 | 0.3798 | -0.2333 | 1.485e-01 |
| **Non-European** | | | | |
| ARIC.AA | 675 | 0.0885 | 0.0565 | 5.554e-01 |
| CARDIA.AA | 132 | 0.099 | -0.0499 | 8.087e-01 |
| CHAP.black | 321 | 0.082 | -0.1777 | 2.179e-01 |
| FBIRN | 275 | 0.2833 | -0.0213 | 8.226e-01 |
| NOMAS.black | 168 | 0.0993 | -0.1115 | 5.422e-01 |
| NOMAS.hispanic | 718 | 0.1965 | 0.0022 | 9.735e-01 |
| SCES | 209 | 0.3607 | -0.0624 | 5.401e-01 |
| SIMES | 201 | 0.3485 | 0.0471 | 6.430e-01 |
| WHICAP.black | 60 | 0.0718 | -0.0488 | 8.908e-01 |
| **Summary European MA** | **74327** | **0.3329** | **-0.037** | **1.615e-11** |
| **Summary African MA** | **1356** | **0.0886** | **-0.034** | **0.6154** |
| **Summary Asian MA** | **1298** | **0.3879** | **-0.037** | **0.3591** |
| **Summary Transancestral MA** | **78088** | **0.3279** | **-0.036** | **1.321e-11** |

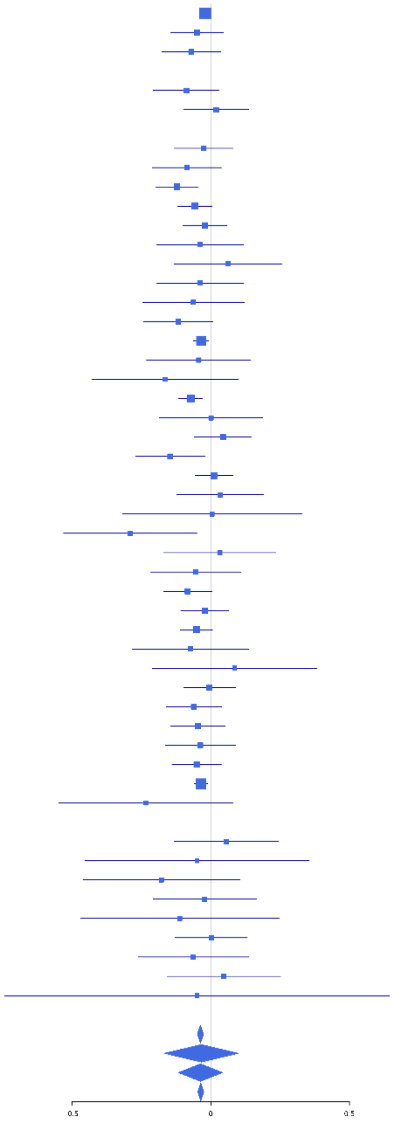

**rs2072235 (T); chr7:50737384; model 1 (I2=3.2, HetP=0.4087)**

| Study | N_total | EAF | Beta | P |
|---|---|---|---|---|
| **Head circumference** | | | | |
| Haworth | 18881 | 0.2454 | 0.0365 | 2.373e-03 |
| HKOS | 888 | 0.5011 | -0.0328 | 4.888e-01 |
| VIDI | 754 | 0.2168 | 0.0321 | 6.079e-01 |
| **CT-based** | | | | |
| ESS | 641 | 0.2263 | 0.0691 | 3.004e-01 |
| UCL | 642 | 0.2077 | 0.0524 | 4.461e-01 |
| **MR-based** | | | | |
| 1000Brains | 751 | 0.2426 | 0.0692 | 2.511e-01 |
| 3C-Bordeaux | 564 | 0.2262 | 0.0475 | 5.042e-01 |
| 3c-Dijon | 1499 | 0.2302 | 0.091 | 3.603e-02 |
| AGES | 2226 | 0.1976 | 0.0959 | 1.086e-02 |
| ARIC.EA | 1390 | 0.2411 | 0.0327 | 4.604e-01 |
| ASPS-Family | 338 | 0.253 | 0.0185 | 8.347e-01 |
| ASRB | 233 | 0.2427 | 0.0406 | 7.073e-01 |
| CARDIA.EA | 349 | 0.2389 | 0.2131 | 1.637e-02 |
| CHAP | 261 | 0.2299 | -0.0336 | 7.467e-01 |
| CHS | 544 | 0.2571 | -0.0396 | 5.681e-01 |
| DHS | 460 | 0.2236 | 0.1765 | 2.571e-02 |
| DNGS | 514 | 0.2724 | 0.0301 | 6.674e-01 |
| ENIGMA | 11373 | 0.2424 | 0.0315 | 4.163e-02 |
| EPOZ | 251 | 0.2429 | 0.0269 | 7.968e-01 |
| ERF | 122 | 0.2715 | 0.0261 | 8.563e-01 |
| FHS | 4710 | 0.2461 | 0.0591 | 1.350e-02 |
| GenR | 265 | 0.2261 | 0.282 | 6.605e-03 |
| HUNT | 851 | 0.2503 | 0.1059 | 5.838e-02 |
| LBC1936 | 577 | 0.2267 | 0.0166 | 8.132e-01 |
| LIFE | 1825 | 0.2583 | 0.0967 | 1.067e-02 |
| LLS | 355 | 0.2327 | -0.0537 | 5.455e-01 |
| MethCT. | 114 | 0.1403 | 0.3038 | 1.111e-01 |
| NOMAS | 141 | 0.3385 | 0.0404 | 7.487e-01 |
| PoznanMS | 204 | 0.3118 | -0.0291 | 7.858e-01 |
| PROSPER | 315 | 0.2127 | 0.0399 | 6.816e-01 |
| RS1 | 1103 | 0.2333 | 0.008 | 8.747e-01 |
| RS2 | 1159 | 0.217 | 0.0679 | 1.776e-01 |
| RS3 | 2582 | 0.2225 | 0.0497 | 1.374e-01 |
| RUSH1 | 184 | 0.2418 | 0.1767 | 1.485e-01 |
| RUSH2 | 106 | 0.1983 | -0.0501 | 7.717e-01 |
| SHIP | 966 | 0.2579 | 0.0367 | 4.804e-01 |
| SHIP-TREND | 858 | 0.2477 | 0.1309 | 1.953e-02 |
| SYS.adolescents | 980 | 0.2759 | 0.0722 | 1.532e-01 |
| SYS.adults | 594 | 0.2861 | -0.0804 | 2.102e-01 |
| TREND | 1084 | 0.2458 | 0.0256 | 6.070e-01 |
| UKB | 14571 | 0.2147 | 0.0536 | 1.700e-04 |
| WHICAP | 83 | 0.3256 | 0.0616 | 7.112e-01 |
| **Non-European** | | | | |
| ARIC.AA | 675 | 0.0458 | -0.02 | 8.776e-01 |
| CARDIA.AA | 132 | 0.044 | 0.3238 | 2.806e-01 |
| CHAP.black | 321 | 0.0438 | -0.3764 | 5.182e-02 |
| FBIRN | 275 | 0.2679 | -0.0998 | 3.009e-01 |
| NOMAS.black | 168 | 0.0656 | 0.4801 | 3.074e-02 |
| NOMAS.hispanic | 718 | 0.1711 | 0.0915 | 1.920e-01 |
| SCES | 209 | 0.5459 | 0.0592 | 5.463e-01 |
| SIMES | 201 | 0.3129 | -0.1198 | 2.658e-01 |
| WHICAP.black | 60 | 0.0601 | 0.0991 | 7.971e-01 |
| **Summary European MA** | **75306** | **0.2363** | **0.048** | **2.504e-15** |
| **Summary African MA** | **1356** | **0.0482** | **0.009** | **0.9233** |
| **Summary Asian MA** | **1298** | **0.4792** | **-0.03** | **0.4423** |
| **Summary Transancestral MA** | **79067** | **0.2364** | **0.046** | **8.431e-15** |

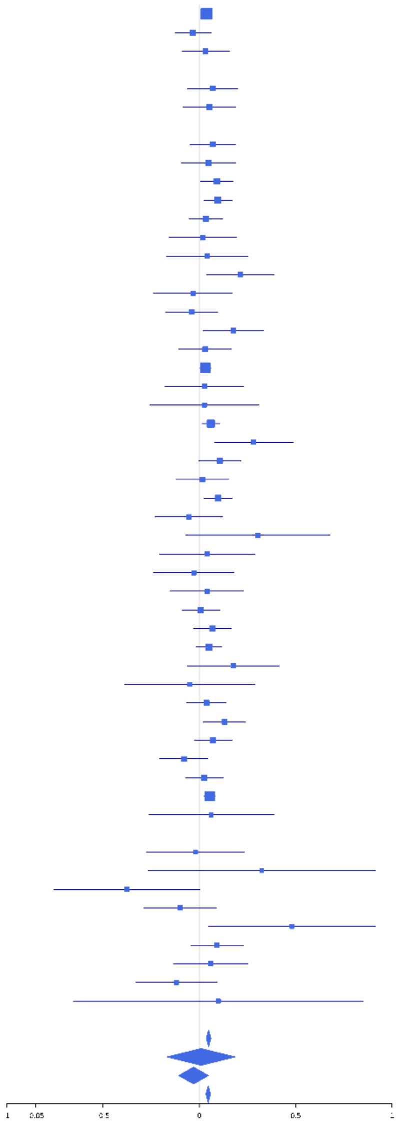

**rs151057105 (T); chr7:54944920; model 1 (I2=29.3, HetP=0.03091)**

| Study | N_total | EAF | Beta | P |
|---|---|---|---|---|
| **Head circumference** | | | | |
| Haworth | 18881 | 0.105 | 0.0418 | 1.255e-02 |
| HKOS | 888 | 0.1368 | 0.1032 | 1.350e-01 |
| VIDI | 731 | 0.1006 | 0.0683 | 4.321e-01 |
| **CT-based** | | | | |
| ESS | 641 | 0.0938 | 0.0201 | 8.334e-01 |
| UCL | 642 | 0.1042 | 0.1125 | 2.178e-01 |
| **MR-based** | | | | |
| 1000Brains | 751 | 0.1024 | 0.1977 | 2.044e-02 |
| 3C-Bordeaux | 564 | 0.1062 | 0.1862 | 5.449e-02 |
| 3c-Dijon | 1499 | 0.1109 | 0.0092 | 8.748e-01 |
| AGES | 2226 | 0.084 | 0.0807 | 1.353e-01 |
| ARIC.EA | 1390 | 0.0921 | 0.2175 | 9.130e-04 |
| ASPS-Family | 338 | 0.0984 | 0.2728 | 3.466e-02 |
| ASRB | 233 | 0.0961 | 0.0354 | 8.222e-01 |
| CARDIA.EA | 349 | 0.0982 | -0.1468 | 2.486e-01 |
| CHAP | 261 | 0.1059 | -0.239 | 9.416e-02 |
| CHS | 544 | 0.1035 | 0.1173 | 2.382e-01 |
| DNGS | 504 | 0.091 | 0.2312 | 3.528e-02 |
| ENIGMA | 11373 | 0.1018 | 0.0431 | 4.940e-02 |
| EPOZ | 251 | 0.0682 | 0.3074 | 8.237e-02 |
| FHS | 4710 | 0.1063 | 0.0874 | 8.947e-03 |
| GenR | 265 | 0.089 | 0.2471 | 1.053e-01 |
| HUNT | 851 | 0.0964 | -0.0801 | 3.294e-01 |
| LBC1936 | 577 | 0.1126 | 0.0386 | 6.790e-01 |
| LIFE | 1825 | 0.1005 | -0.0369 | 5.031e-01 |
| LLS | 355 | 0.0972 | 0.0929 | 4.634e-01 |
| MethCT. | 114 | 0.1461 | -0.3881 | 3.846e-02 |
| NOMAS | 141 | 0.1306 | 0.1386 | 4.344e-01 |
| PoznanMS | 204 | 0.1003 | 0.0945 | 5.669e-01 |
| PROSPER | 315 | 0.1027 | -0.0248 | 8.501e-01 |
| RS1 | 1103 | 0.0871 | 0.2095 | 5.530e-03 |
| RS2 | 1159 | 0.0814 | 0.187 | 1.386e-02 |
| RS3 | 2582 | 0.0839 | -8e-04 | 9.883e-01 |
| RUSH1 | 184 | 0.1363 | -0.2686 | 7.873e-02 |
| RUSH2 | 106 | 0.1064 | 0.2038 | 3.626e-01 |
| SHIP | 966 | 0.0884 | 0.0915 | 2.540e-01 |
| SHIP-TREND | 858 | 0.0912 | -0.0113 | 8.923e-01 |
| SYS.adolescents | 980 | 0.085 | 0.0637 | 4.311e-01 |
| SYS.adults | 594 | 0.0936 | -0.0067 | 9.465e-01 |
| TREND | 1084 | 0.0921 | 0.0022 | 9.772e-01 |
| UKB | 14571 | 0.0922 | 0.1176 | 6.303e-09 |
| WHICAP | 83 | 0.1407 | -0.0429 | 8.479e-01 |
| **Non-European** | | | | |
| ARIC.AA | 675 | 0.0529 | 0.0355 | 7.702e-01 |
| CARDIA.AA | 132 | 0.048 | -0.1529 | 5.953e-01 |
| CHAP.black | 321 | 0.066 | -0.0691 | 6.635e-01 |
| FBIRN | 275 | 0.0768 | 0.07 | 6.627e-01 |
| NOMAS.black | 168 | 0.0374 | -0.1547 | 5.916e-01 |
| NOMAS.hispanic | 718 | 0.0721 | 0.0216 | 8.318e-01 |
| SCES | 209 | 0.0998 | 0.0951 | 5.598e-01 |
| SIMES | 201 | 0.0934 | -0.0127 | 9.375e-01 |
| WHICAP.black | 60 | 0.0544 | 0.4351 | 2.847e-01 |
| **Summary European MA** | **74691** | **0.0985** | **0.067** | **9.25e-15** |
| **Summary African MA** | **1356** | **0.0537** | **-0.011** | **0.8925** |
| **Summary Asian MA** | **1298** | **0.1241** | **0.086** | **0.15** |
| **Summary Transancestral MA** | **78452** | **0.0979** | **0.065** | **1.406e-14** |

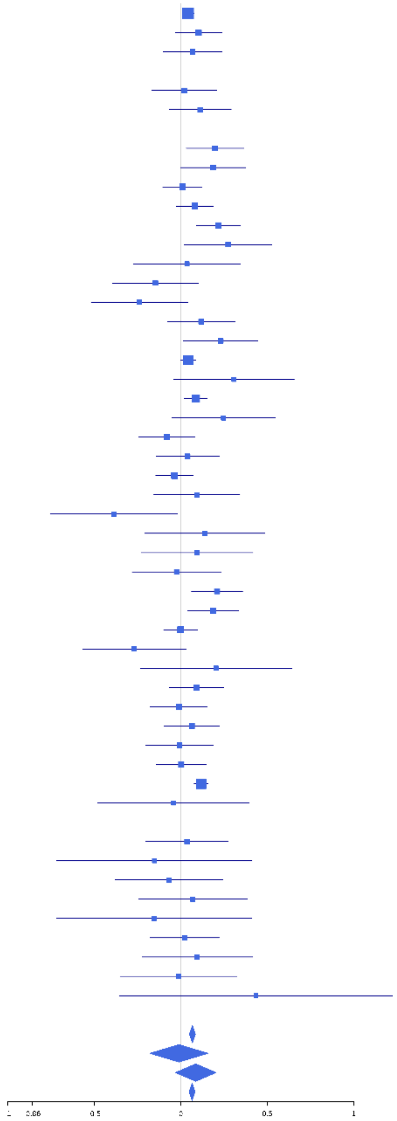

**rs42035 (A); chr7:92239531; model 1 (I2=3.6, HetP=0.4013)**

| Study | N_total | EAF | Beta | P |
|---|---|---|---|---|
| **Head circumference** | | | | |
| Haworth | 18881 | 0.753 | 0.0248 | 3.655e-02 |
| HKOS | 888 | 0.8756 | -0.0747 | 2.986e-01 |
| VIDI | 755 | 0.7756 | 0.176 | 4.426e-03 |
| **CT-based** | | | | |
| ESS | 641 | 0.759 | 0.0112 | 8.644e-01 |
| UCL | 642 | 0.7688 | -0.0735 | 2.667e-01 |
| **MR-based** | | | | |
| 1000Brains | 751 | 0.7557 | 0.0721 | 2.305e-01 |
| 3C-Bordeaux | 564 | 0.7235 | 0.0457 | 4.931e-01 |
| 3c-Dijon | 1499 | 0.7306 | 0.0482 | 2.420e-01 |
| AGES | 2226 | 0.7764 | 0.0138 | 7.007e-01 |
| ARIC.EA | 1390 | 0.7507 | -0.0062 | 8.871e-01 |
| ASPS-Family | 338 | 0.7416 | 0.0014 | 9.872e-01 |
| ASRB | 233 | 0.7532 | -0.013 | 9.040e-01 |
| CARDIA.EA | 349 | 0.7386 | 0.0627 | 4.668e-01 |
| CHAP | 261 | 0.756 | 0.1138 | 2.652e-01 |
| CHS | 544 | 0.7218 | 0.0792 | 2.421e-01 |
| DHS | 460 | 0.7787 | 0.1054 | 1.844e-01 |
| DNGS | 511 | 0.8027 | 0.0442 | 5.741e-01 |
| ENIGMA | 11373 | 0.7555 | 0.0423 | 6.066e-03 |
| EPOZ | 251 | 0.7559 | -0.1096 | 2.913e-01 |
| ERF | 122 | 0.8464 | 0.2698 | 1.284e-01 |
| FHS | 4710 | 0.7435 | 0.0259 | 2.725e-01 |
| GenR | 265 | 0.7706 | 0.0516 | 6.180e-01 |
| HUNT | 851 | 0.7287 | -0.0339 | 5.343e-01 |
| LBC1936 | 577 | 0.76 | -0.0239 | 7.289e-01 |
| LIFE | 1825 | 0.7501 | 0.0503 | 1.884e-01 |
| LLS | 355 | 0.7435 | 0.0533 | 5.354e-01 |
| MethCT. | 114 | 0.8514 | 0.2044 | 2.721e-01 |
| NOMAS | 141 | 0.7504 | -0.1304 | 3.448e-01 |
| PoznanMS | 204 | 0.7522 | 0.2675 | 1.967e-02 |
| PROSPER | 315 | 0.7323 | -0.0303 | 7.363e-01 |
| RS1 | 1103 | 0.742 | -0.0471 | 3.329e-01 |
| RS2 | 1159 | 0.7421 | 0.0029 | 9.512e-01 |
| RS3 | 2582 | 0.7384 | 0.0875 | 5.701e-03 |
| RUSH1 | 184 | 0.7627 | 0.0468 | 7.028e-01 |
| RUSH2 | 106 | 0.7437 | 0.1634 | 3.011e-01 |
| SHIP | 966 | 0.7421 | 0.0609 | 2.416e-01 |
| SHIP-TREND | 858 | 0.7471 | -0.0122 | 8.262e-01 |
| SYS.adolescents | 980 | 0.764 | 0.0472 | 3.750e-01 |
| SYS.adults | 594 | 0.7703 | -0.0576 | 4.039e-01 |
| TREND | 1084 | 0.7399 | 0.0512 | 2.961e-01 |
| UKB | 14571 | 0.7535 | 0.0454 | 8.409e-04 |
| WHICAP | 83 | 0.7224 | 0.3652 | 3.834e-02 |
| **Non-European** | | | | |
| ARIC.AA | 675 | 0.9208 | -0.039 | 6.989e-01 |
| CARDIA.AA | 132 | 0.905 | -0.0162 | 9.388e-01 |
| CHAP.black | 321 | 0.9282 | -0.0023 | 9.884e-01 |
| FBIRN | 275 | 0.7782 | 0.0041 | 9.685e-01 |
| NOMAS.black | 168 | 0.9083 | 0.1423 | 4.525e-01 |
| NOMAS.hispanic | 718 | 0.8077 | 0.1876 | 5.210e-03 |
| SCES | 209 | 0.9151 | -0.0453 | 7.962e-01 |
| SIMES | 201 | 0.861 | -0.1768 | 2.178e-01 |
| WHICAP.black | 60 | 0.8935 | -0.0722 | 8.083e-01 |
| **Summary European MA** | **75304** | **0.7523** | **0.035** | **5.396e-09** |
| **Summary African MA** | **1356** | **0.9183** | **-0.006** | **0.9279** |
| **Summary Asian MA** | **1298** | **0.8797** | **-0.087** | **0.1475** |
| **Summary Transancestral MA** | **79065** | **0.758** | **0.034** | **6.366e-09** |

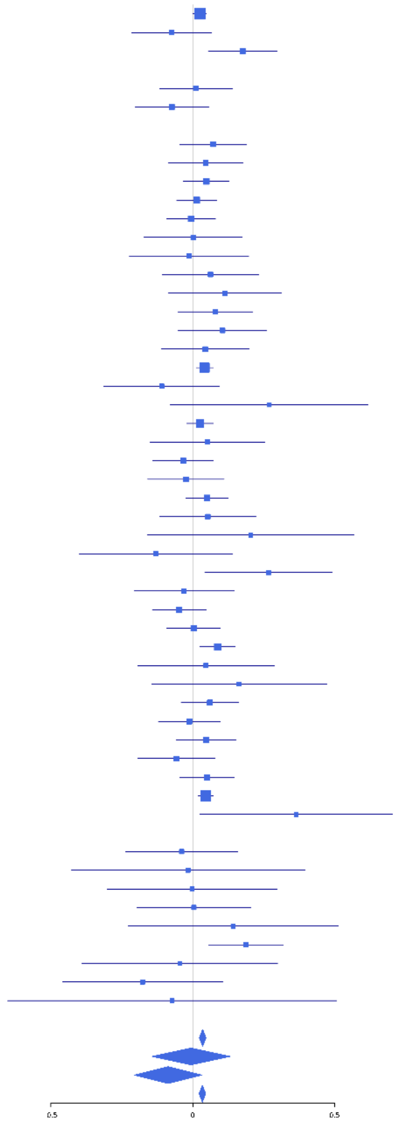

### rs7781436 (T); chr7:92317752; model 1 (I2=7.1, HetP=0.3336)

| Study | N_total | EAF | Beta | P |
|---|---|---|---|---|
| **Head circumference** | | | | |
| Haworth | 18881 | 0.7783 | -0.028 | 2.401e-02 |
| HKOS | 888 | 0.9572 | 0.0114 | 9.226e-01 |
| VIDI | 627 | 0.8008 | -0.149 | 3.541e-02 |
| **CT-based** | | | | |
| ESS | 641 | 0.8121 | -0.0326 | 6.482e-01 |
| UCL | 642 | 0.7735 | 0.0161 | 8.087e-01 |
| **MR-based** | | | | |
| 1000Brains | 751 | 0.8061 | -0.0886 | 1.750e-01 |
| 3C-Bordeaux | 564 | 0.7951 | -0.0509 | 4.903e-01 |
| 3c-Dijon | 1499 | 0.7902 | -0.0486 | 2.782e-01 |
| AGES | 2226 | 0.766 | -0.051 | 1.493e-01 |
| ARIC.EA | 1390 | 0.7827 | -0.1301 | 4.672e-03 |
| ASPS-Family | 338 | 0.784 | -0.0305 | 7.445e-01 |
| ASRB | 233 | 0.7275 | -0.0495 | 6.348e-01 |
| CHAP | 261 | 0.7586 | -0.0652 | 5.249e-01 |
| CHS | 544 | 0.7765 | -0.0266 | 7.145e-01 |
| DHS | 460 | 0.767 | 0.0785 | 3.144e-01 |
| DNGS | 516 | 0.8496 | -0.0247 | 7.768e-01 |
| ENIGMA | 11373 | 0.7925 | -0.0058 | 7.216e-01 |
| EPOZ | 251 | 0.8091 | 0.1114 | 3.267e-01 |
| FHS | 4710 | 0.7809 | -0.0846 | 6.837e-04 |
| GenR | 265 | 0.8165 | -0.127 | 2.578e-01 |
| HUNT | 851 | 0.7961 | -0.113 | 6.027e-02 |
| LBC1936 | 577 | 0.7924 | 0.0504 | 4.872e-01 |
| LIFE | 1825 | 0.8233 | 0.0046 | 9.156e-01 |
| LLS | 355 | 0.7789 | -0.1273 | 1.592e-01 |
| NOMAS | 140 | 0.7964 | 0.0519 | 7.271e-01 |
| PoznanMS | 204 | 0.8717 | -0.22 | 1.373e-01 |
| PROSPER | 315 | 0.827 | -0.0509 | 6.291e-01 |
| RS1 | 1103 | 0.8007 | 0.0082 | 8.782e-01 |
| RS2 | 1159 | 0.7867 | -0.0536 | 2.904e-01 |
| RS3 | 2582 | 0.8037 | -0.0757 | 3.075e-02 |
| RUSH1 | 184 | 0.7989 | -0.0819 | 5.293e-01 |
| RUSH2 | 106 | 0.7594 | -2e-04 | 9.989e-01 |
| SHIP | 966 | 0.8256 | -0.0068 | 9.094e-01 |
| SHIP-TREND | 858 | 0.8205 | 0.0077 | 9.028e-01 |
| SYS.adolescents | 980 | 0.7423 | -0.0603 | 2.433e-01 |
| SYS.adults | 594 | 0.75 | 0.0486 | 4.679e-01 |
| TREND | 1084 | 0.8168 | -0.1697 | 2.287e-03 |
| UKB | 14571 | 0.7802 | -0.0481 | 6.744e-04 |
| WHICAP | 83 | 0.7593 | -0.1514 | 4.067e-01 |
| **Non-European** | | | | |
| ARIC.AA | 675 | 0.9669 | -0.1082 | 4.768e-01 |
| CARDIA.AA | 132 | 0.957 | -0.3707 | 2.216e-01 |
| CHAP.black | 321 | 0.9533 | 0.2 | 2.861e-01 |
| FBIRN | 275 | 0.8362 | -0.0865 | 4.536e-01 |
| NOMAS.black | 166 | 0.9554 | -0.0178 | 9.466e-01 |
| NOMAS.hispanic | 714 | 0.8789 | -0.1336 | 1.000e-01 |
| SCES | 209 | 0.9474 | 0.1601 | 4.646e-01 |
| SIMES | 201 | 0.9826 | 0.0175 | 9.633e-01 |
| WHICAP.black | 60 | 0.9417 | 0.8785 | 2.830e-02 |
| **Summary European MA** | **74709** | **0.7864** | **-0.039** | **4.95e-10** |
| **Summary African MA** | **1354** | **0.9602** | **0.007** | **0.9413** |
| **Summary Asian MA** | **1298** | **0.9596** | **0.039** | **0.6951** |
| **Summary Transancestral MA** | **78350** | **0.7933** | **-0.039** | **5.11e-10** |

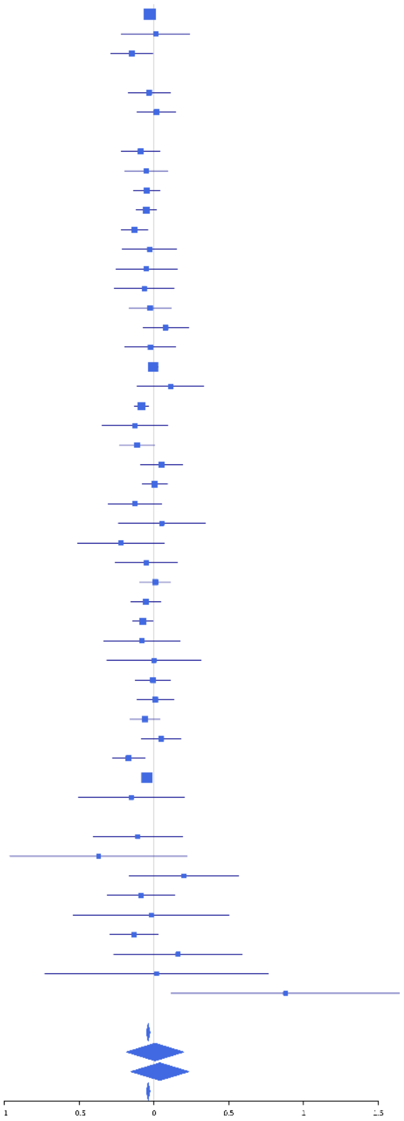

**rs35926075 (T); chr7:92367869; model 1 (I2=14.5, HetP=0.2137)**

| Study | N_total | EAF | Beta | P |
|---|---|---|---|---|
| **Head circumference** | | | | |
| Haworth | 18881 | 0.1699 | -0.0448 | 1.109e-03 |
| VIDI | 661 | 0.1745 | -0.023 | 7.505e-01 |
| **CT-based** | | | | |
| ESS | 641 | 0.1582 | -0.0552 | 4.710e-01 |
| UCL | 642 | 0.1528 | 0.0413 | 5.946e-01 |
| **MR-based** | | | | |
| 1000Brains | 751 | 0.1916 | 0.066 | 3.147e-01 |
| 3C-Bordeaux | 564 | 0.1339 | 0.1326 | 1.297e-01 |
| 3c-Dijon | 1499 | 0.1471 | 0.0388 | 4.520e-01 |
| AGES | 2226 | 0.1758 | -0.106 | 7.094e-03 |
| ARIC.EA | 1390 | 0.165 | -0.1144 | 2.518e-02 |
| ASPS-Family | 338 | 0.1632 | -0.0142 | 8.921e-01 |
| CHAP | 261 | 0.2065 | -0.0255 | 8.133e-01 |
| CHS | 544 | 0.1529 | -0.097 | 2.498e-01 |
| ENIGMA | 11373 | 0.1751 | -0.019 | 2.761e-01 |
| EPOZ | 251 | 0.1795 | -0.1857 | 1.102e-01 |
| FHS | 4710 | 0.1621 | -0.062 | 2.668e-02 |
| GenR | 265 | 0.1293 | -0.0933 | 4.710e-01 |
| HUNT | 851 | 0.1676 | -0.2105 | 1.180e-03 |
| LBC1936 | 577 | 0.183 | -0.047 | 5.374e-01 |
| LIFE | 1825 | 0.1898 | -0.101 | 1.675e-02 |
| LLS | 355 | 0.1838 | 0.0698 | 4.713e-01 |
| NOMAS | 141 | 0.1376 | 0.001 | 9.956e-01 |
| PoznanMS | 204 | 0.2351 | 0.078 | 5.040e-01 |
| PROSPER | 315 | 0.1957 | -0.146 | 1.459e-01 |
| RS1 | 1103 | 0.1853 | -0.0115 | 8.347e-01 |
| RS2 | 1159 | 0.1869 | 0.0269 | 6.140e-01 |
| RS3 | 2582 | 0.1855 | -0.0308 | 3.898e-01 |
| RUSH1 | 184 | 0.2252 | -0.1144 | 3.602e-01 |
| RUSH2 | 106 | 0.2289 | -0.0669 | 6.837e-01 |
| SHIP | 966 | 0.177 | -0.0356 | 5.507e-01 |
| SHIP-TREND | 858 | 0.18 | -0.0155 | 8.056e-01 |
| SYS.adolescents | 980 | 0.1247 | -0.0925 | 1.759e-01 |
| SYS.adults | 594 | 0.1259 | -0.2261 | 9.740e-03 |
| TREND | 1084 | 0.1942 | -0.0677 | 2.127e-01 |
| UKB | 14571 | 0.1699 | -0.033 | 3.435e-02 |
| WHICAP | 83 | 0.116 | -0.3713 | 1.297e-01 |
| **Non-European** | | | | |
| ARIC.AA | 675 | 0.0269 | -0.3494 | 3.777e-02 |
| CHAP.black | 321 | 0.0544 | -0.2133 | 2.211e-01 |
| FBIRN | 275 | 0.1007 | 0.073 | 6.070e-01 |
| NOMAS.black | 168 | 0.0355 | -0.2217 | 4.532e-01 |
| NOMAS.hispanic | 718 | 0.0684 | 0.063 | 5.466e-01 |
| WHICAP.black | 60 | 0.03 | -0.1466 | 7.852e-01 |
| **Summary European MA** | **73535** | **0.1711** | **-0.041** | **2.161e-09** |
| **Summary African MA** | **1224** | **0.0354** | **-0.274** | **0.01216** |
| **Summary Transancestral MA** | **75752** | **0.1677** | **-0.042** | **9.008e-10** |

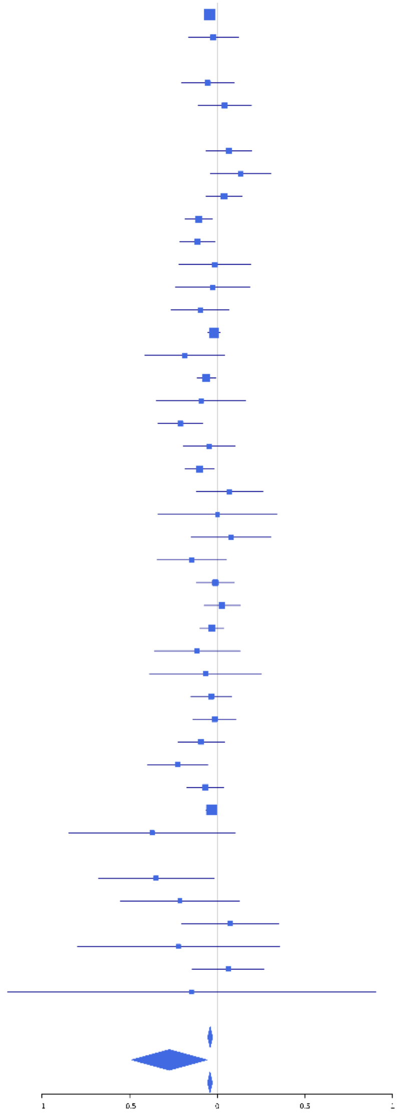

**rs111295525 (A); chr7:135187742; model 1 (I2=8.7, HetP=0.298)**

| Study | N_total | EAF | Beta | P |
|---|---|---|---|---|
| **Head circumference** | | | | |
| Haworth | 18881 | 0.8941 | 0.0528 | 1.612e-03 |
| HKOS | 888 | 0.8806 | -0.0398 | 5.864e-01 |
| VIDI | 747 | 0.8463 | -0.0701 | 3.288e-01 |
| **CT-based** | | | | |
| ESS | 641 | 0.9151 | 0.2125 | 3.401e-02 |
| UCL | 642 | 0.8942 | -0.0095 | 9.161e-01 |
| **MR-based** | | | | |
| 1000Brains | 751 | 0.897 | -0.0156 | 8.540e-01 |
| 3C-Bordeaux | 564 | 0.9015 | -0.2053 | 4.048e-02 |
| 3c-Dijon | 1499 | 0.9015 | 0.0486 | 4.279e-01 |
| AGES | 2226 | 0.8984 | 0.1428 | 3.989e-03 |
| ARIC.EA | 1390 | 0.9017 | 0.1111 | 8.123e-02 |
| ASPS-Family | 338 | 0.8525 | 0.1405 | 1.955e-01 |
| ASRB | 233 | 0.889 | 0.0958 | 5.163e-01 |
| CARDIA.EA | 349 | 0.8889 | 0.0605 | 6.153e-01 |
| CHAP | 261 | 0.866 | -0.1688 | 1.900e-01 |
| CHS | 544 | 0.9079 | 0.0419 | 6.894e-01 |
| DHS | 460 | 0.912 | 0.0973 | 4.033e-01 |
| DNGS | 515 | 0.907 | 0.0191 | 8.590e-01 |
| ENIGMA | 11373 | 0.9025 | 0.0358 | 1.095e-01 |
| EPOZ | 251 | 0.9101 | 0.0136 | 9.306e-01 |
| ERF | 122 | 0.8972 | -0.0403 | 8.488e-01 |
| FHS | 4710 | 0.899 | 0.0347 | 3.107e-01 |
| GenR | 265 | 0.9154 | 0.1686 | 2.801e-01 |
| HUNT | 851 | 0.892 | 0.0132 | 8.661e-01 |
| LBC1936 | 577 | 0.8926 | -0.0413 | 6.641e-01 |
| LIFE | 1825 | 0.8808 | 0.0392 | 4.430e-01 |
| LLS | 355 | 0.9011 | -0.0796 | 5.270e-01 |
| MethCT. | 114 | 0.9038 | 0.0604 | 7.881e-01 |
| NOMAS | 141 | 0.8884 | 0.2351 | 2.160e-01 |
| PoznanMS | 204 | 0.8754 | -0.0526 | 7.258e-01 |
| PROSPER | 315 | 0.9044 | 0.1822 | 1.785e-01 |
| RS1 | 1103 | 0.9116 | 0.0205 | 7.847e-01 |
| RS2 | 1159 | 0.906 | 0.0935 | 1.888e-01 |
| RS3 | 2582 | 0.9056 | 0.0495 | 2.981e-01 |
| RUSH1 | 184 | 0.8886 | 0.0555 | 7.378e-01 |
| RUSH2 | 106 | 0.8949 | 0.1247 | 5.784e-01 |
| SHIP | 966 | 0.8742 | 0.0193 | 7.781e-01 |
| SHIP-TREND | 858 | 0.9017 | 0.0698 | 3.895e-01 |
| SYS.adolescents | 980 | 0.8431 | -0.0426 | 4.928e-01 |
| SYS.adults | 594 | 0.8416 | -0.1212 | 1.273e-01 |
| TREND | 1084 | 0.8808 | 0.0108 | 8.705e-01 |
| UKB | 14571 | 0.9014 | 0.064 | 1.124e-03 |
| WHICAP | 83 | 0.9379 | 0.6509 | 4.646e-02 |
| **Non-European** | | | | |
| ARIC.AA | 675 | 0.9628 | -0.265 | 6.535e-02 |
| CARDIA.AA | 132 | 0.954 | 0.5153 | 7.943e-02 |
| CHAP.black | 321 | 0.9616 | -0.2393 | 2.450e-01 |
| FBIRN | 275 | 0.9181 | -0.0235 | 8.798e-01 |
| NOMAS.black | 168 | 0.9778 | 0.3458 | 3.517e-01 |
| NOMAS.hispanic | 718 | 0.9369 | 0.109 | 3.157e-01 |
| SCES | 209 | 0.9033 | -0.1961 | 2.361e-01 |
| SIMES | 201 | 0.8404 | 0.0666 | 6.208e-01 |
| WHICAP.black | 60 | 0.9519 | 0.7897 | 6.968e-02 |
| **Summary European MA** | **75300** | **0.8965** | **0.046** | **4.257e-08** |
| **Summary African MA** | **1356** | **0.963** | **-0.062** | **0.5424** |
| **Summary Asian MA** | **1298** | **0.878** | **-0.044** | **0.4649** |
| **Summary Transancestral MA** | **79061** | **0.8978** | **0.044** | **1.354e-07** |

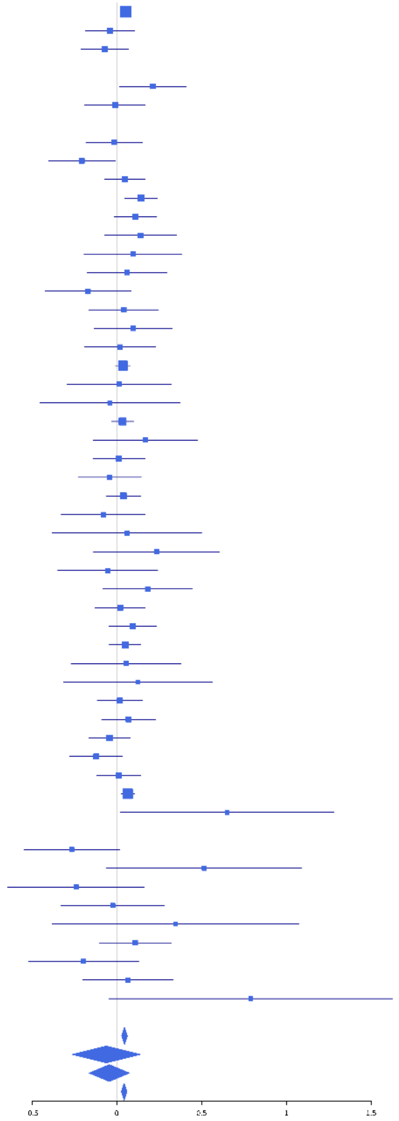

**rs58153210 (T); chr8:26271120; model 1 (I2=15.8, HetP=0.1733)**

| Study | N_total | EAF | Beta | P |
|---|---|---|---|---|
| **Head circumference** | | | | |
| Haworth | 18881 | 0.5828 | 0.0277 | 7.946e-03 |
| HKOS | 888 | 0.8812 | 0.1124 | 1.255e-01 |
| VIDI | 731 | 0.654 | 0.0465 | 3.976e-01 |
| **CT-based** | | | | |
| ESS | 641 | 0.5844 | 0.1383 | 1.468e-02 |
| UCL | 642 | 0.5682 | 0.0177 | 7.536e-01 |
| **MR-based** | | | | |
| 1000Brains | 751 | 0.6116 | 0.0536 | 3.115e-01 |
| 3C-Bordeaux | 564 | 0.5697 | 0.0343 | 5.692e-01 |
| 3c-Dijon | 1499 | 0.5804 | 0.0994 | 7.200e-03 |
| AGES | 2226 | 0.6023 | 0.0208 | 4.971e-01 |
| ARIC.EA | 1390 | 0.5825 | 0.0701 | 6.810e-02 |
| ASPS-Family | 338 | 0.6121 | -0.0223 | 7.783e-01 |
| ASRB | 233 | 0.5845 | -0.0164 | 8.611e-01 |
| CARDIA.EA | 349 | 0.5929 | 0.0263 | 7.333e-01 |
| CHAP | 261 | 0.5712 | 0.0664 | 4.533e-01 |
| CHS | 544 | 0.5511 | 0.1485 | 1.483e-02 |
| DNGS | 504 | 0.6242 | 0.0083 | 8.982e-01 |
| ENIGMA | 11373 | 0.586 | 0.0162 | 2.275e-01 |
| EPOZ | 251 | 0.574 | -0.0155 | 8.632e-01 |
| ERF | 122 | 0.6152 | 0.0128 | 9.229e-01 |
| FHS | 4710 | 0.5682 | 0.0176 | 3.988e-01 |
| GenR | 265 | 0.5598 | 0.0607 | 4.880e-01 |
| HUNT | 851 | 0.6256 | -0.0187 | 7.090e-01 |
| LBC1936 | 577 | 0.5803 | 0.1414 | 1.779e-02 |
| LIFE | 1825 | 0.6219 | 0.036 | 2.920e-01 |
| LLS | 355 | 0.578 | -0.0068 | 9.294e-01 |
| MethCT. | 114 | 0.576 | -0.0736 | 5.828e-01 |
| NOMAS | 141 | 0.5063 | -0.1808 | 1.314e-01 |
| PoznanMS | 204 | 0.6461 | -0.1035 | 3.172e-01 |
| PROSPER | 315 | 0.5896 | -0.1308 | 1.062e-01 |
| RS1 | 1103 | 0.5852 | -0.0531 | 2.196e-01 |
| RS2 | 1159 | 0.5894 | 0.0142 | 7.368e-01 |
| RS3 | 2582 | 0.5874 | 0.0902 | 1.413e-03 |
| RUSH1 | 184 | 0.6078 | 0.1168 | 2.753e-01 |
| RUSH2 | 106 | 0.6369 | -0.0983 | 4.931e-01 |
| SHIP | 966 | 0.6222 | -0.0169 | 7.191e-01 |
| SHIP-TREND | 858 | 0.5952 | 0.0106 | 8.287e-01 |
| SYS.adolescents | 980 | 0.5572 | 0.0696 | 1.257e-01 |
| SYS.adults | 594 | 0.5654 | 0.0283 | 6.293e-01 |
| TREND | 1084 | 0.6111 | -0.0614 | 1.637e-01 |
| UKB | 14571 | 0.5915 | 0.0342 | 4.149e-03 |
| WHICAP | 83 | 0.5251 | -0.0157 | 9.201e-01 |
| **Non-European** | | | | |
| ARIC.AA | 675 | 0.538 | 0.0584 | 2.850e-01 |
| CARDIA.AA | 132 | 0.525 | 0.1078 | 3.817e-01 |
| CHAP.black | 321 | 0.6082 | 0.077 | 3.413e-01 |
| FBIRN | 275 | 0.5717 | -0.1799 | 3.783e-02 |
| NOMAS.black | 168 | 0.5713 | 0.1541 | 1.639e-01 |
| NOMAS.hispanic | 718 | 0.4691 | 0.0554 | 2.954e-01 |
| SCES | 209 | 0.8831 | -0.0863 | 5.707e-01 |
| SIMES | 201 | 0.8056 | 0.0854 | 4.887e-01 |
| WHICAP.black | 60 | 0.541 | 0.3211 | 8.538e-02 |
| **Summary European MA** | **74813** | **0.5879** | **0.029** | **4.849e-08** |
| **Summary African MA** | **1356** | **0.5576** | **0.091** | **0.01918** |
| **Summary Asian MA** | **1298** | **0.8698** | **0.077** | **0.1895** |
| **Summary Transancestral MA** | **78574** | **0.5909** | **0.03** | **8.556e-09** |

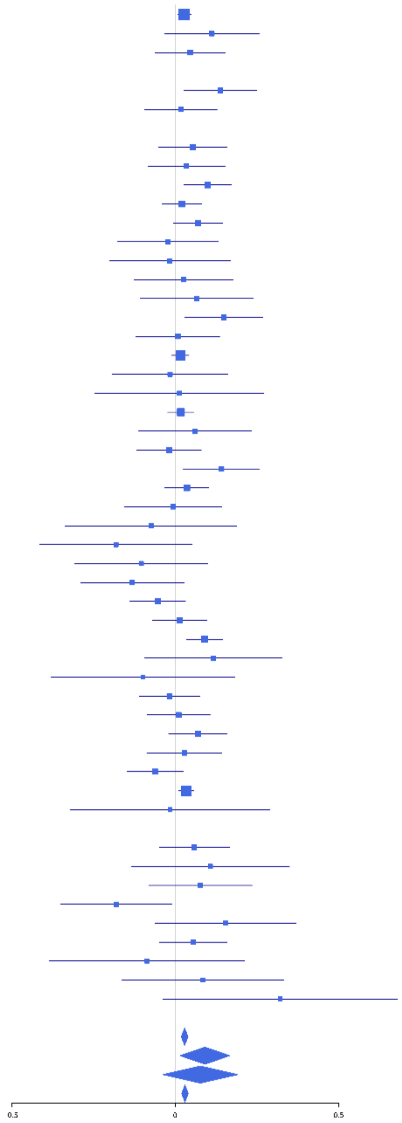

**rs10958478 (A); chr8:57198009; model 1 (I2=0.4, HetP=0.4652)**

| Study | N_total | EAF | Beta | P |
|---|---|---|---|---|
| **Head circumference** | | | | |
| Haworth | 18881 | 0.7754 | -0.0182 | 1.414e-01 |
| HKOS | 888 | 0.8108 | 0.085 | 1.606e-01 |
| VIDI | 730 | 0.7099 | -0.0944 | 1.021e-01 |
| **CT-based** | | | | |
| ESS | 641 | 0.757 | -0.1483 | 2.271e-02 |
| UCL | 642 | 0.7633 | -0.0476 | 4.686e-01 |
| **MR-based** | | | | |
| 1000Brains | 751 | 0.7826 | 0.0469 | 4.538e-01 |
| 3C-Bordeaux | 564 | 0.7794 | -0.0069 | 9.233e-01 |
| 3c-Dijon | 1499 | 0.7851 | -0.0545 | 2.202e-01 |
| AGES | 2226 | 0.7755 | -0.0119 | 7.411e-01 |
| ARIC.EA | 1390 | 0.7682 | -0.0362 | 4.203e-01 |
| ASPS-Family | 338 | 0.7733 | 0.008 | 9.303e-01 |
| ASRB | 233 | 0.7678 | -0.006 | 9.565e-01 |
| CARDIA.EA | 349 | 0.7706 | 0.0051 | 9.544e-01 |
| CHAP | 261 | 0.7627 | -0.2123 | 4.005e-02 |
| CHS | 544 | 0.8054 | 0.1153 | 1.323e-01 |
| ENIGMA | 11373 | 0.7624 | -0.0538 | 5.515e-04 |
| EPOZ | 251 | 0.7448 | -0.2291 | 2.524e-02 |
| FHS | 4710 | 0.7705 | -0.031 | 2.058e-01 |
| GenR | 265 | 0.7809 | -0.1399 | 1.828e-01 |
| HUNT | 851 | 0.7621 | -0.0025 | 9.650e-01 |
| LBC1936 | 577 | 0.7822 | -0.1409 | 4.829e-02 |
| LIFE | 1825 | 0.7659 | -0.0717 | 6.686e-02 |
| LLS | 355 | 0.7254 | -0.2462 | 3.413e-03 |
| MethCT. | 114 | 0.8455 | -0.0764 | 6.765e-01 |
| NOMAS | 141 | 0.7897 | -0.1366 | 3.512e-01 |
| PoznanMS | 204 | 0.7308 | -0.134 | 2.299e-01 |
| PROSPER | 315 | 0.7705 | -0.0192 | 8.392e-01 |
| RS1 | 1103 | 0.7446 | -0.0142 | 7.717e-01 |
| RS2 | 1159 | 0.7488 | -0.0699 | 1.442e-01 |
| RS3 | 2582 | 0.7426 | -0.0401 | 2.074e-01 |
| RUSH1 | 184 | 0.7605 | 0.02 | 8.696e-01 |
| RUSH2 | 106 | 0.7358 | 0.1475 | 3.459e-01 |
| SHIP | 966 | 0.7811 | -0.0166 | 7.634e-01 |
| SHIP-TREND | 858 | 0.7582 | 0.0048 | 9.316e-01 |
| SYS.adolescents | 980 | 0.7828 | 0.0259 | 6.368e-01 |
| SYS.adults | 594 | 0.7886 | -0.0146 | 8.372e-01 |
| TREND | 1084 | 0.7605 | 0.0209 | 6.780e-01 |
| UKB | 14571 | 0.7735 | -0.0415 | 2.996e-03 |
| WHICAP | 83 | 0.8029 | 0.0505 | 7.964e-01 |
| **Non-European** | | | | |
| ARIC.AA | 675 | 0.8962 | 0.0113 | 8.992e-01 |
| CARDIA.AA | 132 | 0.892 | -0.1459 | 4.620e-01 |
| CHAP.black | 321 | 0.8859 | -0.0911 | 4.632e-01 |
| FBIRN | 275 | 0.8345 | 0.0176 | 8.783e-01 |
| NOMAS.black | 168 | 0.9004 | 0.0499 | 7.846e-01 |
| NOMAS.hispanic | 718 | 0.8681 | 0.0468 | 5.487e-01 |
| SCES | 209 | 0.8167 | -0.1708 | 1.766e-01 |
| SIMES | 201 | 0.7963 | -0.0462 | 6.690e-01 |
| WHICAP.black | 60 | 0.85 | -0.0601 | 8.150e-01 |
| **Summary European MA** | **74186** | **0.7691** | **-0.036** | **8.111e-09** |
| **Summary African MA** | **1356** | **0.8918** | **-0.028** | **0.6526** |
| **Summary Asian MA** | **1298** | **0.8095** | **0.022** | **0.6526** |
| **Summary Transancestral MA** | **77947** | **0.7732** | **-0.034** | **2.463e-08** |

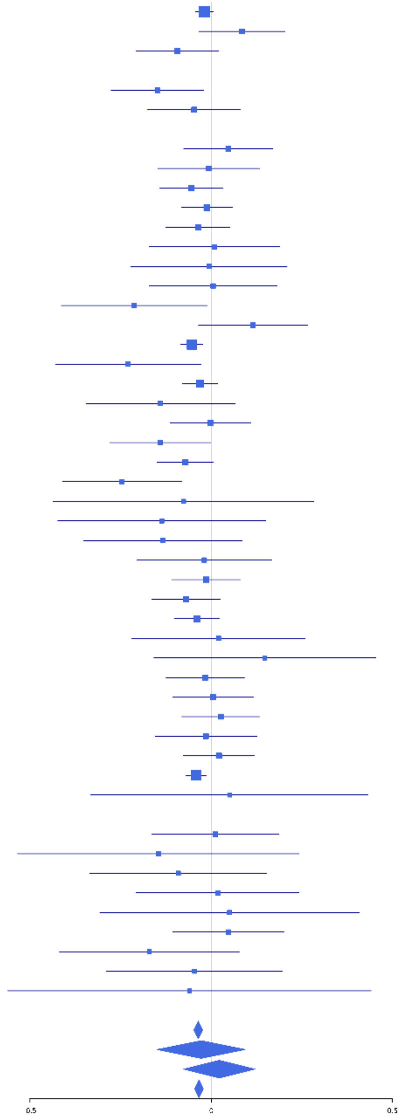

**rs28457693 (A); chr9:98217348; model 1 (I2=0, HetP=0.548)**

| Study | N_total | EAF | Beta | P |
|---|---|---|---|---|
| **Head circumference** | | | | |
| Haworth | 18881 | 0.8932 | -0.0335 | 4.512e-02 |
| HKOS | 888 | 0.7937 | -0.0278 | 6.353e-01 |
| VIDI | 746 | 0.8879 | -0.0852 | 2.994e-01 |
| **CT-based** | | | | |
| ESS | 641 | 0.8962 | -0.1569 | 8.673e-02 |
| UCL | 642 | 0.895 | 0.0496 | 5.856e-01 |
| **MR-based** | | | | |
| 1000Brains | 751 | 0.885 | -0.04 | 6.209e-01 |
| 3C-Bordeaux | 564 | 0.9027 | 0.0563 | 5.757e-01 |
| 3c-Dijon | 1499 | 0.8789 | -0.0873 | 1.190e-01 |
| AGES | 2226 | 0.835 | 0.0439 | 2.774e-01 |
| ARIC.EA | 1390 | 0.8843 | -0.0186 | 7.536e-01 |
| ASPS-Family | 338 | 0.876 | -0.3746 | 1.328e-03 |
| CARDIA.EA | 349 | 0.8966 | -0.0904 | 4.671e-01 |
| CHAP | 261 | 0.875 | -0.1408 | 2.883e-01 |
| CHS | 544 | 0.8905 | 0.0443 | 6.481e-01 |
| ENIGMA | 11373 | 0.8914 | -0.0519 | 1.485e-02 |
| EPOZ | 251 | 0.8893 | -0.0623 | 6.613e-01 |
| FHS | 4710 | 0.8962 | -0.0803 | 1.750e-02 |
| GenR | 265 | 0.8598 | -0.1026 | 4.122e-01 |
| HUNT | 851 | 0.8732 | -0.1246 | 8.704e-02 |
| LIFE | 1825 | 0.871 | -0.0524 | 2.886e-01 |
| LLS | 355 | 0.8776 | 0.0623 | 5.867e-01 |
| MethCT. | 114 | 0.7376 | 0.1769 | 2.400e-01 |
| NOMAS | 141 | 0.8971 | -0.071 | 7.178e-01 |
| PoznanMS | 204 | 0.8223 | 0.028 | 8.293e-01 |
| PROSPER | 315 | 0.8727 | 0.1395 | 2.431e-01 |
| RS1 | 1103 | 0.8908 | -0.1567 | 2.169e-02 |
| RS2 | 1159 | 0.8844 | -0.167 | 1.015e-02 |
| RS3 | 2582 | 0.8927 | 0.021 | 6.413e-01 |
| RUSH1 | 184 | 0.8874 | -0.0168 | 9.192e-01 |
| RUSH2 | 106 | 0.8847 | -0.3301 | 1.279e-01 |
| SHIP | 966 | 0.8748 | 0.0064 | 9.261e-01 |
| SHIP-TREND | 858 | 0.87 | -0.0055 | 9.397e-01 |
| SYS.adolescents | 980 | 0.9038 | -0.0205 | 7.888e-01 |
| SYS.adults | 594 | 0.9036 | -0.0436 | 6.575e-01 |
| TREND | 1084 | 0.8777 | -0.0698 | 2.877e-01 |
| UKB | 14571 | 0.8923 | -0.0621 | 1.010e-03 |
| WHICAP | 83 | 0.8877 | -0.0182 | 9.412e-01 |
| **Non-European** | | | | |
| ARIC.AA | 675 | 0.5822 | -0.0139 | 8.022e-01 |
| CARDIA.AA | 132 | 0.579 | 0.0295 | 8.127e-01 |
| CHAP.black | 321 | 0.583 | 0.0359 | 6.535e-01 |
| FBIRN | 275 | 0.8413 | 0.0546 | 6.404e-01 |
| NOMAS.black | 168 | 0.5838 | 0.0232 | 8.337e-01 |
| NOMAS.hispanic | 718 | 0.7954 | -0.0574 | 3.803e-01 |
| SCES | 209 | 0.7551 | 0.0924 | 4.168e-01 |
| WHICAP.black | 60 | 0.6573 | -0.0792 | 6.821e-01 |
| **Summary European MA** | **73392** | **0.8885** | **-0.047** | **1.051e-08** |
| **Summary African MA** | **1356** | **0.5856** | **0.004** | **0.9177** |
| **Summary Asian MA** | **1097** | **0.7863** | **-0.004** | **0.9424** |
| **Summary Transancestral MA** | **76952** | **0.8805** | **-0.044** | **2.207e-08** |

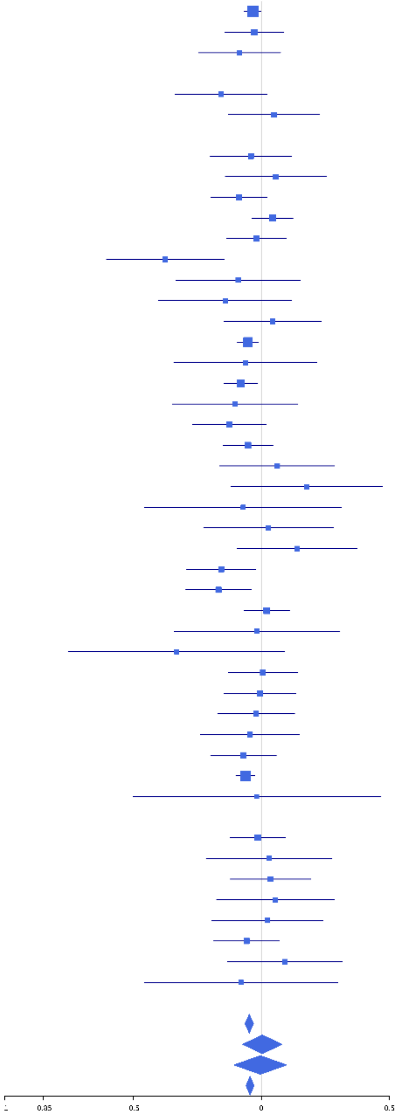

### rs1076148 (T); chr9:136927657; model 1 (I2=0, HetP=0.5931)

| Study | N_total | EAF | Beta | P |
|---|---|---|---|---|
| **Head circumference** | | | | |
| Haworth | 18881 | 0.6926 | -0.0347 | 1.826e-03 |
| HKOS | 888 | 0.4268 | -0.0983 | 4.040e-02 |
| VIDI | 665 | 0.6151 | -0.0105 | 8.517e-01 |
| **CT-based** | | | | |
| ESS | 641 | 0.692 | -0.0113 | 8.516e-01 |
| UCL | 642 | 0.7132 | -0.0479 | 4.380e-01 |
| **MR-based** | | | | |
| 1000Brains | 751 | 0.7162 | -0.0573 | 3.166e-01 |
| 3C-Bordeaux | 564 | 0.7091 | -0.0183 | 7.803e-01 |
| 3c-Dijon | 1499 | 0.6994 | 0.0096 | 8.092e-01 |
| AGES | 2226 | 0.6872 | -0.0322 | 3.191e-01 |
| ARIC.EA | 1390 | 0.7026 | 0.0132 | 7.507e-01 |
| ASPS-Family | 338 | 0.7044 | -0.1967 | 1.961e-02 |
| ASRB | 233 | 0.6866 | 0.0239 | 8.110e-01 |
| CARDIA.EA | 349 | 0.6944 | -0.1549 | 5.945e-02 |
| CHAP | 261 | 0.6992 | -0.0795 | 4.057e-01 |
| CHS | 544 | 0.7169 | 0.0131 | 8.457e-01 |
| DNGS | 516 | 0.6318 | 0.0011 | 9.860e-01 |
| ENIGMA | 11265 | 0.6919 | -0.0259 | 7.309e-02 |
| EPOZ | 251 | 0.6716 | 0.1486 | 1.179e-01 |
| ERF | 122 | 0.7776 | -0.0637 | 6.789e-01 |
| FHS | 4710 | 0.7178 | -0.0227 | 3.205e-01 |
| GenR | 265 | 0.6759 | -0.0304 | 7.428e-01 |
| HUNT | 851 | 0.6792 | -0.075 | 1.487e-01 |
| LBC1936 | 577 | 0.7005 | 0.0163 | 7.997e-01 |
| LIFE | 1825 | 0.6823 | 0.0153 | 6.677e-01 |
| LLS | 355 | 0.6944 | 0.0046 | 9.542e-01 |
| MethCT. | 114 | 0.7077 | -0.1197 | 4.109e-01 |
| NOMAS | 141 | 0.684 | -0.1493 | 2.457e-01 |
| PoznanMS | 204 | 0.7228 | -0.0105 | 9.245e-01 |
| PROSPER | 315 | 0.7143 | 0.0658 | 4.556e-01 |
| RS1 | 1103 | 0.6857 | -0.0698 | 1.280e-01 |
| RS2 | 1159 | 0.6871 | -0.0185 | 6.796e-01 |
| RS3 | 2582 | 0.6893 | -0.0693 | 2.118e-02 |
| RUSH1 | 184 | 0.7223 | -0.0751 | 5.198e-01 |
| RUSH2 | 106 | 0.6887 | 0.3329 | 2.706e-02 |
| SHIP | 966 | 0.6946 | 0 | 1.000e+00 |
| SHIP-TREND | 858 | 0.6894 | -0.0347 | 5.053e-01 |
| SYS.adolescents | 980 | 0.7474 | -0.0874 | 9.281e-02 |
| SYS.adults | 594 | 0.7517 | -0.0569 | 3.962e-01 |
| TREND | 1084 | 0.702 | -0.0524 | 2.648e-01 |
| UKB | 14571 | 0.6965 | -0.0368 | 3.900e-03 |
| WHICAP | 83 | 0.6852 | -0.0986 | 5.568e-01 |
| **Non-European** | | | | |
| ARIC.AA | 675 | 0.6611 | -0.0553 | 3.366e-01 |
| CARDIA.AA | 132 | 0.65 | 0.0375 | 7.708e-01 |
| CHAP.black | 321 | 0.648 | 0.0115 | 8.895e-01 |
| FBIRN | 275 | 0.6843 | 0.1091 | 2.357e-01 |
| NOMAS.black | 168 | 0.6502 | -0.092 | 4.223e-01 |
| NOMAS.hispanic | 718 | 0.7305 | 0.0046 | 9.378e-01 |
| SCES | 209 | 0.4306 | 0.128 | 1.950e-01 |
| SIMES | 201 | 0.5323 | 0.2063 | 4.003e-02 |
| WHICAP.black | 60 | 0.6667 | 0.2481 | 2.059e-01 |
| **Summary European MA** | **74651** | **0.6956** | **-0.031** | **3.043e-08** |
| **Summary African MA** | **1356** | **0.6558** | **-0.022** | **0.5921** |
| **Summary Asian MA** | **1298** | **0.4438** | **-0.014** | **0.7134** |
| **Summary Transancestral MA** | **78412** | **0.691** | **-0.03** | **4.354e-08** |

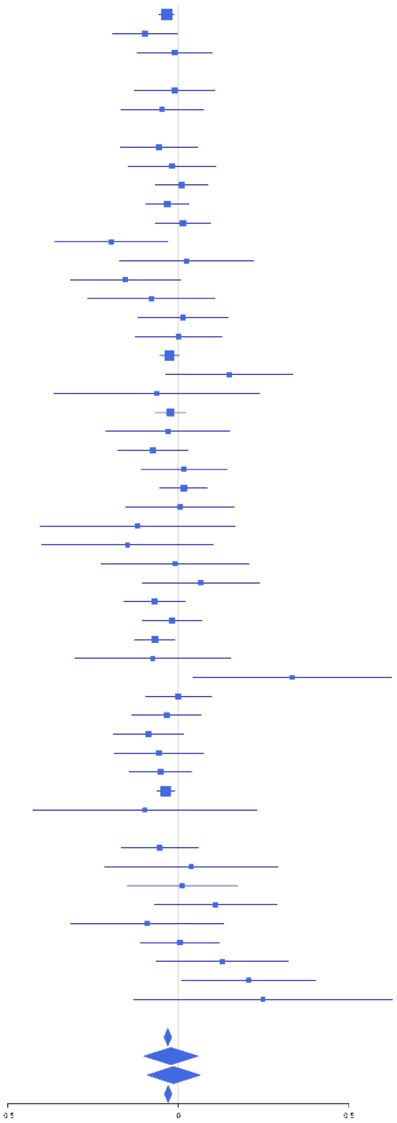

**rs11012732 (A); chr10:21830104; model 1 (I2=24.9, HetP=0.06223)**

| Study | N_total | EAF | Beta | P |
|---|---|---|---|---|
| **Head circumference** | | | | |
| Haworth | 18881 | 0.6424 | 0.0121 | 2.601e-01 |
| VIDI | 755 | 0.6882 | 0.047 | 3.980e-01 |
| **CT-based** | | | | |
| ESS | 641 | 0.6689 | 0.0671 | 2.583e-01 |
| UCL | 642 | 0.663 | -0.0774 | 1.900e-01 |
| **MR-based** | | | | |
| 1000Brains | 751 | 0.6527 | 0.0115 | 8.314e-01 |
| 3C-Bordeaux | 564 | 0.6504 | -0.0432 | 4.894e-01 |
| 3c-Dijon | 1499 | 0.6563 | 0.0237 | 5.380e-01 |
| AGES | 2226 | 0.7149 | 0.0401 | 2.273e-01 |
| ARIC.EA | 1390 | 0.6596 | 0.0443 | 2.687e-01 |
| ASPS-Family | 338 | 0.6462 | 0.233 | 3.772e-03 |
| ASRB | 233 | 0.6537 | 0.0378 | 6.983e-01 |
| CARDIA.EA | 349 | 0.671 | -0.0658 | 4.140e-01 |
| CHAP | 261 | 0.6628 | -0.0485 | 6.006e-01 |
| CHS | 544 | 0.6548 | 0.0362 | 5.706e-01 |
| DNGS | 502 | 0.7604 | -0.0464 | 5.305e-01 |
| ENIGMA | 11373 | 0.6697 | 0.0278 | 4.844e-02 |
| EPOZ | 251 | 0.6813 | -0.1258 | 1.893e-01 |
| ERF | 122 | 0.6239 | -0.1968 | 1.365e-01 |
| FHS | 4710 | 0.6501 | 0.0722 | 8.309e-04 |
| GenR | 265 | 0.6737 | 0.2095 | 2.376e-02 |
| HUNT | 851 | 0.6695 | 0.0276 | 5.923e-01 |
| LBC1936 | 577 | 0.6928 | 0.0425 | 5.056e-01 |
| LIFE | 1825 | 0.6553 | 0.0541 | 1.205e-01 |
| LLS | 355 | 0.6823 | 0.078 | 3.329e-01 |
| MethCT. | 114 | 0.6869 | -0.0437 | 7.596e-01 |
| NOMAS | 141 | 0.6538 | -0.0861 | 4.926e-01 |
| PoznanMS | 204 | 0.6339 | -0.089 | 3.866e-01 |
| PROSPER | 315 | 0.6446 | -0.0169 | 8.388e-01 |
| RS1 | 1103 | 0.6503 | -0.008 | 8.569e-01 |
| RS2 | 1159 | 0.649 | 0.0387 | 3.738e-01 |
| RS3 | 2582 | 0.647 | -0.0198 | 4.972e-01 |
| RUSH1 | 184 | 0.6597 | -0.0575 | 6.018e-01 |
| RUSH2 | 106 | 0.7112 | -0.0761 | 6.171e-01 |
| SHIP | 966 | 0.656 | 0.1118 | 1.982e-02 |
| SHIP-TREND | 858 | 0.6627 | 0.0118 | 8.171e-01 |
| SYS.adolescents | 980 | 0.6368 | -0.0353 | 4.522e-01 |
| SYS.adults | 594 | 0.6448 | 0.1072 | 7.682e-02 |
| TREND | 1084 | 0.654 | -0.014 | 7.566e-01 |
| UKB | 14571 | 0.6698 | 0.0653 | 1.612e-07 |
| WHICAP | 83 | 0.6237 | -0.0011 | 9.942e-01 |
| **Non-European** | | | | |
| ARIC.AA | 675 | 0.7099 | 0.0409 | 4.952e-01 |
| CARDIA.AA | 132 | 0.726 | 0.0951 | 4.911e-01 |
| CHAP.black | 321 | 0.7172 | -0.046 | 6.000e-01 |
| FBIRN | 275 | 0.7065 | 0.0416 | 6.572e-01 |
| NOMAS.black | 168 | 0.6723 | -0.0874 | 4.531e-01 |
| NOMAS.hispanic | 718 | 0.6959 | 0.0279 | 6.269e-01 |
| SCES | 209 | 0.986 | 1.1718 | 4.821e-03 |
| SIMES | 201 | 0.9585 | 0.166 | 4.948e-01 |
| WHICAP.black | 60 | 0.8333 | -0.108 | 6.607e-01 |
| **Summary European MA** | **74835** | **0.6597** | **0.031** | **1.478e-08** |
| **Summary African MA** | **1356** | **0.714** | **0.004** | **0.9327** |
| **Summary Asian MA** | **410** | **0.9725** | **0.532** | **0.01276** |
| **Summary Transcestral MA** | **77708** | **0.6629** | **0.031** | **6.18e-09** |

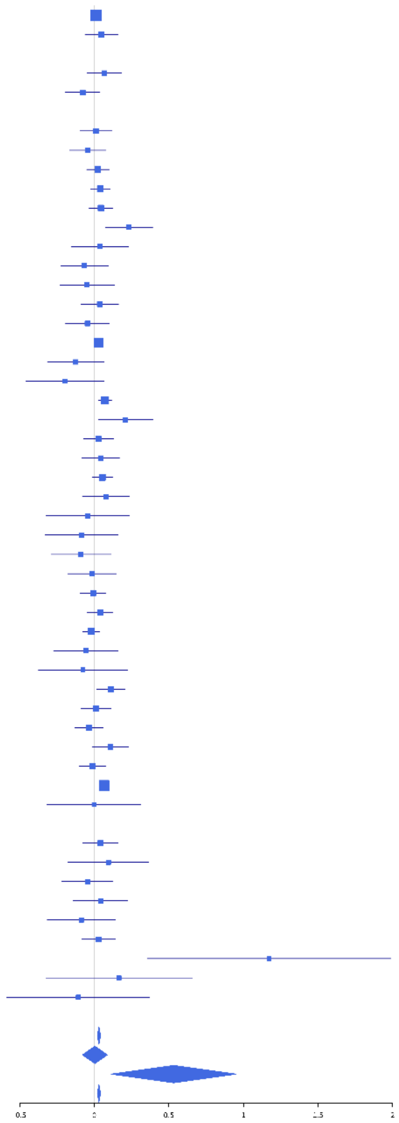

**rs1234214 (A); chr10:89698190; model 1 (I2=0, HetP=0.997)**

| Study | N_total | EAF | Beta | P |
|---|---|---|---|---|
| **Head circumference** | | | | |
| Haworth | 18881 | 0.4153 | -0.0307 | 3.277e-03 |
| HKOS | 888 | 0.4167 | 0.0068 | 8.881e-01 |
| VIDI | 755 | 0.4788 | -0.0134 | 7.940e-01 |
| **CT-based** | | | | |
| ESS | 641 | 0.3974 | -0.032 | 5.752e-01 |
| UCL | 642 | 0.3966 | -0.041 | 4.726e-01 |
| **MR-based** | | | | |
| 1000Brains | 751 | 0.4508 | -0.084 | 1.059e-01 |
| 3C-Bordeaux | 564 | 0.4134 | -0.0447 | 4.596e-01 |
| 3c-Dijon | 1499 | 0.4173 | -0.0666 | 7.235e-02 |
| AGES | 2226 | 0.4128 | -0.0462 | 1.287e-01 |
| ARIC.EA | 1390 | 0.4077 | -0.0219 | 5.698e-01 |
| ASPS-Family | 338 | 0.4218 | -0.0519 | 5.056e-01 |
| ASRB | 233 | 0.4143 | -0.0727 | 4.400e-01 |
| CARDIA.EA | 349 | 0.4076 | -0.0771 | 3.169e-01 |
| CHAP | 261 | 0.4293 | 0.0281 | 7.507e-01 |
| CHS | 544 | 0.4054 | -0.0723 | 2.415e-01 |
| DNGS | 515 | 0.391 | -0.0495 | 4.384e-01 |
| ENIGMA | 11373 | 0.4088 | -0.0136 | 3.147e-01 |
| EPOZ | 251 | 0.4018 | -0.0306 | 7.367e-01 |
| ERF | 122 | 0.4031 | 0.1436 | 2.713e-01 |
| FHS | 4710 | 0.4095 | -0.0289 | 1.675e-01 |
| GenR | 265 | 0.382 | -0.0216 | 8.088e-01 |
| HUNT | 851 | 0.4006 | -0.0625 | 2.063e-01 |
| LBC1936 | 577 | 0.4006 | -0.1512 | 1.185e-02 |
| LIFE | 1825 | 0.4452 | -0.0494 | 1.386e-01 |
| LLS | 355 | 0.4173 | -0.0386 | 6.124e-01 |
| MethCT. | 114 | 0.3176 | -0.1388 | 3.291e-01 |
| NOMAS | 141 | 0.381 | -0.0266 | 8.284e-01 |
| PoznanMS | 204 | 0.4442 | -0.0361 | 7.177e-01 |
| PROSPER | 315 | 0.384 | 0.0984 | 2.298e-01 |
| RS1 | 1103 | 0.4153 | -0.113 | 8.921e-03 |
| RS2 | 1159 | 0.4138 | -0.0761 | 7.129e-02 |
| RS3 | 2582 | 0.4151 | -0.0487 | 8.428e-02 |
| RUSH1 | 184 | 0.3889 | -0.0043 | 9.678e-01 |
| RUSH2 | 106 | 0.4252 | 0.116 | 4.057e-01 |
| SHIP | 966 | 0.4707 | -0.0266 | 5.600e-01 |
| SHIP-TREND | 858 | 0.4748 | -0.0082 | 8.648e-01 |
| SYS.adolescents | 980 | 0.4013 | -0.0227 | 6.230e-01 |
| SYS.adults | 594 | 0.3972 | -0.041 | 4.895e-01 |
| TREND | 1084 | 0.4276 | 0.0082 | 8.510e-01 |
| UKB | 14571 | 0.3894 | -0.0379 | 1.601e-03 |
| WHICAP | 83 | 0.3578 | 0.0113 | 9.441e-01 |
| **Non-European** | | | | |
| ARIC.AA | 675 | 0.183 | -0.0366 | 6.029e-01 |
| CARDIA.AA | 132 | 0.199 | -0.0547 | 7.226e-01 |
| CHAP.black | 321 | 0.1814 | -0.0531 | 6.052e-01 |
| FBIRN | 275 | 0.3737 | -0.0234 | 7.916e-01 |
| NOMAS.black | 168 | 0.1987 | -0.188 | 1.712e-01 |
| NOMAS.hispanic | 718 | 0.2908 | -0.0074 | 8.986e-01 |
| SCES | 209 | 0.4386 | 0.0725 | 4.614e-01 |
| SIMES | 201 | 0.4843 | -0.0252 | 8.003e-01 |
| WHICAP.black | 60 | 0.15 | -0.0665 | 7.956e-01 |
| **Summary European MA** | **74848** | **0.4105** | **-0.034** | **1.15e-10** |
| **Summary African MA** | **1356** | **0.1847** | **-0.063** | **0.2057** |
| **Summary Asian MA** | **1298** | **0.4307** | **0.012** | **0.7548** |
| **Summary Transancestral MA** | **78609** | **0.4055** | **-0.033** | **9.129e-11** |

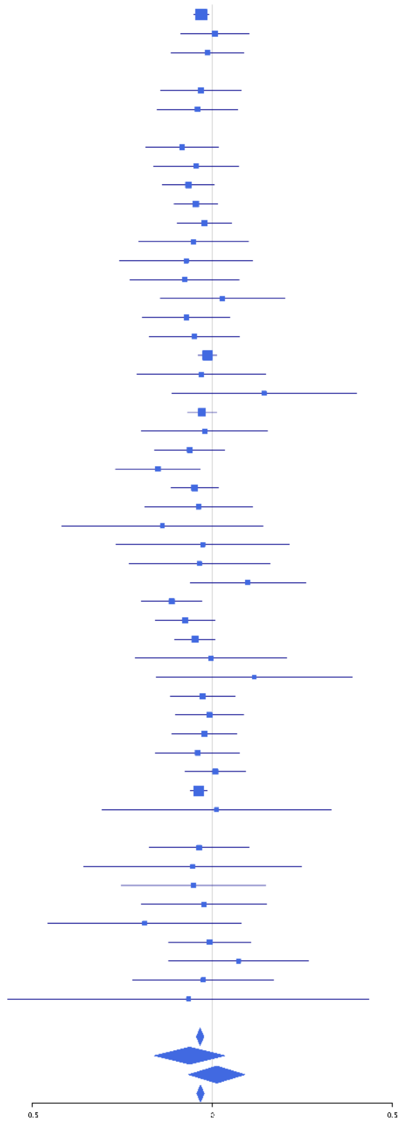

**rs1628768 (T); chr10:105012994; model 1 (I2=23.7, HetP=0.07286)**

| Study | N_total | EAF | Beta | P |
|---|---|---|---|---|
| **Head circumference** | | | | |
| Haworth | 18881 | 0.7642 | -0.032 | 8.163e-03 |
| HKOS | 888 | 0.897 | -0.0165 | 8.320e-01 |
| VIDI | 651 | 0.7612 | -0.0571 | 3.803e-01 |
| **CT-based** | | | | |
| ESS | 641 | 0.7475 | 0.0033 | 9.595e-01 |
| UCL | 642 | 0.7672 | -0.1573 | 1.722e-02 |
| **MR-based** | | | | |
| 1000Brains | 751 | 0.7843 | -0.1704 | 6.762e-03 |
| 3C-Bordeaux | 564 | 0.752 | -0.0658 | 3.399e-01 |
| 3c-Dijon | 1499 | 0.7552 | -0.0677 | 1.107e-01 |
| AGES | 2226 | 0.7501 | -0.1315 | 1.444e-04 |
| ARIC.EA | 1390 | 0.7699 | 0.0388 | 3.897e-01 |
| ASPS-Family | 338 | 0.7984 | 0.0532 | 5.792e-01 |
| ASRB | 233 | 0.735 | -0.1655 | 1.162e-01 |
| CARDIA.EA | 349 | 0.7603 | -0.0804 | 3.646e-01 |
| CHAP | 261 | 0.791 | -0.0072 | 9.468e-01 |
| CHS | 544 | 0.762 | 0.1491 | 3.621e-02 |
| ENIGMA | 11373 | 0.7567 | -0.0502 | 1.160e-03 |
| EPOZ | 251 | 0.7605 | 0.0365 | 7.272e-01 |
| ERF | 122 | 0.8287 | 0.0875 | 6.068e-01 |
| FHS | 4710 | 0.7564 | -0.0666 | 5.531e-03 |
| GenR | 265 | 0.7822 | -0.0963 | 3.603e-01 |
| HUNT | 851 | 0.724 | -0.1225 | 2.387e-02 |
| LBC1936 | 577 | 0.77 | -0.0248 | 7.226e-01 |
| LIFE | 1825 | 0.7734 | -0.1537 | 1.053e-04 |
| LLS | 355 | 0.7686 | -0.1305 | 1.427e-01 |
| MethCT. | 114 | 0.8637 | 0.2258 | 2.419e-01 |
| NOMAS | 141 | 0.7752 | -0.1498 | 2.958e-01 |
| PoznanMS | 204 | 0.7855 | 0.1166 | 3.336e-01 |
| PROSPER | 315 | 0.7523 | -0.0288 | 7.552e-01 |
| RS1 | 1103 | 0.7571 | -0.1215 | 1.439e-02 |
| RS2 | 1159 | 0.7704 | -0.0952 | 5.384e-02 |
| RS3 | 2582 | 0.7606 | -0.0962 | 3.190e-03 |
| RUSH1 | 184 | 0.7736 | -0.074 | 5.534e-01 |
| RUSH2 | 106 | 0.7541 | -0.2938 | 6.845e-02 |
| SHIP | 966 | 0.7677 | -0.0917 | 8.910e-02 |
| SHIP-TREND | 858 | 0.7812 | -0.0531 | 3.631e-01 |
| SYS.adolescents | 980 | 0.7347 | -0.019 | 7.102e-01 |
| SYS.adults | 594 | 0.741 | -0.0234 | 7.232e-01 |
| TREND | 1084 | 0.7641 | -0.0076 | 8.801e-01 |
| UKB | 14571 | 0.7655 | -0.0688 | 6.439e-07 |
| WHICAP | 83 | 0.7218 | -0.3648 | 3.845e-02 |
| **Non-European** | | | | |
| ARIC.AA | 675 | 0.955 | -0.0246 | 8.515e-01 |
| CARDIA.AA | 132 | 0.946 | 0.3175 | 2.438e-01 |
| CHAP.black | 321 | 0.9444 | -0.1352 | 4.332e-01 |
| FBIRN | 275 | 0.8003 | -0.0662 | 5.354e-01 |
| NOMAS.black | 168 | 0.9315 | -0.1706 | 4.306e-01 |
| NOMAS.hispanic | 718 | 0.8634 | -0.0851 | 2.685e-01 |
| SCES | 209 | 0.8683 | -0.0903 | 5.329e-01 |
| SIMES | 201 | 0.8361 | -0.0032 | 9.757e-01 |
| WHICAP.black | 60 | 0.9072 | 0.1686 | 5.943e-01 |
| **Summary European MA** | **74229** | **0.7619** | **-0.058** | **1.709e-21** |
| **Summary African MA** | **1356** | **0.9466** | **-0.027** | **0.7527** |
| **Summary Asian MA** | **1298** | **0.8829** | **-0.027** | **0.6616** |
| **Summary Transancestral MA** | **77990** | **0.7684** | **-0.057** | **2.419e-21** |

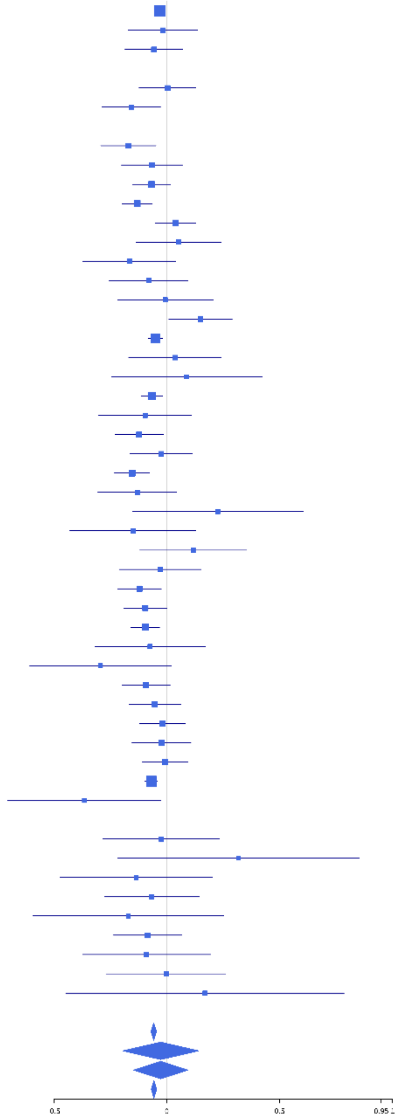

**rs61882777 (T); chr10:111962102; model 1 (I2=9.7, HetP=0.2817)**

| Study | N_total | EAF | Beta | P |
|---|---|---|---|---|
| **Head circumference** | | | | |
| Haworth | 18881 | 0.8968 | 0.0604 | 3.458e-04 |
| HKOS | 888 | 0.6999 | 0.0251 | 6.281e-01 |
| VIDI | 734 | 0.8903 | -0.02 | 8.109e-01 |
| **CT-based** | | | | |
| ESS | 641 | 0.8951 | 0.0734 | 4.210e-01 |
| UCL | 642 | 0.9028 | 0.1566 | 9.655e-02 |
| **MR-based** | | | | |
| 1000Brains | 751 | 0.8953 | -0.0685 | 4.168e-01 |
| 3C-Bordeaux | 564 | 0.8924 | -0.0183 | 8.493e-01 |
| 3c-Dijon | 1499 | 0.8994 | 0.1519 | 1.239e-02 |
| AGES | 2226 | 0.9122 | -0.0218 | 6.808e-01 |
| ARIC.EA | 1390 | 0.9033 | -0.016 | 8.027e-01 |
| ASPS-Family | 338 | 0.894 | -0.0497 | 6.907e-01 |
| ASRB | 233 | 0.9058 | 0.1952 | 2.196e-01 |
| CARDIA.EA | 349 | 0.8911 | 0.086 | 4.787e-01 |
| CHAP | 261 | 0.8961 | 0.0181 | 9.001e-01 |
| CHS | 544 | 0.8871 | 0.0205 | 8.308e-01 |
| ENIGMA | 11373 | 0.9002 | 0.0413 | 6.180e-02 |
| EPOZ | 251 | 0.9136 | -0.0704 | 6.579e-01 |
| ERF | 122 | 0.954 | 0.2091 | 4.941e-01 |
| FHS | 4710 | 0.8875 | 0.0803 | 1.377e-02 |
| GenR | 265 | 0.8845 | 0.0727 | 5.930e-01 |
| HUNT | 851 | 0.9125 | 0.3126 | 2.680e-04 |
| LBC1936 | 577 | 0.9285 | 0.0345 | 7.627e-01 |
| LIFE | 1825 | 0.889 | -0.0407 | 4.402e-01 |
| LLS | 355 | 0.9008 | 0.2118 | 9.152e-02 |
| MethCT. | 114 | 0.8955 | 0.0197 | 9.274e-01 |
| NOMAS | 141 | 0.8406 | 0.1833 | 2.617e-01 |
| PoznanMS | 204 | 0.878 | -5e-04 | 9.977e-01 |
| PROSPER | 315 | 0.9157 | -0.1318 | 3.578e-01 |
| RS1 | 1103 | 0.9052 | 0.0484 | 5.053e-01 |
| RS2 | 1159 | 0.911 | 0.1033 | 1.569e-01 |
| RS3 | 2582 | 0.9054 | 0.0951 | 4.541e-02 |
| RUSH1 | 184 | 0.8994 | -0.0582 | 7.376e-01 |
| RUSH2 | 106 | 0.9262 | -0.0743 | 7.778e-01 |
| SHIP | 966 | 0.8963 | 0.1177 | 1.151e-01 |
| SHIP-TREND | 858 | 0.8957 | -0.0075 | 9.247e-01 |
| SYS.adolescents | 980 | 0.9091 | 0.1146 | 1.447e-01 |
| SYS.adults | 594 | 0.902 | 0.1937 | 4.714e-02 |
| TREND | 1084 | 0.8855 | 0.065 | 3.359e-01 |
| UKB | 14571 | 0.9106 | 0.0101 | 6.220e-01 |
| WHICAP | 83 | 0.8511 | 0.2542 | 2.471e-01 |
| **Non-European** | | | | |
| ARIC.AA | 675 | 0.9633 | 0.2212 | 1.264e-01 |
| CHAP.black | 321 | 0.9747 | -0.3345 | 1.841e-01 |
| FBIRN | 275 | 0.8788 | 0.0854 | 5.135e-01 |
| IMH | 37 | 0.6228 | -0.165 | 4.915e-01 |
| NOMAS.black | 168 | 0.9602 | -0.1691 | 5.451e-01 |
| NOMAS.hispanic | 718 | 0.9353 | -0.1804 | 9.295e-02 |
| SCES | 209 | 0.7255 | 0.0361 | 7.421e-01 |
| SIMES | 201 | 0.6871 | 0.0032 | 9.757e-01 |
| WHICAP.black | 60 | 0.9577 | 0.4935 | 2.816e-01 |
| **Summary European MA** | **74312** | **0.9009** | **0.048** | **2.791e-08** |
| **Summary African MA** | **1224** | **0.9656** | **0.052** | **0.6391** |
| **Summary Asian MA** | **1335** | **0.6998** | **0.018** | **0.6726** |
| **Summary Transancestral MA** | **77978** | **0.8987** | **0.045** | **6.059e-08** |

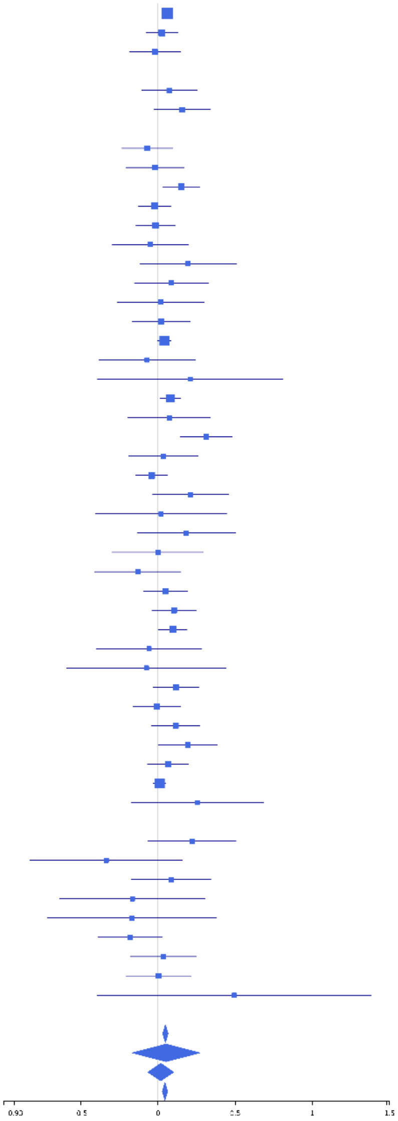

**rs10787231 (A); chr10:112158780; model 1 (I2=9.7, HetP=0.281)**

| Study | N_total | EAF | Beta | P |
|---|---|---|---|---|
| **Head circumference** | | | | |
| Haworth | 18881 | 0.5374 | -0.0359 | 4.885e-04 |
| HKOS | 888 | 0.8108 | -0.0366 | 5.458e-01 |
| VIDI | 742 | 0.5829 | -0.0979 | 6.342e-02 |
| **CT-based** | | | | |
| ESS | 641 | 0.5662 | -0.0402 | 4.753e-01 |
| UCL | 642 | 0.5243 | -0.0093 | 8.675e-01 |
| **MR-based** | | | | |
| 1000Brains | 751 | 0.5343 | -0.0901 | 8.195e-02 |
| 3C-Bordeaux | 564 | 0.5337 | -0.1044 | 8.073e-02 |
| 3c-Dijon | 1499 | 0.529 | 0.0339 | 3.543e-01 |
| AGES | 2226 | 0.5014 | -0.0406 | 1.755e-01 |
| ARIC.EA | 1390 | 0.5228 | 0.0132 | 7.282e-01 |
| ASPS-Family | 338 | 0.5353 | 0.0888 | 2.494e-01 |
| ASRB | 233 | 0.5313 | -0.0946 | 3.089e-01 |
| CARDIA.EA | 349 | 0.5407 | -0.0768 | 3.118e-01 |
| CHAP | 261 | 0.552 | 0.1425 | 1.068e-01 |
| CHS | 544 | 0.542 | -0.0518 | 3.939e-01 |
| DNGS | 508 | 0.6602 | 0.0017 | 9.794e-01 |
| ENIGMA | 11373 | 0.5266 | -0.0301 | 2.353e-02 |
| EPOZ | 251 | 0.5345 | -0.0634 | 4.780e-01 |
| ERF | 122 | 0.4176 | -0.273 | 3.551e-02 |
| FHS | 4710 | 0.5284 | -0.0194 | 3.463e-01 |
| GenR | 265 | 0.6005 | 0.0082 | 9.265e-01 |
| HUNT | 851 | 0.5189 | -0.0881 | 6.950e-02 |
| LBC1936 | 577 | 0.5009 | 0.0737 | 2.110e-01 |
| LIFE | 1825 | 0.537 | -0.0672 | 4.324e-02 |
| LLS | 355 | 0.5 | -0.034 | 6.508e-01 |
| MethCT. | 114 | 0.7167 | -0.1865 | 2.044e-01 |
| NOMAS | 141 | 0.5802 | -0.0533 | 6.591e-01 |
| PoznanMS | 204 | 0.5739 | 0.0397 | 6.912e-01 |
| PROSPER | 315 | 0.5107 | 0.1721 | 3.085e-02 |
| RS1 | 1103 | 0.5307 | -0.0647 | 1.293e-01 |
| RS2 | 1159 | 0.5174 | -0.007 | 8.661e-01 |
| RS3 | 2582 | 0.5334 | -0.0343 | 2.194e-01 |
| RUSH1 | 184 | 0.5462 | 0.0806 | 4.423e-01 |
| RUSH2 | 106 | 0.5082 | -0.1331 | 3.350e-01 |
| SHIP | 966 | 0.5211 | -0.0613 | 1.789e-01 |
| SHIP-TREND | 858 | 0.519 | 0.0105 | 8.275e-01 |
| SYS.adolescents | 980 | 0.5171 | -0.0503 | 2.660e-01 |
| SYS.adults | 594 | 0.516 | -0.0798 | 1.694e-01 |
| TREND | 1084 | 0.5351 | -0.0586 | 1.738e-01 |
| UKB | 14571 | 0.5245 | -0.0232 | 4.753e-02 |
| WHICAP | 83 | 0.5296 | -0.3304 | 3.678e-02 |
| **Non-European** | | | | |
| ARIC.AA | 675 | 0.8175 | -0.0462 | 5.125e-01 |
| CARDIA.AA | 132 | 0.81 | 0.1785 | 2.552e-01 |
| CHAP.black | 321 | 0.7788 | 0.1259 | 1.865e-01 |
| FBIRN | 275 | 0.6433 | -0.1535 | 8.566e-02 |
| NOMAS.black | 168 | 0.8089 | 0.0823 | 5.541e-01 |
| NOMAS.hispanic | 718 | 0.7012 | 0.0294 | 6.102e-01 |
| SCES | 209 | 0.7651 | 0.073 | 5.267e-01 |
| SIMES | 201 | 0.767 | 0.0717 | 5.433e-01 |
| WHICAP.black | 60 | 0.7764 | -0.069 | 7.542e-01 |
| **Summary European MA** | **74828** | **0.5305** | **-0.03** | **4.368e-09** |
| **Summary African MA** | **1356** | **0.8047** | **0.033** | **0.4977** |
| **Summary Asian MA** | **1298** | **0.7967** | **0** | **0.9949** |
| **Summary Transancestral MA** | **78589** | **0.5419** | **-0.029** | **9.441e-09** |

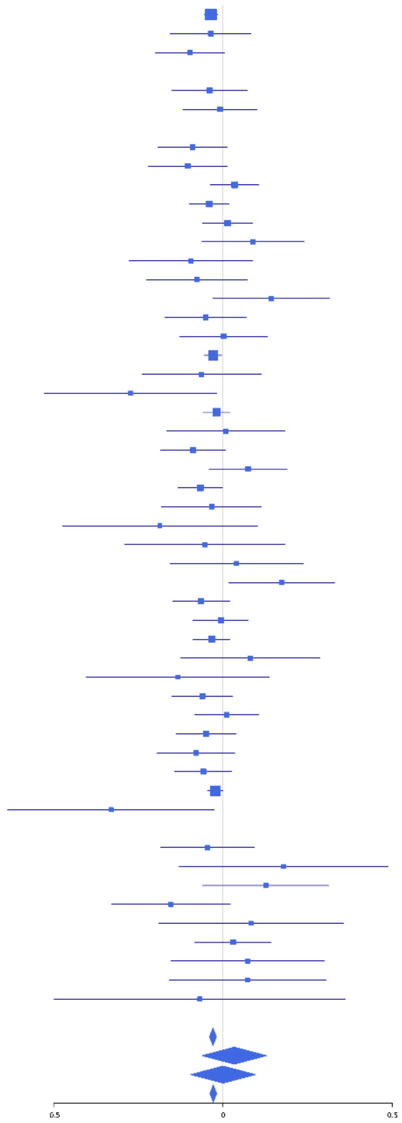

**rs12277225 (T); chr11:68340967; model 1 (I2=3.6, HetP=0.4012)**

| Study | N_total | EAF | Beta | P |
|---|---|---|---|---|
| **Head circumference** | | | | |
| Haworth | 18881 | 0.7369 | 0.0511 | 1.136e-05 |
| HKOS | 888 | 0.7776 | -0.0067 | 9.060e-01 |
| VIDI | 755 | 0.8118 | 0.0686 | 2.977e-01 |
| **CT-based** | | | | |
| ESS | 641 | 0.7353 | 0.0332 | 6.004e-01 |
| UCL | 642 | 0.703 | 0.022 | 7.188e-01 |
| **MR-based** | | | | |
| 1000Brains | 751 | 0.7489 | -0.0127 | 8.308e-01 |
| 3C-Bordeaux | 564 | 0.7245 | 0.0109 | 8.694e-01 |
| 3c-Dijon | 1499 | 0.7323 | -0.0372 | 3.677e-01 |
| AGES | 2226 | 0.7292 | 0.0425 | 2.079e-01 |
| ARIC.EA | 1390 | 0.7338 | 0.0357 | 4.057e-01 |
| ASPS-Family | 338 | 0.7173 | -0.012 | 8.877e-01 |
| ASRB | 233 | 0.7407 | 0.2483 | 1.967e-02 |
| CARDIA.EA | 349 | 0.7486 | -0.056 | 5.212e-01 |
| CHAP | 261 | 0.7298 | 0.1315 | 1.834e-01 |
| CHS | 544 | 0.7472 | 0.1511 | 3.034e-02 |
| DHS | 460 | 0.7417 | 0.0776 | 3.031e-01 |
| DNGS | 516 | 0.7441 | 0.2011 | 5.001e-03 |
| ENIGMA | 11373 | 0.7367 | 0.0213 | 1.573e-01 |
| EPOZ | 251 | 0.7491 | 0.1396 | 1.752e-01 |
| ERF | 122 | 0.763 | -0.0805 | 5.929e-01 |
| FHS | 4710 | 0.7236 | 0.0671 | 3.579e-03 |
| GenR | 265 | 0.7506 | -0.1541 | 1.247e-01 |
| HUNT | 851 | 0.7422 | 0.0513 | 3.543e-01 |
| LBC1936 | 577 | 0.7379 | -0.0082 | 9.018e-01 |
| LIFE | 1825 | 0.7641 | 0.0071 | 8.552e-01 |
| LLS | 355 | 0.7362 | 0.0491 | 5.642e-01 |
| MethCT. | 114 | 0.7811 | -0.2165 | 1.764e-01 |
| NOMAS | 141 | 0.719 | -0.1456 | 2.739e-01 |
| PoznanMS | 204 | 0.7764 | 0.1005 | 3.974e-01 |
| PROSPER | 315 | 0.7442 | 0.0401 | 6.607e-01 |
| RS1 | 1103 | 0.7415 | 0.0515 | 2.902e-01 |
| RS2 | 1159 | 0.7353 | 0.0171 | 7.155e-01 |
| RS3 | 2582 | 0.7455 | 0.001 | 9.755e-01 |
| RUSH1 | 184 | 0.7571 | -0.1307 | 2.836e-01 |
| RUSH2 | 106 | 0.7689 | -0.0026 | 9.873e-01 |
| SHIP | 966 | 0.7458 | 0.023 | 6.597e-01 |
| SHIP-TREND | 858 | 0.7669 | -0.0223 | 6.969e-01 |
| SYS.adolescents | 980 | 0.7501 | 0.0254 | 6.267e-01 |
| SYS.adults | 594 | 0.7583 | -0.0278 | 6.821e-01 |
| TREND | 1084 | 0.7701 | 0.07 | 1.704e-01 |
| UKB | 14571 | 0.7269 | 0.0256 | 5.145e-02 |
| WHICAP | 83 | 0.7902 | 0.0953 | 6.182e-01 |
| **Non-European** | | | | |
| ARIC.AA | 675 | 0.7678 | -0.0976 | 1.299e-01 |
| CARDIA.AA | 132 | 0.752 | 0.1153 | 4.186e-01 |
| CHAP.black | 321 | 0.7449 | -0.0335 | 7.116e-01 |
| FBIRN | 275 | 0.756 | -0.0717 | 4.711e-01 |
| NOMAS.black | 168 | 0.7865 | 0.0493 | 7.116e-01 |
| NOMAS.hispanic | 718 | 0.6917 | 0.0279 | 6.257e-01 |
| SCES | 209 | 0.8158 | 0.2503 | 4.724e-02 |
| SIMES | 201 | 0.7831 | -0.0339 | 7.792e-01 |
| WHICAP.black | 60 | 0.7583 | -0.1874 | 3.832e-01 |
| **Summary European MA** | **75309** | **0.737** | **0.034** | **5.04e-09** |
| **Summary African MA** | **1356** | **0.7627** | **-0.047** | **0.294** |
| **Summary Asian MA** | **1298** | **0.7846** | **0.028** | **0.5564** |
| **Summary Transancestral MA** | **79070** | **0.738** | **0.032** | **2.197e-08** |

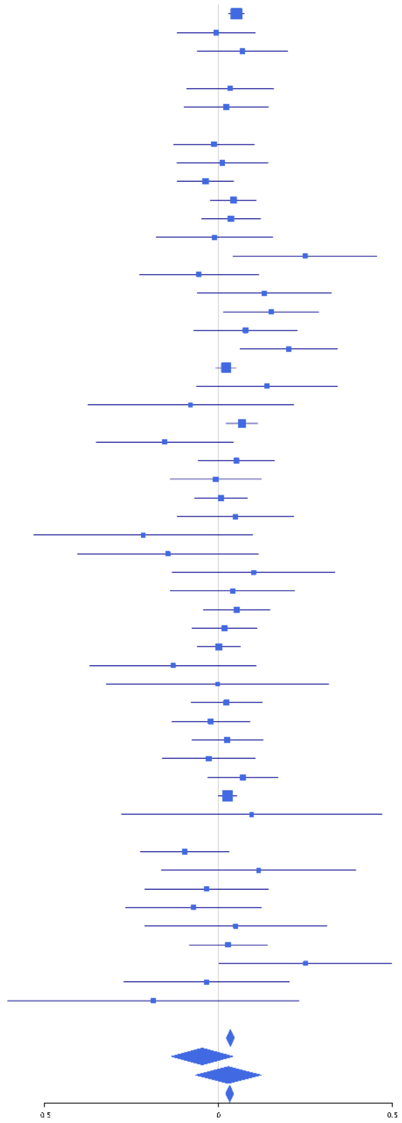

**rs3217870 (T); chr12:4400111; model 1 (I2=0, HetP=0.797)**

| Study | N_total | EAF | Beta | P |
|---|---|---|---|---|
| **Head circumference** | | | | |
| Haworth | 18881 | 0.5993 | -0.0518 | 7.034e-07 |
| HKOS | 888 | 0.75 | 0.0144 | 7.936e-01 |
| VIDI | 699 | 0.6291 | -0.097 | 8.029e-02 |
| **CT-based** | | | | |
| ESS | 641 | 0.6264 | 0.0375 | 5.164e-01 |
| UCL | 642 | 0.5917 | -0.049 | 3.883e-01 |
| **MR-based** | | | | |
| 1000Brains | 751 | 0.6379 | 0.0536 | 3.179e-01 |
| 3C-Bordeaux | 564 | 0.6494 | -0.0205 | 7.426e-01 |
| 3c-Dijon | 1499 | 0.6665 | -0.0095 | 8.065e-01 |
| AGES | 2226 | 0.614 | -0.0639 | 3.803e-02 |
| ARIC.EA | 1390 | 0.6196 | -0.0436 | 2.649e-01 |
| ASPS-Family | 338 | 0.6738 | -0.0479 | 5.589e-01 |
| CHAP | 261 | 0.5933 | -0.0973 | 2.761e-01 |
| CHS | 544 | 0.6289 | -0.1515 | 1.580e-02 |
| ENIGMA | 11373 | 0.6143 | -0.0328 | 1.602e-02 |
| EPOZ | 251 | 0.6419 | -0.1087 | 2.426e-01 |
| ERF | 122 | 0.6814 | -0.0168 | 9.028e-01 |
| FHS | 4710 | 0.6043 | -0.0412 | 5.051e-02 |
| GenR | 265 | 0.6342 | 0.0086 | 9.244e-01 |
| HUNT | 851 | 0.582 | 0.0061 | 9.008e-01 |
| LBC1936 | 577 | 0.6265 | -0.003 | 9.609e-01 |
| LIFE | 1825 | 0.6561 | -0.0028 | 9.365e-01 |
| LLS | 355 | 0.633 | 0.0605 | 4.370e-01 |
| NOMAS | 141 | 0.6169 | 0.0288 | 8.146e-01 |
| PoznanMS | 204 | 0.6799 | -0.1314 | 2.158e-01 |
| PROSPER | 315 | 0.6642 | -0.0138 | 8.698e-01 |
| RS1 | 1103 | 0.6268 | -0.0161 | 7.152e-01 |
| RS2 | 1159 | 0.6322 | -0.0458 | 2.872e-01 |
| RS3 | 2582 | 0.6242 | -0.0574 | 4.583e-02 |
| RUSH1 | 184 | 0.5594 | -0.1909 | 7.075e-02 |
| RUSH2 | 106 | 0.6229 | -0.0179 | 9.000e-01 |
| SHIP | 966 | 0.6731 | -0.0026 | 9.577e-01 |
| SHIP-TREND | 858 | 0.6253 | -0.0911 | 6.799e-02 |
| SYS.adolescents | 980 | 0.6139 | 0.0552 | 2.346e-01 |
| SYS.adults | 594 | 0.6225 | -0.1005 | 9.311e-02 |
| TREND | 1084 | 0.6506 | -0.0916 | 4.218e-02 |
| UKB | 14571 | 0.607 | -0.0363 | 2.506e-03 |
| WHICAP | 83 | 0.6217 | -0.2073 | 1.992e-01 |
| **Non-European** | | | | |
| ARIC.AA | 675 | 0.7568 | -0.0471 | 4.580e-01 |
| CARDIA.AA | 132 | 0.739 | 0.1268 | 3.657e-01 |
| CHAP.black | 321 | 0.7251 | -0.1652 | 6.257e-02 |
| FBIRN | 275 | 0.6502 | -0.1262 | 1.592e-01 |
| IMH | 37 | 0.7456 | -0.2586 | 3.327e-01 |
| NOMAS.black | 168 | 0.7378 | -0.0257 | 8.365e-01 |
| NOMAS.hispanic | 718 | 0.6913 | -0.0272 | 6.334e-01 |
| SCES | 209 | 0.7586 | -0.028 | 8.062e-01 |
| SIMES | 201 | 0.6919 | -0.0335 | 7.283e-01 |
| WHICAP.black | 60 | 0.6923 | 0.1483 | 4.564e-01 |
| **Summary European MA** | **73695** | **0.6141** | **-0.04** | **5.926e-14** |
| **Summary African MA** | **1356** | **0.7424** | **-0.047** | **0.2876** |
| **Summary Asian MA** | **1335** | **0.7425** | **-0.008** | **0.8573** |
| **Summary Transancestral MA** | **77379** | **0.6194** | **-0.04** | **2.478e-14** |

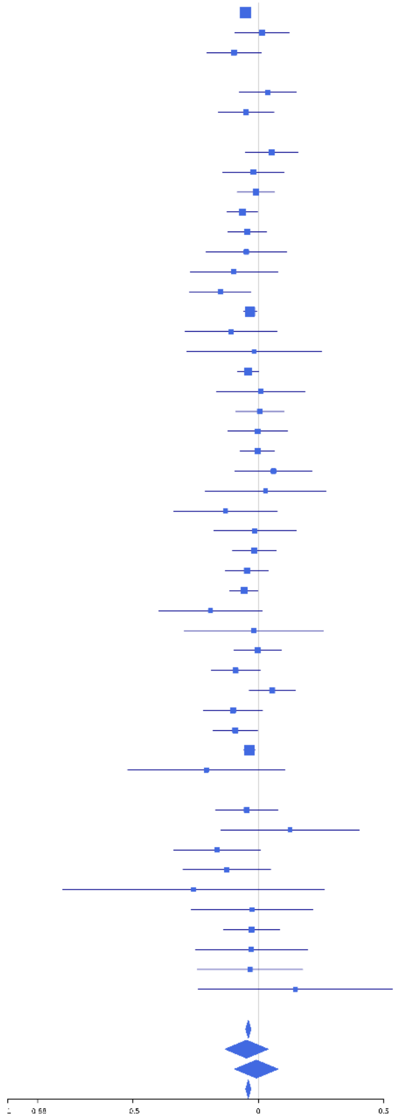

### rs2066827 (T); chr12:12871099; model 1 (I2=0, HetP=0.6487)

| Study | N_total | EAF | Beta | P |
|---|---|---|---|---|
| **Head circumference** | | | | |
| Haworth | 18881 | 0.7594 | 0.0472 | 9.293e-05 |
| HKOS | 888 | 0.9611 | 0.1084 | 3.772e-01 |
| VIDI | 755 | 0.7522 | 0.0243 | 6.836e-01 |
| **CT-based** | | | | |
| ESS | 641 | 0.7396 | 0.0263 | 6.790e-01 |
| UCL | 642 | 0.7813 | 0.0049 | 9.418e-01 |
| **MR-based** | | | | |
| 1000Brains | 751 | 0.762 | 0.135 | 2.615e-02 |
| 3C-Bordeaux | 564 | 0.7653 | 0.1289 | 6.708e-02 |
| 3c-Dijon | 1499 | 0.7632 | 0.0228 | 5.952e-01 |
| AGES | 2226 | 0.7621 | 0.0706 | 4.492e-02 |
| ARIC.EA | 1390 | 0.7763 | 0.0795 | 8.079e-02 |
| ASRB | 233 | 0.7565 | -0.0353 | 7.438e-01 |
| CHAP | 261 | 0.7975 | 0.111 | 3.092e-01 |
| CHS | 544 | 0.7751 | 0.0808 | 2.659e-01 |
| DNGS | 516 | 0.7515 | 0.0254 | 7.243e-01 |
| ENIGMA | 7637 | 0.7698 | 0.0599 | 1.836e-03 |
| EPOZ | 251 | 0.7343 | 0.1037 | 3.049e-01 |
| GenR | 265 | 0.7045 | 0.0412 | 6.648e-01 |
| HUNT | 851 | 0.7408 | 0.0953 | 8.507e-02 |
| LBC1936 | 577 | 0.7968 | 0.1033 | 1.581e-01 |
| LIFE | 1825 | 0.7806 | 0.0029 | 9.428e-01 |
| LLS | 355 | 0.7479 | 0.146 | 9.131e-02 |
| MethCT. | 114 | 0.6465 | 0.0066 | 9.613e-01 |
| PoznanMS | 204 | 0.819 | 0.111 | 3.883e-01 |
| PROSPER | 315 | 0.7492 | 0.0118 | 8.982e-01 |
| RS1 | 1103 | 0.7511 | 0.0566 | 2.507e-01 |
| RS2 | 1159 | 0.7635 | 0.0399 | 4.137e-01 |
| RS3 | 2582 | 0.7604 | 0.0718 | 2.757e-02 |
| RUSH2 | 106 | 0.7925 | -0.0987 | 5.613e-01 |
| SHIP | 966 | 0.7796 | 0.1617 | 3.281e-03 |
| SHIP-TREND | 858 | 0.785 | 0.0143 | 8.083e-01 |
| SYS.adolescents | 980 | 0.7214 | 0.1507 | 2.782e-03 |
| SYS.adults | 594 | 0.7298 | 0.1793 | 6.052e-03 |
| TREND | 1084 | 0.7682 | 0.1203 | 1.830e-02 |
| UKB | 14571 | 0.766 | 0.0553 | 6.334e-05 |
| WHICAP | 83 | 0.7963 | -0.2272 | 2.418e-01 |
| **Non-European** | | | | |
| ARIC.AA | 675 | 0.3339 | -0.0227 | 6.934e-01 |
| CARDIA.AA | 132 | 0.363 | -0.0088 | 9.449e-01 |
| CHAP.black | 321 | 0.324 | 0.0447 | 5.966e-01 |
| NOMAS.black | 168 | 0.4042 | 0.0103 | 9.263e-01 |
| NOMAS.hispanic | 718 | 0.6433 | -0.0457 | 4.066e-01 |
| SCES | 209 | 0.9593 | -0.2028 | 4.131e-01 |
| SIMES | 200 | 0.83 | -0.0047 | 9.724e-01 |
| WHICAP.black | 60 | 0.403 | 0.3959 | 3.807e-02 |
| **Summary European MA** | **65269** | **0.7633** | **0.059** | **2.147e-19** |
| **Summary African MA** | **1356** | **0.3462** | **0.017** | **0.6696** |
| **Summary Asian MA** | **1297** | **0.9406** | **0.032** | **0.6976** |
| **Summary Transancestral MA** | **68754** | **0.757** | **0.055** | **1.311e-18** |

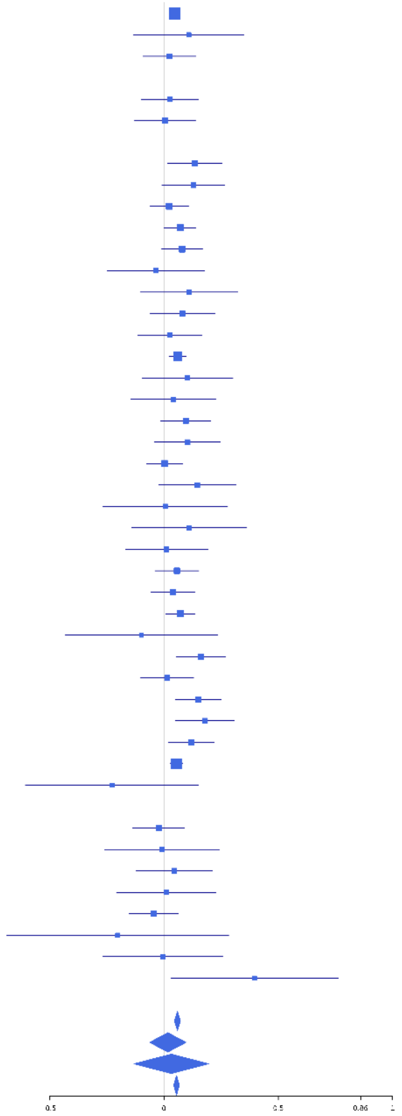

**rs111939932 (T); chr12:53843771; model 1 (I2=11.9, HetP=0.2395)**

| Study | N_total | EAF | Beta | P |
|---|---|---|---|---|
| **Head circumference** | | | | |
| Haworth | 18881 | 0.1912 | 0.054 | 3.546e-05 |
| HKOS | 888 | 0.1498 | 0.0419 | 5.285e-01 |
| VIDI | 746 | 0.1601 | -0.0041 | 9.536e-01 |
| **CT-based** | | | | |
| ESS | 641 | 0.1739 | -0.006 | 9.353e-01 |
| UCL | 642 | 0.1677 | 0.0699 | 3.495e-01 |
| **MR-based** | | | | |
| 1000Brains | 751 | 0.1745 | 0.1343 | 4.852e-02 |
| 3C-Bordeaux | 564 | 0.1693 | 0.178 | 2.533e-02 |
| 3c-Dijon | 1499 | 0.1755 | -0.0731 | 1.281e-01 |
| AGES | 2226 | 0.1686 | 0.0852 | 3.327e-02 |
| ARIC.EA | 1390 | 0.1775 | 0.0672 | 1.758e-01 |
| ASPS-Family | 338 | 0.1766 | 0.0428 | 6.718e-01 |
| ASRB | 233 | 0.1885 | 0.2143 | 7.179e-02 |
| CARDIA.EA | 349 | 0.158 | 0.2213 | 3.291e-02 |
| CHAP | 261 | 0.1824 | -0.0076 | 9.470e-01 |
| CHS | 544 | 0.1689 | 0.0346 | 6.694e-01 |
| DNGS | 511 | 0.1554 | -0.0777 | 3.683e-01 |
| ENIGMA | 11373 | 0.1782 | 0.0494 | 4.382e-03 |
| EPOZ | 251 | 0.1925 | -0.0643 | 5.701e-01 |
| ERF | 122 | 0.1853 | -0.061 | 7.112e-01 |
| FHS | 4710 | 0.17 | 0.0456 | 9.672e-02 |
| GenR | 265 | 0.1682 | 0.1074 | 3.548e-01 |
| HUNT | 851 | 0.195 | 0.1331 | 2.960e-02 |
| LBC1936 | 577 | 0.1896 | -0.0243 | 7.469e-01 |
| LIFE | 1825 | 0.1695 | -0.0103 | 8.147e-01 |
| LLS | 355 | 0.222 | 0.0449 | 6.191e-01 |
| MethCT. | 114 | 0.1466 | 0.0243 | 8.963e-01 |
| NOMAS | 141 | 0.1569 | 0.1619 | 3.245e-01 |
| PoznanMS | 204 | 0.1547 | 0.2193 | 1.091e-01 |
| PROSPER | 315 | 0.1912 | -0.0202 | 8.423e-01 |
| RS1 | 1103 | 0.1892 | 0.0215 | 6.919e-01 |
| RS2 | 1159 | 0.1869 | 0.0082 | 8.776e-01 |
| RS3 | 2582 | 0.1815 | -0.0026 | 9.433e-01 |
| RUSH1 | 184 | 0.1889 | 0.0892 | 5.037e-01 |
| RUSH2 | 106 | 0.2051 | 0.1606 | 3.473e-01 |
| SHIP | 966 | 0.1701 | -0.0233 | 7.002e-01 |
| SHIP-TREND | 858 | 0.1757 | 0.2246 | 4.221e-04 |
| SYS.adolescents | 980 | 0.1141 | -0.0208 | 7.698e-01 |
| SYS.adults | 594 | 0.1187 | 0.0999 | 2.653e-01 |
| TREND | 1084 | 0.1809 | -0.0248 | 6.572e-01 |
| UKB | 14571 | 0.179 | 0.0605 | 7.536e-05 |
| WHICAP | 83 | 0.1662 | 0.0369 | 8.599e-01 |
| **Non-European** | | | | |
| ARIC.AA | 675 | 0.1393 | 0.1201 | 1.265e-01 |
| CARDIA.AA | 132 | 0.138 | 0.1934 | 2.784e-01 |
| CHAP.black | 321 | 0.1422 | 0.1564 | 1.674e-01 |
| FBIRN | 275 | 0.1894 | -0.0665 | 5.418e-01 |
| NOMAS.black | 168 | 0.1282 | 0.3507 | 3.310e-02 |
| NOMAS.hispanic | 718 | 0.1763 | -0.0536 | 4.390e-01 |
| SCES | 209 | 0.1666 | -0.0497 | 7.050e-01 |
| SIMES | 201 | 0.1683 | 0.0735 | 5.758e-01 |
| WHICAP.black | 60 | 0.0936 | -0.2758 | 3.828e-01 |
| **Summary European MA** | **74835** | **0.1794** | **0.048** | **6.864e-13** |
| **Summary African MA** | **1356** | **0.1365** | **0.148** | **0.007943** |
| **Summary Asian MA** | **1298** | **0.1554** | **0.032** | **0.5555** |
| **Summary Transancestral MA** | **78596** | **0.1782** | **0.048** | **2.344e-13** |

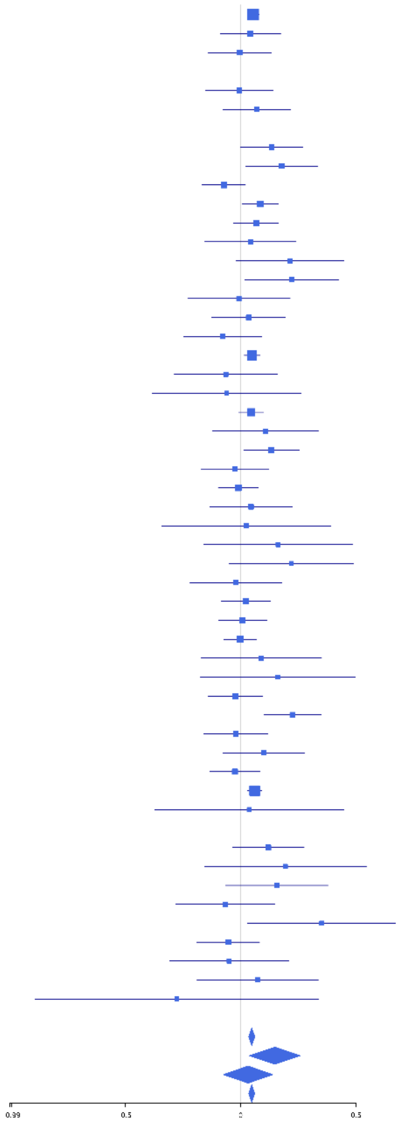

**rs2271194 (A); chr12:56477694; model 1 (I2=23.5, HetP=0.07493)**

| Study | N_total | EAF | Beta | P |
|---|---|---|---|---|
| **Head circumference** | | | | |
| Haworth | 18881 | 0.413 | 0.0411 | 8.032e-05 |
| HKOS | 888 | 0.2359 | -0.0505 | 3.665e-01 |
| VIDI | 750 | 0.4067 | 0.1182 | 2.482e-02 |
| **CT-based** | | | | |
| ESS | 641 | 0.4141 | -9e-04 | 9.872e-01 |
| UCL | 642 | 0.4169 | 0.0731 | 1.968e-01 |
| **MR-based** | | | | |
| 1000Brains | 751 | 0.4005 | 0.1148 | 2.947e-02 |
| 3C-Bordeaux | 564 | 0.4244 | 0.0458 | 4.475e-01 |
| 3c-Dijon | 1499 | 0.4314 | 0.0251 | 4.966e-01 |
| AGES | 2226 | 0.4231 | 0.046 | 1.297e-01 |
| ARIC.EA | 1390 | 0.4173 | 0.0197 | 6.083e-01 |
| ASPS-Family | 338 | 0.4282 | -0.0072 | 9.260e-01 |
| ASRB | 233 | 0.4328 | 0.2144 | 2.275e-02 |
| CARDIA.EA | 349 | 0.4141 | 0.1089 | 1.566e-01 |
| CHAP | 261 | 0.3819 | 0.0637 | 4.801e-01 |
| CHS | 544 | 0.4318 | -0.0646 | 2.912e-01 |
| ENIGMA | 11373 | 0.4151 | 0.025 | 6.286e-02 |
| EPOZ | 251 | 0.41 | -0.2454 | 6.860e-03 |
| ERF | 122 | 0.4676 | -0.0495 | 6.993e-01 |
| FHS | 4710 | 0.4078 | 0.0564 | 7.166e-03 |
| GenR | 265 | 0.4248 | 0.0562 | 5.230e-01 |
| HUNT | 851 | 0.4143 | 0.0461 | 3.491e-01 |
| LBC1936 | 577 | 0.4473 | -0.0191 | 7.467e-01 |
| LIFE | 1825 | 0.3913 | -0.0116 | 7.321e-01 |
| LLS | 355 | 0.4472 | 0.0324 | 6.676e-01 |
| MethCT. | 114 | 0.5006 | -0.085 | 5.207e-01 |
| NOMAS | 141 | 0.4291 | 0.1927 | 1.116e-01 |
| PoznanMS | 204 | 0.3732 | -0.0957 | 3.499e-01 |
| PROSPER | 315 | 0.3922 | -0.083 | 3.094e-01 |
| RS1 | 1103 | 0.4134 | 0.0421 | 3.301e-01 |
| RS2 | 1159 | 0.4262 | 0.0255 | 5.448e-01 |
| RS3 | 2582 | 0.4035 | -0.0115 | 6.846e-01 |
| RUSH1 | 184 | 0.4177 | -0.1675 | 1.147e-01 |
| RUSH2 | 106 | 0.4388 | 0.0933 | 5.021e-01 |
| SHIP | 966 | 0.4027 | 0.0332 | 4.750e-01 |
| SHIP-TREND | 858 | 0.3934 | 0.0209 | 6.731e-01 |
| SYS.adolescents | 980 | 0.3939 | 0.0845 | 6.777e-02 |
| SYS.adults | 594 | 0.4 | -0.0707 | 2.324e-01 |
| TREND | 1084 | 0.4147 | 0.0177 | 6.849e-01 |
| UKB | 14571 | 0.4345 | 0.0632 | 8.937e-08 |
| WHICAP | 83 | 0.4554 | -0.0295 | 8.507e-01 |
| **Non-European** | | | | |
| ARIC.AA | 675 | 0.6672 | 0.0243 | 6.744e-01 |
| CARDIA.AA | 132 | 0.68 | 0.0379 | 7.740e-01 |
| CHAP.black | 321 | 0.6673 | -0.0579 | 4.900e-01 |
| FBIRN | 275 | 0.4113 | -0.0447 | 6.063e-01 |
| NOMAS.black | 168 | 0.6678 | -0.1294 | 2.656e-01 |
| NOMAS.hispanic | 718 | 0.432 | 0.1193 | 2.547e-02 |
| SCES | 209 | 0.1666 | 0.1414 | 2.815e-01 |
| SIMES | 201 | 0.2201 | -0.0872 | 4.567e-01 |
| WHICAP.black | 60 | 0.5475 | 0.0459 | 8.039e-01 |
| **Summary European MA** | **74328** | **0.4172** | **0.038** | **6.367e-13** |
| **Summary African MA** | **1356** | **0.6632** | **-0.012** | **0.7722** |
| **Summary Asian MA** | **1298** | **0.2223** | **-0.029** | **0.5434** |
| **Summary Transancestral MA** | **78089** | **0.4185** | **0.036** | **1.669e-12** |

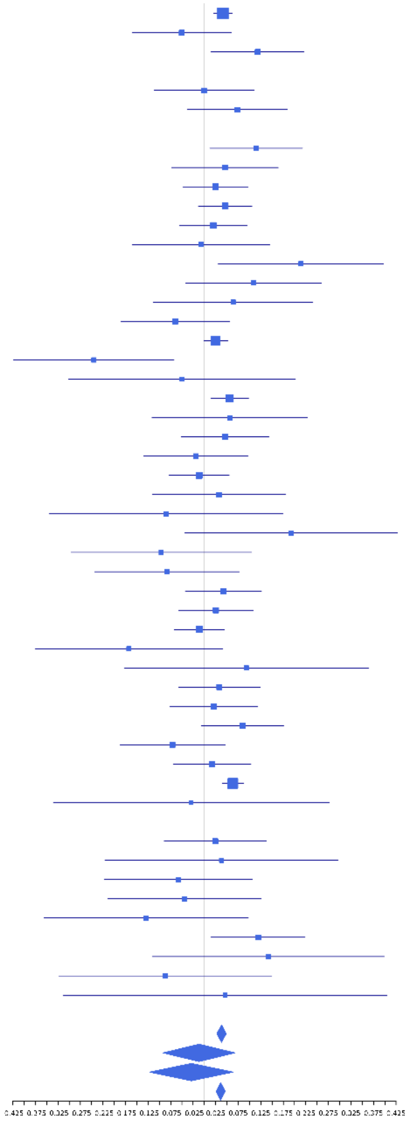

**rs17178006 (T); chr12:65718299; model 1 (I2=5.3, HetP=0.3764)**

| Study | N_total | EAF | Beta | P |
|---|---|---|---|---|
| **Head circumference** | | | | |
| Haworth | 18881 | 0.8859 | 0.0533 | 9.746e-04 |
| VIDI | 755 | 0.9343 | 0.0644 | 5.352e-01 |
| **CT-based** | | | | |
| ESS | 641 | 0.8964 | 0.0591 | 5.192e-01 |
| UCL | 642 | 0.8652 | 0.0597 | 4.657e-01 |
| **MR-based** | | | | |
| 1000Brains | 751 | 0.913 | -0.0626 | 4.941e-01 |
| 3C-Bordeaux | 564 | 0.9048 | 0.1079 | 2.883e-01 |
| 3c-Dijon | 1499 | 0.9101 | 0.1502 | 1.869e-02 |
| AGES | 2226 | 0.9052 | 0.0559 | 2.745e-01 |
| ARIC.EA | 1390 | 0.9 | -0.0027 | 9.666e-01 |
| ASPS-Family | 338 | 0.9249 | 0.2384 | 1.022e-01 |
| CARDIA.EA | 349 | 0.9044 | 0.1954 | 1.289e-01 |
| CHS | 544 | 0.8905 | 0.0957 | 3.243e-01 |
| ENIGMA | 11373 | 0.8908 | 0.0395 | 6.305e-02 |
| EPOZ | 251 | 0.8803 | 0.1844 | 1.800e-01 |
| ERF | 122 | 0.9111 | 0.1286 | 5.671e-01 |
| FHS | 4710 | 0.8992 | 0.0925 | 6.878e-03 |
| GenR | 265 | 0.9254 | 0.2767 | 9.410e-02 |
| HUNT | 851 | 0.9144 | 0.0584 | 5.003e-01 |
| LBC1936 | 577 | 0.8757 | 0.1205 | 1.767e-01 |
| LIFE | 1825 | 0.9145 | -0.0106 | 8.583e-01 |
| LLS | 355 | 0.8892 | 0.0801 | 5.026e-01 |
| NOMAS | 141 | 0.9305 | 0.4683 | 4.751e-02 |
| PoznanMS | 204 | 0.9293 | 0.3205 | 9.693e-02 |
| PROSPER | 315 | 0.9034 | -0.0599 | 6.572e-01 |
| RS1 | 1103 | 0.8956 | -0.0435 | 5.320e-01 |
| RS2 | 1159 | 0.8936 | 0.0292 | 6.644e-01 |
| RS3 | 2582 | 0.8913 | 0.0493 | 2.698e-01 |
| SHIP | 966 | 0.9072 | 0.0597 | 4.464e-01 |
| SHIP-TREND | 858 | 0.8992 | -0.0276 | 7.310e-01 |
| SYS.adolescents | 980 | 0.8888 | -0.159 | 2.683e-02 |
| SYS.adults | 594 | 0.8891 | 0.0507 | 5.832e-01 |
| TREND | 1084 | 0.9133 | -0.1251 | 1.012e-01 |
| UKB | 14571 | 0.8804 | 0.0609 | 7.356e-04 |
| WHICAP | 83 | 0.9318 | -0.0403 | 8.964e-01 |
| **Non-European** | | | | |
| ARIC.AA | 675 | 0.9824 | -0.1238 | 5.498e-01 |
| FBIRN | 275 | 0.9164 | 0.1049 | 4.967e-01 |
| NOMAS.black | 168 | 0.9654 | 0.2531 | 3.975e-01 |
| NOMAS.hispanic | 718 | 0.9437 | -0.0324 | 7.769e-01 |
| WHICAP.black | 60 | 0.9586 | -0.0426 | 9.260e-01 |
| **Summary European MA** | **73549** | **0.8917** | **0.05** | **1.893e-09** |
| **Summary African MA** | **903** | **0.9777** | **-0.028** | **0.8602** |
| **Summary Transancestral MA** | **75445** | **0.8933** | **0.049** | **3.124e-09** |

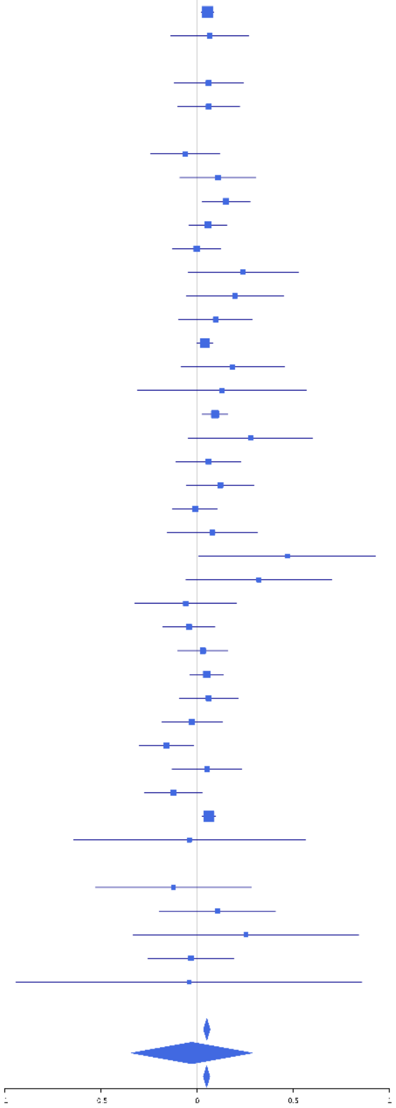

### rs343093 (C); chr12:66255005; model 1 (I2=31.8, HetP=0.01916)

| Study | N_total | EAF | Beta | P |
|---|---|---|---|---|
| **Head circumference** | | | | |
| Haworth | 18881 | 0.8477 | 0.0336 | 1.870e-02 |
| HKOS | 888 | 0.6289 | 0.0399 | 4.164e-01 |
| VIDI | 755 | 0.8636 | -0.0279 | 7.104e-01 |
| **CT-based** | | | | |
| ESS | 641 | 0.8069 | -0.0187 | 7.908e-01 |
| UCL | 642 | 0.8668 | -0.0209 | 7.994e-01 |
| **MR-based** | | | | |
| 1000Brains | 751 | 0.881 | 0.0505 | 5.261e-01 |
| 3C-Bordeaux | 564 | 0.8335 | 0.1607 | 4.487e-02 |
| 3c-Dijon | 1499 | 0.8335 | 0.0935 | 5.656e-02 |
| AGES | 2226 | 0.8694 | 0.0654 | 1.412e-01 |
| ARIC.EA | 1390 | 0.8527 | 0.0194 | 7.177e-01 |
| ASPS-Family | 338 | 0.8506 | -0.2984 | 5.672e-03 |
| ASRB | 233 | 0.7895 | -0.0974 | 3.923e-01 |
| CARDIA.EA | 349 | 0.8522 | -0.0373 | 7.266e-01 |
| CHAP | 261 | 0.8813 | -0.0256 | 8.500e-01 |
| CHS | 544 | 0.8331 | 0.1131 | 1.641e-01 |
| ENIGMA | 11373 | 0.8548 | 0.0479 | 1.098e-02 |
| EPOZ | 251 | 0.8656 | -0.2166 | 9.802e-02 |
| ERF | 122 | 0.7663 | -0.3272 | 3.057e-02 |
| FHS | 4710 | 0.837 | 0.07 | 1.202e-02 |
| GenR | 265 | 0.7365 | 0.0491 | 6.181e-01 |
| HUNT | 851 | 0.8505 | 0.087 | 2.004e-01 |
| LBC1936 | 577 | 0.8564 | -0.056 | 5.047e-01 |
| LIFE | 1825 | 0.8727 | 0.0761 | 1.257e-01 |
| LLS | 355 | 0.8796 | 0.2407 | 3.689e-02 |
| MethCT. | 114 | 0.4026 | -0.0362 | 7.891e-01 |
| NOMAS | 141 | 0.7713 | -0.0509 | 7.204e-01 |
| PoznanMS | 204 | 0.8774 | 0.1262 | 4.031e-01 |
| PROSPER | 315 | 0.8604 | 0.2613 | 2.303e-02 |
| RS1 | 1103 | 0.8546 | 0.08 | 1.855e-01 |
| RS2 | 1159 | 0.8558 | 0.091 | 1.239e-01 |
| RS3 | 2582 | 0.8625 | 0.0766 | 5.792e-02 |
| RUSH1 | 184 | 0.8683 | 0.0669 | 6.647e-01 |
| RUSH2 | 106 | 0.8961 | -0.4398 | 5.349e-02 |
| SHIP | 966 | 0.8843 | 0.0338 | 6.345e-01 |
| SHIP-TREND | 858 | 0.8805 | 0.2507 | 7.919e-04 |
| SYS.adolescents | 980 | 0.7848 | -0.0142 | 7.968e-01 |
| SYS.adults | 594 | 0.795 | 0.208 | 3.810e-03 |
| TREND | 1084 | 0.8802 | -0.0655 | 3.219e-01 |
| UKB | 14571 | 0.86 | 0.051 | 2.518e-03 |
| WHICAP | 83 | 0.7716 | 0.0778 | 6.749e-01 |
| **Non-European** | | | | |
| ARIC.AA | 675 | 0.2087 | 0.0065 | 9.226e-01 |
| CARDIA.AA | 132 | 0.181 | -0.1541 | 3.351e-01 |
| CHAP.black | 321 | 0.2016 | 0.163 | 9.855e-02 |
| FBIRN | 275 | 0.7218 | 0.0967 | 3.106e-01 |
| NOMAS.black | 168 | 0.2053 | 0.0493 | 7.156e-01 |
| NOMAS.hispanic | 718 | 0.5775 | -0.0452 | 3.970e-01 |
| SCES | 209 | 0.658 | 0.1588 | 1.235e-01 |
| SIMES | 201 | 0.5574 | 0.0456 | 6.492e-01 |
| WHICAP.black | 60 | 0.2668 | 0.1152 | 5.794e-01 |
| **Summary European MA** | **74333** | **0.8523** | **0.047** | **1.504e-10** |
| **Summary African MA** | **1356** | **0.2065** | **0.039** | **0.4143** |
| **Summary Asian MA** | **1298** | **0.6225** | **0.059** | **0.1418** |
| **Summary Transancestral MA** | **78094** | **0.8336** | **0.044** | **7.289e-11** |

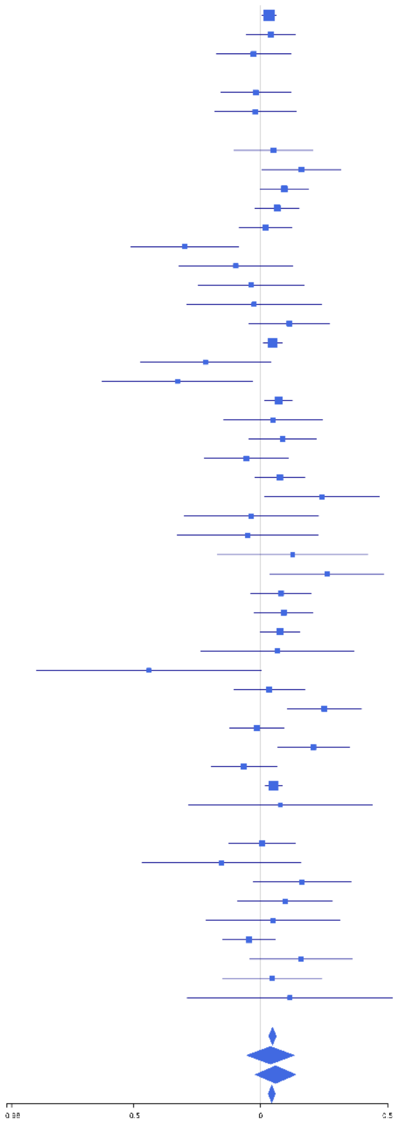

**rs112717745 (T); chr12:66292771; model 1 (I2=0, HetP=0.468)**

| Study | N_total | EAF | Beta | P |
|---|---|---|---|---|
| **Head circumference** | | | | |
| Haworth | 18881 | 0.0873 | -0.0679 | 1.903e-04 |
| VIDI | 724 | 0.0192 | -0.2287 | 2.324e-01 |
| **CT-based** | | | | |
| ESS | 641 | 0.07 | -0.1243 | 2.558e-01 |
| UCL | 642 | 0.0729 | 0.0043 | 9.682e-01 |
| **MR-based** | | | | |
| 1000Brains | 751 | 0.0658 | -0.2635 | 1.153e-02 |
| 3C-Bordeaux | 564 | 0.072 | 0.089 | 4.397e-01 |
| ARIC.EA | 1390 | 0.0704 | 0.0179 | 8.090e-01 |
| ASPS-Family | 338 | 0.0695 | -0.164 | 2.786e-01 |
| EPOZ | 251 | 0.0768 | -0.3025 | 7.107e-02 |
| GenR | 265 | 0.066 | -0.1219 | 4.858e-01 |
| HUNT | 851 | 0.0374 | -0.1354 | 2.890e-01 |
| LIFE | 1825 | 0.0621 | -0.1108 | 1.065e-01 |
| PoznanMS | 204 | 0.0711 | -0.1108 | 5.654e-01 |
| RS1 | 1103 | 0.0719 | 0.009 | 9.134e-01 |
| RS2 | 1159 | 0.0695 | -0.0804 | 3.250e-01 |
| RS3 | 2582 | 0.0748 | -0.0788 | 1.364e-01 |
| SHIP | 966 | 0.0607 | -0.087 | 3.617e-01 |
| SHIP-TREND | 858 | 0.0573 | -0.2402 | 2.095e-02 |
| SYS.adolescents | 980 | 0.0737 | -0.0767 | 3.753e-01 |
| SYS.adults | 594 | 0.0667 | 0.0971 | 4.040e-01 |
| TREND | 1084 | 0.069 | -0.2507 | 3.162e-03 |
| UKB | 14571 | 0.0774 | -0.069 | 1.648e-03 |
| WHICAP | 83 | 0.0394 | 0.1584 | 6.921e-01 |
| **Non-European** | | | | |
| FBIRN | 275 | 0.0563 | 0.2016 | 2.764e-01 |
| **Summary European MA** | **51307** | **0.0768** | **-0.076** | **9.976e-11** |
| **Summary Transancestral MA** | **51582** | **0.0767** | **-0.075** | **1.883e-10** |

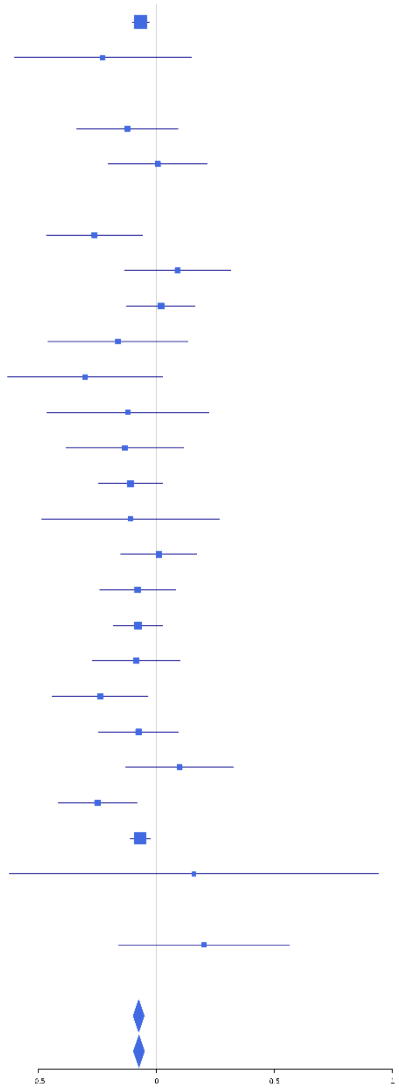

**rs7306710 (T); chr12:66376091; model 1 (I2=22.6, HetP=0.08194)**

| Study | N_total | EAF | Beta | P |
|---|---|---|---|---|
| **Head circumference** | | | | |
| Haworth | 18881 | 0.4707 | 0.0501 | 1.250e-06 |
| HKOS | 888 | 0.0968 | 0.0337 | 6.745e-01 |
| VIDI | 738 | 0.484 | 0.0703 | 1.779e-01 |
| **CT-based** | | | | |
| ESS | 641 | 0.4528 | 0.1515 | 6.938e-03 |
| UCL | 642 | 0.4545 | 0.0331 | 5.550e-01 |
| **MR-based** | | | | |
| 1000Brains | 751 | 0.504 | 0.1135 | 2.813e-02 |
| 3C-Bordeaux | 564 | 0.429 | -0.0803 | 1.825e-01 |
| 3c-Dijon | 1499 | 0.4294 | 0.022 | 5.506e-01 |
| AGES | 2226 | 0.5134 | 0.0891 | 2.979e-03 |
| ARIC.EA | 1390 | 0.4859 | 0.0508 | 1.810e-01 |
| ASPS-Family | 338 | 0.4416 | 0.0922 | 2.342e-01 |
| ASRB | 233 | 0.4581 | 0.0695 | 4.550e-01 |
| CARDIA.EA | 349 | 0.4902 | 0.0223 | 7.686e-01 |
| CHAP | 261 | 0.5046 | 0.139 | 1.135e-01 |
| CHS | 544 | 0.4372 | 0.0522 | 3.934e-01 |
| DNGS | 502 | 0.3638 | -0.0398 | 5.440e-01 |
| ENIGMA | 11373 | 0.485 | 0.0368 | 5.577e-03 |
| EPOZ | 251 | 0.4936 | -0.0763 | 3.927e-01 |
| ERF | 122 | 0.4343 | 0.0948 | 4.629e-01 |
| FHS | 4710 | 0.459 | 0.0636 | 2.085e-03 |
| GenR | 265 | 0.4444 | -0.0183 | 8.341e-01 |
| HUNT | 851 | 0.5143 | 0.1106 | 2.257e-02 |
| LBC1936 | 577 | 0.478 | 0.028 | 6.349e-01 |
| LIFE | 1825 | 0.5097 | 0.0498 | 1.329e-01 |
| LLS | 355 | 0.4918 | -0.0255 | 7.338e-01 |
| MethCT. | 114 | 0.3599 | 0.2773 | 4.443e-02 |
| NOMAS | 141 | 0.3941 | 0.1037 | 3.964e-01 |
| PoznanMS | 204 | 0.547 | 0.0973 | 3.280e-01 |
| PROSPER | 315 | 0.5355 | 0.0036 | 9.639e-01 |
| RS1 | 1103 | 0.4982 | 0.069 | 1.051e-01 |
| RS2 | 1159 | 0.5014 | 0.0638 | 1.245e-01 |
| RS3 | 2582 | 0.4972 | 0.0793 | 4.399e-03 |
| RUSH1 | 184 | 0.4923 | -0.0254 | 8.071e-01 |
| RUSH2 | 106 | 0.4753 | 0.0789 | 5.675e-01 |
| SHIP | 966 | 0.504 | 0.0118 | 7.948e-01 |
| SHIP-TREND | 858 | 0.5191 | 0.0887 | 6.669e-02 |
| SYS.adolescents | 980 | 0.4071 | 0.0903 | 4.952e-02 |
| SYS.adults | 594 | 0.4163 | 0.1789 | 2.364e-03 |
| TREND | 1084 | 0.4856 | 0.1522 | 4.124e-04 |
| UKB | 14571 | 0.4802 | 0.084 | 7.598e-13 |
| WHICAP | 83 | 0.4051 | 0.3034 | 5.867e-02 |
| **Non-European** | | | | |
| ARIC.AA | 675 | 0.4115 | -0.0624 | 2.594e-01 |
| CARDIA.AA | 132 | 0.425 | 0.1148 | 3.563e-01 |
| CHAP.black | 321 | 0.4495 | -0.0973 | 2.211e-01 |
| FBIRN | 275 | 0.3567 | -0.0303 | 7.343e-01 |
| NOMAS.black | 168 | 0.454 | 0.0524 | 6.332e-01 |
| NOMAS.hispanic | 718 | 0.4026 | 0.0466 | 3.868e-01 |
| SCES | 209 | 0.0845 | -0.1685 | 3.382e-01 |
| SIMES | 201 | 0.1288 | -0.191 | 2.016e-01 |
| WHICAP.black | 60 | 0.4034 | 0.0715 | 7.024e-01 |
| **Summary European MA** | **74818** | **0.4772** | **0.061** | **9.792e-32** |
| **Summary African MA** | **1356** | **0.4267** | **-0.033** | **0.3926** |
| **Summary Asian MA** | **1298** | **0.0998** | **-0.035** | **0.5896** |
| **Summary Transancestral MA** | **78579** | **0.4688** | **0.058** | **4.38e-30** |

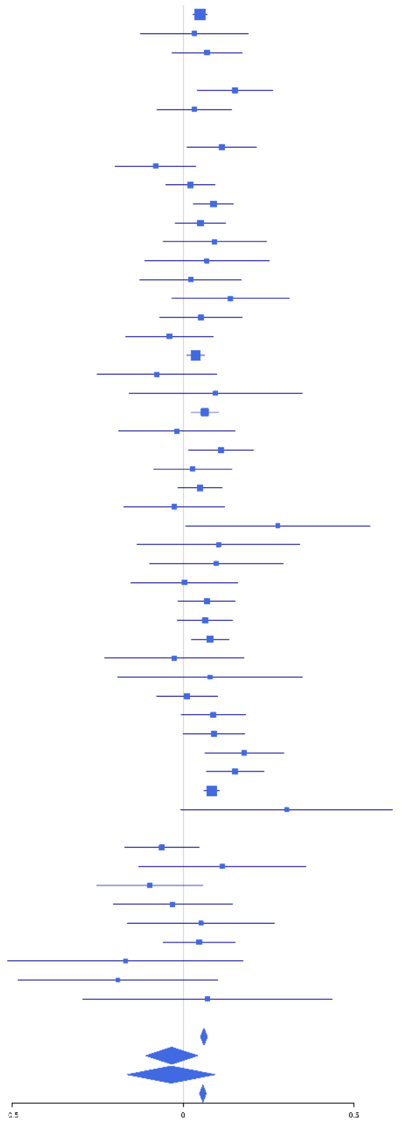

**rs35227403 (A); chr12:68216239; model 1 (I2=24.7, HetP=0.07982)**

| Study | N_total | EAF | Beta | P |
|---|---|---|---|---|
| **Head circumference** | | | | |
| Haworth | 18881 | 0.0597 | -0.0746 | 5.971e-04 |
| HKOS | 888 | 0.0298 | -0.11 | 4.303e-01 |
| VIDI | 723 | 0.0815 | -0.2585 | 7.333e-03 |
| **CT-based** | | | | |
| ESS | 641 | 0.0564 | -0.2049 | 9.067e-02 |
| UCL | 642 | 0.0455 | -0.0021 | 9.870e-01 |
| **MR-based** | | | | |
| 1000Brains | 751 | 0.0486 | -0.0355 | 7.671e-01 |
| 3C-Bordeaux | 564 | 0.0519 | -0.177 | 1.882e-01 |
| 3c-Dijon | 1499 | 0.0518 | -0.1046 | 2.044e-01 |
| AGES | 2226 | 0.0557 | -0.1386 | 3.391e-02 |
| ARIC.EA | 1390 | 0.0588 | -0.2347 | 3.598e-03 |
| ASPS-Family | 338 | 0.0571 | 0.0027 | 9.873e-01 |
| ASRB | 233 | 0.047 | -0.1445 | 5.101e-01 |
| CARDIA.EA | 349 | 0.0533 | -0.0681 | 6.861e-01 |
| CHAP | 261 | 0.0512 | -0.1197 | 5.472e-01 |
| CHS | 544 | 0.0444 | 0.0289 | 8.442e-01 |
| ENIGMA | 11373 | 0.0611 | -0.021 | 4.487e-01 |
| EPOZ | 251 | 0.0502 | 0.1785 | 3.829e-01 |
| ERF | 122 | 0.0778 | 0.3941 | 9.918e-02 |
| FHS | 4710 | 0.0561 | -0.0147 | 7.423e-01 |
| GenR | 265 | 0.0492 | -0.0247 | 9.024e-01 |
| HUNT | 851 | 0.0683 | -0.0135 | 8.881e-01 |
| LBC1936 | 577 | 0.0535 | 0.1429 | 2.750e-01 |
| LIFE | 1825 | 0.0609 | -0.0345 | 6.186e-01 |
| LLS | 355 | 0.0592 | -0.0639 | 6.876e-01 |
| NOMAS | 141 | 0.0458 | 0.039 | 8.915e-01 |
| PoznanMS | 204 | 0.0614 | -0.388 | 5.993e-02 |
| PROSPER | 315 | 0.0654 | -0.3437 | 3.301e-02 |
| RS1 | 1103 | 0.0577 | -0.0128 | 8.885e-01 |
| RS2 | 1159 | 0.0535 | -0.3405 | 2.248e-04 |
| RS3 | 2582 | 0.0537 | -0.1436 | 2.000e-02 |
| RUSH1 | 184 | 0.0601 | -0.0224 | 9.185e-01 |
| SHIP | 966 | 0.0647 | -0.0636 | 4.916e-01 |
| SHIP-TREND | 858 | 0.0714 | -0.1717 | 6.746e-02 |
| SYS.adolescents | 980 | 0.0375 | -0.2655 | 2.553e-02 |
| SYS.adults | 594 | 0.0446 | -0.0649 | 6.444e-01 |
| TREND | 1084 | 0.066 | -0.1107 | 2.009e-01 |
| UKB | 14571 | 0.0601 | -0.0959 | 9.973e-05 |
| WHICAP | 83 | 0.0443 | -0.1166 | 7.581e-01 |
| **Non-European** | | | | |
| FBIRN | 275 | 0.0512 | 0.1825 | 3.464e-01 |
| NOMAS.hispanic | 718 | 0.0415 | 0.0328 | 8.039e-01 |
| WHICAP.black | 60 | 0.028 | 1.0984 | 5.232e-02 |
| **Summary European MA** | **74195** | **0.0587** | **-0.08** | **4.702e-13** |
| **Summary African MA** | **60** | **0.028** | **1.073** | **0.05232** |
| **Summary Asian MA** | **888** | **0.0298** | **-0.11** | **0.4303** |
| **Summary Transancestral MA** | **76136** | **0.0581** | **-0.078** | **1.333e-12** |

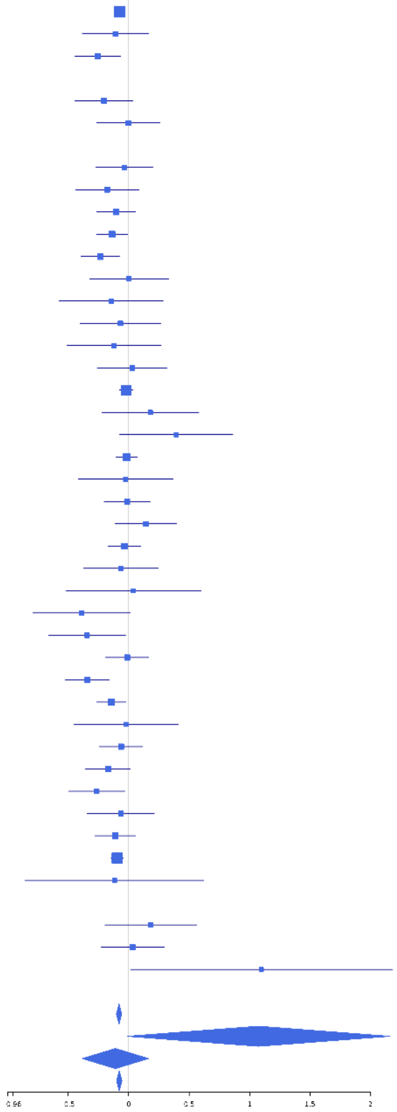

**rs7310309 (T); chr12:79811429; model 1 (I2=2.6, HetP=0.4212)**

| Study | N_total | EAF | Beta | P |
|---|---|---|---|---|
| **Head circumference** | | | | |
| Haworth | 18881 | 0.1531 | -0.0314 | 2.819e-02 |
| HKOS | 888 | 0.0992 | -0.0687 | 3.869e-01 |
| VIDI | 737 | 0.0975 | -0.1207 | 1.693e-01 |
| **CT-based** | | | | |
| ESS | 641 | 0.1892 | -0.0588 | 4.098e-01 |
| UCL | 642 | 0.1489 | -0.0314 | 6.882e-01 |
| **MR-based** | | | | |
| 1000Brains | 751 | 0.1568 | 0.0178 | 8.020e-01 |
| 3C-Bordeaux | 564 | 0.1906 | -0.0675 | 3.734e-01 |
| 3c-Dijon | 1499 | 0.192 | -0.1194 | 9.982e-03 |
| AGES | 2226 | 0.1227 | -0.0232 | 6.116e-01 |
| ARIC.EA | 1390 | 0.1574 | -0.1163 | 2.545e-02 |
| ASPS-Family | 338 | 0.1664 | -0.1481 | 1.516e-01 |
| ASRB | 233 | 0.1929 | 0.0188 | 8.727e-01 |
| CARDIA.EA | 349 | 0.1618 | -0.043 | 6.758e-01 |
| CHAP | 261 | 0.1613 | 0.0444 | 7.096e-01 |
| CHS | 544 | 0.1835 | -0.0135 | 8.638e-01 |
| DNGS | 504 | 0.1896 | -0.0128 | 8.733e-01 |
| ENIGMA | 11373 | 0.1566 | -0.0507 | 5.447e-03 |
| EPOZ | 251 | 0.1744 | 0.0336 | 7.746e-01 |
| ERF | 122 | 0.1999 | 0.4561 | 4.382e-03 |
| FHS | 4710 | 0.1709 | -0.0641 | 1.916e-02 |
| GenR | 265 | 0.2326 | -0.1453 | 1.577e-01 |
| HUNT | 851 | 0.1397 | 0.0259 | 7.115e-01 |
| LBC1936 | 577 | 0.1367 | -0.0272 | 7.506e-01 |
| LIFE | 1825 | 0.1562 | 0.0294 | 5.196e-01 |
| LLS | 355 | 0.1667 | -0.158 | 1.165e-01 |
| MethCT. | 114 | 0.3781 | -5e-04 | 9.965e-01 |
| NOMAS | 141 | 0.2071 | -0.0923 | 5.313e-01 |
| PoznanMS | 204 | 0.1454 | -0.0421 | 7.639e-01 |
| PROSPER | 315 | 0.1536 | -0.1153 | 2.968e-01 |
| RS1 | 1103 | 0.1586 | -0.1519 | 9.164e-03 |
| RS2 | 1159 | 0.1588 | 0.0246 | 6.648e-01 |
| RS3 | 2582 | 0.164 | 0.0222 | 5.553e-01 |
| RUSH1 | 184 | 0.136 | -0.0862 | 5.713e-01 |
| RUSH2 | 106 | 0.1484 | -0.1121 | 5.634e-01 |
| SHIP | 966 | 0.1413 | -0.0444 | 4.976e-01 |
| SHIP-TREND | 858 | 0.1492 | -0.1377 | 4.244e-02 |
| SYS.adolescents | 980 | 0.156 | -0.0486 | 4.355e-01 |
| SYS.adults | 594 | 0.1737 | -0.0351 | 6.472e-01 |
| TREND | 1084 | 0.1384 | -0.0085 | 8.921e-01 |
| UKB | 14571 | 0.1418 | -0.0588 | 4.667e-04 |
| WHICAP | 83 | 0.214 | 0.0666 | 7.257e-01 |
| **Non-European** | | | | |
| ARIC.AA | 675 | 0.6655 | -0.0139 | 8.097e-01 |
| CARDIA.AA | 132 | 0.665 | -0.0323 | 8.045e-01 |
| CHAP.black | 321 | 0.679 | 0.0177 | 8.347e-01 |
| FBIRN | 275 | 0.2082 | 0.1314 | 2.123e-01 |
| NOMAS.black | 168 | 0.6074 | 0.1717 | 1.261e-01 |
| NOMAS.hispanic | 718 | 0.3575 | -0.0314 | 5.684e-01 |
| SCES | 209 | 0.105 | 0.2888 | 7.035e-02 |
| SIMES | 201 | 0.0756 | -0.2045 | 2.753e-01 |
| WHICAP.black | 60 | 0.5897 | 0.0898 | 6.304e-01 |
| **Summary European MA** | **74819** | **0.154** | **-0.044** | **6.04e-10** |
| **Summary African MA** | **1356** | **0.6581** | **0.02** | **0.6213** |
| **Summary Asian MA** | **1298** | **0.0965** | **-0.028** | **0.6754** |
| **Summary Transancestral MA** | **78580** | **0.1641** | **-0.041** | **1.864e-09** |

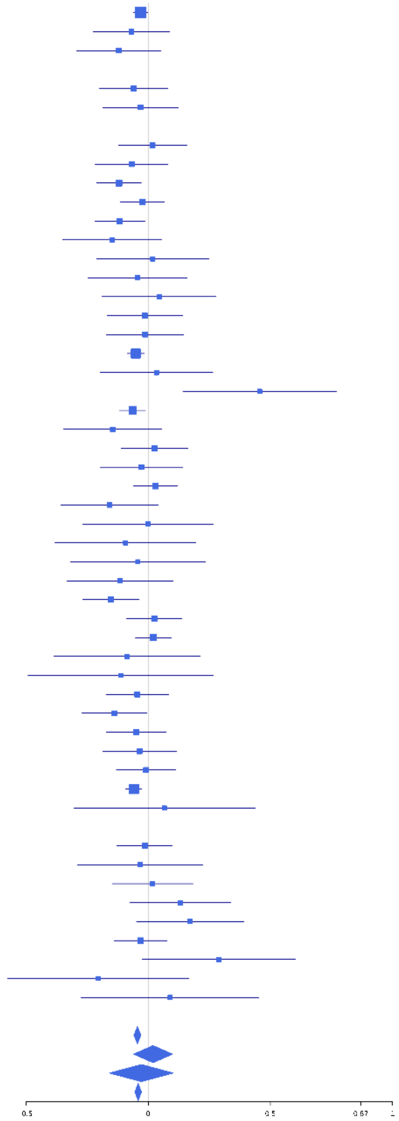

**rs11111293 (T); chr12:102921296; model 1 (I2=4.8, HetP=0.38)**

| Study | N_total | EAF | Beta | P |
|---|---|---|---|---|
| **Head circumference** | | | | |
| Haworth | 18881 | 0.7783 | 0.0169 | 1.708e-01 |
| HKOS | 888 | 0.8789 | 0.0858 | 2.380e-01 |
| VIDI | 656 | 0.8113 | -0.045 | 5.236e-01 |
| **CT-based** | | | | |
| ESS | 641 | 0.7754 | -0.1548 | 2.074e-02 |
| UCL | 642 | 0.8017 | 0.0822 | 2.402e-01 |
| **MR-based** | | | | |
| 1000Brains | 751 | 0.77 | 0.03 | 6.246e-01 |
| 3C-Bordeaux | 564 | 0.8042 | 0.1101 | 1.427e-01 |
| 3c-Dijon | 1499 | 0.7983 | 0.0398 | 3.815e-01 |
| AGES | 2226 | 0.7664 | 0.0562 | 1.128e-01 |
| ARIC.EA | 1390 | 0.7817 | -0.0239 | 6.021e-01 |
| ASPS-Family | 338 | 0.8026 | 0.1005 | 2.985e-01 |
| ASRB | 233 | 0.7843 | 0.0131 | 9.076e-01 |
| CARDIA.EA | 349 | 0.7798 | 0.0353 | 6.989e-01 |
| CHAP | 261 | 0.8103 | 0.1457 | 1.931e-01 |
| CHS | 544 | 0.773 | -0.023 | 7.503e-01 |
| DNGS | 516 | 0.85 | 0.0645 | 4.597e-01 |
| ENIGMA | 11373 | 0.7817 | 0.05 | 1.827e-03 |
| EPOZ | 251 | 0.7605 | -0.0858 | 4.123e-01 |
| FHS | 4710 | 0.7761 | 0.0814 | 9.844e-04 |
| GenR | 265 | 0.819 | 0.118 | 2.957e-01 |
| HUNT | 851 | 0.7558 | -0.0201 | 7.208e-01 |
| LBC1936 | 577 | 0.7771 | 8e-04 | 9.908e-01 |
| LIFE | 1825 | 0.7767 | 0.1289 | 1.199e-03 |
| LLS | 355 | 0.8014 | 0.0017 | 9.858e-01 |
| NOMAS | 141 | 0.8024 | 0.0229 | 8.787e-01 |
| PoznanMS | 204 | 0.745 | -0.037 | 7.442e-01 |
| PROSPER | 315 | 0.7778 | -0.0454 | 6.358e-01 |
| RS1 | 1103 | 0.7819 | 0.0758 | 1.415e-01 |
| RS2 | 1159 | 0.78 | 0.0273 | 5.861e-01 |
| RS3 | 2582 | 0.7821 | 0.0887 | 8.531e-03 |
| RUSH1 | 184 | 0.7518 | 0.2115 | 8.138e-02 |
| RUSH2 | 106 | 0.7783 | 0.2655 | 1.114e-01 |
| SHIP | 966 | 0.7758 | 0.1234 | 2.391e-02 |
| SHIP-TREND | 858 | 0.7681 | 0.0483 | 3.988e-01 |
| SYS.adolescents | 980 | 0.7893 | 0.0744 | 1.792e-01 |
| SYS.adults | 594 | 0.782 | 0.1071 | 1.276e-01 |
| TREND | 1084 | 0.7849 | -0.0084 | 8.728e-01 |
| UKB | 14571 | 0.7802 | 0.0481 | 6.762e-04 |
| WHICAP | 83 | 0.8025 | -0.0871 | 6.561e-01 |
| **Non-European** | | | | |
| ARIC.AA | 675 | 0.9615 | 0.2947 | 3.723e-02 |
| CHAP.black | 321 | 0.9455 | 0.0223 | 8.980e-01 |
| IMH | 37 | 0.8772 | -0.2416 | 4.955e-01 |
| NOMAS.black | 168 | 0.9359 | -0.1434 | 5.204e-01 |
| NOMAS.hispanic | 718 | 0.8981 | 0.0845 | 3.328e-01 |
| SCES | 209 | 0.9163 | -0.0113 | 9.491e-01 |
| SIMES | 201 | 0.8881 | 0.1557 | 3.259e-01 |
| WHICAP.black | 60 | 0.9417 | 0.2766 | 4.806e-01 |
| **Summary European MA** | **74628** | **0.7806** | **0.042** | **1.85e-11** |
| **Summary African MA** | **1224** | **0.9528** | **0.146** | **0.1258** |
| **Summary Asian MA** | **1335** | **0.8861** | **0.073** | **0.2283** |
| **Summary Transancestral MA** | **77905** | **0.7862** | **0.043** | **2.265e-12** |

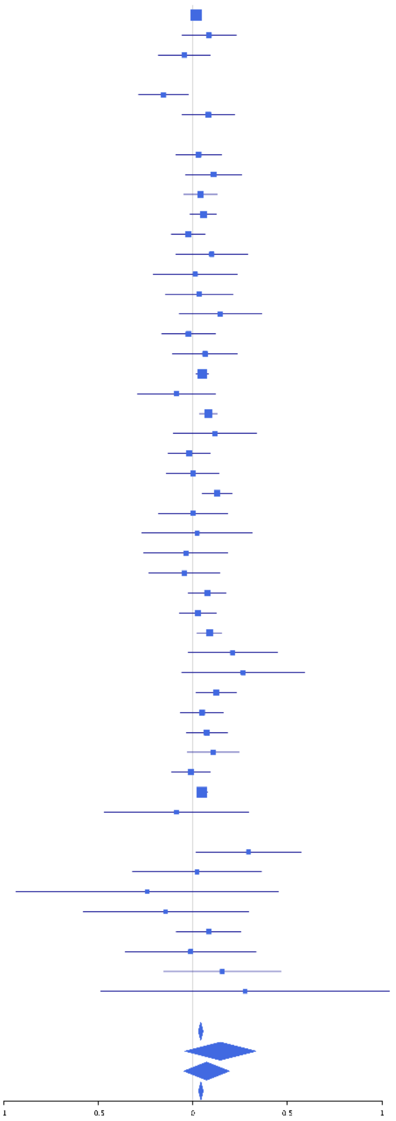

**rs28636834 (A); chr12:123871070; model 1 (I2=3.8, HetP=0.4004)**

| Study | N_total | EAF | Beta | P |
|---|---|---|---|---|
| **Head circumference** | | | | |
| Haworth | 18881 | 0.7809 | -0.07 | 1.881e-08 |
| VIDI | 755 | 0.7906 | -0.0552 | 3.830e-01 |
| **CT-based** | | | | |
| ESS | 641 | 0.7904 | -0.2292 | 8.383e-04 |
| UCL | 642 | 0.7978 | -0.1659 | 1.699e-02 |
| **MR-based** | | | | |
| 1000Brains | 751 | 0.778 | -0.0582 | 3.489e-01 |
| 3C-Bordeaux | 564 | 0.8131 | -0.1231 | 1.074e-01 |
| 3c-Dijon | 1499 | 0.7923 | -0.0946 | 3.557e-02 |
| AGES | 2226 | 0.7508 | -0.0203 | 5.586e-01 |
| ARIC.EA | 1390 | 0.7953 | -0.0643 | 1.714e-01 |
| ASPS-Family | 338 | 0.7713 | -0.1487 | 1.043e-01 |
| ASRB | 233 | 0.791 | -0.0523 | 6.468e-01 |
| CARDIA.EA | 349 | 0.7869 | 0.014 | 8.794e-01 |
| CHAP | 261 | 0.7946 | -0.0754 | 4.873e-01 |
| DNGS | 512 | 0.7953 | -0.0631 | 4.153e-01 |
| ENIGMA | 6782 | 0.78 | -0.0216 | 2.965e-01 |
| EPOZ | 251 | 0.7654 | 0.0252 | 8.112e-01 |
| ERF | 122 | 0.7617 | -0.2601 | 8.354e-02 |
| GenR | 265 | 0.7515 | -0.0915 | 3.628e-01 |
| HUNT | 851 | 0.8012 | -0.0517 | 3.942e-01 |
| LBC1936 | 577 | 0.7895 | -0.0335 | 6.428e-01 |
| LIFE | 1825 | 0.7994 | -0.0582 | 1.596e-01 |
| LLS | 355 | 0.8028 | -0.0242 | 7.974e-01 |
| MethCT. | 114 | 0.6273 | -0.3358 | 1.419e-02 |
| NOMAS | 141 | 0.7835 | -0.0103 | 9.436e-01 |
| PoznanMS | 204 | 0.7543 | -0.2072 | 7.156e-02 |
| PROSPER | 315 | 0.7407 | 0.0085 | 9.254e-01 |
| RS1 | 1103 | 0.7878 | 0.0177 | 7.350e-01 |
| RS2 | 1159 | 0.7922 | 0.0379 | 4.586e-01 |
| RS3 | 2582 | 0.7858 | -0.0021 | 9.510e-01 |
| RUSH1 | 184 | 0.7462 | -0.1412 | 2.399e-01 |
| RUSH2 | 106 | 0.7783 | -0.2708 | 1.046e-01 |
| SHIP | 966 | 0.7947 | 0.0075 | 8.936e-01 |
| SHIP-TREND | 858 | 0.7803 | -0.0911 | 1.184e-01 |
| SYS.adolescents | 980 | 0.7948 | -0.0065 | 9.070e-01 |
| SYS.adults | 594 | 0.8201 | 0.0616 | 4.151e-01 |
| TREND | 1084 | 0.7739 | -0.0097 | 8.506e-01 |
| UKB | 14571 | 0.795 | -0.0499 | 5.820e-04 |
| WHICAP | 83 | 0.759 | -0.0602 | 7.410e-01 |
| **Non-European** | | | | |
| ARIC.AA | 675 | 0.4966 | -0.0132 | 8.081e-01 |
| CARDIA.AA | 132 | 0.55 | -0.1188 | 3.373e-01 |
| CHAP.black | 321 | 0.5372 | -0.0932 | 2.395e-01 |
| FBIRN | 275 | 0.7594 | -0.1191 | 2.337e-01 |
| NOMAS.black | 168 | 0.544 | 0.0504 | 6.461e-01 |
| NOMAS.hispanic | 718 | 0.7272 | -0.0527 | 3.737e-01 |
| WHICAP.black | 60 | 0.525 | 0.164 | 3.739e-01 |
| **Summary European MA** | **65000** | **0.7858** | **-0.051** | **5.011e-14** |
| **Summary African MA** | **1356** | **0.5185** | **-0.027** | **0.4874** |
| **Summary Transancestral MA** | **67463** | **0.7795** | **-0.051** | **8.498e-15** |

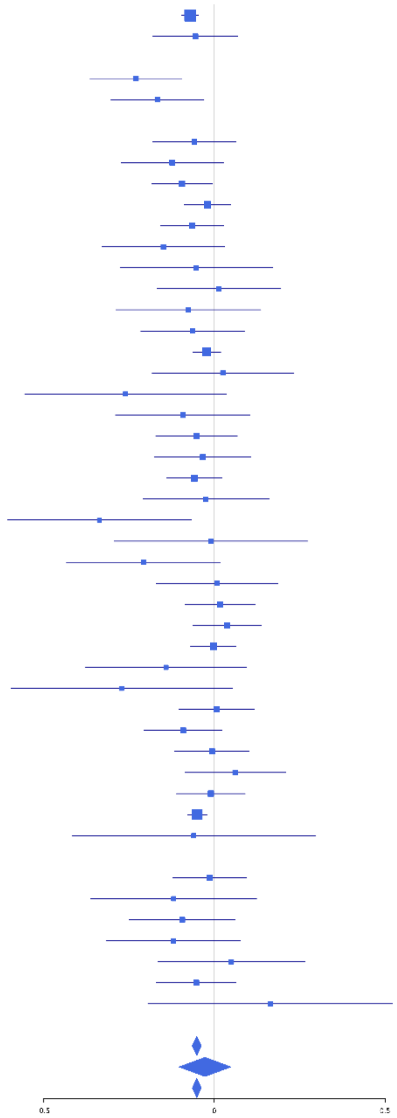

**rs7988627 (A); chr13:81631782; model 1 (I2=9.8, HetP=0.2771)**

| Study | N_total | EAF | Beta | P |
|---|---|---|---|---|
| **Head circumference** | | | | |
| Haworth | 18881 | 0.5783 | -0.031 | 3.073e-03 |
| HKOS | 888 | 0.3874 | -2e-04 | 9.966e-01 |
| VIDI | 708 | 0.633 | -0.1613 | 3.552e-03 |
| **CT-based** | | | | |
| ESS | 641 | 0.5662 | -0.0888 | 1.152e-01 |
| UCL | 642 | 0.5745 | -0.0684 | 2.259e-01 |
| **MR-based** | | | | |
| 1000Brains | 751 | 0.6064 | -0.0399 | 4.501e-01 |
| 3C-Bordeaux | 564 | 0.579 | -0.0368 | 5.415e-01 |
| 3c-Dijon | 1499 | 0.5721 | -0.0251 | 4.971e-01 |
| AGES | 2226 | 0.5183 | -0.108 | 3.160e-04 |
| ARIC.EA | 1390 | 0.562 | -0.0931 | 1.486e-02 |
| ASPS-Family | 338 | 0.5746 | -1e-04 | 9.992e-01 |
| ASRB | 233 | 0.5528 | -0.0539 | 5.629e-01 |
| CARDIA.EA | 349 | 0.5612 | -0.0537 | 4.813e-01 |
| CHAP | 261 | 0.5642 | 0.0621 | 4.828e-01 |
| CHS | 544 | 0.5838 | -0.0934 | 1.288e-01 |
| DNGS | 494 | 0.5807 | 0.0855 | 1.853e-01 |
| ENIGMA | 11373 | 0.5735 | 0.005 | 7.103e-01 |
| EPOZ | 251 | 0.5428 | -0.1793 | 4.535e-02 |
| ERF | 122 | 0.5476 | -0.0394 | 7.597e-01 |
| FHS | 4710 | 0.5792 | -0.0132 | 5.279e-01 |
| GenR | 265 | 0.6183 | 0.131 | 1.430e-01 |
| HUNT | 851 | 0.558 | -0.0114 | 8.153e-01 |
| LBC1936 | 577 | 0.573 | -0.0717 | 2.282e-01 |
| LIFE | 1825 | 0.5885 | -0.038 | 2.588e-01 |
| LLS | 355 | 0.5954 | 0.0651 | 3.949e-01 |
| MethCT. | 114 | 0.728 | 0.1277 | 3.909e-01 |
| NOMAS | 141 | 0.5965 | -0.0296 | 8.079e-01 |
| PoznanMS | 204 | 0.6368 | -0.1073 | 2.973e-01 |
| PROSPER | 315 | 0.5657 | 0.0845 | 2.931e-01 |
| RS1 | 1103 | 0.5564 | -0.0496 | 2.469e-01 |
| RS2 | 1159 | 0.5716 | -0.0408 | 3.314e-01 |
| RS3 | 2582 | 0.5551 | -0.0839 | 2.735e-03 |
| RUSH1 | 184 | 0.5723 | 0.0054 | 9.593e-01 |
| RUSH2 | 106 | 0.5785 | -0.0819 | 5.572e-01 |
| SHIP | 966 | 0.5788 | -0.0566 | 2.192e-01 |
| SHIP-TREND | 858 | 0.5985 | -0.0084 | 8.649e-01 |
| SYS.adolescents | 980 | 0.5629 | -0.0515 | 2.586e-01 |
| SYS.adults | 594 | 0.5601 | 0.0083 | 8.874e-01 |
| TREND | 1084 | 0.5969 | -0.0283 | 5.175e-01 |
| UKB | 14571 | 0.5619 | -0.0328 | 5.444e-03 |
| WHICAP | 83 | 0.59 | 0.0159 | 9.199e-01 |
| **Non-European** | | | | |
| ARIC.AA | 675 | 0.9021 | 0.0386 | 6.734e-01 |
| CARDIA.AA | 132 | 0.895 | 0.1401 | 4.853e-01 |
| CHAP.black | 321 | 0.8675 | 0.0426 | 7.149e-01 |
| FBIRN | 275 | 0.5956 | -0.0481 | 5.802e-01 |
| IMH | 37 | 0.3596 | -0.1567 | 5.178e-01 |
| NOMAS.black | 168 | 0.895 | 0.0897 | 6.153e-01 |
| NOMAS.hispanic | 718 | 0.7154 | -0.0765 | 1.913e-01 |
| SCES | 209 | 0.4469 | 0.0317 | 7.477e-01 |
| SIMES | 201 | 0.399 | 0.0458 | 6.254e-01 |
| WHICAP.black | 60 | 0.8967 | 0.4547 | 1.354e-01 |
| **Summary European MA** | **74780** | **0.5718** | **-0.031** | **2.198e-09** |
| **Summary African MA** | **1356** | **0.8921** | **0.073** | **0.2365** |
| **Summary Asian MA** | **1335** | **0.3977** | **0.008** | **0.8371** |
| **Summary Transancestral MA** | **78578** | **0.576** | **-0.029** | **7.526e-09** |

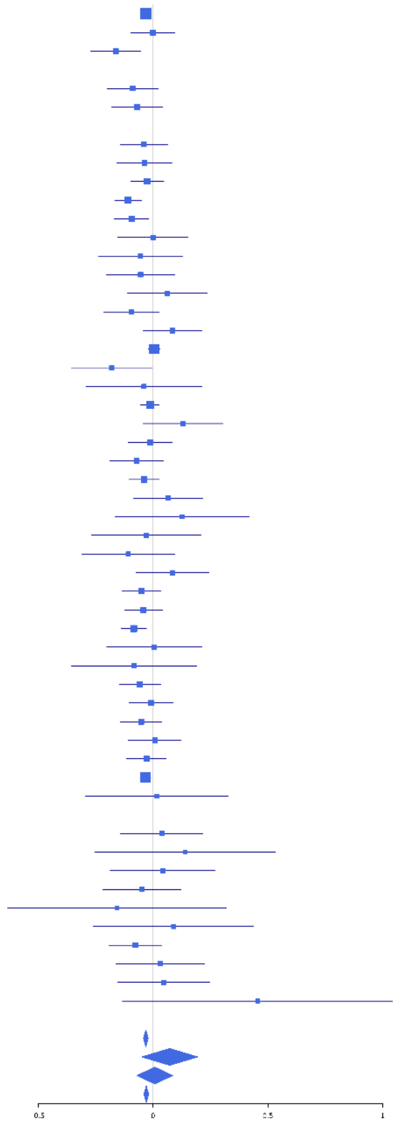

**rs3093872 (T); chr14:20811332; model 1 (I2=4, HetP=0.3941)**

| Study | N_total | EAF | Beta | P |
|---|---|---|---|---|
| **Head circumference** | | | | |
| Haworth | 18881 | 0.0584 | -0.0325 | 1.378e-01 |
| HKOS | 888 | 0.094 | 0.1254 | 1.228e-01 |
| VIDI | 751 | 0.0991 | -0.1743 | 4.390e-02 |
| **CT-based** | | | | |
| UCL | 642 | 0.0478 | -0.0946 | 4.698e-01 |
| **MR-based** | | | | |
| 1000Brains | 751 | 0.0641 | -0.0382 | 7.165e-01 |
| 3C-Bordeaux | 564 | 0.0509 | -0.177 | 1.917e-01 |
| 3c-Dijon | 1499 | 0.0519 | -0.176 | 3.262e-02 |
| AGES | 2226 | 0.0539 | -0.2279 | 5.952e-04 |
| ARIC.EA | 1390 | 0.0561 | 0.0215 | 7.940e-01 |
| ASPS-Family | 338 | 0.0666 | -0.1756 | 2.553e-01 |
| ASRB | 233 | 0.0418 | -0.1143 | 6.215e-01 |
| CARDIA.EA | 349 | 0.0417 | -0.2009 | 2.886e-01 |
| CHAP | 261 | 0.0709 | -0.1615 | 3.445e-01 |
| CHS | 544 | 0.0515 | 0.0062 | 9.640e-01 |
| DNGS | 516 | 0.0644 | 0.0525 | 6.792e-01 |
| ENIGMA | 11373 | 0.0553 | -0.0853 | 3.274e-03 |
| EPOZ | 251 | 0.051 | 0.1118 | 5.817e-01 |
| ERF | 122 | 0.0458 | -0.2242 | 4.645e-01 |
| FHS | 4710 | 0.0416 | -0.0894 | 8.304e-02 |
| GenR | 265 | 0.0508 | 0.3384 | 8.714e-02 |
| HUNT | 851 | 0.0405 | -0.2383 | 5.261e-02 |
| LBC1936 | 577 | 0.0483 | 0.226 | 9.968e-02 |
| LIFE | 1825 | 0.0764 | -0.0573 | 3.579e-01 |
| LLS | 355 | 0.062 | 0.0691 | 6.568e-01 |
| MethCT. | 114 | 0.0724 | 0.0051 | 9.839e-01 |
| NOMAS | 141 | 0.0321 | 0.1162 | 7.315e-01 |
| PoznanMS | 204 | 0.0933 | -0.2014 | 2.368e-01 |
| PROSPER | 315 | 0.0587 | -0.3816 | 2.431e-02 |
| RS1 | 1103 | 0.0507 | -0.068 | 4.832e-01 |
| RS2 | 1159 | 0.0536 | -0.0574 | 5.338e-01 |
| RS3 | 2582 | 0.055 | -0.0414 | 4.972e-01 |
| RUSH1 | 184 | 0.0652 | 0.0137 | 9.479e-01 |
| RUSH2 | 106 | 0.0425 | 0.1175 | 7.306e-01 |
| SHIP | 966 | 0.0637 | 0.0443 | 6.352e-01 |
| SHIP-TREND | 858 | 0.0664 | -0.0145 | 8.805e-01 |
| SYS.adolescents | 980 | 0.0781 | -0.017 | 8.396e-01 |
| SYS.adults | 594 | 0.0724 | 0.0803 | 4.732e-01 |
| TREND | 1084 | 0.063 | -0.1239 | 1.613e-01 |
| UKB | 14571 | 0.0507 | -0.0693 | 9.475e-03 |
| WHICAP | 83 | 0.037 | -0.1007 | 8.075e-01 |
| **Non-European** | | | | |
| ARIC.AA | 675 | 0.0246 | -0.1515 | 3.888e-01 |
| CHAP.black | 321 | 0.0296 | -0.1155 | 6.206e-01 |
| FBIRN | 275 | 0.0819 | -0.2734 | 8.000e-02 |
| NOMAS.black | 168 | 0.0119 | -0.2968 | 5.559e-01 |
| NOMAS.hispanic | 718 | 0.0441 | 0.0384 | 7.650e-01 |
| SCES | 209 | 0.0909 | -0.235 | 1.672e-01 |
| SIMES | 201 | 0.1169 | -0.0722 | 6.419e-01 |
| WHICAP.black | 60 | 0.025 | 0.7864 | 1.845e-01 |
| **Summary European MA** | **74204** | **0.0557** | **-0.063** | **2.127e-08** |
| **Summary African MA** | **1224** | **0.0242** | **-0.108** | **0.4134** |
| **Summary Asian MA** | **1298** | **0.0971** | **0.036** | **0.5898** |
| **Summary Transancestral MA** | **77833** | **0.0559** | **-0.062** | **2.451e-08** |

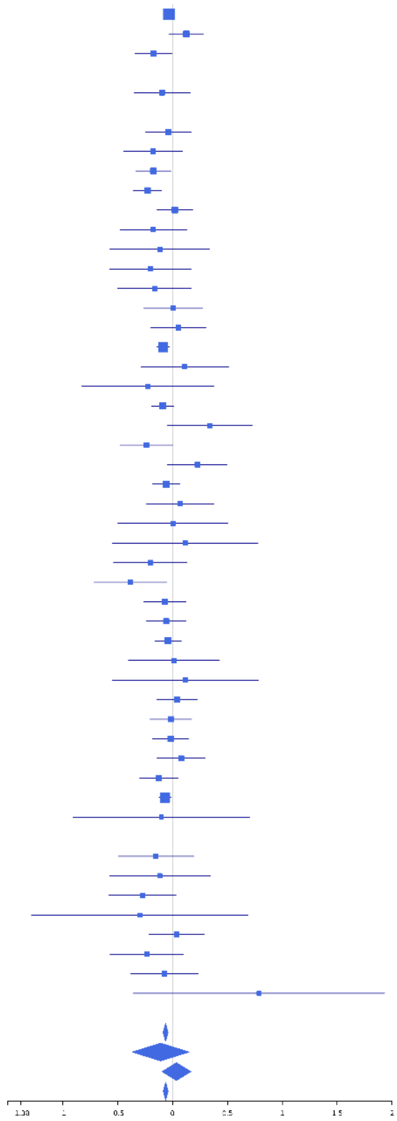

**rs10140304 (A); chr14:23443514; model 1 (I2=23.8, HetP=0.07191)**

| Study | N_total | EAF | Beta | P |
|---|---|---|---|---|
| **Head circumference** | | | | |
| Haworth | 18881 | 0.5974 | 0.0239 | 2.238e-02 |
| HKOS | 888 | 0.7866 | -0.0797 | 1.687e-01 |
| VIDI | 735 | 0.5937 | -0.114 | 3.220e-02 |
| **CT-based** | | | | |
| UCL | 642 | 0.6027 | 0.0416 | 4.654e-01 |
| **MR-based** | | | | |
| 1000Brains | 751 | 0.6121 | 0.0691 | 1.923e-01 |
| 3C-Bordeaux | 564 | 0.6078 | 0.0821 | 1.785e-01 |
| 3c-Dijon | 1499 | 0.6078 | 0.1044 | 5.247e-03 |
| AGES | 2226 | 0.6031 | 0.0204 | 5.063e-01 |
| ARIC.EA | 1390 | 0.5996 | -0.0104 | 7.882e-01 |
| ASPS-Family | 338 | 0.6199 | 0.0013 | 9.869e-01 |
| ASRB | 233 | 0.599 | 0.0125 | 8.948e-01 |
| CARDIA.EA | 349 | 0.6029 | 0.2018 | 9.083e-03 |
| CHAP | 261 | 0.6014 | 0.0123 | 8.901e-01 |
| CHS | 544 | 0.5795 | 0.0159 | 7.960e-01 |
| DNGS | 516 | 0.6418 | 0.0706 | 2.773e-01 |
| ENIGMA | 11373 | 0.6019 | 0.0131 | 3.337e-01 |
| EPOZ | 251 | 0.5933 | -0.2233 | 1.396e-02 |
| ERF | 122 | 0.6021 | 0.1448 | 2.683e-01 |
| FHS | 4710 | 0.5946 | 0.0689 | 1.029e-03 |
| GenR | 265 | 0.6012 | -0.0132 | 8.818e-01 |
| HUNT | 851 | 0.5642 | 0.0443 | 3.647e-01 |
| LBC1936 | 577 | 0.6049 | 0.0416 | 4.894e-01 |
| LIFE | 1825 | 0.605 | -0.0126 | 7.108e-01 |
| LLS | 355 | 0.6193 | 0.0611 | 4.295e-01 |
| MethCT. | 114 | 0.5434 | 0.0443 | 7.390e-01 |
| NOMAS | 141 | 0.5513 | -0.0777 | 5.176e-01 |
| PoznanMS | 204 | 0.6318 | 0.0751 | 4.643e-01 |
| PROSPER | 315 | 0.6056 | -0.083 | 3.085e-01 |
| RS1 | 1103 | 0.6007 | -0.009 | 8.355e-01 |
| RS2 | 1159 | 0.5966 | 0.0257 | 5.447e-01 |
| RS3 | 2582 | 0.6008 | -0.0032 | 9.114e-01 |
| RUSH1 | 184 | 0.6043 | 0.1544 | 1.495e-01 |
| RUSH2 | 106 | 0.6017 | 0.0358 | 7.995e-01 |
| SHIP | 966 | 0.6191 | 0.0204 | 6.632e-01 |
| SHIP-TREND | 858 | 0.6206 | 0.0871 | 8.022e-02 |
| SYS.adolescents | 980 | 0.5913 | -0.0165 | 7.190e-01 |
| SYS.adults | 594 | 0.5893 | 0.0745 | 2.066e-01 |
| TREND | 1084 | 0.6109 | -0.0039 | 9.297e-01 |
| UKB | 14571 | 0.6024 | 0.0586 | 9.876e-07 |
| WHICAP | 83 | 0.5864 | 0.1403 | 3.761e-01 |
| **Non-European** | | | | |
| ARIC.AA | 675 | 0.3841 | 0.0988 | 7.755e-02 |
| CARDIA.AA | 132 | 0.378 | 0.1545 | 2.237e-01 |
| CHAP.black | 321 | 0.4143 | 0.0046 | 9.541e-01 |
| FBIRN | 275 | 0.6229 | -0.1027 | 2.445e-01 |
| NOMAS.black | 168 | 0.3681 | -0.0526 | 6.423e-01 |
| NOMAS.hispanic | 718 | 0.522 | 0.0642 | 2.249e-01 |
| SCES | 209 | 0.7679 | 0.1183 | 3.074e-01 |
| SIMES | 201 | 0.8157 | 0.1585 | 2.188e-01 |
| WHICAP.black | 60 | 0.4 | -0.0829 | 6.579e-01 |
| **Summary European MA** | **74188** | **0.6007** | **0.031** | **5.089e-09** |
| **Summary African MA** | **1356** | **0.3894** | **0.055** | **0.1626** |
| **Summary Asian MA** | **1298** | **0.7881** | **-0.012** | **0.8066** |
| **Summary Transancestral MA** | **77949** | **0.5994** | **0.031** | **3.336e-09** |

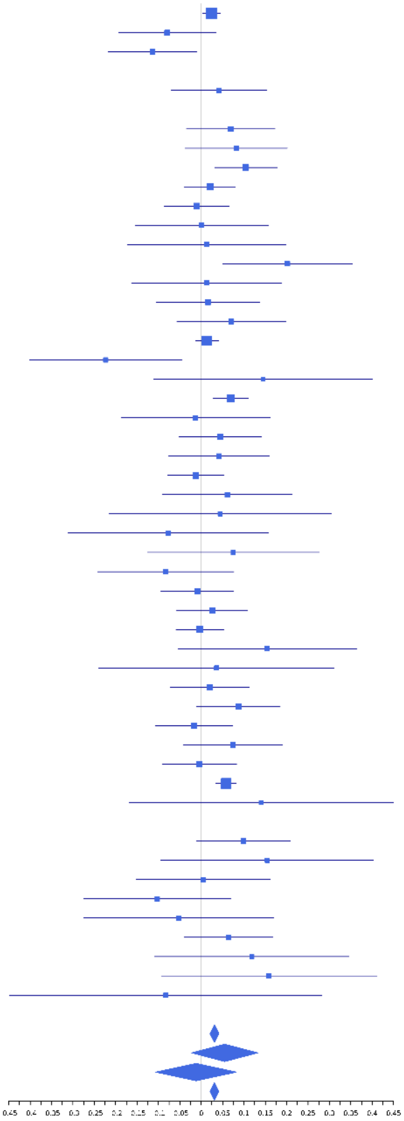

**rs893725 (A); chr15:90128223; model 1 (I2=0, HetP=0.8747)**

| Study | N_total | EAF | Beta | P |
|---|---|---|---|---|
| **Head circumference** | | | | |
| Haworth | 18881 | 0.5959 | 0.0224 | 3.160e-02 |
| HKOS | 888 | 0.5214 | -0.0156 | 7.431e-01 |
| VIDI | 741 | 0.6699 | 0.0494 | 3.718e-01 |
| **CT-based** | | | | |
| UCL | 642 | 0.6152 | 0.0707 | 2.181e-01 |
| **MR-based** | | | | |
| 3C-Bordeaux | 564 | 0.6027 | 0.0711 | 2.428e-01 |
| 3c-Dijon | 1499 | 0.596 | 0.016 | 6.672e-01 |
| AGES | 2226 | 0.556 | 0.0847 | 4.989e-03 |
| ARIC.EA | 1390 | 0.6035 | -0.004 | 9.175e-01 |
| ASPS-Family | 338 | 0.5426 | -0.0671 | 3.846e-01 |
| ASRB | 232 | 0.6111 | 0.0611 | 5.219e-01 |
| CARDIA.EA | 349 | 0.6233 | 0.0037 | 9.617e-01 |
| CHAP | 261 | 0.6054 | 0.1424 | 1.131e-01 |
| CHS | 544 | 0.5719 | 0.0654 | 2.856e-01 |
| DNGS | 516 | 0.5966 | 0.0431 | 4.969e-01 |
| ENIGMA | 11373 | 0.5995 | 0.02 | 1.389e-01 |
| EPOZ | 251 | 0.607 | 5e-04 | 9.961e-01 |
| ERF | 122 | 0.5785 | 0.1193 | 3.576e-01 |
| FHS | 4710 | 0.6048 | 0.0118 | 5.764e-01 |
| GenR | 265 | 0.5939 | 0.0588 | 5.062e-01 |
| HUNT | 851 | 0.5897 | 0.013 | 7.924e-01 |
| LBC1936 | 577 | 0.6131 | -0.0248 | 6.808e-01 |
| LIFE | 1825 | 0.6074 | -0.0173 | 6.094e-01 |
| LLS | 355 | 0.6237 | -0.0045 | 9.540e-01 |
| MethCT | 114 | 0.521 | 0.1369 | 3.014e-01 |
| NOMAS | 141 | 0.6183 | 0.1337 | 2.774e-01 |
| PoznanMS | 204 | 0.6305 | 0.108 | 2.922e-01 |
| PROSPER | 315 | 0.6009 | 0.0584 | 4.726e-01 |
| RS1 | 1103 | 0.6038 | 0.0528 | 2.248e-01 |
| RS2 | 1159 | 0.5994 | 0.1299 | 2.177e-03 |
| RS3 | 2582 | 0.5934 | 0.037 | 1.916e-01 |
| RUSH1 | 184 | 0.6142 | 0.1038 | 3.341e-01 |
| RUSH2 | 106 | 0.6226 | -0.1053 | 4.590e-01 |
| SHIP | 966 | 0.6158 | 0.0311 | 5.062e-01 |
| SHIP-TREND | 858 | 0.5985 | 0.0073 | 8.823e-01 |
| SYS.adolescents | 980 | 0.5875 | 0.0744 | 1.051e-01 |
| SYS.adults | 594 | 0.5803 | 0.1336 | 2.309e-02 |
| TREND | 1084 | 0.6121 | 0.0796 | 7.108e-02 |
| UKB | 14571 | 0.6008 | 0.0414 | 5.349e-04 |
| WHICAP | 83 | 0.6358 | -0.1439 | 3.750e-01 |
| **Non-European** | | | | |
| ARIC.AA | 675 | 0.6357 | -0.0243 | 6.675e-01 |
| CARDIA.AA | 132 | 0.648 | -0.0885 | 4.921e-01 |
| CHAP.black | 321 | 0.6402 | 0.0578 | 4.826e-01 |
| FBIRN | 275 | 0.6212 | 0.0024 | 9.788e-01 |
| IMH | 37 | 0.5526 | 0.152 | 5.154e-01 |
| NOMAS.black | 168 | 0.6265 | 0.1721 | 1.290e-01 |
| NOMAS.hispanic | 718 | 0.6231 | -0.0108 | 8.421e-01 |
| SCES | 209 | 0.511 | 0.059 | 5.468e-01 |
| SIMES | 201 | 0.5163 | -0.0015 | 9.880e-01 |
| WHICAP.black | 60 | 0.6583 | 0.0085 | 9.650e-01 |
| **Summary European MA** | **73442** | **0.5988** | **0.032** | **1.638e-09** |
| **Summary African MA** | **1356** | **0.6378** | **0.015** | **0.7131** |
| **Summary Asian MA** | **1335** | **0.5199** | **0.003** | **0.9414** |
| **Summary Transancestral MA** | **77240** | **0.5984** | **0.031** | **2.515e-09** |

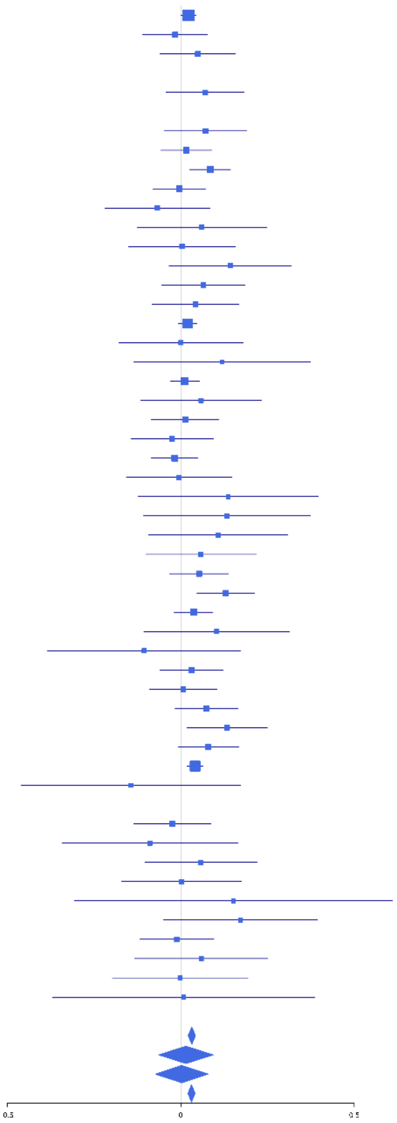

**rs62039480 (A); chr16:15137450; model 1 (I2=5, HetP=0.3753)**

| Study | N_total | EAF | Beta | P |
|---|---|---|---|---|
| **Head circumference** | | | | |
| Haworth | 18881 | 0.3334 | -0.0192 | 7.828e-02 |
| HKOS | 888 | 0.4167 | 0.0062 | 8.977e-01 |
| VIDI | 753 | 0.3214 | -0.0063 | 9.086e-01 |
| **CT-based** | | | | |
| ESS | 641 | 0.3539 | -0.0853 | 1.444e-01 |
| UCL | 642 | 0.337 | -0.078 | 1.866e-01 |
| **MR-based** | | | | |
| 3C-Bordeaux | 564 | 0.3403 | -0.0619 | 3.249e-01 |
| 3c-Dijon | 1499 | 0.3426 | -0.0401 | 2.968e-01 |
| AGES | 2226 | 0.2882 | -0.0119 | 7.200e-01 |
| ARIC.EA | 1390 | 0.3395 | -0.0872 | 2.939e-02 |
| ASPS-Family | 338 | 0.3815 | 0.0306 | 6.986e-01 |
| ASRB | 233 | 0.2841 | 0.1696 | 1.001e-01 |
| CARDIA.EA | 349 | 0.3405 | -0.0476 | 5.515e-01 |
| CHAP | 261 | 0.3262 | 0.0599 | 5.211e-01 |
| CHS | 544 | 0.3538 | 0.0035 | 9.565e-01 |
| ENIGMA | 11373 | 0.3359 | -0.0425 | 2.474e-03 |
| EPOZ | 251 | 0.3364 | 0.1699 | 7.201e-02 |
| ERF | 122 | 0.3328 | 0.0312 | 8.179e-01 |
| FHS | 4710 | 0.3095 | -0.039 | 7.974e-02 |
| GenR | 265 | 0.3396 | 0.0242 | 7.916e-01 |
| HUNT | 851 | 0.3084 | -0.0726 | 1.664e-01 |
| LBC1936 | 577 | 0.3013 | -0.0421 | 5.121e-01 |
| LIFE | 1825 | 0.3536 | 0.0066 | 8.489e-01 |
| LLS | 355 | 0.3338 | -0.0709 | 3.729e-01 |
| MethCT. | 114 | 0.3996 | 0.093 | 4.915e-01 |
| NOMAS | 141 | 0.4127 | 0.0421 | 7.281e-01 |
| PoznanMS | 204 | 0.3631 | -0.0997 | 3.329e-01 |
| PROSPER | 315 | 0.2926 | 0.0362 | 6.793e-01 |
| RS1 | 1103 | 0.3296 | -0.0476 | 2.935e-01 |
| RS2 | 1159 | 0.3269 | -0.0681 | 1.241e-01 |
| RS3 | 2582 | 0.3286 | 0.0268 | 3.649e-01 |
| RUSH1 | 184 | 0.3924 | 0.0629 | 5.565e-01 |
| RUSH2 | 106 | 0.3079 | 0.3397 | 2.458e-02 |
| SHIP | 966 | 0.3817 | -0.0631 | 1.778e-01 |
| SHIP-TREND | 858 | 0.3724 | -0.0536 | 2.833e-01 |
| SYS.adolescents | 980 | 0.3003 | -0.14 | 4.498e-03 |
| SYS.adults | 594 | 0.3216 | -0.0575 | 3.552e-01 |
| TREND | 1084 | 0.3496 | -0.063 | 1.622e-01 |
| UKB | 14571 | 0.3367 | -0.0365 | 3.271e-03 |
| WHICAP | 83 | 0.3481 | 0.2227 | 1.755e-01 |
| **Non-European** | | | | |
| ARIC.AA | 675 | 0.3197 | -0.05 | 3.921e-01 |
| CARDIA.AA | 132 | 0.313 | -0.0936 | 4.811e-01 |
| CHAP.black | 321 | 0.3341 | 0.0038 | 9.635e-01 |
| FBIRN | 275 | 0.3976 | 0.132 | 1.311e-01 |
| NOMAS.black | 168 | 0.3164 | -0.0022 | 9.850e-01 |
| NOMAS.hispanic | 718 | 0.3995 | -0.0929 | 8.492e-02 |
| SCES | 209 | 0.439 | -0.0637 | 5.185e-01 |
| SIMES | 201 | 0.4051 | 0.0605 | 5.458e-01 |
| WHICAP.black | 60 | 0.32 | -0.2266 | 2.520e-01 |
| **Summary European MA** | **73580** | **0.3329** | **-0.031** | **3.008e-08** |
| **Summary African MA** | **1356** | **0.3221** | **-0.043** | **0.2942** |
| **Summary Asian MA** | **1298** | **0.4185** | **0.003** | **0.9323** |
| **Summary Transancestral MA** | **77341** | **0.335** | **-0.03** | **2.377e-08** |

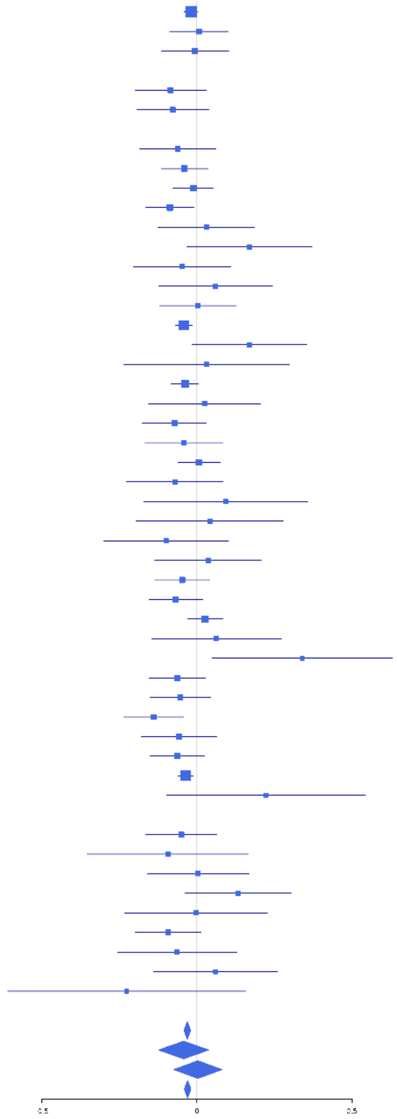

**rs78378222 (T); chr17:7571752; model 1 (I2=0, HetP=0.6119)**

| Study | N_total | EAF | Beta | P |
|---|---|---|---|---|
| **Head circumference** | | | | |
| Haworth | 17737 | 0.9827 | -0.2137 | 1.540e-07 |
| VIDI | 755 | 0.985 | 0.1055 | 6.176e-01 |
| **CT-based** | | | | |
| ESS | 641 | 0.9817 | -0.0874 | 6.751e-01 |
| UCL | 642 | 0.9914 | -0.067 | 8.240e-01 |
| **MR-based** | | | | |
| 3c-Dijon | 1499 | 0.9906 | -0.0727 | 7.008e-01 |
| AGES | 2226 | 0.9816 | -0.1488 | 1.819e-01 |
| ARIC.EA | 1390 | 0.9867 | -0.0938 | 5.716e-01 |
| CHS | 544 | 0.9926 | -0.4731 | 1.826e-01 |
| ENIGMA | 11265 | 0.9847 | -0.1049 | 5.338e-02 |
| ERF | 122 | 0.9776 | -0.3238 | 4.540e-01 |
| HUNT | 851 | 0.9762 | -0.0948 | 5.508e-01 |
| LBC1936 | 577 | 0.9824 | -0.368 | 1.001e-01 |
| LIFE | 1825 | 0.9862 | -0.2646 | 6.269e-02 |
| LLS | 355 | 0.9769 | -0.4377 | 7.964e-02 |
| PROSPER | 315 | 0.9828 | -0.5728 | 6.184e-02 |
| RS1 | 1103 | 0.9815 | -0.2514 | 1.113e-01 |
| RS2 | 1159 | 0.98 | -0.1167 | 4.312e-01 |
| RS3 | 2582 | 0.9813 | -0.1953 | 5.725e-02 |
| SHIP | 966 | 0.984 | -0.3691 | 4.179e-02 |
| SHIP-TREND | 858 | 0.9894 | -0.1698 | 4.718e-01 |
| SYS.adolescents | 980 | 0.9874 | -0.0636 | 7.537e-01 |
| SYS.adults | 594 | 0.9883 | -0.6751 | 1.253e-02 |
| TREND | 1084 | 0.9841 | -0.1835 | 2.859e-01 |
| UKB | 14571 | 0.9874 | -0.1455 | 5.509e-03 |
| WHICAP | 83 | 0.9799 | 0.9938 | 7.631e-02 |
| **Non-European** | | | | |
| **Summary European MA** | **64724** | **0.9846** | **-0.171** | **3.033e-14** |
| **Summary Transancestral MA** | **64724** | **0.9846** | **-0.171** | **3.033e-14** |

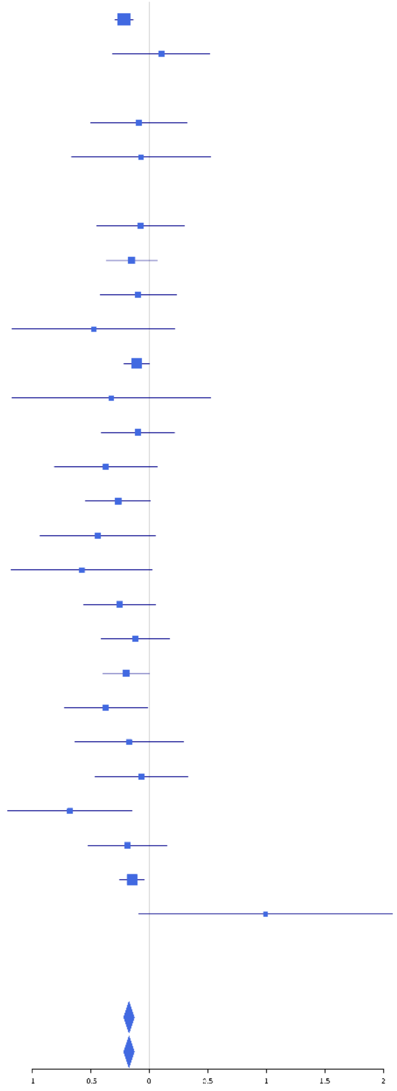

### rs12449730 (A); chr17:27881366; model 1 (I2=3.7, HetP=0.3995)

| Study | N_total | EAF | Beta | P |
|---|---|---|---|---|
| **Head circumference** | | | | |
| Haworth | 18881 | 0.1754 | 0.0292 | 3.102e-02 |
| HKOS | 888 | 0.089 | 0.0164 | 8.436e-01 |
| VIDI | 745 | 0.1745 | 0.0539 | 4.297e-01 |
| **CT-based** | | | | |
| ESS | 641 | 0.1911 | -0.0077 | 9.131e-01 |
| UCL | 642 | 0.1583 | 0.0985 | 1.976e-01 |
| **MR-based** | | | | |
| 3C-Bordeaux | 564 | 0.1734 | -0.0296 | 7.063e-01 |
| 3c-Dijon | 1499 | 0.1791 | 0.0492 | 3.015e-01 |
| AGES | 2226 | 0.1607 | 0.0559 | 1.711e-01 |
| ARIC.EA | 1390 | 0.1702 | 0.0157 | 7.554e-01 |
| ASPS-Family | 338 | 0.2445 | -0.0363 | 6.851e-01 |
| ASRB | 233 | 0.177 | -0.0665 | 5.840e-01 |
| CARDIA.EA | 349 | 0.1851 | 0.0681 | 4.848e-01 |
| CHAP | 261 | 0.1609 | 0.1014 | 3.955e-01 |
| CHS | 544 | 0.1584 | 0.0913 | 2.711e-01 |
| DHS | 460 | 0.148 | -0.096 | 3.009e-01 |
| DNGS | 515 | 0.1569 | 0.2433 | 4.687e-03 |
| ENIGMA | 11373 | 0.1813 | 0.0333 | 5.271e-02 |
| EPOZ | 251 | 0.2038 | -0.0224 | 8.396e-01 |
| ERF | 122 | 0.2001 | -0.121 | 4.495e-01 |
| FHS | 4710 | 0.1706 | 0.0927 | 7.177e-04 |
| GenR | 265 | 0.1795 | 0.0349 | 7.584e-01 |
| HUNT | 851 | 0.1462 | 0.0064 | 9.261e-01 |
| LBC1936 | 577 | 0.1499 | 0.1097 | 1.835e-01 |
| LIFE | 1825 | 0.196 | 0.0739 | 7.642e-02 |
| LLS | 355 | 0.2088 | -0.0582 | 5.286e-01 |
| MethCT. | 114 | 0.1656 | -0.0321 | 8.571e-01 |
| NOMAS | 141 | 0.2025 | 0.1279 | 3.897e-01 |
| PoznanMS | 204 | 0.2321 | -0.0877 | 4.542e-01 |
| PROSPER | 315 | 0.1862 | 0.1543 | 1.316e-01 |
| RS1 | 1103 | 0.1902 | 0.0551 | 3.101e-01 |
| RS2 | 1159 | 0.1854 | 0.0211 | 6.934e-01 |
| RS3 | 2582 | 0.1948 | 0.1354 | 1.171e-04 |
| RUSH1 | 184 | 0.232 | 0.1687 | 1.737e-01 |
| RUSH2 | 106 | 0.2028 | -0.0907 | 5.966e-01 |
| SHIP | 966 | 0.2046 | 0.0863 | 1.261e-01 |
| SHIP-TREND | 858 | 0.1836 | 0.0754 | 2.268e-01 |
| SYS.adolescents | 980 | 0.1858 | -0.0454 | 4.346e-01 |
| SYS.adults | 594 | 0.1939 | -0.1012 | 1.678e-01 |
| TREND | 1084 | 0.2167 | 0.1219 | 1.952e-02 |
| UKB | 14571 | 0.1733 | 0.0612 | 7.620e-05 |
| WHICAP | 83 | 0.1605 | -0.0497 | 8.151e-01 |
| **Non-European** | | | | |
| ARIC.AA | 675 | 0.1568 | 0.0693 | 3.542e-01 |
| CARDIA.AA | 132 | 0.155 | -0.1347 | 4.285e-01 |
| CHAP.black | 321 | 0.1776 | -0.1167 | 2.595e-01 |
| FBIRN | 275 | 0.1911 | 0.1106 | 3.086e-01 |
| IMH | 37 | 0.114 | -0.3866 | 2.905e-01 |
| NOMAS.black | 168 | 0.1753 | 0.1776 | 2.176e-01 |
| NOMAS.hispanic | 718 | 0.1745 | 0.1303 | 6.140e-02 |
| SCES | 209 | 0.0897 | 0.1719 | 3.155e-01 |
| SIMES | 201 | 0.2014 | 0.007 | 9.431e-01 |
| WHICAP.black | 60 | 0.1417 | -0.0741 | 7.780e-01 |
| **Summary European MA** | **74547** | **0.1781** | **0.048** | **1.847e-12** |
| **Summary African MA** | **1356** | **0.1632** | **0.012** | **0.816** |
| **Summary Asian MA** | **1335** | **0.1067** | **0.026** | **0.682** |
| **Summary Transancestral MA** | **78345** | **0.1766** | **0.048** | **6.511e-13** |

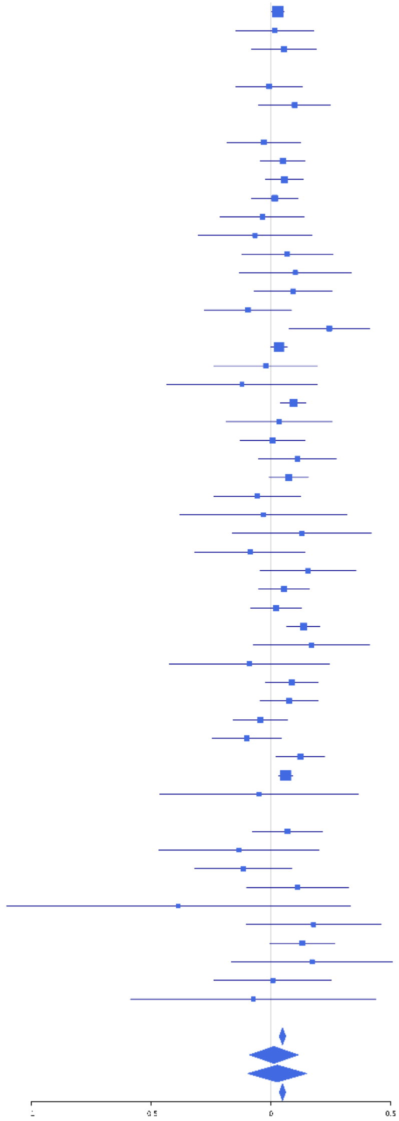

**rs7501777 (T); chr17:42638677; model 1 (I2=7.3, HetP=0.3285)**

| Study | N_total | EAF | Beta | P |
|---|---|---|---|---|
| **Head circumference** | | | | |
| Haworth | 18881 | 0.8331 | 0.0306 | 2.610e-02 |
| HKOS | 888 | 0.8913 | 0.0536 | 4.823e-01 |
| VIDI | 702 | 0.8279 | -0.0021 | 9.765e-01 |
| **CT-based** | | | | |
| ESS | 641 | 0.8312 | 0.1448 | 5.208e-02 |
| UCL | 642 | 0.8511 | 0.0646 | 4.098e-01 |
| **MR-based** | | | | |
| 3C-Bordeaux | 564 | 0.8255 | 0.193 | 1.419e-02 |
| 3c-Dijon | 1499 | 0.8283 | 0.0507 | 2.954e-01 |
| AGES | 2226 | 0.8655 | 0.0823 | 6.091e-02 |
| ARIC.EA | 1390 | 0.839 | -0.0196 | 7.043e-01 |
| ASPS-Family | 338 | 0.7753 | 0.0988 | 2.837e-01 |
| ASRB | 233 | 0.86 | 0.0816 | 5.417e-01 |
| CARDIA.EA | 349 | 0.8451 | 0.0953 | 3.623e-01 |
| CHAP | 261 | 0.8231 | 0.0868 | 4.499e-01 |
| CHS | 544 | 0.8213 | -0.0539 | 4.958e-01 |
| DNGS | 516 | 0.7835 | -0.0011 | 9.891e-01 |
| ENIGMA | 11373 | 0.8421 | -5e-04 | 9.763e-01 |
| EPOZ | 251 | 0.8312 | -0.1724 | 1.480e-01 |
| ERF | 122 | 0.8675 | -0.2478 | 1.894e-01 |
| FHS | 4710 | 0.849 | 0.0375 | 1.920e-01 |
| GenR | 265 | 0.7785 | 0.0925 | 3.765e-01 |
| HUNT | 851 | 0.8559 | 0.0723 | 2.944e-01 |
| LBC1936 | 577 | 0.8721 | 0.2074 | 1.863e-02 |
| LIFE | 1825 | 0.8203 | 0.0882 | 4.095e-02 |
| LLS | 355 | 0.8187 | 0.0272 | 7.804e-01 |
| MethCT. | 114 | 0.7195 | 0.0133 | 9.282e-01 |
| NOMAS | 141 | 0.8333 | 0.1601 | 3.183e-01 |
| PoznanMS | 204 | 0.8047 | 0.1279 | 3.059e-01 |
| PROSPER | 315 | 0.8286 | 0.1703 | 1.073e-01 |
| RS1 | 1103 | 0.8388 | 0.1229 | 3.376e-02 |
| RS2 | 1159 | 0.8289 | 0.0652 | 2.371e-01 |
| RS3 | 2582 | 0.8394 | 0.0713 | 5.997e-02 |
| RUSH1 | 184 | 0.8607 | 0.0343 | 8.200e-01 |
| RUSH2 | 106 | 0.8581 | 0.3537 | 7.544e-02 |
| SHIP | 966 | 0.8409 | 0.0174 | 7.794e-01 |
| SHIP-TREND | 858 | 0.839 | 0.0439 | 5.037e-01 |
| SYS.adolescents | 980 | 0.7895 | -0.001 | 9.857e-01 |
| SYS.adults | 594 | 0.7857 | -0.0724 | 3.057e-01 |
| TREND | 1084 | 0.8286 | 0.0262 | 6.459e-01 |
| UKB | 14571 | 0.8591 | 0.0493 | 3.418e-03 |
| WHICAP | 83 | 0.7535 | 0.1488 | 4.115e-01 |
| **Non-European** | | | | |
| ARIC.AA | 675 | 0.5926 | -0.0408 | 4.620e-01 |
| CARDIA.AA | 132 | 0.599 | 0.0412 | 7.431e-01 |
| CHAP.black | 321 | 0.5941 | -0.1571 | 5.146e-02 |
| FBIRN | 275 | 0.8089 | 0.0044 | 9.669e-01 |
| NOMAS.black | 168 | 0.6458 | 0.2337 | 4.204e-02 |
| NOMAS.hispanic | 718 | 0.7303 | -0.0207 | 7.279e-01 |
| SCES | 209 | 0.8654 | 0.1793 | 2.109e-01 |
| SIMES | 201 | 0.8856 | 0.1752 | 2.642e-01 |
| WHICAP.black | 60 | 0.6833 | -0.0846 | 6.685e-01 |
| **Summary European MA** | **74045** | **0.8404** | **0.039** | **3.96e-08** |
| **Summary African MA** | **1356** | **0.6042** | **-0.029** | **0.46** |
| **Summary Asian MA** | **1298** | **0.8863** | **0.094** | **0.1279** |
| **Summary Transancestral MA** | **77806** | **0.8357** | **0.037** | **5.641e-08** |

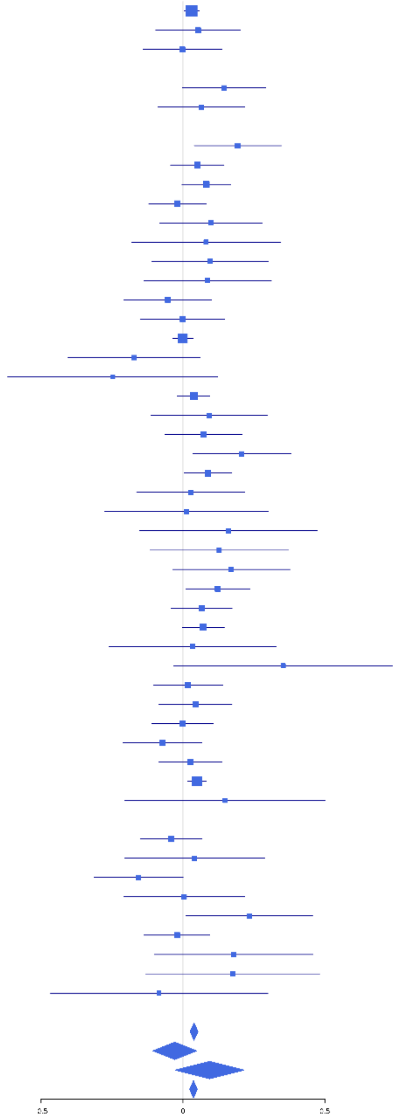

**rs112550936 (A); chr17:43716155; model 1 (I2=0, HetP=0.9059)**

| Study | N_total | EAF | Beta | P |
|---|---|---|---|---|
| **Head circumference** | | | | |
| **CT-based** | | | | |
| **MR-based** | | | | |
| ASPS-Family | 338 | 0.857 | 0.2041 | 6.313e-02 |
| CARDIA.EA | 349 | 0.7865 | 0.221 | 1.670e-02 |
| CHAP | 261 | 0.7935 | 0.0325 | 7.634e-01 |
| CHS | 544 | 0.8341 | 0.0882 | 2.791e-01 |
| ENIGMA | 341 | 0.9013 | 0.1462 | 2.546e-01 |
| FHS | 4710 | 0.8206 | 0.098 | 2.632e-04 |
| LIFE | 1825 | 0.8361 | 0.0852 | 5.677e-02 |
| NOMAS | 141 | 0.8462 | -0.068 | 6.812e-01 |
| RUSH1 | 184 | 0.8017 | 0.1551 | 2.372e-01 |
| RUSH2 | 106 | 0.7641 | 0.1529 | 3.470e-01 |
| SHIP | 966 | 0.8256 | 0.0186 | 7.566e-01 |
| SHIP-TREND | 858 | 0.8397 | 0.0949 | 1.498e-01 |
| SYS.adolescents | 980 | 0.7664 | 0.0768 | 1.503e-01 |
| SYS.adults | 594 | 0.773 | 0.0989 | 1.532e-01 |
| WHICAP | 83 | 0.7411 | 0.1689 | 3.438e-01 |
| **Non-European** | | | | |
| ARIC.AA | 675 | 0.9594 | 6e-04 | 9.970e-01 |
| CHAP.black | 321 | 0.9391 | 0.2094 | 2.055e-01 |
| NOMAS.black | 168 | 0.9527 | 0.5443 | 3.569e-02 |
| NOMAS.hispanic | 718 | 0.8831 | 0.0481 | 5.582e-01 |
| WHICAP.black | 60 | 0.9354 | -0.0698 | 8.519e-01 |
| **Summary European MA** | **12280** | **0.8193** | **0.093** | **1.891e-08** |
| **Summary African MA** | **1224** | **0.952** | **0.131** | **0.1651** |
| **Summary Transancestral MA** | **14222** | **0.8339** | **0.092** | **8.286e-09** |

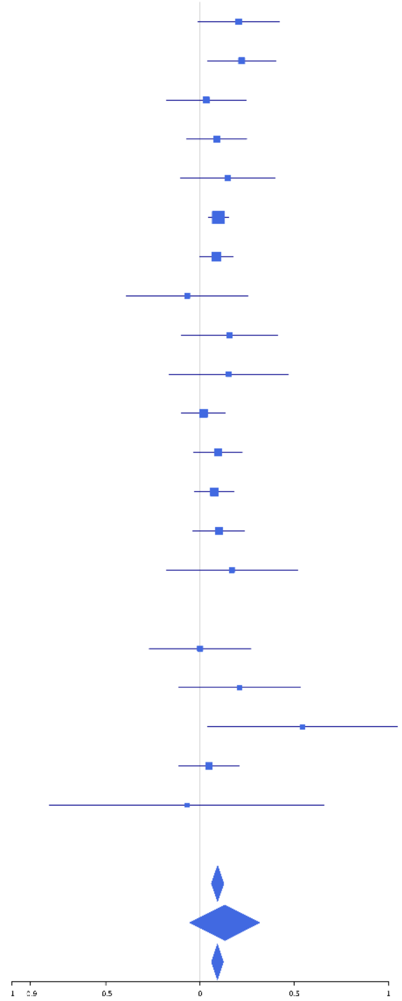

**rs117319001 (A); chr17:43798360; model 1 (I2=24.2, HetP=0.1009)**

| Study | N_total | EAF | Beta | P |
|---|---|---|---|---|
| **Head circumference** | | | | |
| Haworth | 18881 | 0.7337 | 0.0384 | 9.777e-04 |
| VIDI | 476 | 0.0585 | 0.057 | 6.800e-01 |
| **CT-based** | | | | |
| ESS | 641 | 0.799 | 0.1095 | 1.162e-01 |
| UCL | 642 | 0.7649 | 0.0051 | 9.384e-01 |
| **MR-based** | | | | |
| 3C-Bordeaux | 564 | 0.7505 | 0.0021 | 9.750e-01 |
| ARIC.EA | 1390 | 0.7894 | 0.0519 | 2.644e-01 |
| ASPS-Family | 338 | 0.84 | 0.1935 | 6.513e-02 |
| CARDIA.EA | 349 | 0.7871 | 0.2213 | 1.669e-02 |
| CHAP | 261 | 0.7913 | 0.033 | 7.596e-01 |
| CHS | 544 | 0.7681 | 0.0787 | 2.730e-01 |
| DHS | 460 | 0.7796 | 0.0484 | 5.434e-01 |
| ENIGMA | 341 | 0.8821 | 0.1224 | 3.028e-01 |
| EPOZ | 251 | 0.8034 | 0.1484 | 1.865e-01 |
| FHS | 4710 | 0.7679 | 0.0877 | 3.274e-04 |
| GenR | 265 | 0.7937 | 0.2831 | 8.354e-03 |
| HUNT | 851 | 0.7948 | 0.0545 | 3.639e-01 |
| LIFE | 1825 | 0.8143 | 0.0791 | 6.329e-02 |
| NOMAS | 141 | 0.8063 | -0.0967 | 5.217e-01 |
| PoznanMS | 204 | 0.829 | 0.1514 | 2.497e-01 |
| RS1 | 1103 | 0.7692 | 0.0811 | 1.082e-01 |
| RS2 | 1159 | 0.7623 | 0.1546 | 1.528e-03 |
| RS3 | 2582 | 0.7614 | 0.1273 | 9.629e-05 |
| RUSH1 | 184 | 0.8017 | 0.1549 | 2.377e-01 |
| RUSH2 | 106 | 0.7641 | 0.1525 | 3.478e-01 |
| SHIP | 966 | 0.802 | 0.0155 | 7.867e-01 |
| SHIP-TREND | 858 | 0.0012 | -1.1985 | 9.054e-02 |
| SYS.adolescents | 980 | 0.6744 | 0.0608 | 2.072e-01 |
| SYS.adults | 594 | 0.6779 | 0.0933 | 1.330e-01 |
| TREND | 1084 | 0.8068 | 0.1995 | 2.566e-04 |
| WHICAP | 83 | 0.6828 | 0.1009 | 5.470e-01 |
| **Non-European** | | | | |
| ARIC.AA | 675 | 0.9485 | -0.0188 | 8.785e-01 |
| CHAP.black | 321 | 0.9373 | 0.1708 | 2.952e-01 |
| NOMAS.black | 168 | 0.9353 | 0.4495 | 4.430e-02 |
| NOMAS.hispanic | 718 | 0.8423 | 0.0481 | 5.066e-01 |
| WHICAP.black | 60 | 0.9243 | -0.0928 | 7.892e-01 |
| **Summary European MA** | **42833** | **0.7335** | **0.062** | **1.127e-15** |
| **Summary African MA** | **1224** | **0.9426** | **0.096** | **0.2677** |
| **Summary Transancestral MA** | **44775** | **0.741** | **0.062** | **5.322e-16** |

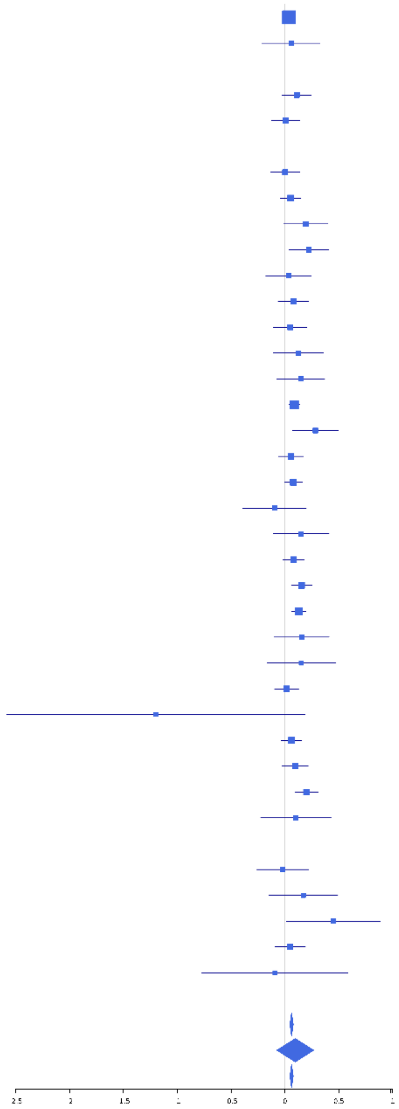

**rs4564621 (C); chr17:43895501; model 1 (I2=8.8, HetP=0.3106)**

| Study | N_total | EAF | Beta | P |
|---|---|---|---|---|
| **Head circumference** | | | | |
| Haworth | 18881 | 0.2202 | -0.0433 | 4.801e-04 |
| VIDI | 755 | 0.0891 | -0.1419 | 1.165e-01 |
| **CT-based** | | | | |
| ESS | 641 | 0.2005 | -0.1122 | 1.076e-01 |
| UCL | 642 | 0.2343 | -0.0045 | 9.454e-01 |
| **MR-based** | | | | |
| 3C-Bordeaux | 564 | 0.2499 | -0.0021 | 9.750e-01 |
| ARIC.EA | 1390 | 0.2112 | -0.0518 | 2.648e-01 |
| ASPS-Family | 338 | 0.1613 | -0.2016 | 5.388e-02 |
| ASRB | 233 | 0.2261 | -0.0771 | 4.868e-01 |
| CARDIA.EA | 349 | 0.2129 | -0.2214 | 1.668e-02 |
| CHAP | 261 | 0.2032 | -0.0375 | 7.303e-01 |
| CHS | 544 | 0.2298 | -0.0755 | 2.946e-01 |
| DHS | 460 | 0.2193 | -0.0492 | 5.375e-01 |
| ENIGMA | 11096 | 0.2155 | -0.0956 | 4.713e-09 |
| EPOZ | 251 | 0.1967 | -0.1461 | 1.932e-01 |
| FHS | 4710 | 0.2291 | -0.0912 | 2.004e-04 |
| GenR | 265 | 0.2066 | -0.2939 | 6.169e-03 |
| HUNT | 851 | 0.1769 | -0.0436 | 4.920e-01 |
| LIFE | 1825 | 0.1863 | -0.0718 | 9.158e-02 |
| MethCT. | 114 | 0.0995 | -0.3768 | 8.864e-02 |
| NOMAS | 141 | 0.1928 | 0.0962 | 5.251e-01 |
| PoznanMS | 204 | 0.1751 | -0.1394 | 2.846e-01 |
| RS1 | 1103 | 0.2335 | -0.0775 | 1.237e-01 |
| RS2 | 1159 | 0.2396 | -0.154 | 1.552e-03 |
| RS3 | 2582 | 0.2408 | -0.1274 | 9.077e-05 |
| RUSH1 | 184 | 0.1983 | -0.1549 | 2.376e-01 |
| RUSH2 | 106 | 0.2357 | -0.1529 | 3.472e-01 |
| SHIP | 966 | 0.196 | -0.0191 | 7.381e-01 |
| SHIP-TREND | 858 | 0.201 | -0.1122 | 6.280e-02 |
| SYS.adolescents | 980 | 0.3328 | -0.0469 | 3.282e-01 |
| SYS.adults | 594 | 0.3282 | -0.0981 | 1.124e-01 |
| TREND | 1084 | 0.1932 | -0.1976 | 2.927e-04 |
| UKB | 14571 | 0.2242 | -0.0923 | 4.942e-11 |
| WHICAP | 83 | 0.3148 | -0.0774 | 6.448e-01 |
| **Non-European** | | | | |
| ARIC.AA | 675 | 0.0444 | -0.0118 | 9.289e-01 |
| CARDIA.AA | 132 | 0.046 | 0.0188 | 9.488e-01 |
| CHAP.black | 321 | 0.0609 | -0.2106 | 2.028e-01 |
| FBIRN | 275 | 0.1485 | 0.102 | 3.957e-01 |
| NOMAS.black | 168 | 0.0566 | -0.4748 | 4.595e-02 |
| NOMAS.hispanic | 718 | 0.1534 | -0.0498 | 4.970e-01 |
| SIMES | 201 | 0.0036 | -0.6613 | 4.066e-01 |
| WHICAP.black | 60 | 0.0649 | 0.0678 | 8.558e-01 |
| **Summary European MA** | **68671** | **0.2203** | **-0.08** | **1.44e-34** |
| **Summary African MA** | **1356** | **0.0509** | **-0.116** | **0.1846** |
| **Summary Asian MA** | **201** | **0.0036** | **-0.691** | **0.4066** |
| **Summary Transancestral MA** | **71335** | **0.2154** | **-0.079** | **5.392e-35** |

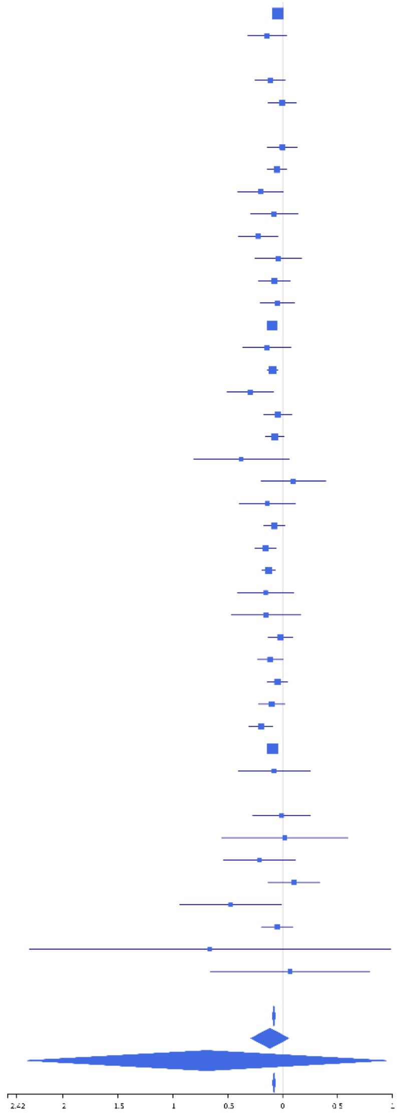

**rs191632554 (A); chr17:43981742; model 1 (I2=0, HetP=0.9465)**

| Study | N_total | EAF | Beta | P |
|---|---|---|---|---|
| **Head circumference** | | | | |
| **CT-based** | | | | |
| **MR-based** | | | | |
| ASPS-Family | 338 | 0.1633 | -0.1989 | 5.593e-02 |
| CARDIA.EA | 349 | 0.219 | -0.2119 | 2.059e-02 |
| CHAP | 261 | 0.2103 | -0.03 | 7.804e-01 |
| CHS | 544 | 0.2288 | -0.0758 | 2.936e-01 |
| DHS | 460 | 0.217 | -0.0558 | 4.853e-01 |
| ENIGMA | 341 | 0.1176 | -0.1328 | 2.639e-01 |
| FHS | 4710 | 0.2279 | -0.0927 | 1.594e-04 |
| LIFE | 1825 | 0.1824 | -0.0785 | 6.733e-02 |
| NOMAS | 141 | 0.1889 | 0.0933 | 5.409e-01 |
| RUSH1 | 184 | 0.2094 | -0.1585 | 2.179e-01 |
| RUSH2 | 106 | 0.2414 | -0.1539 | 3.397e-01 |
| SHIP | 966 | 0.1936 | -0.0348 | 5.454e-01 |
| SHIP-TREND | 858 | 0.1999 | -0.1143 | 5.872e-02 |
| SYS.adolescents | 980 | 0.3338 | -0.0321 | 5.023e-01 |
| SYS.adults | 594 | 0.3295 | -0.0593 | 3.368e-01 |
| WHICAP | 83 | 0.3163 | -0.0798 | 6.343e-01 |
| **Non-European** | | | | |
| ARIC.AA | 675 | 0.0489 | -0.0433 | 7.313e-01 |
| CHAP.black | 321 | 0.0665 | -0.1882 | 2.356e-01 |
| NOMAS.black | 168 | 0.0594 | -0.4494 | 5.329e-02 |
| NOMAS.hispanic | 718 | 0.1526 | -0.0393 | 5.930e-01 |
| WHICAP.black | 60 | 0.0675 | 0.0731 | 8.411e-01 |
| **Summary European MA** | **12740** | **0.2241** | **-0.083** | **3.489e-08** |
| **Summary African MA** | **1224** | **0.0559** | **-0.135** | **0.125** |
| **Summary Transancestral MA** | **14682** | **0.2066** | **-0.082** | **1.209e-08** |

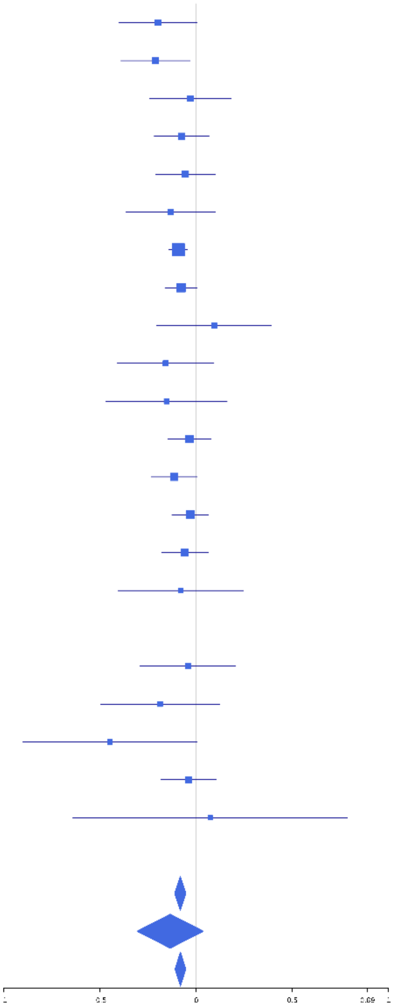

**rs8079695 (T); chr17:44161441; model 1 (I2=4.1, HetP=0.3956)**

| Study | N_total | EAF | Beta | P |
|---|---|---|---|---|
| **Head circumference** | | | | |
| Haworth | 18881 | 0.172 | 0.0347 | 1.086e-02 |
| HKOS | 888 | 0.3429 | 0.0776 | 1.206e-01 |
| VIDI | 743 | 0.2741 | 0.0355 | 5.411e-01 |
| **CT-based** | | | | |
| ESS | 641 | 0.1655 | 0.0411 | 5.842e-01 |
| UCL | 642 | 0.1865 | 0.0281 | 6.949e-01 |
| **MR-based** | | | | |
| 3C-Bordeaux | 564 | 0.1544 | 0.0518 | 5.299e-01 |
| ARIC.EA | 1390 | 0.1743 | 0.0613 | 2.199e-01 |
| ASPS-Family | 338 | 0.1496 | -0.0316 | 7.696e-01 |
| ASRB | 233 | 0.2182 | -0.0616 | 5.833e-01 |
| CARDIA.EA | 349 | 0.1696 | 0.2154 | 3.269e-02 |
| CHAP | 261 | 0.1686 | 0.0392 | 7.378e-01 |
| CHS | 544 | 0.1646 | 0.0548 | 5.027e-01 |
| DHS | 460 | 0.1888 | 0.0155 | 8.541e-01 |
| DNGS | 516 | 0.2327 | -0.0218 | 7.670e-01 |
| ENIGMA | 2339 | 0.1828 | 0.0875 | 2.079e-02 |
| EPOZ | 251 | 0.1956 | -0.0941 | 4.031e-01 |
| FHS | 4710 | 0.1766 | 0.0639 | 1.804e-02 |
| GenR | 265 | 0.1661 | 0.3105 | 7.807e-03 |
| HUNT | 851 | 0.2442 | 0.0473 | 4.019e-01 |
| LIFE | 1825 | 0.1731 | 0.1091 | 1.276e-02 |
| MethCT. | 114 | 0.0657 | 0.2376 | 3.738e-01 |
| NOMAS | 141 | 0.1587 | 0.3258 | 4.765e-02 |
| PoznanMS | 204 | 0.1685 | 0.1337 | 3.119e-01 |
| RS1 | 1103 | 0.1694 | 0.1349 | 1.746e-02 |
| RS2 | 1159 | 0.1651 | 3e-04 | 9.964e-01 |
| RS3 | 2582 | 0.1679 | 0.0583 | 1.171e-01 |
| RUSH1 | 184 | 0.1879 | -0.0137 | 9.183e-01 |
| RUSH2 | 106 | 0.1698 | 0.0494 | 7.877e-01 |
| SHIP | 966 | 0.1734 | -0.0165 | 7.840e-01 |
| SHIP-TREND | 858 | 0.1632 | 0.051 | 4.351e-01 |
| TREND | 1084 | 0.1756 | 0.0021 | 9.695e-01 |
| UKB | 14571 | 0.1884 | 0.0431 | 4.018e-03 |
| WHICAP | 83 | 0.1111 | -0.3678 | 1.407e-01 |
| **Non-European** | | | | |
| ARIC.AA | 675 | 0.0399 | 0.2645 | 5.714e-02 |
| CHAP.black | 321 | 0.0452 | -0.2132 | 2.628e-01 |
| FBIRN | 275 | 0.2218 | -0.0613 | 5.507e-01 |
| IMH | 37 | 0.3947 | -0.0057 | 9.806e-01 |
| NOMAS.black | 168 | 0.0428 | 0.3453 | 2.019e-01 |
| NOMAS.hispanic | 718 | 0.1371 | 0.1175 | 1.262e-01 |
| SCES | 209 | 0.3901 | -0.2223 | 2.661e-02 |
| SIMES | 201 | 0.2293 | 0.042 | 7.210e-01 |
| WHICAP.black | 60 | 0.0812 | -0.2246 | 5.043e-01 |
| **Summary European MA** | **58844** | **0.1794** | **0.046** | **1.109e-09** |
| **Summary African MA** | **1224** | **0.0437** | **0.115** | **0.2444** |
| **Summary Asian MA** | **1335** | **0.3346** | **0.021** | **0.6008** |
| **Summary Transancestral MA** | **62510** | **0.1796** | **0.047** | **2.733e-10** |

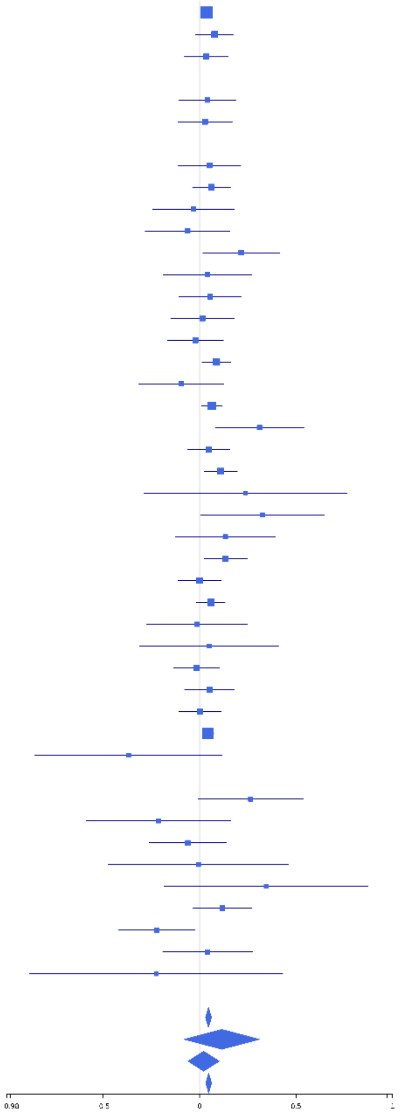

**rs76847569 (A); chr17:47096662; model 1 (I2=13.1, HetP=0.2474)**

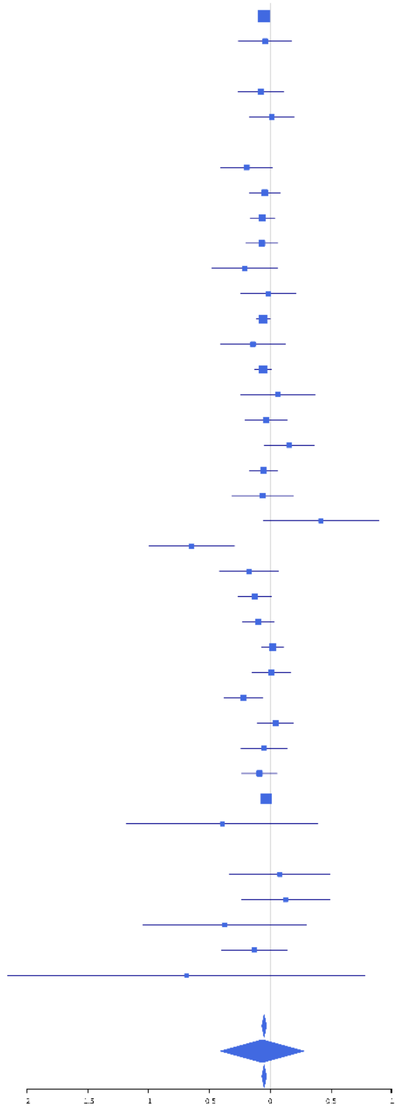

| Study | N_total | EAF | Beta | P |
|---|---|---|---|---|
| **Head circumference** | | | | |
| Haworth | 18881 | 0.1 | -0.051 | 2.899e-03 |
| VIDI | 724 | 0.0589 | -0.0425 | 7.034e-01 |
| **CT-based** | | | | |
| ESS | 641 | 0.0977 | -0.0775 | 4.100e-01 |
| UCL | 642 | 0.1027 | 0.0116 | 8.993e-01 |
| **MR-based** | | | | |
| 3C-Bordeaux | 564 | 0.0817 | -0.1942 | 7.465e-02 |
| 3c-Dijon | 1499 | 0.0844 | -0.0459 | 4.845e-01 |
| AGES | 2226 | 0.0918 | -0.0658 | 2.047e-01 |
| ARIC.EA | 1390 | 0.0874 | -0.0683 | 3.087e-01 |
| ASPS-Family | 338 | 0.0855 | -0.2106 | 1.257e-01 |
| CHS | 544 | 0.075 | -0.016 | 8.894e-01 |
| ENIGMA | 6715 | 0.0955 | -0.0596 | 4.240e-02 |
| EPOZ | 251 | 0.1211 | -0.1426 | 2.972e-01 |
| FHS | 4710 | 0.0913 | -0.0604 | 9.107e-02 |
| GenR | 265 | 0.0849 | 0.0609 | 6.957e-01 |
| HUNT | 851 | 0.0824 | -0.034 | 6.992e-01 |
| LBC1936 | 577 | 0.0877 | 0.1539 | 1.391e-01 |
| LIFE | 1825 | 0.0875 | -0.0558 | 3.409e-01 |
| LLS | 355 | 0.0942 | -0.0633 | 6.220e-01 |
| NOMAS | 141 | 0.0645 | 0.4158 | 8.855e-02 |
| PoznanMS | 204 | 0.0833 | -0.6465 | 3.072e-04 |
| PROSPER | 315 | 0.1163 | -0.1751 | 1.589e-01 |
| RS1 | 1103 | 0.1055 | -0.1264 | 6.830e-02 |
| RS2 | 1159 | 0.1102 | -0.0987 | 1.367e-01 |
| RS3 | 2582 | 0.1081 | 0.0178 | 6.912e-01 |
| SHIP | 966 | 0.0865 | 0.0083 | 9.184e-01 |
| SHIP-TREND | 858 | 0.0975 | -0.2221 | 6.487e-03 |
| SYS.adolescents | 980 | 0.1005 | 0.0425 | 5.724e-01 |
| SYS.adults | 594 | 0.1002 | -0.052 | 5.905e-01 |
| TREND | 1084 | 0.0914 | -0.0891 | 2.325e-01 |
| UKB | 14571 | 0.0931 | -0.033 | 1.012e-01 |
| WHICAP | 83 | 0.039 | -0.3949 | 3.278e-01 |
| **Non-European** | | | | |
| ARIC.AA | 675 | 0.017 | 0.0785 | 7.092e-01 |
| FBIRN | 275 | 0.0563 | 0.1252 | 4.993e-01 |
| NOMAS.black | 168 | 0.0261 | -0.375 | 2.747e-01 |
| NOMAS.hispanic | 718 | 0.039 | -0.1322 | 3.322e-01 |
| WHICAP.black | 60 | 0.0151 | -0.6894 | 3.613e-01 |
| **Summary European MA** | **67638** | **0.0951** | **-0.051** | **4.833e-08** |
| **Summary African MA** | **903** | **0.0186** | **-0.067** | **0.7009** |
| **Summary Transancestral MA** | **69534** | **0.0934** | **-0.051** | **4.2e-08** |

### rs7232135 (T); chr18:13047983; model 1 (I2=0, HetP=0.6482)

| Study | N_total | EAF | Beta | P |
|---|---|---|---|---|
| **Head circumference** | | | | |
| Haworth | 18881 | 0.7998 | 0.0382 | 2.990e-03 |
| HKOS | 888 | 0.5298 | 0.0163 | 7.321e-01 |
| VIDI | 755 | 0.804 | 0.0328 | 6.130e-01 |
| **CT-based** | | | | |
| ESS | 641 | 0.7918 | 0.1337 | 5.200e-02 |
| UCL | 642 | 0.8119 | -0.0418 | 5.576e-01 |
| **MR-based** | | | | |
| 3C-Bordeaux | 564 | 0.7977 | 0.1582 | 3.326e-02 |
| 3c-Dijon | 1499 | 0.7988 | 0.1808 | 7.216e-05 |
| AGES | 2226 | 0.8045 | 0.0936 | 1.324e-02 |
| ARIC.EA | 1390 | 0.7951 | 0.0427 | 3.641e-01 |
| ASPS-Family | 338 | 0.7713 | 0.0272 | 7.667e-01 |
| ASRB | 233 | 0.8147 | 0.0368 | 7.577e-01 |
| CARDIA.EA | 349 | 0.8067 | 0.1395 | 1.457e-01 |
| CHAP | 261 | 0.8004 | 0.2422 | 2.786e-02 |
| CHS | 544 | 0.7885 | 0.0365 | 6.224e-01 |
| DHS | 460 | 0.7868 | 0.0705 | 3.812e-01 |
| DNGS | 516 | 0.727 | 0.0161 | 8.177e-01 |
| ENIGMA | 11373 | 0.7997 | 0.032 | 5.327e-02 |
| EPOZ | 251 | 0.8282 | -0.0101 | 9.324e-01 |
| ERF | 122 | 0.8057 | -0.1069 | 5.089e-01 |
| FHS | 4710 | 0.8003 | 0.0267 | 2.995e-01 |
| GenR | 265 | 0.7837 | 0.0858 | 4.165e-01 |
| HUNT | 851 | 0.8056 | 0.0381 | 5.337e-01 |
| LBC1936 | 577 | 0.8005 | -0.0056 | 9.395e-01 |
| LIFE | 1825 | 0.7817 | 0.0372 | 3.534e-01 |
| LLS | 355 | 0.8382 | -0.0354 | 7.283e-01 |
| MethCT. | 114 | 0.6938 | 0.0649 | 6.514e-01 |
| NOMAS | 141 | 0.7625 | -0.097 | 4.896e-01 |
| PoznanMS | 204 | 0.7333 | -0.2059 | 6.598e-02 |
| PROSPER | 315 | 0.8286 | 0.0689 | 5.143e-01 |
| RS1 | 1103 | 0.8272 | 0.0474 | 4.004e-01 |
| RS2 | 1159 | 0.8208 | 0.0351 | 5.164e-01 |
| RS3 | 2582 | 0.8236 | 0.0595 | 1.033e-01 |
| RUSH1 | 184 | 0.8178 | 0.039 | 7.730e-01 |
| RUSH2 | 106 | 0.7735 | 0.083 | 6.142e-01 |
| SHIP | 966 | 0.7887 | 0.0229 | 6.809e-01 |
| SHIP-TREND | 858 | 0.8015 | 0.1254 | 3.861e-02 |
| SYS.adolescents | 980 | 0.7751 | 0.0032 | 9.527e-01 |
| SYS.adults | 594 | 0.7749 | -0.0985 | 1.562e-01 |
| TREND | 1084 | 0.7952 | 0.02 | 7.071e-01 |
| UKB | 14571 | 0.8115 | 0.0419 | 5.104e-03 |
| WHICAP | 83 | 0.7531 | -0.1301 | 4.717e-01 |
| **Non-European** | | | | |
| ARIC.AA | 675 | 0.7478 | 0.0267 | 6.699e-01 |
| CARDIA.AA | 132 | 0.752 | -0.055 | 6.995e-01 |
| CHAP.black | 321 | 0.7165 | 0.1415 | 1.072e-01 |
| FBIRN | 275 | 0.7713 | -0.0402 | 6.927e-01 |
| NOMAS.black | 168 | 0.8065 | -0.1087 | 4.325e-01 |
| NOMAS.hispanic | 718 | 0.7929 | 0.055 | 3.982e-01 |
| SCES | 209 | 0.5479 | 0.1233 | 2.095e-01 |
| SIMES | 201 | 0.4378 | -0.106 | 2.927e-01 |
| WHICAP.black | 60 | 0.825 | 0.0894 | 7.115e-01 |
| **Summary European MA** | **74558** | **0.802** | **0.042** | **1.318e-10** |
| **Summary African MA** | **1356** | **0.7515** | **0.034** | **0.4439** |
| **Summary Asian MA** | **1298** | **0.5185** | **0.015** | **0.7094** |
| **Summary Transancestral MA** | **78319** | **0.7961** | **0.041** | **8.454e-11** |

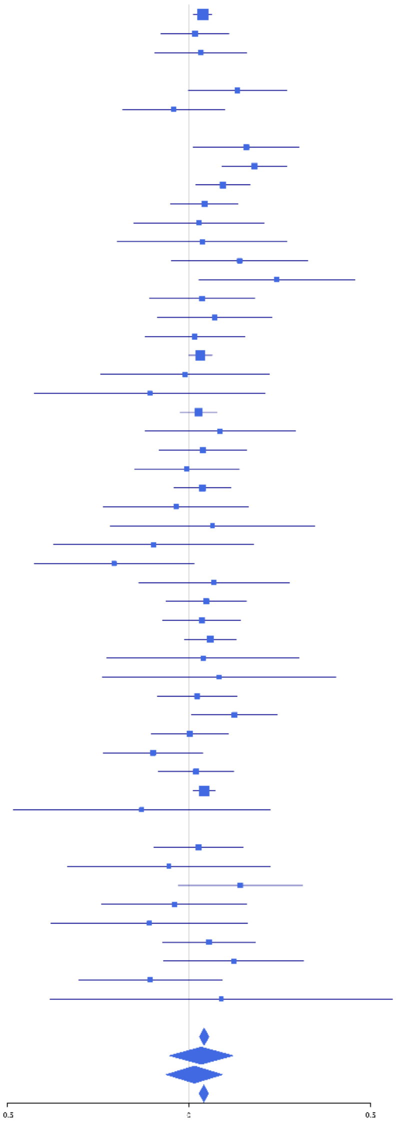

**rs35542154 (A); chr19:12782418; model 1 (I2=30.8, HetP=0.0284)**

| Study | N_total | EAF | Beta | P |
|---|---|---|---|---|
| **Head circumference** | | | | |
| Haworth | 18881 | 0.6226 | 0.0246 | 2.046e-02 |
| HKOS | 888 | 0.8367 | -0.0063 | 9.216e-01 |
| VIDI | 724 | 0.7271 | 0.0415 | 4.822e-01 |
| **CT-based** | | | | |
| UCL | 642 | 0.6309 | -0.0636 | 2.712e-01 |
| **MR-based** | | | | |
| 3C-Bordeaux | 564 | 0.6237 | 0.0516 | 4.019e-01 |
| 3c-Dijon | 1499 | 0.6201 | 0.0627 | 9.566e-02 |
| AGES | 2226 | 0.6923 | 0.0611 | 5.983e-02 |
| ARIC.EA | 1390 | 0.6448 | -0.0662 | 9.486e-02 |
| ASPS-Family | 338 | 0.5863 | -0.2018 | 9.762e-03 |
| ASRB | 233 | 0.64 | 0.1615 | 9.567e-02 |
| CHAP | 261 | 0.6138 | 0.2094 | 2.066e-02 |
| CHS | 544 | 0.6286 | 0.0134 | 8.304e-01 |
| DNGS | 483 | 0.7188 | -0.0057 | 9.361e-01 |
| ENIGMA | 11113 | 0.6261 | 0.0474 | 6.230e-04 |
| EPOZ | 251 | 0.6251 | -0.0998 | 2.793e-01 |
| FHS | 4710 | 0.6032 | 0.0446 | 3.415e-02 |
| GenR | 265 | 0.6729 | -0.0478 | 6.059e-01 |
| HUNT | 851 | 0.6586 | 0.0947 | 6.398e-02 |
| LIFE | 1825 | 0.6108 | -0.0449 | 1.857e-01 |
| LLS | 355 | 0.6089 | -0.1391 | 7.047e-02 |
| MethCT. | 114 | 0.8121 | 0.1767 | 2.972e-01 |
| NOMAS | 141 | 0.5675 | 0.0502 | 6.763e-01 |
| PoznanMS | 204 | 0.5809 | 0.19 | 5.817e-02 |
| PROSPER | 315 | 0.6298 | 0.0688 | 4.042e-01 |
| RS1 | 1103 | 0.6243 | 0.0592 | 1.784e-01 |
| RS2 | 1159 | 0.6393 | 0.0522 | 2.279e-01 |
| RS3 | 2582 | 0.6359 | 0.0632 | 2.893e-02 |
| RUSH1 | 184 | 0.6751 | -0.0433 | 6.977e-01 |
| RUSH2 | 106 | 0.5972 | 0.1511 | 2.834e-01 |
| SHIP | 966 | 0.5946 | 0.0915 | 4.871e-02 |
| SHIP-TREND | 858 | 0.6063 | 0.0284 | 5.651e-01 |
| SYS.adolescents | 980 | 0.6058 | 0.0586 | 2.051e-01 |
| SYS.adults | 594 | 0.6002 | -0.0017 | 9.771e-01 |
| TREND | 1084 | 0.6199 | 0.012 | 7.858e-01 |
| UKB | 14571 | 0.6463 | 0.0338 | 5.836e-03 |
| WHICAP | 83 | 0.6109 | 0.092 | 5.650e-01 |
| **Non-European** | | | | |
| ARIC.AA | 675 | 0.8884 | 0.0516 | 5.507e-01 |
| CARDIA.AA | 132 | 0.874 | 0.02 | 9.140e-01 |
| CHAP.black | 321 | 0.8927 | 0.1112 | 3.837e-01 |
| FBIRN | 275 | 0.7304 | 0.0779 | 4.179e-01 |
| NOMAS.black | 168 | 0.8532 | -0.269 | 8.289e-02 |
| NOMAS.hispanic | 718 | 0.742 | -0.0174 | 7.732e-01 |
| SCES | 209 | 0.8375 | 0.3067 | 2.074e-02 |
| SIMES | 201 | 0.8539 | -0.1299 | 3.432e-01 |
| WHICAP.black | 60 | 0.8954 | 0.4033 | 1.822e-01 |
| **Summary European MA** | **72085** | **0.6308** | **0.032** | **2.754e-09** |
| **Summary African MA** | **1356** | **0.884** | **0.033** | **0.5832** |
| **Summary Asian MA** | **1298** | **0.8395** | **0.025** | **0.6358** |
| **Summary Transancestral MA** | **75846** | **0.6406** | **0.032** | **2.062e-09** |

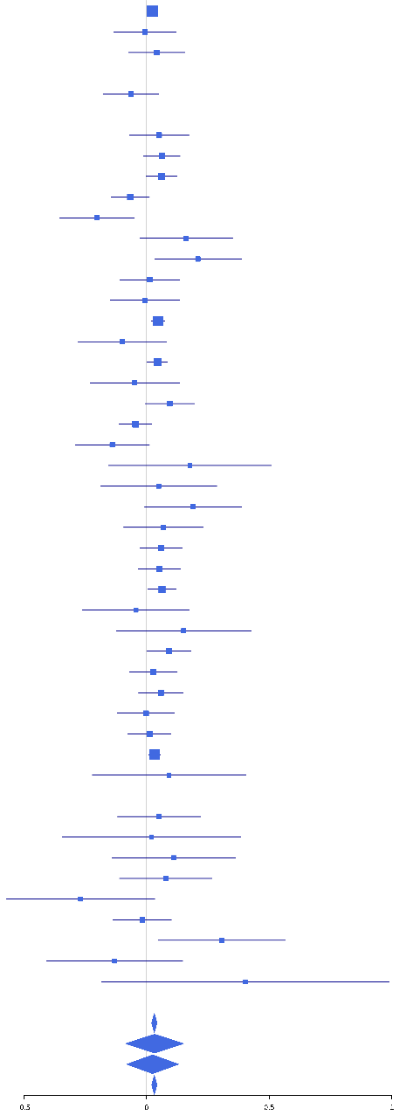

**rs148340480 (C); chr19:13159859; model 1 (I2=3.2, HetP=0.4136)**

| Study | N_total | EAF | Beta | P |
|---|---|---|---|---|
| **Head circumference** | | | | |
| Haworth | 18881 | 0.9165 | 0.0512 | 5.890e-03 |
| HKOS | 888 | 0.9476 | -0.0806 | 4.490e-01 |
| VIDI | 737 | 0.964 | 0.2789 | 4.653e-02 |
| **CT-based** | | | | |
| ESS | 641 | 0.9289 | 0.1734 | 1.105e-01 |
| UCL | 642 | 0.9271 | -0.0061 | 9.547e-01 |
| **MR-based** | | | | |
| 3C-Bordeaux | 564 | 0.9096 | 0.1396 | 1.796e-01 |
| 3c-Dijon | 1499 | 0.9293 | 0.0708 | 3.205e-01 |
| AGES | 2226 | 0.9448 | 0.053 | 4.196e-01 |
| ARIC.EA | 1390 | 0.9298 | 0.0908 | 2.213e-01 |
| ASPS-Family | 338 | 0.9141 | -0.0228 | 8.681e-01 |
| CHAP | 261 | 0.9041 | 0.2609 | 8.047e-02 |
| CHS | 544 | 0.9225 | 0.2365 | 3.709e-02 |
| ENIGMA | 7980 | 0.9232 | 0.0373 | 2.095e-01 |
| EPOZ | 251 | 0.9282 | -0.1011 | 5.585e-01 |
| GenR | 265 | 0.9138 | -0.0816 | 5.983e-01 |
| HUNT | 851 | 0.9568 | -0.0116 | 9.225e-01 |
| LBC1936 | 577 | 0.9516 | 0.0494 | 7.186e-01 |
| LIFE | 1825 | 0.9275 | -0.0788 | 2.175e-01 |
| LLS | 355 | 0.9537 | 0.1342 | 4.527e-01 |
| NOMAS | 141 | 0.931 | 0.5561 | 1.938e-02 |
| PoznanMS | 204 | 0.9273 | 0.1437 | 4.510e-01 |
| PROSPER | 315 | 0.9463 | 0.1155 | 5.134e-01 |
| RS1 | 1103 | 0.9268 | 0.1738 | 3.352e-02 |
| RS2 | 1159 | 0.9235 | 0.0014 | 9.857e-01 |
| RS3 | 2582 | 0.9305 | 0.0236 | 6.662e-01 |
| RUSH2 | 106 | 0.9177 | 0.2107 | 4.015e-01 |
| SHIP | 966 | 0.9373 | 0.1395 | 1.377e-01 |
| SHIP-TREND | 858 | 0.9352 | 0.0948 | 3.339e-01 |
| SYS.adolescents | 980 | 0.9101 | 0.1742 | 2.736e-02 |
| SYS.adults | 594 | 0.9038 | 0.1057 | 2.828e-01 |
| TREND | 1084 | 0.934 | 0.0879 | 3.099e-01 |
| UKB | 14571 | 0.9344 | 0.0847 | 3.435e-04 |
| WHICAP | 83 | 0.9385 | -0.1641 | 6.131e-01 |
| **Non-European** | | | | |
| ARIC.AA | 675 | 0.9899 | 0.5613 | 3.917e-02 |
| CHAP.black | 321 | 0.9846 | -0.4 | 2.129e-01 |
| NOMAS.hispanic | 718 | 0.9481 | 0.0546 | 6.462e-01 |
| **Summary European MA** | **64573** | **0.9268** | **0.065** | **1.252e-09** |
| **Summary African MA** | **996** | **0.9882** | **0.206** | **0.3218** |
| **Summary Asian MA** | **888** | **0.9476** | **-0.081** | **0.449** |
| **Summary Transancestral MA** | **67175** | **0.9282** | **0.064** | **1.582e-09** |

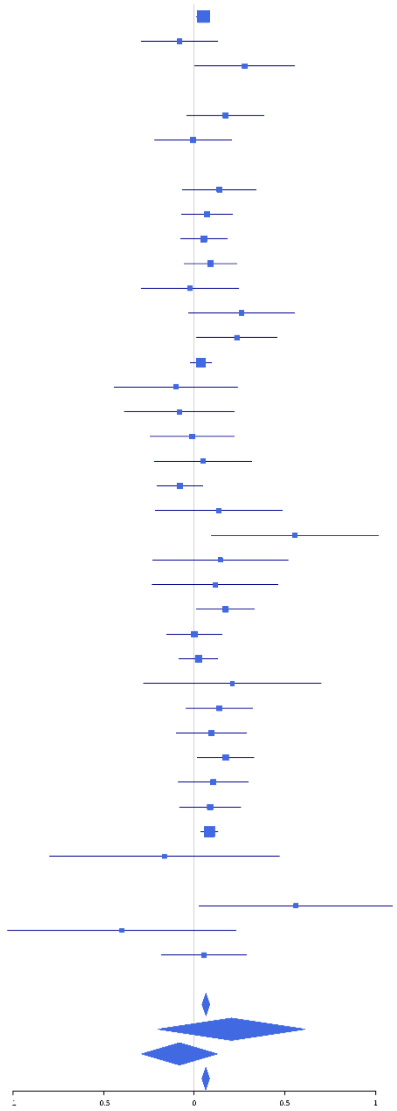

**rs6124328 (A); chr20:39826928; model 1 (I2=0, HetP=0.8734)**

| Study | N_total | EAF | Beta | P |
|---|---|---|---|---|
| **Head circumference** | | | | |
| Haworth | 18881 | 0.6927 | 0.0406 | 2.663e-04 |
| HKOS | 888 | 0.1909 | 0.007 | 9.077e-01 |
| VIDI | 754 | 0.6912 | 2e-04 | 9.979e-01 |
| **CT-based** | | | | |
| ESS | 641 | 0.672 | 0.0479 | 4.211e-01 |
| UCL | 642 | 0.6567 | 0.0487 | 4.072e-01 |
| **MR-based** | | | | |
| 3C-Bordeaux | 564 | 0.6883 | -0.0398 | 5.364e-01 |
| 3c-Dijon | 1499 | 0.6936 | 0.0481 | 2.250e-01 |
| AGES | 2226 | 0.7347 | 0.0252 | 4.582e-01 |
| ARIC.EA | 1390 | 0.6916 | 0.0821 | 4.559e-02 |
| ASPS-Family | 338 | 0.7132 | 0.0646 | 4.474e-01 |
| ASRB | 233 | 0.6742 | 0.1122 | 2.574e-01 |
| CARDIA.EA | 349 | 0.6879 | 0.0457 | 5.757e-01 |
| CHAP | 261 | 0.7092 | 0.0303 | 7.539e-01 |
| CHS | 544 | 0.6845 | 0.1427 | 2.866e-02 |
| DHS | 460 | 0.7038 | 0.0036 | 9.605e-01 |
| DNGS | 516 | 0.5101 | -0.04 | 5.214e-01 |
| ENIGMA | 11373 | 0.6915 | 0.0354 | 1.369e-02 |
| EPOZ | 251 | 0.6596 | -0.0296 | 7.535e-01 |
| ERF | 122 | 0.6953 | -0.0994 | 4.743e-01 |
| FHS | 4710 | 0.6999 | 0.0494 | 2.810e-02 |
| GenR | 265 | 0.6125 | 0.2069 | 2.030e-02 |
| HUNT | 851 | 0.7206 | 0.0898 | 9.622e-02 |
| LBC1936 | 577 | 0.7125 | 0.0321 | 6.211e-01 |
| LIFE | 1825 | 0.6798 | 0.0261 | 4.625e-01 |
| LLS | 355 | 0.6727 | 0.0342 | 6.695e-01 |
| MethCT. | 114 | 0.4525 | -0.1412 | 2.888e-01 |
| NOMAS | 141 | 0.6762 | 0.0979 | 4.433e-01 |
| PoznanMS | 204 | 0.6797 | 0.156 | 1.415e-01 |
| PROSPER | 315 | 0.672 | -0.06 | 4.796e-01 |
| RS1 | 1103 | 0.6785 | -0.005 | 9.130e-01 |
| RS2 | 1159 | 0.6919 | -0.0206 | 6.464e-01 |
| RS3 | 2582 | 0.6934 | -0.0128 | 6.725e-01 |
| RUSH1 | 184 | 0.6759 | 0.0571 | 6.088e-01 |
| RUSH2 | 106 | 0.7448 | 0.2284 | 1.502e-01 |
| SHIP | 966 | 0.6995 | 0.0592 | 2.332e-01 |
| SHIP-TREND | 858 | 0.6783 | 0.0054 | 9.164e-01 |
| SYS.adolescents | 980 | 0.7133 | 0.0134 | 7.885e-01 |
| SYS.adults | 594 | 0.7246 | -0.0386 | 5.525e-01 |
| TREND | 1084 | 0.6941 | -0.0018 | 9.688e-01 |
| UKB | 14571 | 0.6908 | 0.0364 | 4.053e-03 |
| WHICAP | 83 | 0.7436 | -0.0041 | 9.821e-01 |
| **Non-European** | | | | |
| ARIC.AA | 675 | 0.3693 | 0.0125 | 8.253e-01 |
| CARDIA.AA | 132 | 0.355 | -0.0194 | 8.797e-01 |
| CHAP.black | 321 | 0.3339 | -0.0153 | 8.551e-01 |
| FBIRN | 275 | 0.57 | -0.0144 | 8.677e-01 |
| NOMAS.black | 168 | 0.3904 | -0.0265 | 8.128e-01 |
| NOMAS.hispanic | 718 | 0.515 | -0.0448 | 3.960e-01 |
| SCES | 209 | 0.2003 | 0.0229 | 8.514e-01 |
| SIMES | 201 | 0.298 | -0.1625 | 1.368e-01 |
| WHICAP.black | 60 | 0.3166 | 0.3438 | 8.555e-02 |
| **Summary European MA** | **74557** | **0.692** | **0.035** | **6.788e-10** |
| **Summary African MA** | **1356** | **0.3598** | **0.012** | **0.7657** |
| **Summary Asian MA** | **1298** | **0.209** | **-0.02** | **0.6786** |
| **Summary Transancestral MA** | **78318** | **0.6758** | **0.032** | **4.223e-09** |

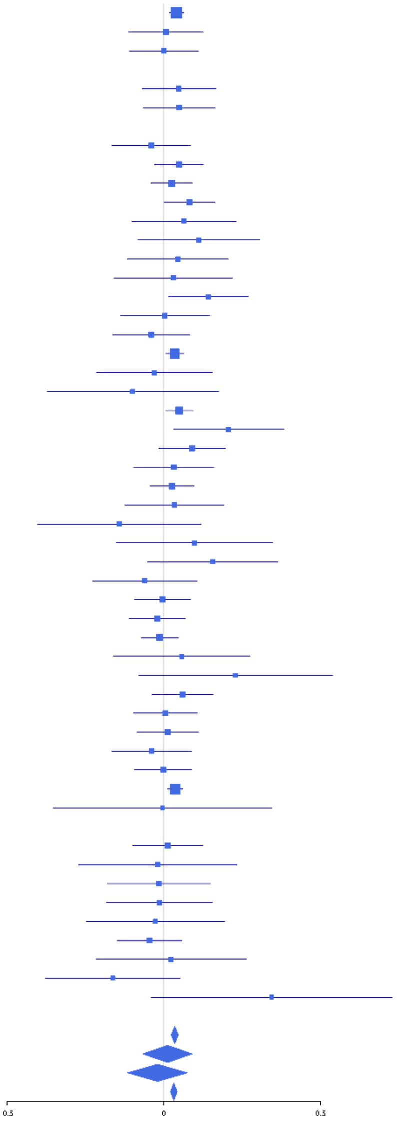

**rs142047625 (T); chr21:38909916; model 1 (I2=56.1, HetP=6.487e-05)**

| Study | N_total | EAF | Beta | P |
|---|---|---|---|---|
| **Head circumference** | | | | |
| Haworth | 17826 | 0.021 | 0.0676 | 6.740e-02 |
| VIDI | 752 | 0.0101 | -0.1338 | 6.049e-01 |
| **CT-based** | | | | |
| ESS | 641 | 0.0227 | -0.1741 | 3.536e-01 |
| UCL | 642 | 0.0259 | -0.3298 | 6.066e-02 |
| **MR-based** | | | | |
| 3C-Bordeaux | 564 | 0.0209 | 0.1222 | 5.576e-01 |
| 3c-Dijon | 1499 | 0.0203 | -0.3446 | 7.827e-03 |
| AGES | 2226 | 0.0134 | -0.1133 | 3.841e-01 |
| ARIC.EA | 1390 | 0.0194 | -0.4675 | 6.849e-04 |
| CHAP | 261 | 0.0114 | -0.55 | 1.834e-01 |
| CHS | 544 | 0.0227 | -0.219 | 2.814e-01 |
| ENIGMA | 11373 | 0.0186 | -0.1454 | 3.049e-03 |
| EPOZ | 251 | 0.0251 | -0.1276 | 6.549e-01 |
| ERF | 122 | 0.0175 | -0.5334 | 2.745e-01 |
| FHS | 4710 | 0.0211 | -0.1559 | 2.979e-02 |
| HUNT | 851 | 0.0117 | -0.1032 | 6.467e-01 |
| LBC1936 | 577 | 0.0135 | 0.353 | 1.663e-01 |
| LIFE | 1825 | 0.0174 | -0.1785 | 1.586e-01 |
| LLS | 355 | 0.0239 | -0.5413 | 2.755e-02 |
| NOMAS | 141 | 0.0269 | 0.0585 | 8.735e-01 |
| PROSPER | 315 | 0.021 | -0.2954 | 2.878e-01 |
| RS1 | 1103 | 0.0254 | -0.0237 | 8.609e-01 |
| RS2 | 1159 | 0.0238 | -0.1623 | 2.339e-01 |
| RS3 | 2582 | 0.022 | -0.105 | 2.685e-01 |
| SHIP | 966 | 0.0162 | -0.2194 | 2.241e-01 |
| SHIP-TREND | 858 | 0.0164 | -0.2529 | 1.831e-01 |
| SYS.adolescents | 980 | 0.0227 | -0.1011 | 5.057e-01 |
| SYS.adults | 594 | 0.0238 | -0.3698 | 5.222e-02 |
| TREND | 1084 | 0.0183 | 0.1639 | 3.063e-01 |
| UKB | 14571 | 0.0204 | -0.171 | 3.710e-05 |
| WHICAP | 83 | 0.0368 | -0.8278 | 4.811e-02 |
| **Non-European** | | | | |
| NOMAS.hispanic | 718 | 0.01 | 0.052 | 8.445e-01 |
| SIMES | 201 | 0.0058 | 1.4549 | 1.318e-03 |
| **Summary European MA** | **70845** | **0.02** | **-0.104** | **4.387e-08** |
| **Summary Asian MA** | **201** | **0.0058** | **2.11** | **0.001318** |
| **Summary Transancestral MA** | **71764** | **0.0198** | **-0.099** | **1.523e-07** |

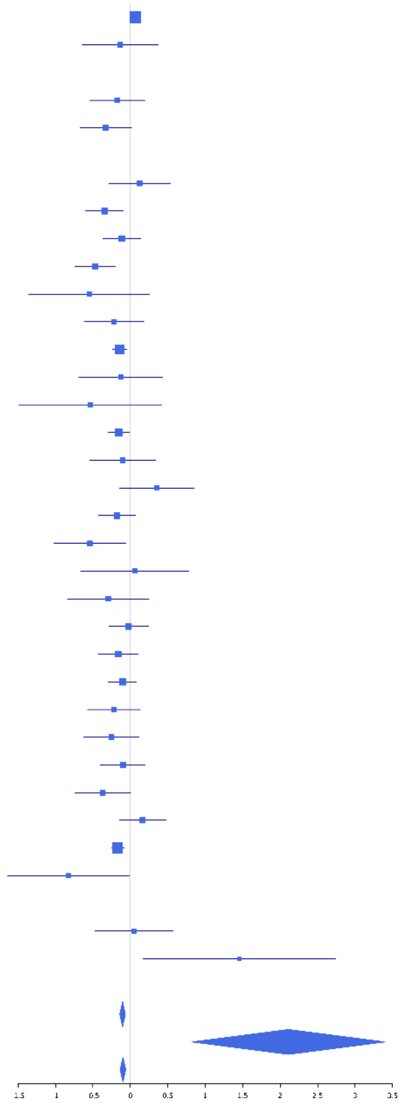

**rs2040167 (T); chr22:22077224; model 1 (I2=0, HetP=0.4866)**

| Study | N_total | EAF | Beta | P |
|---|---|---|---|---|
| **Head circumference** | | | | |
| Haworth | 18881 | 0.0442 | 0.0642 | 1.041e-02 |
| HKOS | 888 | 0.1357 | 0.0272 | 6.944e-01 |
| VIDI | 741 | 0.0402 | 0.1404 | 2.887e-01 |
| **CT-based** | | | | |
| UCL | 642 | 0.0431 | 0.1602 | 2.437e-01 |
| **MR-based** | | | | |
| 3C-Bordeaux | 564 | 0.0318 | 0.2156 | 2.043e-01 |
| 3c-Dijon | 1499 | 0.0358 | 0.0525 | 5.932e-01 |
| AGES | 2226 | 0.0419 | -0.0853 | 2.544e-01 |
| ARIC.EA | 1390 | 0.0396 | 0.1382 | 1.553e-01 |
| ASPS-Family | 338 | 0.0348 | -0.345 | 1.005e-01 |
| ASRB | 233 | 0.0418 | -0.0886 | 7.019e-01 |
| CARDIA.EA | 349 | 0.0363 | -0.1629 | 4.211e-01 |
| CHAP | 261 | 0.0287 | -0.0592 | 8.211e-01 |
| CHS | 544 | 0.0313 | 0.1686 | 3.324e-01 |
| DNGS | 512 | 0.0581 | -0.0216 | 8.714e-01 |
| ENIGMA | 11373 | 0.0423 | 0.0839 | 1.089e-02 |
| EPOZ | 251 | 0.0386 | -0.1096 | 6.359e-01 |
| ERF | 122 | 0.0347 | 0.6804 | 5.168e-02 |
| FHS | 4710 | 0.0387 | 0.0601 | 2.605e-01 |
| GenR | 265 | 0.0468 | 0.0996 | 6.284e-01 |
| HUNT | 851 | 0.0604 | 0.19 | 6.194e-02 |
| LBC1936 | 577 | 0.0452 | 0.0292 | 8.370e-01 |
| LIFE | 1825 | 0.0362 | 0.1321 | 1.363e-01 |
| LLS | 355 | 0.0409 | 0.2061 | 2.768e-01 |
| NOMAS | 141 | 0.042 | -0.0926 | 7.553e-01 |
| PoznanMS | 204 | 0.0413 | 0.1231 | 6.204e-01 |
| PROSPER | 315 | 0.0568 | -0.1017 | 5.544e-01 |
| RS1 | 1103 | 0.0432 | 0.006 | 9.542e-01 |
| RS2 | 1159 | 0.0424 | -0.0101 | 9.217e-01 |
| RS3 | 2582 | 0.042 | 0.0414 | 5.513e-01 |
| RUSH1 | 184 | 0.042 | -0.1481 | 5.693e-01 |
| RUSH2 | 106 | 0.0283 | -0.1474 | 7.224e-01 |
| SHIP | 966 | 0.0455 | 0.1834 | 9.350e-02 |
| SHIP-TREND | 858 | 0.039 | -0.0277 | 8.243e-01 |
| SYS.adolescents | 980 | 0.0551 | 0.1715 | 8.327e-02 |
| SYS.adults | 594 | 0.0545 | 0.4155 | 1.146e-03 |
| TREND | 1084 | 0.0352 | 0.0457 | 6.949e-01 |
| UKB | 14571 | 0.0429 | 0.0877 | 2.404e-03 |
| WHICAP | 83 | 0.0215 | -0.648 | 2.294e-01 |
| **Non-European** | | | | |
| ARIC.AA | 675 | 0.0109 | -0.1612 | 5.387e-01 |
| CHAP.black | 321 | 0.0108 | -0.3826 | 3.173e-01 |
| FBIRN | 275 | 0.0495 | -0.1248 | 5.263e-01 |
| NOMAS.black | 168 | 0.0124 | 0.1075 | 8.277e-01 |
| NOMAS.hispanic | 718 | 0.0326 | 0.0811 | 5.852e-01 |
| SCES | 209 | 0.1395 | 0.249 | 7.770e-02 |
| SIMES | 201 | 0.148 | -0.1358 | 3.321e-01 |
| WHICAP.black | 60 | 0.0165 | -0.5618 | 4.367e-01 |
| **Summary European MA** | **73439** | **0.0426** | **0.071** | **3.335e-08** |
| **Summary African MA** | **1224** | **0.0114** | **-0.202** | **0.289** |
| **Summary Asian MA** | **1298** | **0.1382** | **0.037** | **0.5149** |
| **Summary Transancestral MA** | **76954** | **0.0436** | **0.067** | **8.278e-08** |

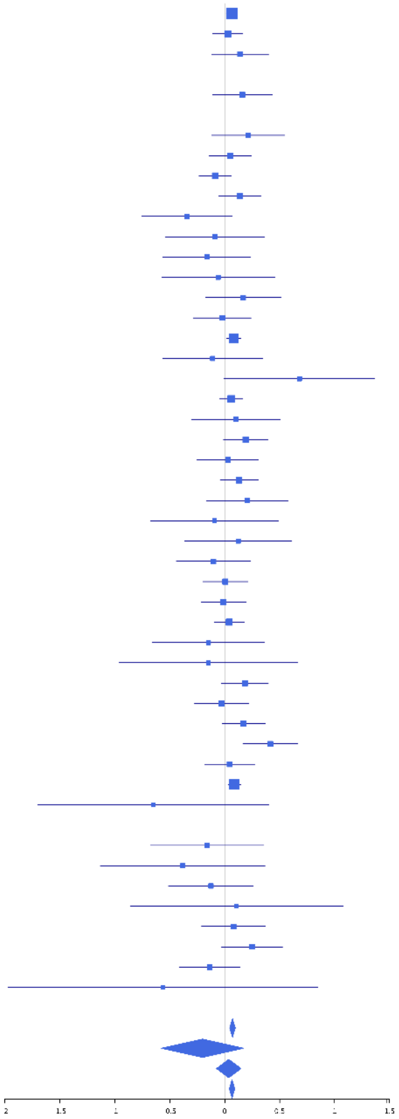

**rs5756181 (T); chr22:22305008; model 1 (I2=0, HetP=0.4929)**

| Study | N_total | EAF | Beta | P |
|---|---|---|---|---|
| **Head circumference** | | | | |
| Haworth | 18881 | 0.7219 | -0.03 | 8.758e-03 |
| HKOS | 888 | 0.7252 | -0.0525 | 3.236e-01 |
| VIDI | 738 | 0.7654 | -5e-04 | 9.935e-01 |
| **CT-based** | | | | |
| ESS | 641 | 0.7468 | 0.0819 | 2.024e-01 |
| UCL | 642 | 0.7179 | -0.1237 | 4.612e-02 |
| **MR-based** | | | | |
| 3C-Bordeaux | 564 | 0.7372 | 0.0988 | 1.445e-01 |
| 3c-Dijon | 1499 | 0.732 | -0.0051 | 9.024e-01 |
| AGES | 2226 | 0.7359 | -0.0045 | 8.939e-01 |
| ARIC.EA | 1390 | 0.737 | -0.0583 | 1.762e-01 |
| ASPS-Family | 338 | 0.7113 | 0.0587 | 4.893e-01 |
| ASRB | 233 | 0.7343 | 0.048 | 6.476e-01 |
| CARDIA.EA | 349 | 0.7374 | 0.0165 | 8.480e-01 |
| CHAP | 261 | 0.7216 | -0.0062 | 9.500e-01 |
| CHS | 544 | 0.7244 | -0.0458 | 4.994e-01 |
| DNGS | 510 | 0.7485 | -0.0317 | 6.606e-01 |
| ENIGMA | 11373 | 0.7345 | -0.0435 | 3.762e-03 |
| EPOZ | 251 | 0.7597 | -0.0758 | 4.679e-01 |
| ERF | 122 | 0.7917 | -0.3386 | 3.171e-02 |
| FHS | 4710 | 0.7228 | -0.0396 | 8.523e-02 |
| GenR | 265 | 0.7443 | -0.0256 | 7.972e-01 |
| HUNT | 851 | 0.7326 | -0.1599 | 3.501e-03 |
| LBC1936 | 577 | 0.7429 | -0.1185 | 7.835e-02 |
| LIFE | 1825 | 0.7223 | -0.0194 | 5.989e-01 |
| LLS | 355 | 0.7304 | -0.0842 | 3.191e-01 |
| MethCT. | 114 | 0.7509 | 5e-04 | 9.977e-01 |
| NOMAS | 141 | 0.7279 | 0.0072 | 9.573e-01 |
| PoznanMS | 204 | 0.6942 | -0.1908 | 7.575e-02 |
| PROSPER | 315 | 0.7222 | 0.0812 | 3.614e-01 |
| RS1 | 1103 | 0.7434 | -0.0243 | 6.178e-01 |
| RS2 | 1159 | 0.738 | 0.0475 | 3.148e-01 |
| RS3 | 2582 | 0.7503 | -0.0176 | 5.842e-01 |
| RUSH1 | 184 | 0.7617 | -0.0015 | 9.906e-01 |
| RUSH2 | 106 | 0.7402 | -0.2492 | 1.148e-01 |
| SHIP | 966 | 0.7342 | -0.0537 | 2.973e-01 |
| SHIP-TREND | 858 | 0.7279 | -0.072 | 1.849e-01 |
| SYS.adolescents | 980 | 0.8043 | -0.0465 | 4.147e-01 |
| SYS.adults | 594 | 0.7919 | -0.0988 | 1.666e-01 |
| TREND | 1084 | 0.723 | -0.0361 | 4.518e-01 |
| UKB | 14571 | 0.7369 | -0.044 | 9.395e-04 |
| WHICAP | 83 | 0.7089 | -0.0501 | 7.701e-01 |
| **Non-European** | | | | |
| ARIC.AA | 675 | 0.8639 | 0.0543 | 4.943e-01 |
| CARDIA.AA | 132 | 0.869 | -0.0792 | 6.645e-01 |
| CHAP.black | 321 | 0.8637 | 0.1958 | 8.979e-02 |
| FBIRN | 275 | 0.7679 | 0.0954 | 3.455e-01 |
| NOMAS.black | 168 | 0.8776 | -0.2086 | 2.120e-01 |
| NOMAS.hispanic | 718 | 0.7857 | -0.0601 | 3.504e-01 |
| SCES | 209 | 0.6902 | -0.0236 | 8.238e-01 |
| SIMES | 201 | 0.6486 | -0.0065 | 9.501e-01 |
| WHICAP.black | 60 | 0.8256 | 0.3361 | 1.683e-01 |
| **Summary European MA** | **74075** | **0.7329** | **-0.035** | **1.825e-09** |
| **Summary African MA** | **1356** | **0.8644** | **0.057** | **0.3064** |
| **Summary Asian MA** | **1298** | **0.7077** | **-0.04** | **0.3521** |
| **Summary Transancestral MA** | **77836** | **0.7354** | **-0.034** | **3.999e-09** |

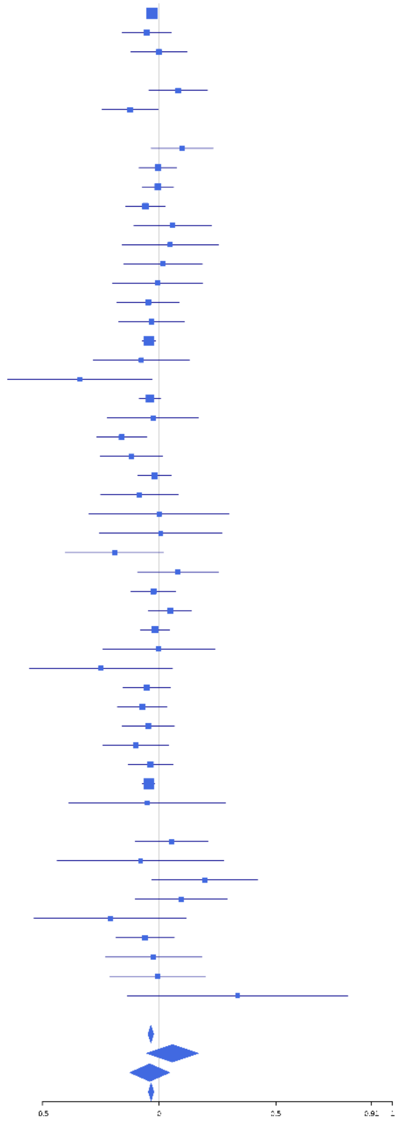

**rs10483213 (A); chr22:42339525; model 1 (I2=25, HetP=0.07449)**

| Study | N_total | EAF | Beta | P |
|---|---|---|---|---|
| **Head circumference** | | | | |
| Haworth | 18881 | 0.0985 | -0.0558 | 1.251e-03 |
| VIDI | 755 | 0.1047 | -0.0113 | 8.926e-01 |
| **CT-based** | | | | |
| ESS | 641 | 0.0902 | 0.0187 | 8.478e-01 |
| UCL | 642 | 0.0776 | -0.3155 | 2.498e-03 |
| **MR-based** | | | | |
| 3C-Bordeaux | 564 | 0.1014 | -0.0389 | 6.939e-01 |
| 3c-Dijon | 1499 | 0.1041 | -0.091 | 1.280e-01 |
| AGES | 2226 | 0.0831 | -0.0124 | 8.188e-01 |
| ARIC.EA | 1390 | 0.0936 | -0.0708 | 2.772e-01 |
| ASPS-Family | 338 | 0.1107 | -0.2868 | 1.926e-02 |
| ASRB | 233 | 0.0885 | -0.3053 | 6.256e-02 |
| CARDIA.EA | 349 | 0.0954 | -0.0752 | 5.593e-01 |
| CHAP | 261 | 0.0785 | 0.1238 | 4.476e-01 |
| CHS | 544 | 0.1094 | -0.017 | 8.612e-01 |
| DNGS | 507 | 0.0504 | -0.1085 | 4.503e-01 |
| ENIGMA | 11032 | 0.0994 | -0.0891 | 7.468e-05 |
| EPOZ | 251 | 0.1102 | 0.3283 | 2.127e-02 |
| ERF | 122 | 0.1276 | 0.1723 | 3.689e-01 |
| FHS | 4710 | 0.0993 | -0.1056 | 2.169e-03 |
| GenR | 265 | 0.0752 | -0.2336 | 1.562e-01 |
| HUNT | 851 | 0.0811 | -0.0968 | 2.756e-01 |
| LBC1936 | 577 | 0.0775 | -0.1888 | 8.642e-02 |
| LIFE | 1825 | 0.1159 | -0.0802 | 1.211e-01 |
| NOMAS | 141 | 0.0721 | -0.3649 | 1.154e-01 |
| PoznanMS | 204 | 0.1205 | 0.0707 | 6.418e-01 |
| PROSPER | 315 | 0.0905 | -0.1303 | 3.483e-01 |
| RS1 | 1103 | 0.0926 | -0.0651 | 3.758e-01 |
| RS2 | 1159 | 0.0973 | 0.0711 | 3.100e-01 |
| RS3 | 2582 | 0.1052 | -0.0355 | 4.339e-01 |
| RUSH2 | 106 | 0.1179 | -0.2202 | 3.038e-01 |
| SHIP | 966 | 0.1118 | -0.0197 | 7.849e-01 |
| SHIP-TREND | 858 | 0.1107 | -0.1546 | 4.485e-02 |
| SYS.adolescents | 980 | 0.0939 | -0.0775 | 3.169e-01 |
| SYS.adults | 594 | 0.0976 | -0.0378 | 6.989e-01 |
| TREND | 1084 | 0.1103 | -0.1685 | 1.418e-02 |
| UKB | 14571 | 0.0889 | -0.0266 | 1.960e-01 |
| WHICAP | 83 | 0.0988 | -0.2632 | 3.148e-01 |
| **Non-European** | | | | |
| ARIC.AA | 675 | 0.0171 | -0.0558 | 7.900e-01 |
| CHAP.black | 321 | 0.014 | 0.1475 | 6.613e-01 |
| FBIRN | 275 | 0.0563 | 0.2745 | 1.389e-01 |
| NOMAS.black | 168 | 0.0288 | 0.5708 | 8.196e-02 |
| NOMAS.hispanic | 718 | 0.0549 | 0.0149 | 8.970e-01 |
| WHICAP.black | 60 | 0.0167 | -0.0748 | 9.169e-01 |
| **Summary European MA** | **73209** | **0.0965** | **-0.062** | **2.192e-12** |
| **Summary African MA** | **1224** | **0.0179** | **0.099** | **0.517** |
| **Summary Transancestral MA** | **75426** | **0.0947** | **-0.059** | **1.66e-11** |

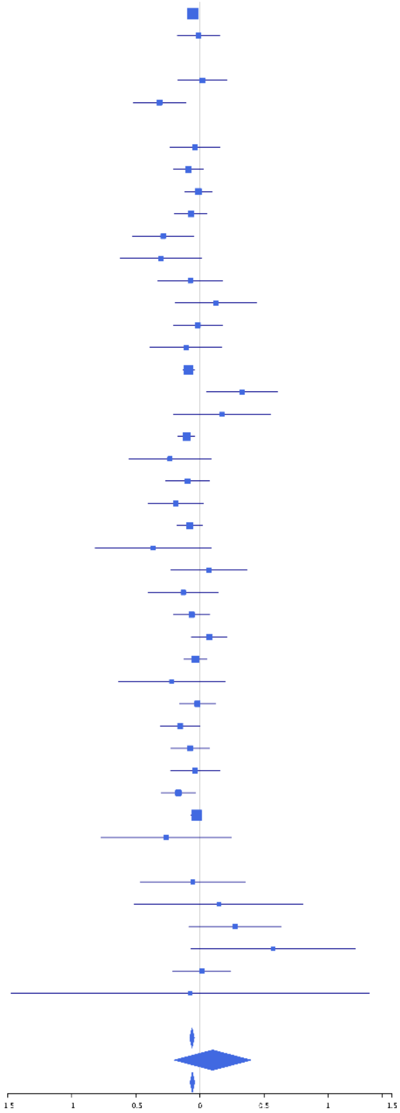
