## Supplementary Figure S2. Locuszoom plots for "Genetic variants for head size share genes and pathways with cancer"

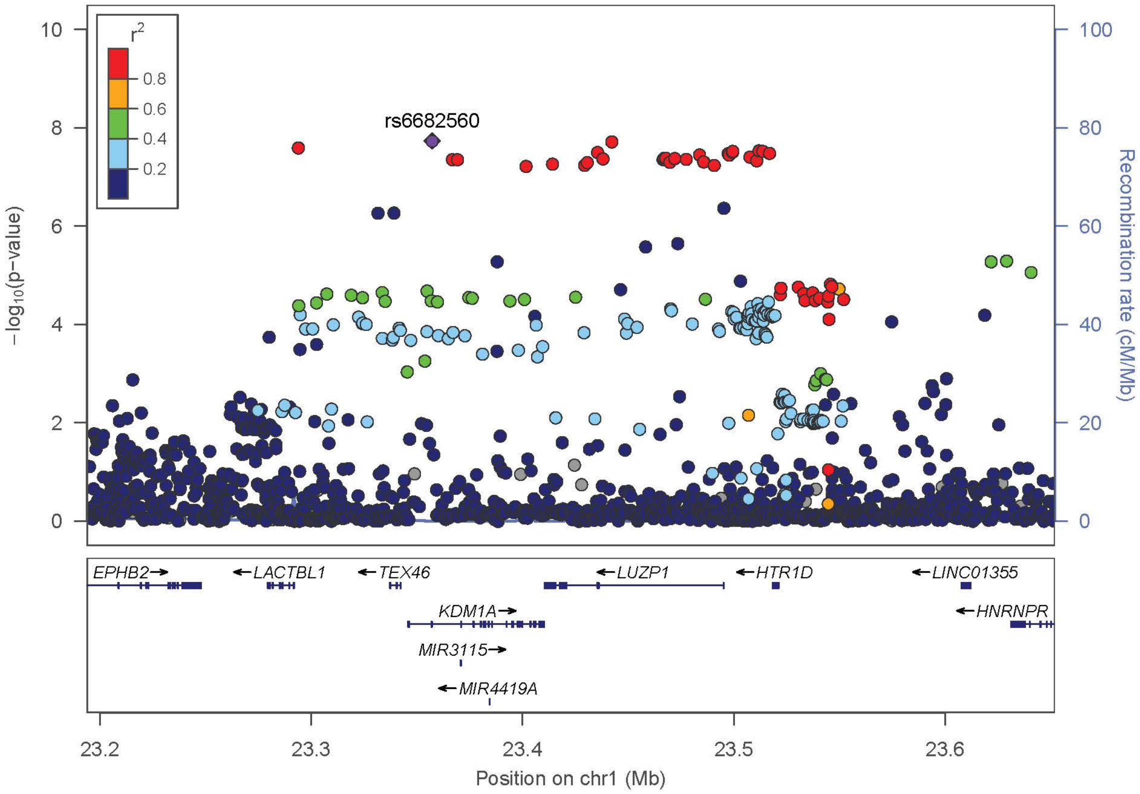
r2
0.8
0.6
0.4
0.2
rs6682560
−log10(p−value)
Recombination rate (cM/Mb)
EPHB2
LACTBL1
TEX46
LUZP1
HTR1D
LINC01355
KDM1A
HNRNPR
MIR3115
MIR4419A
Position on chr1 (Mb)

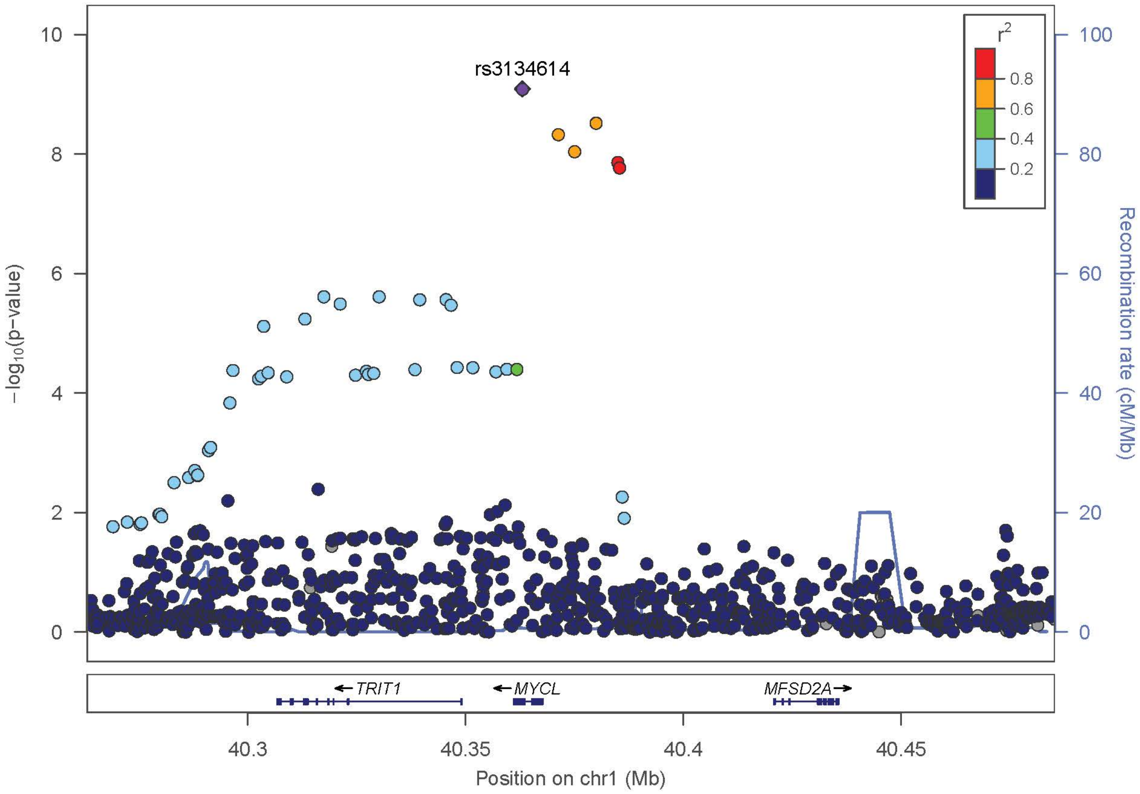
r2
0.8
0.6
0.4
0.2
rs3134614
−log10(p−value)
Recombination rate (cM/Mb)
TRIT1
MYCL
MFSD2A
Position on chr1 (Mb)

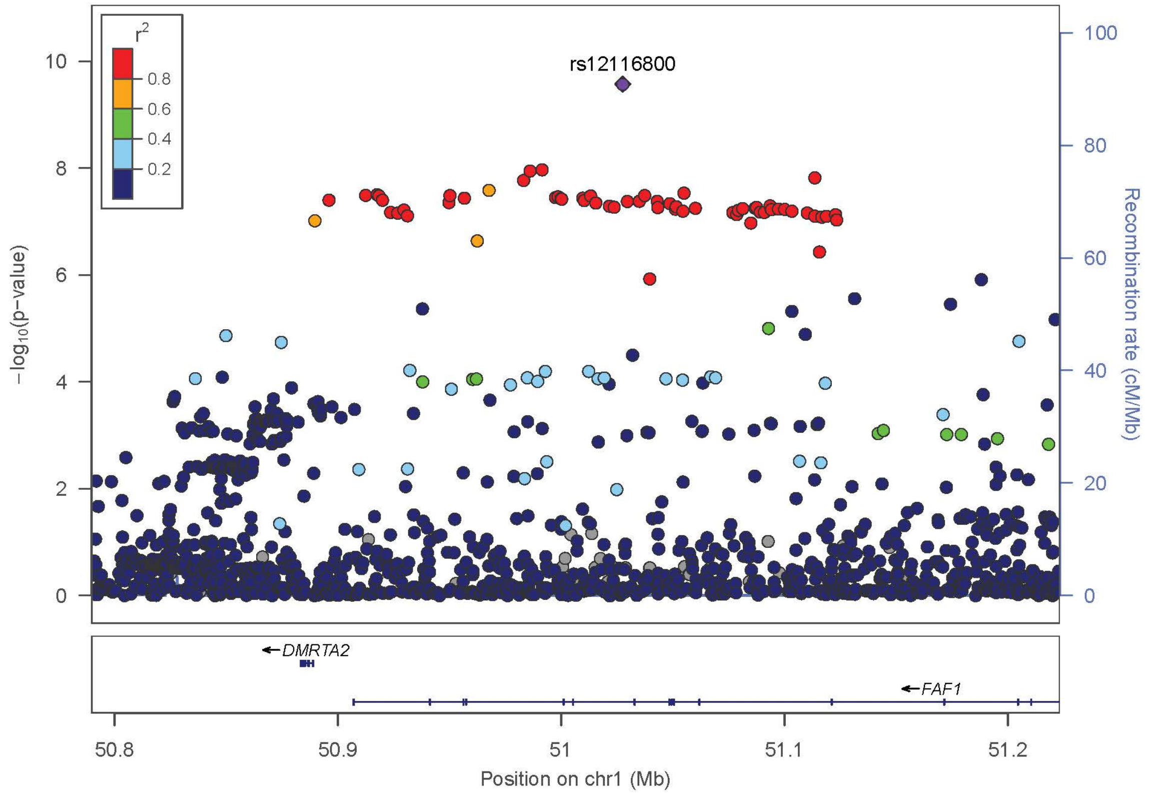

r2
0.8
0.6
0.4
0.2
rs12116800
−log10(p−value)
Recombination rate (cM/Mb)
DMRTA2
FAF1
50.8
50.9
51
51.1
51.2
Position on chr1 (Mb)


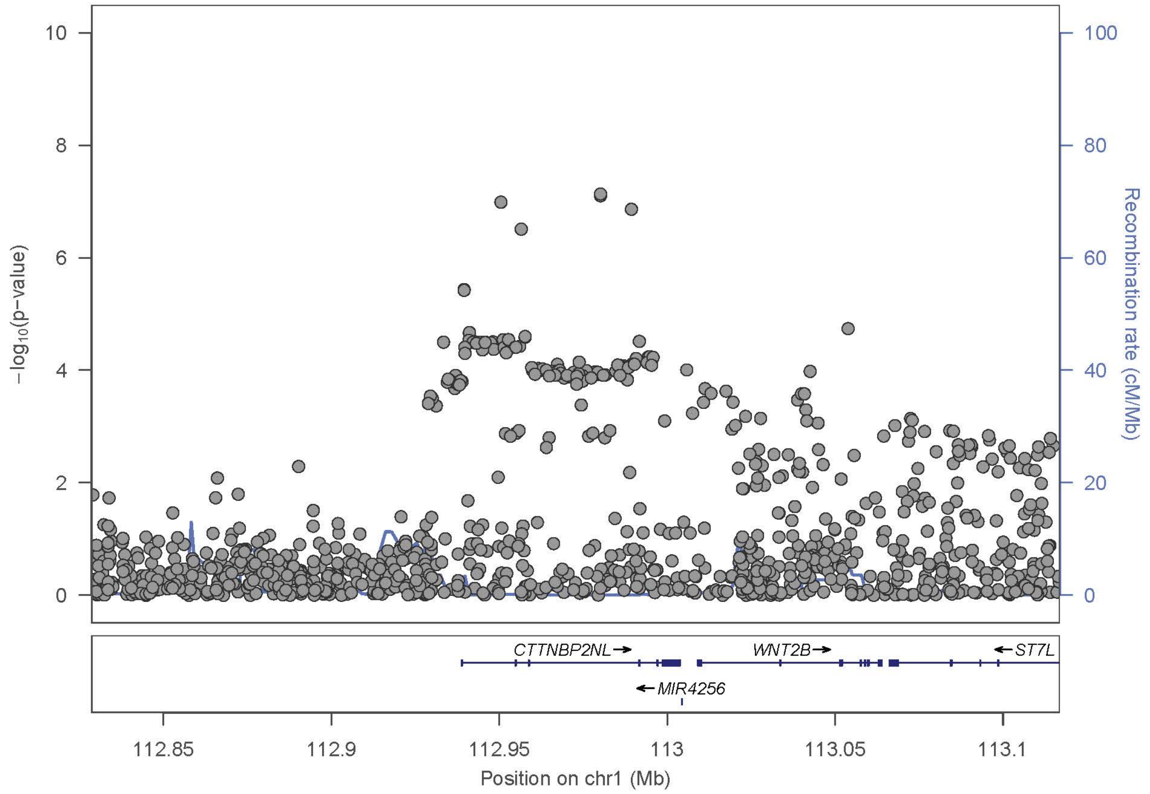

−log10(p−value)
Recombination rate (cM/Mb)
CTTNBP2NL
WNT2B
ST7L
MIR4256
112.85
112.9
112.95
113
113.05
113.1
Position on chr1 (Mb)

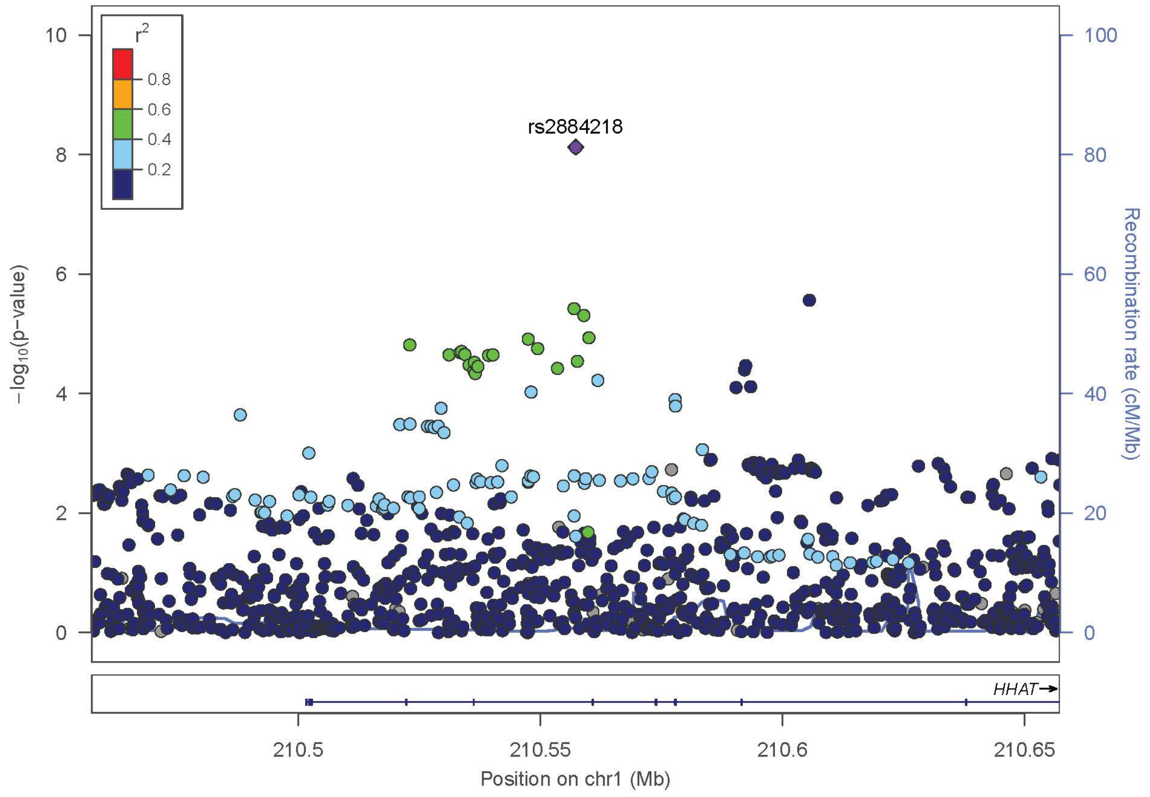

r2
0.8
0.6
0.4
0.2
rs2884218
−log10(p-value)
Recombination rate (cM/Mb)
HHAT
210.5
210.55
210.6
210.65
Position on chr1 (Mb)


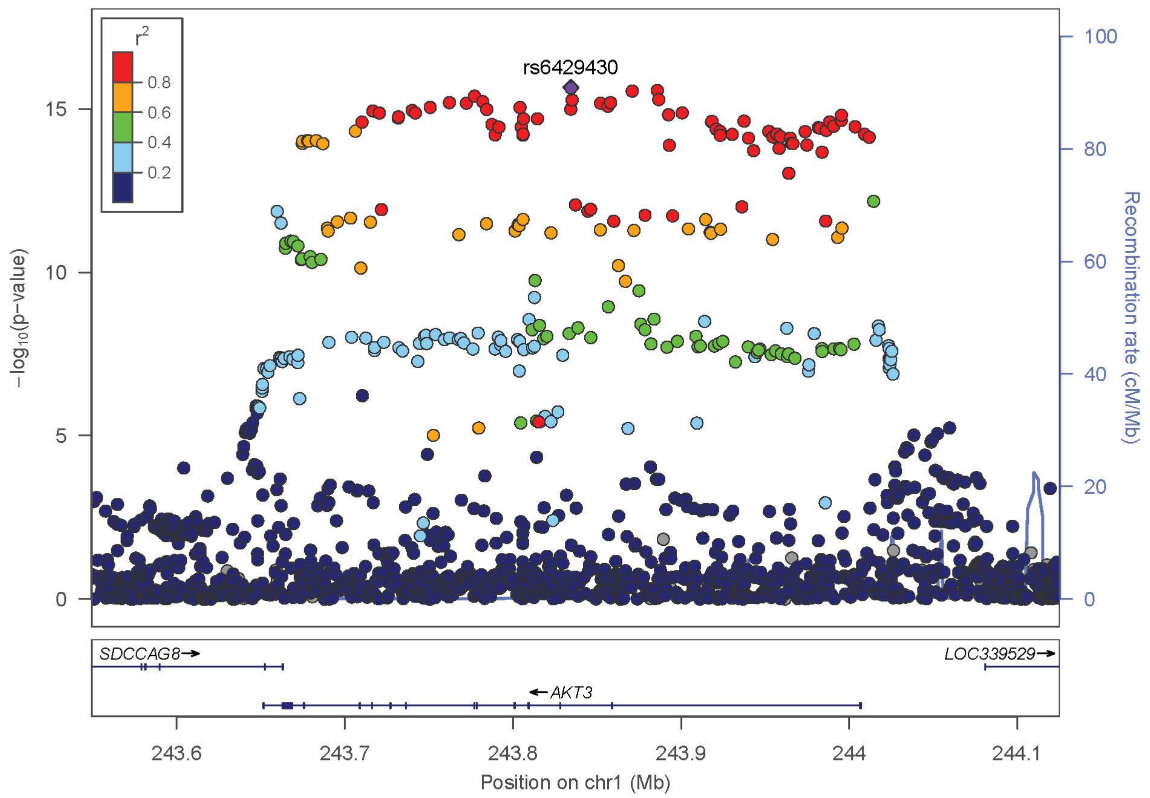

r2
0.8
0.6
0.4
0.2
rs6429430
−log10(p-value)
Recombination rate (cM/Mb)
SDCCAG8
LOC339529
AKT3
243.6
243.7
243.8
243.9
244
244.1
Position on chr1 (Mb)

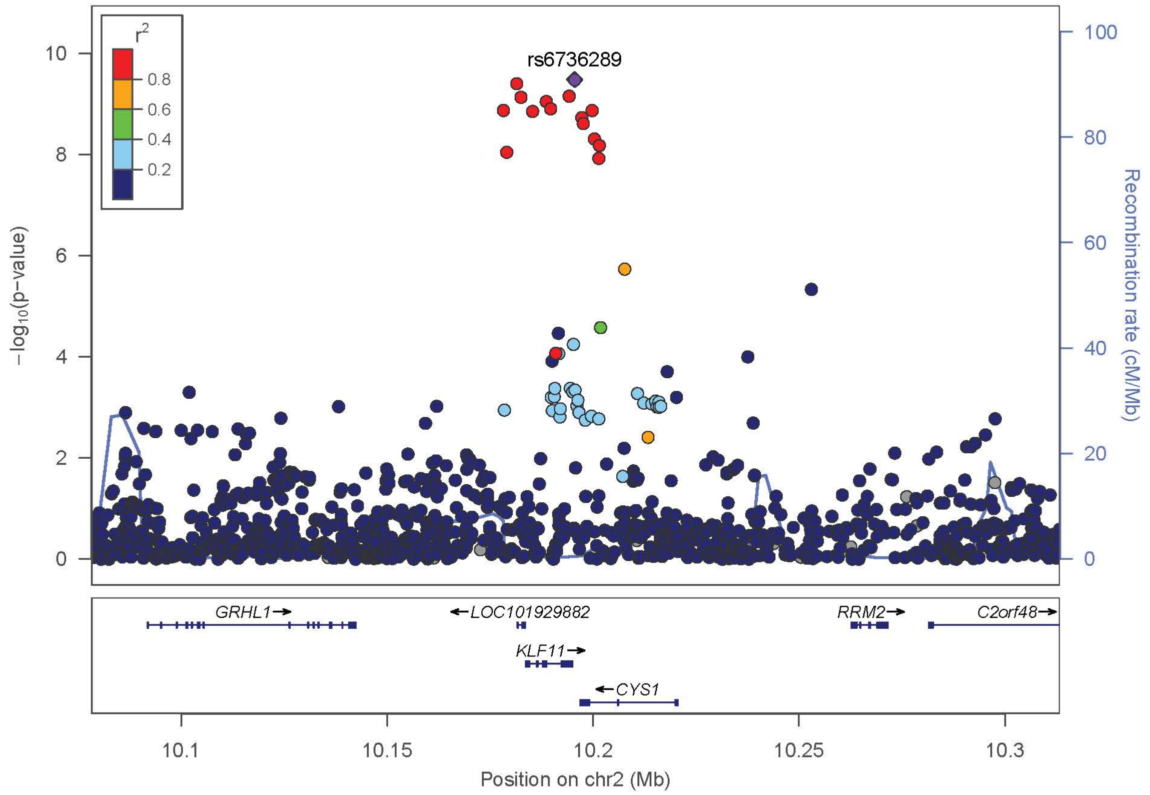
rs6736289
r2
0.8
0.6
0.4
0.2
−log10(p−value)
Recombination rate (cM/Mb)
GRHL1
LOC101929882
KLF11
CYS1
RRM2
C2orf48
10.1
10.15
10.2
10.25
10.3
Position on chr2 (Mb)

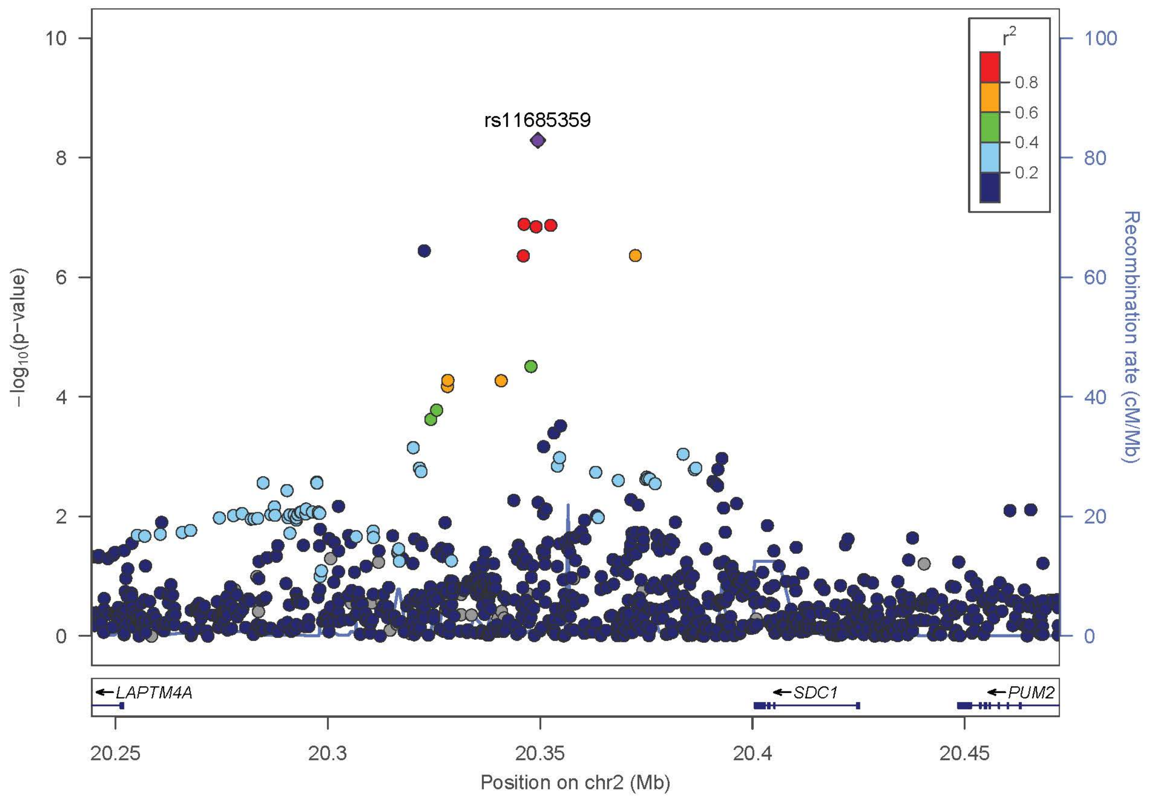
rs11685359
r2
0.8
0.6
0.4
0.2
−log10(p−value)
Recombination rate (cM/Mb)
LAPTM4A
SDC1
PUM2
20.25
20.3
20.35
20.4
20.45
Position on chr2 (Mb)

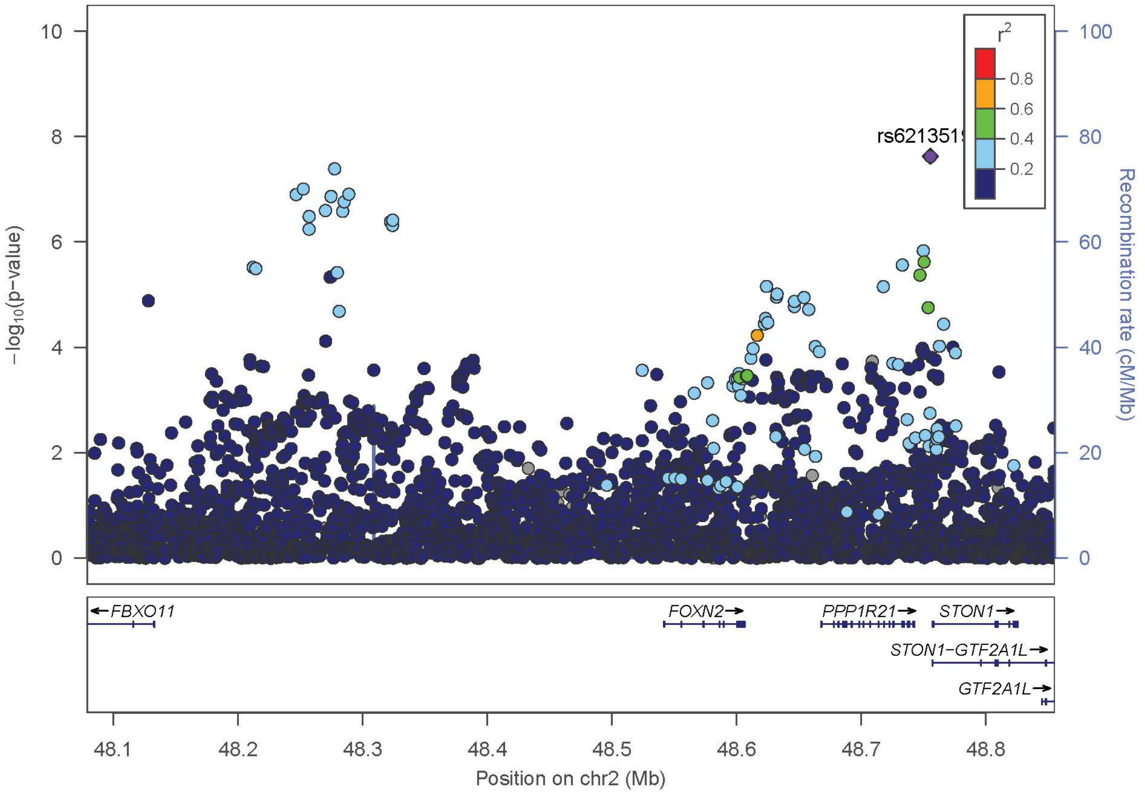
rs6213515
r2
0.8
0.6
0.4
0.2
−log10(p−value)
Recombination rate (cM/Mb)
FBXO11
FOXN2
PPP1R21
STON1
STON1−GTF2A1L
GTF2A1L
48.1
48.2
48.3
48.4
48.5
48.6
48.7
48.8
Position on chr2 (Mb)

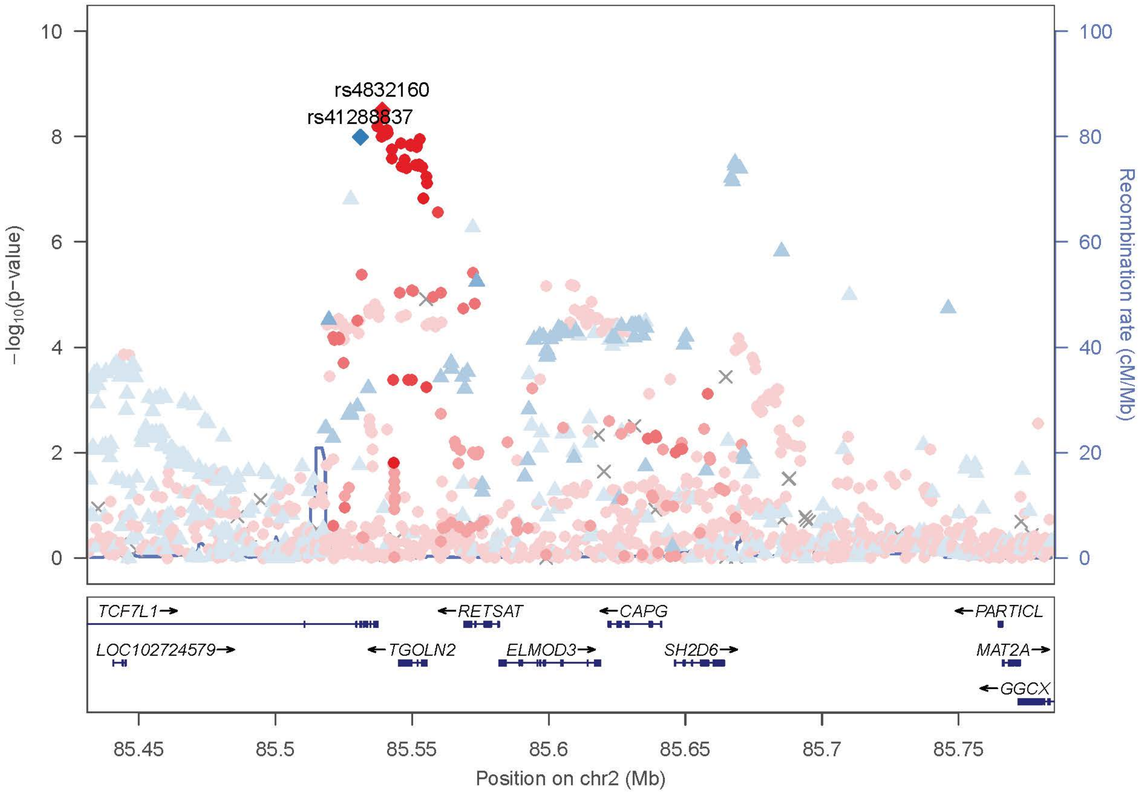
rs4832160
rs41288837
−log10(p−value)
Recombination rate (cM/Mb)
TCF7L1
LOC102724579
TGOLN2
RETSAT
ELMOD3
CAPG
SH2D6
PARTICL
MAT2A
GGCX
85.45
85.5
85.55
85.6
85.65
85.7
85.75
Position on chr2 (Mb)

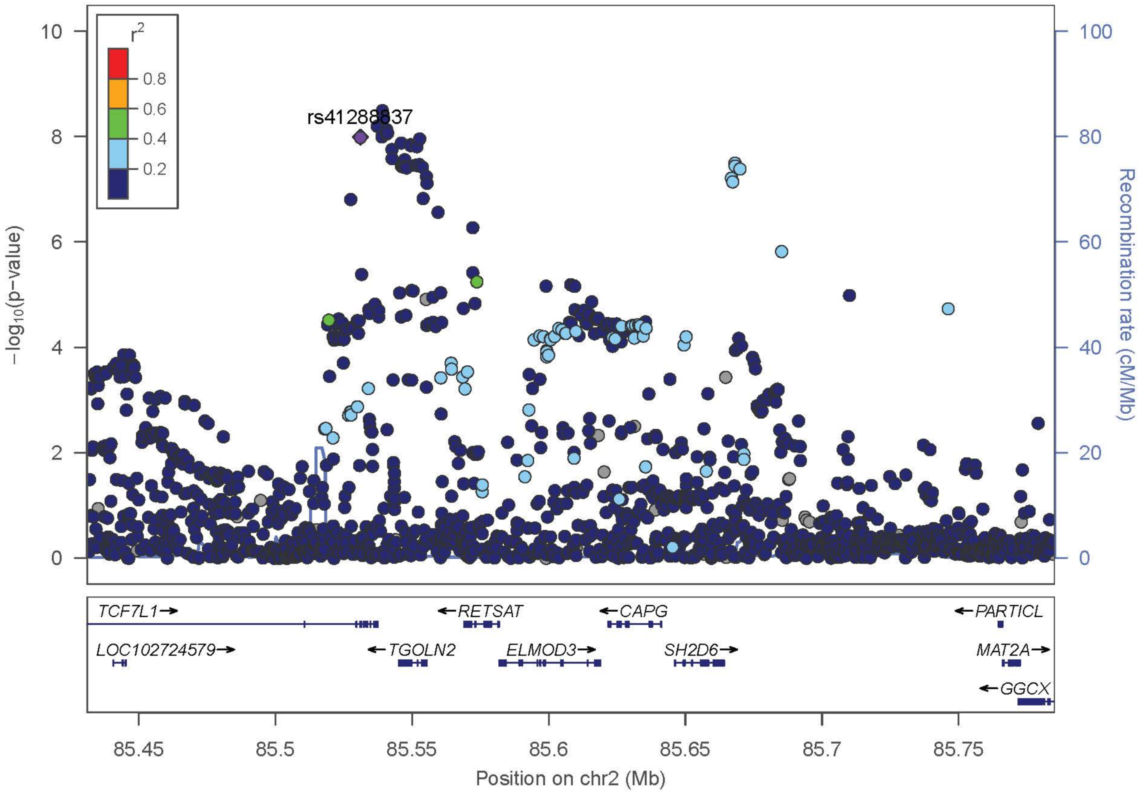
rs41288837
r2
0.8
0.6
0.4
0.2
−log10(p−value)
Recombination rate (cM/Mb)
TCF7L1
RETSAT
CAPG
PARTICL
LOC102724579
TGOLN2
ELMOD3
SH2D6
MAT2A
GGCX
85.45
85.5
85.55
85.6
85.65
85.7
85.75
Position on chr2 (Mb)

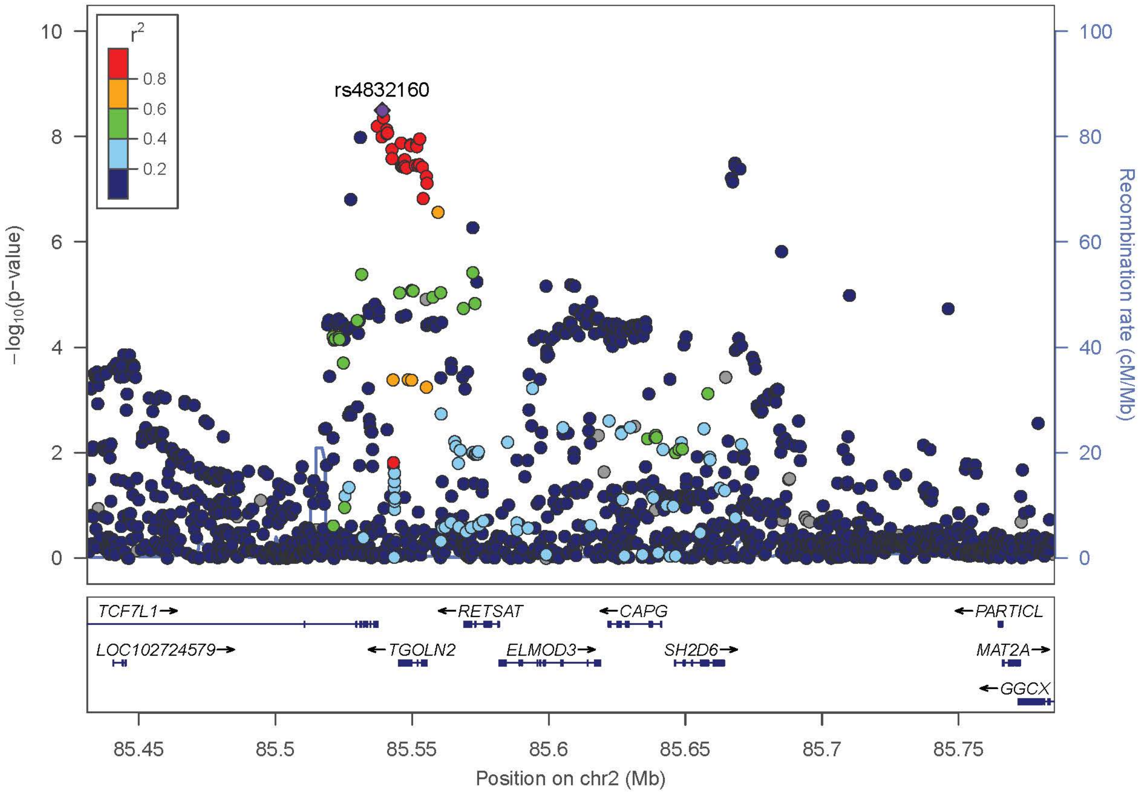
rs4832160
r2
0.8
0.6
0.4
0.2
−log10(p−value)
Recombination rate (cM/Mb)
TCF7L1
RETSAT
CAPG
PARTICL
LOC102724579
TGOLN2
ELMOD3
SH2D6
MAT2A
GGCX
85.45
85.5
85.55
85.6
85.65
85.7
85.75
Position on chr2 (Mb)

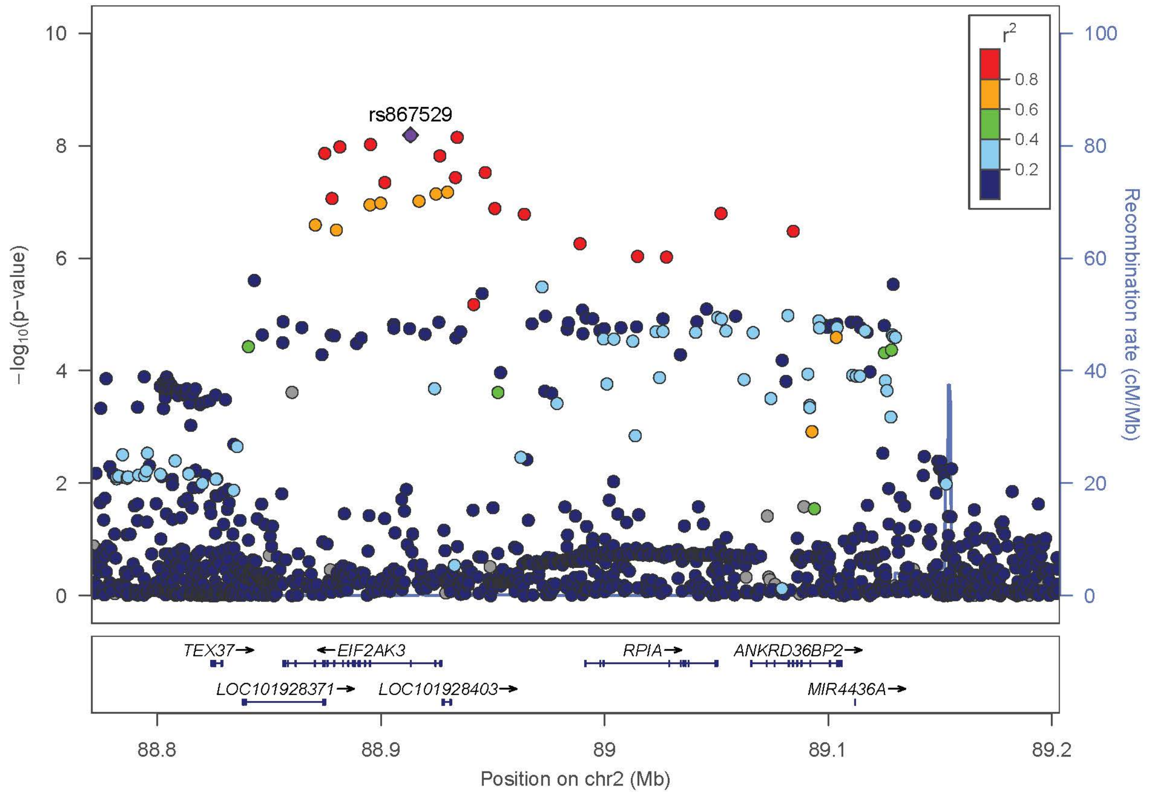
rs867529
r2
0.8
0.6
0.4
0.2
−log10(p−value)
Recombination rate (cM/Mb)
TEX37
EIF2AK3
RPIA
ANKRD36BP2
LOC101928371
LOC101928403
MIR4436A
88.8
88.9
89
89.1
89.2
Position on chr2 (Mb)

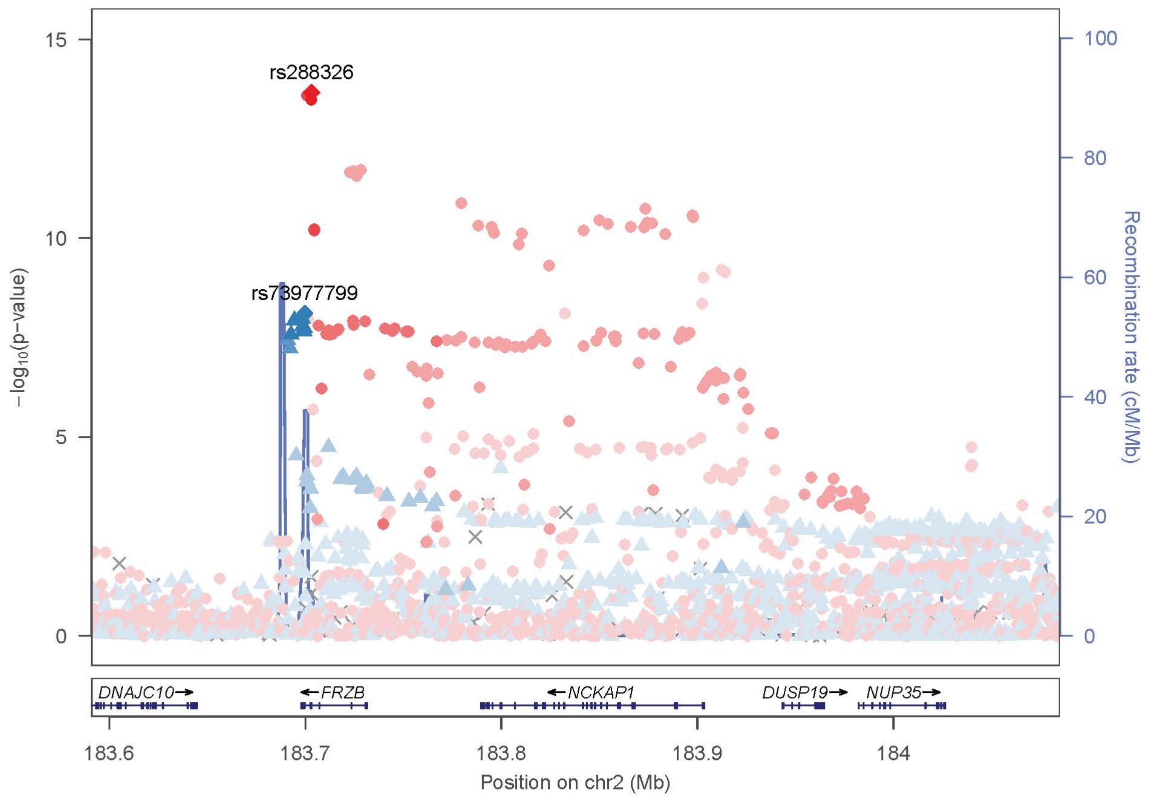
rs288326
rs73977799
−log10(p−value)
Recombination rate (cM/Mb)
DNAJC10
FRZB
NCKAP1
DUSP19
NUP35
183.6
183.7
183.8
183.9
184
Position on chr2 (Mb)

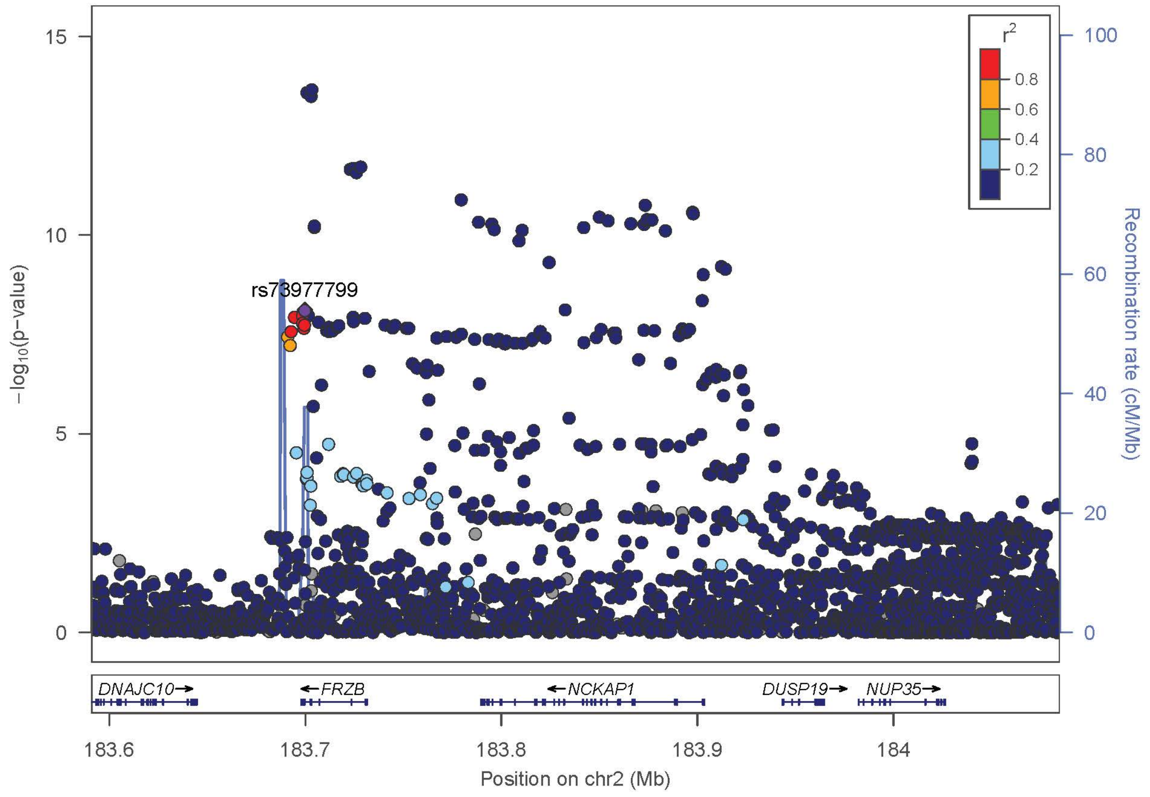
rs73977799
−log10(p−value)
Recombination rate (cM/Mb)
r2
0.8
0.6
0.4
0.2
DNAJC10
FRZB
NCKAP1
DUSP19
NUP35
183.6
183.7
183.8
183.9
184
Position on chr2 (Mb)

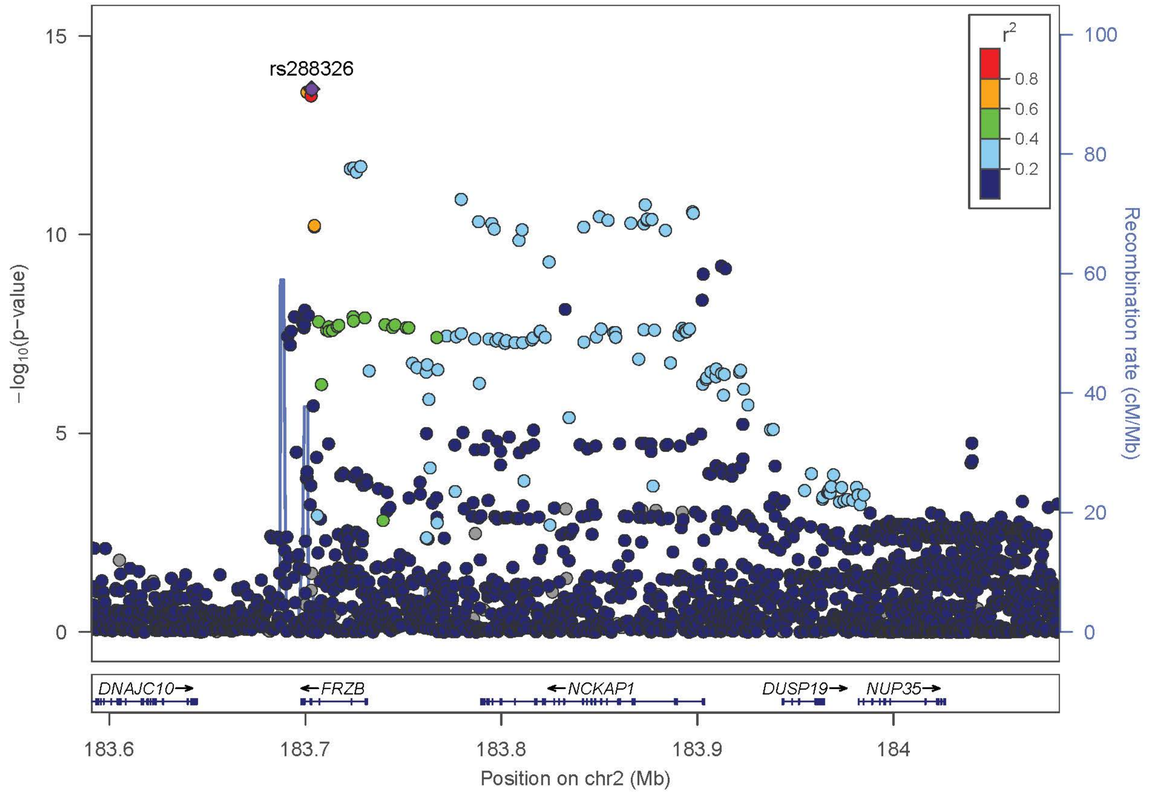
rs288326
−log10(p−value)
Recombination rate (cM/Mb)
r2
0.8
0.6
0.4
0.2
DNAJC10
FRZB
NCKAP1
DUSP19
NUP35
183.6
183.7
183.8
183.9
184
Position on chr2 (Mb)

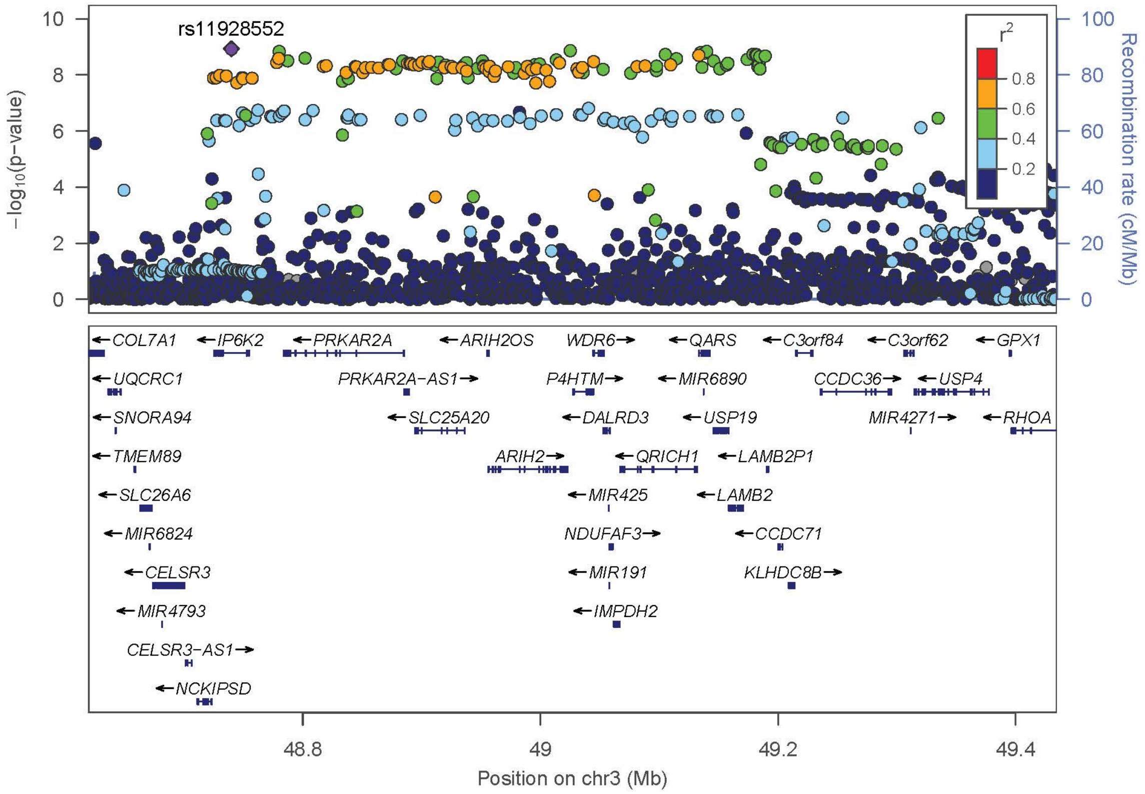
rs11928552
−log10(p−value)
Recombination rate (cM/Mb)
r2
0.8
0.6
0.4
0.2
COL7A1
IP6K2
PRKAR2A
ARIH2OS
WDR6
QARS
C3orf84
C3orf62
GPX1
UQCRC1
PRKAR2A−AS1
P4HTM
MIR6890
CCDC36
USP4
SNORA94
SLC25A20
DALRD3
USP19
MIR4271
RHOA
TMEM89
ARIH2
QRICH1
LAMB2P1
SLC26A6
MIR425
LAMB2
MIR6824
NDUFAF3
CCDC71
CELSR3
MIR191
KLHDC8B
MIR4793
IMPDH2
CELSR3−AS1
NCKIPSD
48.8
49
49.2
49.4
Position on chr3 (Mb)

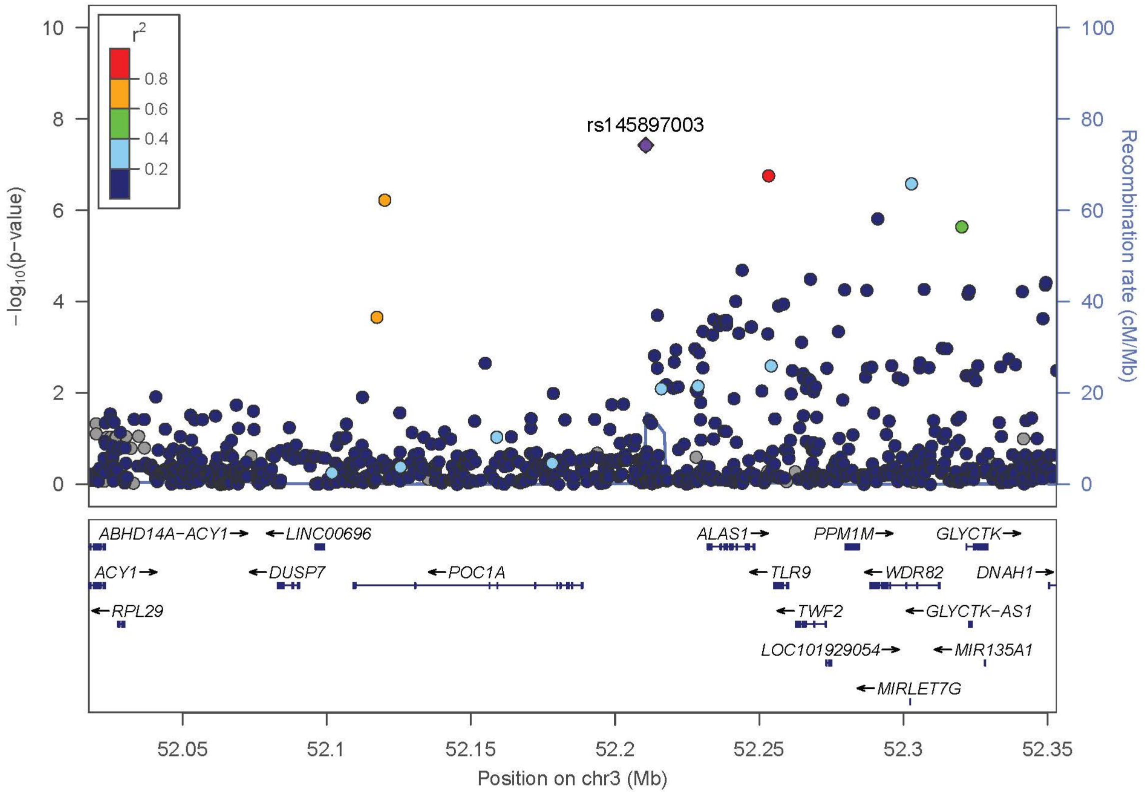
rs145897003
−log10(p−value)
Recombination rate (cM/Mb)
r2
0.8
0.6
0.4
0.2
ABHD14A−ACY1
LINC00696
ALAS1
PPM1M
GLYCTK
ACY1
DUSP7
POC1A
TLR9
WDR82
DNAH1
RPL29
TWF2
GLYCTK−AS1
LOC101929054
MIR135A1
MIRLET7G
52.05
52.1
52.15
52.2
52.25
52.3
52.35
Position on chr3 (Mb)

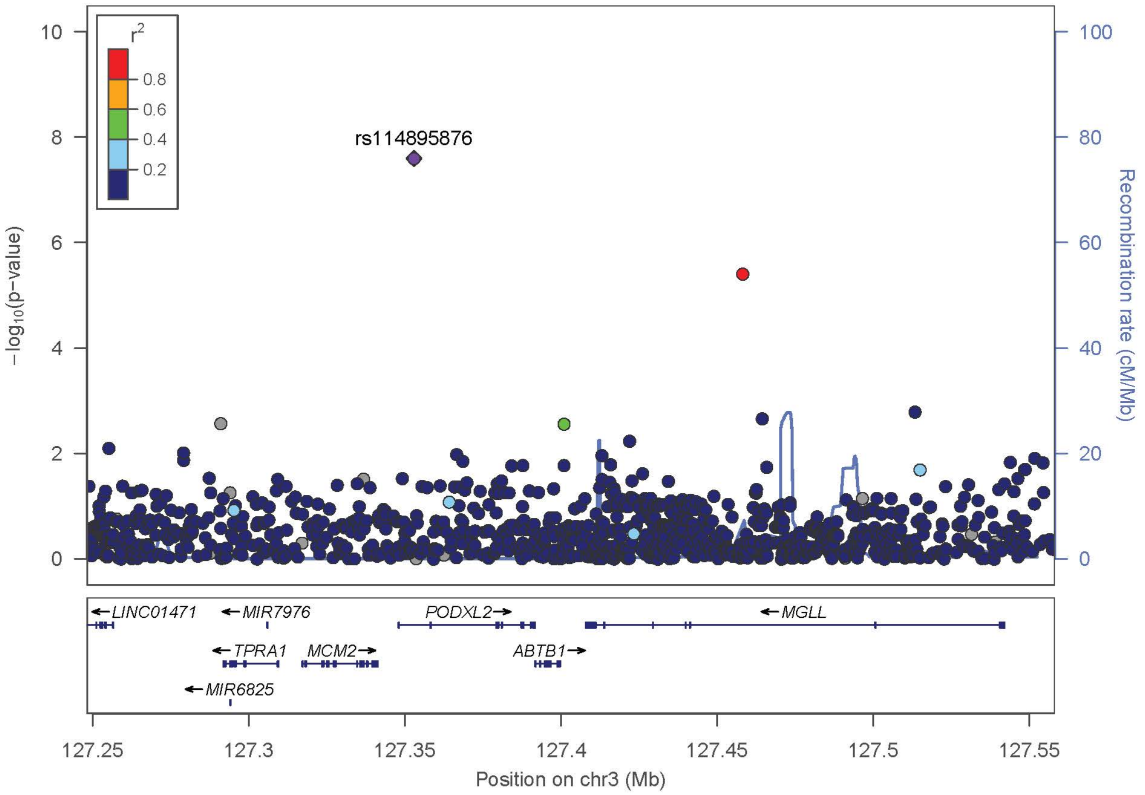

rs114895876
r2
0.8
0.6
0.4
0.2
−log10(p−value)
Recombination rate (cM/Mb)
LINC01471
MIR7976
PODXL2
MGLL
TPRA1
MCM2
ABTB1
MIR6825
127.25
127.3
127.35
127.4
127.45
127.5
127.55
Position on chr3 (Mb)


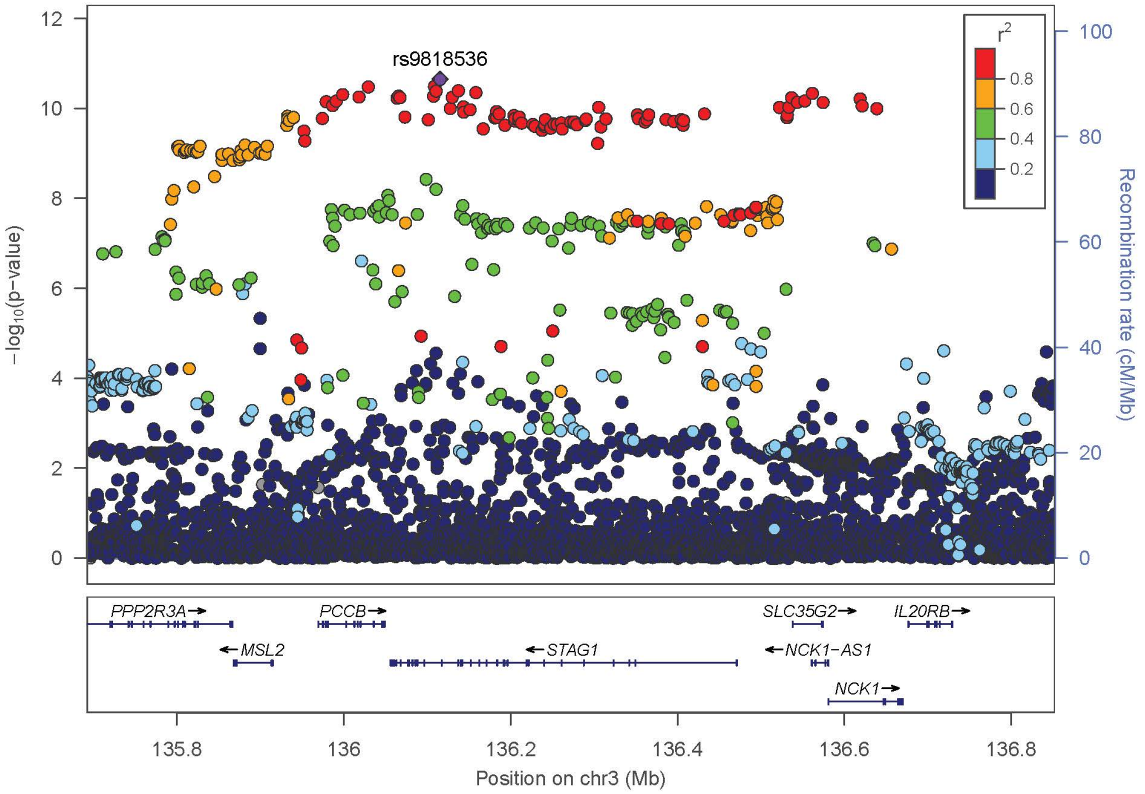

rs9818536
r2
0.8
0.6
0.4
0.2
−log10(p−value)
Recombination rate (cM/Mb)
PPP2R3A
PCCB
SLC35G2
IL20RB
MSL2
STAG1
NCK1−AS1
NCK1
135.8
136
136.2
136.4
136.6
136.8
Position on chr3 (Mb)

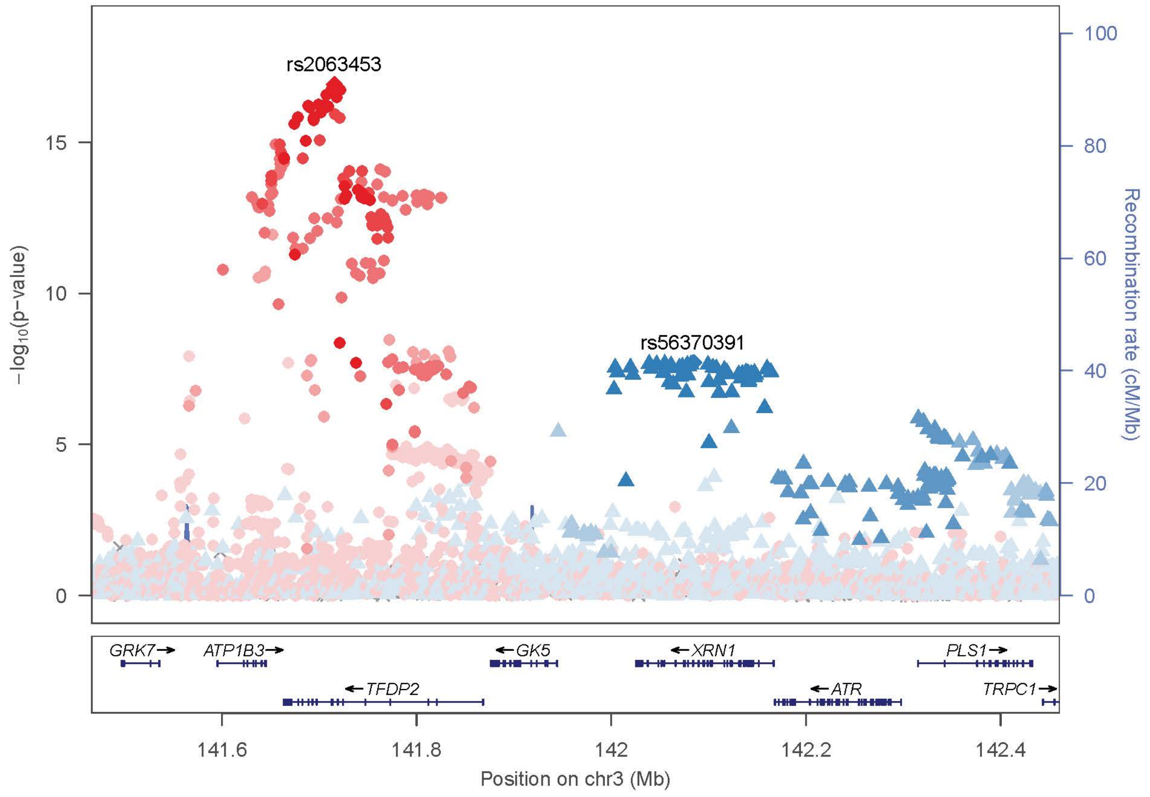

rs2063453
rs56370391
−log10(p−value)
Recombination rate (cM/Mb)
0
5
10
15
0
20
40
60
80
100
GRK7→
ATP1B3→
←GK5
←XRN1
PLS1→
←TFDP2
←ATR
TRPC1→
141.6
141.8
142
142.2
142.4
Position on chr3 (Mb)


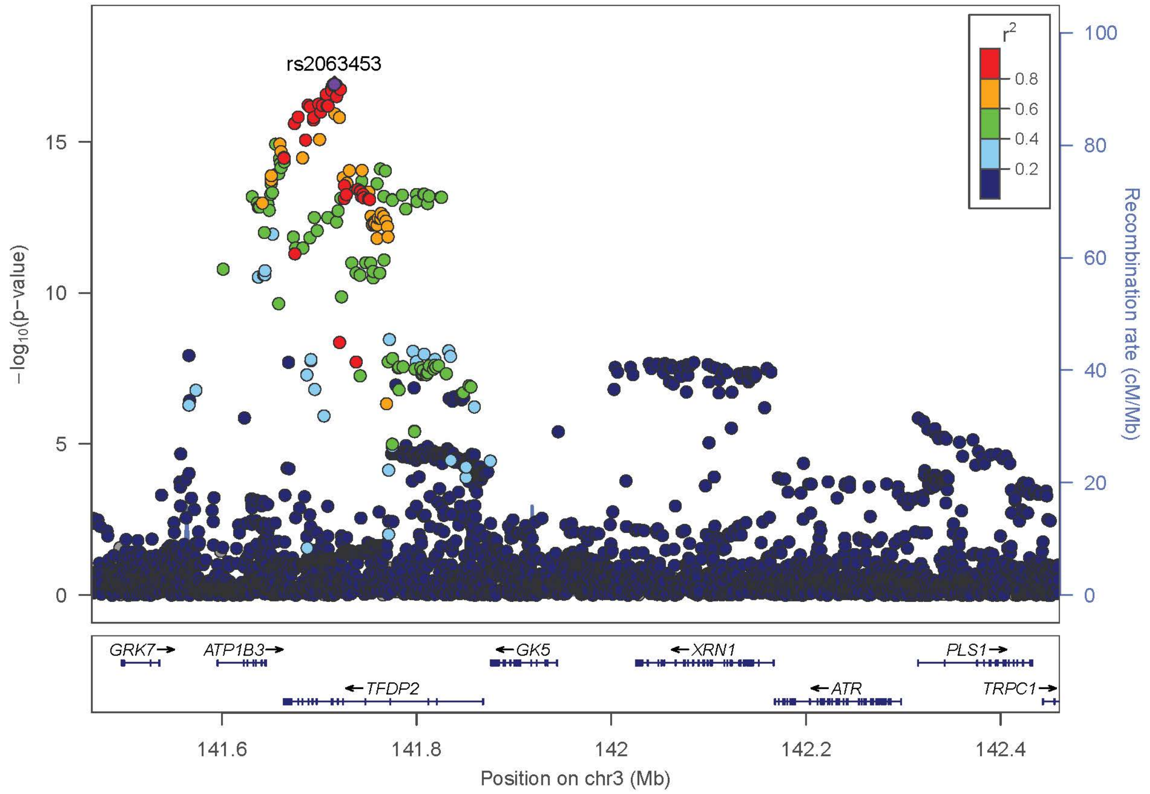

rs2063453
r2
0.8
0.6
0.4
0.2
−log10(p−value)
Recombination rate (cM/Mb)
0
5
10
15
0
20
40
60
80
100
GRK7→
ATP1B3→
←GK5
←XRN1
PLS1→
←TFDP2
←ATR
TRPC1→
141.6
141.8
142
142.2
142.4
Position on chr3 (Mb)

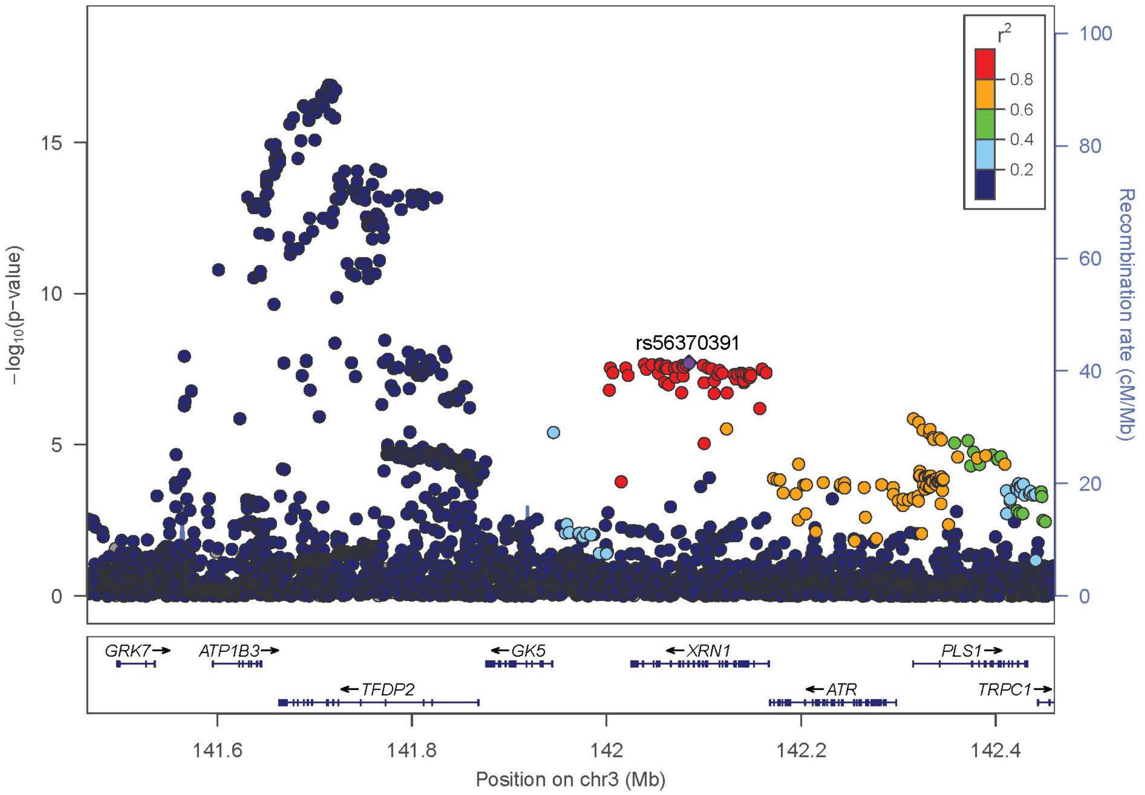

r2
0.8
0.6
0.4
0.2
rs56370391
−log10(p−value)
Recombination rate (cM/Mb)
GRK7
ATP1B3
TFDP2
GK5
XRN1
ATR
PLS1
TRPC1
Position on chr3 (Mb)


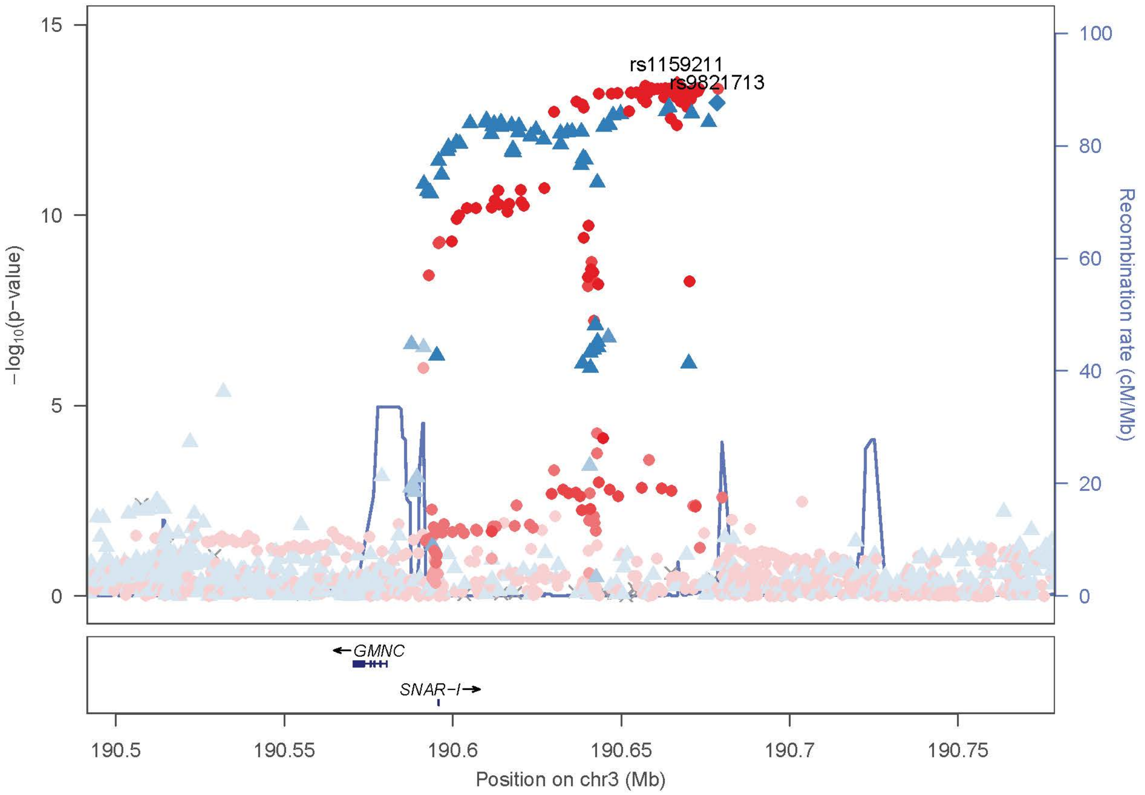

rs1159211
rs9821713
−log10(p−value)
Recombination rate (cM/Mb)
GMNC
SNAR−I
Position on chr3 (Mb)

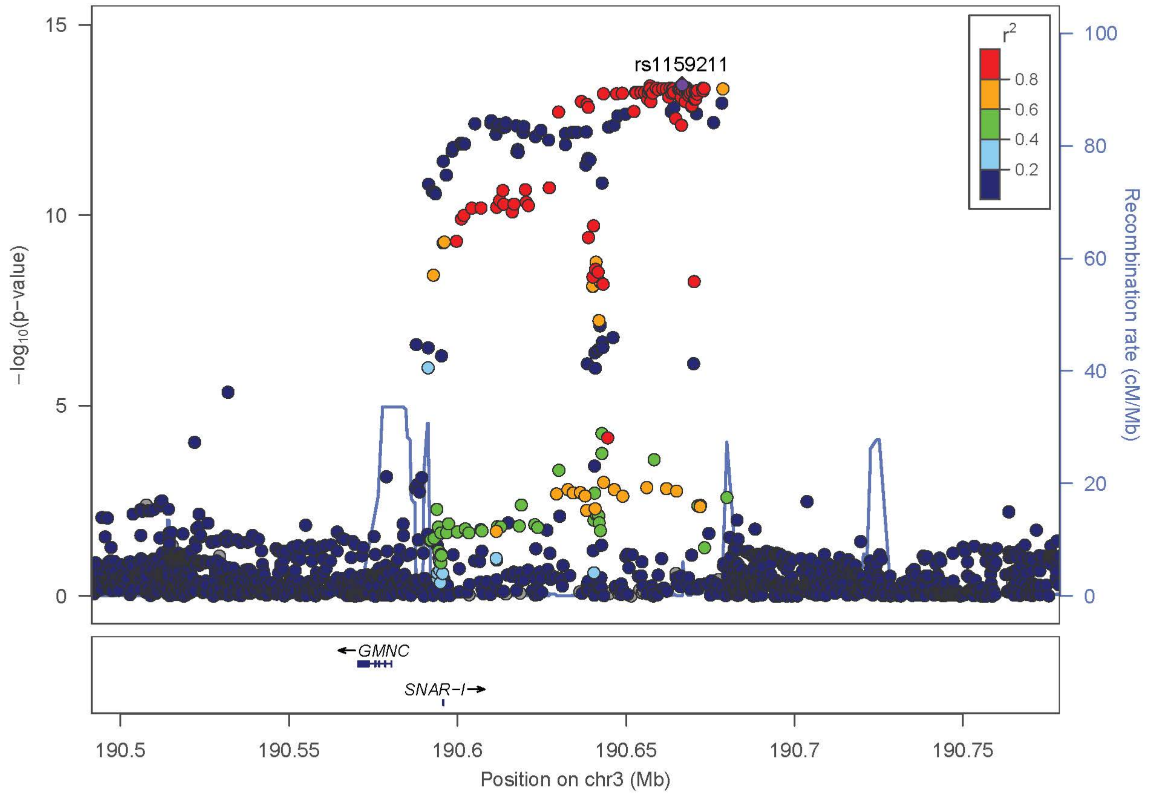
rs1159211
r^2
0.8
0.6
0.4
0.2
−log10(p−value)
Recombination rate (cM/Mb)
GMNC
SNAR−I
190.5
190.55
190.6
190.65
190.7
190.75
Position on chr3 (Mb)

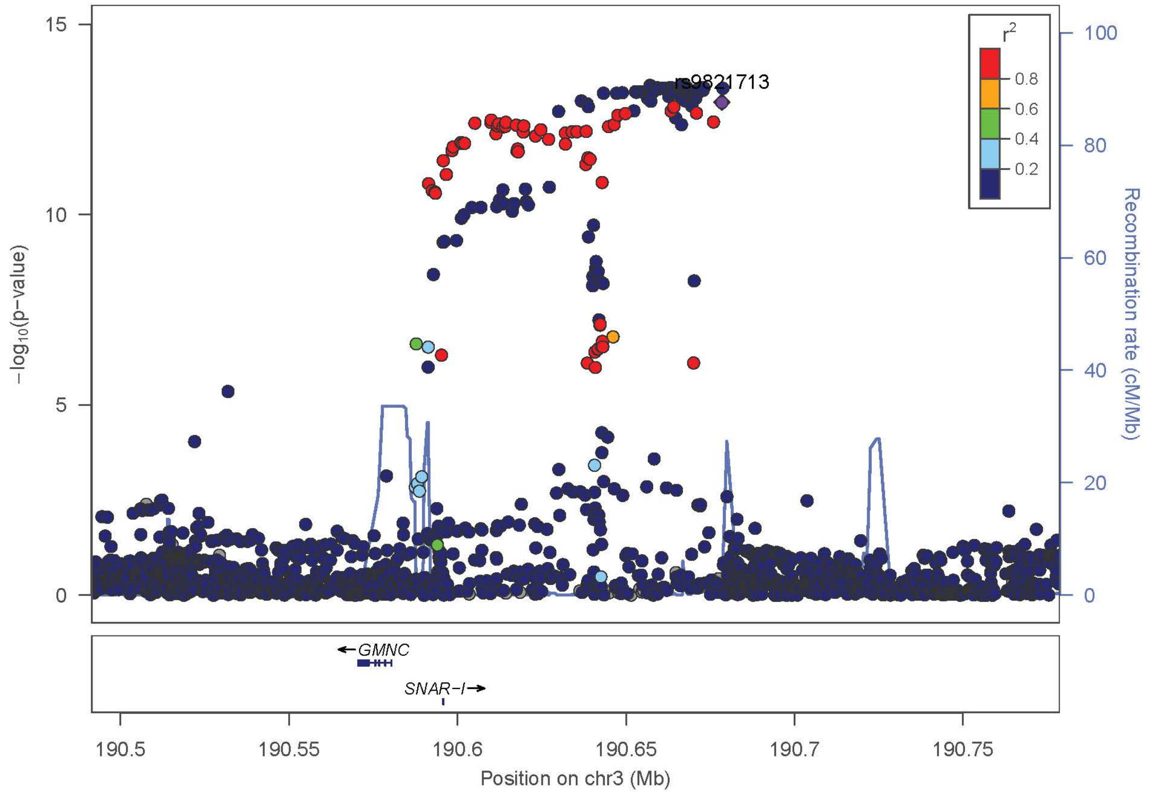
rs9821713
r^2
0.8
0.6
0.4
0.2
−log10(p−value)
Recombination rate (cM/Mb)
GMNC
SNAR−I
190.5
190.55
190.6
190.65
190.7
190.75
Position on chr3 (Mb)

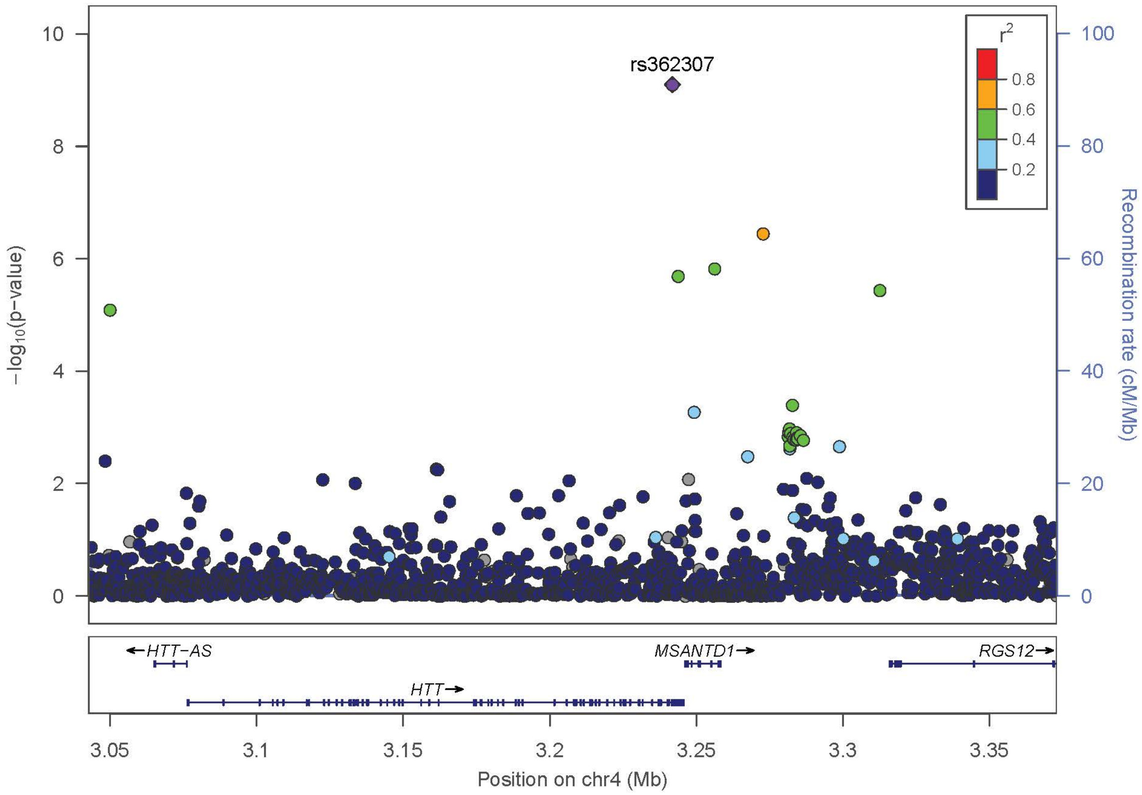
rs362307
r²
0.8
0.6
0.4
0.2
−log10(p−value)
Recombination rate (cM/Mb)
HTT−AS
MSANTD1
RGS12
HTT
Position on chr4 (Mb)

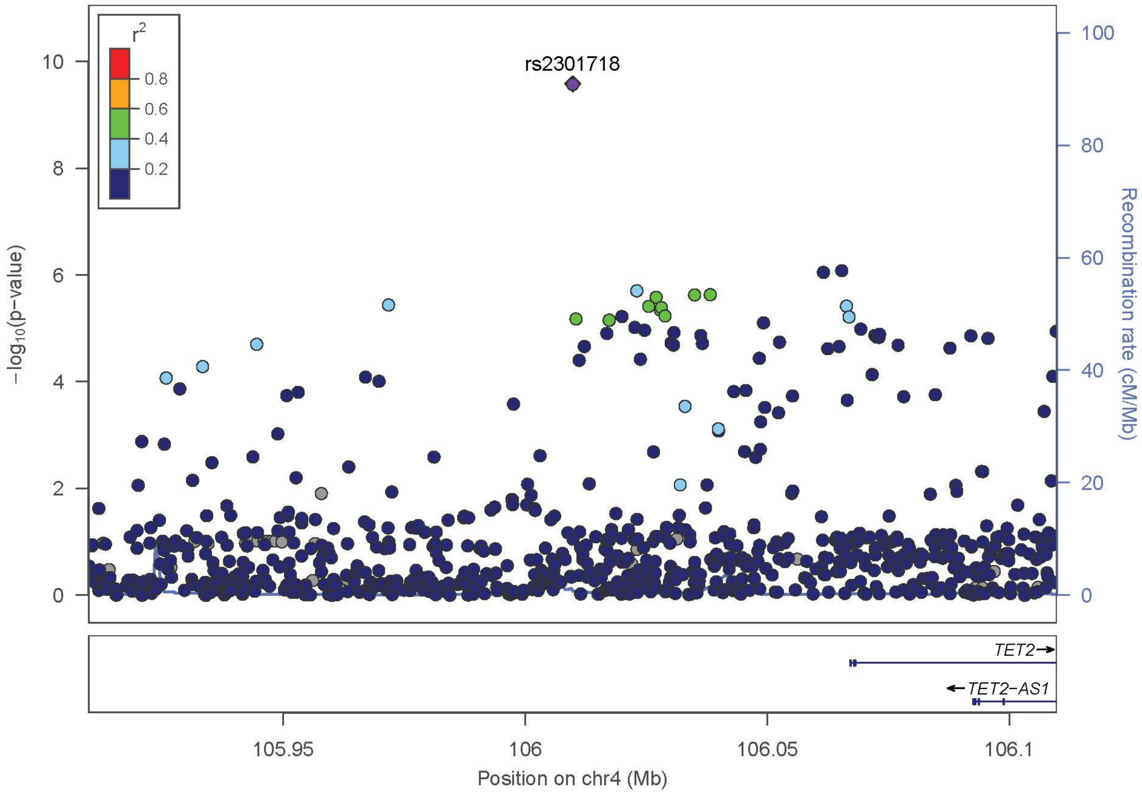
rs2301718
r²
0.8
0.6
0.4
0.2
−log10(p−value)
Recombination rate (cM/Mb)
TET2
TET2−AS1
Position on chr4 (Mb)

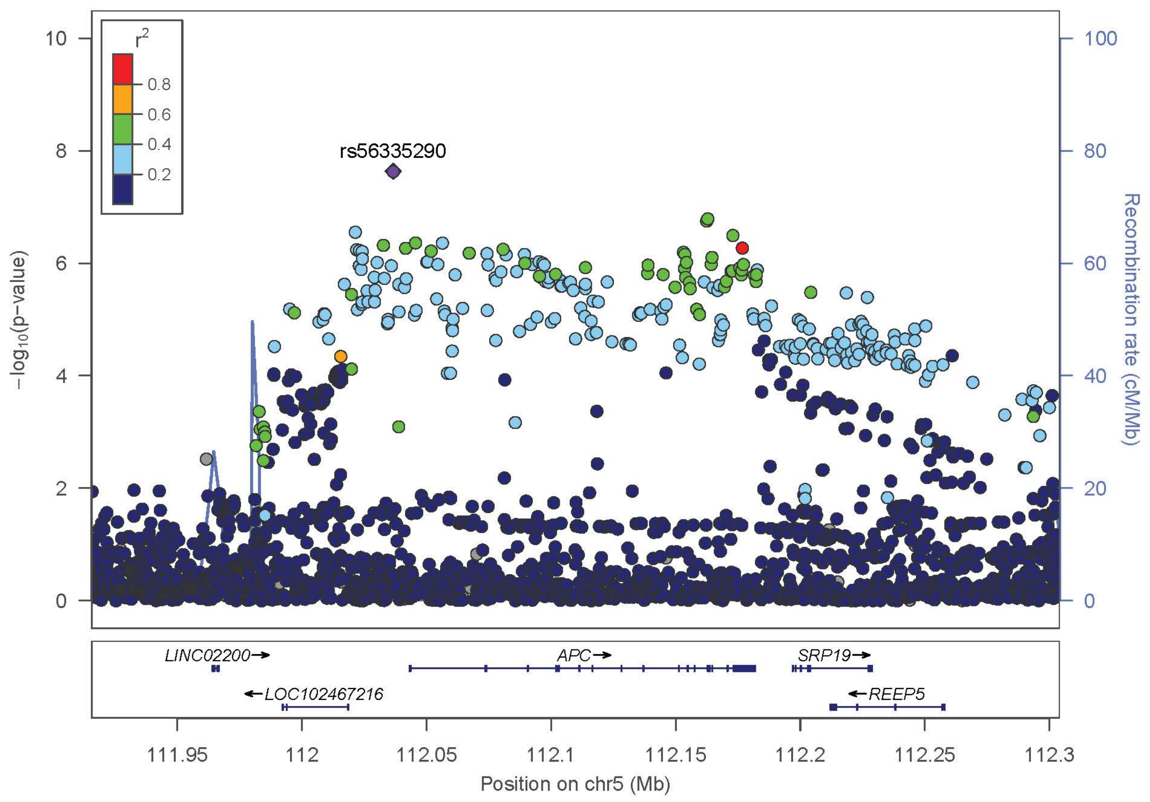
rs56335290
r2
0.8
0.6
0.4
0.2
−log10(p−value)
Recombination rate (cM/Mb)
LINC02200→
APC→
SRP19→
←LOC102467216
←REEP5
111.95
112
112.05
112.1
112.15
112.2
112.25
112.3
Position on chr5 (Mb)

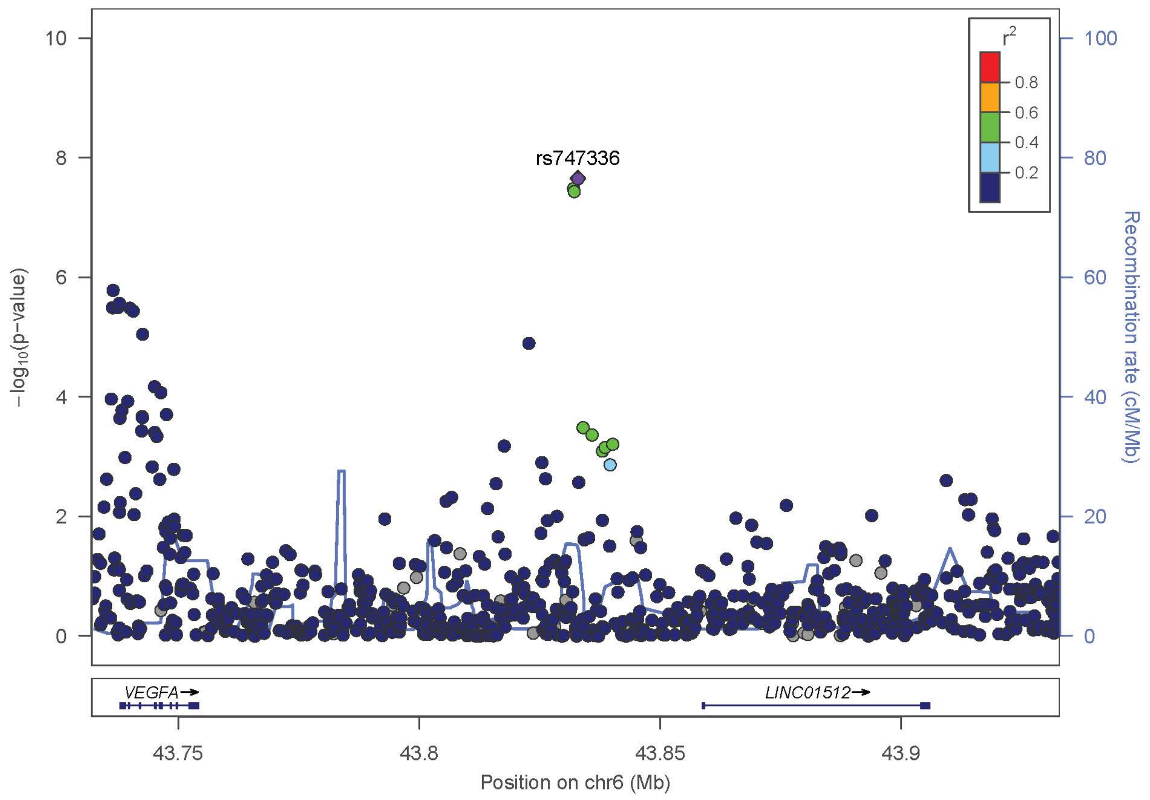
rs747336
r2
0.8
0.6
0.4
0.2
−log10(p−value)
Recombination rate (cM/Mb)
VEGFA→
LINC01512→
43.75
43.8
43.85
43.9
Position on chr6 (Mb)

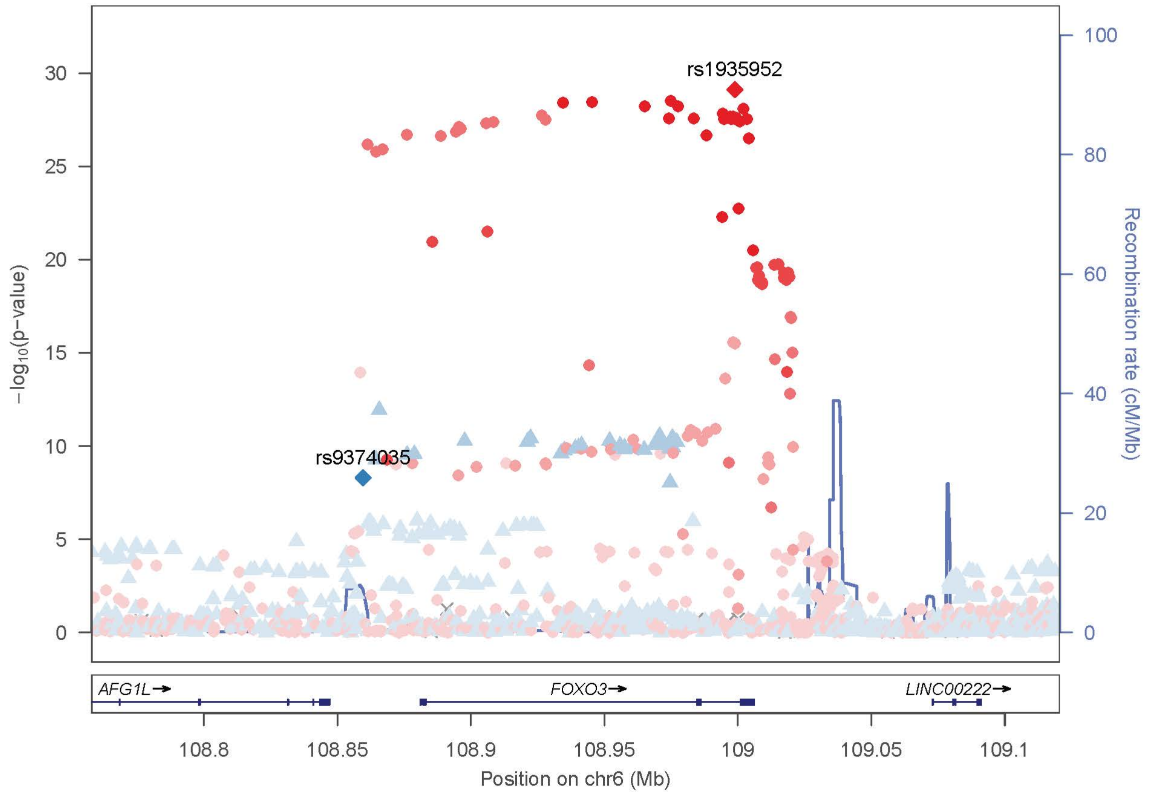
rs1935952
rs9374035
−log10(p−value)
Recombination rate (cM/Mb)
AFG1L
FOXO3
LINC00222
Position on chr6 (Mb)

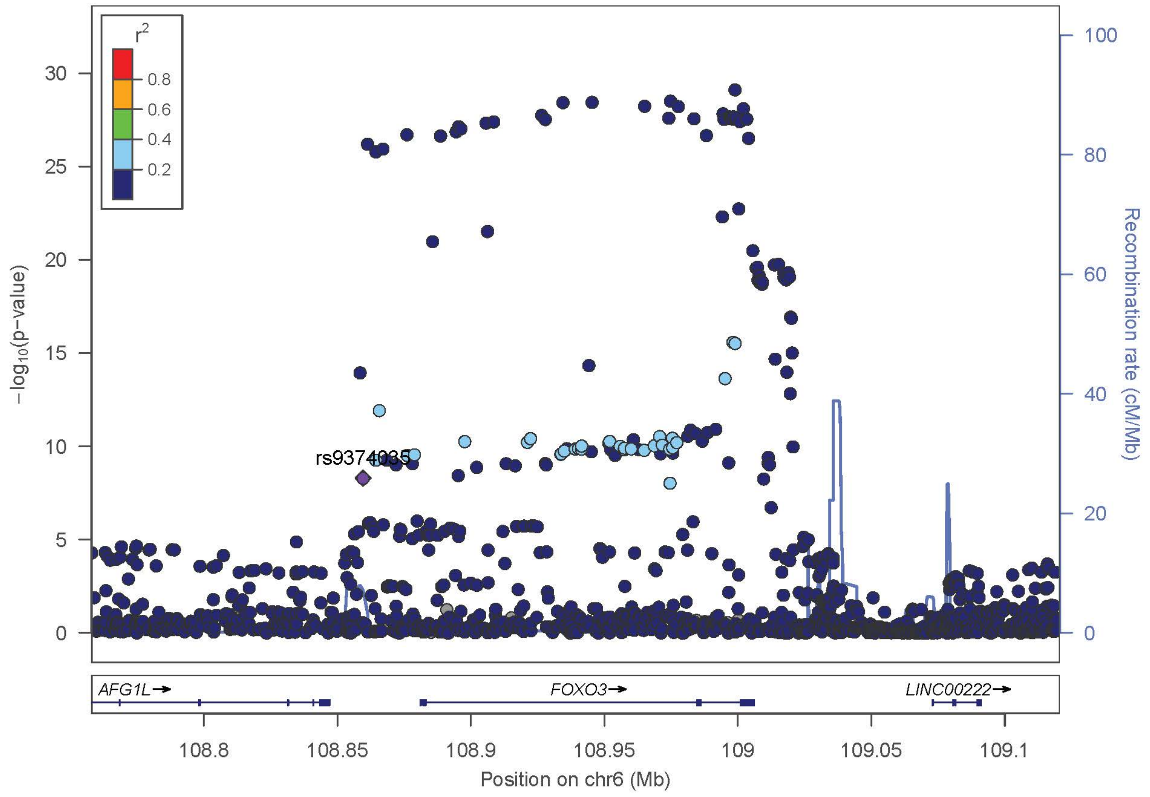
r2
0.8
0.6
0.4
0.2
rs9374035
−log10(p−value)
Recombination rate (cM/Mb)
AFG1L
FOXO3
LINC00222
Position on chr6 (Mb)

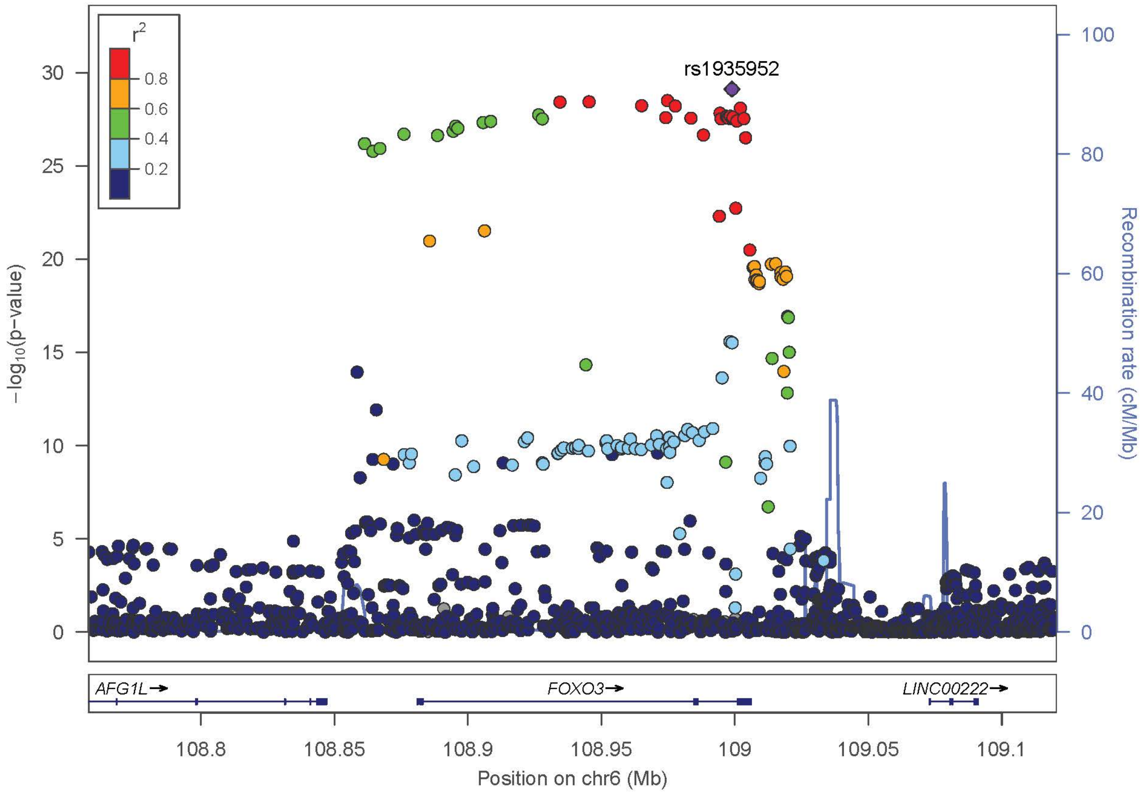

r2
0.8
0.6
0.4
0.2
rs1935952
−log10(p−value)
Recombination rate (cM/Mb)
AFG1L→
FOXO3→
LINC00222→
108.8
108.85
108.9
108.95
109
109.05
109.1
Position on chr6 (Mb)


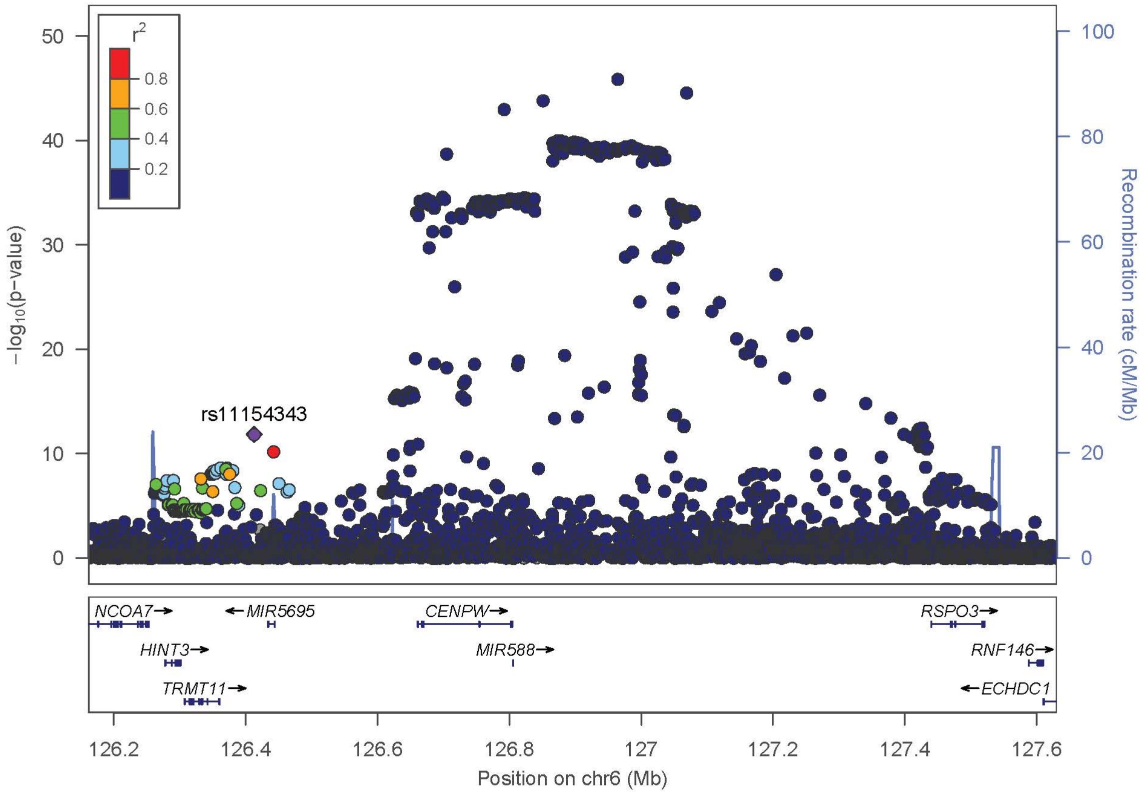

r2
0.8
0.6
0.4
0.2
rs11154343
−log10(p−value)
Recombination rate (cM/Mb)
NCOA7→
←MIR5695
CENPW→
RSPO3→
HINT3→
MIR588→
RNF146→
TRMT11→
←ECHDC1
126.2
126.4
126.6
126.8
127
127.2
127.4
127.6
Position on chr6 (Mb)

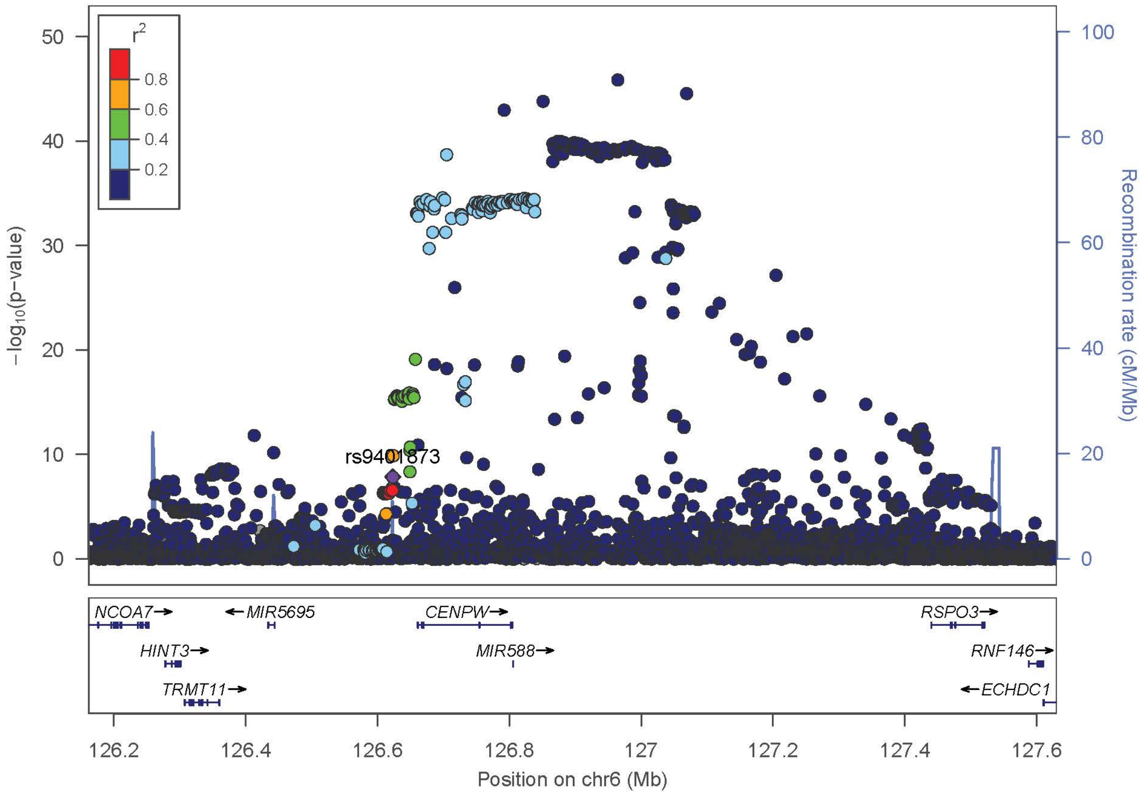

r2
0.8
0.6
0.4
0.2
rs9401873
−log10(p−value)
Recombination rate (cM/Mb)
NCOA7
MIR5695
CENPW
RSPO3
HINT3
MIR588
RNF146
TRMT11
ECHDC1
Position on chr6 (Mb)


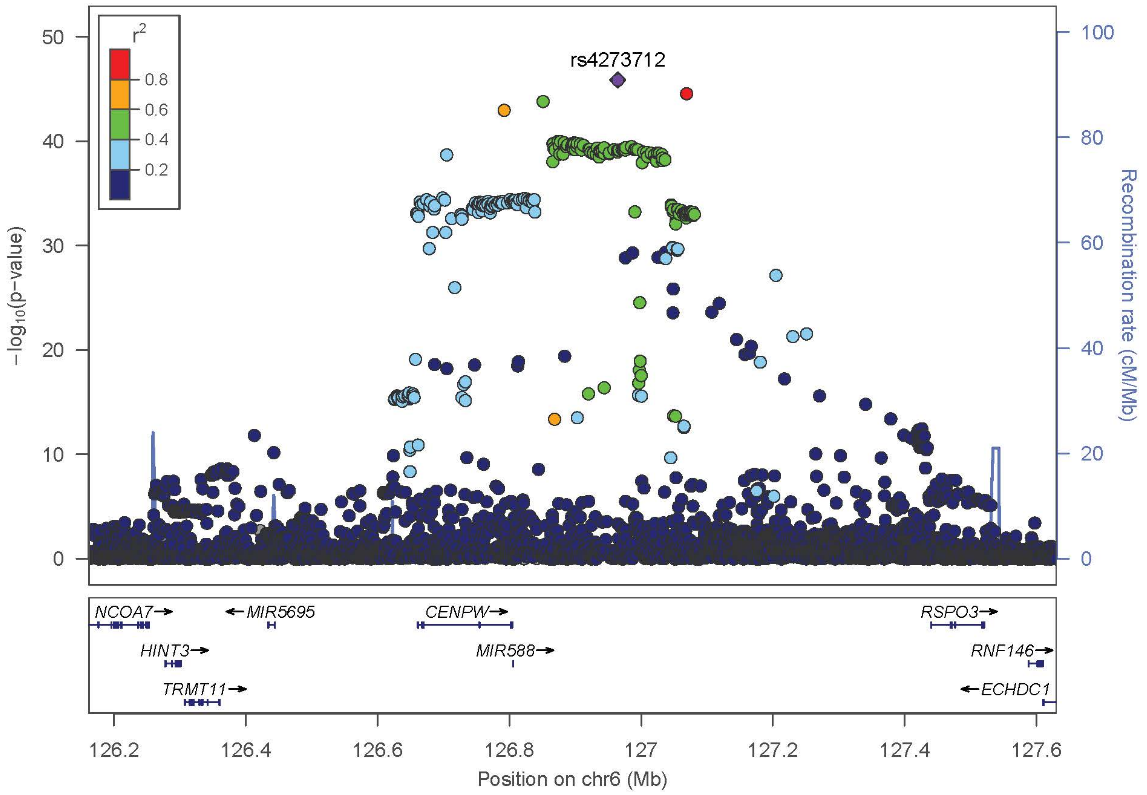

r2
0.8
0.6
0.4
0.2
rs4273712
−log10(p−value)
Recombination rate (cM/Mb)
NCOA7
MIR5695
CENPW
RSPO3
HINT3
MIR588
RNF146
TRMT11
ECHDC1
Position on chr6 (Mb)

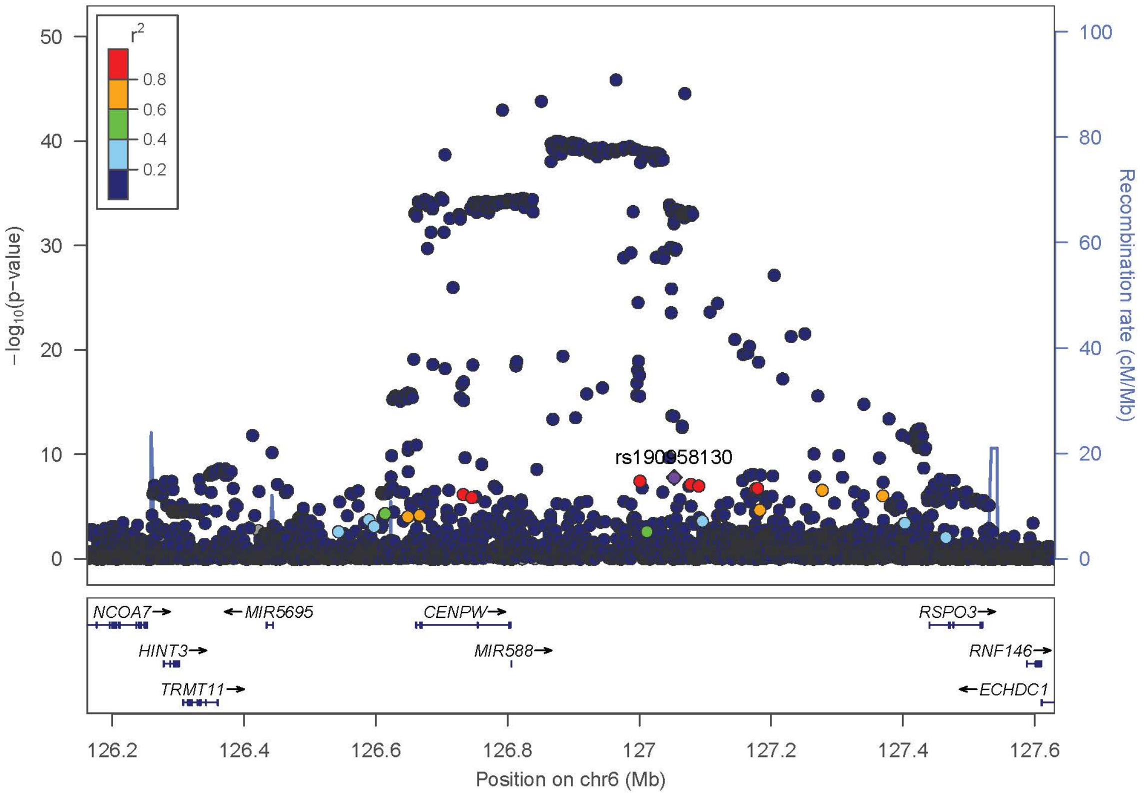
r2
0.8
0.6
0.4
0.2
rs190958130
−log10(p−value)
Recombination rate (cM/Mb)
NCOA7
MIR5695
CENPW
RSPO3
HINT3
MIR588
RNF146
TRMT11
ECHDC1
Position on chr6 (Mb)

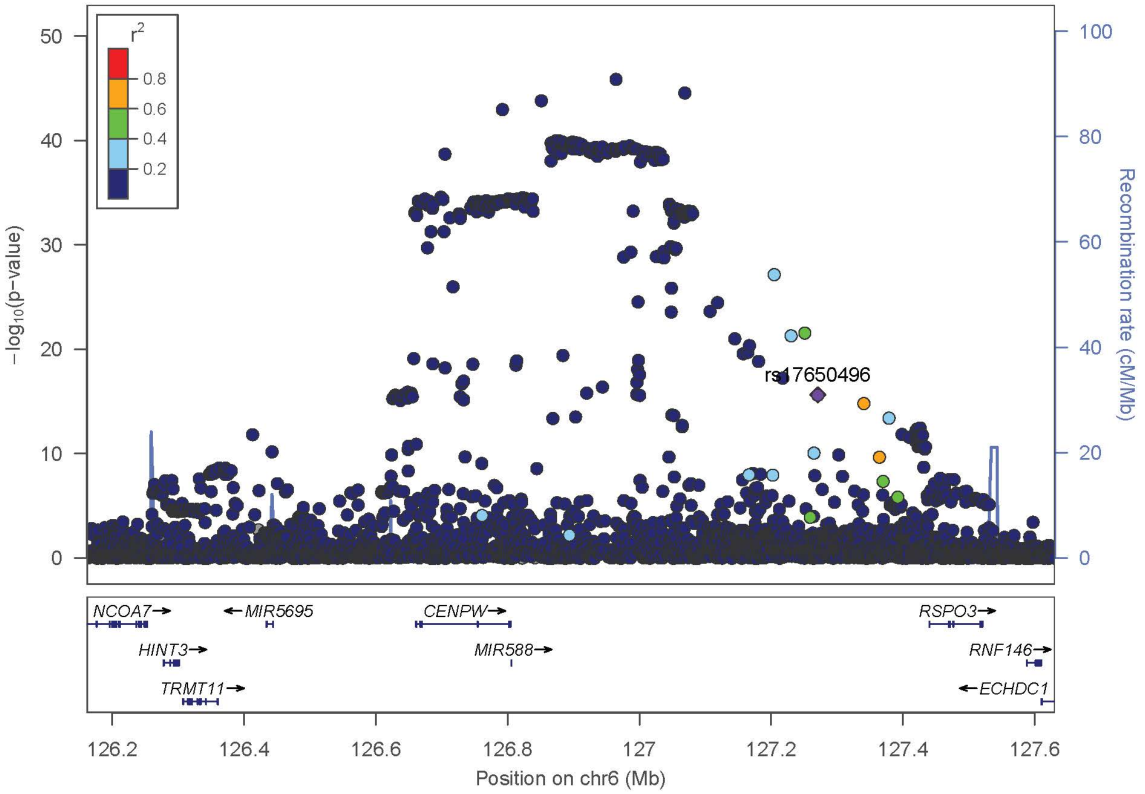
r2
0.8
0.6
0.4
0.2
rs17650496
−log10(p−value)
Recombination rate (cM/Mb)
NCOA7
MIR5695
CENPW
RSPO3
HINT3
MIR588
RNF146
TRMT11
ECHDC1
Position on chr6 (Mb)

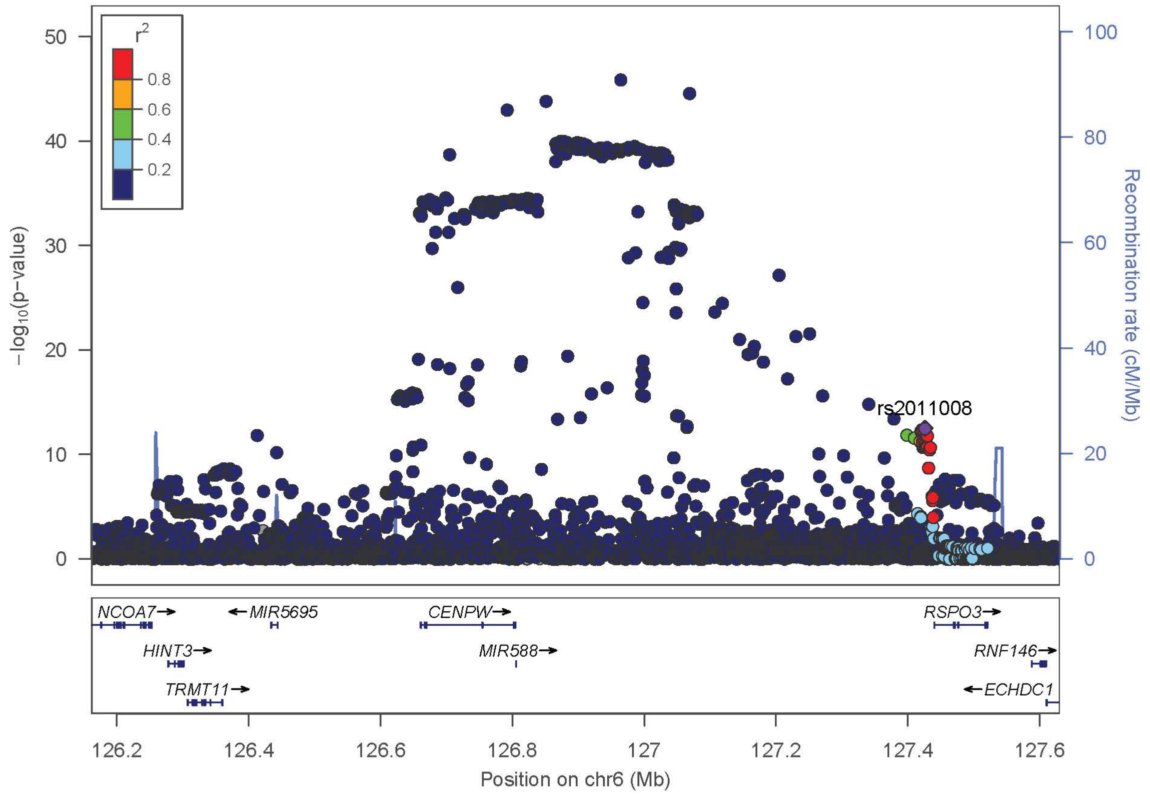

r2
0.8
0.6
0.4
0.2
rs2011008
−log10(p-value)
Recombination rate (cM/Mb)
NCOA7
MIR5695
CENPW
RSPO3
HINT3
MIR588
RNF146
TRMT11
ECHDC1
Position on chr6 (Mb)


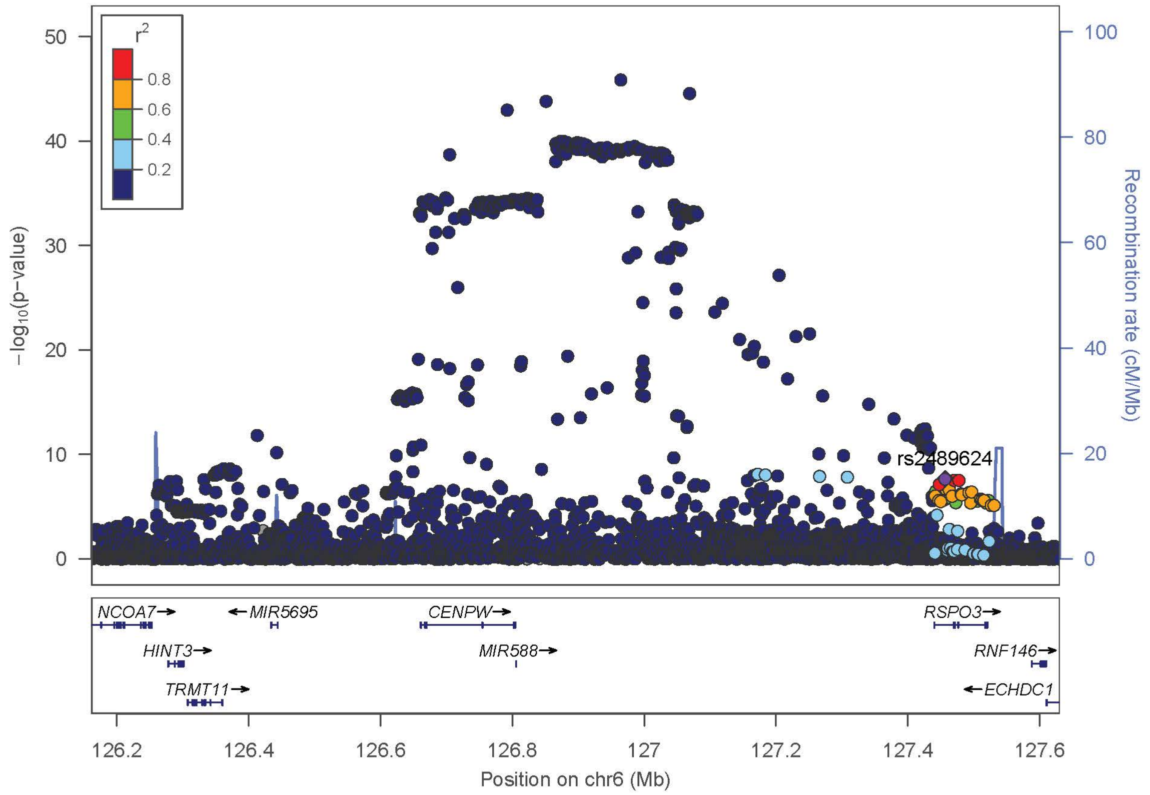

r2
0.8
0.6
0.4
0.2
rs2489624
−log10(p-value)
Recombination rate (cM/Mb)
NCOA7
MIR5695
CENPW
RSPO3
HINT3
MIR588
RNF146
TRMT11
ECHDC1
Position on chr6 (Mb)

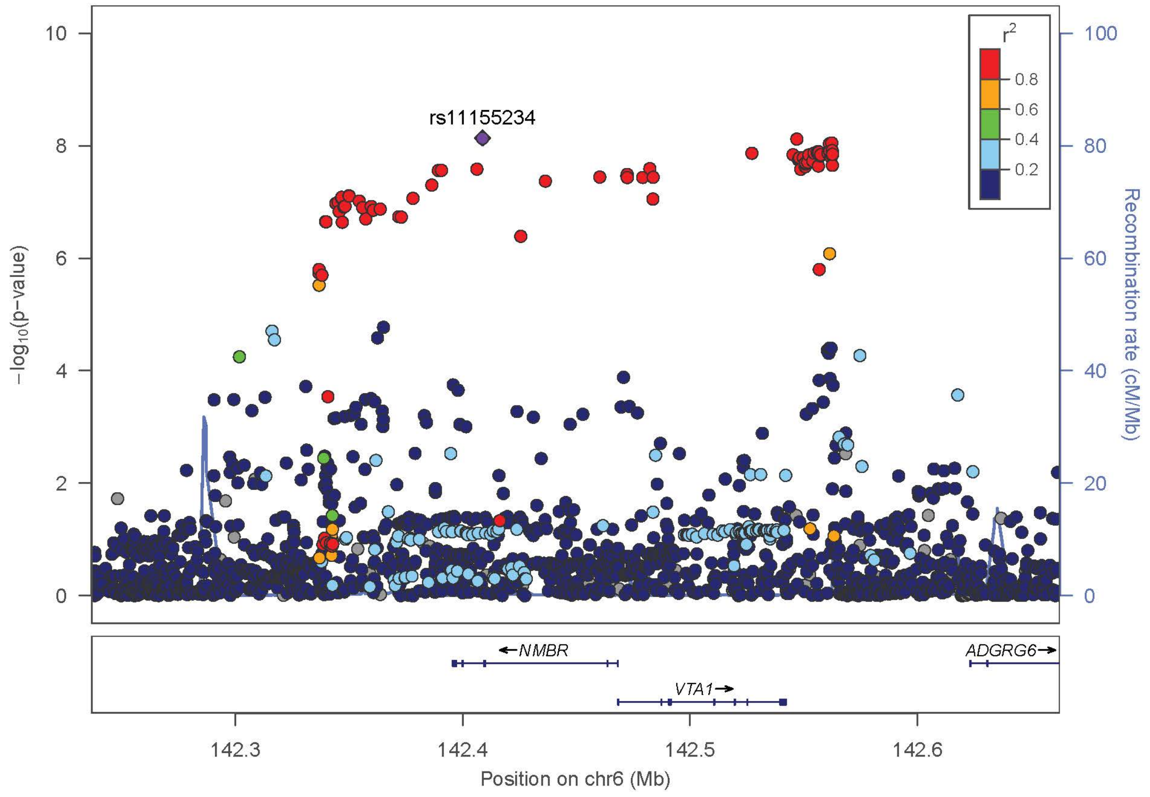
rs11155234
r2
0.8
0.6
0.4
0.2
−log10(p−value)
Recombination rate (cM/Mb)
NMBR
ADGRG6
VTA1
142.3
142.4
142.5
142.6
Position on chr6 (Mb)

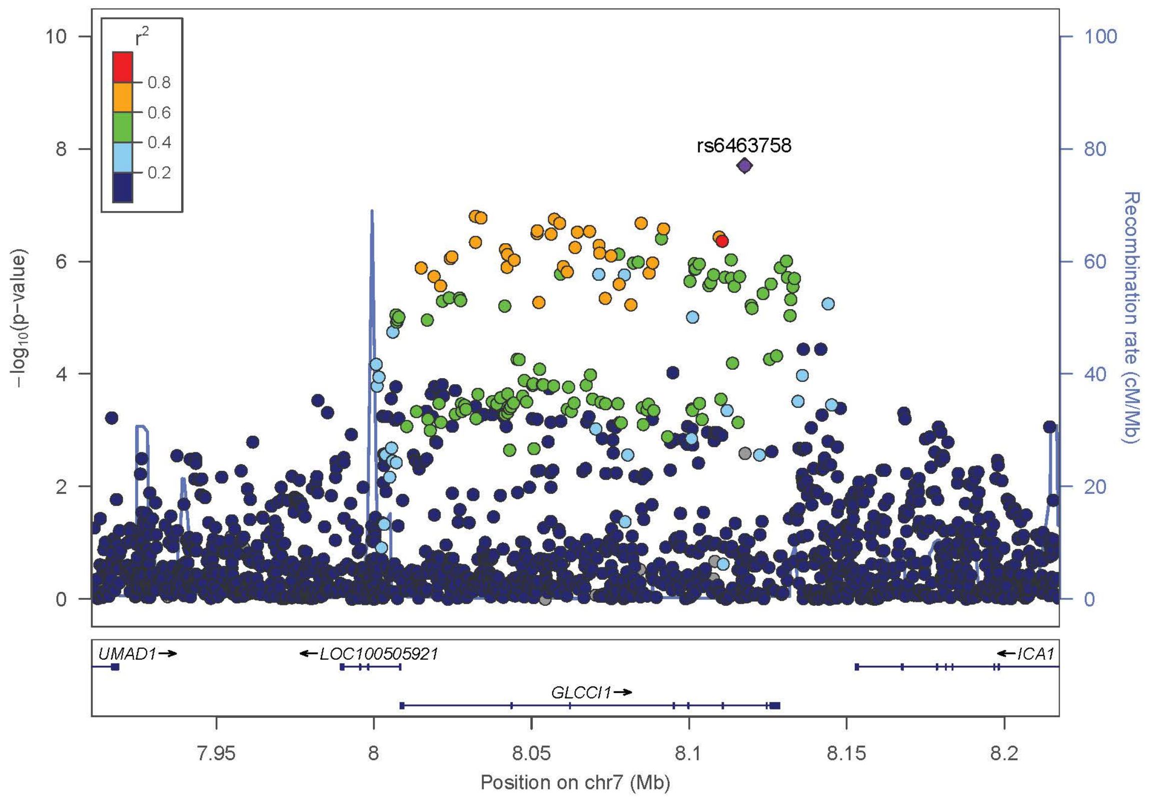
rs6463758
r2
0.8
0.6
0.4
0.2
−log10(p−value)
Recombination rate (cM/Mb)
UMAD1
LOC100505921
ICA1
GLCCI1
7.95
8
8.05
8.1
8.15
8.2
Position on chr7 (Mb)

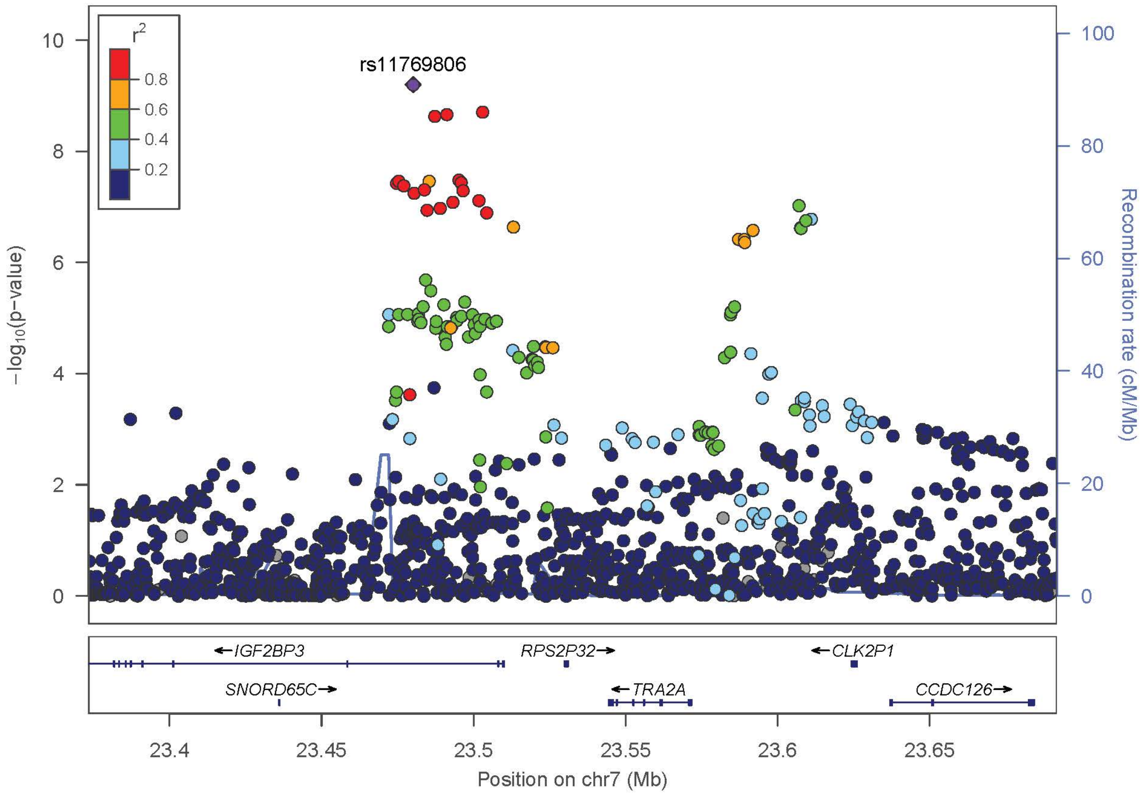

rs11769806
r2
0.8
0.6
0.4
0.2
−log10(p-value)
Recombination rate (cM/Mb)
IGF2BP3
RPS2P32
CLK2P1
SNORD65C
TRA2A
CCDC126
23.4
23.45
23.5
23.55
23.6
23.65
Position on chr7 (Mb)


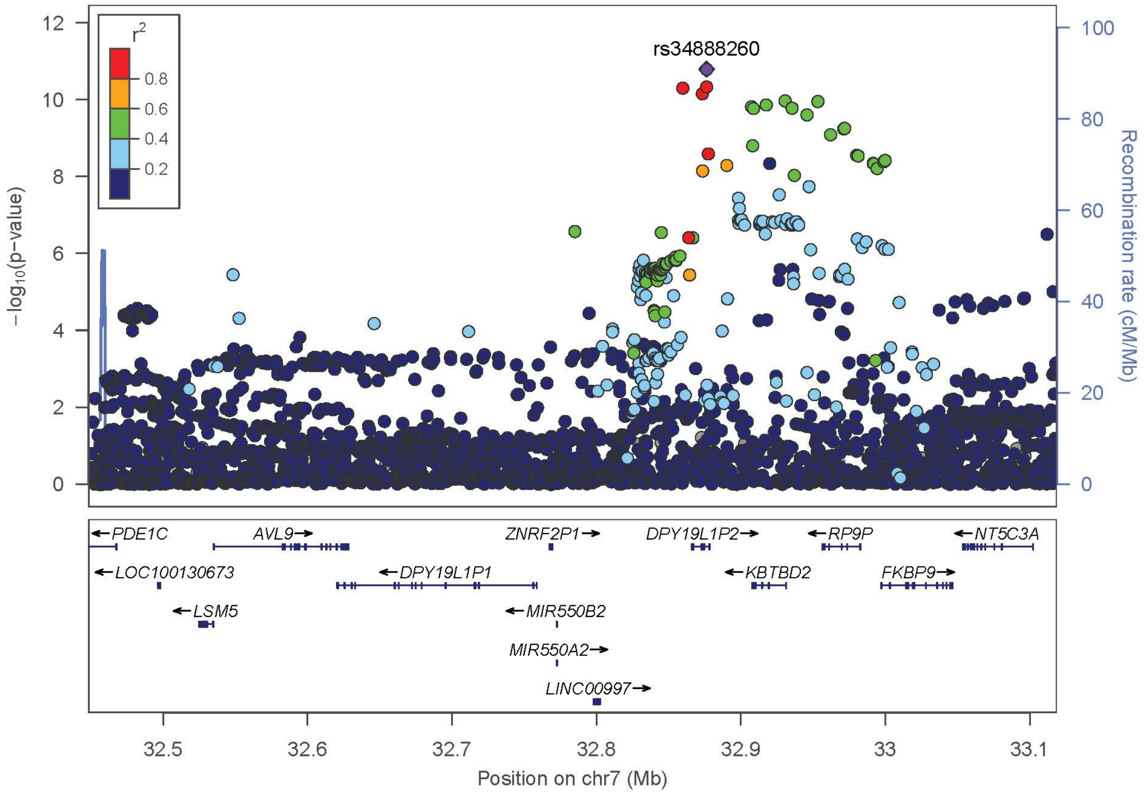

rs34888260
r2
0.8
0.6
0.4
0.2
−log10(p-value)
Recombination rate (cM/Mb)
PDE1C
AVL9
ZNRF2P1
DPY19L1P2
RP9P
NT5C3A
LOC100130673
DPY19L1P1
KBTBD2
FKBP9
LSM5
MIR550B2
MIR550A2
LINC00997
32.5
32.6
32.7
32.8
32.9
33
33.1
Position on chr7 (Mb)

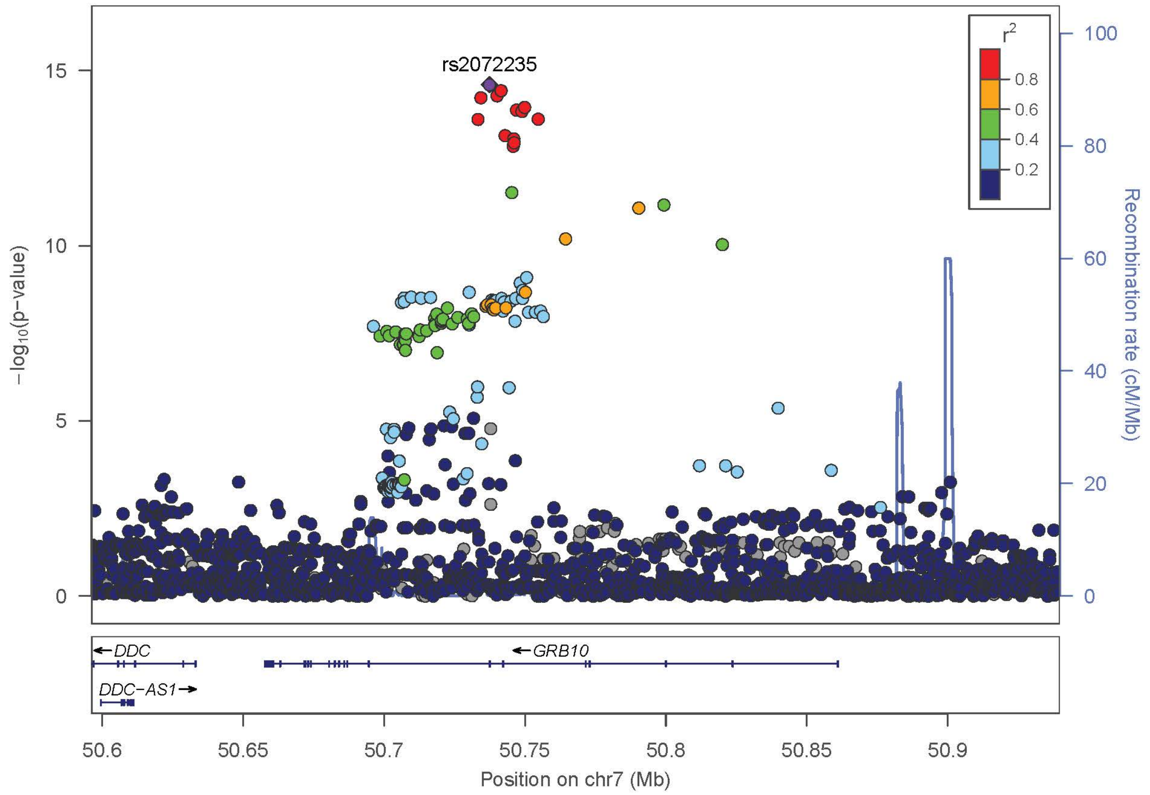
rs2072235
r2
0.8
0.6
0.4
0.2
15
10
5
0
−log10(p−value)
100
80
60
40
20
0
Recombination rate (cM/Mb)
DDC
GRB10
DDC−AS1
50.6
50.65
50.7
50.75
50.8
50.85
50.9
Position on chr7 (Mb)

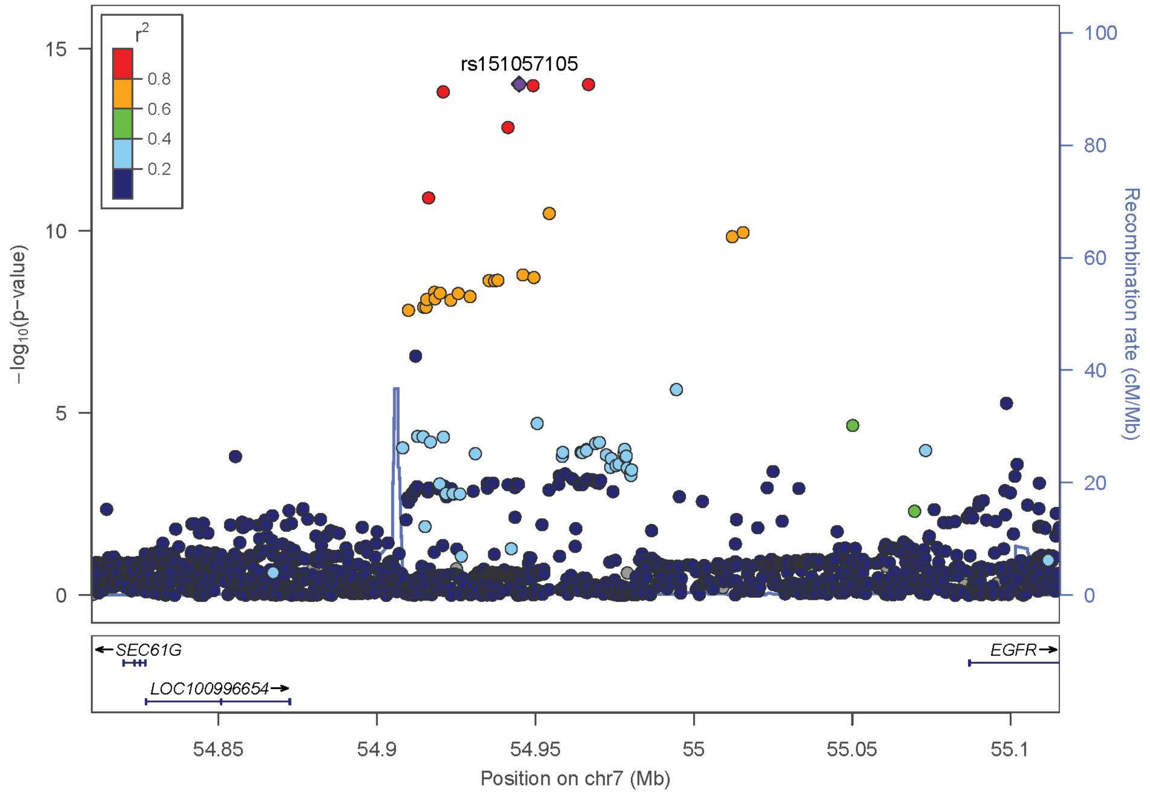
rs151057105
r2
0.8
0.6
0.4
0.2
15
10
5
0
−log10(p−value)
100
80
60
40
20
0
Recombination rate (cM/Mb)
SEC61G
EGFR
LOC100996654
54.85
54.9
54.95
55
55.05
55.1
Position on chr7 (Mb)

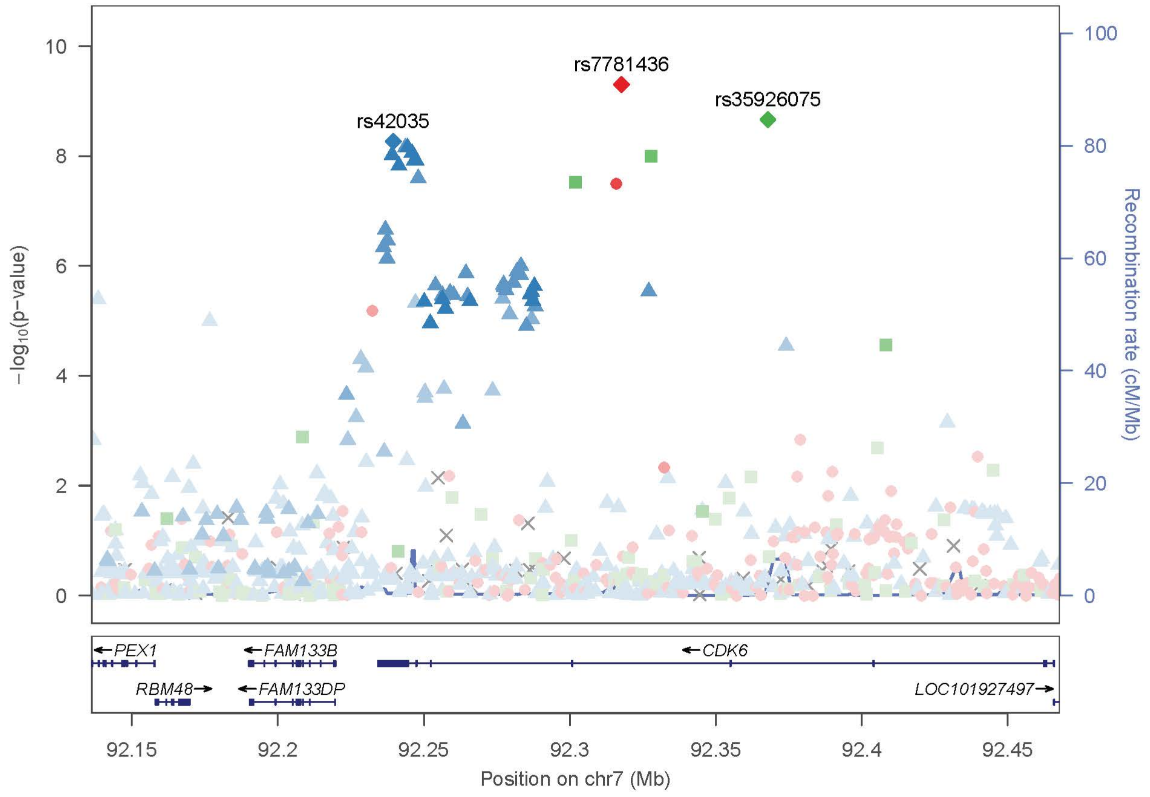
rs7781436
rs35926075
rs42035
−log10(p−value)
Recombination rate (cM/Mb)
PEX1
FAM133B
CDK6
RBM48
FAM133DP
LOC101927497
Position on chr7 (Mb)

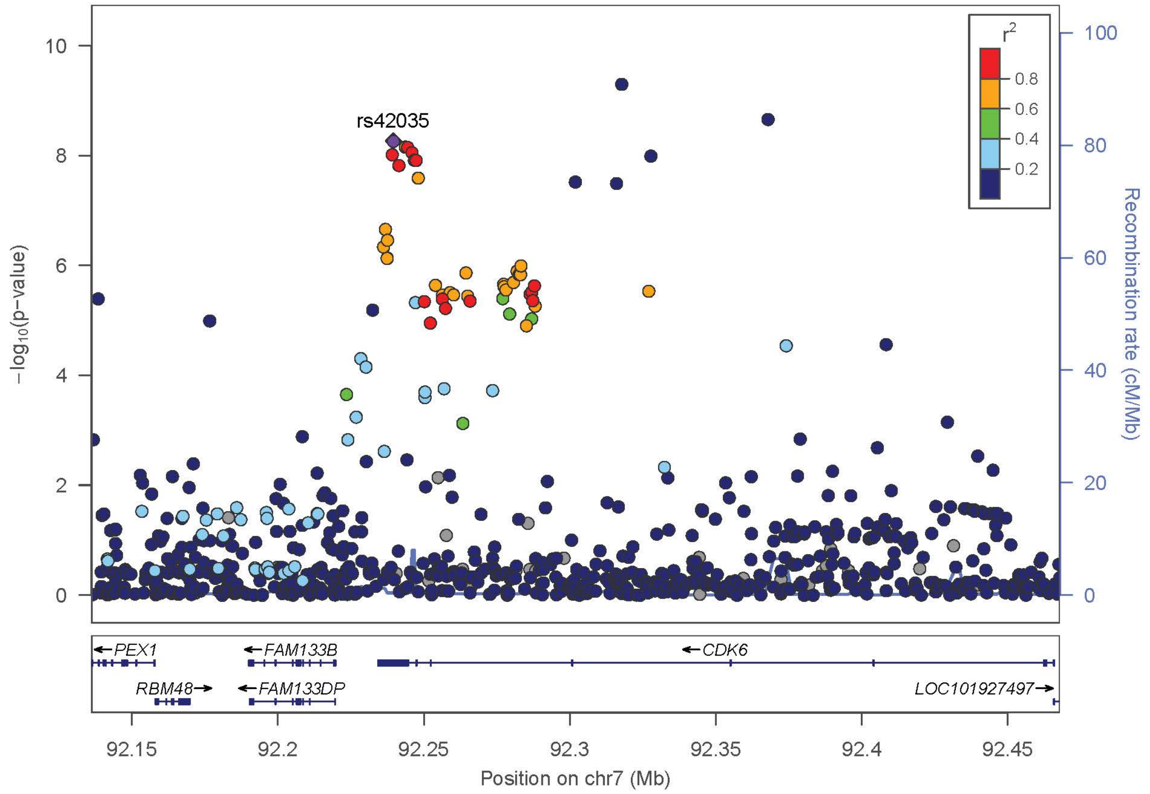
rs42035
r2
0.8
0.6
0.4
0.2
−log10(p−value)
Recombination rate (cM/Mb)
PEX1
FAM133B
CDK6
RBM48
FAM133DP
LOC101927497
Position on chr7 (Mb)

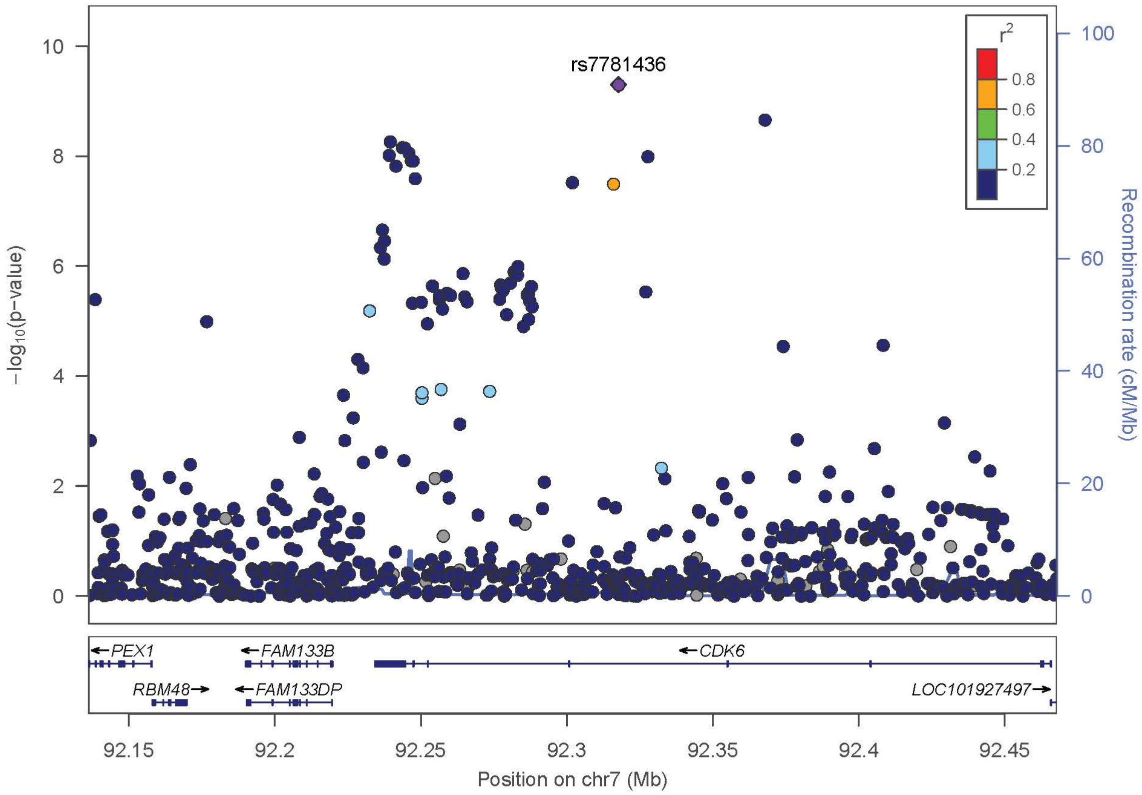

rs7781436
r2
0.8
0.6
0.4
0.2
−log10(p−value)
Recombination rate (cM/Mb)
PEX1
FAM133B
CDK6
RBM48
FAM133DP
LOC101927497
92.15
92.2
92.25
92.3
92.35
92.4
92.45
Position on chr7 (Mb)


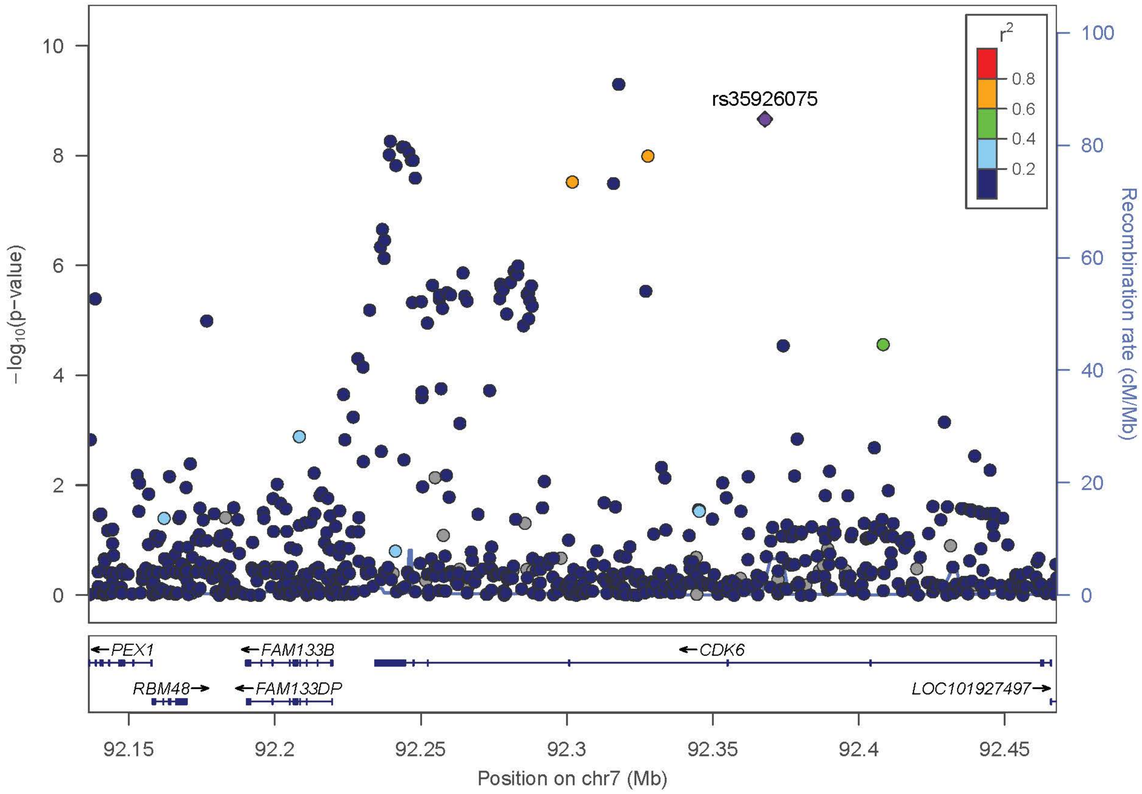

rs35926075
r2
0.8
0.6
0.4
0.2
−log10(p−value)
Recombination rate (cM/Mb)
PEX1
FAM133B
CDK6
RBM48
FAM133DP
LOC101927497
92.15
92.2
92.25
92.3
92.35
92.4
92.45
Position on chr7 (Mb)

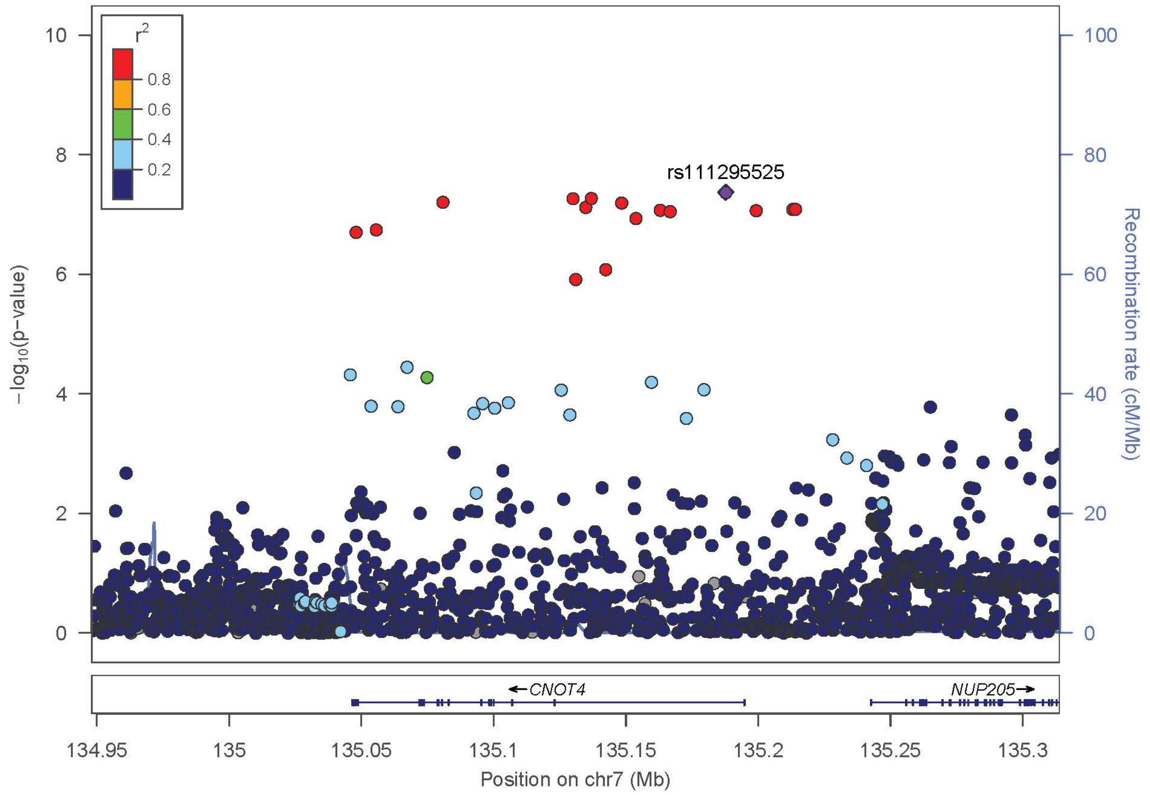

r2
0.8
0.6
0.4
0.2
rs111295525
−log10(p−value)
Recombination rate (cM/Mb)
CNOT4
NUP205
134.95
135
135.05
135.1
135.15
135.2
135.25
135.3
Position on chr7 (Mb)


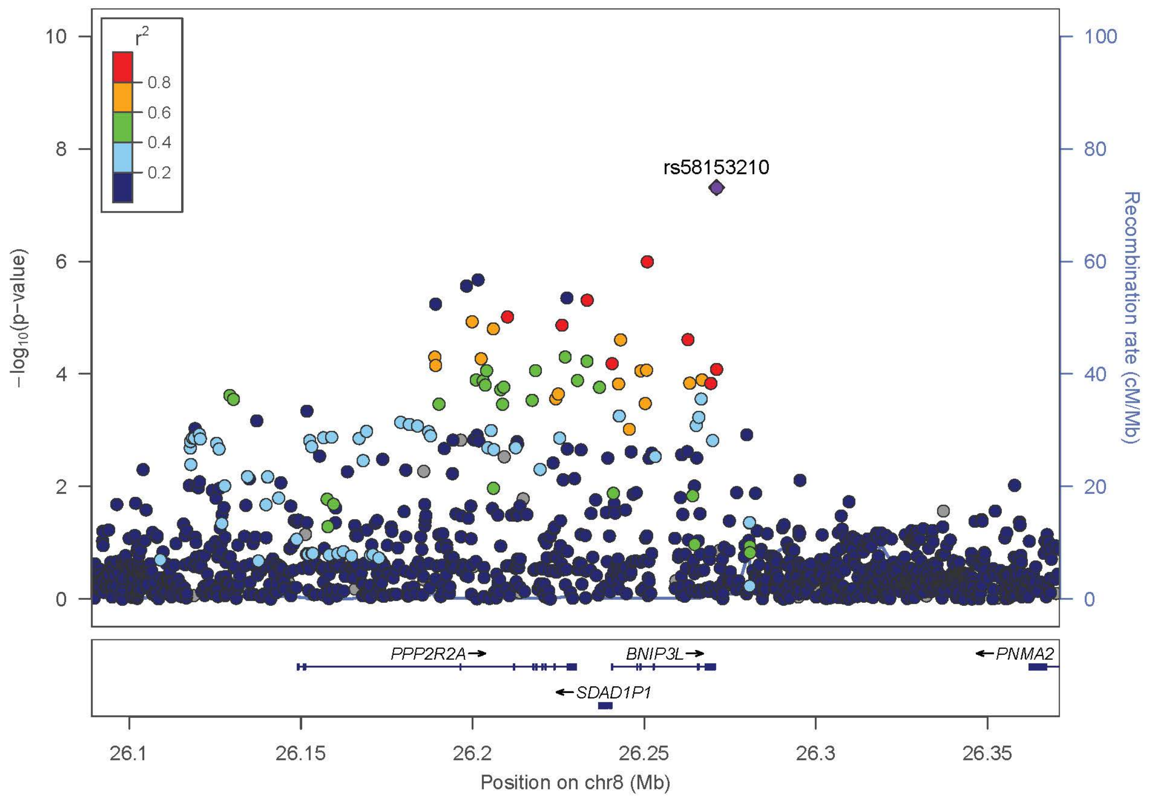

r2
0.8
0.6
0.4
0.2
rs58153210
−log10(p−value)
Recombination rate (cM/Mb)
PPP2R2A
BNIP3L
PNMA2
SDAD1P1
26.1
26.15
26.2
26.25
26.3
26.35
Position on chr8 (Mb)

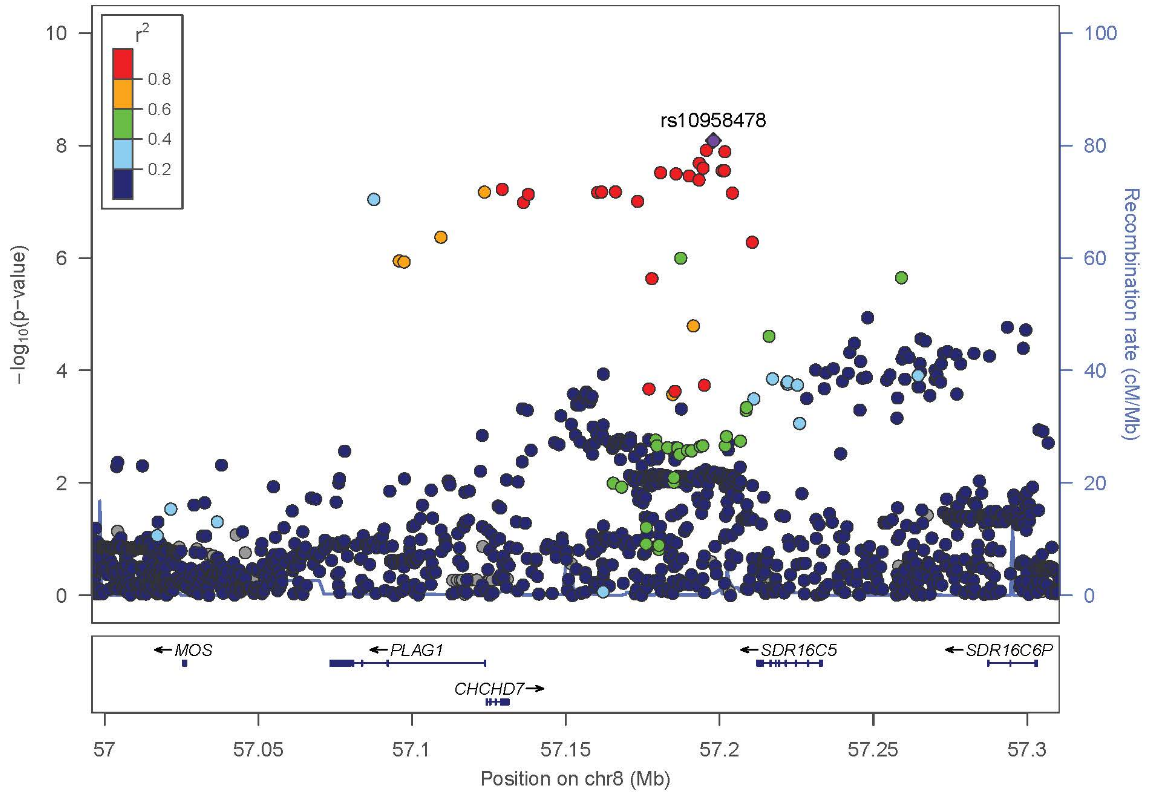
r2
0.8
0.6
0.4
0.2
rs10958478
−log10(p−value)
Recombination rate (cM/Mb)
MOS
PLAG1
CHCHD7
SDR16C5
SDR16C6P
Position on chr8 (Mb)

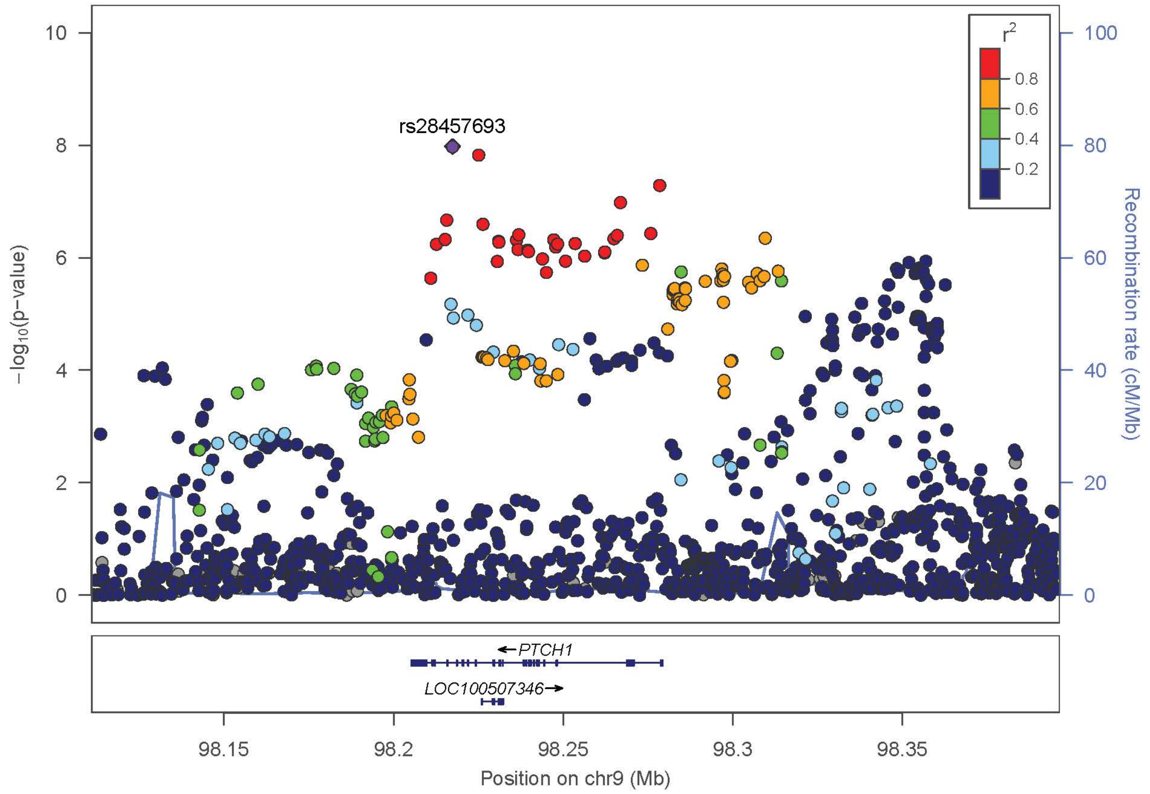
r2
0.8
0.6
0.4
0.2
rs28457693
−log10(p−value)
Recombination rate (cM/Mb)
PTCH1
LOC100507346
Position on chr9 (Mb)

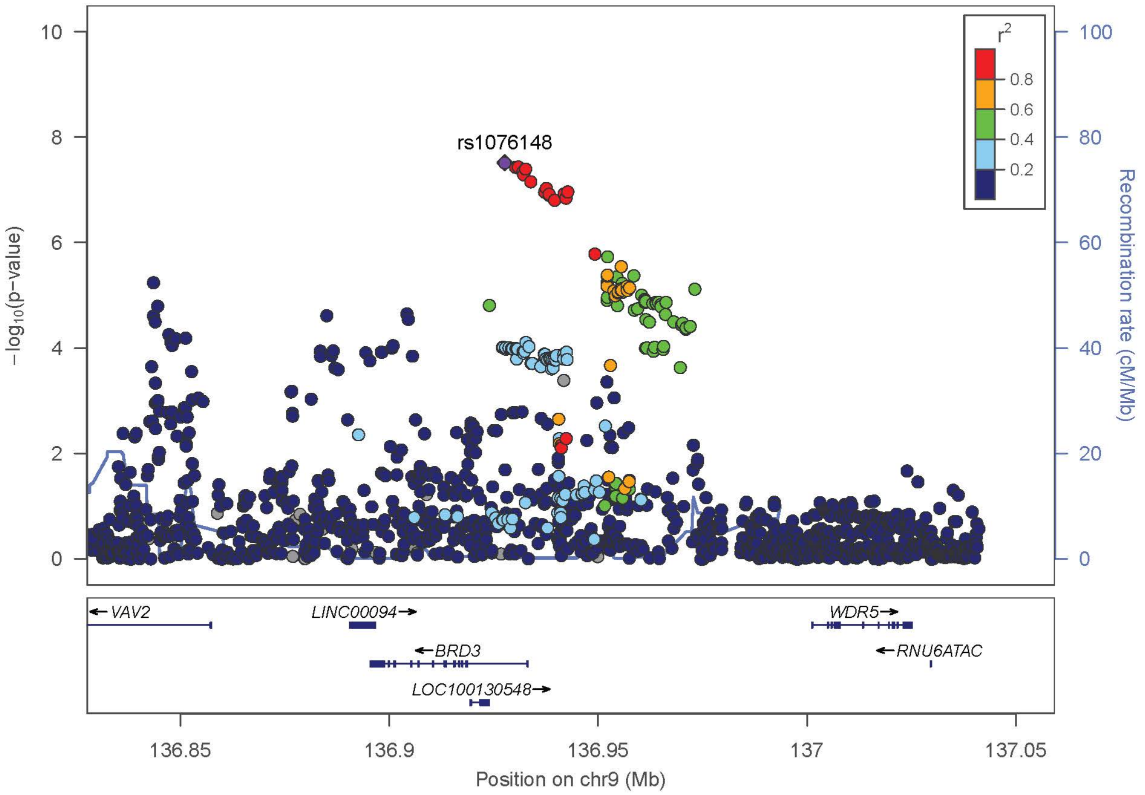

rs1076148
r2
0.8
0.6
0.4
0.2
−log10(p−value)
Recombination rate (cM/Mb)
VAV2
LINC00094
BRD3
LOC100130548
WDR5
RNU6ATAC
136.85
136.9
136.95
137
137.05
Position on chr9 (Mb)


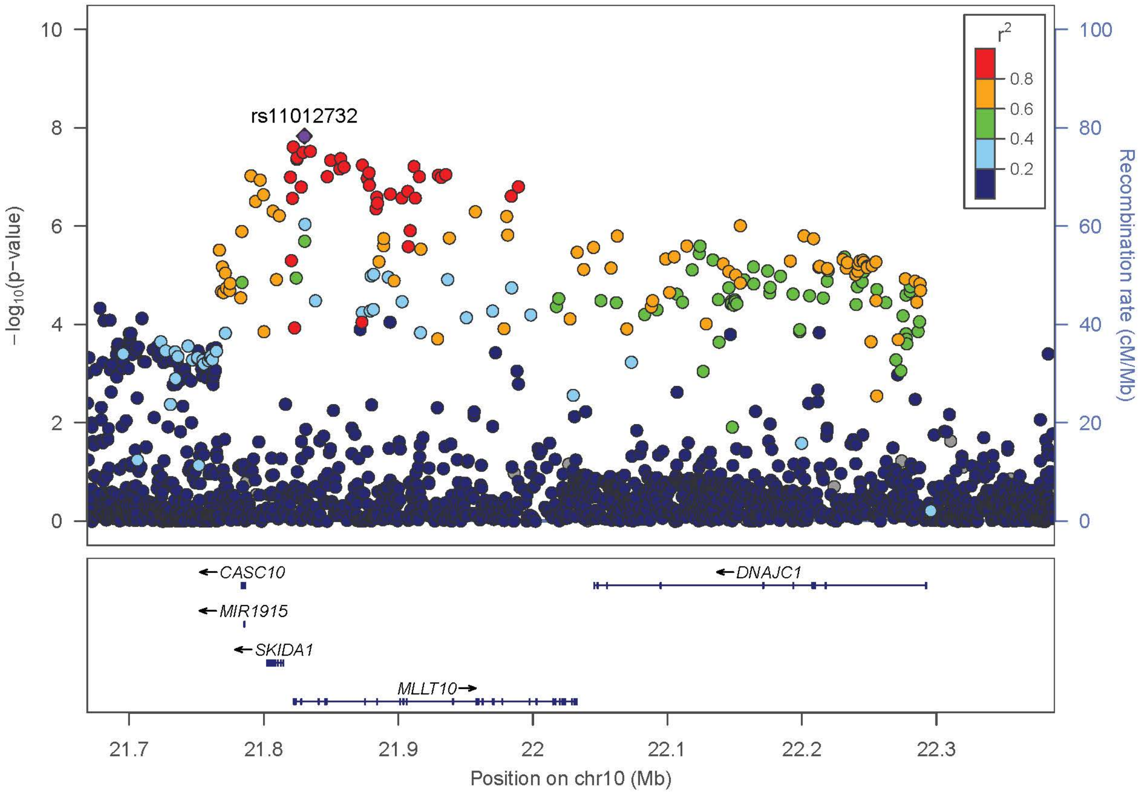

rs11012732
r2
0.8
0.6
0.4
0.2
−log10(p−value)
Recombination rate (cM/Mb)
CASC10
MIR1915
SKIDA1
MLLT10
DNAJC1
21.7
21.8
21.9
22
22.1
22.2
22.3
Position on chr10 (Mb)

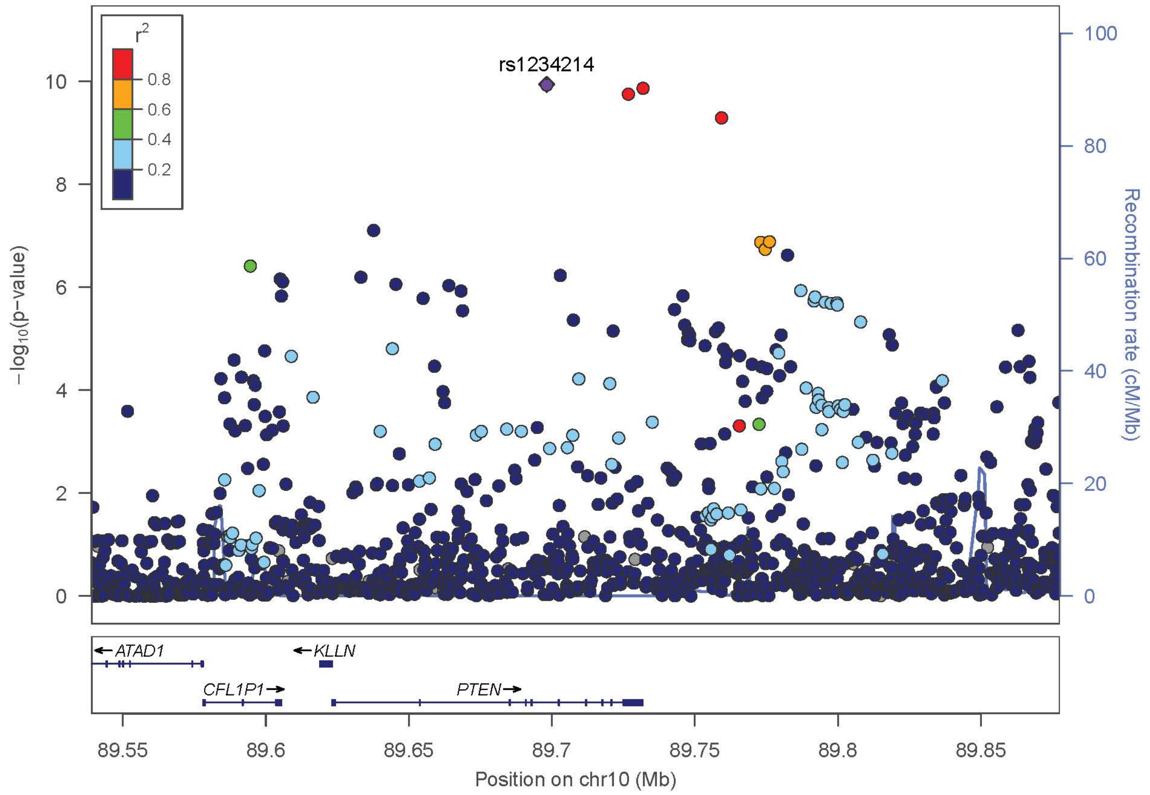
r2
0.8
0.6
0.4
0.2
rs1234214
−log10(p−value)
Recombination rate (cM/Mb)
ATAD1
KLLN
CFL1P1
PTEN
89.55
89.6
89.65
89.7
89.75
89.8
89.85
Position on chr10 (Mb)

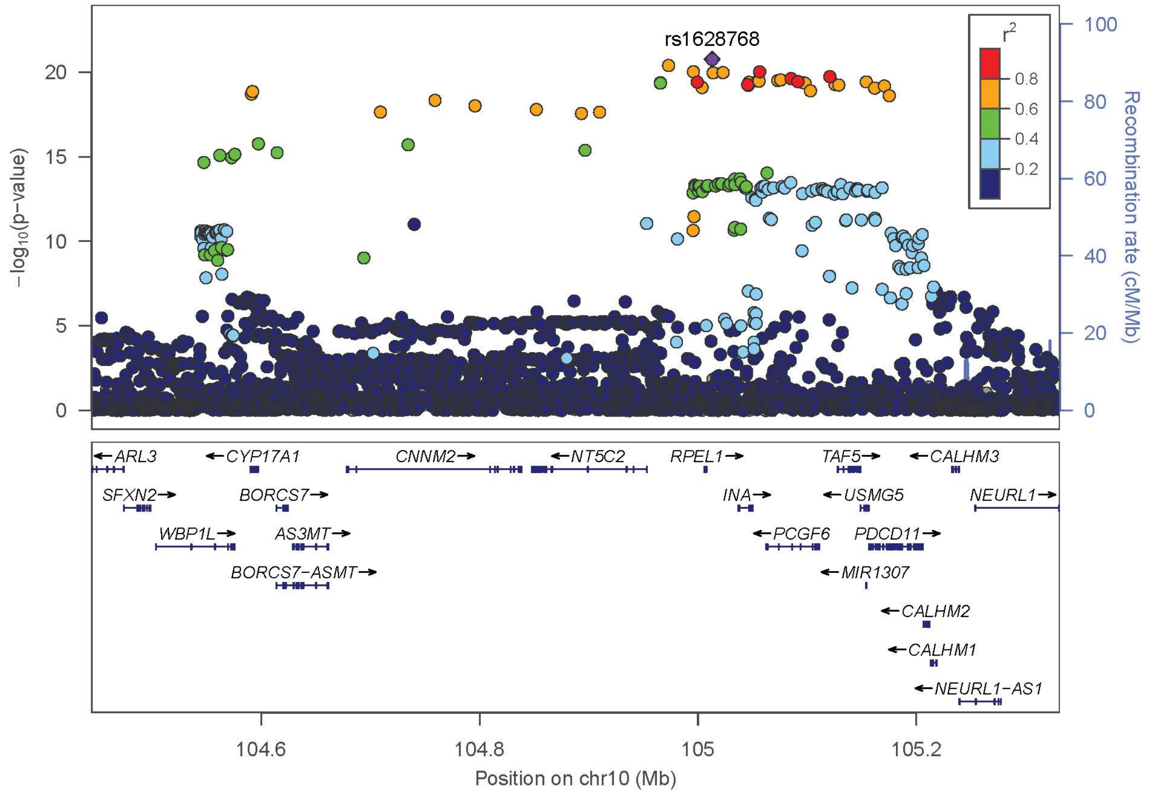
rs1628768
r2
0.8
0.6
0.4
0.2
−log10(p−value)
Recombination rate (cM/Mb)
ARL3
CYP17A1
CNNM2
NT5C2
RPEL1
TAF5
CALHM3
SFXN2
BORCS7
INA
USMG5
NEURL1
WBP1L
AS3MT
PCGF6
PDCD11
BORCS7−ASMT
MIR1307
CALHM2
CALHM1
NEURL1−AS1
104.6
104.8
105
105.2
Position on chr10 (Mb)

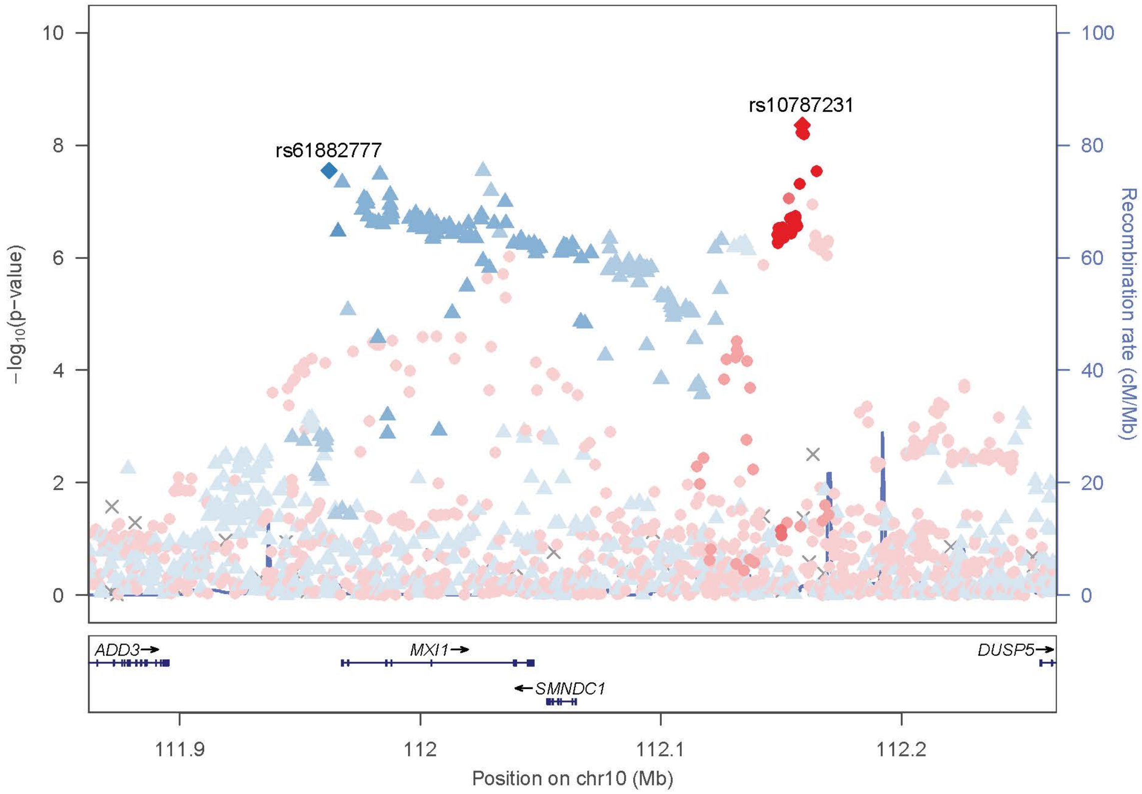
rs10787231
rs61882777
−log10(p−value)
Recombination rate (cM/Mb)
10
8
6
4
2
0
100
80
60
40
20
0
ADD3→
MXI1→
DUSP5→
←SMNDC1
111.9
112
112.1
112.2
Position on chr10 (Mb)

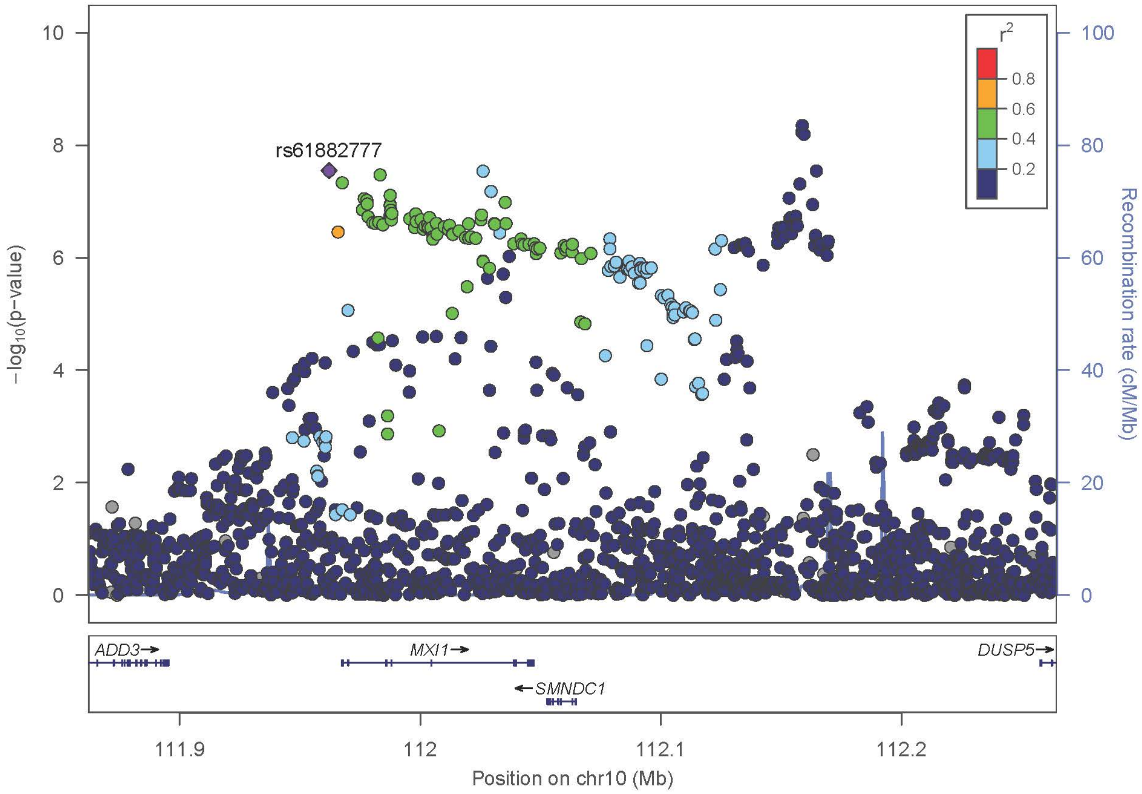
rs61882777
r2
0.8
0.6
0.4
0.2
−log10(p−value)
Recombination rate (cM/Mb)
10
8
6
4
2
0
100
80
60
40
20
0
ADD3→
MXI1→
DUSP5→
←SMNDC1
111.9
112
112.1
112.2
Position on chr10 (Mb)

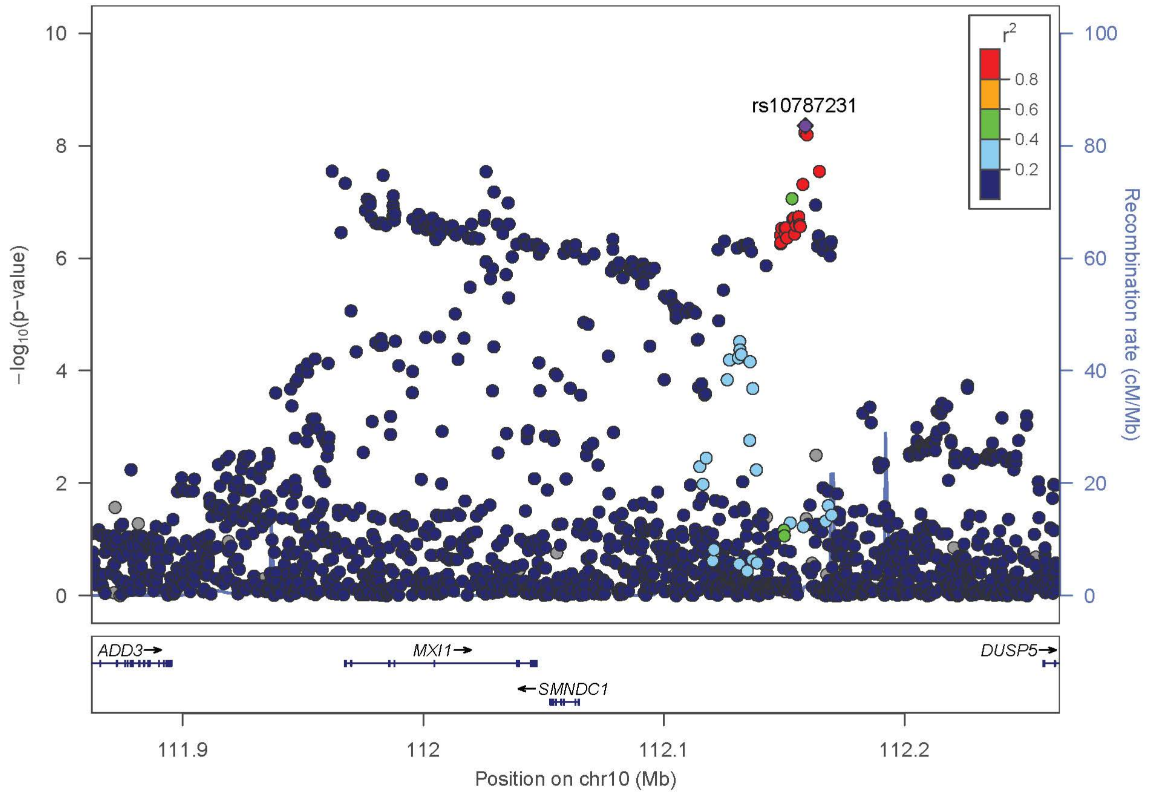
rs10787231
r2
0.8
0.6
0.4
0.2
−log10(p−value)
Recombination rate (cM/Mb)
ADD3
MXI1
SMNDC1
DUSP5
111.9
112
112.1
112.2
Position on chr10 (Mb)

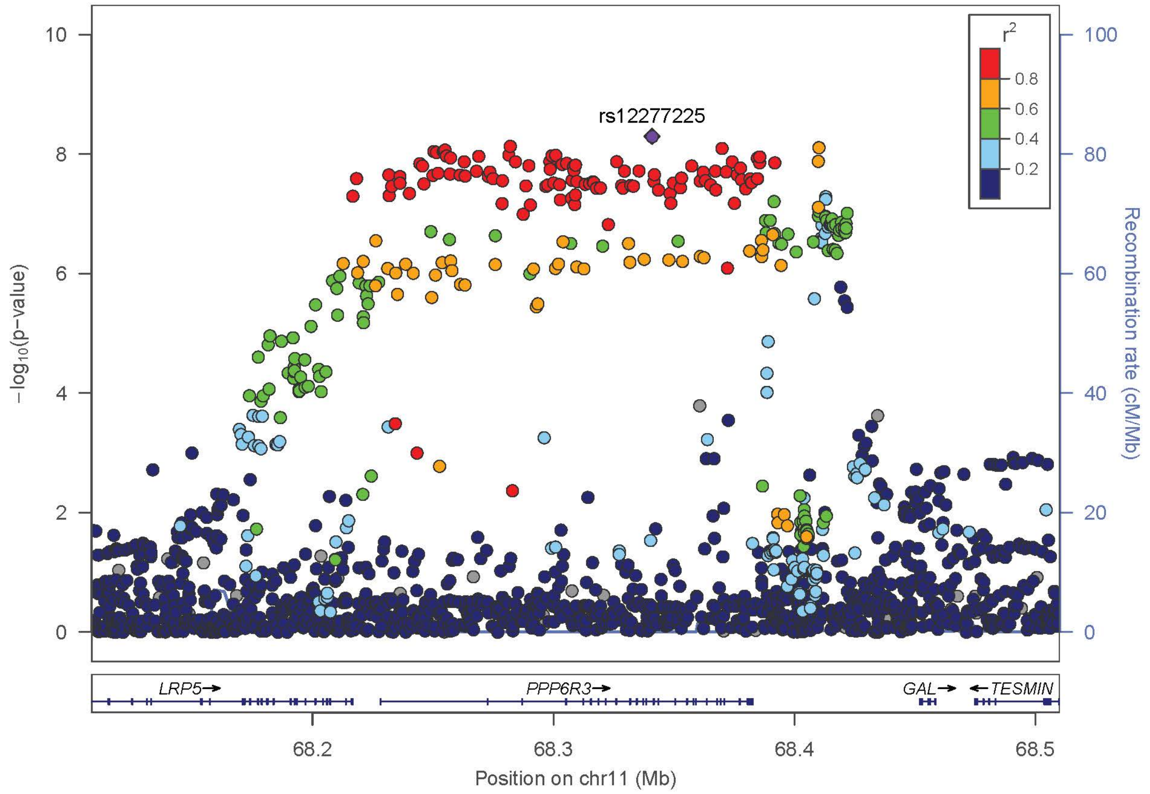
rs12277225
r2
0.8
0.6
0.4
0.2
−log10(p−value)
Recombination rate (cM/Mb)
LRP5
PPP6R3
GAL
TESMIN
68.2
68.3
68.4
68.5
Position on chr11 (Mb)

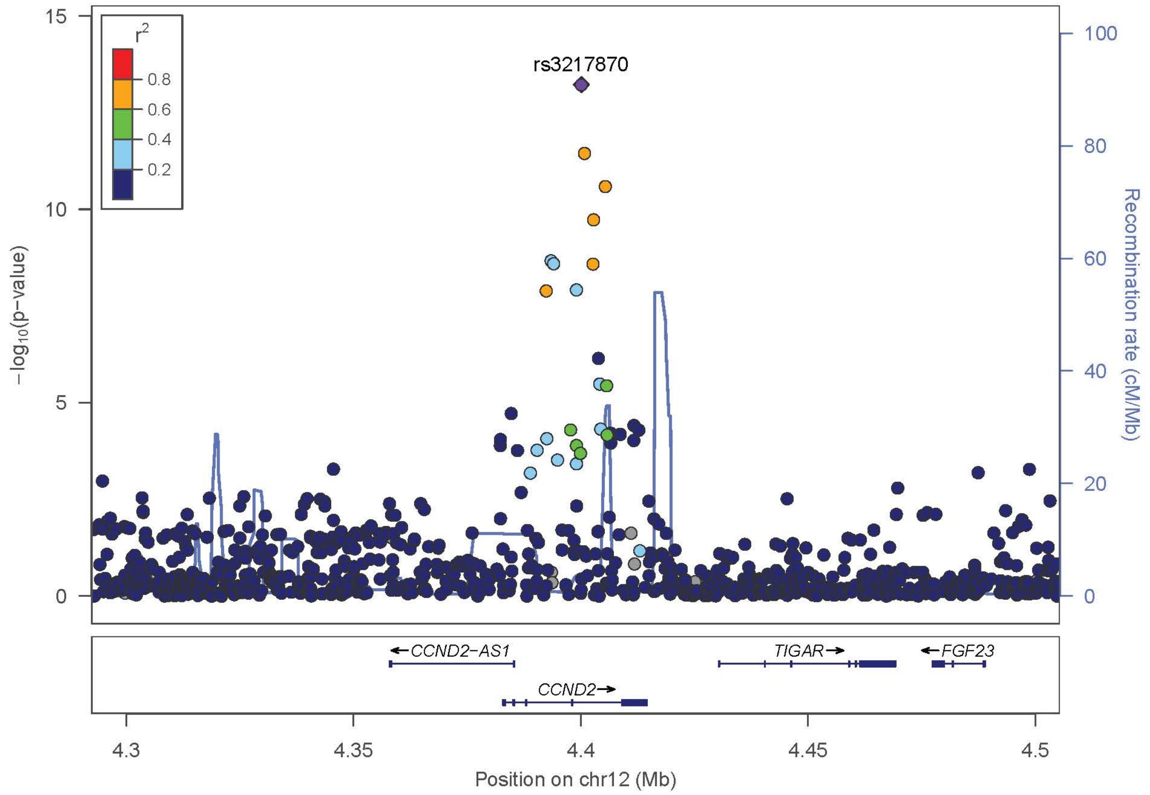

rs3217870
r2
0.8
0.6
0.4
0.2
−log10(p−value)
Recombination rate (cM/Mb)
CCND2−AS1
TIGAR
FGF23
CCND2
4.3
4.35
4.4
4.45
4.5
Position on chr12 (Mb)


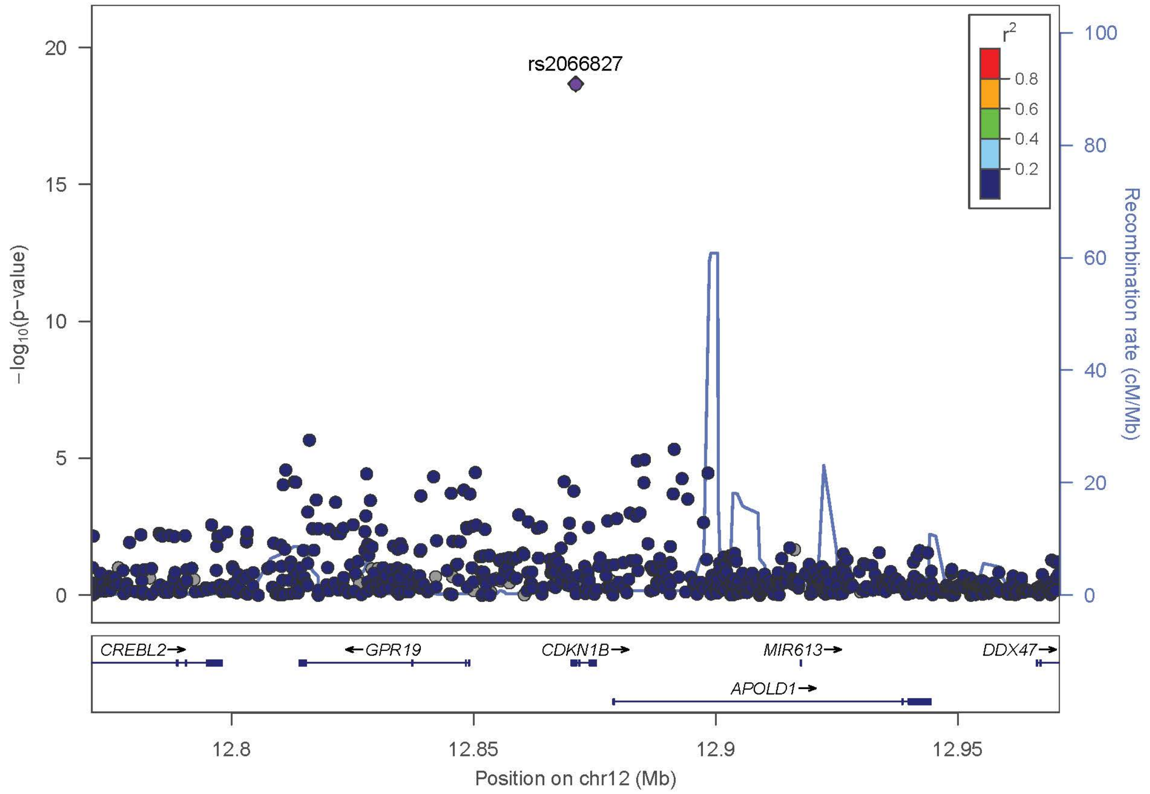

rs2066827
r2
0.8
0.6
0.4
0.2
−log10(p−value)
Recombination rate (cM/Mb)
CREBL2
GPR19
CDKN1B
MIR613
DDX47
APOLD1
12.8
12.85
12.9
12.95
Position on chr12 (Mb)

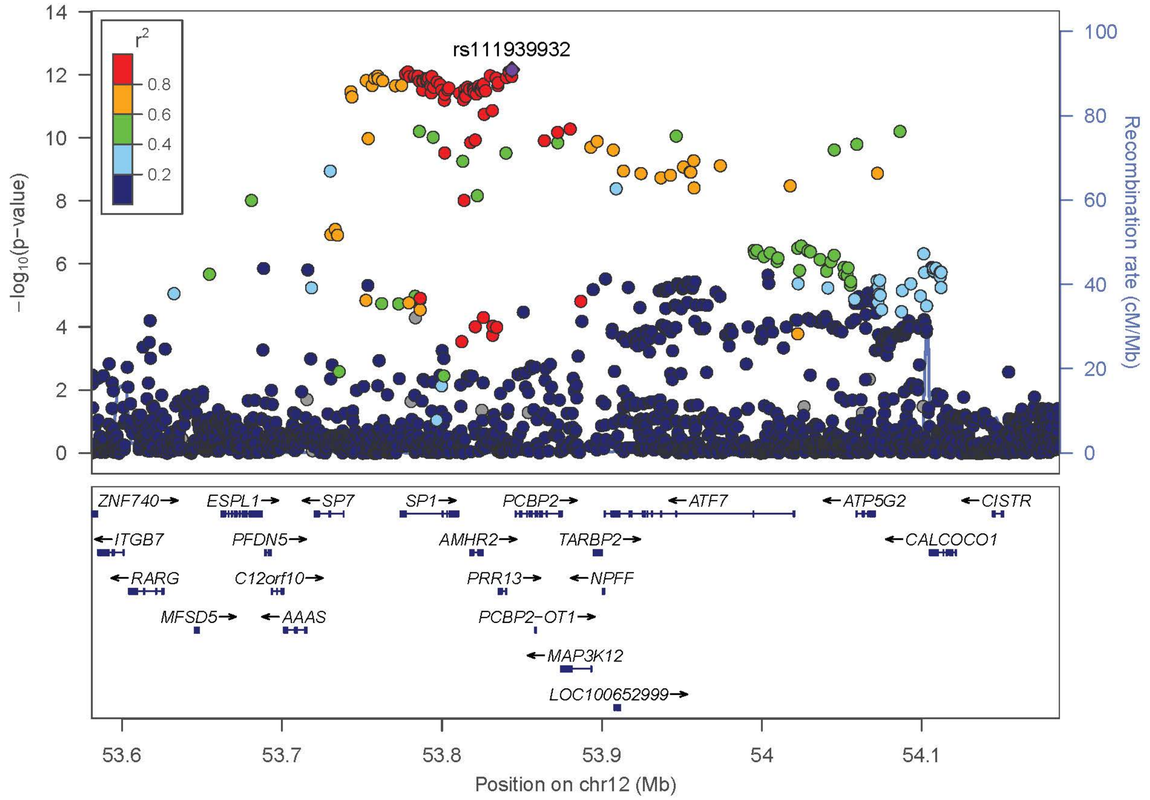
rs111939932
r2
0.8
0.6
0.4
0.2
−log10(p−value)
Recombination rate (cM/Mb)
ZNF740→
ESPL1→
←SP7
SP1→
PCBP2→
←ATF7
←ATP5G2
←CISTR
←ITGB7
PFDN5→
AMHR2→
TARBP2→
←CALCOCO1
←RARG
C12orf10→
PRR13→
←NPFF
MFSD5→
←AAAS
PCBP2−OT1→
←MAP3K12
LOC100652999→
Position on chr12 (Mb)

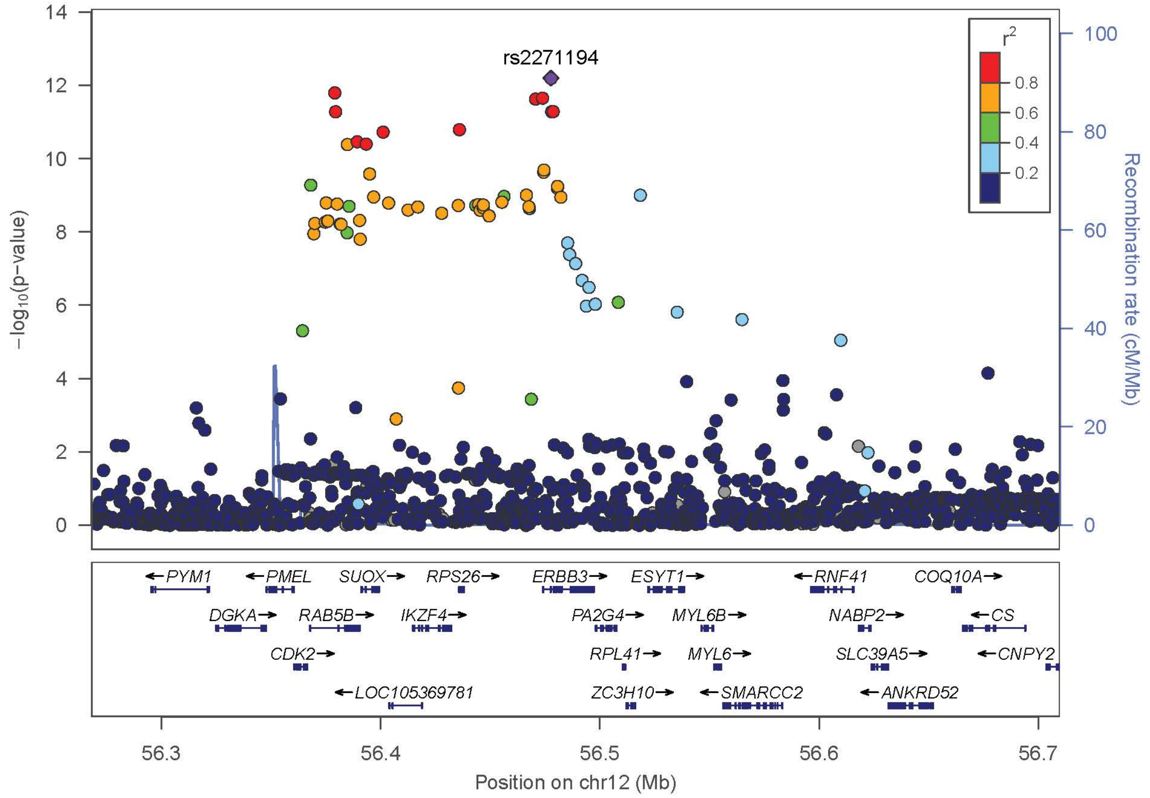
rs2271194
r2
0.8
0.6
0.4
0.2
−log10(p−value)
Recombination rate (cM/Mb)
←PYM1
←PMEL
SUOX→
RPS26→
ERBB3→
ESYT1→
←RNF41
COQ10A→
DGKA→
RAB5B→
IKZF4→
PA2G4→
MYL6B→
NABP2→
←CS
CDK2→
RPL41→
MYL6→
SLC39A5→
←CNPY2
←LOC105369781
ZC3H10→
←SMARCC2
←ANKRD52
Position on chr12 (Mb)

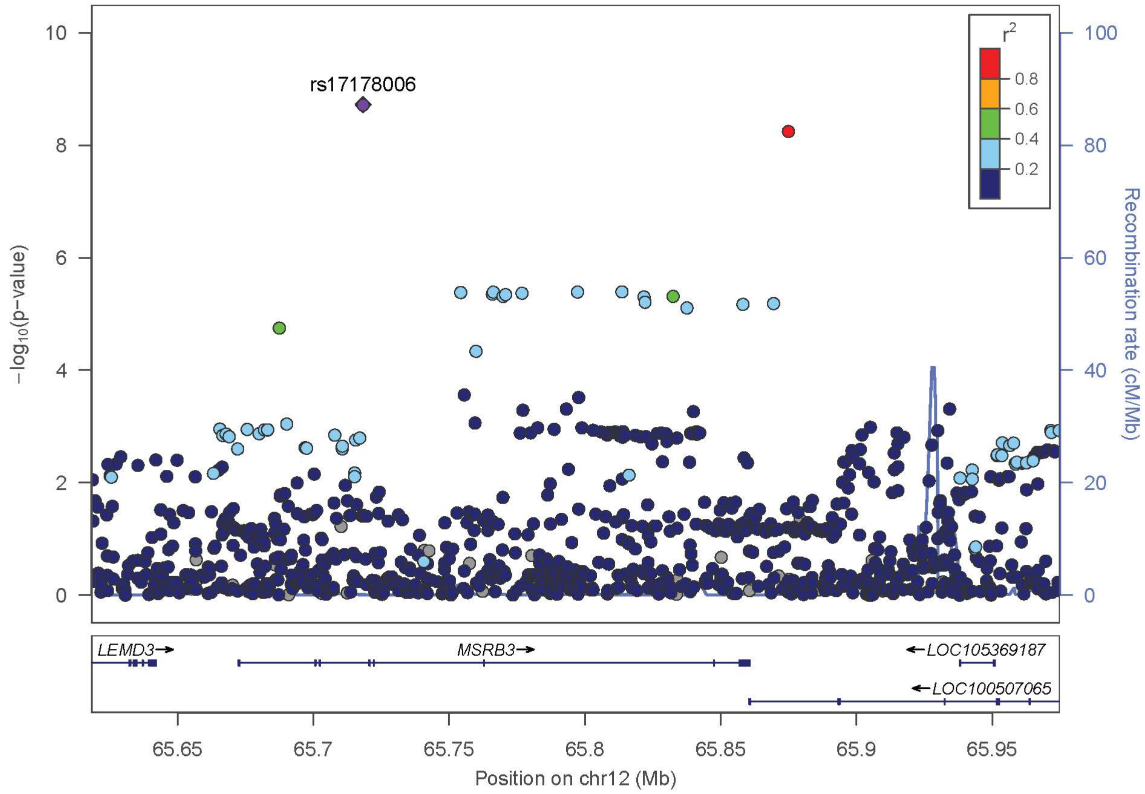

rs17178006
r2
0.8
0.6
0.4
0.2
−log10(p−value)
Recombination rate (cM/Mb)
LEMD3
MSRB3
LOC105369187
LOC100507065
65.65
65.7
65.75
65.8
65.85
65.9
65.95
Position on chr12 (Mb)


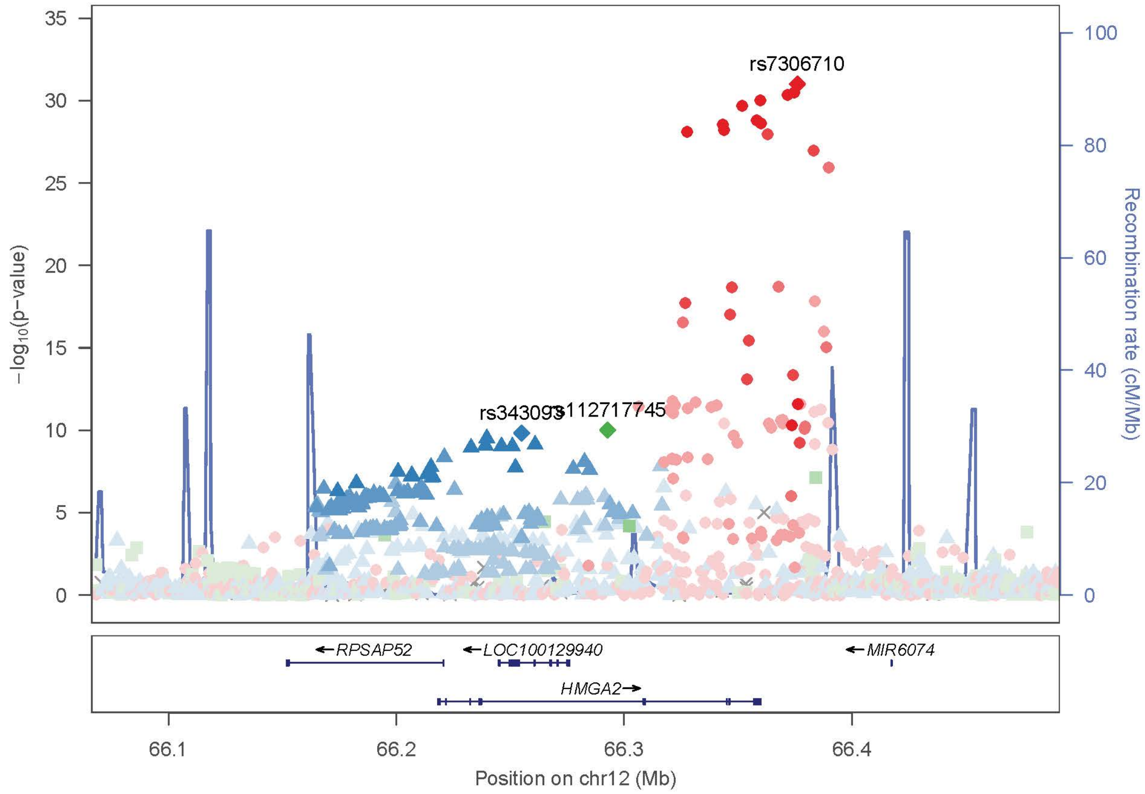

rs7306710
rs343093
rs112717745
−log10(p−value)
Recombination rate (cM/Mb)
RPSAP52
LOC100129940
MIR6074
HMGA2
66.1
66.2
66.3
66.4
Position on chr12 (Mb)

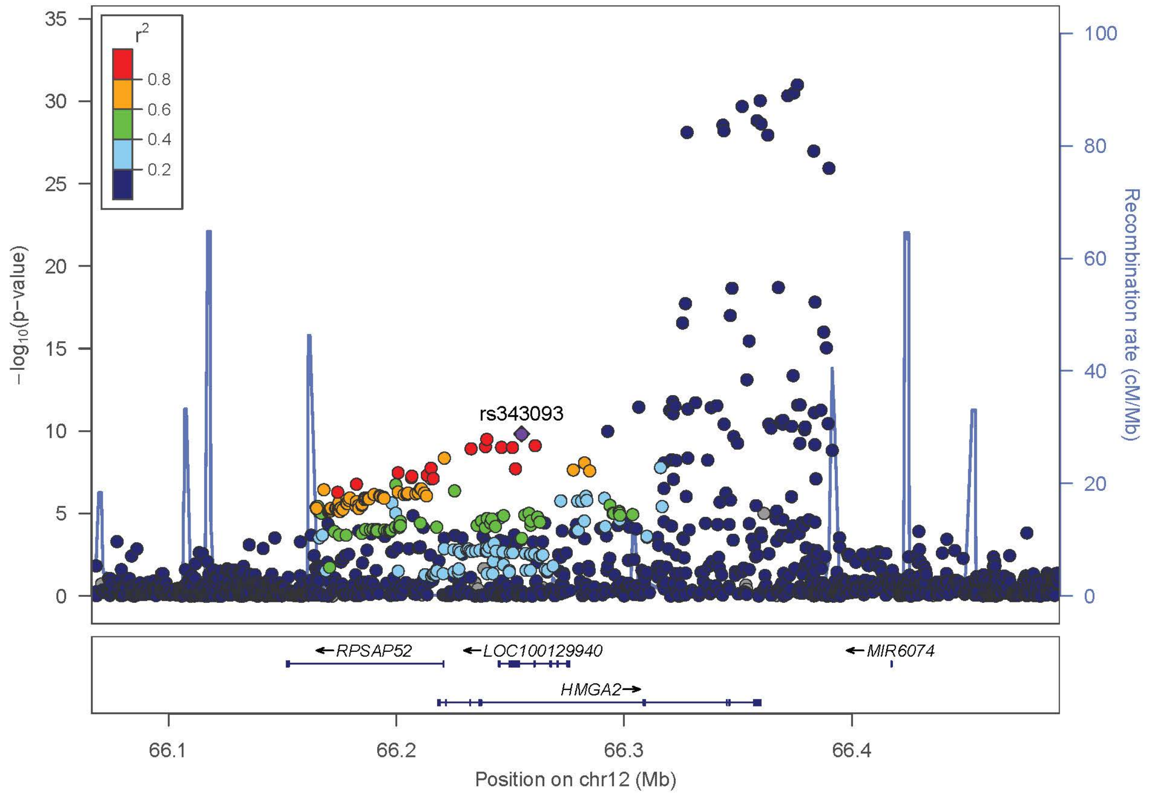
r2
0.8
0.6
0.4
0.2
rs343093
−log10(p−value)
Recombination rate (cM/Mb)
RPSAP52
LOC100129940
MIR6074
HMGA2
Position on chr12 (Mb)

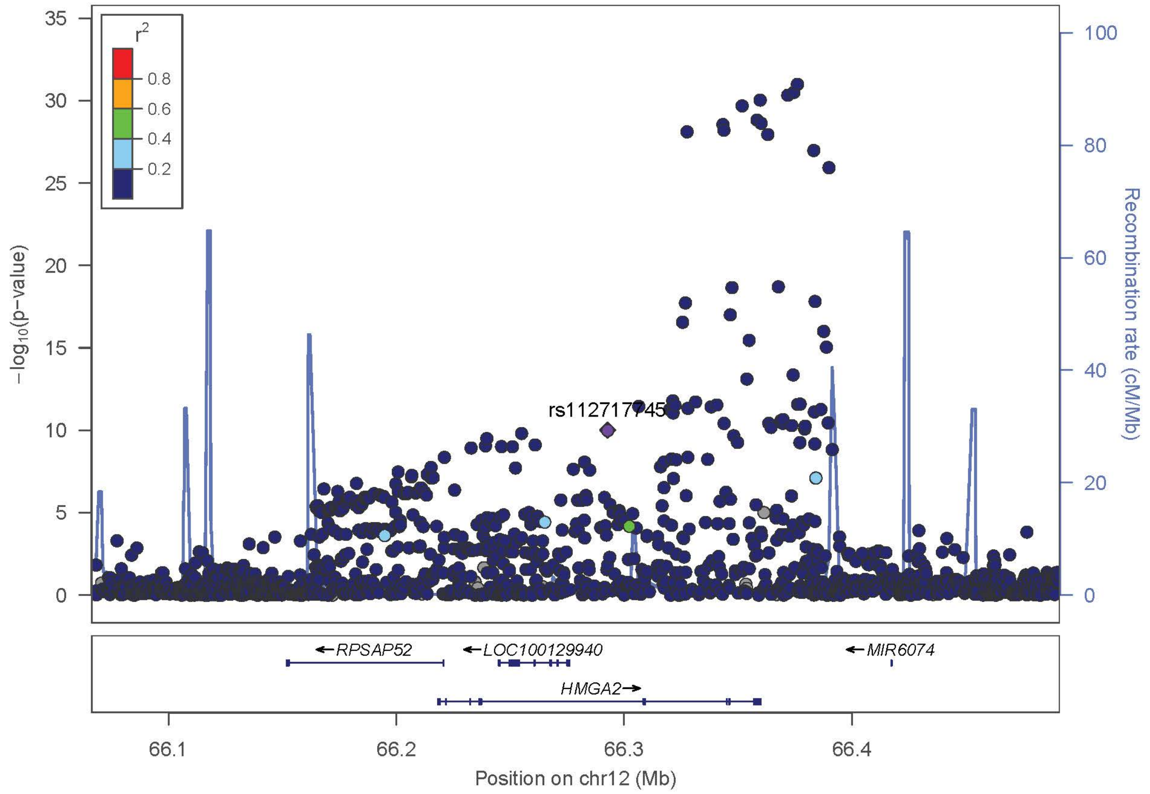
r2
0.8
0.6
0.4
0.2
−log10(p−value)
Recombination rate (cM/Mb)
RPSAP52
LOC100129940
MIR6074
HMGA2
Position on chr12 (Mb)

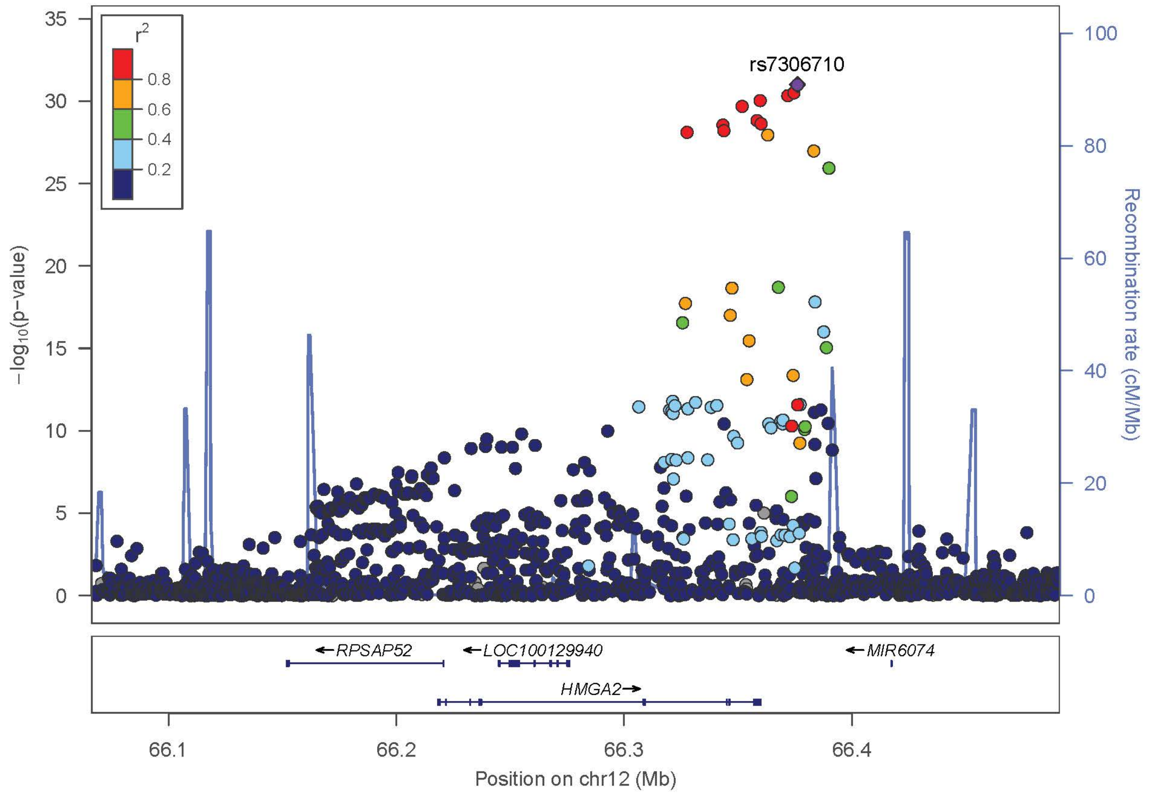

r2
0.8
0.6
0.4
0.2
rs7306710
−log10(p−value)
Recombination rate (cM/Mb)
RPSAP52
LOC100129940
MIR6074
HMGA2
Position on chr12 (Mb)


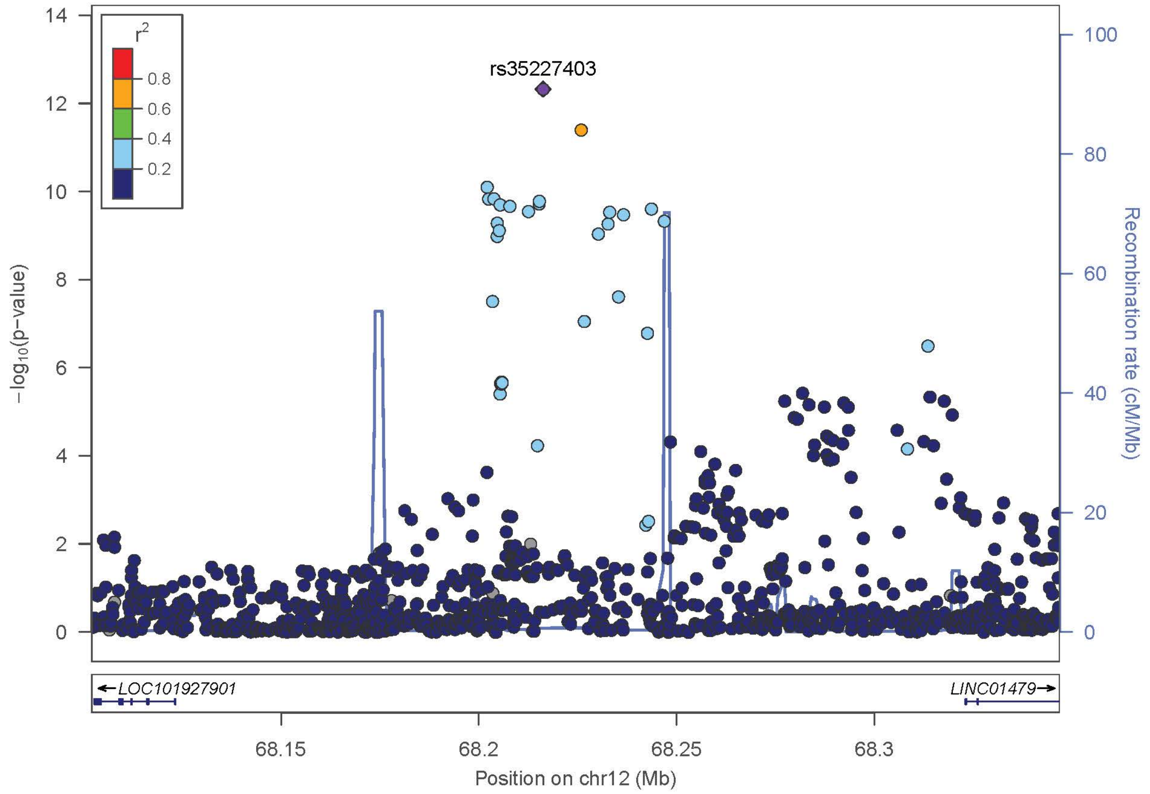

r2
0.8
0.6
0.4
0.2
rs35227403
−log10(p−value)
Recombination rate (cM/Mb)
LOC101927901
LINC01479
Position on chr12 (Mb)

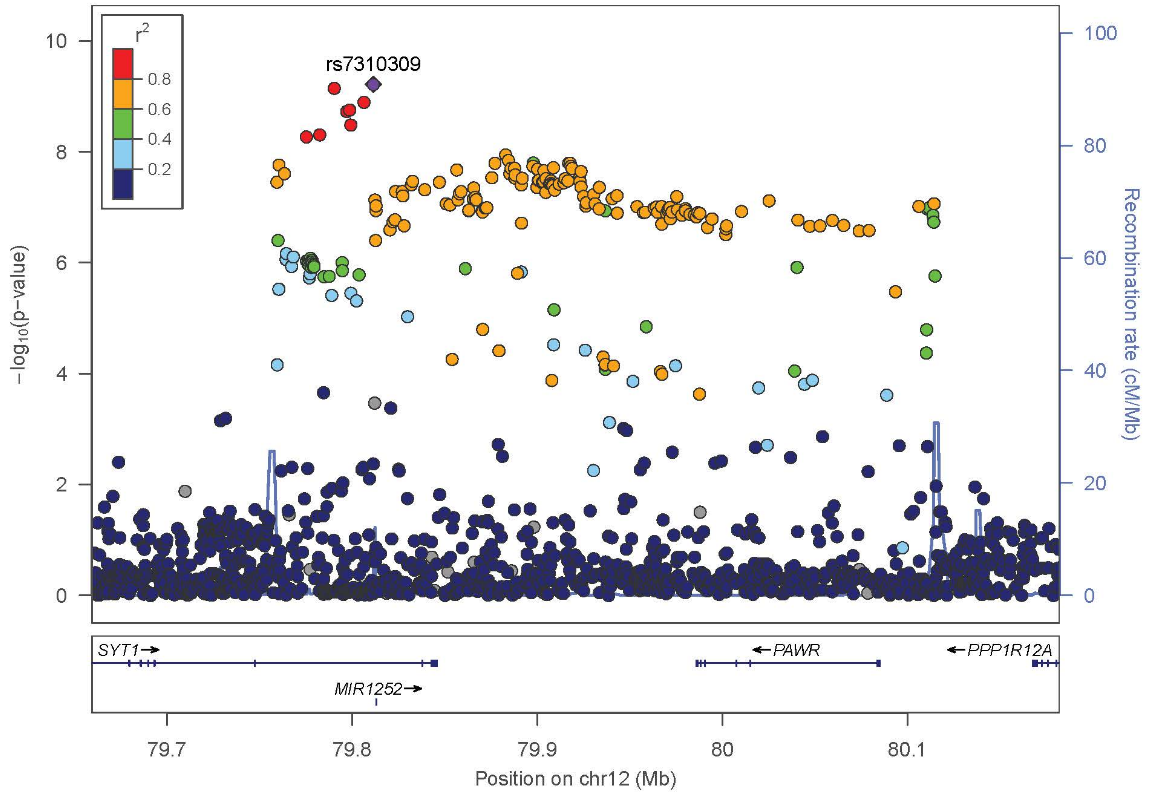
rs7310309
r²
0.8
0.6
0.4
0.2
−log10(p−value)
Recombination rate (cM/Mb)
SYT1→
MIR1252→
←PAWR
←PPP1R12A
79.7
79.8
79.9
80
80.1
Position on chr12 (Mb)

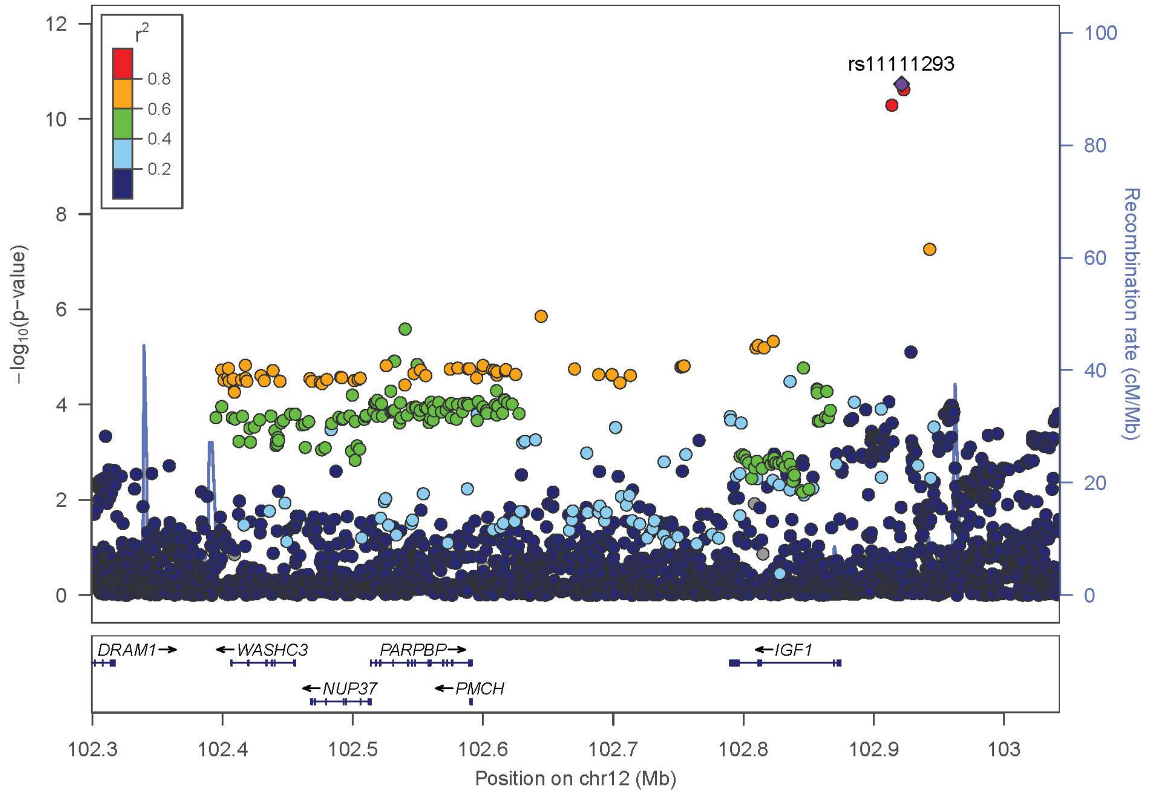
rs11111293
r²
0.8
0.6
0.4
0.2
−log10(p−value)
Recombination rate (cM/Mb)
DRAM1→
←WASHC3
PARPBP→
←NUP37
←PMCH
←IGF1
102.3
102.4
102.5
102.6
102.7
102.8
102.9
103
Position on chr12 (Mb)

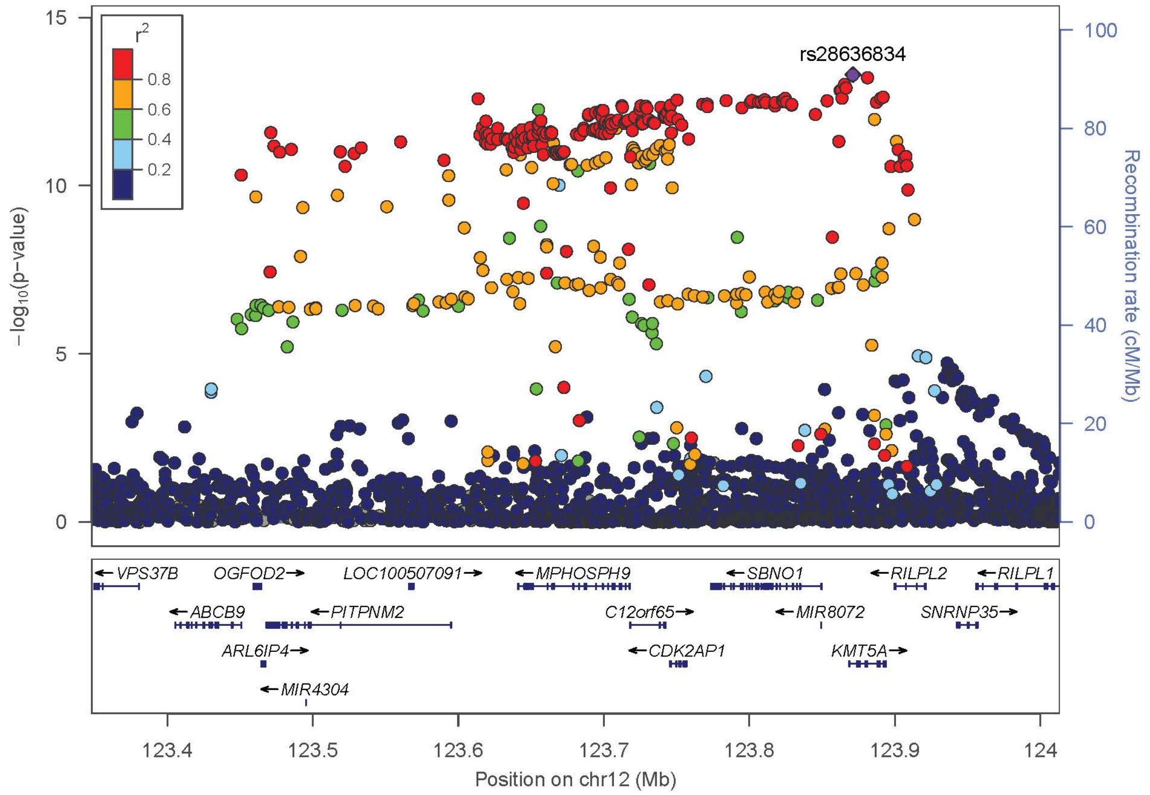

r2
0.8
0.6
0.4
0.2
rs28636834
−log10(p−value)
Recombination rate (cM/Mb)
VPS37B
OGFOD2
LOC100507091
MPHOSPH9
SBNO1
RILPL2
RILPL1
ABCB9
PITPNM2
C12orf65
MIR8072
SNRNP35
ARL6IP4
CDK2AP1
KMT5A
MIR4304
Position on chr12 (Mb)


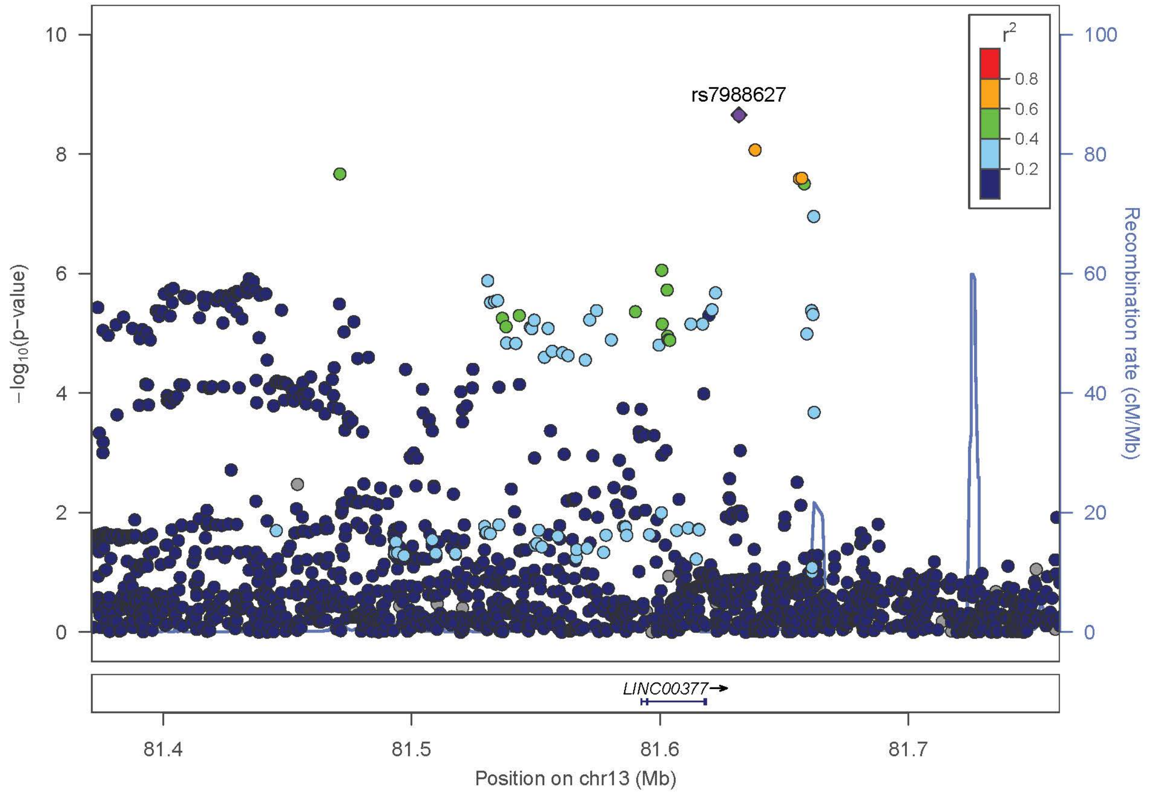

r2
0.8
0.6
0.4
0.2
rs7988627
−log10(p−value)
Recombination rate (cM/Mb)
LINC00377
Position on chr13 (Mb)

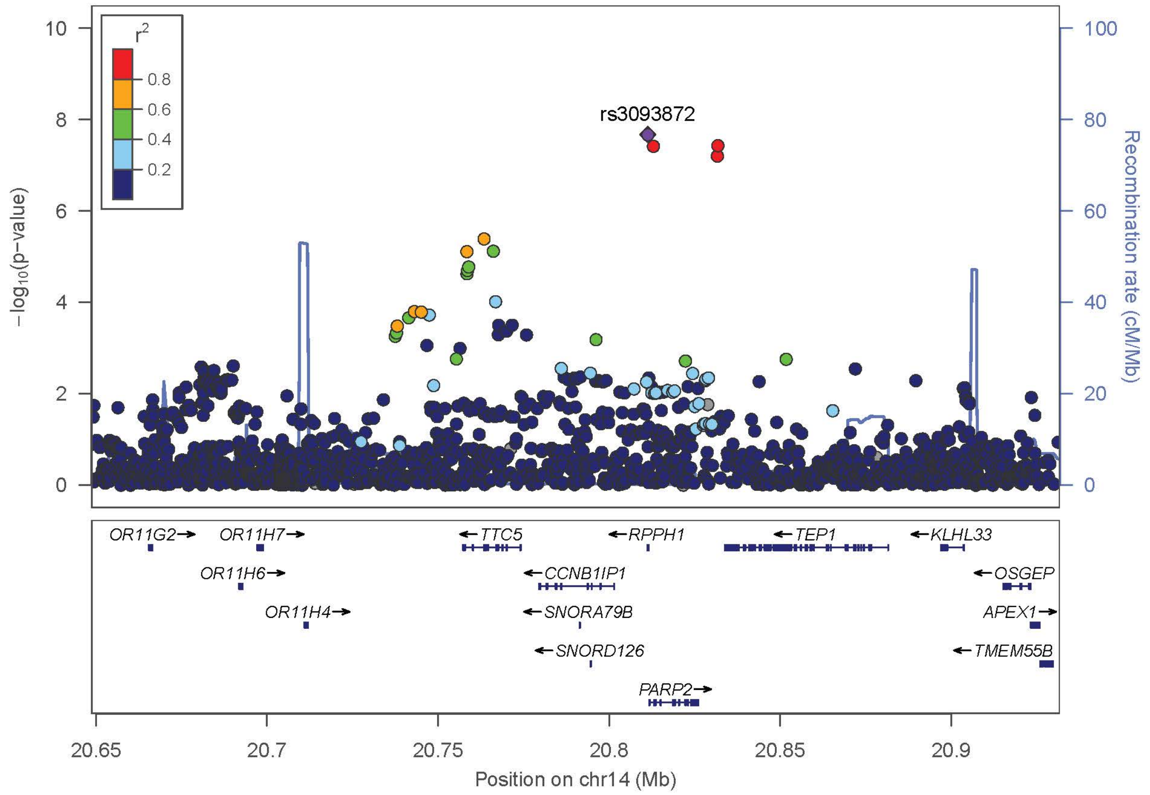
rs3093872
r2
0.8
0.6
0.4
0.2
−log10(p−value)
Recombination rate (cM/Mb)
OR11G2→
OR11H7→
OR11H6→
OR11H4→
←TTC5
←CCNB1IP1
←SNORA79B
←SNORD126
←RPPH1
PARP2→
←TEP1
←KLHL33
←OSGEP
APEX1→
←TMEM55B
20.65
20.7
20.75
20.8
20.85
20.9
Position on chr14 (Mb)

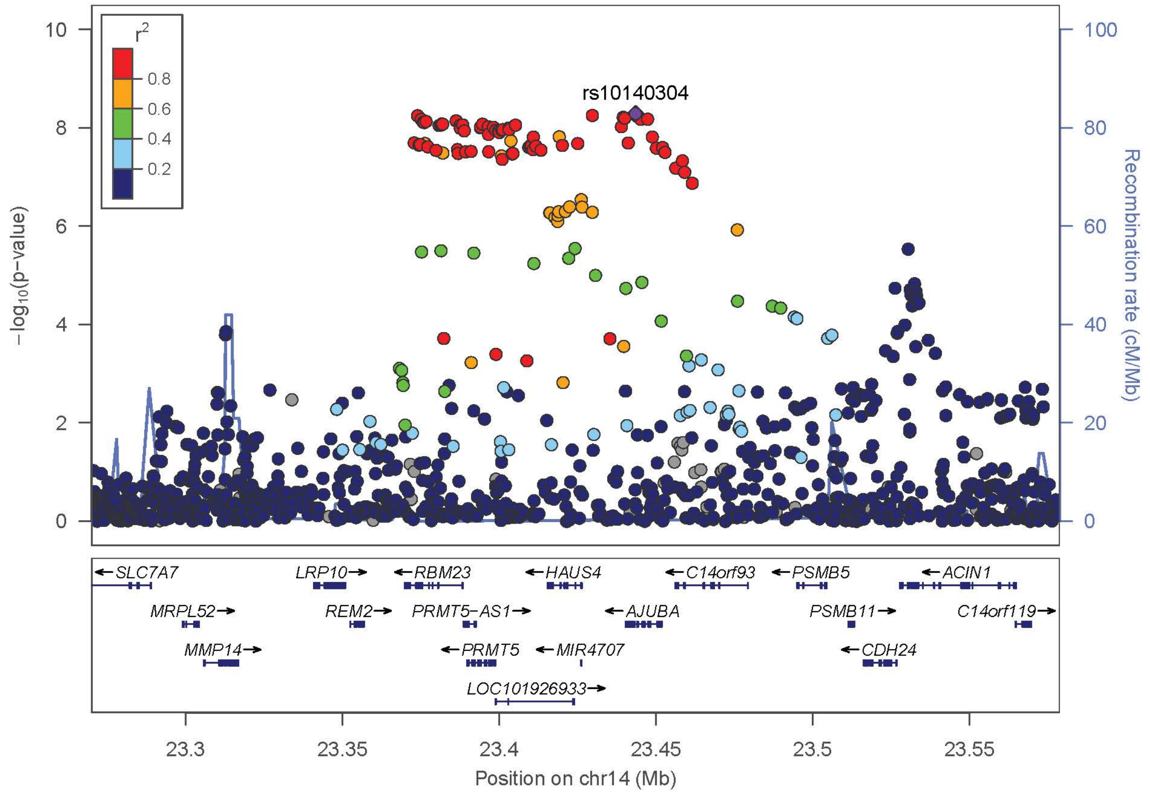
rs10140304
r2
0.8
0.6
0.4
0.2
−log10(p−value)
Recombination rate (cM/Mb)
←SLC7A7
MRPL52→
MMP14→
LRP10→
REM2→
←RBM23
PRMT5−AS1→
←PRMT5
LOC101926933→
←HAUS4
←MIR4707
←AJUBA
←C14orf93
←PSMB5
PSMB11→
←CDH24
←ACIN1
C14orf119→
23.3
23.35
23.4
23.45
23.5
23.55
Position on chr14 (Mb)

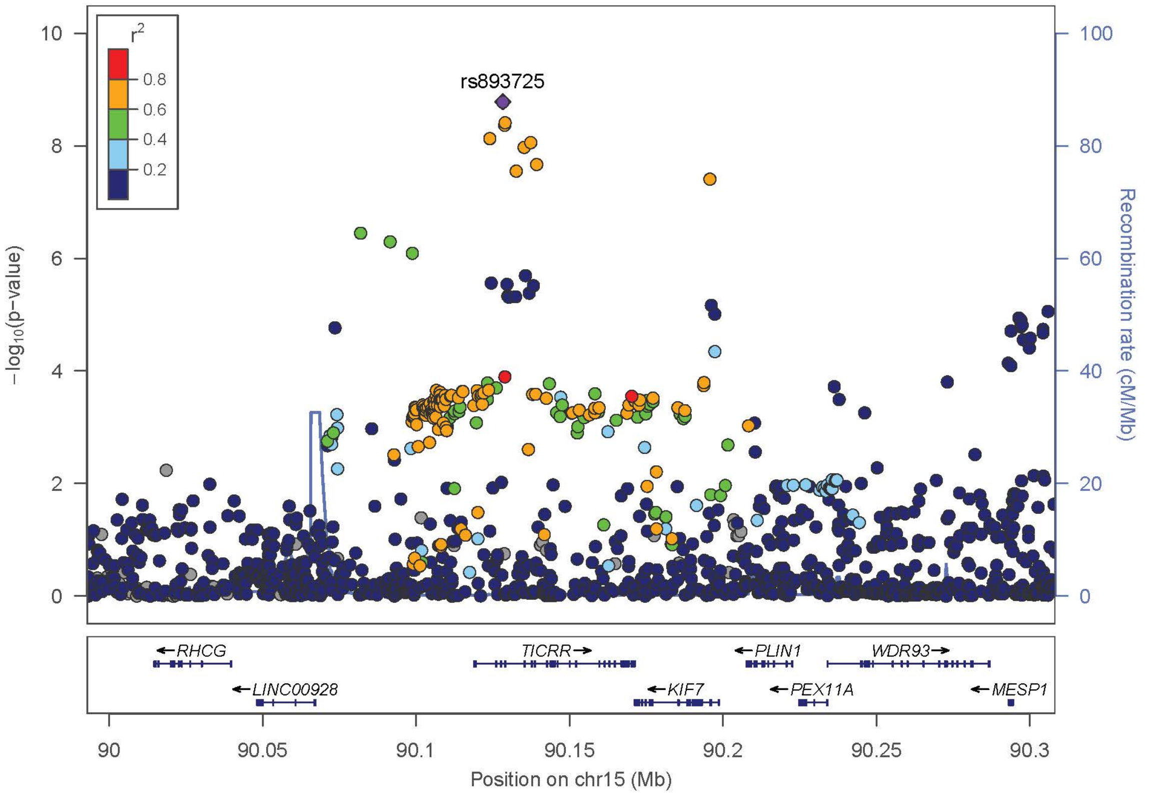

rs893725
r2
0.8
0.6
0.4
0.2
−log10(p−value)
Recombination rate (cM/Mb)
RHCG
LINC00928
TICRR
KIF7
PLIN1
PEX11A
WDR93
MESP1
Position on chr15 (Mb)


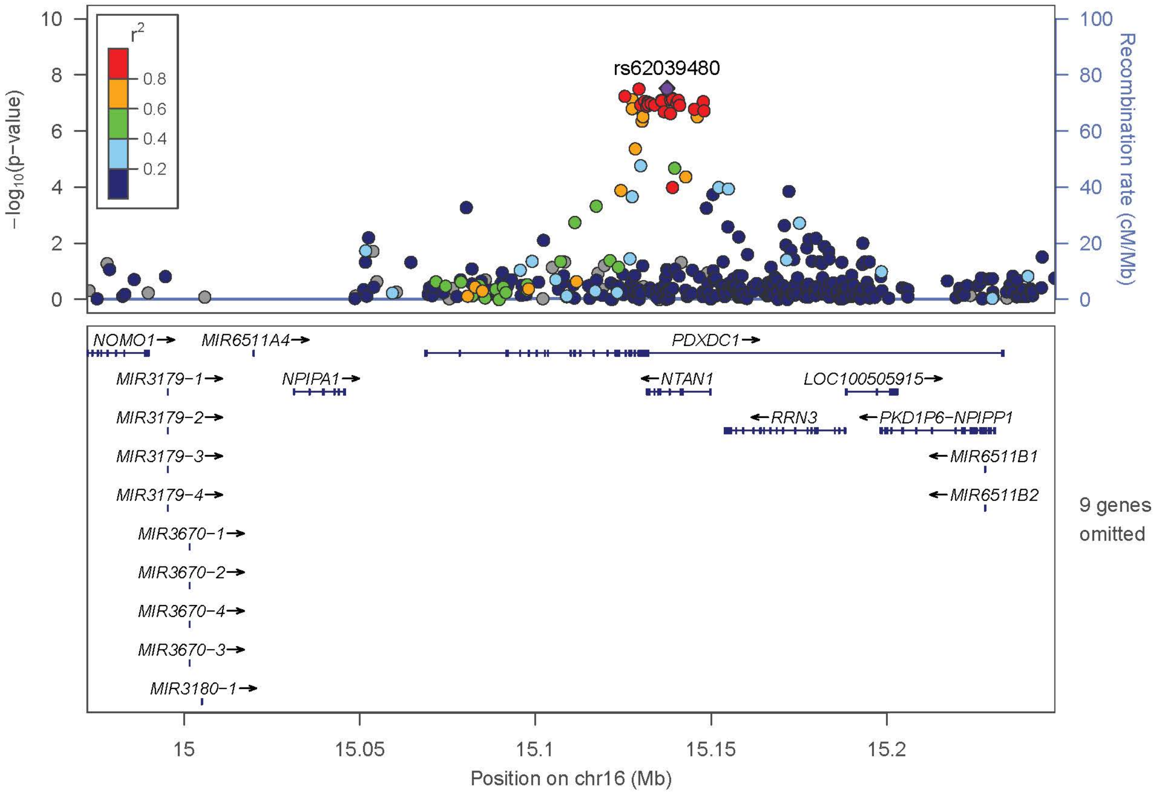

rs62039480
r2
0.8
0.6
0.4
0.2
−log10(p−value)
Recombination rate (cM/Mb)
NOMO1
MIR6511A4
PDXDC1
MIR3179−1
NPIPA1
NTAN1
LOC100505915
MIR3179−2
RRN3
PKD1P6−NPIPP1
MIR3179−3
MIR6511B1
MIR3179−4
MIR6511B2
MIR3670−1
MIR3670−2
MIR3670−4
MIR3670−3
MIR3180−1
9 genes omitted
Position on chr16 (Mb)

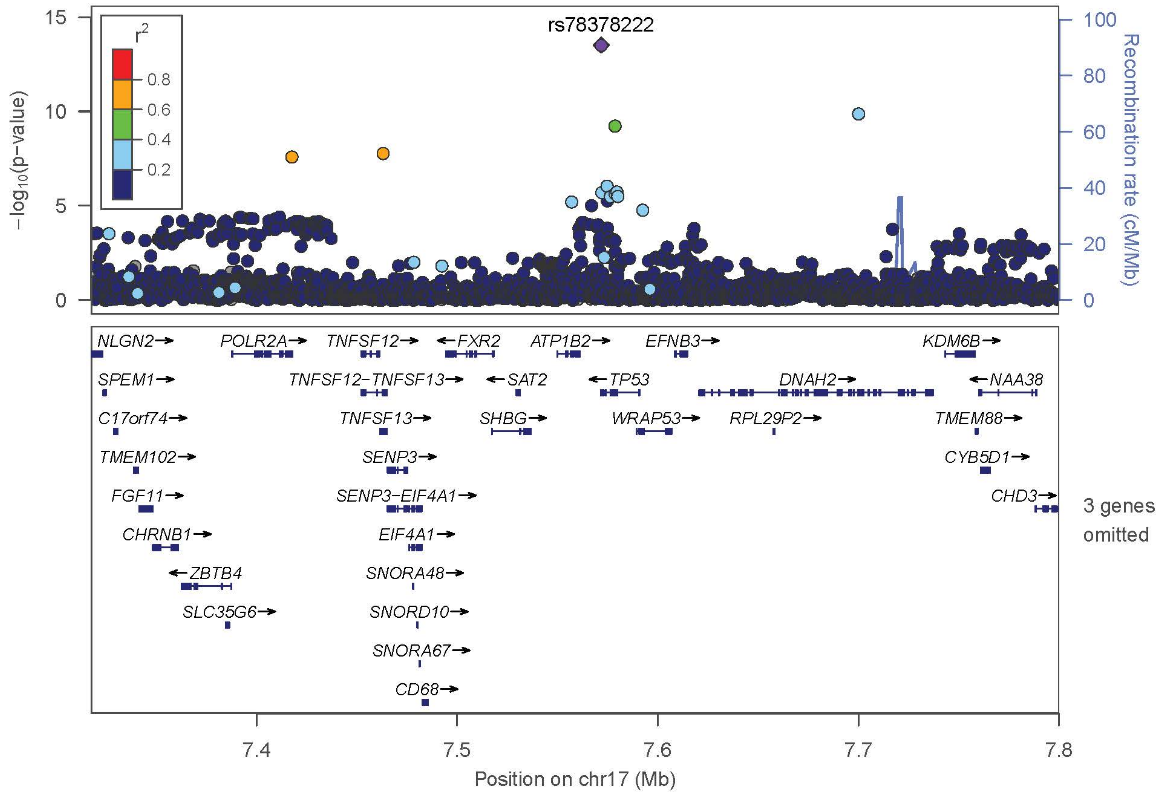
rs78378222
r2
0.8
0.6
0.4
0.2
−log10(p−value)
Recombination rate (cM/Mb)
NLGN2→
POLR2A→
TNFSF12→
←FXR2
ATP1B2→
EFNB3→
KDM6B→
SPEM1→
TNFSF12−TNFSF13→
←SAT2
←TP53
DNAH2→
←NAA38
C17orf74→
TNFSF13→
SHBG→
WRAP53→
RPL29P2→
TMEM88→
TMEM102→
SENP3→
CYB5D1→
FGF11→
SENP3−EIF4A1→
CHD3→
3 genes omitted
CHRNB1→
EIF4A1→
←ZBTB4
SNORA48→
SLC35G6→
SNORD10→
SNORA67→
CD68→
7.4
7.5
7.6
7.7
7.8
Position on chr17 (Mb)

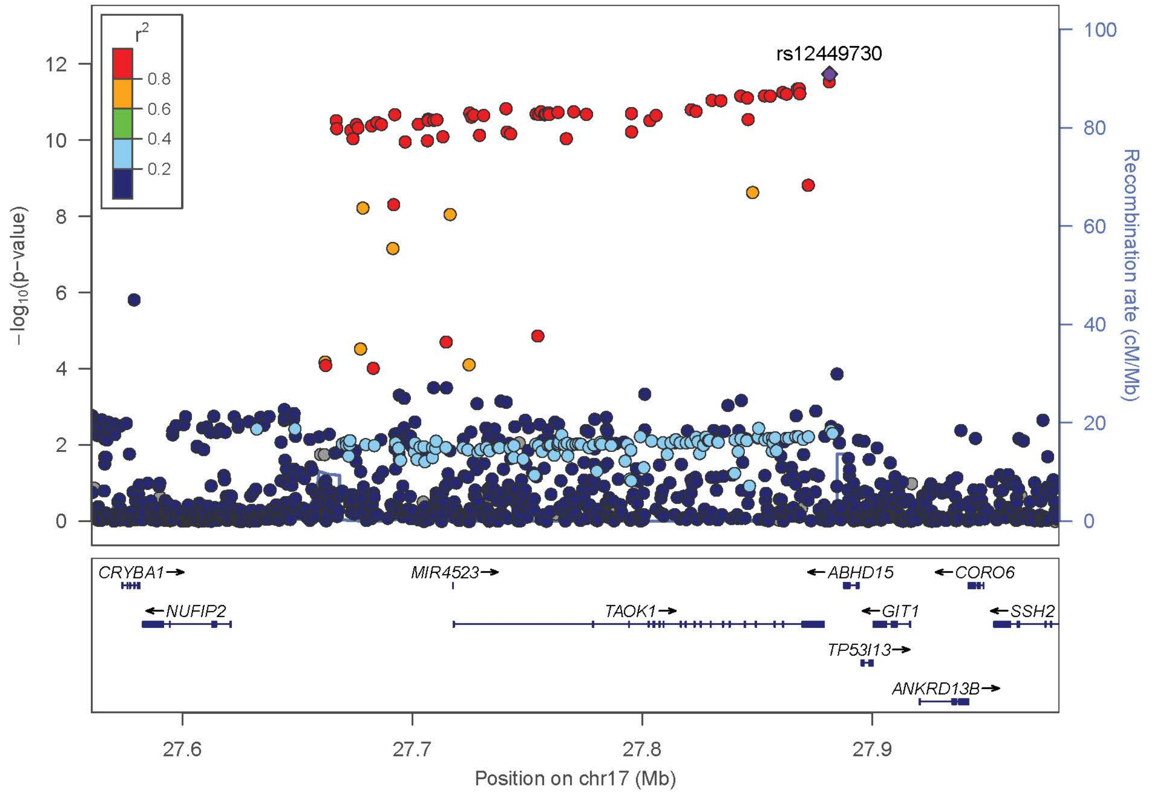
rs12449730
r2
0.8
0.6
0.4
0.2
−log10(p−value)
Recombination rate (cM/Mb)
CRYBA1→
MIR4523→
←ABHD15
←CORO6
←NUFIP2
TAOK1→
←GIT1
←SSH2
TP53I13→
ANKRD13B→
27.6
27.7
27.8
27.9
Position on chr17 (Mb)

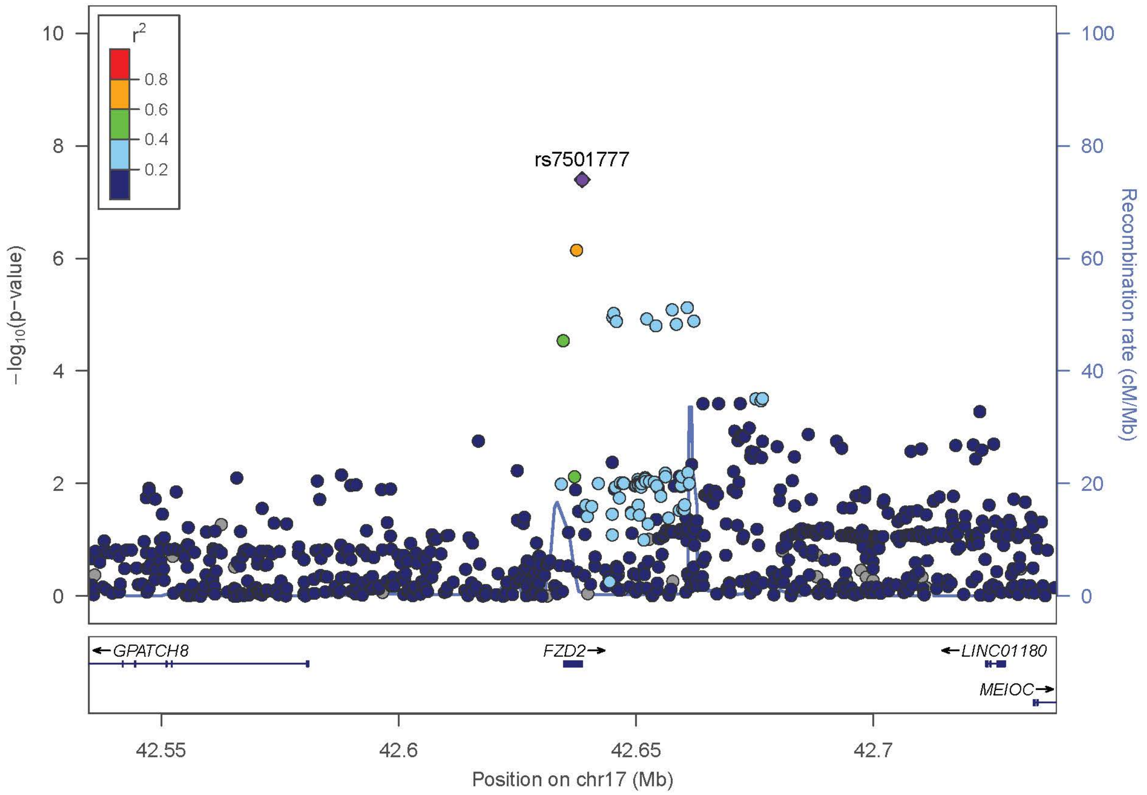
r2
0.8
0.6
0.4
0.2
rs7501777
−log10(p−value)
Recombination rate (cM/Mb)
GPATCH8
FZD2
LINC01180
MEIOC
42.55
42.6
42.65
42.7
Position on chr17 (Mb)

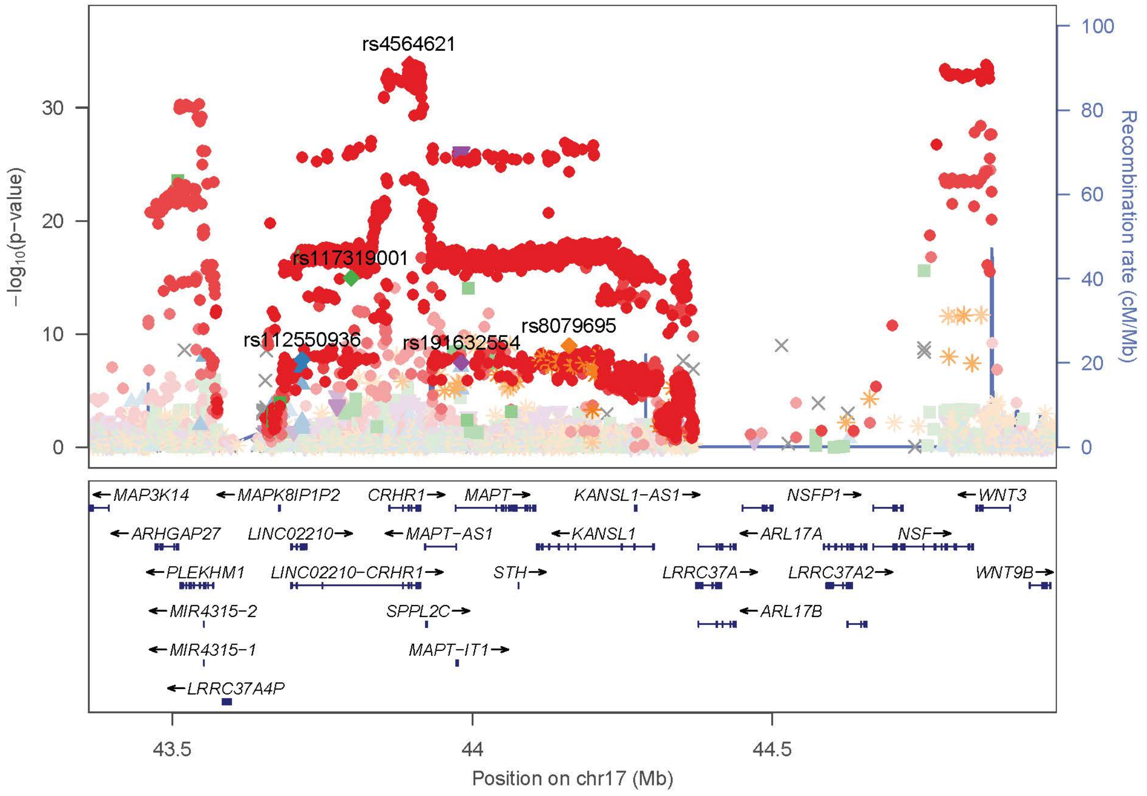
rs4564621
rs117319001
rs112550936
rs191632554
rs8079695
−log10(p−value)
Recombination rate (cM/Mb)
MAP3K14
MAPK8IP1P2
CRHR1
MAPT
KANSL1−AS1
NSFP1
WNT3
ARHGAP27
LINC02210
MAPT−AS1
KANSL1
ARL17A
NSF
PLEKHM1
LINC02210−CRHR1
STH
LRRC37A
LRRC37A2
WNT9B
MIR4315−2
SPPL2C
ARL17B
MIR4315−1
MAPT−IT1
LRRC37A4P
43.5
44
44.5
Position on chr17 (Mb)

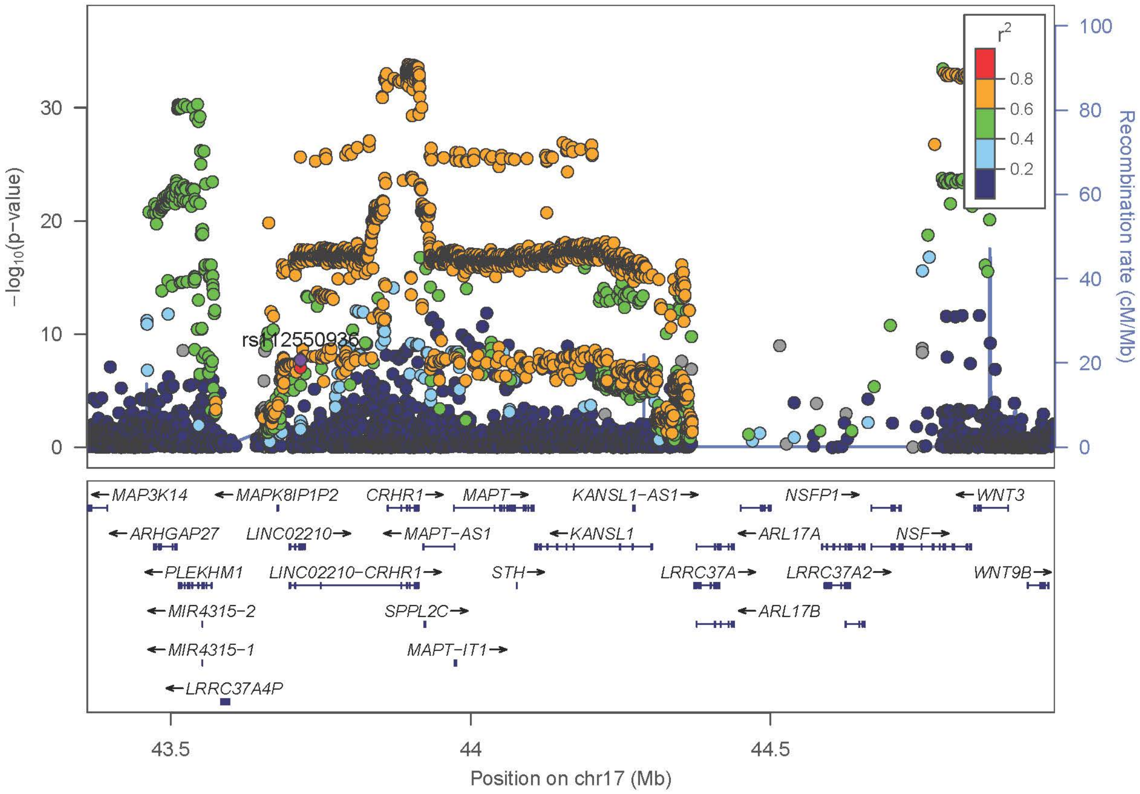

rs112550936
r2
0.8
0.6
0.4
0.2
30
20
10
0
−log10(p−value)
100
80
60
40
20
0
Recombination rate (cM/Mb)
←MAP3K14
←MAPK8IP1P2
CRHR1→
MAPT→
KANSL1−AS1→
NSFP1→
←WNT3
←ARHGAP27
LINC02210→
←MAPT−AS1
←KANSL1
←ARL17A
NSF→
←PLEKHM1
LINC02210−CRHR1→
STH→
LRRC37A→
LRRC37A2→
WNT9B→
←MIR4315−2
SPPL2C→
←ARL17B
←MIR4315−1
MAPT−IT1→
←LRRC37A4P
43.5
44
44.5
Position on chr17 (Mb)


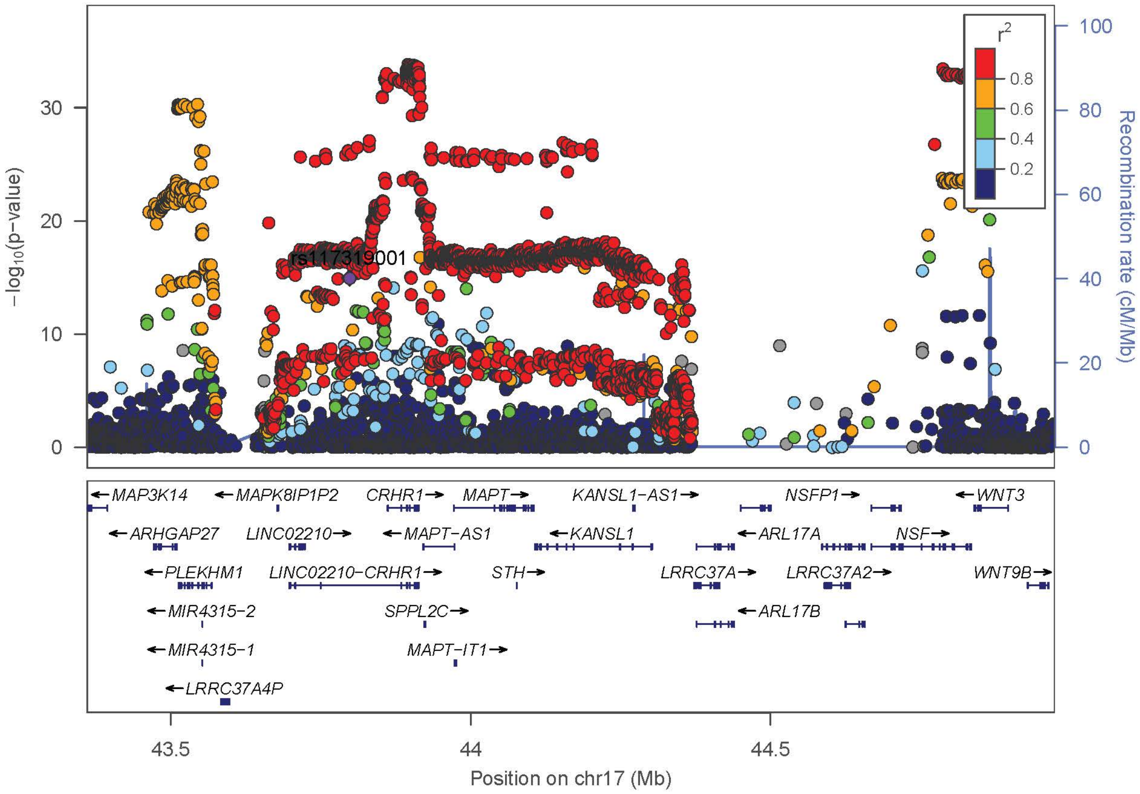

rs117319001
r2
0.8
0.6
0.4
0.2
30
20
10
0
−log10(p−value)
100
80
60
40
20
0
Recombination rate (cM/Mb)
←MAP3K14
←MAPK8IP1P2
CRHR1→
MAPT→
KANSL1−AS1→
NSFP1→
←WNT3
←ARHGAP27
LINC02210→
←MAPT−AS1
←KANSL1
←ARL17A
NSF→
←PLEKHM1
LINC02210−CRHR1→
STH→
LRRC37A→
LRRC37A2→
WNT9B→
←MIR4315−2
SPPL2C→
←ARL17B
←MIR4315−1
MAPT−IT1→
←LRRC37A4P
43.5
44
44.5
Position on chr17 (Mb)

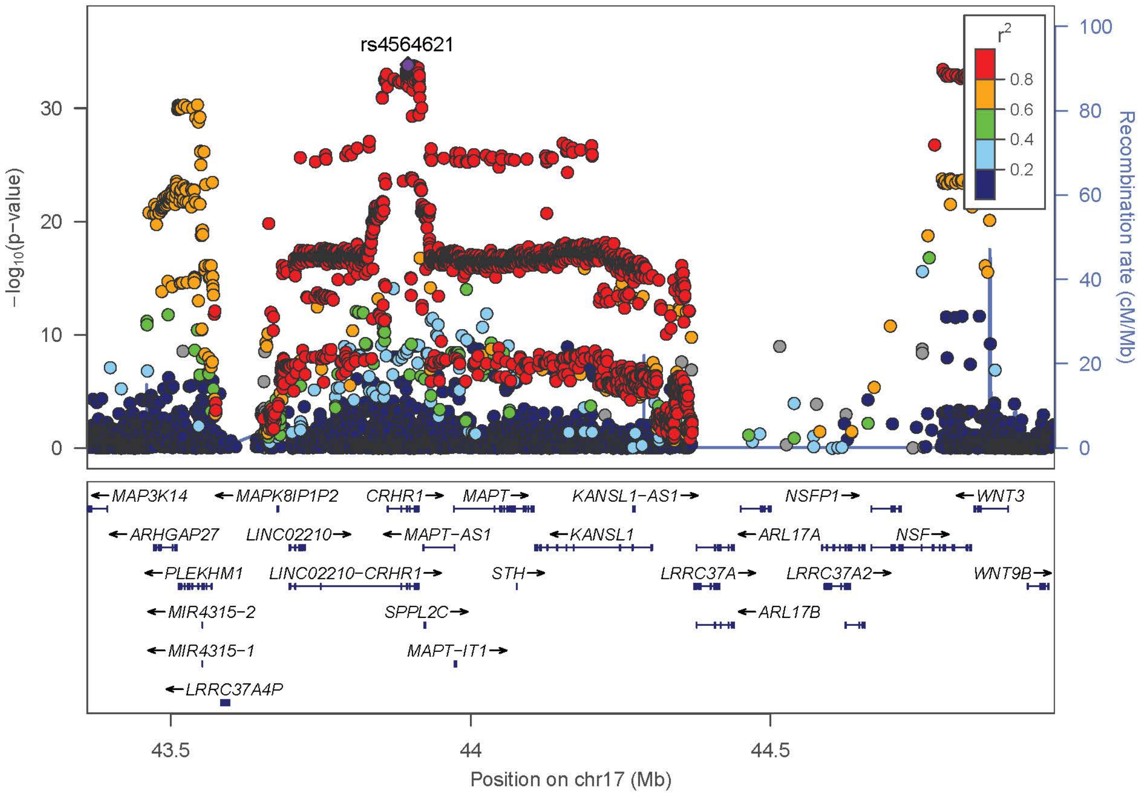

rs4564621
r2
0.8
0.6
0.4
0.2
−log10(p−value)
Recombination rate (cM/Mb)
MAP3K14
MAPK8IP1P2
CRHR1
MAPT
KANSL1−AS1
NSFP1
WNT3
ARHGAP27
LINC02210
MAPT−AS1
KANSL1
ARL17A
NSF
PLEKHM1
LINC02210−CRHR1
STH
LRRC37A
LRRC37A2
WNT9B
MIR4315−2
SPPL2C
ARL17B
MIR4315−1
MAPT−IT1
LRRC37A4P
43.5
44
44.5
Position on chr17 (Mb)


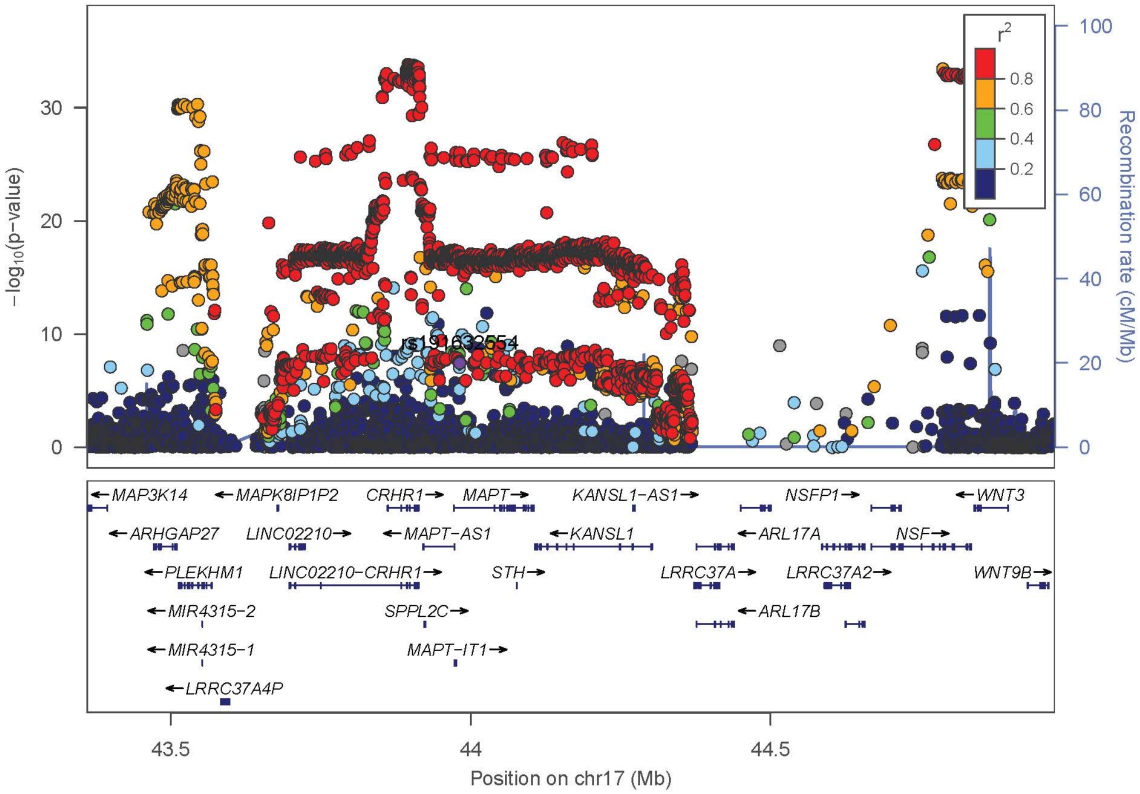

rs191632554
r2
0.8
0.6
0.4
0.2
−log10(p−value)
Recombination rate (cM/Mb)
MAP3K14
MAPK8IP1P2
CRHR1
MAPT
KANSL1−AS1
NSFP1
WNT3
ARHGAP27
LINC02210
MAPT−AS1
KANSL1
ARL17A
NSF
PLEKHM1
LINC02210−CRHR1
STH
LRRC37A
LRRC37A2
WNT9B
MIR4315−2
SPPL2C
ARL17B
MIR4315−1
MAPT−IT1
LRRC37A4P
43.5
44
44.5
Position on chr17 (Mb)

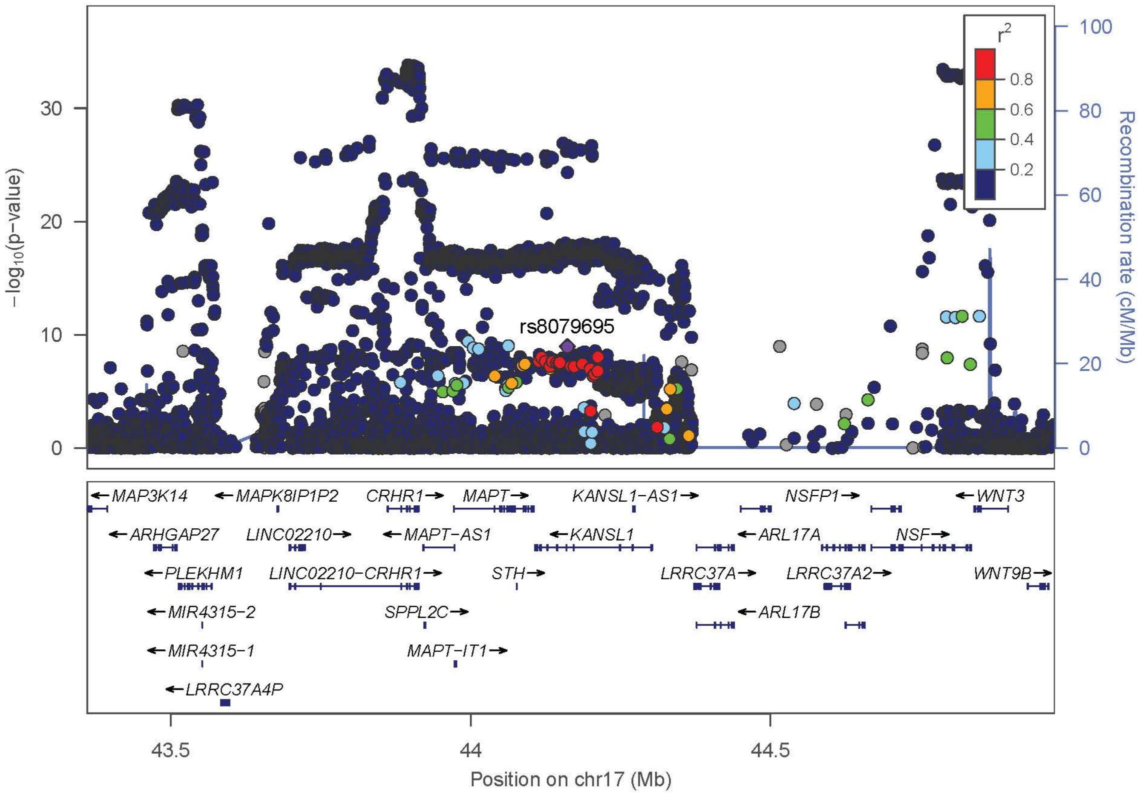
rs8079695
−log10(p−value)
Recombination rate (cM/Mb)
r2
0.8
0.6
0.4
0.2
MAP3K14
MAPK8IP1P2
CRHR1
MAPT
KANSL1−AS1
NSFP1
WNT3
ARHGAP27
LINC02210
MAPT−AS1
KANSL1
ARL17A
NSF
PLEKHM1
LINC02210−CRHR1
STH
LRRC37A
LRRC37A2
WNT9B
MIR4315−2
SPPL2C
ARL17B
MIR4315−1
MAPT−IT1
LRRC37A4P
Position on chr17 (Mb)

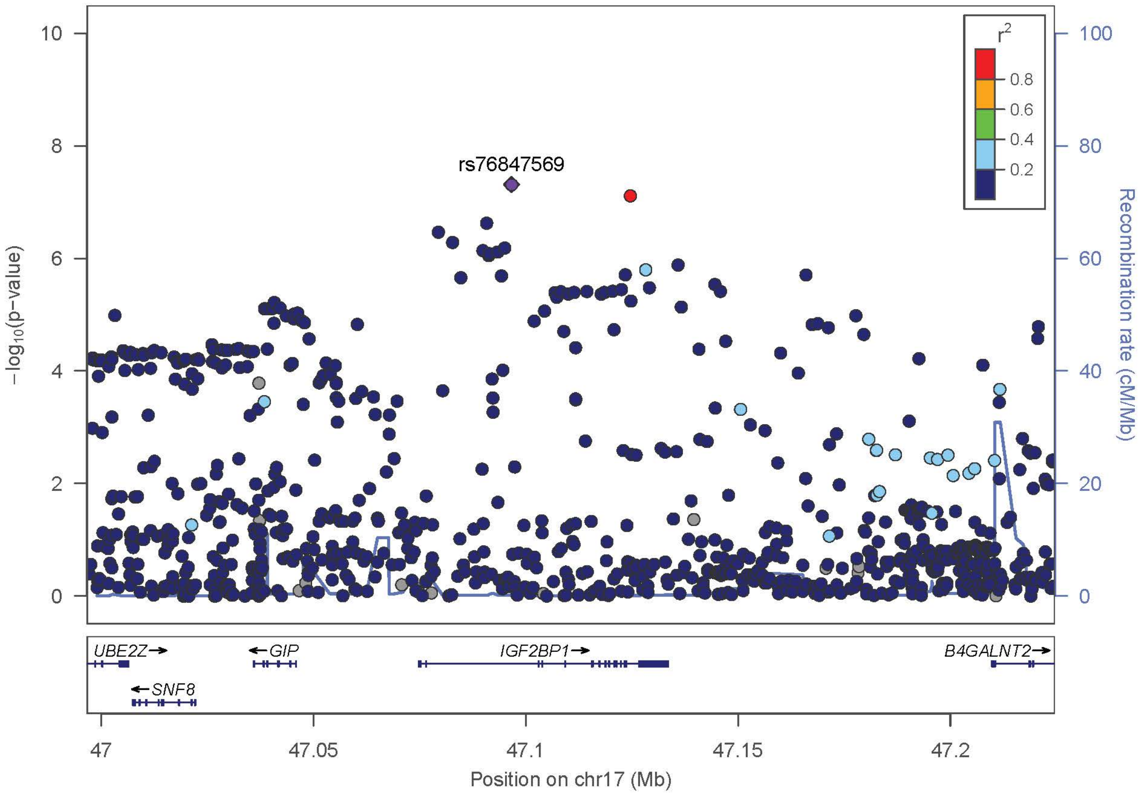
rs76847569
−log10(p−value)
Recombination rate (cM/Mb)
r2
0.8
0.6
0.4
0.2
UBE2Z
GIP
IGF2BP1
B4GALNT2
SNF8
Position on chr17 (Mb)

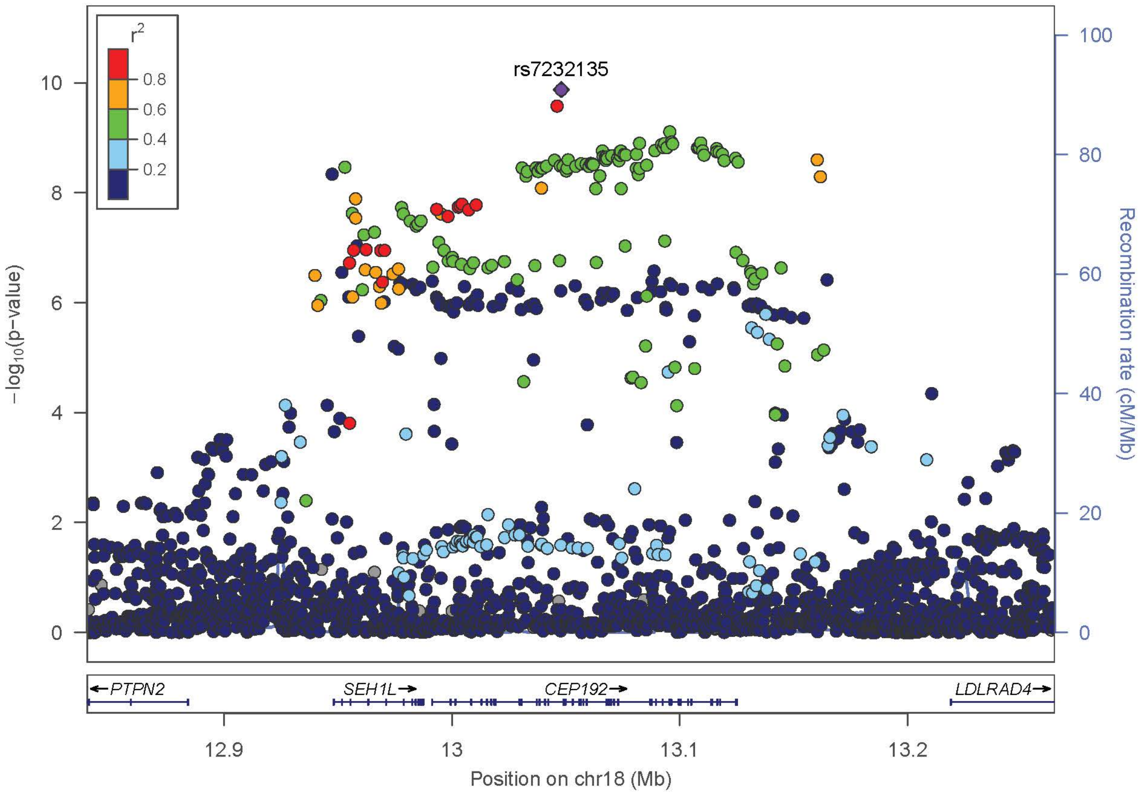

r2
0.8
0.6
0.4
0.2
rs7232135
−log10(p-value)
Recombination rate (cM/Mb)
PTPN2
SEH1L
CEP192
LDLRAD4
12.9
13
13.1
13.2
Position on chr18 (Mb)


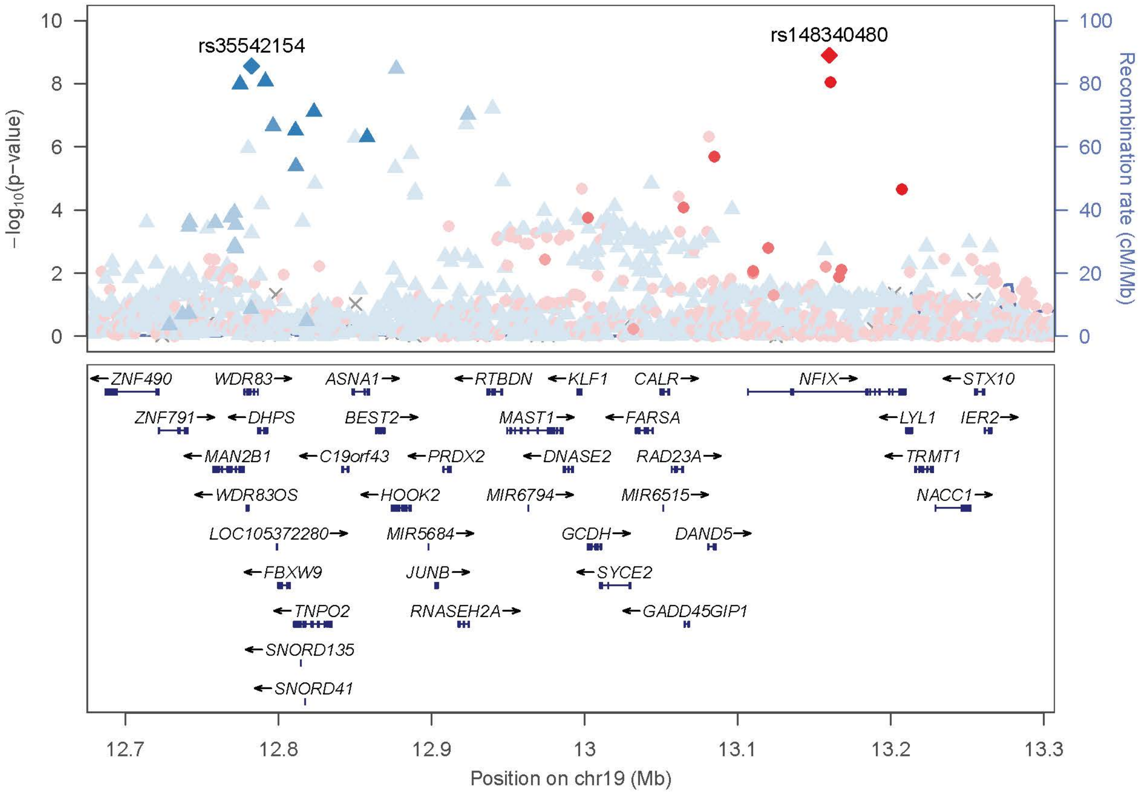

rs35542154
rs148340480
−log10(p-value)
Recombination rate (cM/Mb)
ZNF490
WDR83
ASNA1
RTBDN
KLF1
CALR
NFIX
STX10
ZNF791
DHPS
BEST2
MAST1
FARSA
LYL1
IER2
MAN2B1
C19orf43
PRDX2
DNASE2
RAD23A
TRMT1
WDR83OS
HOOK2
MIR6794
MIR6515
NACC1
LOC105372280
MIR5684
GCDH
DAND5
FBXW9
JUNB
SYCE2
TNPO2
RNASEH2A
GADD45GIP1
SNORD135
SNORD41
12.7
12.8
12.9
13
13.1
13.2
13.3
Position on chr19 (Mb)

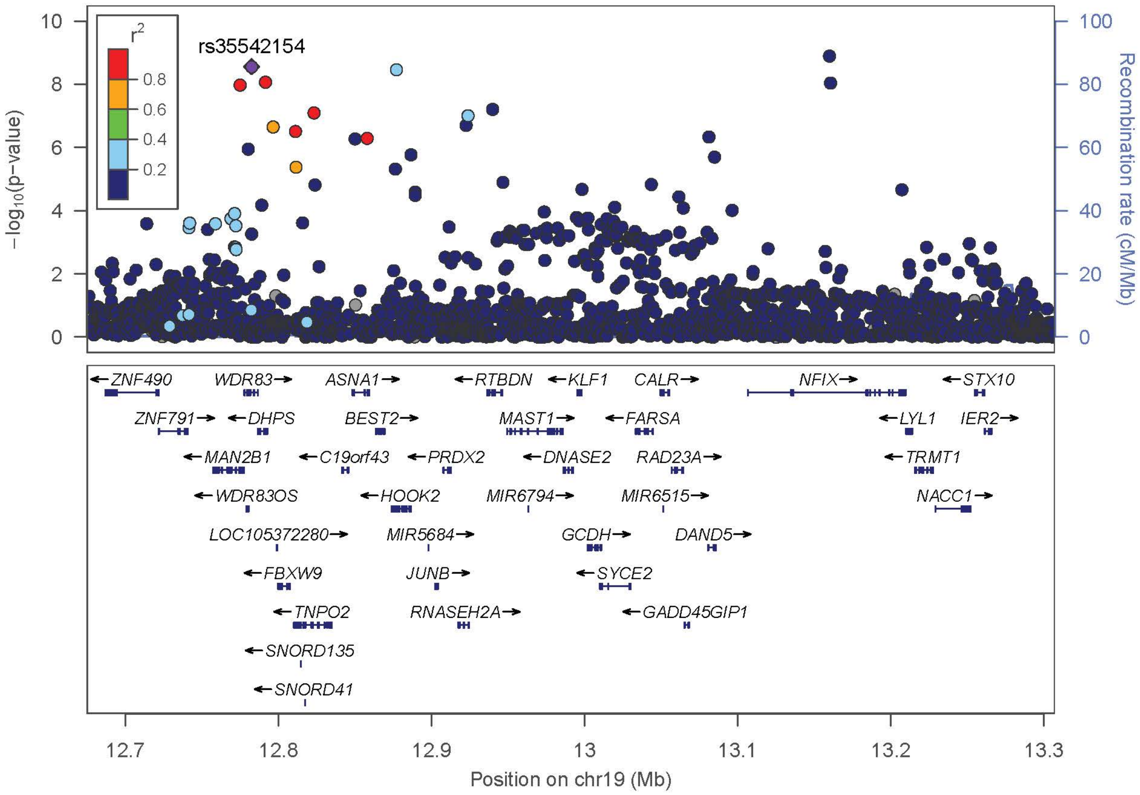
rs35542154
r²
0.8
0.6
0.4
0.2
−log10(p−value)
Recombination rate (cM/Mb)
ZNF490
WDR83
ASNA1
RTBDN
KLF1
CALR
NFIX
STX10
ZNF791
DHPS
BEST2
MAST1
FARSA
LYL1
IER2
MAN2B1
C19orf43
PRDX2
DNASE2
RAD23A
TRMT1
WDR83OS
HOOK2
MIR6794
MIR6515
NACC1
LOC105372280
MIR5684
GCDH
DAND5
FBXW9
JUNB
SYCE2
TNPO2
RNASEH2A
GADD45GIP1
SNORD135
SNORD41
12.7
12.8
12.9
13
13.1
13.2
13.3
Position on chr19 (Mb)

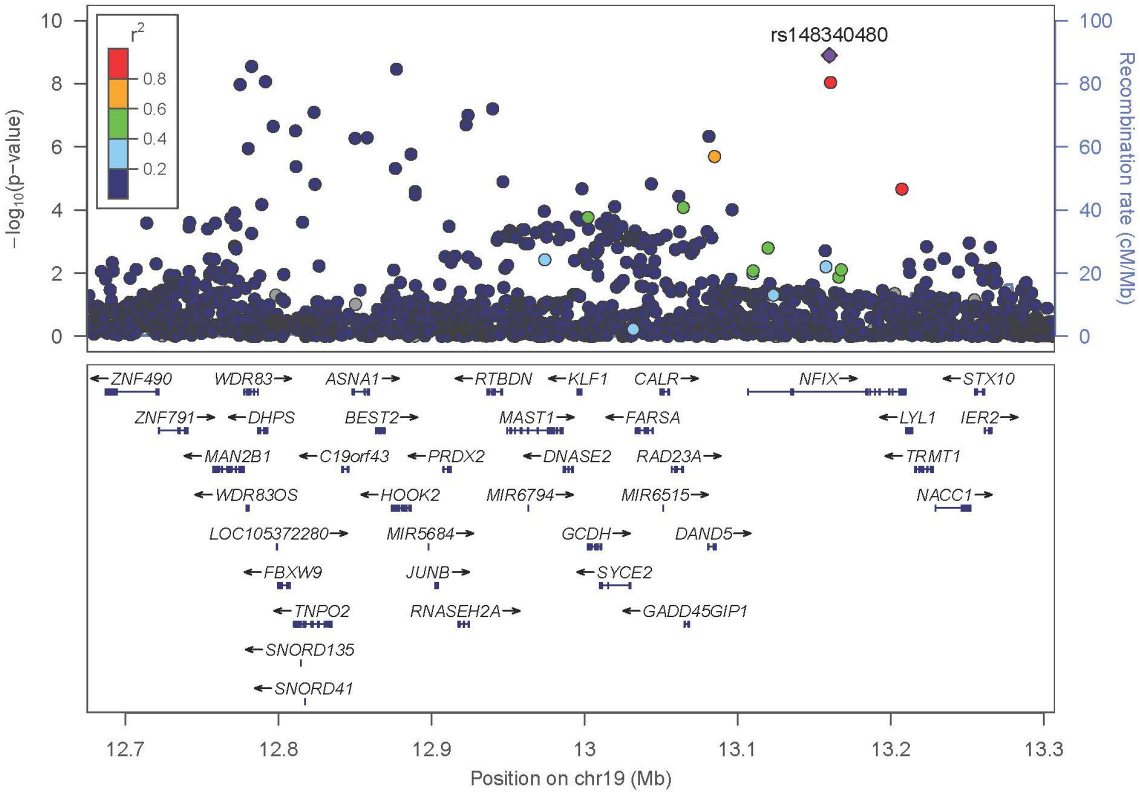
rs148340480
r²
0.8
0.6
0.4
0.2
−log10(p−value)
Recombination rate (cM/Mb)
ZNF490
WDR83
ASNA1
RTBDN
KLF1
CALR
NFIX
STX10
ZNF791
DHPS
BEST2
MAST1
FARSA
LYL1
IER2
MAN2B1
C19orf43
PRDX2
DNASE2
RAD23A
TRMT1
WDR83OS
HOOK2
MIR6794
MIR6515
NACC1
LOC105372280
MIR5684
GCDH
DAND5
FBXW9
JUNB
SYCE2
TNPO2
RNASEH2A
GADD45GIP1
SNORD135
SNORD41
12.7
12.8
12.9
13
13.1
13.2
13.3
Position on chr19 (Mb)

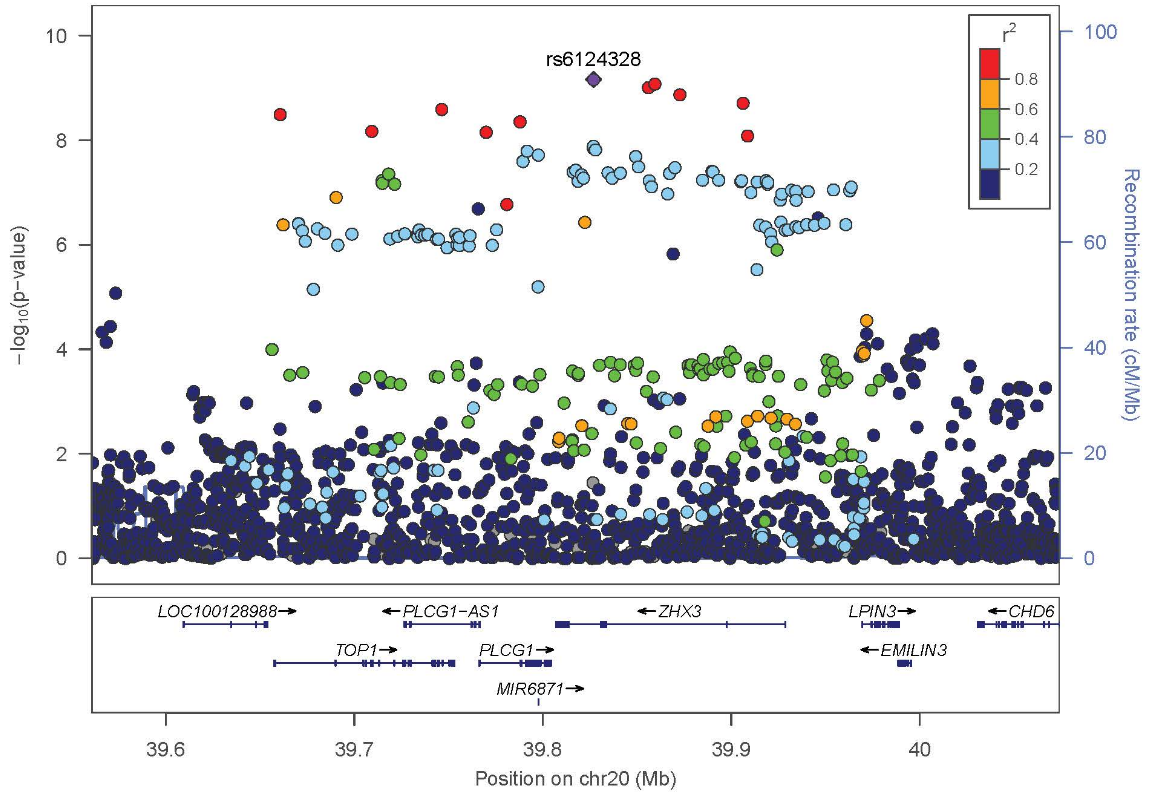

rs6124328
r2
0.8
0.6
0.4
0.2
−log10(p-value)
Recombination rate (cM/Mb)
LOC100128988→
←PLCG1−AS1
←ZHX3
LPIN3→
←CHD6
TOP1→
PLCG1→
←EMILIN3
MIR6871→
39.6
39.7
39.8
39.9
40
Position on chr20 (Mb)


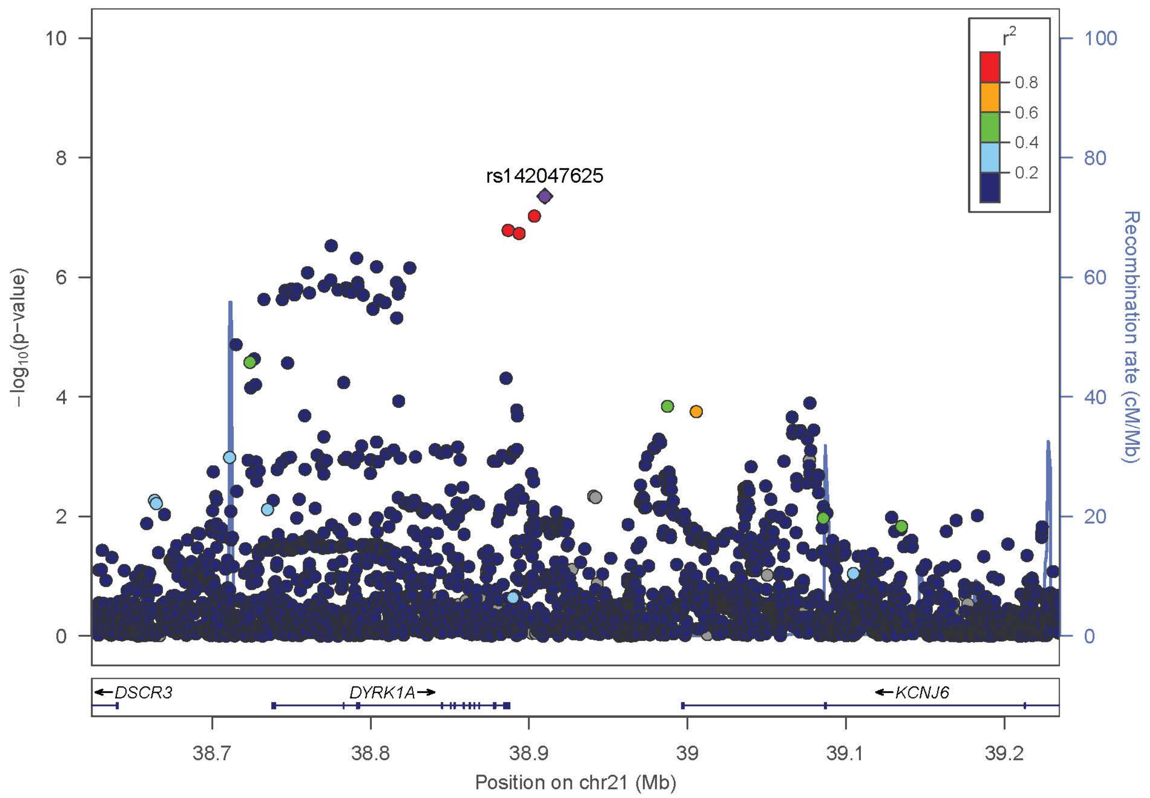

rs142047625
r2
0.8
0.6
0.4
0.2
−log10(p-value)
Recombination rate (cM/Mb)
←DSCR3
DYRK1A→
←KCNJ6
38.7
38.8
38.9
39
39.1
39.2
Position on chr21 (Mb)

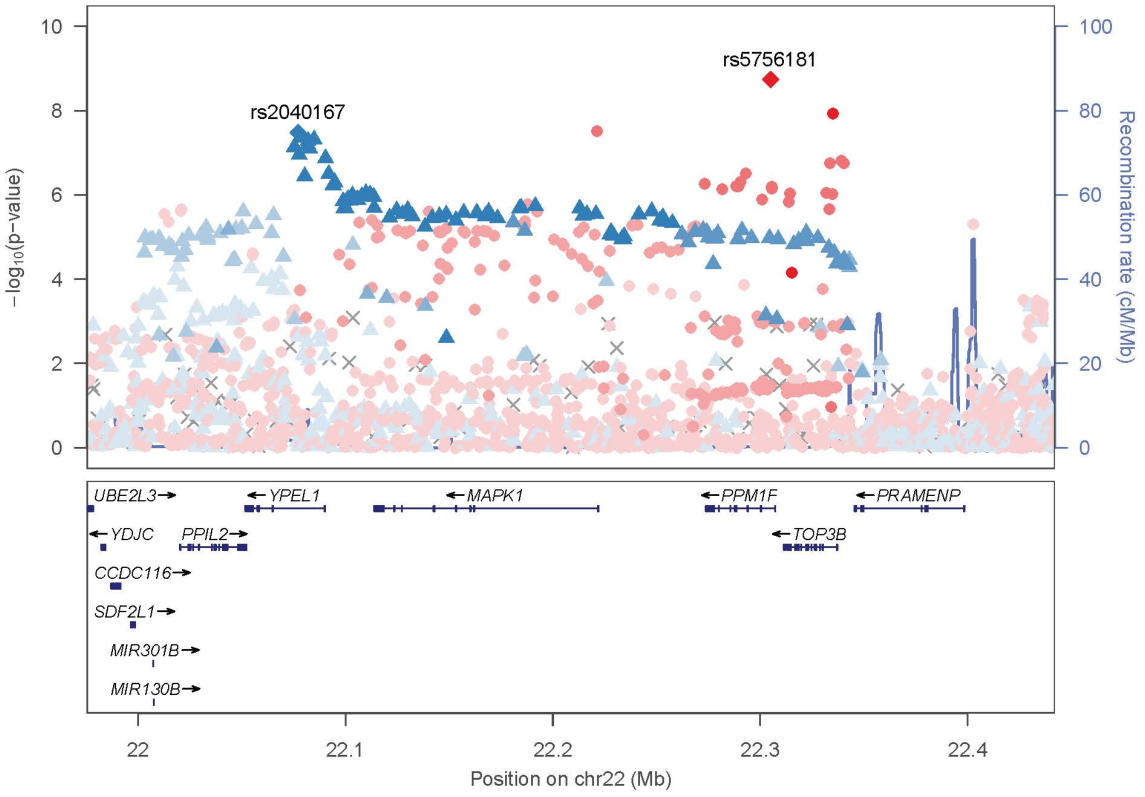
rs5756181
rs2040167
10
8
6
4
2
0
−log10(p−value)
100
80
60
40
20
0
Recombination rate (cM/Mb)
UBE2L3→
←YPEL1
←MAPK1
←PPM1F
←PRAMENP
←YDJC
PPIL2→
←TOP3B
CCDC116→
SDF2L1→
MIR301B→
MIR130B→
22
22.1
22.2
22.3
22.4
Position on chr22 (Mb)

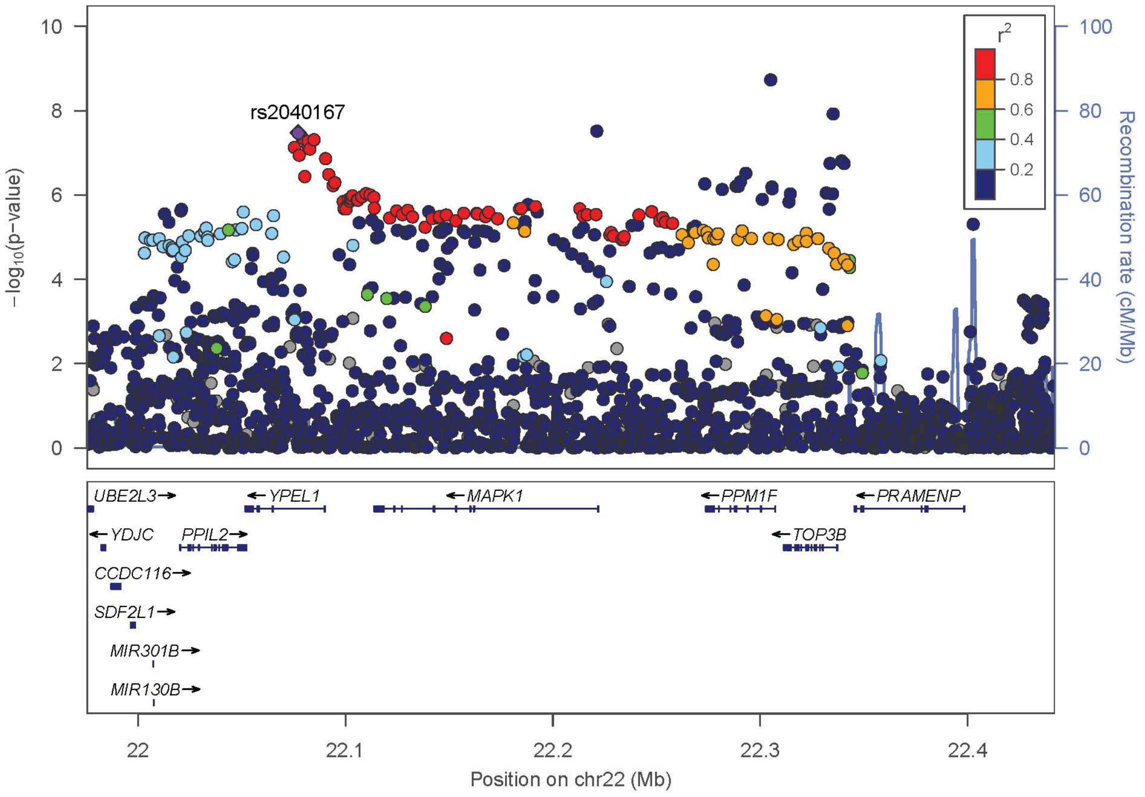
rs2040167
$r^2$
0.8
0.6
0.4
0.2
10
8
6
4
2
0
−log10(p−value)
100
80
60
40
20
0
Recombination rate (cM/Mb)
UBE2L3→
←YPEL1
←MAPK1
←PPM1F
←PRAMENP
←YDJC
PPIL2→
←TOP3B
CCDC116→
SDF2L1→
MIR301B→
MIR130B→
22
22.1
22.2
22.3
22.4
Position on chr22 (Mb)

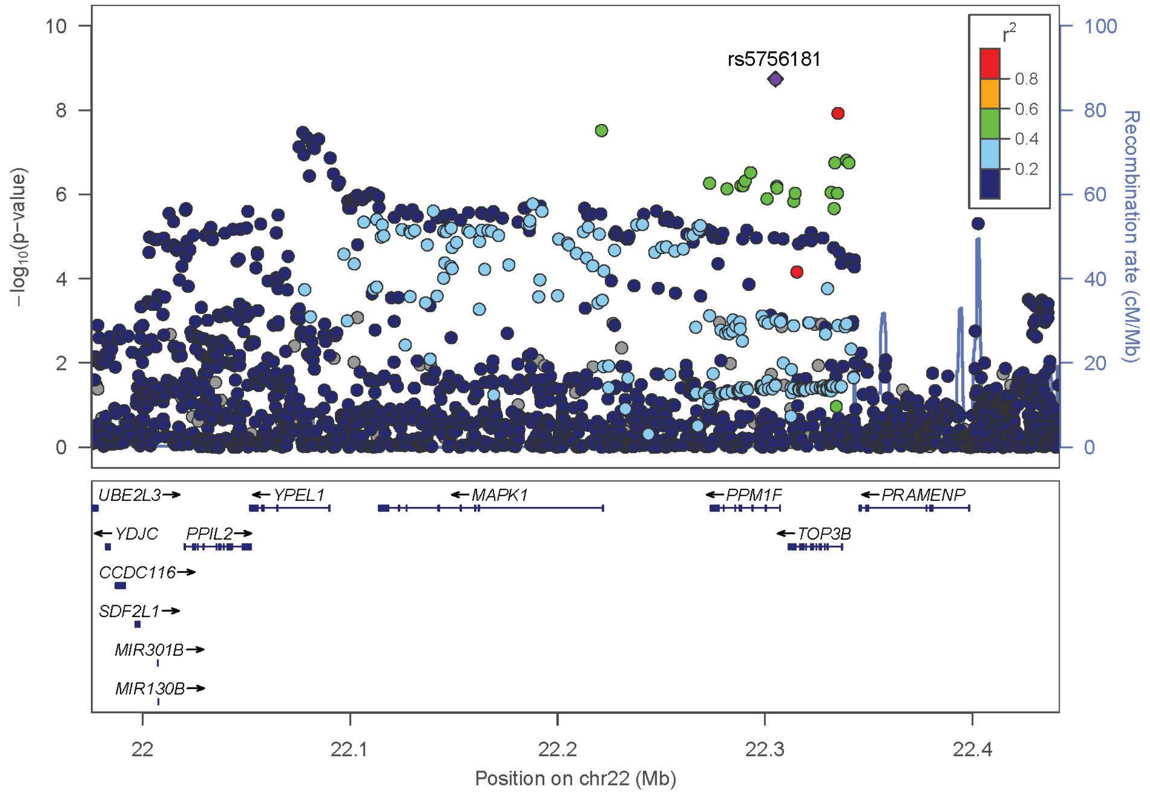
rs5756181
r2
0.8
0.6
0.4
0.2
10
8
6
4
2
0
−log10(p−value)
100
80
60
40
20
0
Recombination rate (cM/Mb)
UBE2L3
YPEL1
MAPK1
PPM1F
PRAMENP
YDJC
PPIL2
TOP3B
CCDC116
SDF2L1
MIR301B
MIR130B
22
22.1
22.2
22.3
22.4
Position on chr22 (Mb)

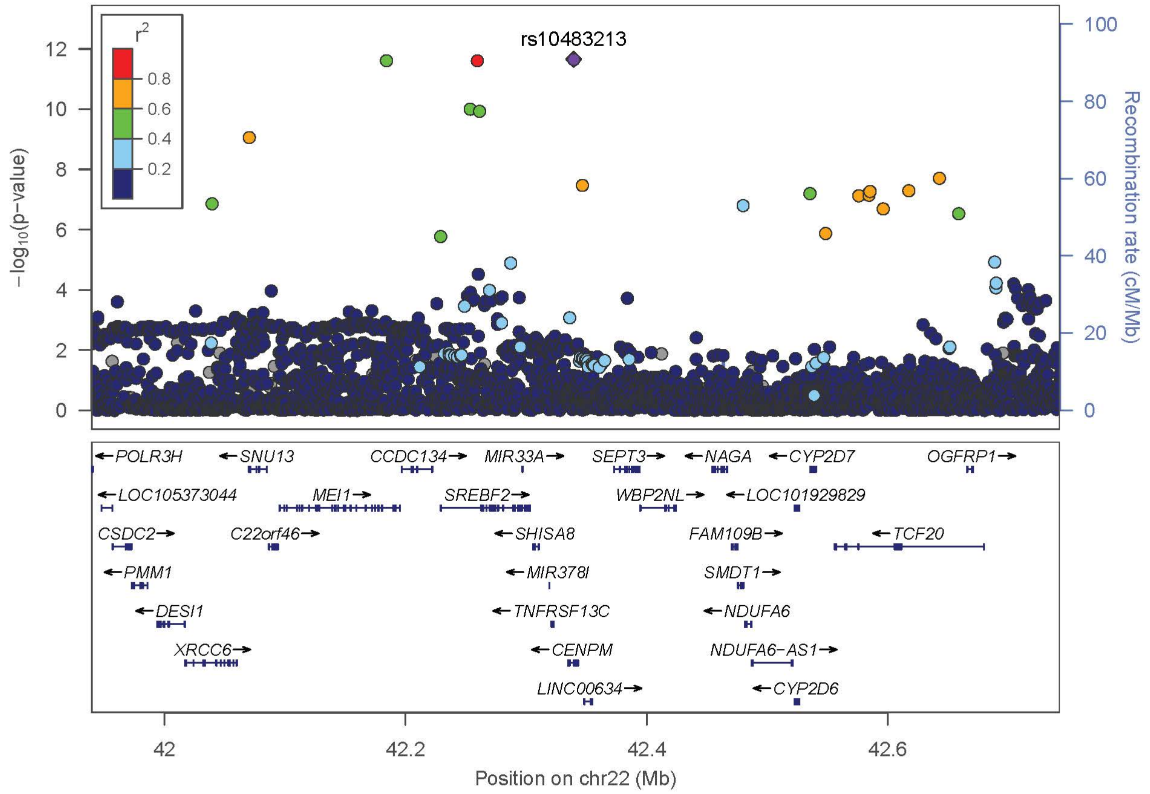
rs10483213
r2
0.8
0.6
0.4
0.2
12
10
8
6
4
2
0
−log10(p−value)
100
80
60
40
20
0
Recombination rate (cM/Mb)
POLR3H
SNU13
CCDC134
MIR33A
SEPT3
NAGA
CYP2D7
OGFRP1
LOC105373044
MEI1
SREBF2
WBP2NL
LOC101929829
CSDC2
C22orf46
SHISA8
FAM109B
TCF20
PMM1
MIR378I
SMDT1
DESI1
TNFRSF13C
NDUFA6
XRCC6
CENPM
NDUFA6−AS1
LINC00634
CYP2D6
42
42.2
42.4
42.6
Position on chr22 (Mb)
